# Defining the role of fibroblasts in skin expansion

**DOI:** 10.1101/2025.03.18.643853

**Authors:** Caroline Aguilera Stewart, Ceyhun Alar, Leslie Bargsted, Ioanna Kaklamanou, Gaia Andrea Gozza, Maria Pia Polito, Elena Enzo, Alejandro Sifrim, Mariaceleste Aragona

**Affiliations:** Novo Nordisk Foundation Center for Stem Cell Medicine (reNEW), Department of Biomedical Sciences (BMI), University of Copenhagen, Copenhagen, Denmark; KU Leuven Institute for Single Cell Omics (LISCO), University of Leuven, KU Leuven, Leuven, Belgium; Department of Human Genetics, University of Leuven, KU Leuven, Leuven, Belgium; Centre for Regenerative Medicine “Stefano Ferrari”, University of Modena and Reggio Emilia, Modena, Italy

## Abstract

Stretch-mediated tissue expansion is commonly used to grow extra skin for reconstructive surgeries. To ensure harmonious growth, the two main skin compartments, the epidermis and the dermis, must both expand in a coordinated manner. Although the epidermal response has been previously described, it remains unclear how fibroblasts, the main supporting cell type, respond to stretching in vivo. Here we map the transcriptional response of the entire skin during stretch-mediated tissue expansion, and we describe the fibroblast response to stretching in vivo. We show an increase in fibroblast volume accompanied by changes in organisation. We demonstrate that stretching forces fibroblasts to exit their quiescent state and restart proliferation. Simultaneously, fibroblasts decrease their collagen content and increase the expression of specific extracellular matrix remodelling factors. By combining data from the in vivo stretching model and an in vitro keratinocyte-fibroblast co-culture system, we demonstrate that changes in fibroblasts promote the self-renewal of the epidermal stem cells, thus coordinating the response of these two compartments during skin expansion. These findings provide valuable insights to guide the design of in vivo stretch-mediated tissue expansion protocols and the production of in vitro skin grafts for clinical application.

## Introduction

The skin serves as the body’s outermost layer, playing a crucial role in protecting against external threats^1^. Within epithelial tissues, mechanical forces are continuously present during homeostasis and wound healing, ensuring the restoration of tissue integrity and the preservation of its intrinsic functions^2–6^. Mechanotransduction converts mechanical stimuli into biochemical signals and gene transcription, thereby influencing not only tissue morphology but also cell behaviour^7,8^. To establish and maintain functional organs, the tissue response to these mechanical forces must be precisely coordinated across different compartments^9^.

Stretch-mediated tissue expansion is a common plastic surgery technique that promotes the growth of extra skin that can then be transplanted for repairing birth defects, scars, or creating space for permanent implants in breast reconstruction following mastectomy^10–12^. Due to its capacity to adapt and respond to external physical stimuli, the skin is an excellent model to study cellular interactions and compartment coordination under mechanical forces^1,13,14^. Few animal models of in vivo skin expansion have been described^15–18^. A model was established in mice, using self-inflating hydrogel expanders, to mimic the expansion process in humans^15^. This model was used to study the effects of stretching on the epidermis, revealing that epidermal stem cells’ fate decisions were transiently biased towards selfrenewal^15^. However, the response of fibroblasts, the main cellular component of the dermis, has not been studied to date^19^. How the main skin compartments, the epidermis and dermis, coordinate their response to stretching and whether fibroblasts provide signals to promote epidermal stem cell self-renewal remains unknown.

## Results

### A transcriptomic atlas of stretch-mediated skin expansion

To study the effect of stretch-mediated skin expansion more broadly on the skin, we performed single-cell RNA sequencing (scRNAseq) in control (CTRL) and expansion (EXP), using the previously described mouse model^15^. All mice underwent surgery, whilst the CTRL mice did not have an expander inserted (Fig. 1a). Live skin cells were isolated by fluorescenceactivated cell sorting (FACS) (Supplementary Fig. 1a). We profiled living cells at day 1 (D1), when the maximal stretch is reached, and at day 2 (D2) and day 4 (D4) when the epidermal stem cells are biased towards self-renewal^15^ (Fig 1. a-c, Supplementary Fig. 1b and Supplementary Table 1). In the entire atlas (n=32 animals, n=22.412 CTRL cells and n=13.904 EXP cells) we uncovered all the different cell types constituting the mouse back skin^20–22^ (Fig. 1b,c, Supplementary Fig. 1b and Supplementary Table 1). Proportionally, the keratinocyte and monocyte clusters contain more EXP cells overall, validating previous data on epidermal expansion and the recruitment of inflammatory cells, whereas the fibroblast cluster contains more CTRL cells (Supplementary Fig. 1c and Supplementary Table 2).

**Figure 1.**
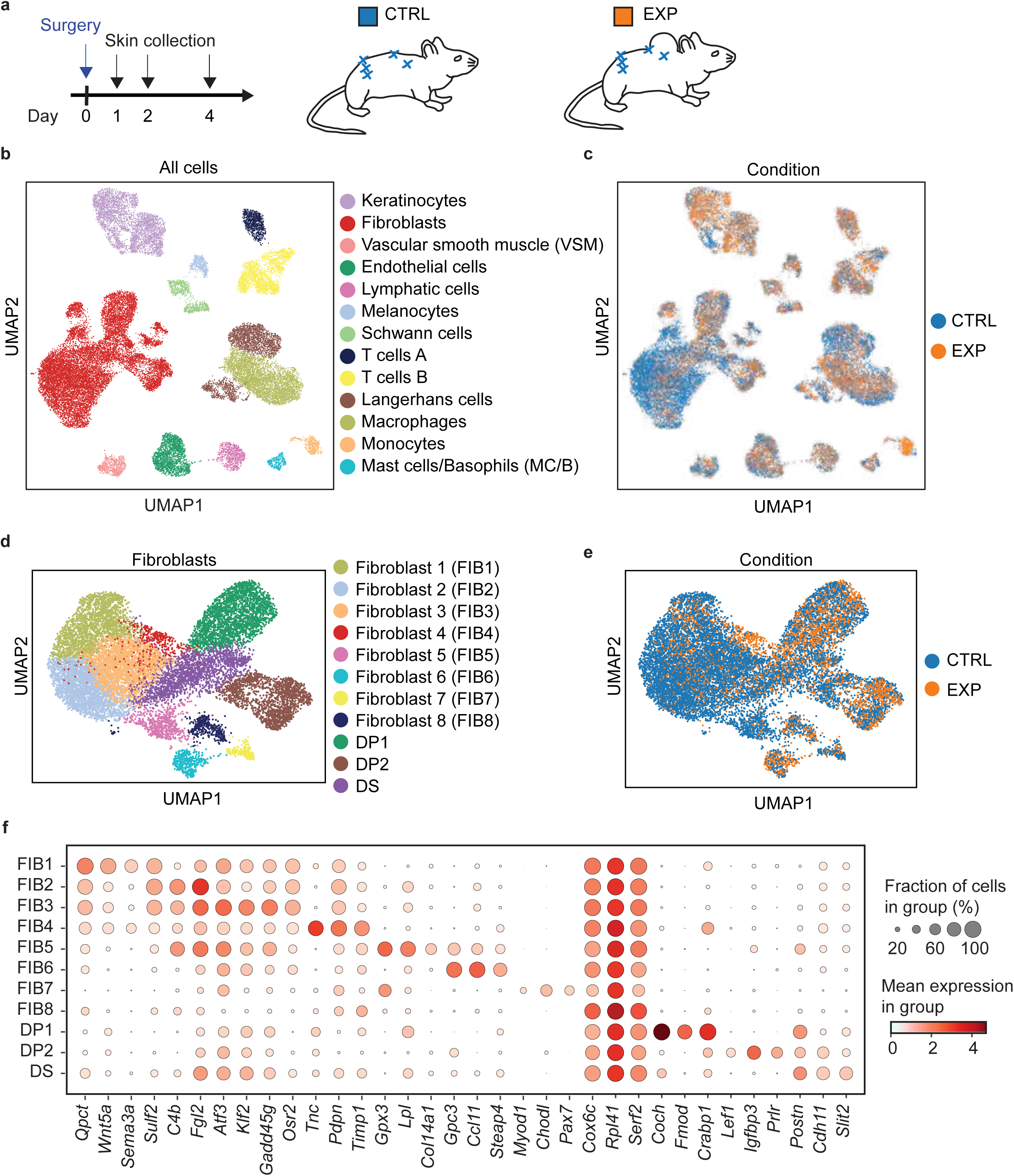
Single-cell RNA sequencing of stretch-mediated skin expansion. **a** – Schematic of the experimental procedure. **b** – UMAP plot of the clustering analysis of scRNAseq data of CTRL (22412 cells, number of mice: D1=4, D2=7, D4=4) and EXP (13904 cells, number of mice: D1=4, D2=7, D4=4) samples. **c** – UMAP plot of the distribution of CTRL and EXP cells within clusters shown in **b**. **d** – UMAP plot of the subclustering analysis of CTRL (11634 cells) and EXP (3785 cells) fibroblast and fibroblast-like cells. **e** – UMAP plot of the distribution of CTRL and EXP cells within subclusters shown in **d**. **f** – Marker gene expression for each of the subclusters shown in **d**.

### scRNAseq of fibroblasts

Given that fibroblasts and other stromal cells are the most important cell types in the dermis, providing structural support and modulating different biological processes^19,23,24^, we decided to investigate this cluster further. In total, there were 15.419 fibroblast and stromal cells (also referred as fibroblast-like cells in the text) in our dataset (Fig. 1d, and Supplementary Fig. 1b,c). Using graph-based clustering, we identified different subpopulations based on marker gene expression as described in previous studies^21,25,26^. We distinguished eight subclusters of fibroblasts (FIB1-8), as well as two dermal papilla (DP1-2) and one dermal sheath subclusters (DS) (Fig. 1d and Supplementary Table 3). The proportions of the different fibroblast subtypes changed during expansion (Fig. 1e, Supplementary Fig. 1d and Supplementary Table 4), with a vast increase of cells from the FIB4 and FIB6 clusters during expansion. Samples from the CTRL and EXP conditions were well distributed between the fibroblast subclusters (Supplementary Fig. 1e-g). Subclusters FIB1-3 were more transcriptionally similar than the other fibroblast subpopulations, FIB5-8 (Fig. 1f). The FIB4 subpopulation, whilst expressing many similar genes to FIB1-3, also shows higher expression of *Pdpn*, *Tnc and Timp1* (Fig. 1f). The transcriptional similarity of FIB4 to FIB1-3 subclusters may indicate that FIB4 is a cell state rather than a new fibroblast subtype, becoming more prominent during expansion (Supplementary Fig. 1d and Supplementary Table 4). The subcluster FIB8 (n=350 cells) has overall lower counts and higher ribosomal gene expression, and we could not identify unique marker genes for this population. Therefore, we excluded it from further analyses.

### Spatial mapping of fibroblast subpopulations

As we are interested in the fibroblast interactions with the keratinocytes, we performed spatial transcriptomics on four D2 CTRL skin samples to determine whether the fibroblast subclusters we identified in the scRNAseq analysis indeed inhabit spatially distinct niches and to determine which clusters are present in the upper dermis closer to the keratinocytes. We used the 10x Genomics Xenium platform to run a panel of 95 custom genes based on the expression patterns of the scRNAseq dataset and 379 genes from the predesigned Xenium mouse tissue atlas. Cells in the spatial dataset were annotated based on the gene expression signatures we found in our scRNAseq dataset (Fig. 1d-f and Supplementary Fig. 2a-e). We concluded that subpopulations FIB1-4 were mostly located in the upper dermis, and subclusters 5-7 were more enriched in the lower regions of the skin (Fig. 2a-c and Supplementary Fig. 2f, g). We identified *Pdpn* as a marker enriched in the upper dermis (Fig. 2d and Supplementary Fig. 2h) whereas expression of *Gpx3* and *Mfap5,* as expected^21^, was mostly restricted to the lower dermis (Fig. 2e,f and Supplementary Fig. 2i,j). The plateletderived growth factor receptor alpha (*Pdgfrα*), which is commonly used as general marker of dermal fibroblasts^27^ was broadly expressed in both upper and lower dermis (Supplementary Fig. 2k). We thus established that in the mouse dorsal skin, the transcriptional heterogeneity reflects at least two spatially distinct anatomical regions of the upper and lower dermis.

**Figure 2.**
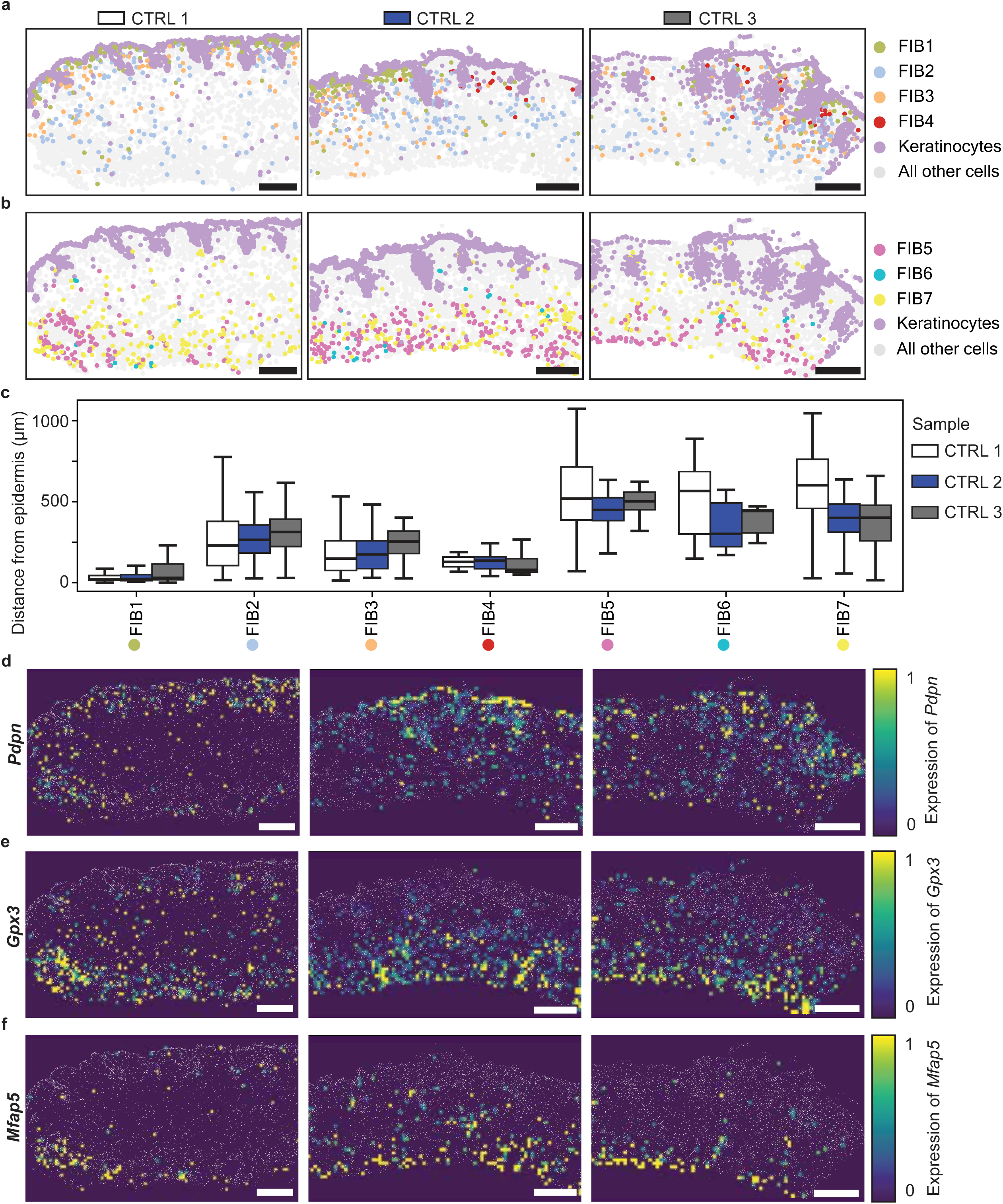
Spatial mapping of fibroblast subpopulations. **a** – Representative image of the spatial distribution of FIB1-4 subpopulations in D2 CTRL skin (n=3 section per mouse). Keratinocytes are shown in purple. **b** – Representative image of the spatial distribution of FIB5-7 subpopulations in D2 CTRL skin (n=3 section per mouse). Keratinocytes are shown in purple. **c** – Fibroblast distance from the epidermis per sample and fibroblast subtype. Data are presented as box and whisker plots, showing minimum, lower quartile, median, upper quartile and maximum values. **d-f** – Representative images of the spatial density plots showing relative expression of *Pdpn*, *Gpx3* and *Mfap5* transcripts. Nuclei are shown in grey. Scale normalized per gene transcript. Bin size=20 μm. Scale bars=200 μm.

### Stretching induces a rapid transcriptional response in fibroblasts

To dissect the response of fibroblasts to stretching, we decided to look further into the gene expression profiles of fibroblasts in our scRNAseq dataset, focusing in particular on the upper dermis. In the expanded condition, we observed upregulation of extracellular matrix (ECM) remodelling glycoprotein genes (*Pdpn*, *Tnc, Timp1*) as well as genes that have been previously associated with an embryonic-like and regenerative fibroblast signature (*Crabp1* and *Mmp13*, also known as collagenase-3)^25,28,29^, and *Pdgfrα* (Fig. 3a and Supplementary Table 5). The transcriptional changes observed were consistent across mice (Supplementary Fig. 3a-c). To validate these results by qPCR, we enriched for fibroblasts via Fluorescence-activated cell sorting (FACS), first through exclusion of vasculature, hematopoietic, melanocyte and epithelial cells (cells expressing Cd31, Cd45, Cd117 and Epcam, respectively) and then inclusion of Cd34+ and Pdgfrα+ cells^27^ (Supplementary Fig. 3d). Selecting for double positive Cd34+ and Pdgfrα+ cells allowed us to collect most upper dermal fibroblasts whilst excluding hair follicle related fibroblasts, such as DP fibroblasts (Supplementary Fig. 3d). Downregulation of Cd34 upon expansion was observed at the transcriptomic level in the scRNAseq (Fig. 3a and Supplementary Table 5) and via qPCR (Supplementary Fig. 3e) and confirmed at the protein level via flow cytometry and immunostaining (Supplementary Fig. 3f-i). Previously, Cd34 has been described as downregulated in neonatal fibroblasts compared to adult fibroblasts^30^. We confirmed the upregulation of *Pdpn*, *Tnc*, *Timp1* and *Mmp13* via qPCR (Fig. 3b-e). The increased expression of Pdpn and Tnc was also confirmed by immunostaining at D2 and D4 (Fig. 3f-i). The matrix metalloproteinases (Mmps) and the tissue inhibitors of metalloproteinases (Timps) are both required in dynamics contexts of active ECM remodelling^31^. The overexpression of these genes, along with the glycoproteins Podoplanin (Pdpn) and Tenascin-C (Tnc) suggests that the ECM composition changes during tissue expansion becoming more compliant and viscoelastic^32^. It has also been previously demonstrated that Pdpn and Tnc are expressed during development, but their expression becomes limited in healthy adult tissues^30,33,34^. Altogether, the upregulation of Pdpn, Tnc, Timp1, and the downregulation of Cd34 suggest that during expansion fibroblasts reacquire a signature that is more similar to an earlier developmental and regenerative state. These data also show that fibroblasts respond to stretching by changing their transcriptional landscape quite rapidly.

**Figure 3.**
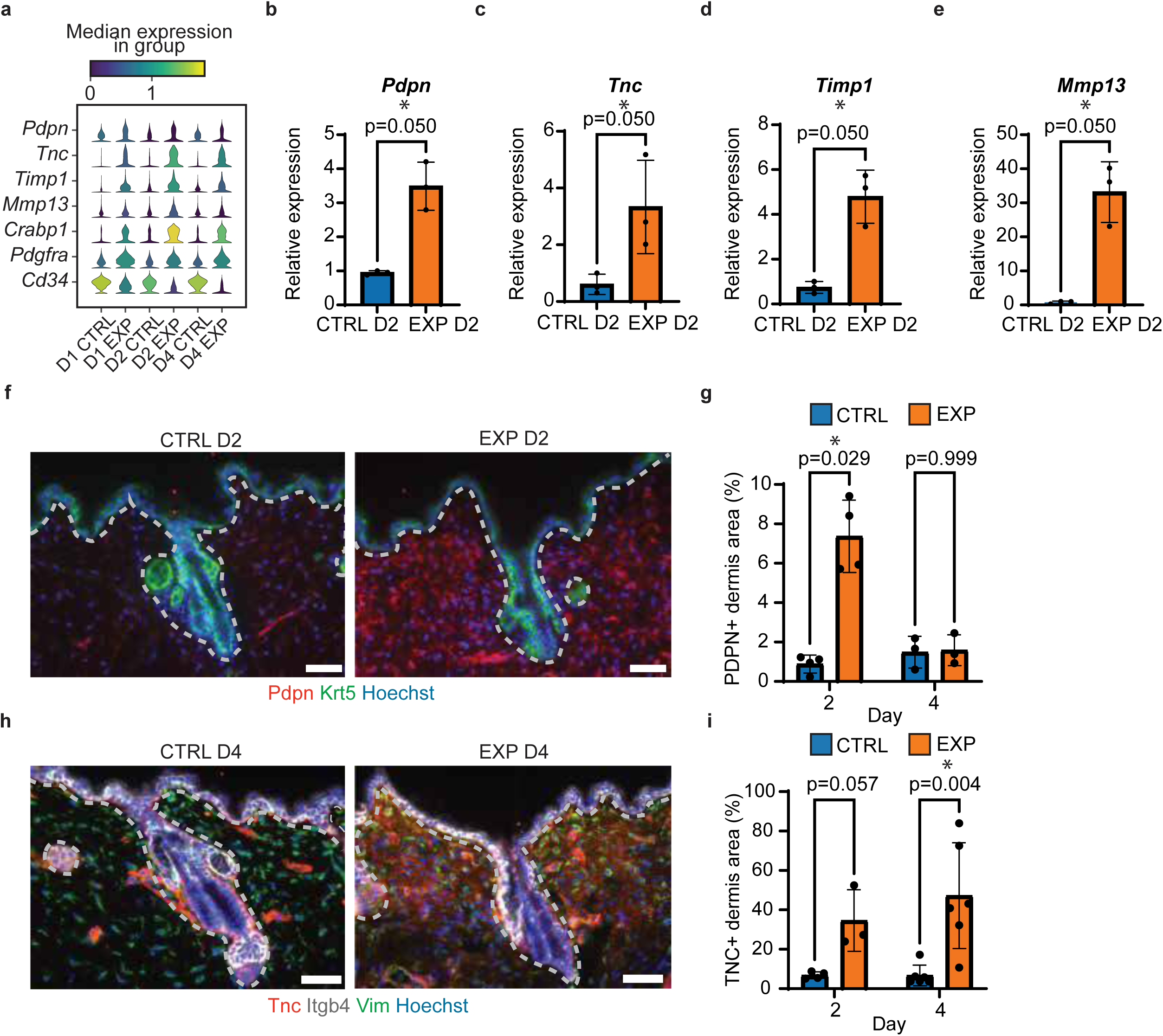
Transcriptional changes in fibroblasts upon expansion. **a** – Stacked violin plot of differentially expressed genes (DEG) in fibroblast and fibroblast-like cells from scRNAseq. **b-e** – Relative expression of the indicated genes in sorted Pdgfrα+ and Cd34+ fibroblasts by qPCR at D2. Unpaired one-tailed Mann-Whitney U-tests, n=3 mice per condition. Data are presented as mean +/-SD. **f** – Immunofluorescence for Pdpn (red), Krt5 (green) and Hoechst (blue) on D2 CTRL and EXP skin. White dotted lines separate the epidermis and hair follicles from the dermis. **g** – Quantification of the Pdpn+ area in the upper dermis, from **f**. Unpaired two-tailed MannWhitney U-tests, n=3-4 mice per condition, the dots represent means of 3 measurements per mouse. Data are presented as mean +/- SD. **h** – Immunofluorescence for Tnc (red), Itgb4 (white), Vim (green) and Hoechst (blue) on D4 CTRL and EXP skin. Dotted white lines delimit the epidermis and hair follicles. **i** – Quantification of the Tnc+ area in the upper dermis from **h**. Unpaired two-tailed MannWhitney U-tests, n=3-6 mice per condition, averages of 6 measurements per mouse are plotted as dots. Data are presented as mean +/- SD. All scale bars=50 μm.

### Fibroblasts undergo a spatial reorganisation during stretch-mediated skin expansion

Given that we identified transcriptional changes in the fibroblast in response to stretching, we wanted to understand whether these changes were associated to differences into fibroblasts appearance. To gain a phenotypic overview of the morphological changes occurring to the fibroblasts during expansion, we traced stromal cells positive for Pdgfra. We used *PdgfrαCre-ERT2;Rosa26^mTmG^* reporter mice to visualise the cell membranes of fibroblasts (Fig. 4a,b and Supplementary Fig. 4a). Using a high dose of induction to study fibroblasts at the population level, we were able to recombine on average 67% of Pdgfrα+ cells, as assessed by flow cytometry (Supplementary Fig. 4b-c). Immunofluorescence staining for Pdgfrα on *PdgfrαCre-ERT2;Rosa26^mTmG^* sections revealed good overlap between the signal of the immunofluorescence for Pdgfrα and the signal of the green fluorescence protein (GFP) form the recombined endogenous Pdgfrα-GFP (Supplementary Fig. 4d). We focused on the upper dermis region for subsequent morphological analyses (as defined in Supplementary Fig. 4a) in Fig. 4c-f and Supplementary Fig. 4e-h, unless otherwise stated, to try to exclude any potential confounding contributions from fascia progenitors^35,36^. Upon stretching, the surface area of the skin increases, as previously measured^15^. However, rather than a thinner dermis, we observe that the dermis thickness remains the same post-expansion (Supplementary Fig. 4e, f), suggesting that new stroma is generated to compensate for the larger area of skin produced. Indeed, the total fibroblast volume in the upper dermis, calculated based on the signal of the Pdgfrα-GFP, increased at D2 and D4 during expansion (Fig. 4b, c). We also observed changes in the organisation of the Pdgfra-GFP signal. To quantify these morphological changes, we took advantage of the ImageJ plugin FracLac that measures complexity of morphology^37^. At D2 and D4, the Pdgfrα-GFP signal had a lower fractal dimension score, indicating a less complex arrangement of the labelled cells (Fig. 4d, e). The overall cell density in the dermis was significantly increased at D4 (Fig. 4f), whilst cell death, as assessed by cleaved Caspase-3 staining, was unchanged (Supplementary Fig. 4g,h).

**Figure 4.**
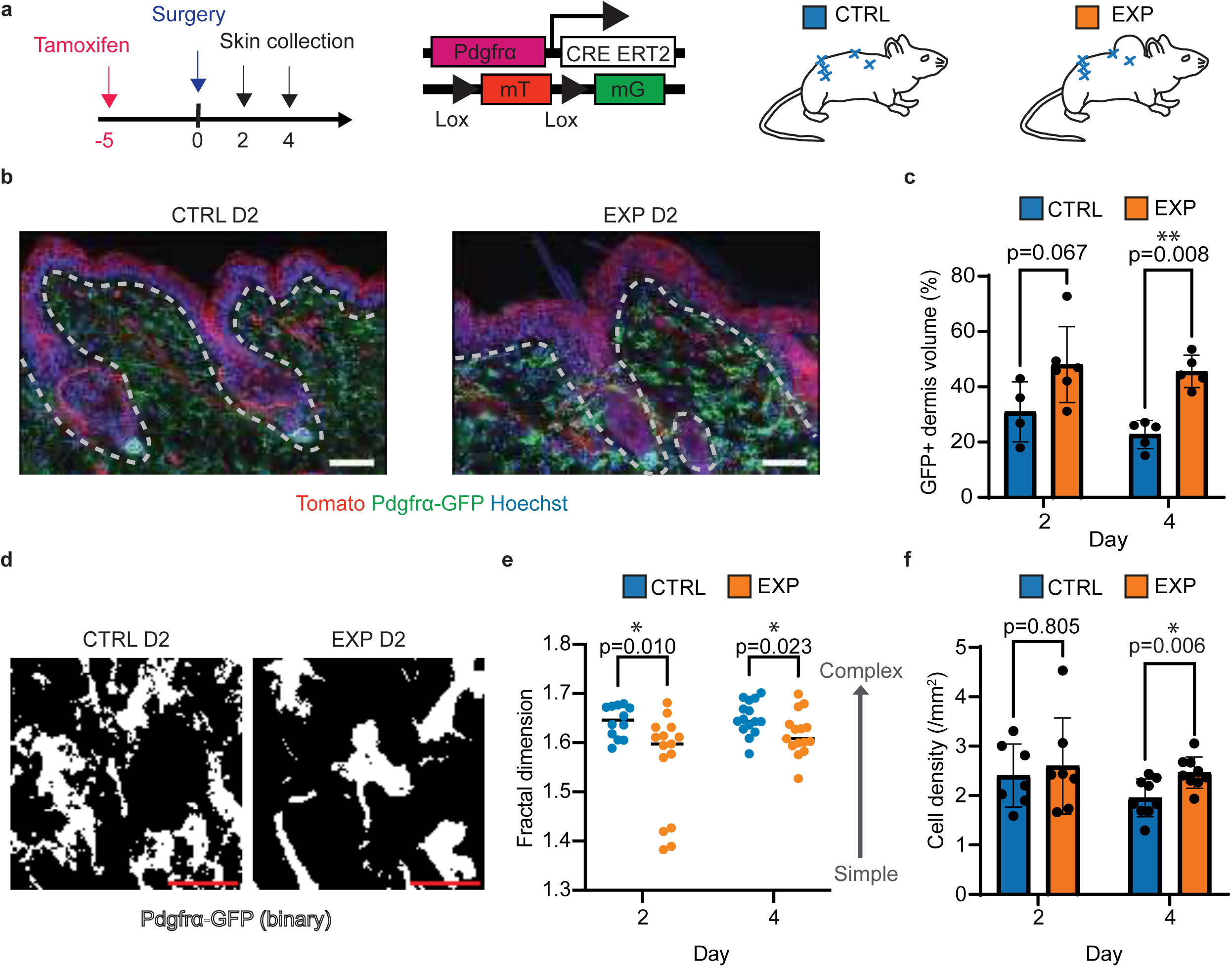
Fibroblasts’ population-level morphological changes upon expansion. **a** – Schematic of the experimental procedure and mouse reporter line used. **b** – Fluorescence for Tomato (red), Pdgfrα-GFP (green) and Hoechst (blue) in D2 CTRL and EXP skin. Dotted lines are used to delimit the epidermis and hair follicles from the dermis. Scale bars=50 μm. **c** – Quantification of Pdgfrα-GFP volume in the upper dermis based on **b**. Unpaired two-tailed Mann-Whitney U-tests, n=4-6 mice per condition, dots represent means of 3 measurements per animal. Data are presented as mean +/- standard deviation (SD). **d** – Representative binary images of Pdgfrα-GFP signal (white) from **b**, used for fractal dimension quantification. Scale bars=20 μm. **e** – Fractal dimension quantification of fibroblast organisation complexity. Unpaired twotailed Mann-Whitney U-tests, n=4-6 mice per condition, 3 measurements per mouse are plotted. Data are presented as individual points and median. **f** – Quantification of total cell density in the upper dermis. Unpaired two-tailed Mann-Whitney U-tests, n=7-9 mice per condition, averages of 3 measurements per animal are plotted. Data are presented as mean +/- SD.

### The volume of individual fibroblasts and the clone size increases upon stretching

To investigate whether the global rearrangements seen at the population level were reflected on the morphology and behaviour of single fibroblasts, we subsequently used the *PdgfrαCreERT2; Rosa26^Confetti^* reporter line recombined at a low level, to visualise and trace individual fibroblasts (Fig. 5a-d and Supplementary Fig. 5a). We focused on CFP+, YFP+ and RFP+ cells to leverage information not only on clone size, but also on fibroblast volume and shape (clone colour frequency per mouse in Supplementary Fig. 5b). D0 mice did not undergo surgery and were used as control to assess clonality. We observed a range of clone sizes (Fig. 5b-e and Supplementary Fig. 5c-f) with a significant increase in clone sizes at D4 post-surgery in the EXP condition (Fig. 5e) and with a clone persistence, calculated as the number of clones per analysed tissue area, that was stable during the time course of the experiment (Supplementary Fig. 5f). The constant persistence with the increase in clone size suggested that more fibroblasts were generated during the tissue expansion procedure.

**Figure 5.**
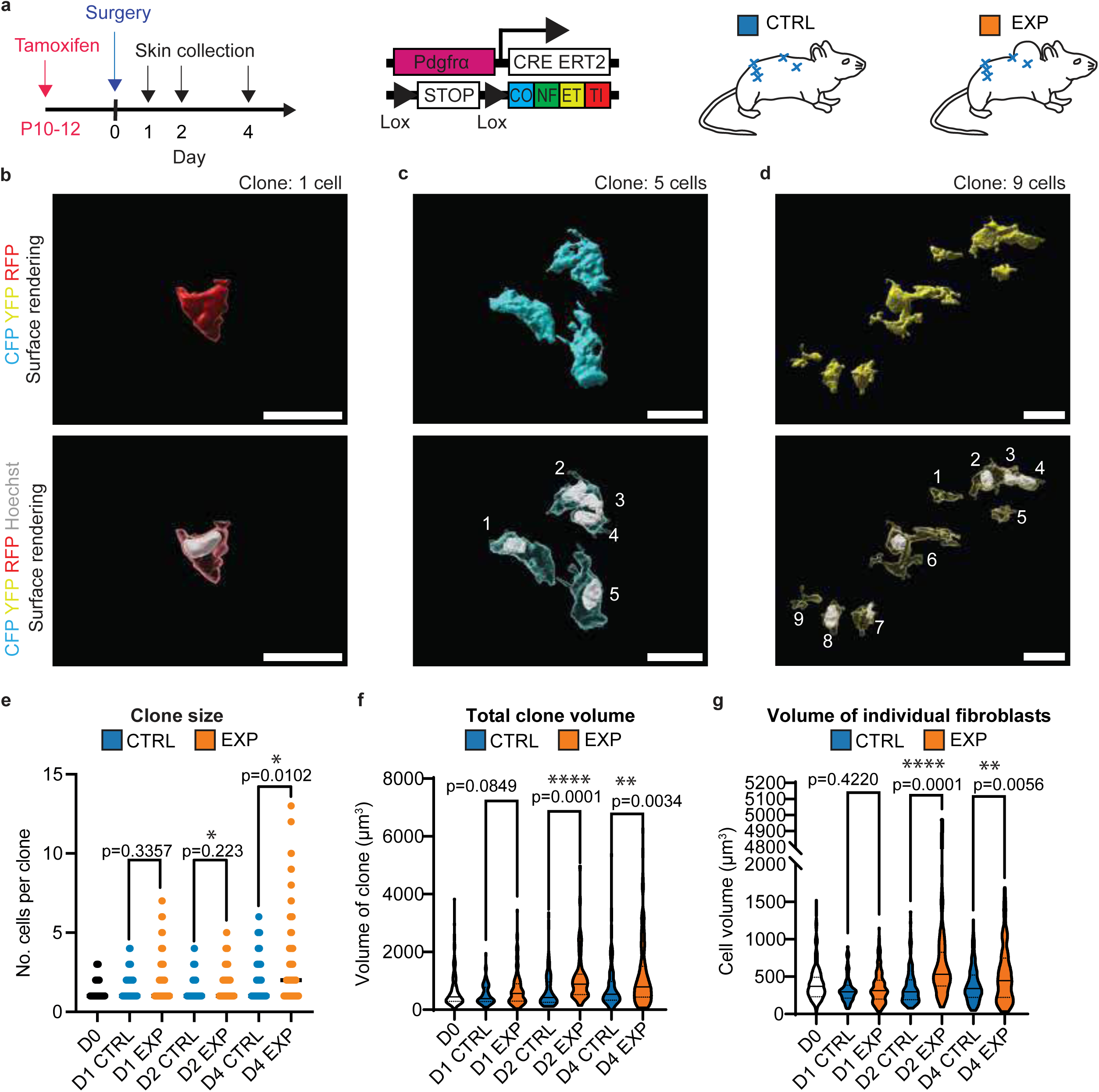
Individual fibroblast morphological changes upon expansion. **a** – Schematic of the experimental procedure and mouse reporter line used. **b-d** – Representative images showing surface rendering of Confetti CFP, YFP and RFP (cyan, yellow and red) and Hoechst (white) fluorescence signal from *PdgfrαCre-ERT2; Rosa26^Confetti^* wholemounts, generated in Imaris. Scale bars=20 μm. In the bottom images the numbers indicate the count of individual cell per clone. **e** – Quantification of clone sizes. Unpaired two-tailed Mann-Whitney U-tests, n=3-4 mice per condition (number of measurements for D0=99, D1 CTRL=73, D1 EXP=124, D2 CTRL=59, D2 EXP=77, D4 CTRL=118, D4 EXP=125). **f** – Quantification of the volume of clones. Unpaired two-tailed Mann-Whitney U-tests, n=34 mice per condition (number of measurements for D0=99, D1 CTRL=73, D1 EXP=124, D2 CTRL=59, D2 EXP=77, D4 CTRL=118, D4 EXP=125). Data are presented as violin plots showing the median and upper and lower quartiles. **g** – Quantification of the volume of individual fibroblasts. Unpaired two-tailed Mann-Whitney U-tests, n=3-4 mice (number of measurements for D0=166, D1 CTRL=90, D1 EXP=198, D2 CTRL=95, D2 EXP=107, D4 CTRL=175, D4 EXP=180). Data are presented as violin plots showing the median and upper and lower quartiles.

It has previously been found that upon loss of fibroblasts due to ageing or wounding, remaining fibroblasts are able to compensate for the loss by extending their membranes into depleted areas^38^. We sought to assess whether fibroblasts changed shape upon expansion and found the shape of fibroblasts (assessed by sphericity) was highly heterogeneous in both CTRL and EXP skin (Supplementary Fig. 5g), however the volumes of clones and individual fibroblasts were increased 2 and 4 days after stretching (Fig. 5f, g). Our data indicate that fibroblasts increase their volume on an individual level upon expansion to compensate for the generation of extra dermis, but also highlight an increase in the clone size, suggesting that an increased number of fibroblasts were produced in response to stretching.

### Downregulation of ECM genes and induction of fibroblast proliferation in expansion

Given the morphological changes happening to fibroblasts upon expansion, we decided to look more closely at the transcriptional changes occurring in the fibroblasts that could be related to their secretory and structural function. In general, in the scRNAseq data, fibroblasts showed downregulation of genes associated with main components of the ECM such as *Col1a1*, *Col1a2*, *Col3a1*, *Col4a1*, *Fn1*, *Lum*, *Fbn1, Eln, Sparc* and *Fbln1* (Fig. 6a, Supplementary Fig. 6a-c and Supplementary Table 5). To investigate in more depth the transcriptional response of the fibroblasts to stretching and expansion, we performed bulk RNA sequencing of fibroblasts, sorted as described previously (Supplementary Fig. 3d). At D1, 440 genes were upregulated and 882 downregulated, at D2, 805 genes were upregulated and 538 downregulated, and at D4, 843 genes were upregulated and 703 downregulated compared to the corresponding controls (Supplementary Fig. 6d-f and Supplementary Table 6). Gene ontology (GO) analysis revealed a downregulation of collagen trimer related genes (Fig. 6b). The downregulation of fibrillar collagen I during expansion was confirmed by multiphoton confocal microscopy, via the acquisition of second harmonic generation (SHG) signal at D4 (Fig. 6c, d). As lower levels of collagen have previously been associated with fibroblast proliferation during development^39^, we investigated the fibroblast proliferation during expansion. In the scRNAseq data, the increase expression of genes associated with proliferation such as *Pcna*, *Nasp*, *Slbp* and *Ctcf* (Fig. 6e) led to the prediction that the proportion of fibroblasts in S phase was increased at all three time points in EXP (Fig. 6f). Additionally, we saw upregulation of genes involved in mitosis in the bulk RNAseq analysis (Fig. 6g and Supplementary Table 7). More than 20 upregulated genes were related to sterol biosynthesis (Supplementary Fig. 6g-j). Given the general membrane remodelling of the entire fibroblast population and the increase in volume of the individual fibroblasts during expansion, the increase in genes related to sterol biosynthesis might be the consequence of cell membrane remodelling. Other upregulated GO terms were related to ribosome biogenesis genes (Supplementary Fig. 6k), changes in Wnt signalling-related genes (Supplementary Fig. 6l and Supplementary Table 8), as well as upregulation of YAP/TAZ targets (Supplementary Fig. 6m). The YAP/TAZ pathway is an important pathway in mechanosignalling^40^ and is upregulated in keratinocytes upon expansion^15,17^. The nuclear relocalisation of YAP was confirmed by immunostaining at D4, as well as the upregulation of YAP target gene *Ctgf* by qPCR (Supplementary Fig. 6m-p).

**Figure 6.**
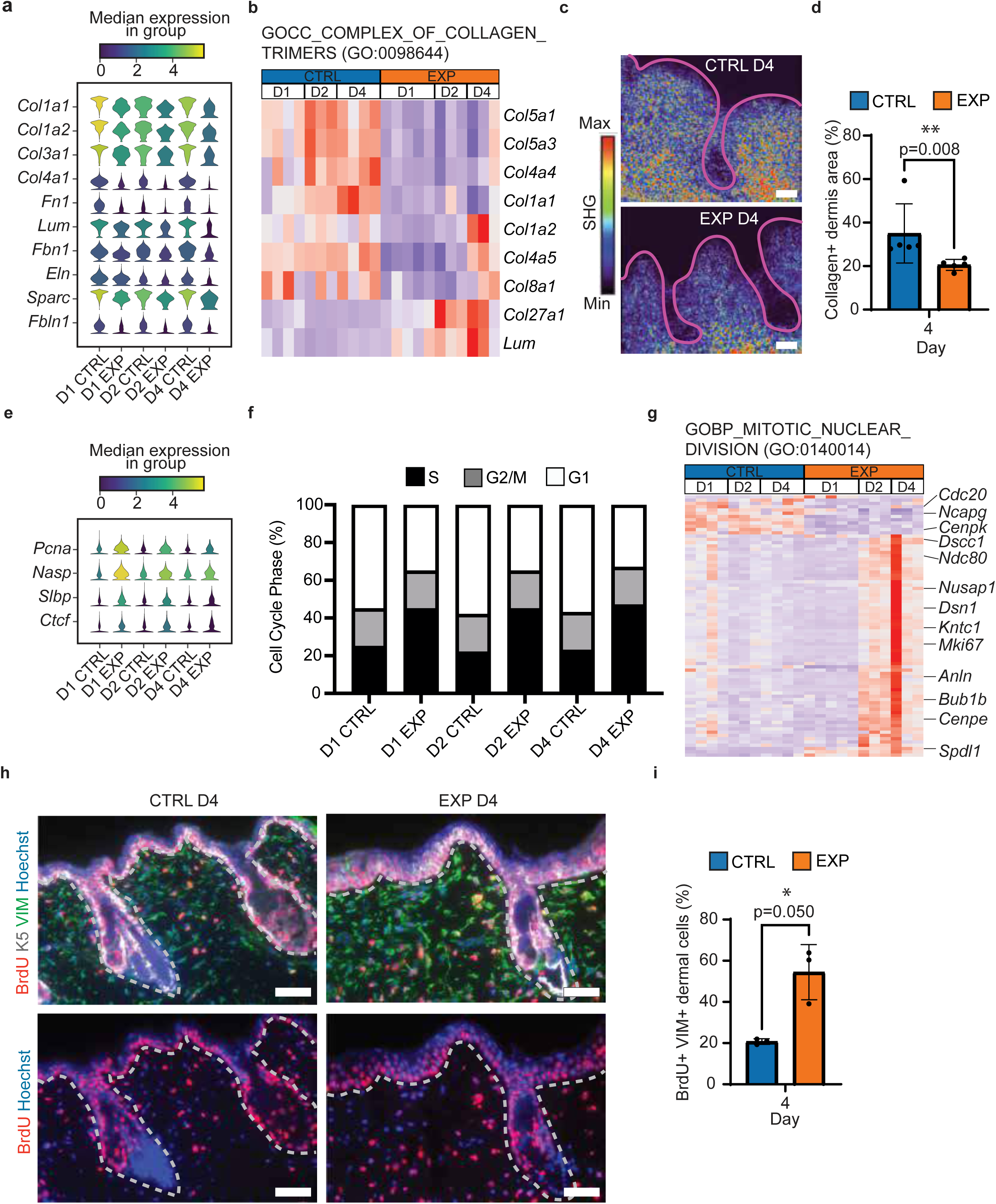
Downregulation of ECM genes upon expansion and upregulation of proliferation. **a** – Stacked violin plot of differentially expressed ECM genes in scRNAseq of fibroblasts and fibroblast-like cells. **b** – Heatmaps of differentially expressed genes (DEG) from bulk RNAseq analysis of sorted Pdgfrα+ Cd34+ fibroblasts. Relative gene expression (Rel. gene exp.) is shown for the indicated gene ontology (GO) term, with red indicating high expression, and blue indicating low expression. **c** – Second harmonic generation (SHG) imaging for visualization of collagen I fibres at D4. Pink lines show the location of the epidermis and hair follicles. The signal intensity is colour coded with red indicating high intensity, blue indicating low intensity, and black indicating no signal. **d** – Quantification of SHG area in the upper dermis from **c**. Unpaired two-tailed MannWhitney U-test, n=5 mice per condition, the averages of 3 measurements per animal are plotted as dots. Data are presented as mean +/- SD. **e** – Stacked violin plot of differentially expressed genes associated to proliferation in scRNAseq of fibroblasts and fibroblast-like cells. **f** – Cell cycle prediction analysis in fibroblasts from scRNAseq data. **g** – Heatmaps of DEG from bulk RNAseq analysis of sorted Pdgfrα+ and Cd34+ fibroblasts. Relative gene expression (Rel. gene exp.) is shown for the indicated gene ontology (GO) term, with red indicating high expression, and blue indicating low expression. **h** – Immunofluorescence for BrdU (red), Krt5 (white), Vim (green) and Hoechst (blue) on D4 CTRL and EXP skin. White lines delimit the dermis from the epidermis and hair follicles. **i** – Quantification of Vim+ and BrdU+ cells in the upper dermis from **h**. Unpaired one-tailed Mann-Whitney U-test, n=3 mice per condition, averages of 3 measurements per animal shown. Data are presented as mean +/- SD. Scale bars=50 μm.

Considering that the identified GO terms could be associated with the cellular responses to cell cycle changes, we sought to assess proliferation in the dermis by assessing 5-Bromo-2ʹdeoxyuridine (BrdU) incorporation (Fig. 6h, i). The percentage of BrdU+ cells increased twofold during expansion at D4 (Fig. 6i), indicating that fibroblasts re-enter the cell cycle. Together, these data suggest that fibroblasts respond to stretching by decreasing ECM production and increasing proliferation. Proliferation and lower expression of collagens are more characteristic of younger fibroblasts^39,41^, which enter quiescence after post-natal day 10 (P10)^27,42^ in the mouse dorsal skin. Altogether, these data suggest that stretching reprograms the fibroblasts to a more immature state.

### Fibroblast downregulation of collagens and upregulation of a proliferation signature play a role in promoting keratinocyte self-renewal

Keratinocytes and fibroblasts interact in vivo and in vitro^41,43–45^. Upon stretching, increased self-renewal of epidermal stem cells is observed^15^ and we hypothesised that fibroblasts provide signals required for this shift in cell fate. To confirm the relevance of the fibroblasts in the keratinocyte proliferative potential, we decided to test this interaction in a wellestablished in vitro co-culture system of fibroblasts and human keratinocytes. Since the 1970s, ex vivo expanded human primary keratinocytes have been extensively used in clinical settings for treating burn patients, corneal transplantation, and gene therapy for genetic skin disorders like epidermolysis bullosa^46–49^. For long-term skin regeneration to be successful, a specific number of self-renewing epidermal stem cells must be included in the graft. The selfrenewal capacity of these cells is strongly influenced by the culture method, which relies on the use of lethally irradiated embryonic fibroblasts as a feeder layer within a specialized culture medium^50^. Notably, normal human keratinocytes cultured without a feeder layer, whether in fresh medium (FM-noFL, grey bars) or feeder-layer-conditioned medium (CMnoFL, green bars), completely lose their proliferative potential after just a few passages (Supplementary Fig. 7a-c) and reduced their markers of stemness such as P63 and FOXM1^50^ (Supplementary Fig. 7d). In contrast, co-culture systems support the maintenance of proliferative potential for dozen of passages^50^. Therefore, this co-culture system can be considered a model of expansion, as it promotes and sustains self-renewal of epidermal stem cells, and a model to further elucidate the role of fibroblasts in supporting stem cells selfrenewal. We thus investigated whether ECM levels influence the clonogenic potential of keratinocytes in the in vitro system. We used two types of feeder layers: standard quality (SQ) feeder layers (SQ-FL) and low quality (LQ) feeder layers (LQ-FL) (see Methods). To confirm the differences between the two feeder layers, we performed mass spectrometry analysis on proteins collected from the two types (Fig. 7a and Supplementary Fig. 7e). The analysis showed that SQ feeder layers had lower expression of ECM-associated proteins, including collagens, and higher expression of proteins linked to cell proliferation compared to LQ feeder layer (Fig. 7a-c and Supplementary Fig. 7f-j), which mimics the fibroblast signature in the in vivo EXP condition. We confirmed these differences in COL4A1 and FN1 by western blot analysis (Fig. 7d) and by immunostaining (Fig. 7e, f and Supplementary Fig. 7k). We subsequently cultured normal human keratinocytes on these two types of feeder layers. Over serial cultivation, keratinocytes cultured on LQ feeder layers exhibited a reduced clonogenic potential, an increased number of aborted colonies, and fewer cells at subconfluence compared to those cultured on SQ feeder layers (Fig. 7g-j). Examination of stem cell-related markers at both the RNA and protein levels revealed a significant decrease in their expression, along with a slight increase in differentiation markers such as Involucrin (IVL) (Fig. 7k, l). This data confirmed that the proliferation capacity of human keratinocytes is dependent on the characteristics of the fibroblasts they are grown on. Together these data indicate that downregulation of ECM and upregulation of markers associated to a proliferative signature in fibroblasts play a role in promoting the self-renewal of epidermal stem cells.

**Figure 7.**
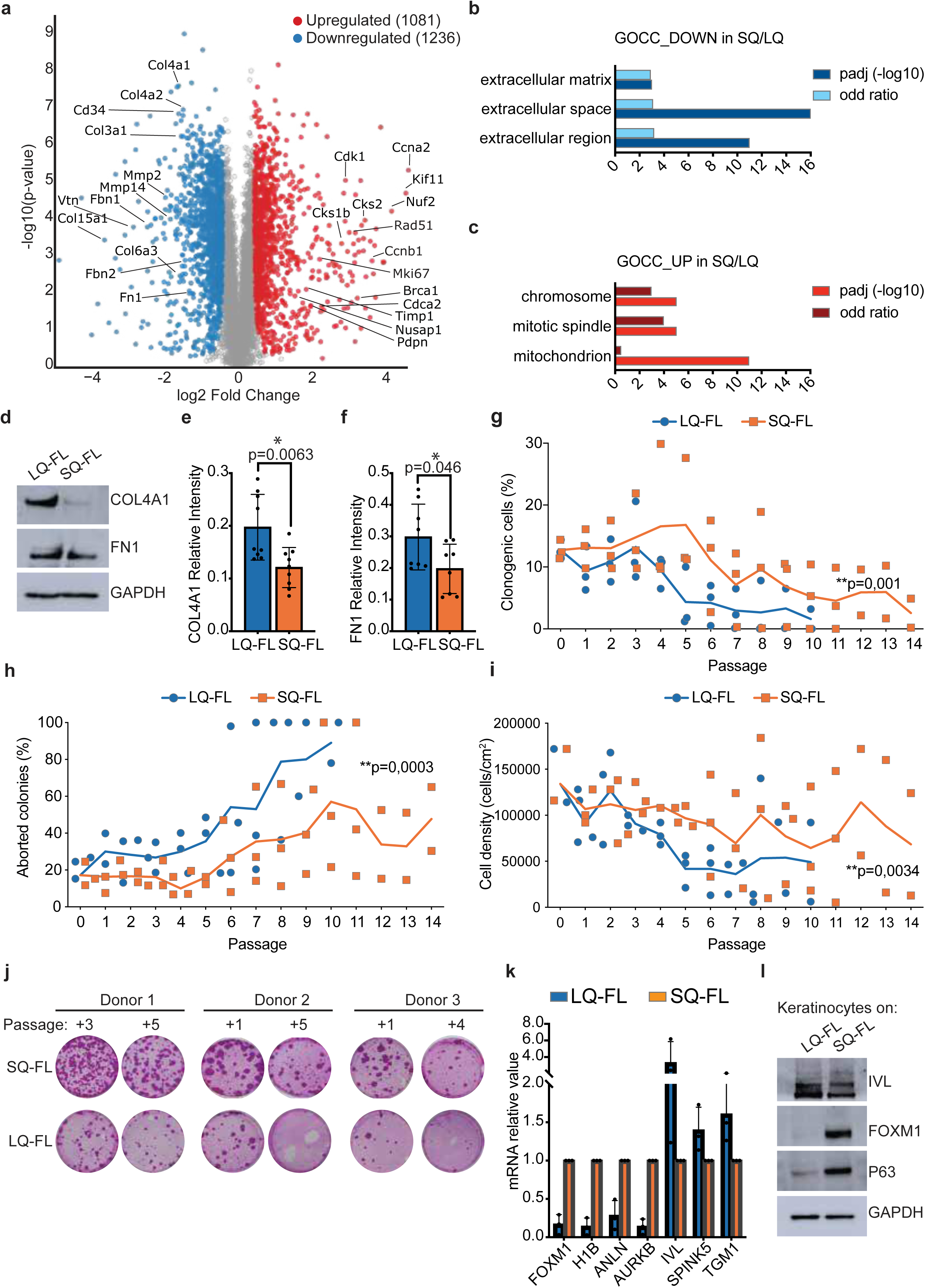
Fibroblast-keratinocyte interactions are important for keratinocyte proliferation. **a** – Volcano plot representation of the proteins identified by mass spectrometry analysis of standard quality feeder layers (SQ-FL) compared to low quality feeder layers (LQ-FL). Significantly downregulated proteins are shown in blue, significantly upregulated proteins are shown in red. x-axis shows fold-change (log2(ratio)), y-axis shows proteins’ significance (as Log10 of the corresponding q-value). **b** – Graph showing downregulated cellular component terms in SQ-FL as compared to LQ-FL. p values are calculated with one-sided Fisher’s Exact test and corrected for multiple tests with the Benjamini–Hochberg method. Data presented as –log10 of the corrected p value (dark blue bars) and corresponding fold changes (light blue bars). **c** – Graph showing upregulated cellular component terms in SQ-FL as compared to LQ-FL. Pvalues are calculated with one-sided Fisher’s Exact test and corrected for multiple tests with the Benjamini–Hochberg method. Data presented as –log10 of the corrected p value (red bars) and corresponding fold changes (dark red bars). **d** – Western analysis of total cell extracts from SQ-FL and LQ-FL. One out of three representative experiments is shown. **e, f** – Quantification of staining intensity related to Supplementary Fig. 7k for Col4a1 (**e**) and Fn1 (**f**). . P value calculated by unpaired *t*-students test. Data are presented as mean +/- SD. **g**, **h** – Serial cultivation of normal human keratinocytes on LQ-FL (blue line) and SQ-FL (orange line); Percentage of clonogenic cells was calculated as the ratio between grown colonies and plated cells (**g**). Percentage of aborted colonies was calculated as the ratio between the colonies scored as aborted and the number of clonogenic cells (**h**). Two-tailed paired Student t-test, n=3 independent biological replicates. Data are presented as mean +/− SEM. **i** – Total amount of cells obtained at subconfluence during serial cultivation of normal human keratinocytes on LQ-FL (blue line) and SQ-FL (orange line). Two-tailed paired Student t-test, n=3 independent biological replicates. Data are presented as mean +/− SEM. **j** – Representative images related to **g**-**i** showing the indicator dishes derived from NHK cultures, coltured on SQ-FL or LQ-FL at the indicated passage. **k** – qRT-PCR quantification of the mRNA levels of clonogenic markers (*FOXM1, H1B, ANLN, AURKB*) and differentiation markers (*IVL, SPINK5, TGM1*) on three human primary keratinocytes cultures grown on SQ-FL (orange bars) and LQ-FL (blue bars). Two-tailed paired Student t-tests, *FOXM1* *p=0.040, *H1B* *p=0.033, *ANLN* p=0.094, *AURKB* *p=0.031, *IVL* p=0.327, *SPINK5* p=0.327, *TGM1* p=0.327. n=3 independent biological replicates. Data are presented as mean +/− SD. **l** – Western analysis of total cell extracts from cultures generated by human primary keratinocytes grown on SQ-FL and LQ-FL. One out of three representative experiments is shown.

## Discussion

We have generated a transcriptomic atlas of stretch-mediated expansion in mouse back skin. We have demonstrated that fibroblasts sense mechanical stretching in vivo and ultimately respond by decreasing their production of ECM and re-entering the cell cycle, reverting to a “young” fibroblast cell state^39,42,51^. Finally, we have shown that this behaviour is important in promoting proliferation of epidermal cells, potentially coupling the response of dermal and epidermal compartments during expansion.

Fibroblast heterogeneity has been described in many contexts^23,52^. Two main fibroblast lineages, papillary and reticular, have been identified in mouse^42,53^ and in human^54,55^. Fibroblast transcriptional heterogeneity has also been described previously, in early development^56^, during homeostasis in mouse^21^, as well as in human skin^57^, and in other contexts, such as wound repair^25,37,58–60^, cancer^61^, ageing^26^ and UV damage^62^. Here we show transcriptional changes during expansion, as well as changes in protein expression. In recent years, spatial transcriptomics datasets of skin have been published^63,64^. Here, we have coupled scRNAseq and spatial transcriptomics to map the locations of fibroblast subtypes and show that fibroblast cellular heterogeneity is associated with specific spatial niches.

ECM plays an important role in cellular behaviour in various contexts, such as development and tumour initiation. Fibroblasts are proliferative in prenatal skin and enter a quiescent state in postnatal skin, which coincide with an increase in collagen deposition^39^. ECM can also influence epithelial progenitor cell behaviour, as observed during postnatal development of sebaceous glands^65^. Collagen levels can affect tumour initiation as well^66^. We observed lower expression of extracellular matrix genes, including collagens, and decreased mature collagen fibres at the protein level in the in vivo EXP condition. From the scRNAseq data we observed increased cellular proliferation in the expanded condition. These findings coincide with an increase in clone sizes at D4 post-surgery in the EXP condition via lineage tracing. The subsequent BrdU incorporation analysis further confirmed this, indicating that the increase in cell density in the dermis is due to proliferation of fibroblasts. However, we cannot rule out that there is also migration of cells from the fascia^35,36^ as we did not trace these populations.

The coordination of different cell types is essential for maintaining the skin’s architecture, integrity, and its barrier function. Many interactions have been described between cell populations in the skin during homeostasis^63,67–69^ and in wound healing^70^. Aberrant interactions can lead to diseases such as psoriasis^71^. In the context of stretching, it has been reported that macrophages provide signals that activate stem cells involved in hair regeneration^16^. Crosstalk can also be indirect, via ECM remodelling^72^. For example, basement membrane remodelling is essential during mouse embryogenesis and interaction between the ECM and cells is required for cell polarization and morphogenesis^72^. It has also been reported that lower substrate stiffness results in increased proliferation of keratinocytes in culture^73–75^. Here we identify the fibroblasts transcriptional changes associated with keratinocyte proliferation in both in vivo and in vitro expansion.

Understanding the interactions required for stretch-mediated expansion could allow for increased rate of growth by targeting pathways involved, thus reducing the months of discomfort patients experience with expanders in place. This study paves the way for future research into the roles of different cell types in coordinating the maintenance of tissue architecture upon expansion. In addition, current in vitro production of grafts for clinical application is limited to the growth of epidermal sheets^48,49^. A deeper understanding of how cell types interact to maintain tissue architecture will improve the production of organised full skin for regenerative therapies.

## Methods

### Animals

The experiments were approved by The Animal Experiments Inspectorate in Denmark, under license number 2021-15-0201-00792. All experiments were performed with male mice of age postnatal day (P) 60-P90, unless otherwise indicated. Mice had access to food and water ad libitum. *PdgfrαCre-ERT2* mice^76^ were obtained from Kim B. Jensen (University of Copenhagen) with permission from Brigid Hogan (Duke University Medical Center). *Rosa26^Confetti^* reporter mice^77^ were obtained from Kim B. Jensen (University of Copenhagen) and *Rosa26^mTmG^* reporter mice^78^ were obtained from Henrik Semb (University of Copenhagen). Where not otherwise specified, SWISS mice (Janvier Labs) were used.

### Expander experiments

Surgery was performed as described previously^15^. Briefly, mice were anaesthetised with 5% xylazine and 10% ketamine in phosphate-buffered saline (PBS) through intraperitoneal injection (200 μL/30g mouse). The back skin was shaved, and mice were injected subcutaneously with a local painkiller, carprofen (5mg/kg), near the incision site. Skin was disinfected with iso-betadine (Meda Pharma) prior to making an incision on the back skin close to the tail. A subcutaneous path from the incision site to the neck region was created using forceps and a self-inflating 4 mL sphere tissue expander (Osmed) was then placed subcutaneously near the neck region. The expanders were kept in place by 3 stitches and the incision site was stitched closed. For CTRL mice, the same procedure was followed minus the expander insertion.

### Lineage tracing experiments

To induce recombination in fluorescent reporter lines, mice were injected intraperitoneally with Tamoxifen (T5648, Sigma-Aldrich) dissolved in corn oil (10616051, Thermo Scientific Chemicals). *Pdgfrα-CreERT2;Rosa26^mTmG^* were injected once 5 days prior to surgery, approximately at P60, with 5mg of tamoxifen per 30g mouse. *PdgfrαCre-ERT2; Rosa26^Confetti^* mice were injected once between P10-P12 with 0.05mg of tamoxifen per 4g mouse. We term D0 mice as those that did not receive surgery.

### Proliferation experiments

Mice were injected twice a day following expander surgery with 5-Bromo-2ʹ-deoxyuridine (BrdU) (B5002, Sigma-Aldrich) dissolved in PBS. A dose of 2mg per 30g mouse was given each time. BrdU was also given in drinking water at a concentration of 1 mg/mL.

### Isolation of live skin cells and fibroblasts

Digestion of skin tissue was performed as previously described^27^. Briefly, skin was shaved, dissected, and minced into small pieces using scissors. The minced tissue was placed in a solution of 2.5 mg/mL Collagenase I (C0130, Sigma-Aldrich) and 0.01M CaCl_2_ (423525000, Thermo Scientific Chemicals) in HBSS (14025100, ThermoFisher Scientific) and incubated at 37°C for 2 hours on a shaker. The digestion was stopped by the addition of 5mM EDTA (15575020, ThermoFisher Scientific), the suspensions were filtered through 40 μm strainers and then centrifuged to obtain cell pellets.

For the isolation of live single cells for scSEQ, cells were stained with 4’,6-diamidino-2phenylindole, dihydrochloride (DAPI) (D1306, ThermoFisher Scientific, 0.3mM). Each sample was also stained with hashing antibodies (TotalSeq™-A03XX anti-mouse Hashtag, BioLegend, 1:100, Supplementary Table 9) for 20 minutes in order to be able to distinguish them once pooled for sequencing. Live cells were sorted into eppendorfs using a BD FACSAria II or BD FACSymphony S6 cell sorter (BD Biosciences), with a 70 μm nozzle.

To sort for fibroblasts for qPCR and bulk RNA sequencing, staining was performed for 30 minutes with primary antibodies, followed by washing with PBS, staining for 30 minutes with secondary antibodies (where required) and washing again. The antibodies used were Cd45PE (12-0451-82, Invitrogen, 1:1200), Cd31-PE (553373, BD Biosciences, 1:2000), Cd117-PE (553355, BD Biosciences, 1:4000), Epcam-PE (118205, BioLegend, 1:800), Pdgfrα-APC (17-1401-81, Invitrogen, 1:100), Cd34-Biotin (14-0341-82, Invitrogen, 1:50) and StrepAlexaFluor750 (S21384, Invitrogen, 1:400). Cells were also stained with 0.3mM DAPI. Sorting was performed using a BD FACSAria II sorter (BD Biosciences) with a 100 μm nozzle. Cells were sorted into Eppendorf tubes and flash frozen in liquid nitrogen.

### Single-cell RNA sequencing

Per sequencing round, we had 2 samples for library preparation – one sample containing pooled control cells and the other containing pooled expanded cells. Single cell libraries were generated using the Chromium Next GEM Single Cell 3’ Reagent Kits v3.1 (Dual Index) (10X Genomics) following the manufacturer’s protocol, with additional steps due to hashing (Supplementary Table 9) as described previously^79,80^. Briefly, a primer (HTO primer, 0.2 mM) was added during the amplification of cDNA to increase the amplification of hashtag barcodes. Following amplification, the hashtag cDNA was separated from the sample cDNA using SPRIselect beads (Beckman Coulter). 30ng of hashtag cDNA was then amplified using the Kapa HiFi HotStart Ready Mix (KK2601, Roche) plus SI-PCR primer (10x Genomics, 0.17 μM) and unique Illumina TruSeq index primers (Illumina, 0.17 μM). Hashtag-related primer sequences can be found in the Supplementary Table 9. All libraries were diluted to 4 nm. The control and expanded cDNA libraries were pooled with their respective hashtag libraries at a ratio of 95:5. Subsequently, these two were pooled at a 50:50 ratio per sequencing round. Sequencing was carried out with a P2 100 kit on a NextSeq2000 sequencer (Illumina).

### scRNAseq analysis

Sequencing reads were aligned to the mm10-2020-A reference genome using CellRanger (v.6.1.2) with default parameters. Filtered count matrices for each sample were subsequently loaded using Scanpy^81^ (v.1.9.3). Cell hashing (Supplementary Table 9) antibody demultiplexing was performed using Seurat^82^ (v.4.3.0), with hashtag-derived oligonucleotides (HTOs) normalized using centred log ratio (CLR) normalization. Demultiplexing was conducted with the HTODemux function, setting the positive quantile to 0.99. Quality control and preprocessing were performed on each individual sample with Scanpy. For each sample, cells passed the following criteria: had more than 500 detected genes, showed expression of fewer than 15% mitochondrial genes, and were not identified as outliers in other metrics using a median absolute deviation (MAD) threshold of 5. Doublets were identified using scDblFinder^83^ (v.1.8.0) in R (v.4.1.3) with default parameters. Cells classified as doublets were removed from further analysis. Normalization was performed by scaling the total counts to the same level for each cell, followed by log1p-transformation. Highly variable genes were selected across batches using Scanpy, taking into account the batch information with the cell-ranger method, retaining the top 5000 variable features. Principle component analysis was performed on normalized data to reduce dimensionality prior to batch correction. Data from different time points and conditions were integrated using HarmonyPy^84^ (v.0.0.9) to correct for batch effects. After this integration, the neighbourhood graph was computed based on the Harmony-corrected principal components for downstream analysis. Leiden clustering was performed at multiple resolutions, with 0.2 chosen for further analysis. Marker genes for each cluster were identified using a Wilcoxon rank-sum test as implemented in Scanpy’s rank_genes_groups function, requiring at least 25% of cells in the cluster to express the marker. Clusters were annotated using known marker genes and manual curation based on prior biological knowledge of mouse skin cell types. Fibroblasts identified in the initial clustering were subset and reclustered at multiple resolutions. Subclusters were annotated as distinct fibroblast subtypes, and marker genes were recalculated to refine subpopulation identification. Differential gene expression analysis was performed using the MAST method in Seurat to investigate differences between both conditions (CTRL vs. EXP), across time points, and for each time point separately. Cell cycle scoring was conducted using Seurat to assign cells into different cell cycle phases. Human cell cycle genes were converted to mouse orthologs using gprofiler2, and cell cycle phases were assigned using CellCycleScoring based on S and G2M phase markers.

### Spatial transcriptomics

Spatial transcriptomics was performed on sections of D2 control skin using the Xenium platform (10X Genomics) following the manufacturer’s protocol at KU Leuven Institute for Single Cell Omics (Leuven, Belgium). A custom panel of 95 genes plus the Xenium Mouse Tissue Atlassing panel (Supplementary Table 10) was run to detect mRNA transcripts. Skin from four mice was used, with 3 technical replicates.

### Spatial transcriptomics analysis

A custom probe set, combined with the Xenium Mouse Tissue Atlassing panel (Supplementary Tables 10), was designed based on our previously obtained scRNAseq data. A deep learning approach was used to prioritise cell type identification and capture within-cell-type expression variability, to optimise gene selection for spatial profiling (https://github.com/sifrimlab/Concrete-Geneselection). Slides were analysed using the Xenium analyser (10X Genomics) with instrument software version 2.0.1.0 and analysis version xenium-2.0.0.10, using default settings for cell segmentation, transcript decoding, and assignment. Filtered output files from the instrument’s pipeline were used for downstream analysis. Spatial transcriptomics data from 10X Xenium was processed using Scanpy (v.1.9.3). Filtered data was loaded using read_10x_h5(), and metadata was integrated. Quality control metrics were computed, and cells with fewer than 10 transcripts were filtered out. Data normalization was performed by scaling the counts with the normalize_total() function followed by log(1+x) transformation. scRNAseq cell type annotation mapping of the spatial data was performed using an adaptation of the PopV package^85^ (v.0.4.2), which incorporates multiple annotation methods such as scVI, scANVI,Scanorama, Harmony, and others^84,86,87^. In addition, two widely used annotation transfer tools, Tangram^88^ and DotR^89^, were applied as well both with default parameters for high resolution data. For integration 10,000 cells per reference cell type were sampled when possible. The final merged spatial dataset contained 85,344 cells and 474 genes. Consensus cell typing, which relies on combining the results of multiple annotation methods to increase accuracy and confidence, was then calculated, and cells for which two or more methods agreed on an annotation were annotated. To calculate fibroblast distances from the epidermis, the interfollicular epidermis was manually annotated. The minimum distance of each cell to the closest epidermis cell was then calculated using the cdist function from scipy.spatial.distance (SciPy v.1.14.1)^90^, applying the Euclidean distance metric. Distances were computed separately for each sample. Density plots were generated by dividing the spatial tissue area into bins, averaging the raw expression values for all spots within each bin, and normalizing these averaged values to a scale of 0 to 1 to allow comparability across samples for the same gene.

### Bulk RNAseq

RNA was extracted from sorted cells using the RNeasy® Micro Kit (74004, QIAGEN), following the manufacturer’s protocol. Ultra-Low Input RNAseq was performed using the Illumina platform (2×150bp) at GENEWIZ (Azenta Life Sciences, New Jersey, USA).

Analysis was performed using R. FASTQ files were aligned to the mouse genome mm39, and counts calculated using RSubread^91^. The counts table was filtered to include only genes where count numbers were 14 or more in at least 3 samples using edgeR^92^. Differential expression analysis was performed using the DESeq2 package^93^, keeping only differentially expressed genes with a p value of <0.05 and a fold change greater than 1.5 or less than -1.5. Gene set enrichment analysis was done using fgsea. Gene lists from gene ontology and Reactome terms were obtained from the MSigDB^94,95^ database using the msigdbr package. To generate heatmaps, gene lists from selected GO and Reactome terms were filtered to include only genes we found to be differentially expressed in our dataset. Heatmaps showing scaled normalized counts of these genes were generated using ComplexHeatmap^96^.

### qPCR

Following RNA extraction using the RNeasy® Micro Kit, cDNA synthesis was performed using the High-Capacity cDNA Reverse Transcription Kit (4368814, ThermoFisher Scientific) following the manufacturer’s protocol. The qPCR reaction was performed using the SolisFAST® SolisGreen® qPCR Kit (Solis BioDyne) according to manufacturer’s instructions and run on a QuantStudio™ 7 Pro Real-Time PCR System (Applied Biosystems). Primer sequences are: *B2m* CTCGGTGACCCTGGTCTTTC (5’-3’), GGATTTCAATGTGAGGCGGG (5’-3’); Cd34 AGGCTGATGCTGGTGCTAG (5’-3’), AGTCTTTCGGGAATAGCTCTG (5’-3’); Ctgf (Ccn2) GAAGCTGACCTGGAGGAAAA (5’-3’), ACTGGCAGAGTGGTGGTTCT (5’-3’); Fdft1 TCCCTGACGTCCTCACCTAC (5’-3’), CCCCTTCCGAATCTTCACTA (5’-3’); Gapdh TGCCAGCCTCGTCCCGTAG (5’-3’), CGGCCTTGACTGTGCCGTTG (5’-3’); Mmp13 CAGTCTCCGAGGAGAAACTATGAT (5’-3’), GGACTTTGTCAAAAAGAGCTCAG (5’-3’); Pdpn AGATAAGAAAGATGGCTTGC (5’-3’), AACAACAATGAAGATCCCTC (5’-3’); Sqle AGTTCGCTGCCTTCTCGGATA (5’-3’), GCTCCTGTTAATGTCGTTTCTGA (5’-3’); Timp1 CGAGACCACCTTATACCAGCG (5’-3’), GGCGTACCGGATATCTGCG (5’-3’); Tnc TCCCCAAGAGAATTTACAGCTACAG (5’-3’), AGATTCATAGACCAGGAGGTATCCA (5’-3’). Relative fold changes in gene expression of samples (run in triplicates) were calculated using the delta-delta Ct method^97^ (ΔΔCt) and normalized to the housekeeping gene *B2m*.

### Tissue preparation, cryosectioning and immunostaining

After dissection, skin samples containing endogenous fluorescence were fixed for 3 hours at room temperature (RT) with 4% PFA (P6148, Sigma-Aldrich) prior to washing and embedding in OCT. Samples without endogenous fluorescence were directly embedded in OCT. Blocks were flash frozen by submersion in Iso-Pentane (10121820, Fisher Chemical), which was cooled to -150°C using liquid nitrogen. Cryosectioning was performed with a HM 560 CryoStar (Thermo Scientific) or CM1950 (Leica Biosystems) at -25°C. Standard sections were cut at a thickness of 8 μm. For analyses of *Pdgfrα-CreERT2;Rosa^mTmG^* and SHG/collagen, wholemount sections of 50 μm were cut, and for *PdgfrαCre-ERT2; Rosa26^Confetti^* sections of 200 μm.

Standard tissue sections were fixed (if the tissue was not already fixed prior to embedding) for 10 minutes at RT with 4% PFA, followed by washing 3 times in PBS for 5 minutes. Tissues were blocked for 1 hour at RT with slide blocking buffer (1% BSA (BP9702-100), 5% horse serum (26-050-088, Gibco), 0.2% Triton X-100 (X100, Sigma-Aldrich) in PBS). Slides were incubated with primary antibodies overnight at 4°C. Slides were washed 3 times in PBS for 5 minutes, followed by incubation with secondary antibodies for 1 hour at RT. Slides were washed again in PBS. For BrdU staining, slides were incubated post-fixation in 1N HCl for 30 minutes at 37°C, followed by washing for 3 times in PBS for 10 minutes before continuing with the staining protocol as described. The following primary antibodies were used: anti-Cd104 (Integrin β4) (553745, BD Biosciences, 1:200), anti-Krt5 (905901, Biolegend, 1:1000), antiVimentin (ab92547, Abcam, 1:2000), anti-Vimentin (ab24525, Abcam, 1:500), anti-Tnc (AB19011, Sigma-Aldrich, 1:100), anti-Pdpn (ab11936, Abcam, 1:80), anti-Yap (14074, Cell Signalling Technology, 1:50), anti-Cd34 (ab81289, Abcam, 1:200), anti-BrdU (ab6326, Abcam, 1:200) and anti-cleaved Caspase-3 (9661, Cell Signalling Technology, 1:100). The following secondary antibodies (Jackson Immunoresearch) were used at a 1:400 dilution: anti-rabbit, anti-chicken, anti-rat conjugated to Rhodamine Red-X, Alexa Fluor 488, Alexa Fluor A647, or Cy5. Nuclei were stained with Hoechst (H3570, ThermoFisher Scientific, 1:1000).

Wholemount tissue sections were blocked for 1 hour at RT with wholemount blocking buffer (1% BSA, 5% horse serum, 0.8% Triton X-100 in PBS), followed by incubation with Hoechst (1:1000) for 3 hours at RT and washing with PBS 3 times for 5 minutes.

For both standard sections and wholemounts, cover slides were mounted with 2.5% 1,4Diazabicyclo[2.2.2]octane (D27802, Sigma-Aldrich) in Glycerol Mounting Medium (C0563, Dako).

### Mouse skin imaging

Z-stack images of wholemounts were acquired using laser-scanning confocal microscopes (LSM 780 (ZEISS), LSM 880 Airyscan (ZEISS) or Stellaris (Leica Biosystems)). To acquire images of collagens using second harmonic generation, a multiphoton microscope was used (SP8 Multiphoton (Leica Biosystems)). Images of standard 8 μm sections were acquired with a widefield microscope (DM5500 (Leica Biosystems)).

### Image analysis

#### PdgfrαCre-ERT2;Rosa26^mTmG^ analysis

Volume quantifications were performed with the image analysis software Imaris (Oxford Instruments) by masking the Pdgfrα-GFP signal to produce 3D surfaces. To account for the larger volume of skin in the expanded condition, GFP volumes from the EXP condition were normalized by multiplying by a factor of 2 at D2 and 2.15 at D4. These values correspond to the measured increase in expander radius from D0 to D2 and D0 to D4^8^.

#### PdgfrαCre-ERT2; Rosa26^Confetti^ analysis

Quantifications of cell volume and sphericity were performed in Imaris by masking the fluorescence signal to produce surfaces. For clonal analysis, clones were defined as surfaces within a 50 μm distance from each other. Surfaces with multiple nuclei were counted as multiple cells within the same clone. Incomplete cells were also counted if their volume exceeded 100 μm^3^. Clone persistence was normalized by dividing the number of clones per image by the area of skin imaged. The area of skin was determined by producing maximum intensity projections of z-stack images and measuring the area manually in ImageJ. Plots of individual fibroblast volume and sphericity show data only from individual cells (surfaces with 1 nucleus).

#### Other analyses

Quantifications of cell numbers were performed manually in ImageJ. Quantifications of signal area and fractal dimension using FracLac^37^ were also performed in ImageJ on binary images. Dermal thickness was measured using QuPath^98^.

### Human Tissues

All human tissue samples were obtained with informed consent for research use, adhering to Italian regulatory standards (Ethics Committee of the Area Vasta Emilia Nord, approval number 178/09 for healthy donor skin samples).

### 3T3-J2 Cell Line used as Feeder Layer

Mouse 3T3-J2 cells, originally provided by Professor Howard Green from Harvard Medical School (Boston, MA, USA), were cultured in DMEM with 10% gamma-irradiated adult bovine serum, penicillin–streptomycin (50 IU/ml), and 4 mM glutamine.

### Primary Human Cell Cultures from Healthy Donors

Primary human keratinocytes were seeded (2.5–3 x 10⁴/cm²) onto lethally irradiated 3T3-J2 cells (2.4 x 10⁴/cm²) whitin 24h from seeding and cultured at 37°C in a humidified atmosphere with 5% CO₂. The medium (KGM) consisted of a 2:1 mixture of Dulbecco’s modified Eagle’s medium (DMEM) and Ham’s F12 media, supplemented with 10% fetal bovine serum (FBS), 50 IU/ml penicillin–streptomycin, 4 mM glutamine, 0.18 mM adenine, 5 μg/ml insulin, 0.1 nM cholera toxin, 0.4 μg/ml hydrocortisone, 2 nM triiodothyronine (Liothyronine Sodium), and 10 ng/ml epidermal growth factor (EGF). Cells were serially passaged upon reaching subconfluency until they reached senescence.

Under the fresh medium (FM)-noFL condition, cells were cultured on plastic surfaces without co-culturing with 3T3-J2 cells, using fresh KGM medium. For the Conditioned Medium (CM)noFL condition, cells were cultured on plastic without co-culturing with 3T3-J2 cells but in KGM medium preconditioned for 24 hours by lethally irradiated 3T3-J2 cells (2.4 x 10⁴/cm²). In the SQ-FL condition, lethally irradiated 3T3-J2 cells (2.4 x 10⁴/cm²) were used as feederlayer within 24 hours from seeding. For the LQ condition, lethally irradiated 3T3-J2 cells (2.4 x 10⁴/cm²) were used as feeder-layer 4 days from seeding.

### Colony-Forming Efficiency, Population Doublings, Growth Rate, and Clone Size

The colony-forming efficiency of keratinocytes during each culture expansion was assessed by plating between 1000 and 5000 cells in parallel on indicator dishes and staining with rhodamine B after 12 days. The total number of colonies was expressed as a percentage of the cells initially seeded. Aborted colonies, consisting of large, flattened, terminally differentiated cells, were quantified as a percentage of the total colonies.

### Western Blot Analysis

Keratinocytes and fibroblasts were harvested by scraping in 1x RIPA buffer (Sigma Aldrich) with phosphatase and protease inhibitor cocktails (Thermo Fisher). Total protein concentrations were determined using Pierce BCA Protein Assay Kits (Thermo Fisher Scientific). Equal amounts of protein were separated on 4–12% NuPAGE Bis-Tris Gels or 3-8% NuPAGE Tris Acetate Gels and transferred onto nitrocellulose membranes (Millipore) at 100 V for 2 hours at 4°C. Membranes were blocked with Everyblot Blocking Solution (Bio-Rad). Primary antibodies were incubated overnight at 4°C or 2 hours at RT. The following primary antibodies were used: anti-IVL 1:5000 from Invitrogen (Ma1-25752, mouse), anti-FOXM1 1:1000 from Cell Signalling Technologies (D3f2b, rabbit), anti-P63α 1:1000 from Cell Signalling Technologies (4892s, rabbit), anti-hGAPDH 1:500 from Sigma-Aldrich (Zrb374, rabbit), antiCOL4 1:1000 from Rockland (600-101-MN4, Goat), anti-FN1 from Proteintech 1:10000 (66042-1-Ig, Mouse), anti-GAPDH 1:500 from Abcam (ab8245, mouse). Secondary antibodies were applied for 1 hour at RT. The following antibodies were used: goat pAb to rabbit IgG (HRP) 1:1000-5000 from Abcam (Ab205718), rabbit pAb to Goat IgG (HRP) 1:2500 from Abcam (Ab6741), and goat pAb to mouse IgG (HRP) 1:20000 from Abcam (Ab6789). Signals were detected using Clarity Western ECL Substrate (Bio-Rad) and visualised with a ChemiDoc system (Bio-Rad) using ImageLab software.

### Immunostaining and image analysis of feeder layer

Lethally irradiated 3t3-J2 cells were plated at 2.4 x 10⁴/cm² onto glass coverslips and fixed with PFA 3% for 10min at room temperature after 24h (SQ-FL) and 4 days (LQ-FL). Cells were then carefully washed with PBS. Permeabilization was performed in PBS/Triton-X 0.5% for 20min at RT. Blocking solution (FBS 5% + BSA 2% in PBS/Triton 0.1%) was added for 30min at 37°C. Primary antibodies were diluted in Blocking solution and added to the samples overnight at 4°C. The following primary antibodies were used: anti-Col4A1 (600-101-MN4, Rockland, 1:400), anti-Fn1 (66042-1-Ig, Proteintech, 1:1000). Secondary antibodie were diluted in Blocking solution and added to the samples for 1h at RT in the dark. The following secondary antibodies were used: donkey anti-goat IgG (H+L) Alexa Fluor 488™ (A11055, Invitrogen, 1:200) and donkey anti-mouse IgG (H+L) Alexa Fluor 568™ (A10037, Invitrogen, 1:2000). Cell nuclei were stained with DAPI (10236276001, Roche). Dako Fluorescence Mounting medium (S3023, Dako) was used to mount coverslips. Zeiss Axio Imager 2 with 20x objective was used to visualize fluorescent signals and acquire images. Quantification of the intensity of fluorescence was performed on randomic fields with the image analysis software ImageJ normalizing the mean gray value on the number of nuclei.

### RNA extraction and real-time qPCR on human samples

Total RNA was extracted from cells using the PureLink RNA Mini Kit (Thermo Fisher Scientific) to perform quantitative real-time PCR. cDNA was synthesized with the SuperScript VILO cDNA Synthesis Kit (Thermo Fisher Scientific). Real-time qPCR experiments were conducted in triplicate using retrotranscribed cDNAs with TaqMan Universal PCR Master Mix on the QuantStudio 12k Flex Real-Time PCR System (Thermo Fisher Scientific). Gene expression levels were normalized to GAPDH as the reference gene. The following TaqMan primers (Thermo Fisher Scientific) were used: ANLN Hs01122612_m1, AURKB Hs00945855_g1, FOXM1 Hs01073586_m1, GAPDH 4352665, H1B Hs00271207_s1, IVL Hs00846307_s1, SPINK5 Hs00928570_m1, TGM1 Hs00165929_m1. Data analysis was carried out using RQ Manager Software 1.2.2.

### Mass spectrometry (MS) sample preparation

SQ and LQ feeder layers were collected by scraping at 24h (SQ-FL) and 4 days (LQ-FL) after plating and pelleted. In two different experiments, we analysed 2 and 3 samples for each condition. Proteomic sample preparation was performed with the EasyPep MS Sample Prep Kit (Thermo Fisher Scientific) following the manufacturer’s instructions. For the first LC-MS experiment, Peptides were separated on a 50 cm μPAC NEO HPLC column (Thermo Scientific), using a Vanquish Neo (Thermo Fisher Scientific) UHPLC. Peptide separation was performed using a 53 min stepped gradient of 4-22.5% solvent B (0.1% formic acid in 80% acetonitrile) for 40 min, 22.5-45% solvent B for 13 min, using a constant flow rate of 350 nL/min. Column temperature was controlled at 50 °C. Upon elution, peptides were injected via a Nanospray Flex ion source and 10-uM emitter (Evosep) into a Tribrid Ascend mass spectrometer (Thermo Scientific). Spray Voltage was set to 2200(V). Data was acquired in data independent mode with the Orbitrap MS resolution 60.000 for full scan range 400-900m/z, AGC target was 250% and maximum injection time set to auto. 42 DIA scans with 12 Th width and 1 Th overlap spanning a mass range of 400-900 m/z were were acquired at 15,000 MS resolution, AGC target 1000% and maximum injection time of 27 ms. HCD fragmentation normalised collision energy (NCE) was set to 30%. For the second LC-MS experiment, peptides were separated on an Aurora (Gen3) 25 cm, 75 μM ID column packed with C18 beads (1.6 μm) (IonOpticks) using a Vanquish Neo (Thermo Fisher Scientific) UHPLC. Peptide separation was performed using a 50 min stepped gradient of 2-17% solvent B (0.1% formic acid in acetonitrile) for 33 min, 1725% solvent B for 11 min, 25-35% solvent B for 6 min, using a constant flow rate of 400 nL/min. Column temperature was controlled at 50 °C. Upon elution, peptides were injected via an EASY-Spray source into a Tribrid Ascend mass spectrometer (Thermo Scientific). Spray Voltage was set to 1700(V). Data was acquired in data independent mode with the Orbitrap MS resolution 60.000 for full scan range 400-900m/z, AGC target was 250% and maximum injection time set to auto. 42 DIA scans with 12 Th width and 1 Th overlap spanning a mass range of 400-900 m/z were were acquired at 15,000 MS resolution, AGC target 1000% and maximum injection time of 27 ms. HCD fragmentation normalised collision energy (NCE) was set to 30%.

### MS data analysis

MS files were processed using DIA-NN v.1.8 in directDIA mode with a library predicted from mouse Uniprot fasta file (UP000000589) covering 55,079 protein isoforms. Highly heuristic protein grouping and Match between runs (MBR) were turned on. Carbamidomethylation of cysteine was specified as fixed modification, Oxidation of methionine, acetylation at the protein N-terminus, and N-terminal methionine excision were set as variable modifications. Maximum missed cleavage was set to 1 and a maximum of 2 variable modifications were allowed. The minimum peptide amino acid length was 7. The protein groups and precursors were filtered at 1% FDR. All other settings were set as default.

All statistical analysis was performed using Proteomelit 1.0.0, an in-house python tool, based on the automated analysis pipeline of the Clinical Knowledge Graph^99^. Intensity values were log2-transformed and features with less than 70% of valid values in at least one group were removed. Remaining missing values were imputed using the MinProb approach using a width of 0.3 and downshift of 1.8^100^. Protein significance was assessed through two-tailed unpaired t-tests with permutation-based False Discovery Rate (FDR) correction for multiple hypothesis testing, where parameters were set to FDR 0.05, Fold-change (FC) 1, s0 1, and 250 permutations. Functional enrichment analysis was performed using Fisheŕs exact test and Benjamini-Hochberg correction (FDR 0.05).

### Statistical analysis

When not otherwise specified, all statistical tests were calculated using GraphPad Prism 10. Statistical significance was defined as p£0.05. The statistical tests used are indicated in the figure legends. For all experiments, normality was tested using D’Agostino & Pearson tests. When normality tests were not passed, nonparametric statistical tests were performed, namely Mann-Whitney U-tests. For human culture and western blot experiments, two-tailed Student t-tests were used. For mass spectrometry experiments, significance was calculated using Fisheŕs exact tests and correction for multiple tests was performed using the Benjamini-Hochberg method.

## Acknowledgements

We thank J. Martin Gonzalez and the Core Facility for Transgenic Mice for mice rederivation and support; the staff of the animal facility (AEM); K.B. Jensen for mice; A. Kalvisa, H. Wollmann, M. Michaut, and the reNEW Genomics Platform for support, technical expertise, and use of instruments; G. Dela Cruz, M. Maimets, and P. van Dieken, and the reNEW Flow Cytometry Platform for support, technical expertise, and use of instruments; J. Bulkescher, J. Viswalingam Bagge and A. Georgantzoglou and the reNEW Microscopy Facility for the help with microscopy and image analysis; D. Andrejeva and the Center for Health Data Science for assistance with bulk RNAseq analysis; T. Voet, K. Vandereyken, N. Vandermeulen and the KU Leuven Institute for Single Cell Omics (LISCO) for the performance of the 10x Genomics Xenium assays. Mass spectrometry based proteomic analysis and data analysis were performed by the Proteomics Research Infrastructure (PRI) at the University of Copenhagen (UCPH), supported by the Novo Nordisk Foundation (NNF) (grant agreement number NNF19SA0059305). We also thank Prof. Michele De Luca (UNIMORE) for providing the 3T3-J2 cells and Prof. Graziella Pellegrini (UNIMORE) for providing human primary epidermal keratinocytes. We are grateful to all the other members of the Aragona lab for fruitful discussions and members of reNEW for constant feedback. Work in the Aragona Lab was funded by the Leo Foundation Research Grant (LC-OC-23001145), the Independent Research Fund Denmark (DFF) (1030-00100B) and the Novo Nordisk Foundation (NNF20OC0064992). C.A.S was supported by the NNF Copenhagen Bioscience PhD program (NNF20SA0035584). The Novo Nordisk Foundation Center for Stem Cell Medicine is supported by the Novo Nordisk Foundation (NNF21CC0073729).

## Author contributions

C.A.S and M.A. designed the experiments and performed data analysis; C.A.S. performed most of the experiments; L.B. helped with experiments; I.K. helped with mice; C.A.S. performed bioinformatic analysis of the bulk RNAseq data; C.A. and A.S. performed all other bioinformatic analyses. G.A.G., M.P.P. and E.E. performed human in vitro experiments. C.A.S and M.A. wrote the manuscript with comments and suggestions from E.E. and A.S. M.A. supervised the work.

## Data and materials availability

Data supporting the findings of this study are available within the article (and its Supplementary tables) and from the corresponding author, Mariaceleste Aragona, on reasonable request. The accession number for the scRNAseq data in GEO is GSE285079. The accession number for the bulk RNAseq data in GEO is GSE284808. The mass spectrometry proteomics data have been deposited in Open Access on the FairDom repository at https://fairdomhub.org/projects/462.

---

**Supplementary Figure 1.**
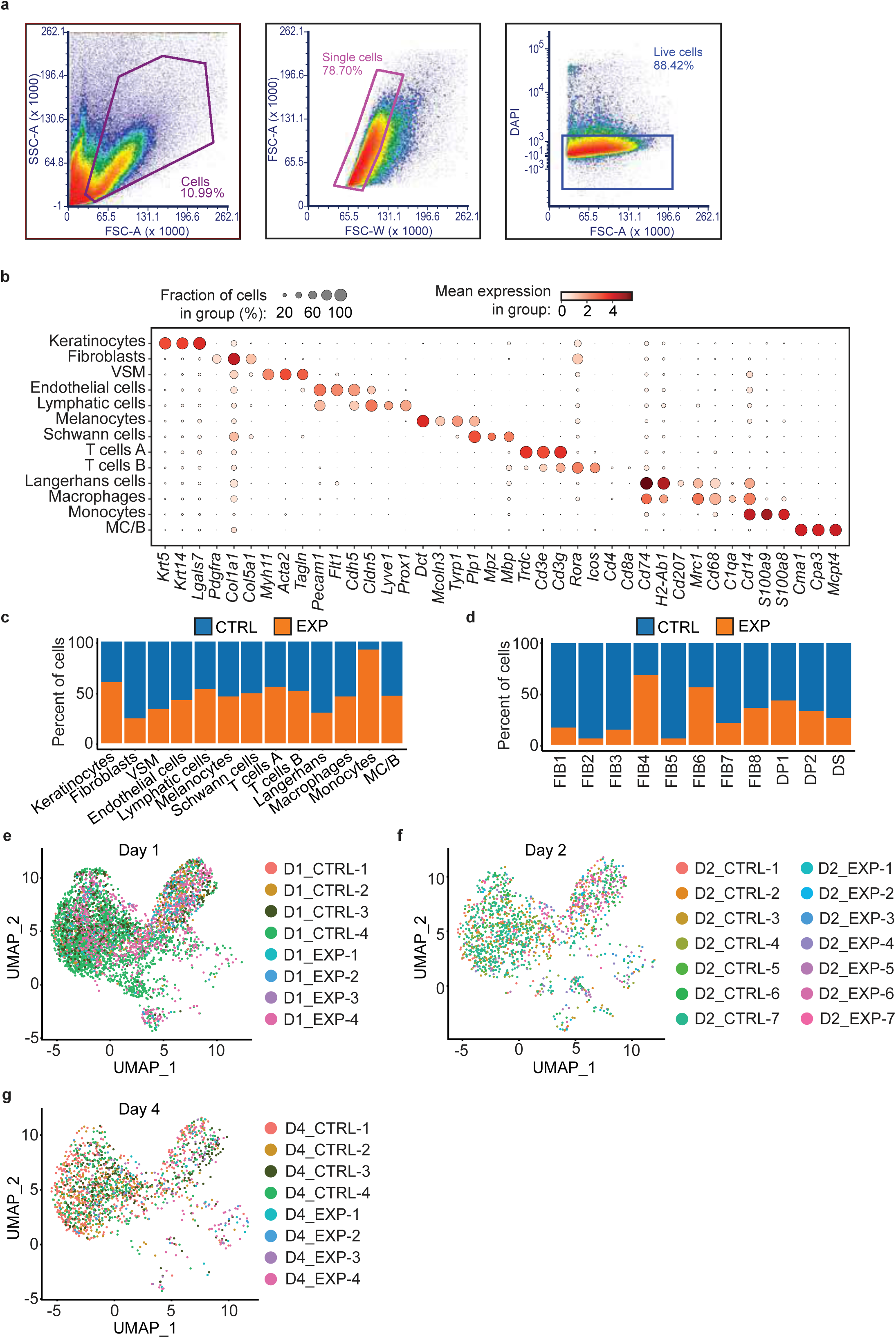
FACS isolation strategy for single-cell RNA sequencing and distribution of biological replicates within the scRNAseq dataset. **a** – FACS isolation strategy of live cells for scRNAseq. **b** – Marker gene expression for each of the clusters shown in Fig. 1b. **c, d** – Proportions of cells from each condition per cluster shown in Fig. 1b, d respectively. **e-g** – Distribution of individual mouse samples within fibroblast subclusters at D1-4, generated through demultiplexing of hashtags.

**Supplementary Figure 2.**
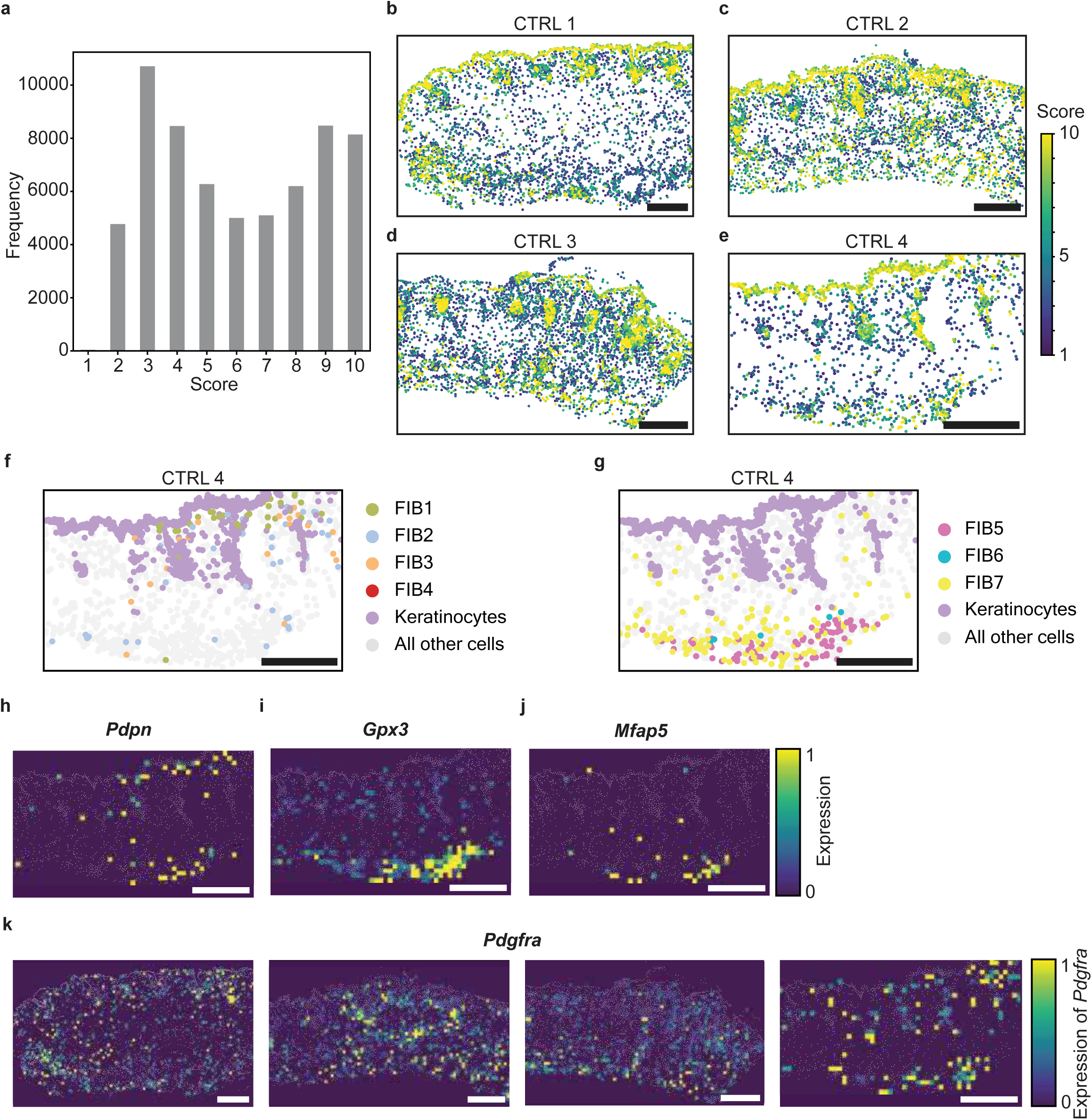
Spatial annotation mapping scores and spatial expression of *Pdgfrα*. **a** – Scoring for annotation mapping of all cell types in the 4 samples. The score corresponds to the number of annotation mapping methods that agree on a cell’s annotation. The frequency (y-axis) shows the number of cells with each score. **b-e** – Spatial plots showing annotation mapping scoring for D2 CTRL samples 1-4 respectively. **f** – Representative image of the spatial distribution of FIB1-4 subpopulations in D2 CTRL sample 4 skin (n=3 sections per mouse). **g** – Representative image of the spatial distribution of FIB5-7 subpopulations in D2 CTRL sample 4 skin (n=3 sections per mouse). **h-j** – Representative images of the spatial density plots showing relative expression of *Pdpn*, *Gpx3* and *Mfap5* transcripts in D2 CTRL sample 4 skin. Scale normalized per gene transcript. Bin size=20 μm. **k** – Representative images of the spatial density plots showing relative expression of *Pdgfrα* transcripts in D2 CTRL 1-4 samples. Scale normalized per gene transcript. Bin size=20 μm. Scale bars=200 μm.

**Supplementary Figure 3.**
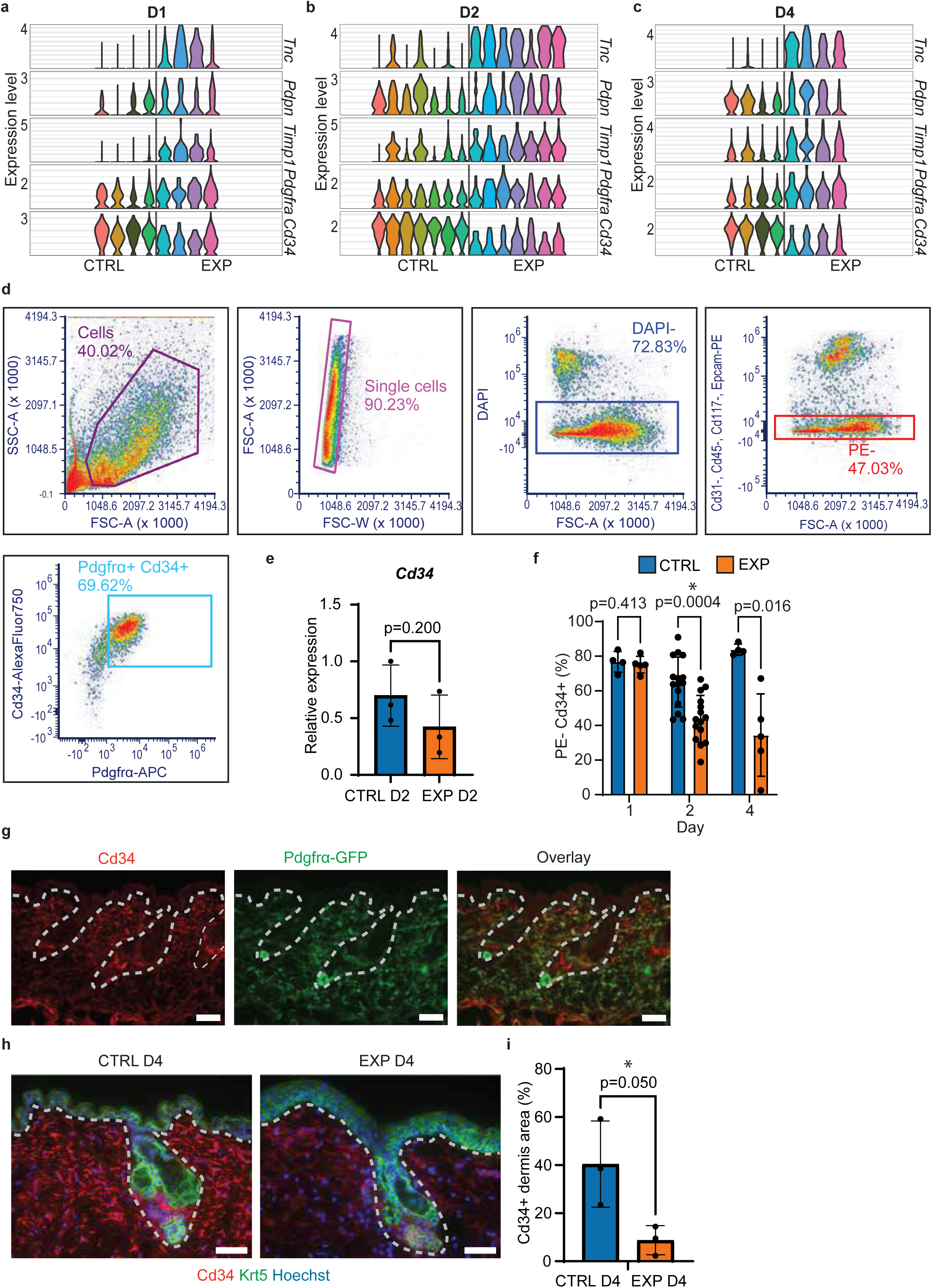
FACS strategy for isolating fibroblasts and expression of *Cd34* in CTRL and EXP skin. **a-c** – Expression of *Tnc, Pdpn, Timp1, Pdgfrα* and *Cd34* in fibroblasts from scRNAseq, per mouse, at D1-D4. **d** – FACS strategy to isolate fibroblasts and fibroblast-like cells from skin cell suspensions. **e** – Relative expression of *Cd34* in sorted fibroblasts by qPCR at D2. Unpaired one-tailed Mann-Whitney U-test, n=3 mice per condition. Data represent mean +/- SD. **f** – Quantification of PE-Cd34+ cells from **d**. Unpaired two-tailed Mann-Whitney U-tests, n=415 mice per condition. Data represent mean +/- SD. **g** – Representative immunofluorescence for Cd34 (red) and Pdgfrα-GFP from *PdgfrαCreERT2;Rosa26^mTmG^* reporter (green). Dotted lines indicate the location of the epidermis and hair follicles. h – Immunofluorescence for Cd34 (red), Krt5 (green) and Hoechst (blue) on D4 CTRL and EXP skin. Dotted lines separate the dermis from the epidermis and hair follicles. **i** – Quantification of the Cd34+ area in the upper dermis, from **h**. Unpaired one-tailed MannWhitney U-test, n=3 mice per condition, means of 3 measurements per animal are plotted as dots. Data represent mean +/- SD. Scale bars=50 μm.

**Supplementary Figure 4.**
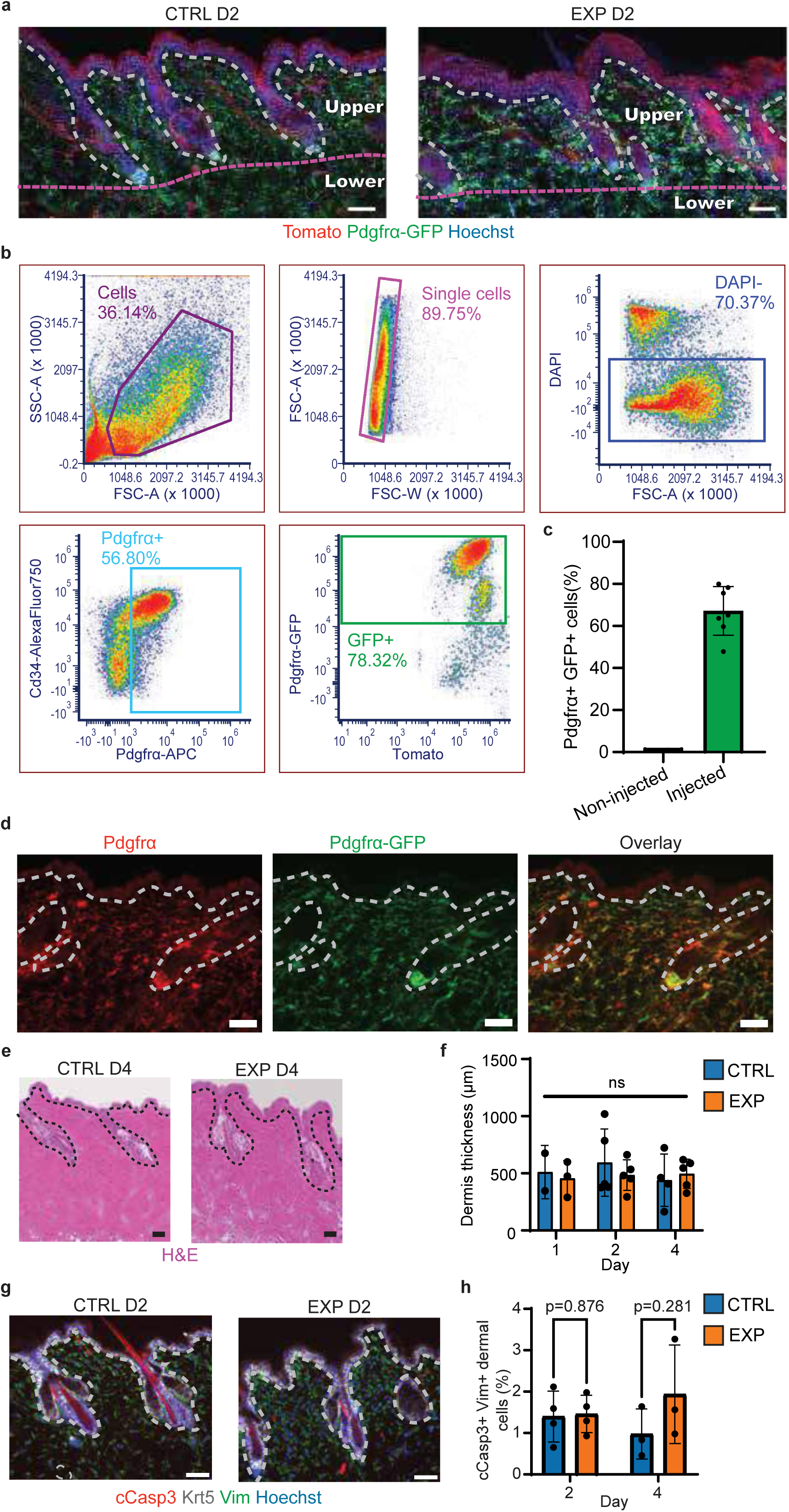
Morphological characterization of expanded skin and *PdgfrαCre-*ERT2;Rosa26^mTmG^ tracing. **a** – Fluorescence for Tomato (red), Pdgfrα-GFP (green) and Hoechst (blue) in D2 CTRL and EXP skin. Upper and lower dermis are delimited by a pink dashed line. The epidermis and hair follicles are delimited by a white dashed line. **b** – Flow cytometry strategy for quantification of *PdgfrαCre-ERT2;Rosa26^mTmG^* recombination. **c** – Quantification of recombination from **b**. n=2-7 mice per condition. Data are presented as mean +/- SD. **d** – Immunofluorescence for Pdgfrα (red) and Pdgfrα-GFP from *PdgfrαCre-ERT2;Rosa26^mTmG^* reporter (green). Epidermis and hair follicle positions are shown by the white line. **e** – Images of Hematoxylin and Eosin (H&E) stained D4 CTRL and EXP skin. Epidermis and hair follicles are delimited by a black dashed line. **f** – Quantification of dermis thickness from **e**. Unpaired two-tailed Mann-Whitney U-tests, n=2-5 mice per condition, means of 3 measurements per animal are plotted as dots. Data represent mean +/- SD. **g** – Immunofluorescence for cleaved Caspase-3 (red), Krt5 (white), Vim (green) and Hoechst (blue) in D2 CTRL and EXP skin. Dotted line delimits the dermis from the epidermis and hair follicles. **h** – Quantification of VIM+ cleaved Caspase-3+ cells in the dermis from **g**. Unpaired two-tailed Mann-Whitney U-test, n=3-4 mice per condition, averages of 3 measurements per animal are plotted. Data represent mean +/- SD. Scale bars=50 μm.

**Supplementary Figure 5.**
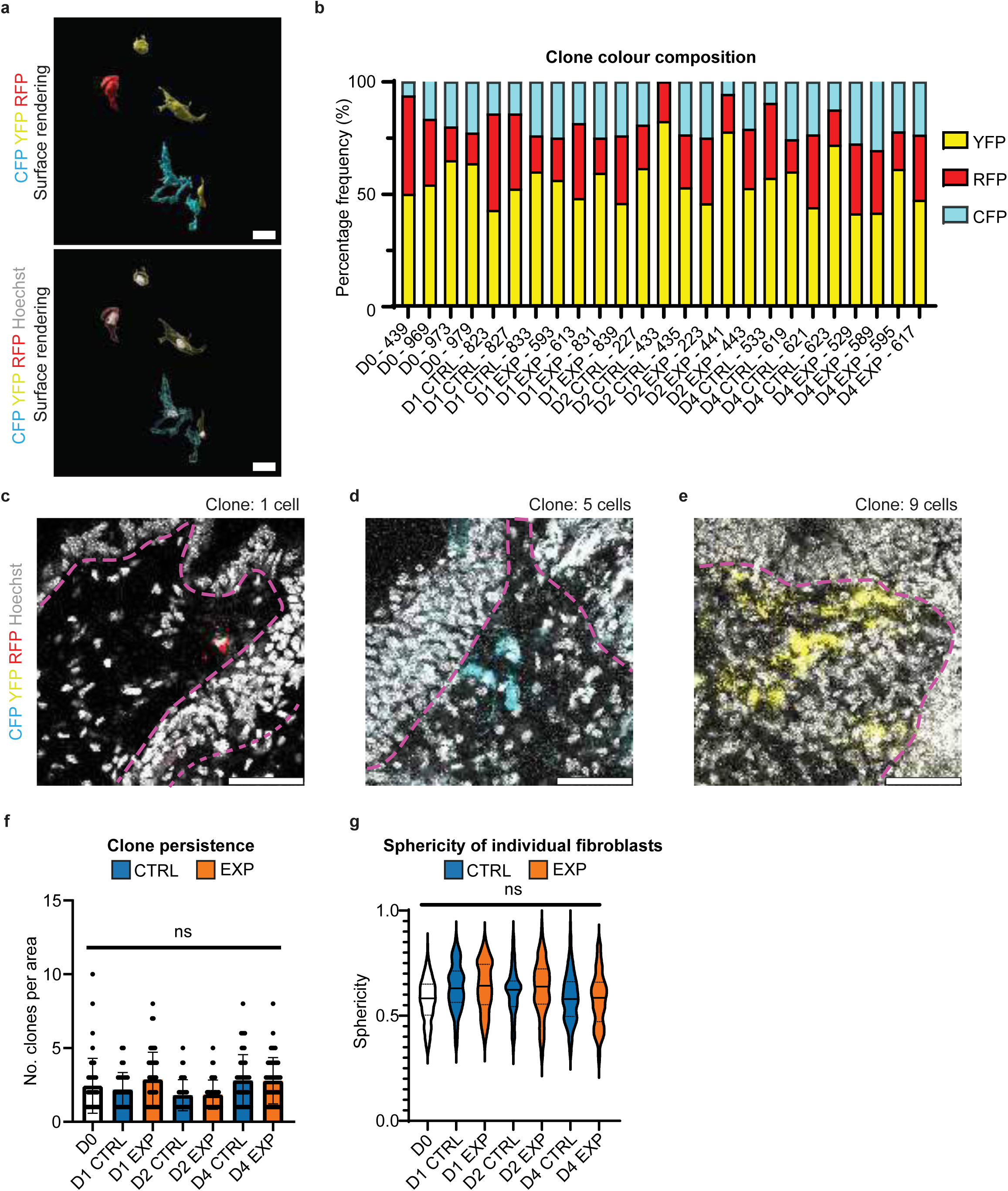
Frequency of recombination of *PdgfrαCre-ERT2; Rosa26^Confetti^* fluorophores and clonal analysis. **a** – Surface rendering of Confetti CFP, YFP and RFP (cyan, yellow and red) and Hoechst (white) fluorescence signal from *PdgfrαCre-ERT2; Rosa26^Confetti^* wholemounts, generated in Imaris. Scale bars=20 μm. **b** – Frequency of YFP, RFP and CFP clone colour per mouse. **c-e** – Representative fluorescence images of labelled cells in skin wholemounts, corresponding to surface rendering images in Fig. 5b**-d**. Pink dotted lines delimit the dermis from the epidermis and hair follicles. Scale bars=50 μm. **f** –Quantification of persistence of clones. Unpaired two-tailed Mann-Whitney U-tests, n=34 mice per condition (number of measurements for D0=47, D1 CTRL=33, D1 EXP=45, D2 CTRL=37, D2 EXP=44, D4 CTRL=47, D4 EXP=47). Data are presented as mean +/- SD. **g** – Quantification of sphericity of individual fibroblasts. Unpaired two-tailed Mann-Whitney U-tests, n=3-4 mice per condition (number of measurements for D0=166, D1 CTRL=90, D1 EXP=198, D2 CTRL=95, D2 EXP=107, D4 CTRL=175, D4 EXP=180). Data are presented as violin plots showing the median and upper and lower quartiles.

**Supplementary Figure 6.**
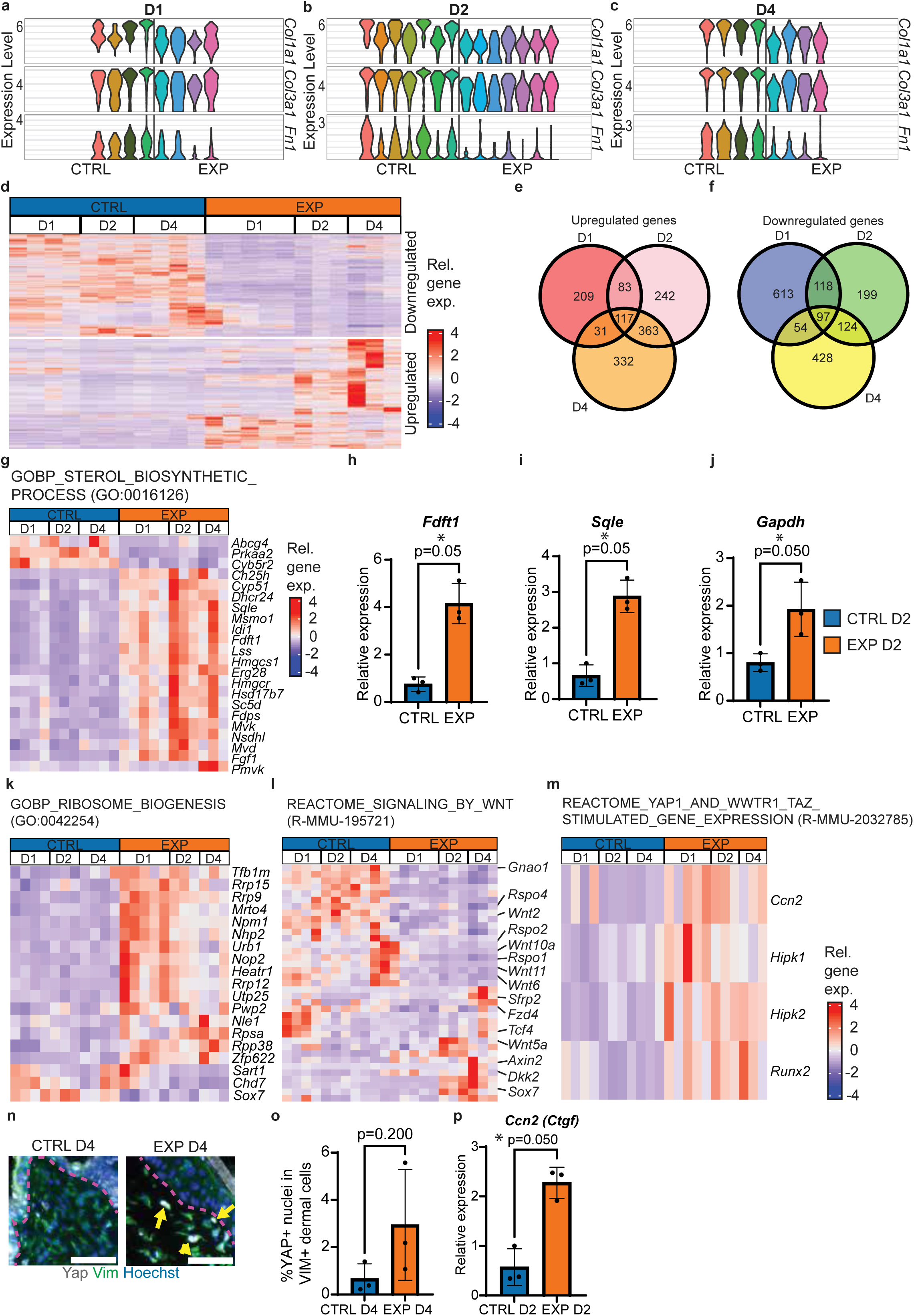
Transcriptional heterogeneity is reproducible across mice, upregulated and downregulated genes from bulk RNAseq and upregulation of YAP target genes. **a-c** – Expression of *Col1a1, Col3a1* and *Fn1* in fibroblasts from scRNAseq, per mouse, at D1D4. **d** – Heatmap showing all differentially expressed genes (DEG) from bulk RNAseq analysis of sorted fibroblasts. Relative gene expression (Rel. gene exp.) is shown. **e,f** – Venn diagrams showing numbers of upregulated and downregulated genes respectively from bulk RNAseq of sorted Pdgfrα+ and Cd34+ fibroblasts in EXP compared to CTRL at D1-4. **g** – Heatmaps of DEG from bulk RNAseq analysis of sorted Pdgfrα+ and Cd34+ fibroblasts. Relative gene expression (Rel. gene exp.) is shown for the indicated gene ontology (GO) term, with red indicating high expression, and blue indicating low expression. **h-j** – Relative expression of the indicated genes in sorted fibroblasts by qPCR at D2. Unpaired one-tailed Mann-Whitney U-tests, n=3 mice per condition. Data are presented as mean +/- SD. **k** – Heatmap of DEG within the gene ontology (GO) term for Ribosome biogenesis. Relative gene expression is shown, with red indicating high expression, and blue indicating low expression. **l** – Heatmap of DEG within the Reactome term for signalling by Wnt. Relative gene expression is shown, with red indicating high expression, and blue indicating low expression. **m** – Heatmap of DEG within the Reactome term for YAP1/TAZ-stimulated gene expression. Relative gene expression is shown, with red indicating high expression, and blue indicating low expression. **n** – Immunofluorescence for Yap (White), Vim (green) and Hoechst (blue) in D4 CTRL and EXP skin. Pink dotted lines separate the dermis from the epidermis and hair follicles. Yellow arrows indicate nuclear Yap nuclear localization. Scale bars=50 μm. **o** – Quantification of nuclear Yap in Vim+ dermal cells from **m**. Unpaired two-tailed MannWhitney U-test, n=3 mice per condition, means of 3 measurements per mouse are plotted as dots. Data are presented as mean +/- SD. **p** – Relative expression of *Ccn2* (*Ctgf*) in sorted fibroblasts by qPCR at D2. Unpaired one-tailed Mann-Whitney U-tests, n=3 mice per condition. Data represent mean +/- SD.

**Supplementary Figure 7.**
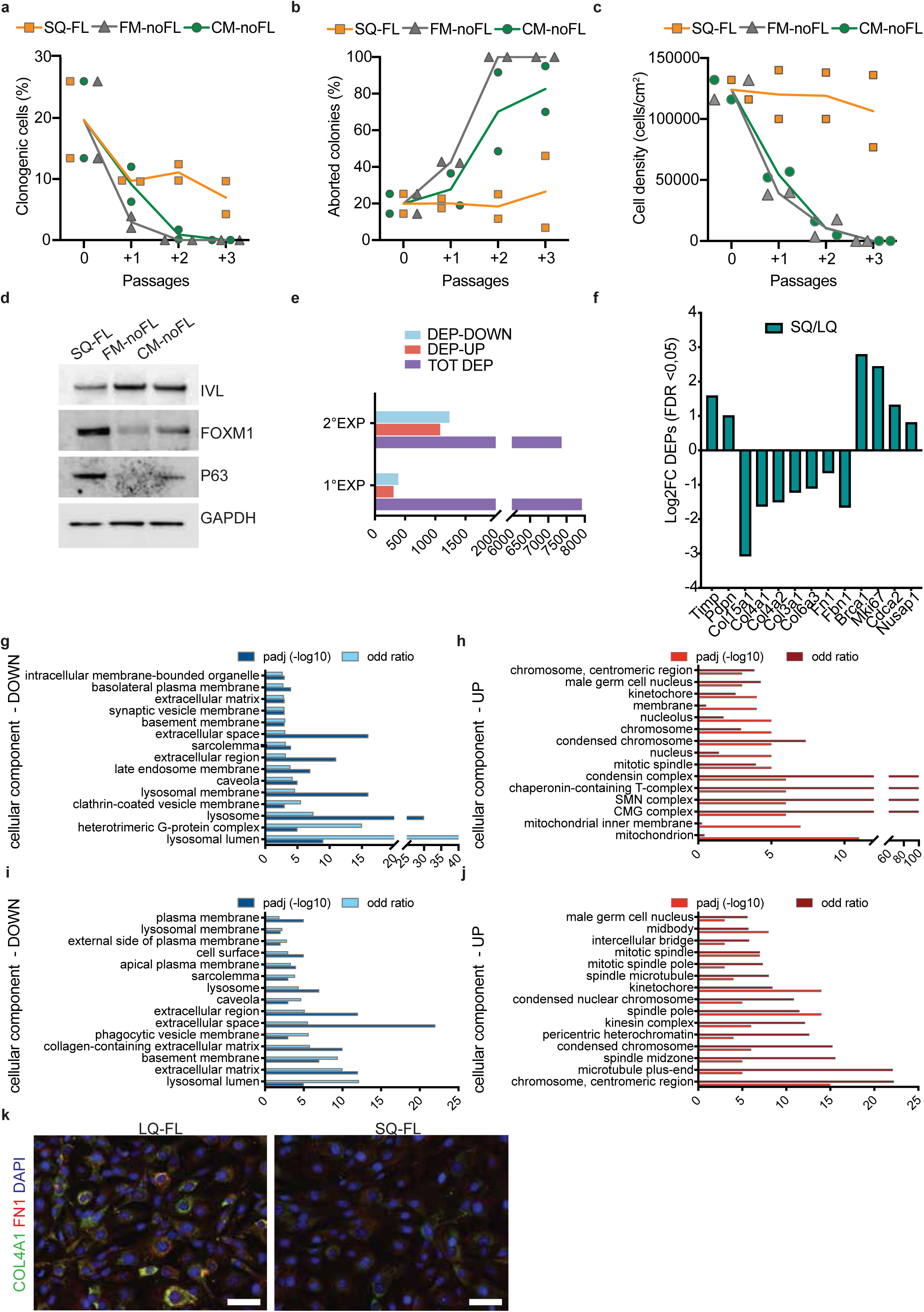
Effect of different keratinocyte-fibroblast co-culture conditions, quality control of mass spectrometry experiments and cellular component term analyses. **a,b** – Serial cultivation of normal human keratinocytes on SQ-FL (orange line), as compared to no feeder layer conditions, with fresh KGM medium (FM-noFL, grey line) or fibroblast conditioned medium (CM-noFL, green line). Percentage of clonogenic cells was calculated as the ratio between grown colonies and plated cells (**a**). Percentage of aborted colonies was calculated as the ratio between the colonies scored as aborted and the number of clonogenic cells (**b**). n=2 independent biological replicates. Data are presented as mean +/− SEM. **c** – Total amount of cells obtained at subconfluence during serial cultivation of normal human keratinocytes (NHK) on SQ-FL (orange line) as compared to FM-noFL (grey line) or CM-noFL, (blue line). n=2 independent biological replicates. Data are presented as mean +/− SEM. **d** – Western analysis of total cell extracts from cultures generated by NHKs on SQ-FL, FM-noFL or CM-noFL. One of two representative experiments is shown. **e** – Numbers of differentially expressed proteins (DEP) from mass spectrometry analysis of two independent experiments (1°EXP and 2°EXP). Upregulated (DEP-UP), downregulated (DEP-DOWN) and total (TOT DEP) protein numbers are plotted. **f** – Bar showing log2 Fold change of some of the downregulated or upregulated proteins of interest. **g**,**h** – Graphs showing downregulated (**g**) and upregulated (**h**) cellular components in mass spectrometry analysis of SQ as compared to LQ feeder layers (2°EXP). p values are calculated with one-sided Fisher’s Exact test and corrected for multiple tests with the Benjamini– Hochberg method. Data presented as –log10 of corrected p value (dark blue and dark red bars respectively) and corresponding fold change (light blue and light red bars respectively). **i**,**j** – Graphs showing downregulated (**i**) and upregulated (**j**) cellular components in mass spectrometry analysis of SQ as compared to LQ feeder layers (1°EXP). p values are calculated with one-sided Fisher’s Exact test and corrected for multiple tests with the Benjamini– Hochberg method. Data presented as –log10 of corrected p value (dark blue and dark red bars respectively) and corresponding fold change (light blue and light red bars respectively). **k** – Representative immunofluorescence of the two type of feeder layer stained for COL4A1 (green) and FN1 (red) and nuceli (blue). Scale bar = 50 μm

## Bibliography

1 Nguyen, T. M. & Aragona, M. Regulation of tissue architecture and stem cell dynamics to sustain homeostasis and repair in the skin epidermis. Semin Cell Dev Biol 130, 79–89 (2022). 10.1016/j.semcdb.2021.09.008

2 Iskratsch, T., Wolfenson, H. & Sheetz, M. P. Appreciating force and shape-the rise of mechanotransduction in cell biology. Nat Rev Mol Cell Biol 15, 825–833 (2014). 10.1038/nrm3903

3 Mammoto, T., Mammoto, A. & Ingber, D. E. Mechanobiology and developmental control. Annu Rev Cell Dev Biol 29, 27–61 (2013). 10.1146/annurev-cellbio-101512-122340

4 LeGoff, L. & Lecuit, T. Mechanical Forces and Growth in Animal Tissues. Cold Spring Harb Perspect Biol 8, a019232 (2015). 10.1101/cshperspect.a019232

5 Dupont, S. & Wickstrom, S. A. Mechanical regulation of chromatin and transcription. Nat Rev Genet 23, 624–643 (2022). 10.1038/s41576-022-00493-6

6 Guillot, C. & Lecuit, T. Mechanics of epithelial tissue homeostasis and morphogenesis. Science 340, 1185–1189 (2013). 10.1126/science.1235249

7 Vining, K. H. & Mooney, D. J. Mechanical forces direct stem cell behaviour in development and regeneration. Nat Rev Mol Cell Biol 18, 728–742 (2017). 10.1038/nrm.2017.108

8 Kirby, T. J. & Lammerding, J. Emerging views of the nucleus as a cellular mechanosensor. Nat Cell Biol 20, 373–381 (2018). 10.1038/s41556-018-0038-y

9 DuFort, C. C., Paszek, M. J. & Weaver, V. M. Balancing forces: architectural control of mechanotransduction. Nat Rev Mol Cell Biol 12, 308–319 (2011). 10.1038/nrm3112

10 Zollner, A. M., Holland, M. A., Honda, K. S., Gosain, A. K. & Kuhl, E. Growth on demand: reviewing the mechanobiology of stretched skin. J Mech Behav Biomed Mater 28, 495–509 (2013). 10.1016/j.jmbbm.2013.03.018

11 Tzolova, N. & Hadjiiski, O. Tissue expansion used as a method of reconstructive surgery in childhood. Ann Burns Fire Disasters 21, 23–30 (2008).

12 Cho, M. J. et al. The current use of tissue expanders in breast reconstruction: device design, features, and technical considerations. Expert Rev Med Devices 21, 27–35 (2024). 10.1080/17434440.2023.2288911

13 Biggs, L. C., Kim, C. S., Miroshnikova, Y. A. & Wickstrom, S. A. Mechanical Forces in the Skin: Roles in Tissue Architecture, Stability, and Function. J Invest Dermatol 140, 284–290 (2020). 10.1016/j.jid.2019.06.137

14 Villeneuve, C. et al. Mechanical forces across compartments coordinate cell shape and fate transitions to generate tissue architecture. Nat Cell Biol 26, 207–218 (2024). 10.1038/s41556-023-01332-4

15 Aragona, M. et al. Mechanisms of stretch-mediated skin expansion at single-cell resolution. Nature 584, 268–273 (2020). 10.1038/s41586-020-2555-7

16 Chu, S. Y. et al. Mechanical stretch induces hair regeneration through the alternative activation of macrophages. Nat Commun 10, 1524 (2019). 10.1038/s41467-019-09402-8

17 Xue, Y. et al. Mechanical tension mobilizes Lgr6(+) epidermal stem cells to drive skin growth. Sci Adv 8, eabl8698 (2022). 10.1126/sciadv.abl8698

18 Ding, J. et al. Macrophages are necessary for skin regeneration during tissue expansion. J Transl Med 17, 36 (2019). 10.1186/s12967-019-1780-z

19 Plikus, M. V. et al. Fibroblasts: Origins, definitions, and functions in health and disease. Cell 184, 3852–3872 (2021). 10.1016/j.cell.2021.06.024

20 Tabula Muris, C., et al. Single-cell transcriptomics of 20 mouse organs creates a Tabula Muris. Nature 562, 367–372 (2018). 10.1038/s41586-018-0590-4

21 Joost, S. et al. The Molecular Anatomy of Mouse Skin during Hair Growth and Rest. Cell Stem Cell 26, 441–457 e447 (2020). 10.1016/j.stem.2020.01.012

22 Liu, Y. et al. Single-Cell Profiling Reveals Divergent, Globally Patterned Immune Responses in Murine Skin Inflammation. iScience 23, 101582 (2020). 10.1016/j.isci.2020.101582

23 Ganier, C., Rognoni, E., Goss, G., Lynch, M. & Watt, F. M. Fibroblast Heterogeneity in Healthy and Wounded Skin. Cold Spring Harb Perspect Biol 14 (2022). 10.1101/cshperspect.a041238

24 Driskell, R. R. & Watt, F. M. Understanding fibroblast heterogeneity in the skin. Trends Cell Biol 25, 92–99 (2015). 10.1016/j.tcb.2014.10.001

25 Phan, Q. M. et al. Lef1 expression in fibroblasts maintains developmental potential in adult skin to regenerate wounds. Elife 9 (2020). 10.7554/eLife.60066

26 Sola, P. et al. Targeting lymphoid-derived IL-17 signaling to delay skin aging. Nat Aging 3, 688–704 (2023). 10.1038/s43587-023-00431-z

27 Salzer, M. C. et al. Identity Noise and Adipogenic Traits Characterize Dermal Fibroblast Aging. Cell 175, 1575–1590 e1522 (2018). 10.1016/j.cell.2018.10.012

28 Phan, Q. M., Sinha, S., Biernaskie, J. & Driskell, R. R. Single-cell transcriptomic analysis of small and large wounds reveals the distinct spatial organization of regenerative fibroblasts. Exp Dermatol 30, 92–101 (2021). 10.1111/exd.14244

29 Sennett, R. et al. An Integrated Transcriptome Atlas of Embryonic Hair Follicle Progenitors, Their Niche, and the Developing Skin. Dev Cell 34, 577–591 (2015). 10.1016/j.devcel.2015.06.023

30 Collins, C. A., Kretzschmar, K. & Watt, F. M. Reprogramming adult dermis to a neonatal state through epidermal activation of beta-catenin. Development 138, 5189–5199 (2011). 10.1242/dev.064592

31 Mittal, R. et al. Intricate Functions of Matrix Metalloproteinases in Physiological and Pathological Conditions. J Cell Physiol 231, 2599–2621 (2016). 10.1002/jcp.25430

32 de Castro Bras, L. E. & Frangogiannis, N. G. Extracellular matrix-derived peptides in tissue remodeling and fibrosis. Matrix Biol 91-92, 176–187 (2020). 10.1016/j.matbio.2020.04.006

33 Midwood, K. S., Chiquet, M., Tucker, R. P. & Orend, G. Tenascin-C at a glance. J Cell Sci 129, 4321–4327 (2016). 10.1242/jcs.190546

34 Asai, J. The Role of Podoplanin in Skin Diseases. Int J Mol Sci 23 (2022). 10.3390/ijms23031310

35 Correa-Gallegos, D. et al. CD201(+) fascia progenitors choreograph injury repair. Nature 623, 792–802 (2023). 10.1038/s41586-023-06725-x

36 Jiang, D. et al. Injury triggers fascia fibroblast collective cell migration to drive scar formation through N-cadherin. Nat Commun 11, 5653 (2020). 10.1038/s41467-020-19425-1

37 Jiang, D. et al. Two succeeding fibroblastic lineages drive dermal development and the transition from regeneration to scarring. Nat Cell Biol 20, 422–431 (2018). 10.1038/s41556-018-0073-8

38 Marsh, E., Gonzalez, D. G., Lathrop, E. A., Boucher, J. & Greco, V. Positional Stability and Membrane Occupancy Define Skin Fibroblast Homeostasis In Vivo. Cell 175, 1620–1633 e1613 (2018). 10.1016/j.cell.2018.10.013

39 Rognoni, E. et al. Fibroblast state switching orchestrates dermal maturation and wound healing. Mol Syst Biol 14, e8174 (2018). 10.15252/msb.20178174

40 Panciera, T., Azzolin, L., Cordenonsi, M. & Piccolo, S. Mechanobiology of YAP and TAZ in physiology and disease. Nat Rev Mol Cell Biol 18, 758–770 (2017). 10.1038/nrm.2017.87

41 Lichtenberger, B. M., Mastrogiannaki, M. & Watt, F. M. Epidermal beta-catenin activation remodels the dermis via paracrine signalling to distinct fibroblast lineages. Nat Commun 7, 10537 (2016). 10.1038/ncomms10537

42 Driskell, R. R. et al. Distinct fibroblast lineages determine dermal architecture in skin development and repair. Nature 504, 277–281 (2013). 10.1038/nature12783

43 Sorrell, J. M., Baber, M. A. & Caplan, A. I. Site-matched papillary and reticular human dermal fibroblasts differ in their release of specific growth factors/cytokines and in their interaction with keratinocytes. J Cell Physiol 200, 134–145 (2004). 10.1002/jcp.10474

44 Wang, Z., Wang, Y., Farhangfar, F., Zimmer, M. & Zhang, Y. Enhanced keratinocyte proliferation and migration in co-culture with fibroblasts. PLoS One 7, e40951 (2012). 10.1371/journal.pone.0040951

45 Rheinwald, J. G. & Green, H. Serial cultivation of strains of human epidermal keratinocytes: the formation of keratinizing colonies from single cells. Cell 6, 331–343 (1975). 10.1016/s0092-8674(75)80001-8

46 Gallico, G. G., 3rd, O’Connor, N. E., Compton, C. C., Kehinde, O. & Green, H. Permanent coverage of large burn wounds with autologous cultured human epithelium. N Engl J Med 311, 448–451 (1984). 10.1056/NEJM198408163110706

47 Rama, P. et al. Limbal stem-cell therapy and long-term corneal regeneration. N Engl J Med 363, 147–155 (2010). 10.1056/NEJMoa0905955

48 Hirsch, T. et al. Regeneration of the entire human epidermis using transgenic stem cells. Nature 551, 327–332 (2017). 10.1038/nature24487

49 De Rosa, L. et al. Hologene 5: A Phase II/III Clinical Trial of Combined Cell and Gene Therapy of Junctional Epidermolysis Bullosa. Front Genet 12, 705019 (2021). 10.3389/fgene.2021.705019

50 Enzo, E. et al. Single-keratinocyte transcriptomic analyses identify different clonal types and proliferative potential mediated by FOXM1 in human epidermal stem cells. Nat Commun 12, 2505 (2021). 10.1038/s41467-021-22779-9

51 Thompson, S. M., Phan, Q. M., Winuthayanon, S., Driskell, I. M. & Driskell, R. R. Parallel Single-Cell Multiomics Analysis of Neonatal Skin Reveals the Transitional Fibroblast States that Restrict Differentiation into Distinct Fates. J Invest Dermatol 142, 1812–1823 e1813 (2022). 10.1016/j.jid.2021.11.032

52 Jiang, D. & Rinkevich, Y. Distinct fibroblasts in scars and regeneration. Curr Opin Genet Dev 70, 7–14 (2021). 10.1016/j.gde.2021.04.005

53 Phan, Q. M. et al. Lineage commitment of dermal fibroblast progenitors is controlled by Kdm6b-mediated chromatin demethylation. EMBO J 42, e113880 (2023). 10.15252/embj.2023113880

54 Korosec, A. et al. Lineage Identity and Location within the Dermis Determine the Function of Papillary and Reticular Fibroblasts in Human Skin. J Invest Dermatol 139, 342–351 (2019). 10.1016/j.jid.2018.07.033

55 Ascension, A. M., Fuertes-Alvarez, S., Ibanez-Sole, O., Izeta, A. & Arauzo-Bravo, M. J. Human Dermal Fibroblast Subpopulations Are Conserved across Single-Cell RNA Sequencing Studies. J Invest Dermatol 141, 1735–1744 e1735 (2021). 10.1016/j.jid.2020.11.028

56 Jacob, T. et al. Molecular and spatial landmarks of early mouse skin development. Dev Cell 58, 2140–2162 e2145 (2023). 10.1016/j.devcel.2023.07.015

57 Philippeos, C. et al. Spatial and Single-Cell Transcriptional Profiling Identifies Functionally Distinct Human Dermal Fibroblast Subpopulations. J Invest Dermatol 138, 811–825 (2018). 10.1016/j.jid.2018.01.016

58 Guerrero-Juarez, C. F. et al. Single-cell analysis reveals fibroblast heterogeneity and myeloid-derived adipocyte progenitors in murine skin wounds. Nat Commun 10, 650 (2019). 10.1038/s41467-018-08247-x

59 Mascharak, S. et al. Preventing Engrailed-1 activation in fibroblasts yields wound regeneration without scarring. Science 372 (2021). 10.1126/science.aba2374

60 Rinkevich, Y. et al. Skin fibrosis. Identification and isolation of a dermal lineage with intrinsic fibrogenic potential. Science 348, aaa2151 (2015). 10.1126/science.aaa2151

61 Forsthuber, A. et al. Cancer-associated fibroblast subtypes modulate the tumor-immune microenvironment and are associated with skin cancer malignancy. Nat Commun 15, 9678 (2024). 10.1038/s41467-024-53908-9

62 Rognoni, E. et al. Role of distinct fibroblast lineages and immune cells in dermal repair following UV radiation-induced tissue damage. Elife 10 (2021). 10.7554/eLife.71052

63 Thrane, K. et al. Single-Cell and Spatial Transcriptomic Analysis of Human Skin Delineates Intercellular Communication and Pathogenic Cells. J Invest Dermatol 143, 2177–2192 e2113 (2023). 10.1016/j.jid.2023.02.040

64 Griffin, M. F. et al. Piezo inhibition prevents and rescues scarring by targeting the adipocyte to fibroblast transition. bioRxiv (2023). 10.1101/2023.04.03.535302

65 Andersen, M. S. et al. Tracing the cellular dynamics of sebaceous gland development in normal and perturbed states. Nat Cell Biol 21, 924–932 (2019). 10.1038/s41556-019-0362-x

66 Bansaccal, N. et al. The extracellular matrix dictates regional competence for tumour initiation. Nature 623, 828–835 (2023). 10.1038/s41586-023-06740-y

67 Tanimura, S. et al. Hair follicle stem cells provide a functional niche for melanocyte stem cells. Cell Stem Cell 8, 177–187 (2011). 10.1016/j.stem.2010.11.029

68 Greco, V. et al. A two-step mechanism for stem cell activation during hair regeneration. Cell Stem Cell 4, 155–169 (2009). 10.1016/j.stem.2008.12.009

69 Rezza, A. et al. Signaling Networks among Stem Cell Precursors, Transit-Amplifying Progenitors, and their Niche in Developing Hair Follicles. Cell Rep 14, 3001–3018 (2016). 10.1016/j.celrep.2016.02.078

70 Shook, B. A. et al. Myofibroblast proliferation and heterogeneity are supported by macrophages during skin repair. Science 362 (2018). 10.1126/science.aar2971

71 Cai, X. et al. Tenascin C(+) papillary fibroblasts facilitate neuro-immune interaction in a mouse model of psoriasis. Nat Commun 14, 2004 (2023). 10.1038/s41467-023-37798-x

72 Kyprianou, C. et al. Basement membrane remodelling regulates mouse embryogenesis. Nature 582, 253–258 (2020). 10.1038/s41586-020-2264-2

73 Trappmann, B. et al. Extracellular-matrix tethering regulates stem-cell fate. Nat Mater 11, 642–649 (2012). 10.1038/nmat3339

74 Zijl, S. et al. Micro-scaled topographies direct differentiation of human epidermal stem cells. Acta Biomater 84, 133–145 (2019). 10.1016/j.actbio.2018.12.003

75 Roig-Rosello, E. et al. Dermal stiffness governs the topography of the epidermis and the underlying basement membrane in young and old human skin. Aging Cell 23, e14096 (2024). 10.1111/acel.14096

76 Chung, M. I., Bujnis, M., Barkauskas, C. E., Kobayashi, Y. & Hogan, B. L. M. Niche-mediated BMP/SMAD signaling regulates lung alveolar stem cell proliferation and differentiation. Development 145 (2018). 10.1242/dev.163014

77 Snippert, H. J. et al. Intestinal crypt homeostasis results from neutral competition between symmetrically dividing Lgr5 stem cells. Cell 143, 134–144 (2010). 10.1016/j.cell.2010.09.016

78 Muzumdar, M. D., Tasic, B., Miyamichi, K., Li, L. & Luo, L. A global double-fluorescent Cre reporter mouse. Genesis 45, 593–605 (2007). 10.1002/dvg.20335

79 Maciag, G. et al. JAK/STAT signaling promotes the emergence of unique cell states in ulcerative colitis. Stem Cell Reports 19, 1172–1188 (2024). 10.1016/j.stemcr.2024.06.006

80 Stoeckius, M. et al. Cell Hashing with barcoded antibodies enables multiplexing and doublet detection for single cell genomics. Genome Biol 19, 224 (2018). 10.1186/s13059-018-1603-1

81 Wolf, F. A., Angerer, P. & Theis, F. J. SCANPY: large-scale single-cell gene expression data analysis. Genome Biol 19, 15 (2018). 10.1186/s13059-017-1382-0

82 Hao, Y. et al. Integrated analysis of multimodal single-cell data. Cell 184, 3573–3587 e3529 (2021). 10.1016/j.cell.2021.04.048

83 Germain, P. L., Lun, A., Garcia Meixide, C., Macnair, W. & Robinson, M. D. Doublet identification in single-cell sequencing data using scDblFinder. F1000Res 10, 979 (2021). 10.12688/f1000research.73600.2

84 Korsunsky, I. et al. Fast, sensitive and accurate integration of single-cell data with Harmony. Nat Methods 16, 1289–1296 (2019). 10.1038/s41592-019-0619-0

85 Ergen, C. et al. Consensus prediction of cell type labels in single-cell data with popV. Nat Genet 56, 2731–2738 (2024). 10.1038/s41588-024-01993-3

86 Gayoso, A. et al. A Python library for probabilistic analysis of single-cell omics data. Nat Biotechnol 40, 163–166 (2022). 10.1038/s41587-021-01206-w

87 Hie, B., Bryson, B. & Berger, B. Efficient integration of heterogeneous single-cell transcriptomes using Scanorama. Nat Biotechnol 37, 685–691 (2019). 10.1038/s41587-019-0113-3

88 Biancalani, T. et al. Deep learning and alignment of spatially resolved single-cell transcriptomes with Tangram. Nat Methods 18, 1352–1362 (2021). 10.1038/s41592-021-01264-7

89 Rahimi, A., Vale-Silva, L. A., Falth Savitski, M., Tanevski, J. & Saez-Rodriguez, J. DOT: a flexible multi-objective optimization framework for transferring features across single-cell and spatial omics. Nat Commun 15, 4994 (2024). 10.1038/s41467-024-48868-z

90 Virtanen, P. et al. SciPy 1.0: fundamental algorithms for scientific computing in Python. Nat Methods 17, 261–272 (2020). 10.1038/s41592-019-0686-2

91 Liao, Y., Smyth, G. K. & Shi, W. The R package Rsubread is easier, faster, cheaper and better for alignment and quantification of RNA sequencing reads. Nucleic Acids Res 47, e47 (2019). 10.1093/nar/gkz114

92 Robinson, M. D., McCarthy, D. J. & Smyth, G. K. edgeR: a Bioconductor package for differential expression analysis of digital gene expression data. Bioinformatics 26, 139–140 (2010). 10.1093/bioinformatics/btp616

93 Love, M. I., Huber, W. & Anders, S. Moderated estimation of fold change and dispersion for RNA-seq data with DESeq2. Genome Biol 15, 550 (2014). 10.1186/s13059-014-0550-8

94 Subramanian, A. et al. Gene set enrichment analysis: a knowledge-based approach for interpreting genome-wide expression profiles. Proc Natl Acad Sci U S A 102, 15545–15550 (2005). 10.1073/pnas.0506580102

95 Castanza, A. S. et al. Extending support for mouse data in the Molecular Signatures Database (MSigDB). Nat Methods 20, 1619–1620 (2023). 10.1038/s41592-023-02014-7

96 Gu, Z., Eils, R. & Schlesner, M. Complex heatmaps reveal patterns and correlations in multidimensional genomic data. Bioinformatics 32, 2847–2849 (2016). 10.1093/bioinformatics/btw313

97 Livak, K. J. & Schmittgen, T. D. Analysis of relative gene expression data using real-time quantitative PCR and the 2(-Delta Delta C(T)) Method. Methods 25, 402–408 (2001). 10.1006/meth.2001.1262

98 Bankhead, P. et al. QuPath: Open source software for digital pathology image analysis. Sci Rep 7, 16878 (2017). 10.1038/s41598-017-17204-5

99 Santos, A. et al. A knowledge graph to interpret clinical proteomics data. Nat Biotechnol 40, 692–702 (2022). 10.1038/s41587-021-01145-6

100 Lazar, C., Gatto, L., Ferro, M., Bruley, C. & Burger, T. Accounting for the Multiple Natures of Missing Values in Label-Free Quantitative Proteomics Data Sets to Compare Imputation Strategies. J Proteome Res 15, 1116–1125 (2016). 10.1021/acs.jproteome.5b00981

