## Supplementary Tables for "Defining the role of fibroblasts in skin expansion"

**Supplementary Table 1 related to Fig. 1b scRNAseq marker genes for all cell types**

| Keratinocytes | Fibroblasts | Vascular smc | Endothelial c | Lymphatic ce | Melanocytes | Schwann cell | T cells A | T cells B | Langerhans c | Macrophages | Monocytes | Mast cells/Basophils |
|---|---|---|---|---|---|---|---|---|---|---|---|---|
| *Lgals7* | *Dcn* | *Igfbp7* | *Pecam1* | *Cldn5* | *Dct* | *Plp1* | *Trdc* | *Srgn* | *Cd74* | *Fcer1g* | *Srgn* | *Serpinb1a* |
| *Fxyd3* | *Bgn* | *Sparcl1* | *Flt1* | *Ifitm3* | *Cd63* | *Cd9* | *Cd3e* | *Ets1* | *H2-Aa* | *Ctss* | *Plek* | *Cma1* |
| *Perp* | *Col6a1* | *Tpm1* | *Cdh5* | *Ecscr* | *Tyrp1* | *Timp3* | *Rgs1* | *Ptprc* | *H2-Ab1* | *Tyrobp* | *Tyrobp* | *Cpa3* |
| *Sfn* | *Col1a2* | *Cald1* | *Ly6c1* | *Egfl7* | *Pax3* | *Cryab* | *Cd3g* | *Cd52* | *Ctss* | *Plek* | *Cxcl2* | *Mcpt4* |
| *Krt5* | *Ctsk* | *Gm13889* | *Ifitm3* | *Prox1* | *Col12a1* | *Qk* | *Rgs2* | *Il2rb* | *Tmsb4x* | *Laptm5* | *Fcer1g* | *Hdc* |
| *Dsp* | *Col5a2* | *Mylk* | *Tm4sf1* | *Timp3* | *Syngr1* | *Gatm* | *Cd7* | *Laptm5* | *H2-DMa* | *Lyz2* | *Cd14* | *Cyp11a1* |
| *Apoe* | *Serpinh1* | *Acta2* | *Plvap* | *Cd9* | *Plp1* | *Cd59a* | *Ctla2a* | *Rora* | *Tyrobp* | *Ftl1* | *Lcp1* | *Srgn* |
| *S100a14* | *Serping1* | *Des* | *Sparcl1* | *Fxyd6* | *Enpp2* | *Pmp22* | *Trat1* | *Rgs1* | *H2-DMb1* | *Slfn2* | *Il1b* | *Ccl7* |
| *Krt14* | *Col1a1* | *Notch3* | *Igfbp7* | *Pecam1* | *Ednrb* | *Kcna1* | *Nkg7* | *Icos* | *Laptm5* | *Atp2b1* | *Lilrb4a* | *Tpsb2* |
| *Dsc3* | *Twist1* | *Mprip* | *Esam* | *Mmrn1* | *Idh2* | *Gpm6b* | *Ptprc* | *D16Ertd472e* | *Srgn* | *Alox5ap* | *Hdc* | *Plek* |
| *Trim29* | *Fstl1* | *Myh11* | *Adgrf5* | *Ramp2* | *Mcoln3* | *Scd2* | *Fermt2* | *Samsn1* | *Ifi30* | *Cxcl2* | *Clec4d* | *Vwa5a* |
| *Rps19* | *Sparc* | *Tpm2* | *Cd93* | *Gng11* | *Phlda1* | *Dbi* | *Cd3d* | *Ifngr1* | *Lsp1* | *Lilrb4a* | *Lilr4b* | *Rgs1* |
| *Rpl13* | *Col3a1* | *Rgs5* | *Egfl7* | *Nsg1* | *Gpnmb* | *Sparc* | *Il2rb* | *Vps37b* | *Cd52* | *Cd14* | *Trem1* | *Ms4a2* |
| *Serpinb5* | *Mmp14* | *Epas1* | *Epas1* | *Lrg1* | *Fabp5* | *Utrn* | *Trdv4* | *Ndfip1* | *Coro1a* | *Mrc1* | *S100a9* | *Kit* |
| *Rpl12* | *Prrx1* | *Myl9* | *Kitl* | *Pard6g* | *Myo5a* | *Mbp* | *Fcer1g* | *Cytip* | *H2-Eb1* | *Atp6v0c* | *Ccr1* | *Fcer1g* |
| *Rpl32* | *Adamts2* | *Tagln* | *Ecscr* | *Ppfibp1* | *Lgals3* | *Art3* | *Arl4c* | *Emb* | *Actb* | *Fcgr3* | *Cebpb* | *Gata2* |
| *Rpl11* | *Mmp2* | *Ebf1* | *Gimap6* | *Rab11a* | *Bcl2* | *Cnp* | *Sytl3* | *Septin1* | *Napsa* | *Cd68* | *Btg1* | *Gnai1* |
| *Rps5* | *Pcolce* | *Fermt2* | *Eng* | *Ccl21a* | *Kit* | *Dag1* | *Vgll4* | *Cotl1* | *Ptprc* | *Lgmn* | *Fxyd5* | *H2afz* |
| *Rps26* | *Fbln2* | *Mustn1* | *Cyyr1* | *Aplp2* | *Eef1a1* | *Mal* | *Rgcc* | *Rgs2* | *Fcer1g* | *Cstb* | *S100a8* | *Mrgprb1* |
| *Urah* | *Col6a2* | *Gucy1a1* | *Ece1* | *Lbp* | *Malat1* | *Pmp* | *Vps37b* | *Crem* | *Rel* | *Ctsb* | *Clec4e* | *Samsn1* |
| *Rpl4* | *Timp2* | *Gucy1b1* | *Rasip1* | *Gimap6* | *Acot1* | *Malat1* | *Cd2* | *Vgll4* | *Alox5ap* | *Ms4a6c* | *Ptprc* | *Osbpl8* |
| *Rps8* | *Nbl1* | *Gng11* | *Mmrn2* | *Arl4a* | *Rpl14* | *Itgb8* | *Neurl3* | *Fam107b* | *Cytip* | *Ms4a6d* | *Acod1* | *Pakap-1* |
| *Rpl10a* | *Ccdc80* | *Rbpms* | *Scarb1* | *Flt4* | *Mitf* | *Pou3f1* | *Itk* | *Il2rg* | *Plek* | *Lilr4b* | *Nlrp3* | *Ccl2* |
| *Rpl8* | *Lum* | *Ifitm3* | *Emcn* | *Cavin2* | *Adgrg1* | *Sorbs1* | *Prkar1a* | *Stk17b* | *H2afz* | *Psap* | *Ccrl2* | *Suco* |
| *Rps7* | *Htra1* | *Pdgfa* | *Ptprb* | *Tm4sf1* | *Dbi* | *Limch1* | *Prkacb* | *Trbc2* | *Gm2a* | *Cd53* | *Pim1* | *Slc18a2* |
| *Rpl6* | *Ppic* | *Pde3a* | *Ctla2a* | *Cdh5* | *Tm4sf1* | *Fxyd1* | *Acpp* | *Neurl3* | *Syngr2* | *Srgn* | *Sgms2* | *Cma2* |
| *Rps18* | *Zbtb20* | *Itgb1* | *Cav1* | *F11r* | *Atp1a1* | *Secisbp2l* | *Ankrd11* | *Rac2* | *Lcp1* | *Cd44* | *Fth1* | *Stk17b* |
| *Krt15* | *Cd63* | *Sncg* | *Slfn5* | *Ece1* | *Nav2* | *Sptbn1* | *Fosl2* | *Dennd4a* | *Arl4c* | *Zeb2* | *Cd53* | *Lat2* |
| *Rpl5* | *Plpp3* | *Rgs4* | *Ly6e* | *Arhgap29* | *Ncald* | *Cadm4* | *Cd247* | *Itpkb* | *Cd83* | *Ctsc* | *Mxd1* | *Tph1* |
| *Rps15* | *Antxr1* | *Phlda1* | *Ramp2* | *Lcn2* | *Mgll* | *Dst* | *Fxyd5* | *Tmsb4x* | *Actg1* | *Spi1* | *Ets2* | *Csf2rb* |
| *Dmkn* | *Aspn* | *Crispld2* | *Adgrl4* | *Nr2f2* | *Pmel* | *Megf9* | *Srgn* | *Mbnl1* | *Cst3* | *Wfdc17* | *Cd44* | *Rab27b* |
| *Nfib* | *Aebp1* | *Id3* | *Abcg2* | *Kdr* | *mt-Nd1* | *Emp2* | *Ctsw* | *Il7r* | *Sh3bgrl3* | *Efhd2* | *Tpd52* | *Hs6st2* |
| *Rps20* | *Clmp* | *Mef2c* | *Sptbn1* | *Nfat5* | *Auts2* | *Csrp1* | *Laptm5* | *Cd3g* | *Ctsz* | *Cyba* | *Samsn1* | *Slc6a4* |
| *Rps16* | *Col5a1* | *Chchd2* | *Tcf4* | *Rhoj* | *St3gal6* | *Egfl8* | *Slc38a1* | *Itk* | *Spi1* | *Mafb* | *Lmnb1* | *Fam107b* |
| *Rps27a* | *Nupr1* | *Timp3* | *Mast4* | *Apold1* | *Marcks* | *Mpz* | *Itgae* | *Ikzf3* | *Csf2ra* | *Calm1* | *Slc15a3* | *Ndrg1* |
| *Rpl23* | *Olfml3* | *Myl6* | *Arhgap31* | *Lyve1* | *Myo10* | *Sema3b* | *Myo1e* | *Crip1* | *Ucp2* | *Npc2* | *Hcar2* | *Csf2rb2* |
| *Rpl27a* | *Fbn1* | *Adamts1* | *Podxl* | *Rgs16* | *Gm47316* | *Itga6* | *Zeb1* | *Skap1* | *Cd86* | *Ccl9* | *Il1r2* | *Cd55* |
| *Rpl7* | *Tnfaip6* | *Pcp4l1* | *Ly6a* | *Gpm6a* | *Mlana* | *Map1b* | *Runx3* | *Itgae* | *Cyba* | *Sdcbp* | *Alox5ap* | *Adora3* |
| *Rpl18* | *Gsn* | *S1pr3* | *S1pr1* | *Ptma* | *Ppia* | *Vim* | *Dennd4a* | *Ptpn22* | *Efhd2* | *Clec4a2* | *Emilin2* | *Tmem64* |
| *Rpl13a* | *Dpt* | *Kitl* | *Tspan13* | *Bcr* | *Sox6* | *Ank3* | *Stk17b* | *Tnfaip3* | *Dennd4a* | *Arl4c* | *Ifitm1* | *Sla* |
| *Rps17* | *Lrp1* | *Serpine2* | *Ets1* | *Fth1* | *mt-Atp6* | *Scd1* | *Smad7* | *Ppp1r16b* | *Atp2b1* | *Pira2* | *Ifitm2* | *Basp1* |
| *Rpl15* | *Mfap2* | *Filip1l* | *Ebf1* | *Sptbn1* | *Rexo2* | *Gldn* | *Gna13* | *P2ry10* | *S100a11* | *Mcl1* | *Pik3ap1* | *Ago2* |
| *Rps10* | *Rcn3* | *Ptp4a3* | *Nfib* | *Tshz2* | *Mlph* | *Cldn19* | *Tcrg-C1* | *Aebp2* | *Lyz2* | *Tmsb4x* | *Csf3r* | *Arhgdib* |
| *Rpl28* | *Igfbp6* | *Tinagl1* | *Tshz2* | *Klf2* | *Cadm1* | *Cdh19* | *Grap2* | *Nabp1* | *Arhgdib* | *Ctsz* | *Il1rn* | *Gask1b* |
| *Jup* | *Rarres2* | *Cystm1* | *Tspan7* | *Emcn* | *mt-Co3* | *Col5a3* | *Gem* | *Fxyd5* | *Cd53* | *Csf1r* | *Slc7a11* | *Rgs2* |
| *Rpl3* | *Rnase4* | *Flna* | *Pdlim1* | *Reln* | *Syt4* | *Cnn3* | *Klrd1* | *Akap13* | *Plbd1* | *Sirpa* | *Grina* | *Prkacb* |
| *Rpl30* | *Ddr2* | *Map3k20* | *Kdr* | *Serinc3* | *Cdk2* | *Lgi4* | *Klrb1b* | *Rnf125* | *Pfn1* | *Ifi207* | *Cd52* | *Plgrkt* |
| *Eef1b2* | *Scara5* | *Slc25a4* | *Cdh13* | *Scn1b* | *Rpl17* | *Arpc1a* | *Ets1* | *Btg1* | *Mgl2* | *Lcp1* | *Gmfg* | *Gfpt1* |
| *Rps12* | *Lpar1* | *Zfhx3* | *Apold1* | *Dusp2* | *Rpl13* | *Col28a1* | *Avil* | *Traf1* | *Csf2rb* | *Dab2* | *Marcksl1* | *Cited2* |
| *Col17a1* | *Col6a3* | *Ppp1r12a* | *Hspb1* | *Gngt2* | *Mgat4b* | *Dmd* | *Ubald2* | *Ptprcap* | *Vim* | *Vim* | *Wfdc17* | *Chpt1* |

**Supplementary Table 2 related to Supplementary Fig. 1c cell types proportion**

| Timepoint | Celltype | cell_count_Control | total_cells_Control | proportion_Control | cell_count_Expande | total_cells_Expande | proportion_Expanded | p_value | p_signif |
|---|---|---|---|---|---|---|---|---|---|
| Day1 | CD4+/CD8+ T cells | 491 | 10768 | 0.0455980683506686 | 858 | 7060 | 0.121529745042493 | 3,41E-64 | *** |
| Day2 | CD4+/CD8+ T cells | 462 | 6139 | 0.0752565564424173 | 212 | 4108 | 0.0516066212268744 | 2,70E+08 | *** |
| Day4 | CD4+/CD8+ T cells | 234 | 5505 | 0.0425068119891008 | 195 | 2736 | 0.0712719298245614 | 4,18E+06 | *** |
| Day1 | CD4- CD8- T cells | 208 | 10768 | 0.0193164933135215 | 267 | 7060 | 0.0378186968838527 | 8,98E+00 | *** |
| Day2 | CD4- CD8- T cells | 165 | 6139 | 0.0268773415865776 | 166 | 4108 | 0.0404089581304771 | 0.000184093418008187 | *** |
| Day4 | CD4- CD8- T cells | 55 | 5505 | 0.00999091734786558 | 98 | 2736 | 0.0358187134502924 | 5,81E-02 | *** |
| Day1 | Endothelial cells | 530 | 10768 | 0.0492199108469539 | 611 | 7060 | 0.0865439093484419 | 3,18E-09 | *** |
| Day2 | Endothelial cells | 289 | 6139 | 0.047076071021339 | 109 | 4108 | 0.0265335929892892 | 1,77E+07 | *** |
| Day4 | Endothelial cells | 433 | 5505 | 0.0786557674841054 | 204 | 2736 | 0.0745614035087719 | 0.540782434635887 | |
| Day1 | Fibroblasts | 6230 | 10768 | 0.578566121842496 | 2056 | 7060 | 0.291218130311615 | 1,70E-295 | *** |
| Day2 | Fibroblasts | 2725 | 6139 | 0.443883368626812 | 1036 | 4108 | 0.252190847127556 | 1,80E-72 | *** |
| Day4 | Fibroblasts | 2679 | 5505 | 0.486648501362398 | 693 | 2736 | 0.253289473684211 | 2,54E-77 | *** |
| Day1 | Keratinocytes | 908 | 10768 | 0.0843239227340268 | 1792 | 7060 | 0.253824362606232 | 4,87E-195 | *** |
| Day2 | Keratinocytes | 402 | 6139 | 0.0654829776836618 | 284 | 4108 | 0.0691333982473223 | 0.493809393310025 | |
| Day4 | Keratinocytes | 343 | 5505 | 0.0623069936421435 | 429 | 2736 | 0.156798245614035 | 1,84E-29 | *** |
| Day1 | Langerhans cells | 845 | 10768 | 0.0784732540861813 | 364 | 7060 | 0.0515580736543909 | 3,40E+02 | *** |
| Day2 | Langerhans cells | 585 | 6139 | 0.0952923928978661 | 307 | 4108 | 0.0747322297955209 | 0.000340475175338264 | *** |
| Day4 | Langerhans cells | 540 | 5505 | 0.0980926430517711 | 177 | 2736 | 0.0646929824561404 | 5,04E+07 | *** |
| Day1 | Lymphatic cells | 134 | 10768 | 0.012444279346211 | 222 | 7060 | 0.0314447592067989 | 1,20E-04 | *** |
| Day2 | Lymphatic cells | 164 | 6139 | 0.026714448607265 | 139 | 4108 | 0.0338364167478092 | 0.0427397569652064 | * |
| Day4 | Lymphatic cells | 92 | 5505 | 0.0167120799273388 | 83 | 2736 | 0.0303362573099415 | 7,53E+09 | *** |
| Day1 | Macrophages | 1008 | 10768 | 0.0936106983655275 | 457 | 7060 | 0.0647308781869688 | 7,97E+02 | *** |
| Day2 | Macrophages | 808 | 6139 | 0.131617527284574 | 1230 | 4108 | 0.299415774099318 | 2,34E-82 | *** |
| Day4 | Macrophages | 701 | 5505 | 0.127338782924614 | 447 | 2736 | 0.163377192982456 | 1,01E+09 | *** |
| Day1 | Mast cells | 90 | 10768 | 0.00835809806835067 | 94 | 7060 | 0.013314447592068 | 0.00176816556202027 | ** |
| Day2 | Mast cells | 64 | 6139 | 0.0104251506760059 | 53 | 4108 | 0.0129016553067186 | 0.288451645955805 | |
| Day4 | Mast cells | 29 | 5505 | 0.00526793823796549 | 13 | 2736 | 0.00475146198830409 | 0.884054440938706 | |
| Day1 | Melanocytes | 50 | 10768 | 0.00464338781575037 | 76 | 7060 | 0.0107648725212465 | 2,86E+08 | *** |
| Day2 | Melanocytes | 153 | 6139 | 0.0249226258348265 | 87 | 4108 | 0.0211781888997079 | 0.24538918634822 | |
| Day4 | Melanocytes | 48 | 5505 | 0.00871934604904632 | 50 | 2736 | 0.0182748538011696 | 0.000251562833697127 | *** |
| Day1 | Monocytes | 5 | 10768 | 0.000464338781575037 | 17 | 7060 | 0.00240793201133144 | 0.000681051577079814 | *** |
| Day2 | Monocytes | 17 | 6139 | 0.00276918064831406 | 264 | 4108 | 0.0642648490749757 | 2,25E-63 | *** |
| Day4 | Monocytes | 15 | 5505 | 0.00272479564032698 | 148 | 2736 | 0.054093567251462 | 1,83E-41 | *** |
| Day1 | Schwann cells | 93 | 10768 | 0.00863670133729569 | 157 | 7060 | 0.0222379603399433 | 6,98E+00 | *** |
| Day2 | Schwann cells | 163 | 6139 | 0.0265515556279524 | 126 | 4108 | 0.0306718597857838 | 0.240472063239752 | |
| Day4 | Schwann cells | 186 | 5505 | 0.0337874659400545 | 144 | 2736 | 0.0526315789473684 | 5,14E+09 | *** |
| Day1 | Vascular smooth muscle | 176 | 10768 | 0.0163447251114413 | 89 | 7060 | 0.0126062322946176 | 0.0506845442690732 | |
| Day2 | Vascular smooth muscle | 142 | 6139 | 0.023130803062388 | 95 | 4108 | 0.0231256085686465 | 0.999999999999999 | |
| Day4 | Vascular smooth muscle | 150 | 5505 | 0.0272479564032698 | 55 | 2736 | 0.0201023391812865 | 0.0592517190214648 | |

**Supplemantary Table 3 related to Fig. 1d scRNAseq marker genes for all fibroblast cluster**

| FIB1 | FIB2 | FIB3 | FIB4 | FIB5 | FIB6 | FIB7 | FIB8 | DS | DP1 | DP2 |
|---|---|---|---|---|---|---|---|---|---|---|
| Qpct | Aebp1 | Sparc | Tnc | Cygb | Apod | Cxcl1 | Rpl41 | Pmepa1 | Coch | Ptma |
| Col3a1 | Col1a2 | Col1a1 | Pdpn | Serpinf1 | Ccl11 | Runx1 | Rps24 | Col6a3 | Crabp1 | Igfbp3 |
| Wnt5a | Col1a1 | Aebp1 | Txn1 | Clec3b | Gpc3 | Gpx3 | Rps26 | Col4a1 | Gldn | Serpine2 |
| S100a4 | Col3a1 | Lum | Sod2 | Gsn | Gsn | Crlf1 | Rps12 | Slit2 | Gpm6b | Wif1 |
| Thbs2 | Sparc | Ctsk | Pkm | Gpx3 | Cp | Chodl | Rps4x | Cd9 | Tnmd | Bcl2 |
| Col5a1 | Fgl2 | Rnase4 | Ifitm3 | Serping1 | Dcn | Tnfrsf12a | Rpl13 | Postn | Fmod | Ifitm1 |
| Mmp2 | Rnase4 | Col3a1 | Nme1 | Nfib | Igfbp4 | Des | Rps8 | Cdh11 | Trps1 | Prlr |
| Abi3bp | Pcolce2 | Cpz | Ran | Ly6a | Ebf2 | Pakap-1 | Rplp1 | Bgn | Prelp | Ebf1 |
| Nbl1 | Lum | Col1a2 | Slc39a14 | Cxcl12 | Fth1 | Ncl | Rps3a1 | Dkk2 | Tcf4 | Malat1 |
| Tnxb | Uap1 | Pcolce2 | Esd | C1s1 | Apoe | Flnc | Rps20 | Col4a2 | Cd9 | Zfand5 |
| Tgfbi | Ctsk | Gadd45g | Plaur | Selenop | Smoc2 | Slc43a3 | Serf2 | Gas1 | Col11a1 | Slc26a7 |
| Grem1 | Mmp2 | Atf3 | Arpc3 | Dpep1 | Timp3 | Pax7 | Rpl34 | Neat1 | Igsf10 | Actb |
| Sparc | Mt1 | Ctsh | Rplp2 | Ifi205 | Nr2f2 | Asb5 | Rpl39 | Vim | Masp1 | Cfl1 |
| Efemp1 | Col5a2 | Tgfbi | Fst | Ly6c1 | Steap4 | Apoe | Rpsa | Fgfr1 | Igfbp7 | Inhba |
| Rnase4 | Tgfbi | Clec3b | Arpc1b | Lpl | Vit | Cfh | Fth1 | Vcan | Ogn | Npm1 |
| Fbln1 | Mt2 | Osr2 | Rpl27 | C7 | Smim41 | Nop58 | Rpl37a | Cyp1b1 | Pi16 | Hnrnpa2b1 |
| Fam102b | Psap | Rarres2 | Mmp3 | Ifi211 | Egr1 | Cux1 | Fau | Col6a1 | Nav3 | H3f3b |
| Igfbp6 | Tnxb | Meg3 | Rbm3 | Ogn | Cxcl1 | Hmcn2 | Rpl32 | Ptma | Dkk2 | a |
| Ly6a | Gpnmb | Man1a | Rpl24 | Dpt | C1s1 | Pdlim4 | Rpl23 | Lmna | Igfbp4 | Fgfr1 |
| Sema3a | Cpz | Gpnmb | Psma7 | Slfn5 | Ecm1 | Peg3 | Nme2 | Cyth3 | Mxra8 | Pappa2 |
| Col1a1 | Timp2 | Klf2 | Rack1 | Gfpt2 | Matn2 | Sdc4 | Rps10 | Nrp1 | Fbln5 | Hsp90ab1 |
| Loxl2 | Sulf2 | Npc2 | Fxyd5 | Ebf1 | H3f3b | Gm14636 | Rps5 | Cilp | Srpx2 | Rbm39 |
| Apcdd1 | Adamts2 | Mt1 | Mrpl52 | Slc43a3 | Vwa1 | Pde4b | Rpl11 | Thy1 | Sparcl1 | Lef1 |
| Nt5e | Gas6 | Sulf2 | Rpl23 | C3 | Plxdc2 | Notch3 | Rps3 | Pdgfra | Lepr | Tmem176b |
| Aebp1 | Serpina3n | Gas6 | Rps3 | Apod | Ifitm3 | Arid5b | Rpl30 | Tpbg | Odc1 | Actg1 |
| Cdh4 | Ctsh | Sectm1a | Uqcrq | Egfr | Sox9 | Erfe | Rps2 | Col6a2 | Syne1 | Mdk |
| Cd47 | Ccdc80 | Serpina3n | Sh3pxd2b | Ifi204 | Mgp | Cxcl2 | Rps16 | F2r | Rspo4 | Tpm2 |
| Dcn | Dcn | Jun | Eef1g | Itm2b | Sned1 | Tanc2 | Rps23 | Igf1 | Enpp2 | Rsrp1 |
| Ccbe1 | Clec3b | Fos | Btf3 | Ecm1 | Rasgrp2 | Vcam1 | Rplp2 | Aqp1 | Hsd11b1 | Srrm2 |
| Gda | C4b | Zfp36 | Gapdh | Igfbp6 | Ifitm2 | Srrm2 | Rpl27a | Serpinf1 | Megf6 | Rps19 |
| Timp2 | Npc2 | Mt2 | Psmb3 | Dcn | Jun | Sqstm1 | Ifitm3 | Lama2 | Abca8b | Spon1 |
| Lum | Tnfaip6 | Bhlhe40 | Clic1 | Lbp | Sulf1 | Pmepa1 | Tpt1 | Skil | Plxdc1 | Enpp2 |
| Col1a2 | Selenop | Rian | Psmb5 | Gas6 | Tshz2 | Ddx5 | Rplp0 | Ubc | Maff | Apoe |
| Sulf2 | Fstl1 | Creb3l1 | Mpp6 | Zfhx4 | Phlda1 | Uck2 | Cox6c | Lox | Fscn1 | Hnrnpu |
| Cadm3 | Gns | Il1r2 | Rps19 | Lgi2 | Abca8a | Cd82 | Rpl10a | Tuba1a | Rps16 | Ppia |
| Slc27a6 | Col5a1 | C1qtnf3 | Timp1 | Tnxb | Egr3 | Itm2a | Rps15a | Mafb | Gfra1 | Tmem176a |
| Pcolce2 | Man1a | Ly6c1 | Cstb | C4b | Pdzd2 | Eif5 | Rpl35a | Crip1 | Itga11 | Ncam1 |
| Cd109 | Itgbl1 | Creb5 | Rpsa | Cxcl1 | Fos | Gnl3 | Rpl6 | Mfap4 | Tmsb4x | Runx3 |
| Gpnmb | Gja1 | Ndufa4l2 | Glrx | Hk2 | Cldn1 | Hnrnpa2b1 | Tmsb10 | Dpysl3 | Crispld2 | Trps1 |
| Entpd1 | Efemp1 | Prdx6 | Cd44 | Ace | Plk2 | Tubb2b | Rps11 | Loxl1 | Abca8a | Slc6a6 |
| Col6a2 | Serping1 | Cldn10 | Ppia | Ccdc80 | Foxd1 | Odc1 | Rpl28 | H3f3b | Ctsl | Eif4a1 |
| Igfbp2 | Cd34 | Selenop | Cd63 | Scn7a | Spp1 | Rbm39 | Rpl19 | Pten | Mafb | Srsf3 |
| Col6a1 | Hexa | Serpinh1 | Rps2 | Gpc3 | Tm4sf1 | Tenm4 | Rpl9 | Sgk1 | Sdc4 | Ndnf |
| Aldh3a1 | Fbn1 | C1qtnf6 | Psma2 | Serpinb6a | Cfh | Tln2 | Lgals1 | Igfbp7 | Postn | Dek |
| Fxyd6 | Cdh4 | Dcn | Psmd11 | Phlda1 | Sdc2 | Cdh15 | Rps19 | Pcdh7 | Asap1 | Srsf7 |
| Pam | Creb3l1 | Ccdc80 | Sdc4 | Col14a1 | Lsamp | Srsf7 | S100a6 | Ltbp2 | Abca9 | Rps8 |
| Ppic | Adgrd1 | Socs3 | Riok3 | Adamts5 | Tgfbi | Ncam1 | Rpl14 | Ppia | Kera | Igfbp4 |
| Igfbp5 | Fkbp9 | Cd34 | Psma6 | Cxcl14 | Pabpc1 | Txnrd1 | Rpl22 | Hsp90ab1 | Lmna | Cox4i1 |
| F13a1 | Adamts4 | Ppic | Fth1 | F3 | Trf | Cd44 | Rpl37 | Mmp16 | Thy1 | Col23a1 |
| Man1a | Pam | Dapk1 | Mapkapk2 | Mgst1 | Csmd1 | Sulf1 | Rps27a | Lgals3 | Pth1r | Set |

**Supplementary  Table 4 related to Supplementary Fig. 1d fibroblasts cluster proportions**

| Timepoint | Celltype | cell_count_Control | total_cells_Control | proportion_Control | cell_count_Expande | total_cells_Expande | proportion_Expanded | p_value | p_signif |
|---|---|---|---|---|---|---|---|---|---|
| Day1 | Fibroblast 1 | 1042 | 6230 | 0.167255216693419 | 230 | 2056 | 0.111867704280156 | 1,90E+05 | *** |
| Day2 | Fibroblast 1 | 294 | 2725 | 0.107889908256881 | 91 | 1036 | 0.0878378378378378 | 0.079748563635882 | |
| Day4 | Fibroblast 1 | 399 | 2679 | 0.148936170212766 | 32 | 693 | 0.0461760461760462 | 8,20E+01 | *** |
| Day1 | Fibroblast 2 | 1371 | 6230 | 0.220064205457464 | 94 | 2056 | 0.0457198443579767 | 6,36E-58 | *** |
| Day2 | Fibroblast 2 | 355 | 2725 | 0.130275229357798 | 38 | 1036 | 0.0366795366795367 | 8,57E-03 | *** |
| Day4 | Fibroblast 2 | 489 | 2679 | 0.182530795072788 | 17 | 693 | 0.0245310245310245 | 5,64E-11 | *** |
| Day1 | Fibroblast 3 | 1438 | 6230 | 0.230818619582665 | 292 | 2056 | 0.142023346303502 | 1,14E-03 | *** |
| Day2 | Fibroblast 3 | 454 | 2725 | 0.166605504587156 | 75 | 1036 | 0.0723938223938224 | 1,68E+01 | *** |
| Day4 | Fibroblast 3 | 543 | 2679 | 0.202687569988802 | 57 | 693 | 0.0822510822510823 | 2,25E+01 | *** |
| Day1 | Fibroblast 4 | 25 | 6230 | 0.00401284109149278 | 154 | 2056 | 0.0749027237354086 | 3,43E-67 | *** |
| Day2 | Fibroblast 4 | 67 | 2725 | 0.0245871559633028 | 83 | 1036 | 0.0801158301158301 | 1,58E+00 | *** |
| Day4 | Fibroblast 4 | 39 | 2679 | 0.0145576707726764 | 52 | 693 | 0.075036075036075 | 6,36E-04 | *** |
| Day1 | Lower dermis Fibroblast 1 | 320 | 6230 | 0.0513643659711075 | 22 | 2056 | 0.0107003891050584 | 1,55E-01 | *** |
| Day2 | Lower dermis Fibroblast 1 | 189 | 2725 | 0.0693577981651376 | 9 | 1036 | 0.00868725868725869 | 1,82E+01 | *** |
| Day4 | Lower dermis Fibroblast 1 | 69 | 2679 | 0.0257558790593505 | 8 | 693 | 0.0115440115440115 | 0.0366390637117865 | * |
| Day1 | Lower dermis Fibroblast 2 | 89 | 6230 | 0.0142857142857143 | 161 | 2056 | 0.0783073929961089 | 1,54E-34 | *** |
| Day2 | Lower dermis Fibroblast 2 | 48 | 2725 | 0.0176146788990826 | 33 | 1036 | 0.0318532818532819 | 0.0104197196504766 | * |
| Day4 | Lower dermis Fibroblast 2 | 31 | 2679 | 0.0115714818962299 | 25 | 693 | 0.0360750360750361 | 1,48E+09 | *** |
| Day1 | Lower dermis Fibroblast 3 | 41 | 6230 | 0.00658105939004815 | 12 | 2056 | 0.00583657587548638 | 0.835502866281306 | |
| Day2 | Lower dermis Fibroblast 3 | 87 | 2725 | 0.0319266055045872 | 19 | 1036 | 0.0183397683397683 | 0.0324379468602921 | * |
| Day4 | Lower dermis Fibroblast 3 | 51 | 2679 | 0.019036954087346 | 18 | 693 | 0.025974025974026 | 0.317691068650309 | |
| Day1 | Lower dermis Fibroblast 4 | 45 | 6230 | 0.007223113964687 | 29 | 2056 | 0.0141050583657588 | 0.00612725165539221 | ** |
| Day2 | Lower dermis Fibroblast 4 | 147 | 2725 | 0.0539449541284404 | 81 | 1036 | 0.0781853281853282 | 0.00679893896526701 | ** |
| Day4 | Lower dermis Fibroblast 4 | 36 | 2679 | 0.013437849944009 | 20 | 693 | 0.0288600288600289 | 0.00770071120505213 | ** |
| Day1 | DP1 | 725 | 6230 | 0.116372391653291 | 633 | 2056 | 0.307879377431907 | 1,14E-77 | *** |
| Day2 | DP1 | 336 | 2725 | 0.123302752293578 | 277 | 1036 | 0.267374517374517 | 2,00E-12 | *** |
| Day4 | DP1 | 273 | 2679 | 0.101903695408735 | 118 | 693 | 0.17027417027417 | 7,65E+07 | *** |
| Day1 | DP2 | 271 | 6230 | 0.0434991974317817 | 97 | 2056 | 0.0471789883268482 | 0.521805757898221 | |
| Day2 | DP2 | 447 | 2725 | 0.164036697247706 | 212 | 1036 | 0.204633204633205 | 0.00400507734030606 | ** |
| Day4 | DP2 | 434 | 2679 | 0.162000746547219 | 270 | 693 | 0.38961038961039 | 3,86E-25 | *** |
| Day1 | DS | 863 | 6230 | 0.138523274478331 | 332 | 2056 | 0.16147859922179 | 0.0113136615966318 | * |
| Day2 | DS | 301 | 2725 | 0.11045871559633 | 118 | 1036 | 0.113899613899614 | 0.809074076232504 | |
| Day4 | DS | 315 | 2679 | 0.117581187010078 | 76 | 693 | 0.10966810966811 | 0.607691441547101 | |

**Supplemantary Table 5 related to Fig. 3a differentially expressed genes in CTRL and EXP**

| Gene | p_val | avg_log2FC | pct.1 | pct.2 | p_val_adj |
|---|---|---|---|---|---|
| Ablim1 | 0 | -0.413915834 | 0,16458333 | 0,38125 | 0 |
| Aebp1 | 0 | -0.492724921 | 0,49861111 | 0,59513889 | 0 |
| Anxa5 | 0 | -0.278790080 | 0,59652778 | 0,06597222 | 0 |
| Arpc1b | 0 | 0.334404518 | 0,63958333 | 0,57777778 | 0 |
| Ccdc80 | 0 | -0.501127484 | 0,48541667 | 0,6125 | 0 |
| Cd302 | 0 | 0.303902300 | 0,63402778 | 0,56388889 | 0 |
| Cd34 | 0 | -0.491785958 | 0,41319444 | 0,57013889 | 0 |
| Cirbp | 0 | 0.363072834 | 0,41805556 | 0,16319444 | 0 |
| Clec3b | 0 | -0.795340682 | 0,22638889 | 00.07 | 0 |
| Col15a1 | 0 | -0.516787379 | 0,31319444 | 0,5375 | 0 |
| Col1a1 | 0 | -0.526758006 | 0,61180556 | 0,67430556 | 0 |
| Col1a2 | 0 | -0.491766657 | 0,62708333 | 0,67708333 | 0 |
| Col3a1 | 0 | -0.520837787 | 0,57361111 | 0,66805556 | 0 |
| Col5a1 | 0 | -0.431886407 | 0,45972222 | 0,575 | 0 |
| Col5a2 | 0 | -0.342668006 | 0,05902778 | 0,64444444 | 0 |
| Cpz | 0 | -0.570495404 | 0,18541667 | 0,40763889 | 0 |
| Creb3l1 | 0 | -0.426391593 | 0,38263889 | 0,51736111 | 0 |
| Creb5 | 0 | -0.547960710 | 0,15555556 | 0,39236111 | 0 |
| Ctsh | 0 | -0.535039820 | 0,24097222 | 0,45277778 | 0 |
| Efemp1 | 0 | -0.529939487 | 00.39 | 0,46736111 | 0 |
| Fcgr2b | 0 | 0.376633280 | 0,20763889 | 0.052 | 0 |
| Fgl2 | 0 | -0.544369013 | 0,41944444 | 0,52152778 | 0 |
| Fkbp9 | 0 | -0.362534560 | 0,44930556 | 0,55069444 | 0 |
| Glt8d2 | 0 | -0.396903789 | 0,10902778 | 0,32152778 | 0 |
| Hexa | 0 | -0.460516878 | 00.53 | 0,52708333 | 0 |
| Hnrnpa3 | 0 | 0.301971130 | 0,64791667 | 0,55902778 | 0 |
| Ifitm1 | 0 | 0.597438780 | 0,36319444 | 0,13680556 | 0 |
| Ifitm2 | 0 | 0.252738988 | 0,68402778 | 0,06736111 | 0 |
| Ifitm3 | 0 | 0.289108264 | 0,68125 | 0,66597222 | 0 |
| Itgbl1 | 0 | -0.521792652 | 0,31458333 | 0,51666667 | 0 |
| Lum | 0 | -0.501227347 | 0,47291667 | 0,59375 | 0 |
| Man1a | 0 | -0.411568752 | 0,04305556 | 0,54375 | 0 |
| Meg3 | 0 | -0.776330767 | 0,26875 | 0,49375 | 0 |
| Mmp27 | 0 | -0.416347415 | 0,12916667 | 0,33958333 | 0 |
| Mrpl52 | 0 | 0.331674014 | 0,575 | 0,46041667 | 0 |
| Ndufa4l2 | 0 | -0.547705254 | 0,27361111 | 0,48888889 | 0 |
| Pcolce2 | 0 | -0.759636902 | 0,225 | 0,48680556 | 0 |
| Pdgfrl | 0 | -0.409083292 | 0,27361111 | 0,45625 | 0 |
| Pkm | 0 | 0.374201123 | 0,63263889 | 0,54791667 | 0 |
| Ppia | 0 | 0.258905565 | 0,68819444 | 0,67777778 | 0 |
| Psap | 0 | -0.301281756 | 0,61319444 | 0,64722222 | 0 |
| Psmb3 | 0 | 0.352586586 | 0,58402778 | 0,425 | 0 |
| Rbm3 | 0 | 0.479578091 | 0,68333333 | 0,62986111 | 0 |
| Rcn3 | 0 | -0.376541455 | 0,4625 | 0,57847222 | 0 |
| Rian | 0 | -0.653295138 | 0,15486111 | 0,38402778 | 0 |
| Rnase4 | 0 | -0.388062321 | 0,58958333 | 0,62569444 | 0 |
| Runx1 | 0 | 0.438559478 | 0,05694444 | 0,37013889 | 0 |
| Selenop | 0 | -0.456031362 | 0,48888889 | 0,59166667 | 0 |
| Sema3b | 0 | -0.349206100 | 0,08055556 | 0,27708333 | 0 |
| Serpinh1 | 0 | -0.360311264 | 00.09 | 0,67083333 | 0 |
| Slc6a6 | 0 | 0.512089833 | 0,49513889 | 0,21180556 | 0 |
| Sod2 | 0 | 0.552774544 | 0,06180556 | 0,47013889 | 0 |
| Sparc | 0 | -0.388791069 | 0,64236111 | 0,68194444 | 0 |
| Spon2 | 0 | -0.361487862 | 0.062 | 0,26527778 | 0 |
| Sulf2 | 0 | -0.494393929 | 0,21041667 | 0,42638889 | 0 |
| Timp1 | 0 | 0.585834948 | 0,04444444 | 0,23541667 | 0 |
| Tnc | 0 | 0.947745955 | 0,39791667 | 0,09305556 | 0 |
| Tnxb | 0 | -0.516066034 | 0,28333333 | 0,47777778 | 0 |
| Txn1 | 0 | 0.276601454 | 0,06736111 | 0,65138889 | 0 |
| Uap1 | 0 | -0.503993037 | 0,39930556 | 0,53541667 | 0 |
| Vkorc1 | 0 | -0.377420281 | 0,38819444 | 0,5375 | 0 |
| Gpnmb | 1,36E-304 | -0.492375330 | 0,28263889 | 0,04722222 | 4,40E-300 |
| Nme1 | 1,84E-304 | 0.357283602 | 0,53194444 | 0,36527778 | 5,93E-300 |
| Gas6 | 2,52E-303 | -0.560561854 | 0,30138889 | 0,49861111 | 8,12E-299 |
| Dapk1 | 1,48E-300 | -0.415304849 | 0,23680556 | 0,42638889 | 4,78E-296 |
| Slc25a5 | 4,15E-295 | 0.303448976 | 0,62222222 | 0,51944444 | 1,34E-290 |
| Uqcrq | 3,48E-294 | 0.267663299 | 0,60347222 | 0,05347222 | 1,12E-288 |
| Cdh4 | 5,05E-292 | -0.424757334 | 00.24 | 0,36736111 | 1,63E-287 |
| C1qtnf3 | 8,48E-289 | -0.491330960 | 0,13958333 | 0,3625 | 2,74E-285 |
| Mirg | 1,45E-288 | -0.407720643 | 0.063 | 0,225 | 4,68E-284 |
| Fn1 | 1,78E-286 | -0.518706818 | 0,26527778 | 0,45416667 | 5,74E-282 |
| Esd | 1,42E-282 | 0.317728630 | 0,05416667 | 0,45 | 4,60E-278 |
| Celf2 | 3,48E-282 | -0.366687119 | 0,38263889 | 0,52569444 | 1,12E-277 |
| Cfl1 | 4,82E-282 | 0.260705650 | 0,66736111 | 0,06180556 | 1,56E-277 |
| Snhg3 | 6,61E-282 | 0.332058170 | 0,44791667 | 0,22361111 | 2,13E-277 |
| B830012L14I | 7,66E-282 | -0.442312654 | 0.087 | 0,25486111 | 2,47E-277 |
| Hsd11b1 | 1,18E-282 | 0.440809363 | 0,47777778 | 0,30347222 | 3,80E-277 |
| Mfap2 | 1,19E-280 | -0.378200341 | 0,41041667 | 0,54930556 | 3,83E-276 |
| Npc2 | 4,15E-279 | -0.251040358 | 0,60486111 | 0,63958333 | 1,34E-274 |
| Psma7 | 1,69E-277 | 0.303629640 | 0,58472222 | 0,48263889 | 5,47E-272 |
| Ppic | 1,55E-274 | -0.294502314 | 0,54305556 | 0,06111111 | 4,99E-271 |
| Gapdh | 2,25E-270 | 0.256851782 | 0,64236111 | 0,6 | 7,27E-266 |
| Psma2 | 1,69E-268 | 0.310689760 | 0,57986111 | 0,43263889 | 5,46E-264 |
| a | 1,07E-267 | 0.429926931 | 0,2625 | 0,0875 | 3,46E-263 |

| | | | | | |
|---|---|---|---|---|---|
| Crabp1 | 1,47E-266 | 0.608538751 | 0,44097222 | 0,22291667 | 4,76E-262 |
| Oxct1 | 1,82E-265 | -0.349141605 | 0,25902778 | 0,42222222 | 5,87E-261 |
| Mif | 5,93E-263 | 0.333445461 | 0,58611111 | 0,44652778 | 1,91E-258 |
| Fbn1 | 7,47E-264 | -0.346351169 | 0,51041667 | 0,05833333 | 2,41E-258 |
| Fxyd5 | 1,95E-262 | 0.403175696 | 0,4875 | 0,28263889 | 6,30E-258 |
| Nfia | 4,94E-262 | -0.35608924 | 0,41388889 | 0,55069444 | 1,60E-257 |
| Mrc2 | 4,01E-260 | -0.321261495 | 0,22430556 | 0,37708333 | 1,29E-255 |
| Psmb5 | 8,47E-260 | 0.328453667 | 0,52847222 | 0,34305556 | 2,73E-255 |
| Hnrnpd | 4,31E-251 | 0.319499377 | 0,55 | 0,35694444 | 1,39E-247 |
| Cldn10 | 3,31E-250 | -0.37269948 | 0,13263889 | 0,33194444 | 1,07E-245 |
| Cyp2f2 | 4,19E-250 | -0.495347648 | 0,12430556 | 0,32847222 | 1,35E-245 |
| Scpep1 | 3,41E-248 | -0.30411730 | 0,3375 | 0,46180556 | 1,10E-243 |
| P4ha2 | 1,69E-247 | -0.32412804 | 0,2 | 0,35625 | 5,45E-243 |
| Opcml | 2,41E-245 | -0.27159678 | 0.079 | 0,21805556 | 7,80E-241 |
| Pam | 1,02E-244 | -0.311591645 | 0,42291667 | 0,51458333 | 3,31E-240 |
| Osr2 | 3,20E-243 | -0.42492655 | 0,22986111 | 0,41319444 | 1,03E-239 |
| Adgrd1 | 6,96E-244 | -0.38121630 | 0,18402778 | 0,36736111 | 2,25E-238 |
| Itgb5 | 9,48E-242 | -0.30843865 | 0,36666667 | 0,49027778 | 3,06E-237 |
| Fbln1 | 2,23E-241 | -0.41296703 | 0,30833333 | 0,48541667 | 7,20E-237 |
| C1qtnf6 | 4,47E-238 | -0.33253863 | 0,26527778 | 0,39444444 | 1,44E-233 |
| Fstl1 | 5,10E-237 | -0.257904545 | 0,59791667 | 0,06458333 | 1,65E-232 |
| Srsf2 | 3,95E-236 | 0.294552303 | 0,60902778 | 0,48055556 | 1,27E-231 |
| Il1r2 | 3,83E-235 | -0.41083860 | 0,23194444 | 0,42013889 | 1,24E-230 |
| Gxylt2 | 5,45E-234 | -0.36572341 | 0,24444444 | 0,42777778 | 1,76E-229 |
| Mmp23 | 2,14E-232 | -0.31698727 | 0,20347222 | 0,36527778 | 6,90E-228 |
| Tgfbi | 1,94E-231 | -0.42551696 | 0,32569444 | 0,47847222 | 6,28E-227 |
| Ldha | 9,31E-232 | 0.289519521 | 0,60347222 | 0,50902778 | 3,01E-226 |
| Gns | 1,20E-228 | -0.27204620 | 0,42291667 | 0,49375 | 3,88E-224 |
| Ar | 2,70E-228 | -0.35435480 | 0,27708333 | 0,45416667 | 8,71E-224 |
| Coch | 6,76E-227 | 0.750012448 | 0,33541667 | 0,14513889 | 2,18E-222 |
| Odc1 | 1,06E-226 | 0.379250057 | 0,52291667 | 0,35902778 | 3,41E-222 |
| Ecm2 | 2,50E-225 | -0.38624588 | 0,32708333 | 0,5125 | 8,06E-221 |
| Cpne8 | 8,90E-224 | 0.312855744 | 0,42291667 | 0,21875 | 2,87E-219 |
| Hnrnpa1 | 9,55E-224 | 0.271951129 | 0,61527778 | 0,52152778 | 3,08E-221 |
| Hmox1 | 6,17E-221 | 0.394849359 | 0,43333333 | 0,24583333 | 1,99E-216 |
| Flot1 | 5,68E-220 | 0.299867972 | 0,45972222 | 0,28263889 | 1,83E-215 |
| Ran | 1,39E-218 | 0.294428659 | 0,58888889 | 0,47013889 | 4,49E-214 |
| Has1 | 3,46E-215 | -0.44680297 | 0,36180556 | 0,49236111 | 1,12E-210 |
| Fndc1 | 9,31E-215 | -0.37016547 | 0,18611111 | 00.05 | 3,01E-210 |
| Tmed3 | 1,87E-214 | -0.25803970 | 0,47638889 | 0,55208333 | 6,05E-210 |
| Snrpf | 7,14E-214 | 0.291466318 | 0,54305556 | 0,38263889 | 2,31E-209 |
| Gfra1 | 8,65E-213 | 0.277436355 | 0,21875 | 0.089 | 2,79E-208 |
| Ugdh | 1,35E-212 | -0.34169173 | 0,48680556 | 0,54930556 | 4,37E-207 |
| Psma4 | 1,05E-207 | 0.286531779 | 0,54375 | 0,3875 | 3,40E-203 |
| Rcn1 | 1,81E-205 | -0.26888682 | 0,36180556 | 0,46111111 | 5,85E-201 |
| Gsn | 2,90E-203 | -0.25855507 | 0,06388889 | 0,67083333 | 9,38E-199 |
| Dpt | 1,55E-202 | -0.37074861 | 0,41527778 | 0,55347222 | 5,00E-199 |
| Slc38a10 | 1,89E-202 | -0.25080889 | 0,31736111 | 0,40763889 | 6,11E-197 |
| Htra1 | 7,34E-201 | -0.28813259 | 0,52430556 | 0,61875 | 2,37E-196 |
| Cntfr | 7,71E-200 | 0.293604203 | 0,20486111 | 0,08125 | 2,49E-195 |
| Set | 1,48E-199 | 0.275181172 | 0,61388889 | 0,49236111 | 4,79E-195 |
| Cpq | 1,66E-199 | -0.28095306 | 00.04 | 0,40763889 | 5,37E-195 |
| Bax | 1,70E-198 | 0.284375115 | 0,50347222 | 0,32708333 | 5,50E-194 |
| Srsf3 | 2,24E-198 | 0.265841803 | 0,06458333 | 0,55 | 7,22E-194 |
| Loxl2 | 1,02E-197 | -0.33236275 | 0,33888889 | 0,48333333 | 3,30E-193 |
| Ppp1r14a | 2,52E-197 | -0.32405745 | 0,10972222 | 0,28194444 | 8,13E-193 |
| C4b | 3,60E-196 | -0.46850246 | 0,18333333 | 0,34583333 | 1,16E-191 |
| Hpgd | 7,64E-196 | -0.30485769 | 0.094 | 00.33 | 2,47E-192 |
| Prok2 | 1,08E-195 | 0.263241024 | 0,11597222 | 0.021 | 3,47E-193 |
| Igfbp6 | 4,64E-194 | -0.33971848 | 0,45069444 | 0,55625 | 1,50E-189 |
| Aldh3a1 | 4,34E-193 | -0.35298319 | 00.23 | 00.49 | 1,40E-188 |
| Fdps | 7,16E-194 | 0.258154675 | 0,27986111 | 0,10902778 | 2,31E-188 |
| Ace | 1,43E-192 | -0.32220574 | 0,08888889 | 0,23819444 | 4,63E-188 |
| Lbp | 1,27E-190 | 0.381365543 | 0,26111111 | 0,10069444 | 4,10E-186 |
| Epas1 | 5,37E-189 | 0.280863495 | 0,28888889 | 0,11736111 | 1,73E-184 |
| Mafb | 5,16E-187 | 0.398795678 | 0,45486111 | 0,30486111 | 1,67E-182 |
| Aspn | 1,49E-184 | -0.34799786 | 0,45902778 | 0,60138889 | 4,81E-180 |
| Cox5b | 2,03E-182 | 0.257211241 | 0,58055556 | 0,4625 | 6,56E-178 |
| Prdx5 | 2,32E-180 | 0.264472819 | 0,56597222 | 0,46388889 | 7,48E-176 |
| Psmc4 | 4,77E-180 | 0.254283141 | 0,44791667 | 0,2625 | 1,54E-175 |
| Cd55 | 8,16E-180 | -0.30367949 | 0,24027778 | 0,38541667 | 2,63E-175 |
| Id1 | 6,07E-178 | 0.286390062 | 0,28888889 | 0,11944444 | 1,96E-173 |
| Ucp2 | 1,05E-176 | -0.29394633 | 0,16111111 | 00.46 | 3,40E-173 |
| Psmd4 | 1,80E-176 | 0.261082215 | 0,52986111 | 0,37152778 | 5,80E-172 |
| Cox7b | 2,36E-175 | 0.256925285 | 0,58194444 | 0,48125 | 7,61E-171 |
| Cd248 | 8,17E-176 | -0.30817051 | 0,22013889 | 00.54 | 2,64E-170 |
| Tnfrsf1a | 2,94E-174 | 0.263917088 | 0,52152778 | 0,37083333 | 9,49E-170 |
| H2afj | 1,45E-173 | 0.269667717 | 0,53402778 | 0,39236111 | 4,69E-169 |
| Masp1 | 2,03E-174 | 0.352790847 | 0,45625 | 0,32083333 | 6,54E-169 |
| Dkk2 | 1,27E-172 | 0.364405899 | 0,4 | 0,21388889 | 4,11E-168 |
| Tnfaip6 | 1,02E-169 | -0.37686680 | 0,04791667 | 0,59375 | 3,31E-165 |
| Steap4 | 1,33E-169 | 0.304360194 | 0,25625 | 0,10069444 | 4,30E-165 |
| Tppp3 | 1,81E-167 | -0.35594782 | 0,25763889 | 0,40555556 | 5,85E-163 |
| Qpct | 3,00E-167 | -0.36546866 | 00.42 | 0,45625 | 9,68E-163 |
| Kazald1 | 6,95E-167 | -0.27765626 | 0,14097222 | 0,29305556 | 2,24E-162 |
| Psmb6 | 1,95E-163 | 0.250371571 | 0,55972222 | 0,41736111 | 6,29E-159 |
| Serpinf1 | 2,64E-163 | -0.30620008 | 0,44652778 | 0,53125 | 8,51E-159 |

| | | | | | |
|---|---|---|---|---|---|
| Ier2 | 4,73E-164 | 0.304765323 | 0,62083333 | 0,52222222 | 1,53E-158 |
| 1110004F10I | 5,07E-162 | 0.253584570 | 0,49097222 | 0,33472222 | 1,64E-157 |
| Egr1 | 8,33E-162 | 0.298073810 | 0,64444444 | 0,05972222 | 2,69E-158 |
| Ndufa12 | 2,86E-161 | 0.250995269 | 0,48194444 | 0,31388889 | 9,23E-157 |
| Penk | 3,58E-162 | -0.29983337 | 0,11458333 | 0,28055556 | 1,16E-155 |
| Jdp2 | 4,03E-160 | -0.28562045 | 0,31111111 | 0,4 | 1,30E-155 |
| Igfbp4 | 3,04E-160 | 0.396368904 | 0,32569444 | 0,15625 | 9,82E-155 |
| Podn | 1,89E-158 | -0.26831159 | 0,33958333 | 0,44513889 | 6,12E-152 |
| Fos | 6,51E-156 | 0.348204560 | 0,64513889 | 0,58472222 | 2,10E-151 |
| Nceh1 | 1,85E-155 | -0.26868271 | 0,17222222 | 0,30833333 | 5,97E-151 |
| Dpp4 | 8,22E-155 | -0.27711484 | 0,14444444 | 0,30763889 | 2,65E-150 |
| Gnl2 | 1,70E-155 | 0.252023571 | 0,38125 | 0,20763889 | 5,50E-150 |
| Tubb5 | 4,62E-152 | 0.260221331 | 0,59375 | 0,50486111 | 1,49E-147 |
| Ier3 | 5,03E-152 | 0.251024337 | 0,6375 | 0,59930556 | 1,62E-147 |
| Akap12 | 9,03E-152 | -0.29860770 | 0,13611111 | 0,28611111 | 2,92E-147 |
| Tspan4 | 1,10E-151 | 0.251063809 | 0,51319444 | 00.57 | 3,56E-147 |
| Aldh1a1 | 1,02E-150 | -0.32195514 | 0,17430556 | 00.49 | 3,28E-146 |
| Cadm3 | 1,00E-149 | -0.30823230 | 0,29444444 | 0,04652778 | 3,24E-145 |
| Col23a1 | 1,02E-149 | 0.269593060 | 0,25347222 | 00.15 | 3,28E-145 |
| Rasd1 | 2,18E-149 | 0.285840367 | 0,21597222 | 0,07916667 | 7,04E-145 |
| Ccl7 | 1,63E-148 | -0.46524897 | 0,26180556 | 0,41875 | 5,26E-144 |
| Sectm1a | 3,16E-148 | -0.30144031 | 0,17638889 | 0,34861111 | 1,02E-143 |
| Gfpt2 | 4,26E-147 | -0.29808472 | 0,35972222 | 0,04513889 | 1,37E-142 |
| Clec11a | 4,01E-140 | -0.25555587 | 00.39 | 0,38541667 | 1,30E-135 |
| Ptges | 2,35E-139 | 0.265773964 | 0,31597222 | 0,17569444 | 7,60E-135 |
| Mfap4 | 4,02E-140 | -0.42688081 | 0,31388889 | 0,44791667 | 1,30E-134 |
| Fzd1 | 1,70E-138 | 0.264005037 | 0,37430556 | 0,21458333 | 5,50E-134 |
| Errfi1 | 7,85E-137 | -0.25372564 | 0,05902778 | 0,64375 | 2,53E-132 |
| Hexim1 | 1,20E-135 | 0.255395894 | 0,51597222 | 0,36736111 | 3,87E-131 |
| Fst | 1,29E-132 | 0.371906419 | 0,30694444 | 0,20416667 | 4,16E-128 |
| Srpx | 8,59E-132 | -0.27921754 | 0,36736111 | 0,49375 | 2,77E-127 |
| C1qtnf1 | 1,76E-131 | 0.262988754 | 0,41875 | 0,27916667 | 5,69E-127 |
| Spon1 | 5,61E-130 | 0.295306144 | 0,26736111 | 0,12361111 | 1,81E-125 |
| Csf1 | 3,81E-130 | -0.30435090 | 0,30416667 | 0,40902778 | 1,23E-124 |
| Gldn | 4,00E-128 | 0.343659556 | 0,44097222 | 0,28263889 | 1,29E-122 |
| Igfbp5 | 1,02E-125 | -0.35025891 | 0,43541667 | 0,55069444 | 3,28E-121 |
| Pdgfra | 1,28E-122 | 0.250153673 | 0,05277778 | 0,43194444 | 4,13E-118 |
| Nfib | 3,33E-122 | -0.27859802 | 0,18333333 | 0,31319444 | 1,08E-118 |
| Apod | 2,78E-120 | 0.318063662 | 0,56388889 | 0,52986111 | 8,96E-116 |
| C7 | 3,38E-119 | -0.26388141 | 0.071 | 0,15347222 | 1,09E-114 |
| Cyp26b1 | 1,66E-117 | 0.309723179 | 0,53194444 | 0,44027778 | 5,37E-113 |
| Cfb | 3,33E-116 | 0.306020986 | 0,26041667 | 00.21 | 1,07E-111 |
| Ctla2a | 4,44E-115 | -0.42527992 | 0,10763889 | 0,24027778 | 1,43E-110 |
| Ifi205 | 6,72E-109 | -0.27189881 | 0,11388889 | 0,21805556 | 2,17E-104 |
| Saa3 | 9,60E-109 | 0.437401067 | 0,12152778 | 0.054 | 3,10E-104 |
| Rarres2 | 4,15E-103 | -0.27550787 | 0,48333333 | 0,57222222 | 1,34E-98 |
| Slc5a3 | 5,32E-103 | 0.253589398 | 00.55 | 0,23541667 | 1,72E-98 |
| Ifi203 | 1,96E-101 | -0.27049462 | 0,16597222 | 0,27916667 | 6,32E-97 |
| Abi3bp | 1,54E-98 | -0.25800955 | 0,36736111 | 0,50555556 | 4,98E-94 |
| Hk2 | 7,08E-97 | -0.26541913 | 0,31736111 | 0,39236111 | 2,29E-92 |
| Inhba | 4,50E-94 | 0.267817882 | 0,26111111 | 0,13541667 | 1,45E-87 |
| Serpine1 | 8,91E-87 | 0.254734968 | 0,04444444 | 0,31597222 | 2,88E-82 |
| Lepr | 3,59E-84 | 0.254861200 | 0,23402778 | 00.18 | 1,16E-79 |
| Adamts4 | 1,62E-83 | -0.27777957 | 0,35763889 | 0,42013889 | 5,23E-79 |
| Enpp2 | 1,26E-76 | 0.257022890 | 0,46388889 | 0,04236111 | 4,07E-71 |
| Cxcl1 | 1,98E-73 | 0.351222190 | 00.47 | 0,20972222 | 6,38E-69 |
| Ccl8 | 1,79E-72 | -0.25367159 | 0,19305556 | 0,31458333 | 5,77E-68 |
| Lpl | 2,85E-72 | -0.29217238 | 0,20763889 | 0,31666667 | 9,20E-68 |
| Serpine2 | 4,58E-72 | 0.317037541 | 0,21388889 | 0,11527778 | 1,48E-67 |
| Wif1 | 7,50E-65 | 0.255164677 | 0,20625 | 0,10694444 | 2,42E-60 |
| Gbp2 | 5,00E-64 | -0.26277423 | 0,21805556 | 00.44 | 1,61E-58 |
| Postn | 1,39E-41 | 0.265798675 | 0,34444444 | 0,25555556 | 4,49E-37 |

**Supplementary Table 6 related to Supplementary Fig. 6d heatmap differentially expressed genes**

| Category | Ensembl ID | D1-CTRL_1 | D1-CTRL_2 | D1-CTRL_3 | D1-CTRL_4 | D2-CTRL_1 | D2-CTRL_2 | D2-CTRL_3 | D4-CTRL_1 | D4-CTRL_2 | D4-CTRL_3 | D4-CTRL_4 | D1-EXP_1 | D1-EXP_2 | D1-EXP_3 | D1-EXP_4 | D1-EXP_5 | D2-EXP_1 | D2-EXP_2 | D2-EXP_3 | D4-EXP_1 | D4-EXP_2 | D4-EXP_3 |
|---|---|---|---|---|---|---|---|---|---|---|---|---|---|---|---|---|---|---|---|---|---|---|---|
| Downregulated | ENSMUSG00000023011 | 2,114950705 | 47,869377 | 17,24719 | 48,1783712 | 13,9943375 | 9,28878453 | 7,95125211 | 1,22222602 | 7,4743178 | 4,31478604 | 0 | 3,503399 | 19,0333728 | 1,870957 | 0 | 1,823091581 | 8,272368 | 0 | 0 | 0 | 0 | 11,6657122 |
| Downregulated | ENSMUSG00000036040 | 415,5878135 | 727,614531 | 293,202229 | 1780,6726 | 122,1324 | 99,5226914 | 301,011687 | 587,890716 | 217,822976 | 256,729769 | 158,992316 | 46,42003 | 291,511131 | 260,063 | 110,7128 | 84,77375853 | 94,305 | 144,6643174 | 209,1246 | 39,99938 | 446,4069 | 371,508066 |
| Downregulated | ENSMUSG00000049112 | 131,1269437 | 110,631449 | 65,3577725 | 362,301351 | 123,404612 | 69,00239937 | 24,9896495 | 15,8889383 | 21,3551937 | 18,3378407 | 11,0411331 | 43,79248 | 14,0245905 | 33,67722 | 3,18598 | 4,557728953 | 11,58132 | 68,36875273 | 10,45623 | 0 | 7,971552 | 42,1760364 |
| Downregulated | ENSMUSG00000106634 | 19,03455634 | 31,912918 | 9,0774684 | 42,3969667 | 5,08884999 | 17,25059984 | 24,9896495 | 13,4444862 | 4,27103874 | 11,8656616 | 1,10411331 | 9,634347 | 9,01580818 | 3,741914 | 10,35443 | 10,93854949 | 3,308947 | 7,92681191 | 4,752832 | 0 | 1,771456 | 0 |
| Downregulated | ENSMUSG00000048814 | 16,91960564 | 22,3390426 | 9,0774684 | 50,105506 | 1,2722125 | 7,961815312 | 27,2614358 | 1,22222602 | 9,60983717 | 20,4952337 | 2,20822662 | 1,751699 | 0 | 0 | 0 | 8,203912116 | 0,827237 | 2,972554466 | 4,752832 | 3,999938 | 15,9431 | 2,69208743 |
| Downregulated | ENSMUSG00000031712 | 47,58639085 | 35,1042098 | 16,3394431 | 82,8667985 | 21,6276125 | 39,80907656 | 37,4844742 | 22,0000684 | 28,8295115 | 49,6200394 | 12,1452464 | 2,627549 | 8,01405172 | 4,677392 | 14,33691 | 0,911545791 | 0,827237 | 26,7529902 | 5,703399 | 7,333219 | 53,14368 | 34,9971366 |
| Downregulated | ENSMUSG00000099624 | 20,09203169 | 19,1477508 | 15,4316963 | 32,7612924 | 13,9943375 | 1,326969219 | 24,9896495 | 3,66667807 | 10,6775969 | 19,4165372 | 22,0822662 | 1,751699 | 3,00526939 | 4,677392 | 7,168455 | 5,469274744 | 8,272368 | 8,917663399 | 11,4068 | 9,333188 | 6,200096 | 21,5366995 |
| Downregulated | ENSMUSG00000086596 | 62,39104579 | 63,825836 | 62,634532 | 176,332839 | 15,26655 | 11,94272297 | 13,6307179 | 25,6667465 | 29,8972712 | 77,6661487 | 32,019286 | 19,26869 | 15,026347 | 36,48366 | 27,87732 | 51,95811007 | 31,435 | 11,89021787 | 8,555098 | 1,333313 | 4,42864 | 33,2024117 |
| Downregulated | ENSMUSG00000028031 | 1912,972912 | 2809,40055 | 1613,06613 | 4360,14259 | 1704,76475 | 1549,900047 | 2503,50852 | 1932,33934 | 1655,02751 | 3361,21832 | 1517,05169 | 1482,814 | 724,269924 | 1274,122 | 1698,127 | 2061,916578 | 1719,825 | 1969,81276 | 1135,927 | 207,9968 | 489,8076 | 1334,378 |
| Downregulated | ENSMUSG00000059659 | 37,01163733 | 54,2519606 | 19,9704305 | 31,797725 | 20,3554 | 27,86635359 | 6,81535895 | 17,1111643 | 18,1519147 | 25,8887162 | 7,72879316 | 4,379248 | 13,022834 | 4,677392 | 10,35443 | 26,43482793 | 19,02645 | 14,86277233 | 14,2585 | 11,99981 | 22,1432 | 9,87098725 |
| Downregulated | ENSMUSG00000020491 | 78,25317607 | 90,4199344 | 45,387342 | 57,8140454 | 30,5331 | 59,71361484 | 17,0383974 | 20,7778424 | 22,4229534 | 60,4070045 | 15,4575863 | 15,76529 | 21,0368858 | 19,64505 | 18,31938 | 10,93854949 | 15,7175 | 22,78958424 | 24,71473 | 41,33269 | 24,80038 | 32,3050492 |
| Downregulated | ENSMUSG00000000056 | 985,5670284 | 1336,0875 | 788,832004 | 691,84141 | 739,155462 | 456,4774112 | 673,584643 | 482,779279 | 405,748681 | 714,097089 | 550,952541 | 378,3671 | 184,323189 | 391,03 | 331,3419 | 338,1834883 | 383,8379 | 321,0358824 | 516,1576 | 483,9924 | 727,1827 | 618,282747 |
| Downregulated | ENSMUSG00000068373 | 39,12658804 | 89,3561705 | 29,9556457 | 26,0163204 | 15,26655 | 0 | 11,3589316 | 11,0000342 | 10,6775969 | 26,9674127 | 20,9781529 | 0,87585 | 6,01053879 | 0 | 19,91237 | 13,67318686 | 0 | 26,7529902 | 4,752832 | 4,666594 | 8,85728 | 8,0762623 |
| Downregulated | ENSMUSG00000059824 | 64,50599649 | 131,906728 | 18,1549368 | 33,7248598 | 26,7164625 | 13,26969219 | 24,9896495 | 7,33335613 | 9,60983717 | 37,7543778 | 36,4357392 | 14,88944 | 3,00526939 | 16,83861 | 10,35443 | 30,08101109 | 30,60776 | 12,88106935 | 13,30793 | 12,66647 | 12,40019 | 16,1525246 |
| Downregulated | ENSMUSG00000026188 | 6,344852114 | 37,2317377 | 33,5866331 | 8,67210682 | 11,4499125 | 13,26969219 | 20,4460768 | 19,5556163 | 3,20327906 | 19,4165372 | 18,7699262 | 2,627549 | 0 | 1,870957 | 10,35443 | 1,823091581 | 2,481711 | 1,981702978 | 2,851699 | 5,999906 | 0 | 3,58944991 |
| Downregulated | ENSMUSG00000081096 | 9,517278171 | 40,4230295 | 35,4021268 | 27,9434553 | 15,26655 | 14,5966614 | 5,67946579 | 26,8889725 | 8,54207749 | 17,2591441 | 14,353473 | 0,87585 | 1,00175646 | 0 | 0 | 9,115457906 | 0 | 1,981702978 | 0,950566 | 0 | 0 | 8,0762623 |
| Downregulated | ENSMUSG00000033282 | 95,17278171 | 309,555305 | 212,412761 | 135,863007 | 97,9603624 | 237,5274901 | 135,171286 | 232,222944 | 137,740999 | 114,34183 | 213,093869 | 40,28909 | 32,0562069 | 27,12887 | 32,65629 | 40,10801479 | 41,36184 | 92,14918846 | 91,25438 | 107,3317 | 102,7444 | 87,9415228 |
| Downregulated | ENSMUSG00000026489 | 176,5983838 | 229,77301 | 173,379646 | 83,8303659 | 115,771337 | 141,9857064 | 69,2894827 | 157,667157 | 74,743178 | 62,5643975 | 143,53473 | 36,78569 | 69,1211961 | 47,7094 | 62,9231 | 30,08101109 | 38,05289 | 65,39619826 | 47,52832 | 106,665 | 70,85824 | 116,657122 |
| Downregulated | ENSMUSG00000059713 | 325,7024085 | 448,90838 | 445,703698 | 243,782558 | 211,187275 | 240,1814286 | 251,032388 | 281,111985 | 152,689635 | 350,576365 | 290,3818 | 171,6665 | 98,1721335 | 96,35427 | 133,0147 | 106,6508575 | 160,4839 | 244,7403177 | 286,1205 | 415,3269 | 251,5468 | 194,727658 |
| Downregulated | ENSMUSG00000025218 | 131,1269437 | 184,031161 | 163,394431 | 115,628091 | 129,765675 | 126,0620758 | 92,0073458 | 113,66702 | 39,5071084 | 115,420526 | 79,4961582 | 42,91663 | 54,0948491 | 25,25792 | 58,14413 | 41,93110637 | 77,76026 | 85,21322804 | 107,414 | 129,3313 | 103,6302 | 41,278674 |
| Downregulated | ENSMUSG00000073792 | 167,0811057 | 291,471318 | 244,1839 | 170,551434 | 270,981262 | 192,4105367 | 214,683807 | 240,778526 | 215,687457 | 255,651073 | 216,406209 | 88,46082 | 90,1580818 | 38,35461 | 47,7897 | 79,30448379 | 124,0855 | 138,7192084 | 158,7446 | 249,9961 | 211,689 | 170,498871 |
| Downregulated | ENSMUSG00000041912 | 16,91960564 | 39,3592656 | 23,6014178 | 18,3077811 | 27,988675 | 18,57756906 | 21,58197 | 19,5556163 | 14,9486356 | 23,7313232 | 11,0411331 | 3,503399 | 1,00175646 | 0 | 1,59299 | 3,646183163 | 7,445132 | 9,908514888 | 2,851699 | 27,33291 | 10,62874 | 16,1525246 |
| Downregulated | ENSMUSG00000116903 | 115,2648134 | 204,242675 | 106,20638 | 131,04517 | 83,9660249 | 49,09786109 | 118,132888 | 140,555993 | 105,708209 | 122,971402 | 102,682538 | 66,56458 | 41,072015 | 18,70957 | 29,47031 | 61,07356797 | 63,69724 | 164,4813471 | 126,4253 | 25,33294 | 31,88621 | 59,2259235 |
| Downregulated | ENSMUSG00000042453 | 33,83921127 | 95,7387541 | 44,4795952 | 97,3203098 | 11,4499125 | 95,54178374 | 30,6691153 | 63,5557531 | 50,1847052 | 48,5413429 | 45,2686457 | 52,55098 | 14,0245905 | 10,29026 | 26,28433 | 16,40782423 | 42,18908 | 63,41449528 | 35,17096 | 0 | 16,82883 | 17,0498871 |
| Downregulated | ENSMUSG00000071633 | 11,63222888 | 55,3157246 | 7,26197472 | 52,9962083 | 29,2608875 | 19,90453828 | 26,1255426 | 35,4445546 | 18,1519147 | 22,6526267 | 9,93701978 | 8,758497 | 9,01580818 | 0 | 2,389485 | 0,911545791 | 4,136184 | 13,87192084 | 0 | 13,33313 | 10,62874 | 25,1261494 |
| Downregulated | ENSMUSG00000026303 | 50,75881691 | 144,671895 | 62,634532 | 96,3567424 | 81,4215999 | 33,17423047 | 78,3766279 | 85,5558215 | 24,5584728 | 87,3744172 | 98,2660845 | 25,39964 | 45,0790409 | 17,77409 | 25,48784 | 37,37337742 | 24,81711 | 60,44194082 | 41,82493 | 15,33309 | 15,9431 | 78,967898 |
| Downregulated | ENSMUSG00000072541 | 42,29901409 | 82,9735868 | 52,6493167 | 50,105506 | 47,0718625 | 31,84726125 | 61,3382305 | 40,3334587 | 58,7267827 | 76,5874522 | 28,706946 | 17,51699 | 16,0281034 | 14,03218 | 17,52289 | 28,25791951 | 11,58132 | 38,64320806 | 15,20906 | 7,333219 | 23,02893 | 29,6129618 |
| Downregulated | ENSMUSG00000029211 | 49,70134156 | 164,88341 | 74,4352409 | 81,903231 | 80,1493874 | 115,446322 | 156,753256 | 92,8891777 | 72,6076586 | 140,230546 | 94,9537446 | 3,503399 | 14,0245905 | 28,06435 | 9,557939 | 51,95811007 | 10,75408 | 93,14003995 | 36,12153 | 0 | 0 | 42,1760364 |
| Downregulated | ENSMUSG00000029869 | 1095,544465 | 1181,84173 | 925,901777 | 1148,57237 | 1170,4355 | 1293,794988 | 1106,35994 | 1142,78133 | 839,259113 | 1339,74106 | 806,002715 | 278,5202 | 231,405743 | 173,999 | 209,4782 | 332,7142136 | 328,413 | 540,0140614 | 473,3821 | 197,3303 | 143,4879 | 499,8309 |
| Downregulated | ENSMUSG00000021835 | 1738,489479 | 2418,99919 | 1409,73084 | 1972,42252 | 1866,33574 | 2192,153149 | 1937,83373 | 1569,33821 | 1354,98704 | 1632,06782 | 1266,41797 | 541,2751 | 268,470733 | 318,9981 | 438,8687 | 508,6425512 | 322,6224 | 1035,439806 | 408,7436 | 59,99906 | 225,8606 | 732,247782 |
| Downregulated | ENSMUSG00000031431 | 7410,787269 | 8454,79575 | 4538,7342 | 5161,83069 | 6484,46711 | 5192,430552 | 6840,3486 | 6672,13185 | 4578,55353 | 7121,55435 | 5722,61928 | 3832,718 | 2066,62359 | 2126,342 | 2797,29 | 2233,287187 | 2226,922 | 4302,277164 | 1716,723 | 638,6567 | 1159,418 | 4198,75903 |
| Downregulated | ENSMUSG00000021388 | 14367,91761 | 23941,0711 | 20053,9432 | 28717,1999 | 25879,3466 | 16593,75008 | 22302,1263 | 32483,101 | 22479,5447 | 29471,0673 | 26365,1217 | 7492,018 | 10241,9581 | 7169,506 | 3351,651 | 4702,664734 | 4243,725 | 17962,15579 | 8897,302 | 3467,946 | 10863,45 | 20402,4333 |
| Downregulated | ENSMUSG00000058914 | 19475,52356 | 34678,7042 | 18857,5329 | 47036,5438 | 37156,2382 | 26431,89987 | 42271,128 | 31614,0983 | 28385,3235 | 40290,3933 | 31811,7127 | 6022,342 | 14174,854 | 7214,409 | 5434,485 | 7659,719279 | 9772,976 | 24278,83403 | 10870,68 | 1122,649 | 6856,421 | 30661,9785 |
| Downregulated | ENSMUSG00000030255 | 1250,993342 | 1513,73608 | 1206,39555 | 1795,12611 | 1298,92896 | 1442,415541 | 1561,85309 | 1818,67232 | 1241,80451 | 1822,9971 | 811,523282 | 414,2769 | 465,816756 | 232,9341 | 295,4996 | 302,6332025 | 515,3686 | 1099,845153 | 584,5984 | 565,9912 | 519,0366 | 928,770164 |
| Downregulated | ENSMUSG00000045658 | 2792,792405 | 2996,623 | 2803,12224 | 3288,65562 | 2310,3379 | 2742,845375 | 3274,77998 | 3308,56584 | 2177,162 | 3594,21677 | 2671,95421 | 685,7903 | 711,24709 | 577,1902 | 610,1151 | 495,8809101 | 624,5638 | 1037,421509 | 945,8137 | 373,3275 | 1085,017 | 1696,01508 |
| Downregulated | ENSMUSG00000053477 | 5826,689191 | 7086,79533 | 5933,03335 | 8317,514 | 6943,73582 | 6744,984538 | 7819,4885 | 7114,57767 | 6054,19742 | 9834,47607 | 6480,04101 | 3234,513 | 2122,72195 | 3167,53 | 2990,839 | 2959,789182 | 3463,641 | 4700,599463 | 3297,515 | 1632,641 | 2200,148 | 3847,8903 |
| Downregulated | ENSMUSG00000024084 | 5547,515698 | 14032,1101 | 8797,88237 | 11441,3996 | 16219,4371 | 11486,24556 | 17140,6278 | 13522,7087 | 17600,9507 | 12738,3271 | 13709,775 | 3711,851 | 4747,32389 | 4255,491 | 3768,218 | 5181,226274 | 2086,291 | 8208,213733 | 4171,086 | 459,3262 | 3942,375 | 8680,18724 |
| Downregulated | ENSMUSG00000086283 | 59,21861973 | 149,990715 | 107,114127 | 135,863007 | 131,037887 | 136,6778295 | 153,345576 | 118,555924 | 142,012038 | 125,128795 | 109,307218 | 37,66154 | 47,0825538 | 44,90296 | 33,45279 | 44,66574374 | 16,54474 | 60,44194082 | 31,36869 | 21,333 | 57,57232 | 107,683497 |
| Downregulated | ENSMUSG00000020102 | 204,092743 | 455,290964 | 237,829672 | 238,001154 | 347,314012 | 258,7589976 | 321,457764 | 453,445854 | 429,239394 | 373,228992 | 325,713426 | 67,44043 | 221,388179 | 45,83844 | 81,24248 | 106,6508575 | 77,76026 | 176,371565 | 144,4861 | 29,33288 | 96,54435 | 253,056219 |
| Downregulated | ENSMUSG00000057766 | 872,4171657 | 1381,82935 | 1011,22998 | 908,644081 | 1452,86667 | 1076,172036 | 1266,52087 | 1431,22667 | 1578,14882 | 1460,55507 | 1137,23671 | 245,2379 | 326,572607 | 272,2242 | 156,9095 | 249,7635466 | 103,4046 | 504,3434078 | 270,9115 | 141,3311 | 329,4908 | 664,048233 |
| Downregulated | ENSMUSG00000032849 | 2449,112916 | 3491,27323 | 1942,57824 | 2296,18117 | 4125,78513 | 3253,728524 | 3277,05176 | 4325,45789 | 3359,17197 | 3740,91949 | 3060,60209 | 1188,528 | 1131,9848 | 944,8332 | 801,2739 | 929,7767064 | 686,6066 | 1461,505946 | 1590,298 | 543,3249 | 1027,444 | 2445,31275 |
| Downregulated | ENSMUSG00000028565 | 3631,37036 | 4950,75735 | 4588,66028 | 5428,73887 | 4952,72326 | 5058,406661 | 4805,96395 | 6370,24203 | 6125,73732 | 5731,11455 | 4926,55358 | 2740,534 | 2548,46845 | 1560,378 | 1772,998 | 1618,905324 | 871,0804 | 2346,336326 | 1617,864 | 1761,973 | 1988,459 | 2239,81674 |
| Downregulated | ENSMUSG00000025666 | 3456,886927 | 4708,21917 | 3997,71708 | 3566,16304 | 3845,89838 | 3744,707135 | 4010,83874 | 5525,68384 | 4986,43773 | 5335,23293 | 3580,63946 | 2246,554 | 1812,17744 | 1333,992 | 1749,899 | 2380,046059 | 1678,464 | 2328,500999 | 1653,035 | 788,6544 | 721,8683 | 2250,58509 |
| Downregulated | ENSMUSG00000018899 | 6172,483631 | 7434,64613 | 5684,31071 | 7823,20392 | 8799,89385 | 7768,077806 | 8313,60203 | 9165,47294 | 9700,59674 | 9177,5499 | 7156,86247 | 4007,888 | 2422,24713 | 2264,793 | 3581,041 | 3412,82744 | 3563,736 | 4244,807778 | 3697,704 | 2029,968 | 1722,741 | 4844,86002 |
| Downregulated | ENSMUSG00000068854 | 13,74717958 | 15,956459 | 26,3246584 | 14,4535114 | 22,899825 | 7,961815312 | 19,3101837 | 17,1111643 | 10,6775969 | 15,1017511 | 13,2493597 | 4,379248 | 0 | 2,806435 | 5,575465 | 0,911545791 | 14,06303 | 13,87192084 | 0,950566 | 3,333281 | 23,02893 | 8,0762623 |
| Downregulated | ENSMUSG00000020435 | 44,4139648 | 39,3592656 | 95,3134182 | 57,8140454 | 109,410275 | 57,0596764 | 92,0073458 | 85,5558215 | 40,5748681 | 79,8235417 | 27,6028327 | 23,64794 | 23,0403987 | 16,83861 | 14,33691 | 12,76164107 | 23,16263 | 25,76213871 | 42,77549 | 28,66622 | 98,31581 | 69,9942732 |
| Downregulated | ENSMUSG00000054545 | 27,49435916 | 32,976682 | 28,140152 | 25,052753 | 20,3554 | 15,92363062 | 30,6691153 | 30,5556505 | 28,8295115 | 25,8887162 | 30,9151726 | 14,0136 | 9,01580818 | 1,870957 | 7,964949 | 8,203912116 | 10,75408 | 11,89021787 | 18,06076 | 13,33313 | 28,3433 | 10,7683497 |
| Downregulated | ENSMUSG00000097616 | 40,18406339 | 46,8056131 | 67,1732662 | 31,797725 | 41,9830125 | 3,980907656 | 37,4844742 | 39,1112327 | 38,4393487 | 37,7543778 | 33,1233993 | 14,0136 | 13,022834 | 17,77409 | 11,94742 | 8,203912116 | 9,099605 | 14,86277233 | 30,41813 | 10,6665 | 31,88621 | 9,87098725 |
| Downregulated | ENSMUSG00000045333 | 390,208405 | 265,940983 | 296,833217 | 211,984833 | 236,631525 | 96,86875296 | 269,206679 | 166,222739 | 248,788007 | 201,716247 | 91,6414046 | 43,79248 | 111,194968 | 49,58035 | 72,48104 | 69,27748009 | 38,88013 | 102,0577033 | 76,99589 | 274,6624 | 426,0352 | 306,897967 |
| Downregulated | ENSMUSG00000079494 | 105,7475352 | 35,1042098 | 70,8042535 | 49,1419386 | 40,7108 | 37,15513812 | 61,3382305 | 56,222397 | 100,36941 | 77,6661487 | 50,7892122 | 8,758497 | 39,0685021 | 15,90313 | 14,33691 | 10,93854949 | 11,58132 | 56,47853486 | 45,62719 | 49,33256 | 113,3732 | 69,9942732 |
| Downregulated | ENSMUSG00000041216 | 39,12658804 | 5,31881967 | 29,9556457 | 9,63567424 | 6,36106249 | 10,61575375 | 21,58197 | 12,2222602 | 22,4229534 | 8,62957207 | 7,72879316 | 6,130948 | 3,00526939 | 0,935478 | 1,59299 | 2,734637372 | 15,7175 | 7,92681191 | 3,802266 | 1,999969 | 27,45757 | 13,4604372 |
| Downregulated | ENSMUSG00000021662 | 469,5190564 | 366,998557 | 417,563546 | 158,025058 | 110,682487 | 47,77089187 | 450,949584 | 67,2224312 | 256,262325 | 351,655062 | 240,696701 | 85,83327 | 136,238879 | 14,03218 | 16,72639 | 25,52328214 | 118,2949 | 50,53342593 | 195,8167 | 99,99844 | 350,7483 | 204,598645 |
| Downregulated | ENSMUSG00000022679 | 51,81629226 | 29,7853902 | 50,833823 | 33,7248598 | 40,7108 | 14,5966614 | 38,6203674 | 67,2224312 | 20,287434 | 30,2035023 | 12,1452464 | 4,379248 | 12,0210776 | 3,741914 | 7,168455 | 10,0270037 | 36,39842 | 43,59746551 | 12,35736 | 51,99919 | 76,17261 | 25,1261494 |
| Downregulated | ENSMUSG00000014633 | 96,23025706 | 136,161784 | 177,010634 | 158,025058 | 99,2325749 | 90,23390687 | 130,627713 | 129,555958 | 77,9464571 | 102,476168 | 58,5180054 | 34,15814 | 55,0966056 | 30,87079 | 85,22496 | 29,1694653 | 90,16882 | 106,0211093 | 139,7333 | 155,3309 | 196,6316 | 97,81251 |
| Downregulated | ENSMUSG00000039883 | 2103,318476 | 2599,83905 | 2684,20741 | 4315,81849 | 1606,80439 | 1694,539692 | 1826,5162 | 2484,7855 | 1224,72036 | 2108,85168 | 1054,42821 | 99,84686 | 263,46195 | 112,2574 | 122,6602 | 106,6508575 | 893,4158 | 1458,533392 | 1234,786 | 3689,276 | 4492,412 | 1796,51968 |
| Downregulated | ENSMUSG00000079559 | 29,60930986 | 24,4665705 | 52,6493167 | 43,3605341 | 10,1777 | 17,25059984 | 23,8537563 | 31,7778766 | 16,0163953 | 17,2591441 | 29,8110593 | 2,627549 | 13,022834 | 5,61287 | 7,168455 | 10,0270037 | 19,85368 | 8,917663399 | 12,35736 | 21,99966 | 39,85776 | 22,4340619 |
| Downregulated | ENSMUSG00000113585 | 30,66678522 | 9,57387541 | 32,6788862 | 34,6884273 | 7,63327499 | 5,307876874 | 19,3101837 | 20,7778424 | 9,60983717 | 14,0230546 | 11,0411331 | 0,87585 | 6,01053879 | 2,806435 | 8,761444 | 3,646183163 | 8,272368 | 13,87192084 | 6,653965 | 17,99972 | 16,82883 | 1,79472495 |
| Downregulated | ENSMUSG00000028268 | 829,0606762 | 763,782505 | 1221,82725 | 290,033795 | 552,140224 | 810,7781926 | 516,831387 | 393,556779 | 461,272184 | 1136,94612 | 932,975746 | 310,0508 | 157,275765 | 38,35461 | 371,1666 | 303,5447483 | 606,3646 | 431,0203976 | 403,0402 | 90,66525 | 425,1494 | 351,766091 |
| Downregulated | ENSMUSG00000017830 | 157,5638275 | 158,500826 | 301,371951 | 86,7210682 | 40,7108 | 98,19572218 | 93,143239 | 37,8890067 | 52,3202246 | 170,434048 | 158,992316 | 33,28229 | 25,0439116 | 11,22574 | 122,6602 | 80,21602958 | 187,7828 | 58,46023784 | 61,78682 | 52,66584 | 152,3452 | 61,9180109 |
| Downregulated | ENSMUSG00000067297 | 58,16114438 | 61,6983082 | 60,8190383 | 37,5791295 | 15,26655 | 39,80907656 | 29,5332221 | 13,4444862 | 1,06775969 | 110,027044 | 12,1452464 | 10,5102 | 6,01053879 | 6,548349 | 27,08083 | 28,25791951 | 26,47158 | 22,78958424 | 21,86303 | 1,999969 | 38,0863 | 11,6657122 |
| Downregulated | ENSMUSG00000032816 | 258,023986 | 138,289311 | 256,892356 | 78,0489613 | 76,3327499 | 124,7351066 | 68,1535895 | 136,889314 | 102,50493 | 240,549322 | 262,778967 | 56,05438 | 62,1089008 | 39,29009 | 15,1334 | 11,85009528 | 17,37197 | 88,1857825 | 38,97323 | 149,9977 | 174,4884 | 115,75976 |
| Downregulated | ENSMUSG00000033491 | 101,5176338 | 24,4665705 | 159,763444 | 48,1783712 | 7,63327499 | 0 | 5,67946579 | 0 | 14,9486356 | 255,651073 | 97,1619712 | 0 | 4,00702586 | 0 | 0,796495 | 0 | 0 | 25,76213871 | 15,20906 | 28,66622 | 47,82931 | 96,9151476 |
| Downregulated | ENSMUSG00000056174 | 108,9199613 | 76,5910032 | 85,328203 | 85,7575007 | 54,7051374 | 38,48210734 | 51,1151921 | 42,7779108 | 44,8459068 | 235,155839 | 104,890764 | 9,634347 | 13,022834 | 14,03218 | 9,557939 | 18,23091581 | 7,445132 | 21,79873275 | 3,802266 | 31,9995 | 106,2874 | 76,2758106 |
| Downregulated | ENSMUSG00000052544 | 10,57475352 | 14,8926951 | 26,3246584 | 19,2713485 | 20,3554 | 13,26969219 | 9,08714527 | 7,33335613 | 14,9486356 | 35,5969848 | 4,41645324 | 7,882647 | 3,00526939 | 6,548349 | 0 | 0,911545791 | 19,02645 | 10,89936638 | 1,901133 | 5,999906 | 4,42864 | 17,0498871 |
| Downregulated | ENSMUSG00000046807 | 4,229901409 | 8,51011147 | 31,7711394 | 2,89070227 | 6,36106249 | 5,307876874 | 6,81535895 | 6,11113011 | 4,27103874 | 24,8100197 | 2,20822662 | 0,87585 | 1,00175646 | 0 | 0 | 1,823091581 | 2,481711 | 0,990851489 | 4,752832 | 9,333188 | 0 | 18,844612 |
| Downregulated | ENSMUSG00000021904 | 26,43688381 | 5,31881967 | 48,1105825 | 26,9798879 | 16,5387625 | 5,307876874 | 23,8537563 | 23,2222944 | 26,6939921 | 33,4395918 | 7,72879316 | 7,882647 | 4,00702586 | 0 | 10,35443 | 5,469274744 | 4,963421 | 26,7529902 | 10,45623 | 7,999875 | 2,657184 | 9,87098725 |
| Downregulated | ENSMUSG00000045613 | 6,344852114 | 7,44634754 | 21,7859242 | 12,5263765 | 5,08884999 | 2,653938437 | 4,54357263 | 15,8889383 | 10,6775969 | 22,6526267 | 4,41645324 | 1,751699 | 0 | 0 | 5,575465 | 1,823091581 | 0 | 13,87192084 | 0 | 17,33306 | 2,657184 | 5,38417486 |
| Downregulated | ENSMUSG00000024245 | 56,04619367 | 24,4665705 | 42,6641015 | 21,1984833 | 12,722125 | 17,25059984 | 2,27178632 | 25,6667465 | 2,13551937 | 33,4395918 | 22,0822662 | 0,87585 | 10,0175646 | 9,354784 | 1,59299 | 0 | 9,926842 | 7,92681191 | 2,851699 | 0 | 3,542912 | 8,97362477 |
| Downregulated | ENSMUSG00000095105 | 25,37940846 | 7,44634754 | 15,4316963 | 16,3806462 | 7,63327499 | 2,653938437 | 0 | 9,77780817 | 4,27103874 | 14,0230546 | 6,62467985 | 0 | 0 | 0 | 0,796495 | 0 | 1,654474 | 0 | 0 | 0 | 5,314368 | 1,79472495 |
| Downregulated | ENSMUSG00000022340 | 164,966155 | 93,6112262 | 144,331748 | 49,1419386 | 13,9943375 | 50,42483031 | 13,6307179 | 45,2223628 | 24,5584728 | 96,0039893 | 20,9781529 | 46,42003 | 24,0421551 | 53,32227 | 18,31938 | 38,28492321 | 22,33539 | 38,64320806 | 29,46756 | 1,999969 | 9,743008 | 27,8182368 |
| Downregulated | ENSMUSG00000028551 | 352,1392923 | 285,088734 | 406,670584 | 224,51121 | 152,6655 | 168,5250908 | 132,8995 | 190,667259 | 174,044829 | 213,581909 | 140,22239 | 78,82647 | 96,1686206 | 61,74157 | 42,21423 | 46,48883532 | 39,70737 | 95,12174293 | 129,277 | 87,99863 | 140,8308 | 184,85667 |
| Downregulated | ENSMUSG00000075273 | 201,9777923 | 117,014033 | 138,885267 | 102,138147 | 40,7108 | 80,94512234 | 68,1535895 | 59,8890751 | 9,60983717 | 64,7217906 | 23,1863795 | 15,76529 | 26,0456681 | 3,741914 | 17,52289 | 20,05400739 | 41,36184 | 20,80788127 | 25,6653 | 31,9995 | 43,40067 | 43,9707614 |
| Downregulated | ENSMUSG00000053835 | 689,4739297 | 540,392078 | 646,31575 | 156,097923 | 124,676825 | 257,4320284 | 249,896495 | 158,889383 | 113,182527 | 270,752824 | 125,868917 | 19,26869 | 78,1370042 | 6,548349 | 35,84227 | 71,10057167 | 58,73382 | 63,41449528 | 95,05665 | 23,33297 | 67,31533 | 117,554485 |
| Downregulated | ENSMUSG00000026222 | 794,1639896 | 407,421587 | 691,703092 | 343,030003 | 143,760012 | 323,7804893 | 163,568615 | 300,667601 | 317,124627 | 193,086675 | 144,638843 | 78,82647 | 191,335485 | 67,35444 | 270,0118 | 350,9451294 | 129,8762 | 102,0577033 | 228,136 | 229,9964 | 478,2931 | 115,75976 |
| Downregulated | ENSMUSG00000031767 | 114,2073381 | 45,7418492 | 100,759899 | 33,7248598 | 22,899825 | 29,19332281 | 37,4844742 | 18,3333903 | 20,287434 | 47,4626464 | 37,5398525 | 6,130948 | 27,0474245 | 17,77409 | 23,09835 | 33,72719425 | 33,91671 | 8,917663399 | 41,82493 | 63,999 | 55,80086 | 39,483949 |
| Downregulated | ENSMUSG00000001240 | 1559,776145 | 769,101324 | 1606,71191 | 607,047477 | 282,431175 | 444,5346882 | 216,955593 | 447,334724 | 498,643773 | 775,58279 | 705,528404 | 228,5968 | 403,707855 | 250,7082 | 189,5658 | 179,5745208 | 172,8925 | 216,0056246 | 272,8126 | 312,6618 | 820,1841 | 359,842353 |
| Downregulated | ENSMUSG00000035775 | 65,56347184 | 24,4665705 | 81,6972156 | 15,4170788 | 2,544425 | 0 | 5,67946579 | 25,6667465 | 3,20327906 | 8,62957207 | 17,6658129 | 0 | 1,00175646 | 0 | 0 | 0 | 0 | 0 | 7,604532 | 2,666625 | 29,22902 | 6,28153734 |
| Downregulated | ENSMUSG00000053399 | 158,6213028 | 3,1912918 | 206,96628 | 0,96356742 | 0 | 0 | 4,54357263 | 9,77780817 | 0 | 1,07869651 | 0 | 0 | 1,00175646 | 0 | 0 | 0 | 0 | 7,92681191 | 16,15963 | 0 | 24,80038 | 23,3314244 |
| Downregulated | ENSMUSG00000026259 | 134,2993697 | 60,6345442 | 125,269064 | 39,5062644 | 58,5217749 | 2,653938437 | 26,1255426 | 36,6667807 | 19,2196743 | 29,1248057 | 16,5616996 | 14,0136 | 66,1159267 | 8,419306 | 28,67382 | 24,61173635 | 4,963421 | 19,81702978 | 24,71473 | 13,33313 | 69,97251 | 41,278674 |
| Downregulated | ENSMUSG00000047246 | 101,5176338 | 62,7620721 | 91,6824308 | 39,5062644 | 29,2608875 | 3,980907656 | 32,9409016 | 13,4444862 | 14,9486356 | 20,4952337 | 12,1452464 | 14,88944 | 18,0316164 | 15,90313 | 6,37196 | 5,469274744 | 19,85368 | 4,954257444 | 19,01133 | 15,99975 | 38,97203 | 23,3314244 |
| Downregulated | ENSMUSG00000018102 | 2233,387944 | 1504,1622 | 2142,28254 | 938,514671 | 589,034387 | 272,0286898 | 814,435394 | 242,000752 | 158,028433 | 462,760802 | 464,831703 | 367,8569 | 359,630571 | 468,6747 | 100,3584 | 148,5819639 | 372,2566 | 204,1154067 | 697,7158 | 371,3275 | 1478,28 | 957,485763 |
| Downregulated | ENSMUSG00000068246 | 68,7358979 | 42,5505574 | 74,4352409 | 18,3077811 | 17,810975 | 37,15513812 | 15,9025042 | 4,88890409 | 3,20327906 | 49,6200394 | 19,8740396 | 0,87585 | 5,00878232 | 2,806435 | 23,89485 | 10,0270037 | 35,57118 | 1,981702978 | 31,36869 | 38,66606 | 31,88621 | 15,2551621 |
| Downregulated | ENSMUSG00000057346 | 57,10366903 | 57,4432524 | 75,3429877 | 21,1984833 | 1,2722125 | 17,25059984 | 17,0383974 | 2,44445204 | 1,06775969 | 44,2265569 | 11,0411331 | 4,379248 | 4,00702586 | 3,741914 | 33,45279 | 6,380820534 | 33,08947 | 5,945108933 | 11,4068 | 17,99972 | 18,60029 | 6,28153734 |
| Downregulated | ENSMUSG00000042408 | 322,5299825 | 231,900538 | 255,076862 | 97,3203098 | 110,682487 | 124,7351066 | 195,373623 | 113,66702 | 130,266682 | 168,276655 | 101,578424 | 96,34347 | 59,1036314 | 32,74174 | 126,6427 | 41,93110637 | 86,85987 | 105,0302578 | 117,8702 | 142,6644 | 65,54387 | 107,683497 |

| | | | | | | | | | | | | | | | | | | | | | | | |
|---|---|---|---|---|---|---|---|---|---|---|---|---|---|---|---|---|---|---|---|---|---|---|---|
| Downregulated | ENSMUSG00000073096 | 69,79337325 | 48,933141 | 37,2176204 | 27,9434553 | 24,1720375 | 23,88544594 | 32,9409016 | 15,8889383 | 16,0163953 | 37,7543778 | 35,3316259 | 14,88944 | 16,0281034 | 6,548349 | 29,47031 | 6,380820534 | 42,18908 | 21,79873275 | 27,56643 | 27,99956 | 22,1432 | 21,5366995 |
| Downregulated | ENSMUSG00000009092 | 99,40268312 | 23,4028065 | 69,8965067 | 21,1984833 | 12,722125 | 13,26969219 | 15,9025042 | 0 | 3,20327906 | 1,07869651 | 2,20822662 | 23,64794 | 29,0509375 | 2,806435 | 15,9299 | 10,93854949 | 28,12605 | 7,92681191 | 8,555098 | 5,33325 | 3,542912 | 8,97362477 |
| Downregulated | ENSMUSG00000069755 | 119,4947148 | 113,822741 | 145,239494 | 68,4132871 | 30,5331 | 22,55847672 | 20,4460768 | 2,44445204 | 0 | 44,2265569 | 34,2275126 | 33,28229 | 58,1018749 | 24,32244 | 48,58619 | 30,08101109 | 42,18908 | 22,78958424 | 76,99589 | 19,33303 | 10,62874 | 21,5366995 |
| Downregulated | ENSMUSG00000044456 | 72,96579931 | 44,6780852 | 37,2176204 | 41,4333992 | 13,9943375 | 27,86635359 | 15,9025042 | 20,7778424 | 7,4743178 | 19,4165372 | 35,3316259 | 12,2619 | 5,00878232 | 11,22574 | 2,389485 | 21,87709898 | 0 | 10,89936638 | 3,802266 | 23,99963 | 46,05786 | 16,1525246 |
| Downregulated | ENSMUSG00000029428 | 2069,479264 | 1661,59926 | 1743,78168 | 1225,65776 | 1216,23515 | 1021,766298 | 984,819368 | 1113,44791 | 766,651454 | 1643,93348 | 1341,49767 | 695,4247 | 641,124137 | 425,6427 | 452,4091 | 565,1583902 | 597,265 | 852,1322804 | 846,9547 | 941,9853 | 1670,483 | 1060,68245 |
| Downregulated | ENSMUSG00000030064 | 268,5987395 | 235,091829 | 162,486684 | 112,737389 | 66,1550499 | 167,1981215 | 94,2791321 | 154,000479 | 104,640449 | 145,624029 | 140,22239 | 74,44722 | 45,0790409 | 20,58052 | 35,84227 | 39,196469 | 13,23579 | 62,4236438 | 109,3151 | 65,99897 | 209,0318 | 160,627883 |
| Downregulated | ENSMUSG00000058385 | 124,7820916 | 85,1011147 | 64,4500256 | 37,5791295 | 39,4385875 | 0 | 42,0280469 | 9,77780817 | 0 | 25,8887162 | 24,2904928 | 23,64794 | 5,00878232 | 28,06435 | 0,796495 | 13,67318686 | 31,435 | 7,92681191 | 40,87436 | 18,66638 | 99,20154 | 37,6892241 |
| Downregulated | ENSMUSG00000069268 | 57,10366903 | 41,4867934 | 38,1253673 | 11,5628091 | 5,08884999 | 6,634846093 | 5,67946579 | 1,22222602 | 1,06775969 | 4,31478604 | 7,72879316 | 7,006798 | 5,00878232 | 1,870957 | 0,796495 | 1,823091581 | 13,23579 | 1,981702978 | 16,15963 | 4,666594 | 31,00048 | 18,844612 |
| Downregulated | ENSMUSG00000097828 | 44,4139648 | 15,956459 | 31,7711394 | 4,81783712 | 0 | 6,634846093 | 0 | 1,22222602 | 0 | 0 | 4,41645324 | 8,758497 | 0 | 0 | 4,77897 | 0 | 8,272368 | 0 | 7,604532 | 0 | 19,48602 | 17,0498871 |
| Downregulated | ENSMUSG00000085558 | 35,95416198 | 52,1244328 | 33,5866331 | 22,1620508 | 10,1777 | 13,26969219 | 3,40767947 | 14,6667123 | 27,7617518 | 3,23608953 | 8,83290647 | 6,130948 | 15,026347 | 6,548349 | 7,168455 | 3,646183163 | 6,617895 | 7,92681191 | 18,06076 | 19,33303 | 21,25747 | 30,5103242 |
| Downregulated | ENSMUSG00000109812 | 161,7937289 | 157,437062 | 215,136001 | 124,300198 | 67,4272624 | 116,7732912 | 165,840401 | 59,8890751 | 36,3038293 | 28,0461092 | 107,098991 | 103,3503 | 37,0649892 | 15,90313 | 40,62124 | 25,52328214 | 112,5042 | 58,46023784 | 126,4253 | 21,333 | 116,0304 | 39,483949 |
| Downregulated | ENSMUSG00000039253 | 28,55183451 | 31,912918 | 34,4943799 | 31,797725 | 25,44425 | 11,94272297 | 22,7178632 | 9,77780817 | 11,7453565 | 10,7869651 | 29,8110593 | 4,379248 | 16,0281034 | 4,677392 | 13,54041 | 10,93854949 | 14,89026 | 22,78958424 | 38,02266 | 25,99959 | 16,82883 | 20,639337 |
| Downregulated | ENSMUSG00000028555 | 35,95416198 | 25,5303344 | 52,6493167 | 31,797725 | 29,2608875 | 34,50119968 | 3,40767947 | 13,4444862 | 6,40655811 | 7,55087556 | 14,353473 | 15,76529 | 6,01053879 | 7,483827 | 9,557939 | 0 | 19,85368 | 6,935960422 | 13,30793 | 25,99959 | 1,771456 | 21,5366995 |
| Downregulated | ENSMUSG00000056185 | 98,34520777 | 51,0606688 | 117,099342 | 55,8869106 | 76,3327499 | 92,8878453 | 46,5716195 | 56,222397 | 46,9814262 | 46,3839499 | 45,2686457 | 25,39964 | 22,0386422 | 9,354784 | 30,26681 | 4,557728953 | 33,91671 | 43,59746551 | 53,23172 | 57,99909 | 46,94358 | 28,7155993 |
| Downregulated | ENSMUSG00000082644 | 86,71297889 | 43,6143213 | 123,45357 | 14,4535114 | 0 | 1,326969219 | 1,13589316 | 33,0001026 | 7,4743178 | 30,2035023 | 8,83290647 | 0,87585 | 4,00702586 | 1,870957 | 1,59299 | 9,115457906 | 0 | 1,981702978 | 6,653965 | 0 | 1,771456 | 7,17889982 |
| Downregulated | ENSMUSG00000078672 | 211,4950705 | 123,396616 | 475,659344 | 76,1218265 | 7,63327499 | 5,307876874 | 5,67946579 | 113,66702 | 10,6775969 | 101,397472 | 27,6028327 | 5,255098 | 27,0474245 | 9,354784 | 5,575465 | 7,292366325 | 0 | 6,935960422 | 26,61586 | 0 | 7,085824 | 36,7918616 |
| Downregulated | ENSMUSG00000066154 | 40,18406339 | 32,976682 | 99,8521524 | 8,67210682 | 2,544425 | 13,26969219 | 0 | 31,7778766 | 9,60983717 | 23,7313232 | 7,72879316 | 0 | 5,00878232 | 3,741914 | 0,796495 | 0,911545791 | 10,75408 | 6,935960422 | 4,752832 | 0,666656 | 6,200096 | 39,483949 |
| Downregulated | ENSMUSG00000059336 | 19,03455634 | 17,0202229 | 38,1253673 | 0,96356742 | 2,544425 | 0 | 1,13589316 | 0 | 0 | 7,55087556 | 0 | 0 | 0 | 0 | 0 | 0 | 0 | 11,89021787 | 4,752832 | 9,333188 | 0,885728 | 7,17889982 |
| Downregulated | ENSMUSG00000041117 | 71,90832396 | 61,6983082 | 78,0662282 | 98,2838772 | 24,1720375 | 47,77089187 | 54,5228716 | 52,5557189 | 38,4393487 | 56,0922185 | 52,9974388 | 26,27549 | 20,0351293 | 19,64505 | 39,82475 | 22,78864477 | 35,57118 | 34,67980211 | 35,17096 | 69,99891 | 32,77194 | 26,9208743 |
| Downregulated | ENSMUSG00000042363 | 577,3815424 | 603,154151 | 833,311599 | 873,955654 | 428,735612 | 452,4965035 | 556,587648 | 601,335203 | 453,797866 | 811,179775 | 525,557935 | 299,5406 | 158,277521 | 194,5795 | 301,0751 | 244,2942719 | 430,9904 | 437,9563581 | 472,4315 | 690,6559 | 372,0058 | 362,534441 |
| Downregulated | ENSMUSG00000045733 | 181,8857606 | 255,303344 | 280,493774 | 290,997362 | 44,5274375 | 147,2935833 | 181,742905 | 343,445512 | 97,1661314 | 146,702725 | 94,9537446 | 15,76529 | 35,0614763 | 11,22574 | 9,557939 | 65,63129693 | 87,68711 | 51,52427742 | 107,414 | 134,6646 | 71,74397 | 139,091184 |
| Downregulated | ENSMUSG00000055745 | 426,162567 | 248,920761 | 356,744508 | 331,467194 | 265,892412 | 163,2172139 | 184,014692 | 256,667465 | 163,367232 | 262,123252 | 107,098991 | 91,08837 | 18,0316164 | 31,80627 | 125,8462 | 35,55028583 | 123,2583 | 111,9662182 | 227,1854 | 149,331 | 98,31581 | 130,117559 |
| Downregulated | ENSMUSG00000034110 | 155,4488768 | 99,9938098 | 149,778229 | 145,498681 | 66,1550499 | 75,63724546 | 106,773957 | 100,222534 | 117,453565 | 151,017511 | 54,1015521 | 56,05438 | 35,0614763 | 28,99983 | 59,73712 | 47,40038111 | 109,1953 | 70,35045571 | 90,30382 | 72,66553 | 46,05786 | 76,2758106 |
| Downregulated | ENSMUSG00000019813 | 76,13822537 | 55,3157246 | 91,6824308 | 81,903231 | 69,9716874 | 58,38664562 | 70,4253758 | 62,3335271 | 74,743178 | 59,328308 | 41,9563057 | 13,13775 | 14,0245905 | 54,25775 | 15,1334 | 7,292366325 | 38,05289 | 34,67980211 | 48,47889 | 77,33213 | 30,11475 | 38,5865865 |
| Downregulated | ENSMUSG00000037108 | 25,37940846 | 15,956459 | 33,5866331 | 21,1984833 | 16,5387625 | 9,28878453 | 15,9025042 | 25,6667465 | 19,2196743 | 15,1017511 | 9,93701978 | 1,751699 | 1,00175646 | 15,90313 | 7,168455 | 4,557728953 | 4,136184 | 2,972554466 | 8,555098 | 18,66638 | 9,743008 | 14,3577996 |
| Downregulated | ENSMUSG00000008226 | 255,9090353 | 230,836774 | 249,630381 | 227,401912 | 109,410275 | 159,2363062 | 182,878798 | 216,334006 | 177,248108 | 127,286188 | 56,3097787 | 51,67513 | 68,1194396 | 118,8058 | 57,34764 | 49,22347269 | 76,93303 | 87,19493102 | 73,19362 | 174,6639 | 130,202 | 131,014922 |
| Downregulated | ENSMUSG00000032109 | 286,5758205 | 286,152498 | 250,538128 | 271,726014 | 169,204262 | 213,6420442 | 191,965944 | 171,111643 | 111,047007 | 242,706715 | 161,200543 | 77,07477 | 88,1545689 | 102,9026 | 93,9864 | 84,77375853 | 71,96961 | 88,1857825 | 137,8321 | 285,3289 | 124,0019 | 133,707009 |
| Downregulated | ENSMUSG00000028344 | 219,9548733 | 155,309534 | 213,320507 | 132,008737 | 119,587975 | 96,86875296 | 165,840401 | 144,222671 | 96,0983717 | 135,91576 | 96,0578579 | 35,90984 | 39,0685021 | 63,61253 | 61,33011 | 75,65830062 | 64,52447 | 92,14918846 | 47,52832 | 179,3305 | 91,22999 | 91,5309727 |
| Downregulated | ENSMUSG00000055660 | 436,7373205 | 379,763724 | 371,268458 | 275,580283 | 188,28745 | 164,5441831 | 157,889149 | 217,556232 | 161,231713 | 245,942804 | 135,805937 | 53,42683 | 92,1615947 | 97,28975 | 162,485 | 42,84265216 | 110,0225 | 145,6551689 | 119,7714 | 231,3297 | 186,8886 | 117,554485 |
| Downregulated | ENSMUSG00000042647 | 62,39104579 | 62,7620721 | 67,1732662 | 50,105506 | 34,3497375 | 34,50119968 | 43,16394 | 33,0001026 | 40,5748681 | 49,6200394 | 27,6028327 | 13,13775 | 22,0386422 | 3,741914 | 29,47031 | 3,646183163 | 18,19921 | 25,76213871 | 19,01133 | 69,33225 | 40,74349 | 25,1261494 |
| Downregulated | ENSMUSG00000025933 | 142,7591726 | 184,031161 | 237,829672 | 156,097923 | 77,6049624 | 132,6969219 | 127,220034 | 139,333766 | 127,063403 | 124,050099 | 145,742957 | 16,64114 | 57,1001185 | 17,77409 | 19,11588 | 18,23091581 | 12,40855 | 55,48768337 | 45,62719 | 95,33185 | 106,2874 | 72,6863607 |
| Downregulated | ENSMUSG00000021120 | 246,3917571 | 292,535082 | 349,482533 | 285,215958 | 222,637187 | 221,6038595 | 232,858097 | 188,222807 | 199,671061 | 305,271112 | 227,447342 | 91,96422 | 112,196724 | 98,22523 | 105,1373 | 93,88921644 | 165,4474 | 123,8564361 | 177,7559 | 221,9965 | 203,7174 | 208,188095 |
| Downregulated | ENSMUSG00000053226 | 68,7358979 | 78,7185311 | 102,575393 | 49,1419386 | 27,988675 | 50,42483031 | 46,5716195 | 31,7778766 | 59,7945424 | 63,643094 | 36,4357392 | 26,27549 | 37,0649892 | 19,64505 | 23,89485 | 12,76164107 | 39,70737 | 27,74384169 | 33,26983 | 54,66581 | 28,3433 | 35,8944991 |
| Downregulated | ENSMUSG00000001663 | 991,9118805 | 1015,89456 | 970,381372 | 1107,13897 | 728,977762 | 1167,732912 | 729,243408 | 508,446025 | 478,356339 | 567,394364 | 446,061777 | 299,5406 | 712,248846 | 320,8691 | 188,7693 | 253,4097298 | 574,9296 | 802,5897059 | 717,6777 | 439,3265 | 1387,05 | 853,391716 |
| Downregulated | ENSMUSG00000100594 | 20,09203169 | 17,0202229 | 26,3246584 | 13,4899439 | 10,1777 | 21,2315075 | 5,67946579 | 0 | 1,06775969 | 19,4165372 | 7,72879316 | 0 | 5,00878232 | 5,61287 | 0,796495 | 2,734637372 | 9,926842 | 9,908514888 | 10,45623 | 0 | 26,57184 | 2,69208743 |
| Downregulated | ENSMUSG00000072476 | 60,27609508 | 43,6143213 | 40,8486078 | 48,1783712 | 40,7108 | 37,15513812 | 26,1255426 | 7,33335613 | 26,6939921 | 29,1248057 | 15,4575863 | 10,5102 | 23,0403987 | 14,03218 | 17,52289 | 15,49627844 | 14,06303 | 12,88106935 | 19,9619 | 23,99963 | 19,48602 | 14,3577996 |
| Downregulated | ENSMUSG00000015850 | 5218,640864 | 2831,73959 | 3394,97318 | 2847,34174 | 2646,202 | 2890,138958 | 2731,82305 | 2434,67424 | 1657,16303 | 2282,52181 | 1761,06073 | 995,8411 | 1356,37825 | 793,2857 | 597,3712 | 672,7207935 | 908,3061 | 1214,783925 | 1537,066 | 803,9875 | 1714,769 | 1651,14696 |
| Downregulated | ENSMUSG00000025350 | 142,7591726 | 101,057574 | 109,837368 | 128,154467 | 105,593637 | 100,8496606 | 99,9585979 | 69,6668832 | 41,6426277 | 79,8235417 | 39,7480791 | 30,65474 | 28,049181 | 14,03218 | 13,54041 | 25,52328214 | 38,05289 | 49,54257444 | 52,28116 | 35,99944 | 11,51446 | 27,8182368 |
| Downregulated | ENSMUSG00000035126 | 97,28773241 | 40,4230295 | 87,1436966 | 72,2675568 | 36,8941625 | 58,38664562 | 36,3485811 | 70,8891093 | 7,4743178 | 30,2035023 | 16,5616996 | 7,006798 | 11,0193211 | 3,741914 | 6,37196 | 2,734637372 | 5,790658 | 21,79873275 | 27,56643 | 21,99966 | 24,80038 | 5,38417486 |
| Downregulated | ENSMUSG00000059479 | 108,9199613 | 95,7387541 | 161,578938 | 134,899439 | 82,6938124 | 82,27209155 | 70,4253758 | 67,2224312 | 40,5748681 | 48,5413429 | 17,6658129 | 11,38605 | 13,022834 | 21,516 | 36,63877 | 42,84265216 | 29,78053 | 43,59746551 | 46,57776 | 51,99919 | 28,3433 | 31,4076867 |
| Downregulated | ENSMUSG00000086877 | 35,95416198 | 47,869377 | 95,3134182 | 59,7411803 | 29,2608875 | 61,04058406 | 40,8921537 | 30,5556505 | 25,6262325 | 21,5739302 | 28,706946 | 10,5102 | 21,0368858 | 11,22574 | 25,48784 | 10,0270037 | 19,85368 | 19,81702978 | 16,15963 | 21,99966 | 28,3433 | 28,7155993 |
| Downregulated | ENSMUSG00000036502 | 25,37940846 | 18,0839869 | 57,1880509 | 18,3077811 | 19,0831875 | 38,48210734 | 19,3101837 | 24,4445204 | 19,2196743 | 20,4952337 | 14,353473 | 1,751699 | 18,0316164 | 5,61287 | 1,59299 | 0 | 7,445132 | 1,981702978 | 10,45623 | 11,99981 | 8,85728 | 11,6657122 |
| Downregulated | ENSMUSG00000086158 | 62,39104579 | 72,3359475 | 124,361317 | 48,1783712 | 40,7108 | 78,2911839 | 49,979299 | 45,2223628 | 42,7103874 | 60,4070045 | 33,1233993 | 11,38605 | 20,0351293 | 24,32244 | 29,47031 | 14,58473265 | 13,23579 | 15,85362382 | 26,61586 | 52,66584 | 42,51494 | 32,3050492 |
| Downregulated | ENSMUSG00000116097 | 37,01163733 | 36,1679738 | 54,4648104 | 28,9070227 | 12,722125 | 34,50119968 | 9,08714527 | 25,6667465 | 21,3551937 | 8,62957207 | 26,4987194 | 10,5102 | 18,0316164 | 0,935478 | 7,168455 | 0 | 6,617895 | 10,89936638 | 11,4068 | 13,33313 | 25,68611 | 4,48681239 |
| Downregulated | ENSMUSG00000029798 | 343,6794895 | 269,132275 | 434,810736 | 142,607979 | 150,121075 | 240,1814286 | 112,453423 | 255,445239 | 238,11041 | 162,883173 | 136,91005 | 63,06118 | 88,1545689 | 67,35444 | 138,5901 | 71,10057167 | 169,5836 | 103,0485548 | 111,2163 | 71,99888 | 123,1162 | 109,478222 |
| Downregulated | ENSMUSG00000008822 | 211,4950705 | 225,517954 | 236,921925 | 140,680844 | 143,760012 | 217,6229519 | 153,345576 | 128,333732 | 167,638271 | 115,420526 | 98,2660845 | 83,20572 | 141,247662 | 93,54784 | 75,66702 | 77,4813922 | 39,70737 | 83,23152506 | 35,17096 | 66,66563 | 110,716 | 81,6599854 |
| Downregulated | ENSMUSG00000089967 | 9,517278171 | 10,6376393 | 8,16972156 | 22,1620508 | 5,08884999 | 9,28878453 | 4,54357263 | 11,0000342 | 9,60983717 | 16,1804476 | 12,1452464 | 0,87585 | 5,00878232 | 2,806435 | 3,18598 | 1,823091581 | 9,926842 | 4,954257444 | 4,752832 | 12,66647 | 15,9431 | 8,97362477 |
| Downregulated | ENSMUSG00000022044 | 17,97708099 | 8,51011147 | 15,4316963 | 31,797725 | 2,544425 | 6,634846093 | 4,54357263 | 22,0000684 | 24,5584728 | 51,7774324 | 15,4575863 | 3,503399 | 13,022834 | 5,61287 | 2,389485 | 0,911545791 | 14,89026 | 0 | 0,950566 | 7,999875 | 30,11475 | 8,97362477 |
| Downregulated | ENSMUSG00000018740 | 97,28773241 | 45,7418492 | 82,6049624 | 131,04517 | 55,9773499 | 51,75179953 | 60,2023374 | 107,55589 | 102,50493 | 116,499223 | 61,8303453 | 33,28229 | 21,0368858 | 31,80627 | 7,964949 | 33,72719425 | 70,31513 | 94,13089144 | 29,46756 | 67,33228 | 97,43008 | 89,7362477 |
| Downregulated | ENSMUSG00000045915 | 2,114950705 | 5,31881967 | 6,35422788 | 36,6155621 | 17,810975 | 6,634846093 | 0 | 20,7778424 | 20,287434 | 19,4165372 | 20,9781529 | 4,379248 | 0 | 2,806435 | 0 | 0 | 0 | 0 | 0 | 1,999969 | 5,314368 | 3,58944991 |
| Downregulated | ENSMUSG00000002012 | 0 | 7,44634754 | 2,72324052 | 25,052753 | 8,90548749 | 14,5966614 | 3,40767947 | 18,3333903 | 21,3551937 | 19,4165372 | 7,72879316 | 0 | 1,00175646 | 1,870957 | 0,796495 | 0 | 0 | 4,954257444 | 7,604532 | 3,999938 | 22,1432 | 8,97362477 |
| Downregulated | ENSMUSG00000055963 | 153,3339261 | 239,346885 | 383,069166 | 453,840257 | 181,926387 | 302,5489818 | 299,875794 | 343,445512 | 254,126805 | 337,632007 | 193,219829 | 66,56458 | 147,2582 | 134,7089 | 74,07403 | 47,40038111 | 95,13224 | 167,4539016 | 192,0144 | 251,3294 | 343,6625 | 248,569406 |
| Downregulated | ENSMUSG00000045211 | 431,4499437 | 324,448 | 389,423394 | 461,548796 | 312,964275 | 285,298382 | 322,593657 | 328,7788 | 291,498394 | 403,432494 | 226,343228 | 107,7295 | 90,1580818 | 145,9346 | 74,87053 | 122,1471359 | 139,803 | 201,1428522 | 192,0144 | 267,9958 | 476,5217 | 313,179505 |
| Downregulated | ENSMUSG00000028737 | 139,5867465 | 143,608131 | 186,995849 | 213,911968 | 115,771337 | 86,25299921 | 97,6868116 | 162,556061 | 98,2338911 | 140,230546 | 76,1838183 | 61,30948 | 53,0930926 | 51,45131 | 75,66702 | 21,87709898 | 112,5042 | 81,24982208 | 79,84759 | 95,9985 | 181,5742 | 149,859534 |
| Downregulated | ENSMUSG00000025656 | 37,01163733 | 49,9969049 | 57,1880509 | 83,8303659 | 44,5274375 | 39,80907656 | 103,366277 | 132,00041 | 105,708209 | 129,443581 | 29,8110593 | 24,52379 | 15,026347 | 26,19339 | 9,557939 | 15,49627844 | 46,32526 | 49,54257444 | 64,63852 | 79,99875 | 89,45853 | 81,6599854 |
| Downregulated | ENSMUSG00000027955 | 472,6914825 | 1259,4965 | 1168,27018 | 1955,0783 | 763,327499 | 857,2221152 | 1672,03473 | 2195,11794 | 1179,87445 | 2382,84059 | 1532,50927 | 530,7649 | 471,827295 | 335,8367 | 302,6681 | 484,9423606 | 687,4338 | 1060,211093 | 952,4676 | 1383,978 | 1128,417 | 1844,97725 |
| Downregulated | ENSMUSG00000054293 | 162,8512043 | 279,769915 | 285,032508 | 296,778767 | 328,230825 | 116,7732912 | 282,837396 | 545,112806 | 344,886378 | 416,376853 | 442,749437 | 108,6054 | 92,1615947 | 73,90279 | 56,55114 | 34,63874004 | 186,9555 | 180,334971 | 288,0216 | 216,6633 | 414,5207 | 453,168051 |
| Downregulated | ENSMUSG00000047757 | 9,517278171 | 14,8926951 | 35,4021268 | 37,5791295 | 20,3554 | 50,42483031 | 26,1255426 | 37,8890067 | 38,4393487 | 39,9117708 | 29,8110593 | 10,5102 | 8,01405172 | 5,61287 | 9,557939 | 6,380820534 | 40,53461 | 14,86277233 | 27,56643 | 77,99878 | 47,82931 | 53,8417486 |
| Downregulated | ENSMUSG00000039167 | 6,344852114 | 0 | 0 | 28,9070227 | 0 | 0 | 0 | 7,33335613 | 21,3551937 | 12,9443581 | 0 | 0 | 0 | 0 | 0 | 0 | 0,827237 | 0 | 0 | 48,66591 | 23,02893 | 24,2287869 |
| Downregulated | ENSMUSG00000033910 | 23,26445775 | 6,3825836 | 0,90774684 | 5,78140454 | 1,2722125 | 3,980907656 | 1,13589316 | 15,8889383 | 30,9650309 | 16,1804476 | 3,31233993 | 0 | 1,00175646 | 2,806435 | 0 | 0 | 12,40855 | 0,990851489 | 1,901133 | 26,66625 | 12,40019 | 27,8182368 |
| Downregulated | ENSMUSG00000020268 | 12,68970423 | 12,7651672 | 29,9556457 | 18,3077811 | 13,9943375 | 25,21241515 | 10,2230384 | 66,0002052 | 52,3202246 | 4,31478604 | 29,8110593 | 0 | 10,0175646 | 7,483827 | 2,389485 | 4,557728953 | 3,308947 | 7,92681191 | 12,35736 | 99,33178 | 2,657184 | 11,6657122 |
| Downregulated | ENSMUSG00000024027 | 14,80465493 | 5,31881967 | 11,8007089 | 23,1256182 | 17,810975 | 1,326969219 | 6,81535895 | 58,666849 | 11,7453565 | 22,6526267 | 12,1452464 | 0 | 5,00878232 | 0 | 0,796495 | 1,823091581 | 0 | 13,87192084 | 9,505665 | 53,3325 | 7,971552 | 27,8182368 |
| Downregulated | ENSMUSG00000064115 | 5,287376762 | 35,1042098 | 19,0626836 | 10,5992417 | 10,1777 | 5,307876874 | 13,6307179 | 50,1112669 | 39,5071084 | 44,2265569 | 9,93701978 | 0 | 3,00526939 | 4,677392 | 2,389485 | 0 | 2,481711 | 6,935960422 | 10,45623 | 45,33263 | 3,542912 | 8,97362477 |
| Downregulated | ENSMUSG00000042251 | 26,43688381 | 107,440157 | 27,2324052 | 53,9597757 | 58,5217749 | 90,23390687 | 159,025042 | 51,3334929 | 32,0327906 | 155,332297 | 90,5372913 | 11,38605 | 23,0403987 | 19,64505 | 9,557939 | 30,08101109 | 12,40855 | 59,45108933 | 40,87436 | 135,9979 | 151,4595 | 163,319971 |
| Downregulated | ENSMUSG00000033906 | 125,8395669 | 159,56459 | 119,822583 | 107,919551 | 52,1607124 | 90,23390687 | 223,770952 | 125,88928 | 102,50493 | 194,165372 | 152,367637 | 24,52379 | 52,0913362 | 70,16088 | 13,54041 | 48,3119269 | 63,69724 | 101,0668519 | 132,1287 | 131,9979 | 112,4875 | 141,783271 |
| Downregulated | ENSMUSG00000029456 | 94,11530636 | 99,9938098 | 98,0366587 | 83,8303659 | 69,9716874 | 99,5226914 | 131,763606 | 108,778116 | 104,640449 | 172,591441 | 45,2686457 | 31,53059 | 47,0825538 | 10,29026 | 27,08083 | 26,43482793 | 69,48789 | 65,39619826 | 120,7219 | 142,6644 | 87,68707 | 61,0206485 |
| Downregulated | ENSMUSG00000086968 | 40,18406339 | 32,976682 | 41,7563546 | 40,4698318 | 29,2608875 | 15,92363062 | 40,8921537 | 33,0001026 | 13,8808759 | 86,2957207 | 34,2275126 | 5,255098 | 10,0175646 | 4,677392 | 19,91237 | 7,292366325 | 8,272368 | 13,87192084 | 47,52832 | 65,33231 | 48,71504 | 25,1261494 |
| Downregulated | ENSMUSG00000066721 | 27,49435916 | 32,976682 | 28,140152 | 42,3969667 | 29,2608875 | 37,15513812 | 30,6691153 | 19,5556163 | 26,6939921 | 36,6756813 | 34,2275126 | 13,13775 | 3,00526939 | 12,16122 | 15,1334 | 8,203912116 | 18,19921 | 16,84447531 | 36,12153 | 55,33247 | 36,31485 | 17,0498871 |
| Downregulated | ENSMUSG00000031840 | 70,85084861 | 62,7620721 | 106,20638 | 149,352951 | 76,3327499 | 84,92602999 | 99,9585979 | 55,000171 | 64,0655811 | 119,735313 | 70,6632518 | 55,17853 | 27,0474245 | 28,99983 | 31,8598 | 11,85009528 | 59,56105 | 36,66150509 | 70,34192 | 127,3313 | 77,05834 | 53,8417486 |
| Downregulated | ENSMUSG00000036411 | 47,58639085 | 15,956459 | 39,940861 | 32,7612924 | 34,3497375 | 47,77089187 | 21,58197 | 50,1112669 | 39,5071084 | 71,1939696 | 41,9563057 | 4,379248 | 9,01580818 | 2,806435 | 7,964949 | 5,469274744 | 9,099605 | 8,917663399 | 8,555098 | 63,33234 | 15,05738 | 7,17889982 |
| Downregulated | ENSMUSG00000027009 | 16,91960564 | 6,3825836 | 20,8781773 | 23,1256182 | 17,810975 | 22,55847672 | 18,1742905 | 34,2223286 | 18,1519147 | 34,5182883 | 24,2904928 | 0 | 3,00526939 | 1,870957 | 0 | 3,646183163 | 0 | 10,89936638 | 0 | 25,99959 | 3,542912 | 5,38417486 |
| Downregulated | ENSMUSG00000043557 | 251,6791339 | 268,068511 | 306,818432 | 299,669469 | 377,847112 | 379,5131965 | 602,023374 | 429,001334 | 257,330084 | 624,565279 | 436,124757 | 102,4744 | 78,1370042 | 64,54801 | 56,55114 | 127,6164107 | 119,1221 | 220,959882 | 264,2575 | 417,3268 | 313,5477 | 284,463905 |
| Downregulated | ENSMUSG00000026589 | 943,2680143 | 613,79179 | 837,850333 | 571,395482 | 1050,84752 | 1169,059882 | 1277,8798 | 1091,44784 | 714,33123 | 1095,95565 | 1064,36523 | 349,464 | 261,458437 | 114,1284 | 66,10908 | 94,80076223 | 290,3601 | 611,3553686 | 564,6365 | 1507,976 | 896,3568 | 883,004678 |
| Downregulated | ENSMUSG00000002265 | 817,4284473 | 493,586465 | 1140,13003 | 527,071381 | 973,242561 | 1117,308082 | 940,519535 | 1431,22667 | 627,842695 | 1307,38017 | 971,619712 | 340,7055 | 252,442629 | 225,4503 | 175,2289 | 193,2477076 | 335,8582 | 313,1090705 | 547,5263 | 1093,316 | 608,4951 | 1026,58267 |
| Downregulated | ENSMUSG00000039542 | 299,2655247 | 325,511764 | 652,669978 | 507,800032 | 256,986925 | 314,4917048 | 386,203674 | 369,112259 | 440,98475 | 550,13522 | 351,108032 | 138,3843 | 124,217802 | 57,06418 | 48,58619 | 112,1201322 | 128,2217 | 261,584793 | 181,5582 | 1070,65 | 286,9759 | 338,305654 |
| Downregulated | ENSMUSG00000029563 | 80,36812678 | 120,205325 | 101,667646 | 56,850478 | 110,682487 | 120,7541989 | 92,0073458 | 127,111506 | 85,4207749 | 134,837064 | 65,1426852 | 22,77209 | 36,0632327 | 9,354784 | 40,62124 | 30,99255688 | 47,1525 | 44,588317 | 83,64985 | 215,33 | 71,74397 | 70,8916357 |
| Downregulated | ENSMUSG00000029716 | 9,517278171 | 17,0202229 | 32,6788862 | 17,3442136 | 13,9943375 | 45,11695343 | 18,1742905 | 37,8890067 | 14,9486356 | 24,8100197 | 5,52056654 | 0,87585 | 3,00526939 | 9,354784 | 0,796495 | 6,380820534 | 3,308947 | 14,86277233 | 6,653965 | 79,99875 | 12,40019 | 6,28153734 |
| Downregulated | ENSMUSG00000028197 | 51,81629226 | 14,8926951 | 34,4943799 | 20,2349159 | 15,26655 | 37,15513812 | 52,2510853 | 90,4447256 | 10,6775969 | 79,8235417 | 24,2904928 | 14,88944 | 2,00351293 | 0 | 0 | 0 | 0 | 36,66150509 | 35,17096 | 141,3311 | 58,45805 | 40,3813115 |
| Downregulated | ENSMUSG00000031216 | 172,3684824 | 144,671895 | 115,283849 | 113,700956 | 57,2495624 | 82,27209155 | 71,561269 | 160,111609 | 84,3530152 | 145,624029 | 82,8084982 | 33,28229 | 45,0790409 | 12,16122 | 62,12661 | 54,69274744 | 86,85987 | 102,0577033 | 107,414 | 287,3288 | 148,8023 | 109,478222 |
| Downregulated | ENSMUSG00000056832 | 187,1731374 | 103,185102 | 202,427545 | 173,442136 | 125,949037 | 71,65633781 | 139,714858 | 278,667533 | 177,248108 | 248,100197 | 112,619557 | 35,03399 | 76,1334913 | 34,6127 | 80,44599 | 41,01956058 | 99,26842 | 164,4813471 | 98,85892 | 419,3268 | 230,2893 | 140,885909 |
| Downregulated | ENSMUSG00000047497 | 689,4739297 | 845,692327 | 1001,24476 | 628,24596 | 554,684649 | 686,043086 | 646,323207 | 1009,55869 | 723,941067 | 1300,90799 | 692,279045 | 176,0458 | 380,667457 | 249,7727 | 142,5726 | 171,3706086 | 335,0309 | 554,8768337 | 647,3358 | 1658,641 | 1332,135 | 1144,13716 |
| Downregulated | ENSMUSG00000033595 | 157,5638275 | 192,541272 | 137,97752 | 148,389383 | 245,537012 | 348,9929045 | 415,736896 | 525,557189 | 338,47982 | 374,307689 | 327,921653 | 13,13775 | 53,0930926 | 37,41914 | 40,62124 | 20,96555318 | 16,54474 | 43,59746551 | 53,23172 | 1110,649 | 705,9252 | 238,698419 |
| Downregulated | ENSMUSG00000022367 | 2607,734219 | 1439,2726 | 3106,30969 | 1110,02967 | 1234,04612 | 1285,833173 | 1601,60935 | 2198,78461 | 1960,40678 | 2185,43913 | 1540,23807 | 657,7631 | 1562,74008 | 547,2549 | 468,339 | 370,087591 | 637,7996 | 977,9704195 | 918,2472 | 5889,241 | 4591,614 | 1791,1355 |
| Downregulated | ENSMUSG00000074793 | 43,35648945 | 73,3997114 | 60,8190383 | 91,5389053 | 43,255225 | 107,4845067 | 106,773957 | 106,333664 | 50,1847052 | 97,0826858 | 25,3946061 | 22,77209 | 25,0439116 | 31,80627 | 20,70887 | 15,49627844 | 20,68092 | 47,56087146 | 39,92379 | 99,99844 | 75,28688 | 64,6100984 |
| Downregulated | ENSMUSG00000001493 | 5,287376762 | 10,6376393 | 16,3394431 | 32,7612924 | 29,2608875 | 39,80907656 | 17,0383974 | 36,6667807 | 21,3551937 | 33,4395918 | 2,20822662 | 0,87585 | 2,00351293 | 0 | 1,59299 | 1,823091581 | 0 | 0 | 6,653965 | 46,66594 | 15,9431 | 9,87098725 |
| Downregulated | ENSMUSG00000068299 | 12,68970423 | 9,57387541 | 4,5387342 | 21,1984833 | 12,722125 | 18,57756906 | 22,7178632 | 18,3333903 | 7,4743178 | 31,2821988 | 5,52056654 | 0 | 8,01405172 | 0 | 10,35443 | 17,31937002 | 1,654474 | 2,972554466 | 4,752832 | 41,99934 | 20,37174 | 17,0498871 |
| Downregulated | ENSMUSG00000094526 | 2288,376662 | 2339,21689 | 2111,41915 | 2416,6271 | 2643,65757 | 2308,92644 | 3714,37063 | 3105,67632 | 2828,49541 | 2395,78495 | 2180,62378 | 581,5642 | 596,045096 | 770,8342 | 446,0372 | 826,7720321 | 1153,168 | 1319,814183 | 1596,001 | 3467,946 | 4152,293 | 3061,80077 |

| | | | | | | | | | | | | | | | | | | | | | | | |
|---|---|---|---|---|---|---|---|---|---|---|---|---|---|---|---|---|---|---|---|---|---|---|---|
| Downregulated | ENSMUSG00000028497 | 510,7605952 | 601,026623 | 524,677673 | 515,508572 | 614,478637 | 683,3891476 | 492,977631 | 526,779415 | 489,033936 | 574,945239 | 423,979511 | 47,29588 | 119,209019 | 23,38696 | 30,26681 | 39,196469 | 129,0489 | 393,3680411 | 327,9454 | 661,323 | 635,9527 | 589,567148 |
| Downregulated | ENSMUSG00000026837 | 57510,79704 | 56919,8806 | 54785,245 | 62892,0458 | 65041,864 | 64704,34607 | 69829,0319 | 68494,7685 | 50880,8845 | 82785,6423 | 73649,8783 | 16904,77 | 21564,8114 | 23652,64 | 15092,78 | 23049,34686 | 13570,82 | 32736,74234 | 28156,73 | 62721,69 | 77771,35 | 53120,2692 |
| Downregulated | ENSMUSG00000020150 | 116,3222888 | 159,56459 | 231,475444 | 296,778767 | 161,570987 | 264,0668745 | 227,178632 | 235,889622 | 259,465604 | 222,211481 | 83,9126115 | 50,79928 | 88,1545689 | 42,09653 | 50,17918 | 27,34637372 | 86,85987 | 146,6460203 | 137,8321 | 366,6609 | 371,12 | 132,809647 |
| Downregulated | ENSMUSG00000024228 | 12,68970423 | 7,44634754 | 8,16972156 | 22,1620508 | 12,722125 | 15,92363062 | 15,9025042 | 6,11113011 | 11,7453565 | 7,55087556 | 11,0411331 | 0 | 2,00351293 | 0 | 8,761444 | 0 | 0 | 1,981702978 | 2,851699 | 25,33294 | 24,80038 | 7,17889982 |
| Downregulated | ENSMUSG00000057596 | 99,40268312 | 43,6143213 | 70,8042535 | 39,5062644 | 21,6276125 | 2,653938437 | 12,4948247 | 62,3335271 | 69,4043796 | 48,5413429 | 2,20822662 | 3,503399 | 0 | 7,483827 | 19,11588 | 21,87709898 | 80,24197 | 19,81702978 | 19,9619 | 20,66634 | 7,085824 | 8,97362477 |
| Downregulated | ENSMUSG00000105403 | 48,64386621 | 43,6143213 | 40,8486078 | 38,542697 | 2,544425 | 18,57756906 | 17,0383974 | 29,3334245 | 11,7453565 | 24,8100197 | 13,2493597 | 7,006798 | 2,00351293 | 14,96765 | 16,72639 | 17,31937002 | 50,46145 | 20,80788127 | 48,47889 | 19,99969 | 5,314368 | 11,6657122 |
| Downregulated | ENSMUSG00000000901 | 3373,346374 | 1033,97854 | 2466,34816 | 1052,21563 | 798,949449 | 859,8760537 | 843,968617 | 785,891332 | 927,883167 | 2309,48923 | 1433,13907 | 798,7749 | 464,815 | 763,3504 | 651,5329 | 778,4601052 | 3522,374 | 2920,039338 | 2613,107 | 1252,647 | 1377,307 | 1199,77363 |
| Downregulated | ENSMUSG00000051200 | 28,55183451 | 2,12752787 | 37,2176204 | 0,96356742 | 0 | 0 | 0 | 0 | 5,33879843 | 12,9443581 | 0 | 0 | 6,01053879 | 0 | 0 | 15,49627844 | 35,57118 | 3,963405955 | 56,08342 | 0 | 0 | 0 |
| Downregulated | ENSMUSG00000047759 | 64,50599649 | 20,2115147 | 13,6162026 | 11,5628091 | 6,36106249 | 19,90453828 | 3,40767947 | 6,11113011 | 14,9486356 | 21,5739302 | 17,6658129 | 42,04079 | 16,0281034 | 4,677392 | 26,28433 | 2,734637372 | 9,926842 | 7,92681191 | 3,802266 | 1,999969 | 3,542912 | 3,58944991 |
| Downregulated | ENSMUSG00000073125 | 300,3230001 | 7,44634754 | 80,7894688 | 39,5062644 | 77,6049624 | 18,57756906 | 60,2023374 | 35,4445546 | 93,9628523 | 98,1613823 | 161,200543 | 27,15134 | 22,0386422 | 30,87079 | 25,48784 | 44,66574374 | 9,926842 | 44,588317 | 42,77549 | 30,66619 | 46,94358 | 52,0470237 |
| Downregulated | ENSMUSG00000032489 | 80,36812678 | 24,4665705 | 32,6788862 | 26,0163204 | 10,1777 | 5,307876874 | 28,397329 | 17,1111643 | 21,3551937 | 29,1248057 | 26,4987194 | 5,255098 | 23,0403987 | 20,58052 | 18,31938 | 3,646183163 | 15,7175 | 11,89021787 | 27,56643 | 21,333 | 20,37174 | 18,844612 |
| Downregulated | ENSMUSG00000056091 | 348,9668663 | 154,24577 | 114,376102 | 145,498681 | 58,5217749 | 22,55847672 | 14,7666111 | 52,5557189 | 28,8295115 | 157,48969 | 87,2249514 | 86,70912 | 156,274008 | 57,06418 | 11,15093 | 34,63874004 | 36,39842 | 126,8289906 | 127,3759 | 22,66631 | 93,88717 | 85,2494354 |
| Downregulated | ENSMUSG00000041351 | 31,72426057 | 13,8289311 | 10,8929621 | 9,63567424 | 12,722125 | 22,55847672 | 13,6307179 | 4,88890409 | 5,33879843 | 5,39348255 | 1,10411331 | 4,379248 | 2,00351293 | 0 | 7,168455 | 7,292366325 | 0 | 9,908514888 | 9,505665 | 13,33313 | 7,085824 | 3,58944991 |
| Downregulated | ENSMUSG00000038637 | 145,9315986 | 102,121338 | 82,6049624 | 54,9233432 | 71,2438999 | 92,8878453 | 82,9202006 | 42,7779108 | 54,455744 | 55,013522 | 27,6028327 | 25,39964 | 15,026347 | 41,16105 | 20,70887 | 30,08101109 | 37,22566 | 63,41449528 | 61,78682 | 45,99928 | 98,31581 | 33,2024117 |
| Downregulated | ENSMUSG00000028453 | 333,104736 | 189,34998 | 253,261368 | 105,992417 | 160,298775 | 127,389045 | 92,0073458 | 94,1114037 | 84,3530152 | 169,355352 | 90,5372913 | 69,19213 | 62,1089008 | 28,99983 | 78,05651 | 66,54284272 | 138,1486 | 139,7100599 | 178,7065 | 177,9972 | 110,716 | 92,4283352 |
| Downregulated | ENSMUSG00000036181 | 912,601229 | 427,633101 | 525,58542 | 504,90933 | 358,763925 | 327,761397 | 331,680802 | 278,667533 | 116,385806 | 319,294167 | 211,989755 | 93,71592 | 242,425064 | 161,8378 | 147,3516 | 105,7393117 | 205,982 | 325,9901398 | 447,7168 | 363,9943 | 397,6919 | 324,845217 |
| Downregulated | ENSMUSG00000037921 | 242,1618557 | 86,1648786 | 190,626836 | 106,955984 | 38,166375 | 141,9857064 | 170,383974 | 111,222568 | 104,640449 | 110,027044 | 86,1208381 | 15,76529 | 39,0685021 | 13,0967 | 77,26001 | 66,54284272 | 112,5042 | 44,588317 | 81,74872 | 46,66594 | 93,00144 | 41,278674 |
| Downregulated | ENSMUSG00000032356 | 366,9439473 | 60,6345442 | 186,995849 | 30,8341576 | 75,0605374 | 78,2911839 | 151,07379 | 25,6667465 | 62,9978215 | 89,5318103 | 64,0385719 | 27,15134 | 10,0175646 | 4,677392 | 30,26681 | 17,31937002 | 129,0489 | 5,945108933 | 110,2657 | 41,99934 | 15,05738 | 18,844612 |
| Downregulated | ENSMUSG00000020638 | 352,1392923 | 249,984524 | 475,659344 | 219,693373 | 138,671162 | 367,5704736 | 139,714858 | 124,667054 | 158,028433 | 476,783857 | 134,701824 | 44,66833 | 39,0685021 | 13,0967 | 196,7343 | 138,5549602 | 86,85987 | 51,52427742 | 103,6117 | 64,66566 | 77,94407 | 76,2758106 |
| Downregulated | ENSMUSG00000023341 | 322,5299825 | 178,712341 | 298,64871 | 122,373063 | 111,9547 | 248,1432439 | 104,502171 | 67,2224312 | 57,659023 | 290,169361 | 69,5591385 | 74,44722 | 30,0526939 | 5,61287 | 143,3691 | 94,80076223 | 130,7034 | 62,4236438 | 45,62719 | 10,6665 | 69,08679 | 47,5602113 |
| Downregulated | ENSMUSG00000033355 | 1065,935155 | 1096,74062 | 1476,90411 | 570,431915 | 519,062699 | 1319,007403 | 406,649751 | 349,556642 | 444,188029 | 1075,46042 | 762,942296 | 230,3485 | 163,286304 | 89,80593 | 521,7042 | 466,7114448 | 440,09 | 308,154813 | 308,9341 | 54,66581 | 592,552 | 308,692692 |
| Downregulated | ENSMUSG00000082292 | 192,4605141 | 185,094925 | 157,040203 | 79,0125288 | 76,3327499 | 207,0071981 | 26,1255426 | 68,4446572 | 70,4721393 | 146,702725 | 64,0385719 | 7,006798 | 13,022834 | 10,29026 | 36,63877 | 30,08101109 | 15,7175 | 9,908514888 | 2,851699 | 0 | 11,51446 | 27,8182368 |
| Downregulated | ENSMUSG00000090628 | 60,27609508 | 15,956459 | 49,9260762 | 26,0163204 | 8,90548749 | 55,73270718 | 23,8537563 | 20,7778424 | 2,13551937 | 34,5182883 | 12,1452464 | 13,13775 | 2,00351293 | 0 | 12,74392 | 6,380820534 | 8,272368 | 11,89021787 | 4,752832 | 3,333281 | 0 | 0 |
| Downregulated | ENSMUSG00000050541 | 21,14950705 | 11,7014033 | 9,0774684 | 0,96356742 | 8,90548749 | 22,55847672 | 3,40767947 | 6,11113011 | 0 | 19,4165372 | 4,41645324 | 0,87585 | 3,00526939 | 0 | 0 | 0 | 0 | 4,954257444 | 1,901133 | 3,333281 | 7,085824 | 6,28153734 |
| Downregulated | ENSMUSG00000028088 | 211,4950705 | 189,34998 | 121,638077 | 67,4497197 | 128,493462 | 203,0262904 | 69,2894827 | 55,000171 | 80,0819764 | 179,063621 | 162,304656 | 53,42683 | 42,0737715 | 42,09653 | 39,02825 | 19,1424616 | 12,40855 | 56,47853486 | 73,19362 | 42,666 | 76,17261 | 116,657122 |
| Downregulated | ENSMUSG00000033170 | 801,5663171 | 293,598846 | 374,899445 | 210,057698 | 234,0871 | 329,0883662 | 170,383974 | 189,445033 | 270,1432 | 302,035023 | 191,011602 | 169,039 | 209,367101 | 79,51566 | 87,61444 | 94,80076223 | 62,87 | 89,17663399 | 143,5355 | 56,66578 | 234,7179 | 207,290732 |
| Downregulated | ENSMUSG00000091017 | 47,58639085 | 10,6376393 | 24,5091647 | 21,1984833 | 26,7164625 | 38,48210734 | 7,95125211 | 7,33335613 | 10,6775969 | 20,4952337 | 15,4575863 | 15,76529 | 11,0193211 | 8,419306 | 13,54041 | 0,911545791 | 4,963421 | 0,990851489 | 2,851699 | 7,333219 | 3,542912 | 12,5630747 |
| Downregulated | ENSMUSG00000094066 | 151,2189754 | 25,5303344 | 123,45357 | 40,4698318 | 21,6276125 | 26,53938437 | 68,1535895 | 135,667088 | 82,2174958 | 55,013522 | 29,8110593 | 22,77209 | 28,049181 | 30,87079 | 18,31938 | 18,23091581 | 14,89026 | 29,72554466 | 39,92379 | 129,998 | 115,1446 | 62,8153734 |
| Downregulated | ENSMUSG00000078721 | 120,5521902 | 17,0202229 | 111,652861 | 26,0163204 | 7,63327499 | 42,463015 | 38,6203674 | 74,5557873 | 54,455744 | 23,7313232 | 27,6028327 | 14,0136 | 26,0456681 | 12,16122 | 15,9299 | 11,85009528 | 15,7175 | 25,76213871 | 38,97323 | 125,3314 | 59,34378 | 33,2024117 |
| Downregulated | ENSMUSG00000078746 | 137,4717958 | 24,4665705 | 148,870482 | 23,1256182 | 22,899825 | 42,463015 | 34,0767947 | 94,1114037 | 81,1497361 | 46,3839499 | 34,2275126 | 21,89624 | 37,0649892 | 20,58052 | 17,52289 | 24,61173635 | 15,7175 | 33,68895062 | 55,13286 | 110,6649 | 106,2874 | 42,1760364 |
| Downregulated | ENSMUSG00000093996 | 130,0694683 | 15,956459 | 155,22471 | 54,9233432 | 7,63327499 | 33,17423047 | 42,0280469 | 127,111506 | 68,3366199 | 21,5739302 | 33,1233993 | 28,90304 | 19,0333728 | 16,83861 | 32,65629 | 16,40782423 | 14,89026 | 22,78958424 | 53,23172 | 112,6649 | 90,34426 | 27,8182368 |
| Downregulated | ENSMUSG00000038286 | 273,8861163 | 223,390426 | 273,231799 | 242,818991 | 190,831875 | 212,315075 | 153,345576 | 185,778355 | 169,77379 | 179,063621 | 142,430617 | 27,15134 | 97,1703771 | 76,70923 | 39,82475 | 39,196469 | 148,9026 | 146,6460203 | 178,7065 | 273,9957 | 253,3182 | 162,422608 |
| Downregulated | ENSMUSG00000042156 | 190,3455634 | 161,692118 | 205,150786 | 145,498681 | 105,593637 | 177,8138753 | 157,889149 | 155,222705 | 85,4207749 | 145,624029 | 146,84707 | 28,02719 | 74,1299784 | 57,99966 | 73,27754 | 71,10057167 | 74,45132 | 111,9662182 | 138,7827 | 191,997 | 171,8312 | 106,786135 |
| Downregulated | ENSMUSG00000059901 | 788,8766128 | 375,508669 | 1308,0632 | 513,581437 | 454,179862 | 666,1385477 | 661,089818 | 768,780168 | 599,013184 | 1072,22433 | 659,155645 | 133,1292 | 63,1106573 | 110,3864 | 62,9231 | 149,4935097 | 171,238 | 216,0056246 | 353,6107 | 941,9853 | 1335,678 | 560,851548 |
| Downregulated | ENSMUSG00000056481 | 15640,06046 | 6238,97547 | 19831,5452 | 11052,1184 | 7667,62473 | 11528,70857 | 11229,4398 | 8914,9166 | 6836,86527 | 11583,0431 | 6086,97667 | 1977,669 | 3426,00711 | 1644,571 | 1290,322 | 2082,882132 | 2878,784 | 4335,966115 | 5354,541 | 10298,51 | 19222,07 | 6936,61195 |
| Downregulated | ENSMUSG00000041809 | 78,25317607 | 37,2317377 | 94,4056714 | 65,5225848 | 43,255225 | 54,40573796 | 47,7075126 | 33,0001026 | 53,3879843 | 62,5643975 | 55,2056654 | 10,5102 | 38,0667457 | 19,64505 | 23,89485 | 10,93854949 | 26,47158 | 29,72554466 | 43,72606 | 71,33222 | 101,8587 | 38,5865865 |
| Downregulated | ENSMUSG00000024420 | 918,9460812 | 536,137023 | 1015,76871 | 560,796241 | 688,266962 | 883,7614996 | 486,162272 | 697,891058 | 584,064548 | 886,688531 | 378,710865 | 271,5134 | 305,535722 | 383,5461 | 232,5765 | 119,4124986 | 354,8846 | 480,5629721 | 603,6097 | 815,9873 | 1049,588 | 718,787344 |
| Downregulated | ENSMUSG00000067158 | 181,8857606 | 115,950269 | 222,397976 | 60,7047477 | 45,79965 | 167,1981215 | 67,0176963 | 178,444999 | 73,6754183 | 124,050099 | 175,554016 | 26,27549 | 92,1615947 | 41,16105 | 25,48784 | 37,37337742 | 19,02645 | 12,88106935 | 57,98456 | 53,99916 | 132,8592 | 163,319971 |
| Downregulated | ENSMUSG00000070371 | 119,4947148 | 60,6345442 | 111,652861 | 58,7776129 | 30,5331 | 66,34846093 | 60,2023374 | 66,0002052 | 38,4393487 | 69,0365766 | 81,7043849 | 19,26869 | 56,098362 | 15,90313 | 40,62124 | 20,96555318 | 56,25211 | 44,588317 | 71,29249 | 46,66594 | 80,60125 | 87,0441603 |
| Downregulated | ENSMUSG00000030882 | 16,91960564 | 3,1912918 | 19,0626836 | 14,4535114 | 12,722125 | 25,21241515 | 35,2126879 | 45,2223628 | 21,3551937 | 10,7869651 | 32,019286 | 2,627549 | 4,00702586 | 0,935478 | 0 | 5,469274744 | 20,68092 | 14,86277233 | 20,91246 | 25,33294 | 10,62874 | 8,97362477 |
| Downregulated | ENSMUSG00000028845 | 75,08075001 | 37,2317377 | 48,1105825 | 50,105506 | 39,4385875 | 37,15513812 | 43,16394 | 68,4446572 | 79,0142167 | 62,5643975 | 78,3920449 | 31,53059 | 15,026347 | 7,483827 | 10,35443 | 16,40782423 | 34,74395 | 57,46938635 | 51,33059 | 59,33241 | 41,62922 | 24,2287869 |
| Downregulated | ENSMUSG00000025407 | 119,4947148 | 10,6376393 | 182,457115 | 72,2675568 | 47,0718625 | 18,57756906 | 54,5228716 | 182,111677 | 193,264503 | 74,4300591 | 81,7043849 | 23,64794 | 31,0544504 | 28,06435 | 20,70887 | 20,05400739 | 28,12605 | 77,28641613 | 67,49022 | 16,66641 | 46,05786 | 65,5074609 |
| Downregulated | ENSMUSG00000050765 | 82,48307748 | 61,6983082 | 146,147241 | 102,138147 | 31,8053125 | 94,21481452 | 59,0664442 | 144,222671 | 96,0983717 | 58,2496115 | 87,2249514 | 24,52379 | 35,0614763 | 13,0967 | 12,74392 | 8,203912116 | 49,63421 | 83,23152506 | 117,8702 | 38,66606 | 78,82979 | 72,6863607 |
| Downregulated | ENSMUSG00000045287 | 368,0014226 | 278,706151 | 348,574787 | 242,818991 | 339,680737 | 217,6229519 | 357,806345 | 537,77945 | 428,171634 | 394,802922 | 288,173574 | 54,30268 | 130,22834 | 73,90279 | 145,7586 | 107,5624033 | 118,2949 | 147,6368718 | 335,55 | 276,6623 | 302,0333 | 243,185231 |
| Downregulated | ENSMUSG00000090145 | 537,197479 | 426,569338 | 564,618534 | 333,394329 | 463,08535 | 331,7423047 | 511,151921 | 728,446709 | 581,929029 | 607,306135 | 496,850989 | 159,4046 | 178,312651 | 118,8058 | 82,83547 | 111,2085865 | 143,9392 | 353,7339815 | 371,6715 | 307,3285 | 660,7531 | 538,417486 |
| Downregulated | ENSMUSG00000000037 | 83,54055283 | 74,4634754 | 85,328203 | 50,105506 | 40,7108 | 70,32936859 | 62,4741237 | 116,111472 | 82,2174958 | 77,6661487 | 70,6632518 | 25,39964 | 19,0333728 | 20,58052 | 21,50536 | 10,93854949 | 43,84355 | 63,41449528 | 64,63852 | 93,33188 | 48,71504 | 84,3520729 |
| Downregulated | ENSMUSG00000047996 | 307,7253275 | 432,951921 | 326,788862 | 370,973458 | 237,903737 | 309,1838279 | 313,506512 | 562,22397 | 510,38913 | 261,044555 | 215,302095 | 67,44043 | 107,187942 | 127,2251 | 109,9163 | 120,3240444 | 208,4637 | 302,2097041 | 237,6416 | 313,9951 | 320,6335 | 304,20588 |
| Downregulated | ENSMUSG00000034842 | 234,7595282 | 237,219357 | 245,091647 | 176,332839 | 108,138062 | 236,2005209 | 156,753256 | 474,223696 | 357,699495 | 249,178894 | 140,22239 | 40,28909 | 92,1615947 | 86,06401 | 21,50536 | 32,81564846 | 128,2217 | 143,6734659 | 143,5355 | 140,6645 | 416,2922 | 312,282142 |
| Downregulated | ENSMUSG00000031465 | 225,24225 | 198,923856 | 132,531039 | 132,972305 | 95,4159374 | 76,96421468 | 63,6100169 | 78,2224654 | 77,9464571 | 96,0039893 | 221,926775 | 62,18533 | 64,1124137 | 20,58052 | 39,82475 | 50,13501849 | 36,39842 | 91,15833697 | 69,39135 | 59,99906 | 60,22951 | 157,935796 |
| Downregulated | ENSMUSG00000086233 | 41,24153874 | 49,9969049 | 18,1549368 | 40,4698318 | 36,8941625 | 17,25059984 | 20,4460768 | 24,4445204 | 21,3551937 | 37,7543778 | 41,9563057 | 0,87585 | 18,0316164 | 1,870957 | 13,54041 | 10,0270037 | 11,58132 | 18,82617829 | 26,61586 | 23,99963 | 27,45757 | 42,1760364 |
| Downregulated | ENSMUSG00000068747 | 87,77045424 | 9,57387541 | 19,9704305 | 48,1783712 | 30,5331 | 18,57756906 | 20,4460768 | 31,7778766 | 24,5584728 | 35,5969848 | 48,5809856 | 12,2619 | 19,0333728 | 0 | 1,59299 | 8,203912116 | 3,308947 | 32,69809913 | 11,4068 | 29,33288 | 38,97203 | 69,9942732 |
| Downregulated | ENSMUSG00000055421 | 185,0581867 | 79,782295 | 76,2507346 | 57,8140454 | 0 | 58,38664562 | 0 | 147,889349 | 50,1847052 | 36,6756813 | 7,72879316 | 2,627549 | 13,022834 | 15,90313 | 15,9299 | 2,734637372 | 25,64434 | 6,935960422 | 64,63852 | 67,99894 | 37,20058 | 18,844612 |
| Downregulated | ENSMUSG00000004328 | 43,35648945 | 10,6376393 | 26,3246584 | 8,67210682 | 10,1777 | 11,94272297 | 5,67946579 | 42,7779108 | 6,40655811 | 15,1017511 | 14,353473 | 0 | 0 | 0 | 6,37196 | 0,911545791 | 0 | 5,945108933 | 2,851699 | 0 | 3,542912 | 8,0762623 |
| Downregulated | ENSMUSG00000028488 | 28,55183451 | 7,44634754 | 9,98521524 | 18,3077811 | 0 | 1,326969219 | 0 | 33,0001026 | 6,40655811 | 14,0230546 | 7,72879316 | 0 | 7,01229525 | 2,806435 | 3,982475 | 0 | 4,136184 | 7,92681191 | 0 | 1,333313 | 0,885728 | 10,7683497 |
| Downregulated | ENSMUSG00000029650 | 71,90832396 | 54,2519606 | 31,7711394 | 49,1419386 | 55,9773499 | 22,55847672 | 45,4357263 | 62,3335271 | 20,287434 | 40,9904673 | 40,8521924 | 5,255098 | 10,0175646 | 2,806435 | 23,09835 | 24,61173635 | 19,85368 | 32,69809913 | 6,653965 | 6,666563 | 31,88621 | 29,6129618 |
| Downregulated | ENSMUSG00000022949 | 46,5289155 | 47,869377 | 31,7711394 | 30,8341576 | 38,166375 | 21,2315075 | 23,8537563 | 41,5556847 | 6,40655811 | 24,8100197 | 46,372759 | 16,64114 | 24,0421551 | 2,806435 | 2,389485 | 19,1424616 | 14,89026 | 29,72554466 | 9,505665 | 12,66647 | 21,25747 | 10,7683497 |
| Downregulated | ENSMUSG00000109293 | 32,78173592 | 25,5303344 | 25,4169115 | 28,9070227 | 36,8941625 | 22,55847672 | 10,2230384 | 35,4445546 | 13,8808759 | 24,8100197 | 32,019286 | 16,64114 | 7,01229525 | 3,741914 | 3,18598 | 7,292366325 | 19,02645 | 15,85362382 | 14,2585 | 3,333281 | 22,1432 | 18,844612 |
| Downregulated | ENSMUSG00000042436 | 38481,52807 | 48262,9697 | 20261,8172 | 65718,189 | 38126,9364 | 16848,52817 | 40993,2482 | 111836,125 | 36229,0861 | 60232,2557 | 47619,3029 | 9537,127 | 17204,1655 | 15972,36 | 12932,69 | 20217,17409 | 6852,003 | 17521,22688 | 12174,86 | 3074,619 | 15571,98 | 40456,6899 |
| Downregulated | ENSMUSG00000022376 | 31,72426057 | 29,7853902 | 6,35422788 | 54,9233432 | 24,1720375 | 15,92363062 | 10,2230384 | 59,8890751 | 21,3551937 | 0 | 28,706946 | 3,503399 | 5,00878232 | 0,935478 | 7,168455 | 1,823091581 | 0 | 0 | 0,950566 | 0 | 0,885728 | 15,2551621 |
| Downregulated | ENSMUSG00000048368 | 715,9108135 | 756,336157 | 673,548155 | 651,371579 | 357,491712 | 217,6229519 | 457,764943 | 979,003043 | 427,103874 | 698,995338 | 431,708304 | 91,96422 | 260,456681 | 83,25758 | 14,33691 | 29,1694653 | 78,5875 | 410,2125164 | 269,9609 | 111,3316 | 357,8341 | 821,984029 |
| Downregulated | ENSMUSG00000042254 | 40308,84548 | 49639,4802 | 18456,3087 | 41622,2584 | 17728,2812 | 11959,97357 | 23096,1156 | 33869,1053 | 12519,4823 | 34634,7875 | 13895,266 | 10559,24 | 7796,67056 | 7794,406 | 5739,543 | 7185,715468 | 13452,53 | 12450,04896 | 14823,13 | 3609,277 | 15258,44 | 19634,291 |
| Downregulated | ENSMUSG00000037138 | 442,0246973 | 546,774662 | 300,464204 | 638,845202 | 268,436837 | 309,1838279 | 366,89349 | 392,334553 | 244,516968 | 423,927728 | 295,902367 | 64,81288 | 81,1422736 | 91,67688 | 62,9231 | 90,24303327 | 130,7034 | 185,2892284 | 179,6571 | 85,332 | 305,5762 | 318,563679 |
| Downregulated | ENSMUSG00000037251 | 474,8064332 | 343,595751 | 340,405065 | 406,625453 | 255,714712 | 197,7184136 | 266,934892 | 337,334382 | 170,84155 | 488,649519 | 277,132441 | 137,5084 | 149,261713 | 112,2574 | 172,0429 | 113,031678 | 196,0551 | 231,8592484 | 273,7632 | 265,9958 | 330,3765 | 264,721931 |
| Downregulated | ENSMUSG00000035042 | 35,95416198 | 22,3390426 | 27,2324052 | 26,9798879 | 17,810975 | 19,90453828 | 17,0383974 | 23,2222944 | 8,54207749 | 32,3608953 | 9,93701978 | 3,503399 | 9,01580818 | 5,61287 | 3,18598 | 4,557728953 | 8,272368 | 16,84447531 | 15,20906 | 13,99978 | 24,80038 | 18,844612 |
| Downregulated | ENSMUSG00000091373 | 212,5525458 | 204,242675 | 181,549368 | 210,057698 | 8,90548749 | 94,21481452 | 141,986645 | 130,778184 | 81,1497361 | 190,929282 | 100,474311 | 88,46082 | 110,193211 | 49,58035 | 6,37196 | 92,97767064 | 66,17895 | 46,57001997 | 109,3151 | 30,66619 | 3,542912 | 109,478222 |
| Downregulated | ENSMUSG00000067212 | 8998,057773 | 8501,60136 | 8767,92673 | 7265,29838 | 1890,50777 | 5977,99633 | 6205,38432 | 8015,35825 | 5312,10444 | 8951,02363 | 5089,96235 | 1700,024 | 4127,23663 | 1593,12 | 657,1083 | 2359,080506 | 1498,126 | 2323,546741 | 2602,651 | 921,9856 | 607,6094 | 4664,49016 |
| Downregulated | ENSMUSG00000031906 | 97,28773241 | 126,587908 | 146,147241 | 155,134355 | 10,1777 | 46,44392265 | 9,08714527 | 99,0003078 | 13,8808759 | 182,29971 | 39,7480791 | 42,91663 | 30,0526939 | 0 | 4,77897 | 21,87709898 | 9,099605 | 0 | 29,46756 | 8,666531 | 69,08679 | 91,5309727 |
| Downregulated | ENSMUSG00000078670 | 855,49756 | 886,115357 | 1032,10816 | 1491,60237 | 430,007825 | 269,3747514 | 736,058767 | 294,556471 | 402,545401 | 1304,14408 | 469,248156 | 287,2787 | 111,194968 | 198,3214 | 142,5726 | 184,1322497 | 140,6303 | 288,3377832 | 172,0525 | 39,99938 | 86,80135 | 167,806783 |
| Downregulated | ENSMUSG00000028621 | 190,3455634 | 135,09802 | 165,209925 | 179,223541 | 95,4159374 | 83,59906077 | 130,627713 | 95,3336297 | 84,3530152 | 208,188426 | 68,4550251 | 44,66833 | 65,1141702 | 56,1287 | 33,45279 | 25,52328214 | 50,46145 | 68,36875273 | 44,67663 | 73,99885 | 75,28688 | 96,0177851 |
| Downregulated | ENSMUSG00000032852 | 673,6117994 | 524,435619 | 532,847395 | 666,788657 | 398,202512 | 179,1408445 | 533,869784 | 462,001436 | 248,788007 | 1061,43737 | 395,272565 | 373,112 | 48,0843103 | 35,54818 | 109,1198 | 198,7169824 | 141,4575 | 406,2491104 | 255,7024 | 17,33306 | 83,25843 | 327,537304 |
| Downregulated | ENSMUSG00000042961 | 1553,431293 | 2031,78911 | 1763,75211 | 1500,27448 | 898,182024 | 1100,057482 | 1399,42037 | 1195,33705 | 1081,64056 | 2290,07269 | 1123,98735 | 980,0758 | 527,925657 | 233,8696 | 463,5601 | 452,1267122 | 1003,438 | 967,0710531 | 789,9208 | 125,998 | 247,1181 | 956,588401 |
| Downregulated | ENSMUSG00000063698 | 65,56347184 | 86,1648786 | 91,6824308 | 89,6117704 | 58,5217749 | 46,44392265 | 48,8434058 | 58,666849 | 59,7945424 | 56,0922185 | 23,1863795 | 14,88944 | 31,0544504 | 12,16122 | 30,26681 | 11,85009528 | 10,75408 | 17,8353268 | 34,22039 | 21,99966 | 26,57184 | 27,8182368 |
| Downregulated | ENSMUSG00000059325 | 248,5067078 | 275,514859 | 395,777622 | 337,248598 | 232,814887 | 149,9475217 | 216,955593 | 411,890169 | 258,397844 | 215,739302 | 96,0578579 | 26,27549 | 99,17389 | 69,2254 | 99,56187 | 72,01211746 | 69,48789 | 76,29556464 | 86,50155 | 29,99953 | 79,71552 | 139,988546 |
| Downregulated | ENSMUSG00000032652 | 129,011993 | 185,094925 | 152,501469 | 156,097923 | 183,1986 | 126,0620758 | 93,143239 | 157,667157 | 87,5562942 | 166,119262 | 122,556577 | 51,67513 | 27,0474245 | 39,29009 | 82,03898 | 42,84265216 | 71,14237 | 55,48768337 | 57,03399 | 66,66563 | 48,71504 | 92,4283352 |
| Downregulated | ENSMUSG00000030643 | 734,9453699 | 1214,81841 | 933,163751 | 1059,92417 | 1114,45815 | 769,6421468 | 726,971621 | 917,891742 | 829,649276 | 727,041447 | 420,667171 | 228,5968 | 177,310894 | 233,8696 | 489,0479 | 312,6602062 | 408,655 | 568,7487546 | 597,9063 | 317,995 | 357,8341 | 690,071745 |
| Downregulated | ENSMUSG00000028036 | 3480,151384 | 3817,84876 | 3846,12336 | 3591,21579 | 3052,03778 | 2233,289195 | 3055,5526 | 2770,78639 | 1986,03302 | 3112,03943 | 1826,20341 | 691,0454 | 637,117111 | 650,1575 | 2257,267 | 1053,746934 | 2437,867 | 1363,411649 | 1844,099 | 897,986 | 432,2353 | 1390,91184 |
| Downregulated | ENSMUSG00000070348 | 1298,579733 | 1510,54479 | 1673,88517 | 1745,98417 | 1207,32966 | 613,059779 | 1585,70685 | 986,3364 | 675,891881 | 1364,55108 | 665,780325 | 306,5474 | 226,396961 | 192,7085 | 414,1774 | 457,5959869 | 857,0174 | 1118,671331 | 932,5057 | 498,6589 | 755,526 | 1379,24613 |
| Downregulated | ENSMUSG00000035062 | 80,36812678 | 95,7387541 | 120,73033 | 116,591658 | 97,9603624 | 45,11695343 | 103,366277 | 83,1113695 | 77,9464571 | 118,656616 | 82,8084982 | 10,5102 | 20,0351293 | 21,516 | 31,0633 | 15,49627844 | 43,84355 | 68,36875273 | 40,87436 | 70,66556 | 86,80135 | 70,8916357 |
| Downregulated | ENSMUSG00000033544 | 2072,651691 | 1829,67397 | 2868,48001 | 2884,92087 | 2601,67456 | 1208,868958 | 2356,9783 | 1232,00383 | 853,139989 | 2848,83748 | 1354,74703 | 130,5016 | 485,851885 | 162,7732 | 154,52 | 120,3240444 | 118,2949 | 986,8880829 | 373,5726 | 824,6538 | 1040,73 | 1029,27476 |
| Downregulated | ENSMUSG00000019437 | 242,1618557 | 299,981429 | 305,910685 | 514,545004 | 367,669412 | 331,7423047 | 428,231721 | 380,112293 | 269,075441 | 341,946793 | 226,343228 | 96,34347 | 118,207263 | 151,5475 | 123,4567 | 101,1815828 | 200,1913 | 272,4841594 | 209,1246 | 183,3305 | 232,9465 | 318,563679 |
| Downregulated | ENSMUSG00000002908 | 212,5525458 | 202,115147 | 243,276153 | 280,39812 | 234,0871 | 203,0262904 | 257,847747 | 242,000752 | 149,486356 | 234,077142 | 120,348351 | 52,55098 | 92,1615947 | 142,1927 | 62,9231 | 57,42738481 | 180,3376 | 171,4173076 | 117,8702 | 83,99869 | 224,9749 | 178,575133 |
| Downregulated | ENSMUSG00000070284 | 540,369905 | 625,493193 | 641,777016 | 981,875205 | 488,529599 | 566,6158564 | 687,215361 | 562,22397 | 483,695138 | 654,768781 | 447,16589 | 205,8247 | 382,670969 | 217,031 | 277,9767 | 132,1741396 | 390,4558 | 322,0267339 | 340,3028 | 436,6599 | 477,4074 | 467,525851 |
| Downregulated | ENSMUSG00000096210 | 2842,493747 | 3003,00559 | 2928,39131 | 4212,71678 | 1782,36971 | 2047,513504 | 2640,95159 | 2437,11869 | 986,60995 | 2298,70226 | 1119,5709 | 559,668 | 726,273437 | 720,3184 | 976,5028 | 997,231095 | 1460,073 | 1326,750144 | 1828,89 | 1975,303 | 1894,572 | 1669,99157 |
| Downregulated | ENSMUSG00000032606 | 350,0243416 | 403,166531 | 374,899445 | 439,386745 | 290,06445 | 378,1862273 | 368,029383 | 297,000923 | 301,108231 | 240,549322 | 228,551455 | 89,33667 | 117,205506 | 110,3864 | 112,3058 | 91,15457906 | 148,9026 | 206,0971097 | 198,6684 | 277,329 | 299,3761 | 197,419745 |
| Downregulated | ENSMUSG00000079671 | 83,54055283 | 206,370203 | 121,638077 | 198,494889 | 117,04355 | 96,86875296 | 137,443072 | 108,778116 | 90,7595733 | 128,364885 | 107,098991 | 14,88944 | 40,0702586 | 3,741914 | 19,91237 | 23,70019056 | 40,53461 | 78,27726762 | 37,07209 | 53,99916 | 95,65863 | 68,1995483 |
| Downregulated | ENSMUSG00000090256 | 152,2764507 | 94,6749901 | 54,4648104 | 14,4535114 | 63,6106249 | 47,77089187 | 70,4253758 | 134,444862 | 18,1519147 | 73,3513626 | 17,6658129 | 11,38605 | 7,01229525 | 17,77409 | 10,35443 | 9,115457906 | 8,272368 | 13,87192084 | 27,56643 | 17,33306 | 1,771456 | 31,4076867 |
| Downregulated | ENSMUSG00000089694 | 142,7591726 | 117,014033 | 112,560608 | 73,2311242 | 109,410275 | 84,92602999 | 103,366277 | 127,111506 | 40,5748681 | 77,6661487 | 77,2879316 | 18,39284 | 31,0544504 | 17,77409 | 10,35443 | 18,23091581 | 12,40855 | 43,59746551 | 44,67663 | 11,33316 | 24,80038 | 34,9971366 |

| | | | | | | | | | | | | | | | | | | | | | | | |
|---|---|---|---|---|---|---|---|---|---|---|---|---|---|---|---|---|---|---|---|---|---|---|---|
| Downregulated | ENSMUSG00000111080 | 261,196412 | 198,923856 | 220,582482 | 102,138147 | 183,1986 | 140,6587372 | 209,004341 | 233,44517 | 122,792364 | 202,794944 | 108,203104 | 84,08157 | 86,151056 | 63,61253 | 75,66702 | 27,34637372 | 49,63421 | 80,25897059 | 89,35325 | 133,3313 | 74,40115 | 90,6336102 |
| Downregulated | ENSMUSG00000028518 | 343,6794895 | 416,995462 | 205,150786 | 166,697164 | 236,631525 | 201,6993212 | 194,23773 | 212,667328 | 225,297294 | 190,929282 | 257,258401 | 73,57137 | 104,182672 | 43,96748 | 124,2532 | 52,86965586 | 65,35171 | 117,9113272 | 82,69928 | 23,33297 | 50,4865 | 139,091184 |
| Downregulated | ENSMUSG00000046856 | 1270,027898 | 1572,24309 | 362,190989 | 266,908176 | 547,051374 | 534,7685951 | 679,264109 | 684,446572 | 633,181494 | 281,539789 | 644,802172 | 163,7839 | 167,29333 | 75,77375 | 48,58619 | 56,51583902 | 0,827237 | 119,8930301 | 56,08342 | 26,66625 | 96,54435 | 202,80392 |
| Downregulated | ENSMUSG00000097792 | 59,21861973 | 48,933141 | 27,2324052 | 25,052753 | 19,0831875 | 17,25059984 | 17,0383974 | 29,3334245 | 16,0163953 | 15,1017511 | 22,0822662 | 9,634347 | 17,0298599 | 0 | 12,74392 | 8,203912116 | 12,40855 | 9,908514888 | 11,4068 | 5,33325 | 5,314368 | 13,4604372 |
| Downregulated | ENSMUSG00000040795 | 117,3797641 | 118,077797 | 102,575393 | 64,5590174 | 38,166375 | 61,04058406 | 79,5125211 | 50,1112669 | 54,455744 | 88,4531138 | 47,4768723 | 22,77209 | 30,0526939 | 43,96748 | 45,40021 | 14,58473265 | 39,70737 | 32,69809913 | 67,49022 | 64,66566 | 85,91562 | 45,7654863 |
| Downregulated | ENSMUSG00000041609 | 198,8053662 | 159,56459 | 185,180355 | 161,879327 | 78,8771749 | 76,96421468 | 148,802004 | 100,222534 | 68,3366199 | 155,332297 | 77,2879316 | 39,41324 | 30,0526939 | 57,99966 | 43,01073 | 20,96555318 | 70,31513 | 35,6706536 | 73,19362 | 75,99881 | 102,7444 | 31,4076867 |
| Downregulated | ENSMUSG00000053334 | 389,1509297 | 330,830583 | 223,305723 | 238,964721 | 157,75435 | 132,6969219 | 211,276127 | 206,556198 | 129,198922 | 173,670138 | 100,474311 | 75,32307 | 61,1071443 | 70,16088 | 191,9553 | 107,5624033 | 159,6567 | 113,9479212 | 149,2389 | 127,998 | 168,2883 | 119,349209 |
| Downregulated | ENSMUSG00000059708 | 75,08075001 | 104,248866 | 59,0035446 | 81,903231 | 40,7108 | 41,13604578 | 45,4357263 | 63,5557531 | 65,1333408 | 83,0596312 | 27,6028327 | 13,13775 | 35,0614763 | 44,90296 | 24,69134 | 15,49627844 | 66,17895 | 30,71639615 | 40,87436 | 41,99934 | 29,22902 | 63,7127359 |
| Downregulated | ENSMUSG00000037032 | 498,0708909 | 346,787042 | 441,164964 | 365,192054 | 437,6411 | 459,1313496 | 257,847747 | 272,556403 | 371,580371 | 457,36732 | 293,69414 | 84,08157 | 64,1124137 | 174,9345 | 161,6885 | 135,8203228 | 166,2746 | 137,7283569 | 243,345 | 298,662 | 180,6885 | 199,21447 |
| Downregulated | ENSMUSG00000113204 | 119,4947148 | 86,1648786 | 91,6824308 | 74,1946916 | 123,404612 | 118,1002605 | 80,6484142 | 94,1114037 | 75,8109377 | 86,2957207 | 49,6850989 | 21,02039 | 25,0439116 | 34,6127 | 53,36516 | 21,87709898 | 44,67079 | 52,51512891 | 43,72606 | 93,99853 | 41,62922 | 65,5074609 |
| Downregulated | ENSMUSG00000020135 | 117,3797641 | 51,0606688 | 50,833823 | 71,3039894 | 52,1607124 | 119,4272297 | 60,2023374 | 92,8891777 | 52,3202246 | 88,4531138 | 71,7673651 | 17,51699 | 4,00702586 | 4,677392 | 6,37196 | 15,49627844 | 17,37197 | 11,89021787 | 41,82493 | 52,66584 | 14,17165 | 18,844612 |
| Downregulated | ENSMUSG00000019320 | 33,83921127 | 23,4028065 | 24,5091647 | 28,9070227 | 16,5387625 | 29,19332281 | 10,2230384 | 22,0000684 | 13,8808759 | 22,6526267 | 6,62467985 | 9,634347 | 4,00702586 | 0,935478 | 10,35443 | 2,734637372 | 10,75408 | 15,85362382 | 6,653965 | 19,33303 | 8,85728 | 15,2551621 |
| Downregulated | ENSMUSG00000097287 | 60,27609508 | 42,5505574 | 47,2028357 | 34,6884273 | 21,6276125 | 65,02149171 | 32,9409016 | 37,8890067 | 19,2196743 | 52,8561289 | 27,6028327 | 16,64114 | 12,0210776 | 23,38696 | 22,30186 | 3,646183163 | 10,75408 | 39,63405955 | 30,41813 | 49,33256 | 23,02893 | 34,0997741 |
| Downregulated | ENSMUSG00000046312 | 830,1181516 | 718,040655 | 694,426333 | 545,379162 | 601,756512 | 545,3843489 | 637,236062 | 661,224278 | 353,428456 | 585,732204 | 373,190298 | 183,0526 | 246,43209 | 106,6445 | 177,6184 | 103,9162201 | 306,0776 | 190,2434859 | 416,3481 | 451,9929 | 462,35 | 660,458783 |
| Downregulated | ENSMUSG00000073609 | 184,0007113 | 172,329757 | 163,394431 | 97,3203098 | 145,032225 | 120,7541989 | 109,045743 | 161,333835 | 72,6076586 | 177,984924 | 80,6002715 | 35,03399 | 50,0878232 | 32,74174 | 74,87053 | 44,66574374 | 86,85987 | 98,09429739 | 149,2389 | 134,6646 | 84,14416 | 100,504597 |
| Downregulated | ENSMUSG00000059554 | 72,96579931 | 55,3157246 | 54,4648104 | 22,1620508 | 49,6162874 | 70,32936859 | 44,2998332 | 47,6668148 | 40,5748681 | 59,328308 | 44,1645324 | 3,503399 | 21,0368858 | 24,32244 | 23,09835 | 15,49627844 | 21,50816 | 25,76213871 | 38,02266 | 49,99922 | 40,74349 | 37,6892241 |
| Downregulated | ENSMUSG00000021245 | 170,2535317 | 123,396616 | 137,97752 | 88,648203 | 90,3270874 | 153,9284294 | 120,404675 | 105,111438 | 76,8786974 | 122,971402 | 54,1015521 | 18,39284 | 43,075528 | 72,03184 | 62,12661 | 29,1694653 | 69,48789 | 38,64320806 | 116,9197 | 103,9984 | 85,02989 | 66,4048233 |
| Downregulated | ENSMUSG00000020793 | 86,71297889 | 19,1477508 | 19,0626836 | 39,5062644 | 15,26655 | 19,90453828 | 63,6100169 | 30,5556505 | 26,6939921 | 62,5643975 | 32,019286 | 14,88944 | 2,00351293 | 0 | 2,389485 | 0 | 4,136184 | 17,8353268 | 14,2585 | 5,999906 | 20,37174 | 13,4604372 |
| Downregulated | ENSMUSG00000020614 | 839,6354297 | 680,808918 | 587,312205 | 666,788657 | 459,268712 | 404,7256117 | 1135,89316 | 363,001128 | 416,426277 | 880,216351 | 465,935816 | 218,9624 | 151,265226 | 254,4501 | 117,0848 | 186,8668871 | 258,9251 | 360,6699419 | 334,5994 | 432,6599 | 583,6948 | 302,411155 |
| Downregulated | ENSMUSG00000046834 | 149,1040247 | 3,1912918 | 5,44648104 | 177,296406 | 0 | 147,2935833 | 1,13589316 | 2,44445204 | 22,4229534 | 4,31478604 | 3,31233993 | 7,882647 | 13,022834 | 14,96765 | 7,168455 | 17,31937002 | 5,790658 | 0,990851489 | 6,653965 | 0 | 4,42864 | 1,79472495 |
| Downregulated | ENSMUSG00000033765 | 72,96579931 | 0 | 0 | 77,0853939 | 0 | 79,61815312 | 0 | 0 | 37,371589 | 0 | 0 | 0 | 0 | 0 | 0 | 0 | 0 | 0 | 0 | 0 | 10,62874 | 0 |
| Downregulated | ENSMUSG00000067594 | 39,12658804 | 0 | 0 | 132,008737 | 0 | 82,27209155 | 0 | 0 | 41,6426277 | 0 | 0 | 0 | 0 | 0 | 0 | 0 | 0 | 0 | 0 | 0 | 5,314368 | 0 |
| Downregulated | ENSMUSG00000057454 | 53,93124297 | 6,3825836 | 0 | 84,7939333 | 0 | 38,48210734 | 0 | 9,77780817 | 30,9650309 | 2,15739302 | 5,52056654 | 0 | 4,00702586 | 0 | 3,18598 | 4,557728953 | 0 | 0 | 0,950566 | 0 | 41,62922 | 0 |
| Downregulated | ENSMUSG00000061527 | 22,2069824 | 1,06376393 | 0 | 119,482361 | 0 | 65,02149171 | 1,13589316 | 28,1111985 | 54,455744 | 0 | 0 | 5,255098 | 11,0193211 | 0,935478 | 1,59299 | 5,469274744 | 26,47158 | 0 | 0,950566 | 0 | 61,11523 | 0,89736248 |
| Downregulated | ENSMUSG00000020682 | 24,3219331 | 9,57387541 | 26,3246584 | 33,7248598 | 3,8166375 | 19,90453828 | 2,27178632 | 9,77780817 | 6,40655811 | 7,55087556 | 5,52056654 | 12,2619 | 10,0175646 | 1,870957 | 7,168455 | 9,115457906 | 9,099605 | 5,945108933 | 8,555098 | 0 | 9,743008 | 17,9472495 |
| Downregulated | ENSMUSG00000051705 | 26,43688381 | 24,4665705 | 23,6014178 | 24,0891856 | 10,1777 | 35,8281689 | 5,67946579 | 20,7778424 | 6,40655811 | 7,55087556 | 1,10411331 | 13,13775 | 6,01053879 | 5,61287 | 0,796495 | 0 | 7,445132 | 6,935960422 | 22,8136 | 31,9995 | 15,05738 | 2,69208743 |
| Downregulated | ENSMUSG00000034308 | 43,35648945 | 58,5070164 | 38,1253673 | 54,9233432 | 31,8053125 | 39,80907656 | 38,6203674 | 36,6667807 | 24,5584728 | 11,8656616 | 22,0822662 | 7,006798 | 21,0368858 | 8,419306 | 7,964949 | 0,911545791 | 5,790658 | 4,954257444 | 25,6653 | 47,99925 | 25,68611 | 39,483949 |
| Downregulated | ENSMUSG00000054418 | 42,29901409 | 34,0404459 | 33,5866331 | 44,3241015 | 16,5387625 | 41,13604578 | 23,8537563 | 40,3334587 | 22,4229534 | 12,9443581 | 29,8110593 | 6,130948 | 14,0245905 | 26,19339 | 5,575465 | 0 | 13,23579 | 8,917663399 | 18,06076 | 27,33291 | 36,31485 | 25,1261494 |
| Downregulated | ENSMUSG00000022075 | 92,00035565 | 38,2955016 | 85,328203 | 81,903231 | 61,0661999 | 65,02149171 | 52,2510853 | 42,7779108 | 67,2688602 | 21,5739302 | 44,1645324 | 20,14454 | 33,0579633 | 28,06435 | 27,08083 | 21,87709898 | 52,11592 | 31,70724764 | 19,01133 | 101,3318 | 36,31485 | 45,7654863 |
| Downregulated | ENSMUSG00000069633 | 76,13822537 | 36,1679738 | 49,0183294 | 50,105506 | 52,1607124 | 66,34846093 | 51,1151921 | 17,1111643 | 34,1683099 | 44,2265569 | 44,1645324 | 14,0136 | 21,0368858 | 14,03218 | 11,15093 | 12,76164107 | 35,57118 | 27,74384169 | 41,82493 | 60,66572 | 39,85776 | 39,483949 |
| Downregulated | ENSMUSG00000039063 | 42,29901409 | 23,4028065 | 24,5091647 | 49,1419386 | 24,1720375 | 62,36755328 | 30,6691153 | 24,4445204 | 43,7781471 | 8,62957207 | 38,6439658 | 7,882647 | 18,0316164 | 2,806435 | 1,59299 | 1,823091581 | 19,85368 | 25,76213871 | 32,31926 | 51,99919 | 24,80038 | 36,7918616 |
| Downregulated | ENSMUSG00000020513 | 132,184419 | 151,054479 | 88,0514435 | 119,482361 | 101,777 | 58,38664562 | 68,1535895 | 91,6669516 | 60,8623021 | 45,3052534 | 54,1015521 | 32,40644 | 37,0649892 | 70,16088 | 44,60372 | 28,25791951 | 61,21553 | 87,19493102 | 52,28116 | 81,33206 | 90,34426 | 63,7127359 |
| Downregulated | ENSMUSG00000056367 | 92,00035565 | 67,0171278 | 69,8965067 | 79,0125288 | 43,255225 | 57,0596764 | 22,7178632 | 70,8891093 | 28,8295115 | 40,9904673 | 14,353473 | 18,39284 | 27,0474245 | 25,25792 | 23,09835 | 21,87709898 | 27,29882 | 41,61576253 | 37,07209 | 48,66591 | 69,97251 | 50,2522987 |
| Downregulated | ENSMUSG00000024592 | 99,40268312 | 79,782295 | 80,7894688 | 68,4132871 | 54,7051374 | 72,98330702 | 46,5716195 | 91,6669516 | 36,3038293 | 30,2035023 | 25,3946061 | 7,006798 | 19,0333728 | 43,96748 | 39,82475 | 17,31937002 | 28,12605 | 35,6706536 | 38,02266 | 69,33225 | 97,43008 | 59,2259235 |
| Downregulated | ENSMUSG00000049086 | 52,87376762 | 43,6143213 | 39,0331141 | 55,8869106 | 25,44425 | 39,80907656 | 26,1255426 | 42,7779108 | 45,9136665 | 23,7313232 | 8,83290647 | 5,255098 | 6,01053879 | 21,516 | 33,45279 | 14,58473265 | 20,68092 | 27,74384169 | 33,26983 | 37,99941 | 33,65766 | 17,0498871 |
| Downregulated | ENSMUSG00000051098 | 47,58639085 | 28,7216262 | 29,9556457 | 46,2512364 | 29,2608875 | 37,15513812 | 46,5716195 | 39,1112327 | 18,1519147 | 25,8887162 | 19,8740396 | 11,38605 | 5,00878232 | 6,548349 | 18,31938 | 23,70019056 | 23,16263 | 16,84447531 | 35,17096 | 27,33291 | 28,3433 | 36,7918616 |
| Downregulated | ENSMUSG00000041849 | 47,58639085 | 12,7651672 | 19,0626836 | 40,4698318 | 12,722125 | 25,21241515 | 13,6307179 | 23,2222944 | 1,06775969 | 4,31478604 | 2,20822662 | 10,5102 | 12,0210776 | 4,677392 | 7,964949 | 5,469274744 | 0 | 5,945108933 | 19,9619 | 18,66638 | 4,42864 | 17,9472495 |
| Downregulated | ENSMUSG00000107741 | 51,81629226 | 18,0839869 | 21,7859242 | 32,7612924 | 5,08884999 | 37,15513812 | 18,1742905 | 12,2222602 | 29,8972712 | 15,1017511 | 13,2493597 | 7,882647 | 5,00878232 | 10,29026 | 9,557939 | 4,557728953 | 13,23579 | 12,88106935 | 12,35736 | 23,99963 | 25,68611 | 17,0498871 |
| Downregulated | ENSMUSG00000019761 | 501,243317 | 127,651672 | 286,848001 | 490,455819 | 108,138062 | 359,6086582 | 155,617363 | 125,88928 | 188,993464 | 160,72578 | 141,326504 | 28,02719 | 72,1264654 | 79,51566 | 56,55114 | 62,89665955 | 37,22566 | 102,0577033 | 84,60042 | 80,66541 | 118,6876 | 77,1731731 |
| Downregulated | ENSMUSG00000030030 | 40,18406339 | 5,31881967 | 10,8929621 | 31,797725 | 22,899825 | 25,21241515 | 10,2230384 | 6,11113011 | 14,9486356 | 23,7313232 | 2,20822662 | 0,87585 | 0 | 20,58052 | 2,389485 | 0 | 3,308947 | 5,945108933 | 3,802266 | 4,666594 | 18,60029 | 15,2551621 |
| Downregulated | ENSMUSG00000086541 | 98,34520777 | 109,567685 | 113,468355 | 90,5753379 | 57,2495624 | 112,7923836 | 138,578965 | 86,7780475 | 100,36941 | 104,633561 | 109,307218 | 28,90304 | 81,1422736 | 14,03218 | 15,1334 | 15,49627844 | 17,37197 | 28,73469318 | 28,51699 | 47,33259 | 131,0877 | 111,272947 |
| Downregulated | ENSMUSG00000029413 | 911,5437537 | 848,883619 | 1227,27373 | 824,813715 | 763,327499 | 893,0502841 | 1146,1162 | 1192,8926 | 1017,57498 | 969,748162 | 766,254636 | 282,0236 | 396,69556 | 332,0948 | 202,3097 | 333,6257594 | 307,7321 | 392,3771896 | 469,5798 | 786,6544 | 1426,022 | 652,382521 |
| Downregulated | ENSMUSG00000026401 | 86,71297889 | 89,3561705 | 127,992304 | 76,1218265 | 91,5992999 | 116,7732912 | 137,443072 | 101,44476 | 136,67324 | 147,781422 | 100,474311 | 19,26869 | 16,0281034 | 20,58052 | 12,74392 | 23,70019056 | 11,58132 | 21,79873275 | 39,92379 | 55,99913 | 129,3163 | 72,6863607 |
| Downregulated | ENSMUSG00000026399 | 2840,378796 | 2658,34607 | 4299,08903 | 1964,71398 | 2859,9337 | 3268,325185 | 4543,57263 | 3779,12286 | 4024,38626 | 4081,78759 | 2977,79359 | 742,7205 | 912,600139 | 791,4147 | 589,4063 | 622,585775 | 310,2138 | 1017,604479 | 892,5819 | 1932,637 | 4660,701 | 2118,67281 |
| Downregulated | ENSMUSG00000031613 | 2570,722581 | 1928,60401 | 5389,29299 | 1759,47412 | 2733,98466 | 3407,656953 | 3990,39266 | 3369,67714 | 2672,60249 | 3380,63486 | 3773,85929 | 390,629 | 1239,17275 | 636,1253 | 518,5182 | 257,9674588 | 513,7141 | 1488,258936 | 2305,124 | 1579,975 | 4145,207 | 2242,50883 |
| Downregulated | ENSMUSG00000020895 | 414,5303381 | 270,196039 | 442,980458 | 398,916914 | 348,586225 | 493,6325493 | 327,13723 | 529,223867 | 463,407704 | 336,553311 | 355,524485 | 69,19213 | 83,1457866 | 74,83827 | 62,9231 | 74,74675483 | 77,76026 | 179,3441195 | 102,6612 | 189,997 | 329,4908 | 165,114696 |
| Downregulated | ENSMUSG00000021643 | 287,6332958 | 336,149403 | 446,611445 | 406,625453 | 395,658087 | 411,3604578 | 297,604007 | 559,779518 | 490,101696 | 263,201948 | 399,689018 | 49,04758 | 214,375883 | 164,6442 | 109,9163 | 68,3659343 | 81,06921 | 216,996476 | 153,0412 | 205,9968 | 426,0352 | 236,903694 |
| Downregulated | ENSMUSG00000086915 | 31,72426057 | 27,6578623 | 59,0035446 | 50,105506 | 36,8941625 | 49,09786109 | 49,979299 | 70,8891093 | 76,8786974 | 56,0922185 | 25,3946061 | 7,882647 | 20,0351293 | 25,25792 | 8,761444 | 3,646183163 | 9,099605 | 20,80788127 | 17,1102 | 48,66591 | 73,51543 | 25,1261494 |
| Downregulated | ENSMUSG00000036782 | 442,0246973 | 586,133928 | 644,500256 | 504,90933 | 695,900237 | 899,6851302 | 480,482806 | 1096,33674 | 1090,18264 | 841,383277 | 723,194217 | 289,0304 | 575,008211 | 173,0635 | 121,0672 | 138,5549602 | 127,3945 | 286,3560803 | 269,9609 | 583,3242 | 1036,302 | 571,619898 |
| Downregulated | ENSMUSG00000085042 | 22,2069824 | 15,956459 | 30,8633926 | 25,052753 | 21,6276125 | 61,04058406 | 36,3485811 | 51,3334929 | 36,3038293 | 15,1017511 | 6,62467985 | 1,751699 | 9,01580818 | 2,806435 | 0,796495 | 8,203912116 | 0 | 22,78958424 | 20,91246 | 33,33281 | 53,14368 | 24,2287869 |
| Downregulated | ENSMUSG00000042918 | 31,72426057 | 17,0202229 | 58,0957978 | 25,052753 | 36,8941625 | 102,1766298 | 72,6971621 | 36,6667807 | 29,8972712 | 33,4395918 | 17,6658129 | 1,751699 | 9,01580818 | 6,548349 | 5,575465 | 5,469274744 | 11,58132 | 31,70724764 | 13,30793 | 23,33297 | 54,02941 | 21,5366995 |
| Downregulated | ENSMUSG00000052676 | 43,35648945 | 38,2955016 | 47,2028357 | 41,4333992 | 31,8053125 | 62,36755328 | 62,4741237 | 39,1112327 | 45,9136665 | 12,9443581 | 48,5809856 | 9,634347 | 14,0245905 | 12,16122 | 11,15093 | 4,557728953 | 3,308947 | 19,81702978 | 32,31926 | 33,33281 | 41,62922 | 26,0235118 |
| Downregulated | ENSMUSG00000072640 | 28,55183451 | 17,0202229 | 47,2028357 | 58,7776129 | 47,0718625 | 50,42483031 | 65,8818032 | 57,444623 | 48,0491859 | 25,8887162 | 27,6028327 | 17,51699 | 25,0439116 | 25,25792 | 17,52289 | 0,911545791 | 9,099605 | 13,87192084 | 26,61586 | 47,33259 | 32,77194 | 25,1261494 |
| Downregulated | ENSMUSG00000021747 | 1,057475352 | 22,3390426 | 10,8929621 | 14,4535114 | 17,810975 | 13,26969219 | 32,9409016 | 20,7778424 | 16,0163953 | 8,62957207 | 25,3946061 | 0,87585 | 4,00702586 | 0 | 1,59299 | 0 | 3,308947 | 18,82617829 | 8,555098 | 10,6665 | 15,9431 | 1,79472495 |
| Downregulated | ENSMUSG00000039798 | 14,80465493 | 31,912918 | 29,9556457 | 33,7248598 | 26,7164625 | 31,84726125 | 47,7075126 | 25,6667465 | 14,9486356 | 28,0461092 | 43,060419 | 12,2619 | 6,01053879 | 4,677392 | 9,557939 | 8,203912116 | 10,75408 | 30,71639615 | 7,604532 | 7,999875 | 18,60029 | 18,844612 |
| Downregulated | ENSMUSG00000072066 | 19,03455634 | 25,5303344 | 55,3725572 | 54,9233432 | 26,7164625 | 27,86635359 | 101,094491 | 63,5557531 | 46,9814262 | 32,3608953 | 36,4357392 | 4,379248 | 8,01405172 | 0 | 16,72639 | 1,823091581 | 12,40855 | 46,57001997 | 29,46756 | 39,99938 | 46,05786 | 40,3813115 |
| Downregulated | ENSMUSG00000043587 | 769,8420565 | 948,877429 | 856,00527 | 1137,00956 | 790,043962 | 805,4703157 | 916,665779 | 982,669721 | 579,793509 | 1052,80779 | 560,889561 | 316,1817 | 260,456681 | 173,0635 | 317,8015 | 296,252382 | 579,0658 | 553,8859822 | 615,9671 | 1035,984 | 651,8958 | 520,470237 |
| Downregulated | ENSMUSG00000054675 | 1355,683402 | 1285,02683 | 2285,70654 | 2965,86053 | 2109,32832 | 1555,207924 | 2645,49517 | 2258,67369 | 1720,16085 | 2636,33427 | 1374,62107 | 360,8501 | 283,497079 | 201,1279 | 332,9349 | 272,5521914 | 728,7957 | 1100,836004 | 1176,801 | 2519,294 | 1251,534 | 1490,51908 |
| Downregulated | ENSMUSG00000020623 | 26,43688381 | 48,933141 | 31,7711394 | 91,5389053 | 49,6162874 | 51,75179953 | 81,7843074 | 79,4446914 | 56,5912633 | 49,6200394 | 36,4357392 | 20,14454 | 20,0351293 | 6,548349 | 16,72639 | 3,646183163 | 54,59763 | 36,66150509 | 48,47889 | 58,66575 | 32,77194 | 55,6364736 |
| Downregulated | ENSMUSG00000038354 | 20,09203169 | 47,869377 | 69,8965067 | 79,0125288 | 96,6881499 | 83,59906077 | 63,6100169 | 88,0002736 | 30,9650309 | 66,8791836 | 33,1233993 | 5,255098 | 15,026347 | 0,935478 | 32,65629 | 1,823091581 | 19,85368 | 25,76213871 | 3,802266 | 49,33256 | 29,22902 | 43,0733989 |
| Downregulated | ENSMUSG00000035561 | 15,86213028 | 25,5303344 | 12,7084558 | 51,0690735 | 26,7164625 | 31,84726125 | 37,4844742 | 50,1112669 | 11,7453565 | 24,8100197 | 6,62467985 | 3,503399 | 24,0421551 | 0 | 11,94742 | 3,646183163 | 4,136184 | 13,87192084 | 12,35736 | 16,66641 | 17,71456 | 9,87098725 |
| Downregulated | ENSMUSG00000021032 | 32,78173592 | 0 | 35,4021268 | 41,4333992 | 20,3554 | 46,44392265 | 69,2894827 | 90,4447256 | 21,3551937 | 84,1383277 | 48,5809856 | 34,15814 | 12,0210776 | 0 | 23,09835 | 16,40782423 | 9,099605 | 22,78958424 | 27,56643 | 13,33313 | 9,743008 | 14,3577996 |
| Downregulated | ENSMUSG00000107182 | 17,97708099 | 6,3825836 | 0,90774684 | 17,3442136 | 6,36106249 | 26,53938437 | 19,3101837 | 31,7778766 | 9,60983717 | 30,2035023 | 6,62467985 | 3,503399 | 6,01053879 | 0 | 8,761444 | 0 | 9,099605 | 5,945108933 | 13,30793 | 4,666594 | 3,542912 | 0,89736248 |
| Downregulated | ENSMUSG00000042846 | 62,39104579 | 77,6547672 | 55,3725572 | 129,118035 | 63,6106249 | 222,9308287 | 181,742905 | 242,000752 | 122,792364 | 182,29971 | 55,2056654 | 21,89624 | 28,049181 | 5,61287 | 11,15093 | 14,58473265 | 40,53461 | 25,76213871 | 58,93512 | 25,99959 | 188,6601 | 115,75976 |
| Downregulated | ENSMUSG00000076617 | 1,057475352 | 11,7014033 | 11,8007089 | 15,4170788 | 7,63327499 | 120,7541989 | 47,7075126 | 183,333903 | 45,9136665 | 106,790954 | 38,6439658 | 5,255098 | 0 | 0 | 0 | 7,292366325 | 0 | 2,972554466 | 13,30793 | 1,333313 | 36,31485 | 54,7391111 |
| Downregulated | ENSMUSG00000027489 | 15,86213028 | 31,912918 | 23,6014178 | 32,7612924 | 40,7108 | 106,1575375 | 88,5996663 | 111,222568 | 46,9814262 | 92,7678998 | 47,4768723 | 8,758497 | 23,0403987 | 0,935478 | 11,94742 | 23,70019056 | 28,12605 | 24,77128722 | 29,46756 | 7,333219 | 29,22902 | 64,6100984 |
| Downregulated | ENSMUSG00000005045 | 38,06911268 | 82,9735868 | 75,3429877 | 95,393175 | 249,35365 | 256,1050592 | 330,544909 | 237,111848 | 164,434992 | 226,526267 | 128,077144 | 56,05438 | 19,0333728 | 12,16122 | 20,70887 | 4,557728953 | 62,04276 | 43,59746551 | 40,87436 | 4,666594 | 9,743008 | 52,9443862 |
| Downregulated | ENSMUSG00000026726 | 29,60930986 | 118,077797 | 53,5570636 | 55,8869106 | 198,46515 | 330,4153354 | 257,847747 | 266,445273 | 133,469961 | 289,090664 | 92,7455179 | 31,53059 | 2,00351293 | 10,29026 | 19,11588 | 81,12757537 | 53,77039 | 67,37790124 | 51,33059 | 53,3325 | 29,22902 | 79,8652605 |
| Downregulated | ENSMUSG00000049409 | 28,55183451 | 25,5303344 | 29,0478989 | 13,4899439 | 61,0661999 | 79,61815312 | 79,5125211 | 92,8891777 | 38,4393487 | 65,8004871 | 5,52056654 | 19,26869 | 14,0245905 | 28,06435 | 23,89485 | 17,31937002 | 32,26224 | 14,86277233 | 18,06076 | 35,99944 | 23,02893 | 27,8182368 |
| Downregulated | ENSMUSG00000112358 | 11,63222888 | 1,06376393 | 19,0626836 | 5,78140454 | 11,4499125 | 38,48210734 | 19,3101837 | 14,6667123 | 16,0163953 | 23,7313232 | 25,3946061 | 0 | 1,00175646 | 0 | 4,77897 | 9,115457906 | 9,099605 | 8,917663399 | 13,30793 | 5,999906 | 1,771456 | 7,17889982 |
| Downregulated | ENSMUSG00000085794 | 9,517278171 | 4,25505574 | 7,26197472 | 1,92713485 | 2,544425 | 23,88544594 | 20,4460768 | 6,11113011 | 12,8131162 | 20,4952337 | 17,6658129 | 2,627549 | 6,01053879 | 0 | 0,796495 | 3,646183163 | 0 | 14,86277233 | 3,802266 | 0 | 2,657184 | 3,58944991 |
| Downregulated | ENSMUSG00000085971 | 11,63222888 | 20,2115147 | 19,0626836 | 39,5062644 | 31,8053125 | 79,61815312 | 63,6100169 | 26,8889725 | 23,4907131 | 26,9674127 | 20,9781529 | 10,5102 | 18,0316164 | 1,870957 | 11,15093 | 2,734637372 | 13,23579 | 17,8353268 | 20,91246 | 4,666594 | 14,17165 | 18,844612 |
| Downregulated | ENSMUSG00000030256 | 33,83921127 | 49,9969049 | 67,1732662 | 109,846686 | 128,493462 | 266,7208129 | 212,412021 | 105,111438 | 57,659023 | 116,499223 | 51,8933255 | 44,66833 | 35,0614763 | 19,64505 | 50,17918 | 24,61173635 | 11,58132 | 120,8838816 | 18,06076 | 11,33316 | 22,1432 | 42,1760364 |
| Downregulated | ENSMUSG00000050671 | 68,7358979 | 65,9533639 | 43,5718483 | 47,2148038 | 122,1324 | 272,0286898 | 326,001336 | 102,666986 | 92,8950927 | 139,15185 | 33,1233993 | 7,006798 | 7,01229525 | 8,419306 | 4,77897 | 9,115457906 | 9,926842 | 2,972554466 | 21,86303 | 3,999938 | 2,657184 | 38,5865865 |
| Downregulated | ENSMUSG00000055312 | 0 | 5,31881967 | 4,5387342 | 10,5992417 | 11,4499125 | 39,80907656 | 38,6203674 | 18,3333903 | 4,27103874 | 8,62957207 | 12,1452464 | 1,751699 | 2,00351293 | 1,870957 | 7,168455 | 2,734637372 | 0 | 1,981702978 | 0 | 4,666594 | 8,85728 | 0,89736248 |
| Downregulated | ENSMUSG00002075232 | 29,60930986 | 10,6376393 | 17,24719 | 35,6519947 | 24,1720375 | 76,96421468 | 37,4844742 | 24,4445204 | 17,084155 | 30,2035023 | 22,0822662 | 11,38605 | 10,0175646 | 1,870957 | 5,575465 | 4,557728953 | 22,33539 | 22,78958424 | 6,653965 | 2,666625 | 4,42864 | 17,9472495 |
| Downregulated | ENSMUSG00000003872 | 9,517278171 | 2,12752787 | 5,44648104 | 17,3442136 | 30,5331 | 61,04058406 | 31,8050084 | 20,7778424 | 13,8808759 | 46,3839499 | 16,5616996 | 0,87585 | 3,00526939 | 0 | 0 | 2,734637372 | 6,617895 | 11,89021787 | 13,30793 | 0 | 0,885728 | 9,87098725 |
| Downregulated | ENSMUSG00000035513 | 28,55183451 | 30,8491541 | 17,24719 | 29,8705901 | 53,4329249 | 110,1384451 | 61,3382305 | 42,7779108 | 18,1519147 | 51,7774324 | 64,0385719 | 18,39284 | 28,049181 | 39,29009 | 6,37196 | 17,31937002 | 16,54474 | 28,73469318 | 15,20906 | 16,66641 | 15,05738 | 23,3314244 |
| Downregulated | ENSMUSG00000085925 | 63,44852114 | 18,0839869 | 24,5091647 | 19,2713485 | 31,8053125 | 140,6587372 | 93,143239 | 69,6668832 | 33,1005503 | 55,013522 | 48,5809856 | 13,13775 | 16,0281034 | 24,32244 | 15,1334 | 17,31937002 | 1,654474 | 30,71639615 | 4,752832 | 19,33303 | 7,971552 | 24,2287869 |
| Downregulated | ENSMUSG00000098202 | 224,1847747 | 81,9098229 | 125,269064 | 169,587867 | 223,9094 | 690,0239937 | 533,869784 | 242,000752 | 169,77379 | 369,992903 | 391,960225 | 81,45402 | 30,0526939 | 32,74174 | 73,27754 | 30,99255688 | 47,97974 | 122,8655846 | 73,19362 | 51,33253 | 42,51494 | 126,528109 |
| Downregulated | ENSMUSG00000102657 | 591,1287219 | 84,0373508 | 332,235343 | 290,033795 | 370,213837 | 1027,074175 | 791,717531 | 504,779347 | 294,701673 | 543,663041 | 694,487271 | 174,2941 | 121,212532 | 97,28975 | 136,2006 | 52,86965586 | 27,29882 | 286,3560803 | 93,15552 | 113,9982 | 82,37271 | 173,190958 |
| Downregulated | ENSMUSG00000048782 | 21,14950705 | 15,956459 | 26,3246584 | 57,8140454 | 50,8884999 | 42,463015 | 45,4357263 | 35,4445546 | 26,6939921 | 72,2726661 | 3,31233993 | 1,751699 | 0 | 14,03218 | 10,35443 | 0 | 6,617895 | 50,53342593 | 23,76416 | 3,333281 | 4,42864 | 51,1496612 |
| Downregulated | ENSMUSG00000030228 | 12,68970423 | 30,8491541 | 27,2324052 | 13,4899439 | 39,4385875 | 43,78998421 | 20,4460768 | 33,0001026 | 8,54207749 | 45,3052534 | 15,4575863 | 0 | 2,00351293 | 2,806435 | 0 | 0 | 0 | 6,935960422 | 6,653965 | 3,999938 | 7,971552 | 29,6129618 |
| Downregulated | ENSMUSG00000030701 | 20,09203169 | 15,956459 | 16,3394431 | 25,052753 | 35,62195 | 39,80907656 | 17,0383974 | 36,6667807 | 4,27103874 | 31,2821988 | 12,1452464 | 6,130948 | 6,01053879 | 0 | 5,575465 | 1,823091581 | 0,827237 | 7,92681191 | 18,06076 | 0 | 5,314368 | 31,4076867 |
| Downregulated | ENSMUSG00000085915 | 6,344852114 | 26,5940983 | 21,7859242 | 123,33663 | 72,5161124 | 91,56087608 | 19,3101837 | 92,8891777 | 50,1847052 | 58,2496115 | 17,6658129 | 5,255098 | 2,00351293 | 2,806435 | 7,964949 | 4,557728953 | 8,272368 | 40,62491104 | 19,9619 | 0 | 7,085824 | 40,3813115 |
| Downregulated | ENSMUSG00000022523 | 6,344852114 | 94,6749901 | 44,4795952 | 270,762446 | 118,315762 | 207,0071981 | 107,90985 | 118,555924 | 91,827333 | 217,896695 | 90,5372913 | 8,758497 | 43,075528 | 11,22574 | 29,47031 | 31,90410267 | 43,01632 | 67,37790124 | 13,30793 | 9,999844 | 38,97203 | 79,8652605 |
| Downregulated | ENSMUSG00000033227 | 6,344852114 | 9,57387541 | 9,0774684 | 32,7612924 | 16,5387625 | 37,15513812 | 14,7666111 | 18,3333903 | 6,40655811 | 25,8887162 | 7,72879316 | 14,0136 | 8,01405172 | 6,548349 | 4,77897 | 2,734637372 | 3,308947 | 5,945108933 | 2,851699 | 1,333313 | 1,771456 | 0 |

| | | | | | | | | | | | | | | | | | | | | | | | |
|---|---|---|---|---|---|---|---|---|---|---|---|---|---|---|---|---|---|---|---|---|---|---|---|
| Downregulated | ENSMUSG00000043702 | 2205,893585 | 1886,05345 | 2337,44811 | 1631,31965 | 2101,69505 | 3354,578185 | 2334,26044 | 2280,67376 | 1930,50951 | 1959,99156 | 1751,12371 | 915,2629 | 843,478943 | 514,5131 | 624,452 | 582,4777602 | 313,5228 | 912,5742212 | 617,8682 | 385,3273 | 525,2367 | 919,796539 |
| Downregulated | ENSMUSG00000010362 | 360,5990951 | 384,01878 | 344,943799 | 332,430761 | 297,697725 | 488,3246725 | 441,862439 | 349,556642 | 334,208782 | 259,965859 | 220,822662 | 63,93703 | 96,1686206 | 41,16105 | 58,14413 | 92,06612485 | 29,78053 | 127,8198421 | 46,57776 | 69,33225 | 138,1736 | 123,836022 |
| Downregulated | ENSMUSG00000107369 | 927,405884 | 1011,6395 | 878,698941 | 768,926804 | 1202,24081 | 1422,511002 | 970,052757 | 794,446914 | 845,665671 | 790,684541 | 745,276483 | 399,3875 | 547,960786 | 350,8044 | 387,0965 | 319,0410267 | 411,1367 | 445,88317 | 374,5232 | 195,9969 | 565,0945 | 477,396838 |
| Downregulated | ENSMUSG00000058135 | 4359,970878 | 4563,54728 | 4068,52134 | 3913,04731 | 5487,05251 | 6628,211247 | 4963,8531 | 4611,45878 | 4330,83329 | 4368,72086 | 3488,99806 | 2029,344 | 2716,76353 | 1681,055 | 2023,097 | 1578,797309 | 1912,572 | 2285,894385 | 1751,894 | 1017,984 | 2967,189 | 2675,03755 |
| Downregulated | ENSMUSG00000053117 | 282,3459191 | 330,830583 | 478,382585 | 220,65694 | 365,124987 | 594,4822099 | 312,370619 | 262,778595 | 255,194565 | 273,988913 | 358,836825 | 71,81967 | 87,1528124 | 79,51566 | 124,2532 | 70,18902588 | 48,80697 | 74,31386166 | 43,72606 | 9,999844 | 77,05834 | 85,2494354 |
| Downregulated | ENSMUSG00000034949 | 89,88540495 | 56,3794885 | 87,1436966 | 59,7411803 | 77,6609624 | 114,1193528 | 59,0664442 | 53,777945 | 46,9814262 | 78,7448452 | 71,7673651 | 8,758497 | 32,0562069 | 2,806435 | 6,37196 | 34,63874004 | 10,75408 | 12,88106935 | 18,06076 | 10,6665 | 19,48602 | 24,2287869 |
| Downregulated | ENSMUSG00000037321 | 2058,904511 | 1565,86051 | 1804,60072 | 1904,9728 | 1779,82529 | 2845,022005 | 1591,38631 | 1659,78294 | 1652,89199 | 2599,65859 | 2021,63147 | 609,5914 | 577,011724 | 540,7065 | 1055,356 | 814,010391 | 938,9138 | 889,784637 | 581,7467 | 317,3284 | 754,6403 | 835,444466 |
| Downregulated | ENSMUSG00000070867 | 244,2768064 | 212,752787 | 413,932559 | 240,891856 | 287,520025 | 390,1289503 | 386,203674 | 202,88952 | 349,157417 | 333,317221 | 309,151726 | 145,391 | 91,1598383 | 193,644 | 152,1305 | 133,0856854 | 73,62408 | 189,2526344 | 104,5623 | 49,33256 | 108,0588 | 134,604372 |
| Downregulated | ENSMUSG00000025738 | 170,2535317 | 125,524144 | 181,549368 | 167,660732 | 236,631525 | 248,1432439 | 210,140234 | 125,88928 | 154,825154 | 204,952337 | 156,78409 | 126,1224 | 61,1071443 | 123,4831 | 74,87053 | 137,6434144 | 56,25211 | 104,0394063 | 110,2657 | 32,66616 | 89,45853 | 46,6628488 |
| Downregulated | ENSMUSG00000022330 | 13205,7522 | 13825,7398 | 12121,1436 | 12141,9131 | 18809,6618 | 24187,99492 | 18778,5857 | 12382,3718 | 9240,39232 | 14597,9999 | 17817,0765 | 9268,241 | 4814,44157 | 6239,641 | 5168,456 | 6542,164139 | 6560,815 | 7972,391079 | 6585,525 | 3675,943 | 4689,044 | 7900,37925 |
| Downregulated | ENSMUSG00000016356 | 154,3914014 | 95,7387541 | 158,855697 | 129,118035 | 180,654175 | 289,2792897 | 232,858097 | 122,222602 | 81,1497361 | 156,410994 | 158,992316 | 84,08157 | 75,1317348 | 68,28992 | 44,60372 | 65,63129693 | 27,29882 | 80,25897059 | 40,87436 | 7,333219 | 12,40019 | 43,9707614 |
| Downregulated | ENSMUSG00000026628 | 21426,56559 | 17731,881 | 22932,4084 | 25077,8058 | 32080,1104 | 48293,71774 | 33656,5143 | 19416,2826 | 16901,5681 | 26446,4023 | 20577,3597 | 9472,314 | 10698,759 | 11085,42 | 10711,26 | 7183,892376 | 5878,345 | 7675,135632 | 8283,236 | 6565,231 | 10798,8 | 18624,7582 |
| Downregulated | ENSMUSG00000058297 | 200,9203169 | 202,115147 | 240,552913 | 263,053907 | 228,99825 | 416,6683346 | 355,534559 | 267,667499 | 242,381449 | 234,077142 | 260,570741 | 65,68873 | 86,151056 | 98,22523 | 91,59692 | 118,5009528 | 24,81711 | 77,28641613 | 87,45212 | 62,66569 | 80,60125 | 187,548758 |
| Downregulated | ENSMUSG00000041444 | 1075,452433 | 722,295711 | 898,669372 | 570,431915 | 1277,30135 | 1835,198429 | 1739,05243 | 1095,11452 | 1028,25258 | 1622,35955 | 1229,98223 | 419,532 | 511,897553 | 471,4811 | 437,2757 | 542,3697454 | 153,0388 | 388,4137836 | 346,9568 | 120,6648 | 414,5207 | 567,133086 |
| Downregulated | ENSMUSG00000062591 | 964,4175213 | 643,57718 | 905,023599 | 868,174249 | 2181,84444 | 2239,924041 | 2412,63707 | 851,891537 | 1639,01112 | 2038,7364 | 1752,22782 | 465,0762 | 294,516401 | 454,6425 | 556,75 | 575,1853939 | 172,0653 | 558,8402397 | 361,2153 | 101,9984 | 332,148 | 480,986288 |
| Downregulated | ENSMUSG00000026121 | 869,2447396 | 543,58337 | 677,179143 | 399,880481 | 1352,36189 | 1762,215122 | 1580,02738 | 832,335921 | 800,819764 | 878,058958 | 1252,06449 | 398,5116 | 276,484784 | 65,48349 | 291,5172 | 260,7020961 | 156,3478 | 358,6882389 | 189,1627 | 217,9966 | 90,34426 | 354,458179 |
| Downregulated | ENSMUSG00000110206 | 287,6332958 | 165,947174 | 212,412761 | 175,369271 | 414,741275 | 748,4106393 | 387,339567 | 361,778902 | 401,477642 | 340,868097 | 409,626038 | 67,44043 | 78,1370042 | 41,16105 | 54,95815 | 58,3389306 | 27,29882 | 178,353268 | 73,19362 | 54,66581 | 54,02941 | 82,5573479 |
| Downregulated | ENSMUSG00000034266 | 171,3110071 | 197,860092 | 297,740964 | 246,673261 | 502,523937 | 658,1767324 | 412,329216 | 314,112088 | 433,510432 | 419,612942 | 374,294412 | 136,6326 | 129,226584 | 153,4185 | 202,3097 | 118,5009528 | 111,677 | 247,7128722 | 88,40268 | 21,333 | 86,80135 | 144,475359 |
| Downregulated | ENSMUSG00000037379 | 5640,573529 | 2912,58565 | 8841,45422 | 4313,89136 | 13157,2217 | 16137,27267 | 14024,8728 | 10105,3647 | 9253,20544 | 12712,4384 | 4795,1641 | 234,7277 | 1403,46081 | 523,8679 | 105,9338 | 151,3166012 | 82,72368 | 932,391251 | 903,9887 | 783,3211 | 3091,191 | 3052,82715 |
| Downregulated | ENSMUSG00000042404 | 142,7591726 | 157,437062 | 196,981064 | 183,077811 | 246,809225 | 301,2220126 | 315,778298 | 169,889417 | 170,84155 | 220,054088 | 143,53473 | 88,46082 | 33,0579633 | 88,87045 | 82,83547 | 68,3659343 | 85,20539 | 82,24067357 | 87,45212 | 151,331 | 119,5733 | 82,5573479 |
| Downregulated | ENSMUSG00000028807 | 66,6209472 | 62,7620721 | 93,4979245 | 50,105506 | 64,8828374 | 98,19572218 | 106,773957 | 74,5557873 | 48,0491859 | 76,5874522 | 51,8933255 | 14,88944 | 22,0386422 | 9,354784 | 31,8598 | 28,25791951 | 41,36184 | 32,69809913 | 43,72606 | 47,33259 | 40,74349 | 55,6364736 |
| Downregulated | ENSMUSG00000028931 | 561,5194121 | 524,435619 | 948,595448 | 606,08391 | 732,794399 | 740,448824 | 1226,76461 | 823,780339 | 533,879843 | 1037,70604 | 489,122196 | 192,6869 | 95,1668641 | 164,6442 | 360,0157 | 319,9525725 | 489,7242 | 488,489784 | 467,6787 | 45,33263 | 84,14416 | 390,352678 |
| Downregulated | ENSMUSG00000048402 | 398,6682078 | 276,578623 | 718,935497 | 391,208374 | 427,4634 | 627,6564404 | 680,400002 | 504,779347 | 465,543223 | 737,828412 | 403,001358 | 104,2261 | 38,0667457 | 115,9993 | 220,6291 | 311,7486604 | 282,915 | 422,1027342 | 272,8126 | 136,6645 | 185,1172 | 202,80392 |
| Downregulated | ENSMUSG00000046862 | 785,7041868 | 1024,40467 | 1231,81246 | 904,789811 | 1166,61886 | 1324,31528 | 1452,80735 | 1296,78181 | 879,833981 | 1551,16558 | 724,298331 | 787,3889 | 388,681508 | 298,4176 | 736,7578 | 520,4926465 | 239,8987 | 363,6424964 | 334,5994 | 186,6638 | 298,4903 | 356,252904 |
| Downregulated | ENSMUSG00000023034 | 28158,45368 | 26873,8683 | 37464,5276 | 34626,7589 | 47078,2235 | 46086,96793 | 56203,9935 | 35605,8885 | 32788,7644 | 41720,7449 | 29622,256 | 17577,43 | 11262,7479 | 11343,61 | 30723,2 | 19417,74843 | 18393,61 | 19830,9017 | 14602,6 | 10533,17 | 8171,727 | 15379,8955 |
| Downregulated | ENSMUSG00000055022 | 28,55183451 | 307,427777 | 222,397976 | 183,077811 | 315,5087 | 305,2029203 | 310,098832 | 270,111951 | 304,31151 | 343,02549 | 175,554016 | 99,84686 | 44,0772844 | 75,77375 | 133,0147 | 129,4395023 | 28,95329 | 95,12174293 | 115,9691 | 17,99972 | 22,1432 | 98,7098725 |
| Downregulated | ENSMUSG00000084910 | 12,68970423 | 37,2317377 | 36,3098736 | 24,0891856 | 44,5274375 | 47,77089187 | 31,8050084 | 33,0001026 | 25,6262325 | 35,5969848 | 20,9781529 | 11,38605 | 10,0175646 | 3,741914 | 22,30186 | 13,67318686 | 7,445132 | 17,8353268 | 10,45623 | 2,666625 | 9,743008 | 23,3314244 |
| Downregulated | ENSMUSG00000027858 | 97,28773241 | 149,990715 | 131,623292 | 109,846686 | 236,631525 | 419,3222731 | 266,934892 | 204,111746 | 244,516968 | 369,992903 | 231,863795 | 21,02039 | 43,075528 | 38,35461 | 39,82475 | 14,58473265 | 23,16263 | 60,44194082 | 31,36869 | 2,666625 | 20,37174 | 70,8916357 |
| Downregulated | ENSMUSG00000033207 | 476,9213839 | 887,179121 | 819,695396 | 633,063798 | 1281,11799 | 1621,556385 | 1790,16762 | 1134,22575 | 1222,58484 | 2282,52181 | 1468,4707 | 240,8587 | 172,302112 | 159,0313 | 140,9796 | 183,2207039 | 86,85987 | 276,4475654 | 105,5129 | 92,66522 | 168,2883 | 331,126754 |
| Downregulated | ENSMUSG00000000305 | 3102,632684 | 4096,55491 | 3323,26118 | 4556,71035 | 5802,56121 | 7274,445256 | 7716,12222 | 5150,46046 | 5252,30989 | 7129,10523 | 5539,33647 | 1106,198 | 1181,07087 | 887,769 | 1084,03 | 1310,802847 | 731,2774 | 2630,710703 | 1373,569 | 80,66541 | 824,6128 | 2651,70612 |
| Downregulated | ENSMUSG00000024600 | 1059,590303 | 1640,32399 | 1796,431 | 1510,87372 | 2234,00515 | 2262,482518 | 3136,20101 | 2047,22859 | 2070,38603 | 2432,46063 | 2026,04792 | 473,8347 | 209,367101 | 333,0303 | 521,7042 | 525,0503754 | 524,4682 | 734,2209532 | 518,0587 | 28,66622 | 119,5733 | 716,095257 |
| Downregulated | ENSMUSG00000021294 | 217,8399226 | 197,860092 | 322,250128 | 367,119189 | 555,956862 | 663,4846093 | 620,197664 | 451,001402 | 335,276541 | 460,235281 | 357,732712 | 34,15814 | 40,0702586 | 29,93531 | 43,80722 | 77,4813922 | 47,1525 | 122,8655846 | 101,7106 | 8,666531 | 53,14368 | 157,935796 |
| Downregulated | ENSMUSG00000030047 | 198,8053662 | 191,477508 | 227,844457 | 225,474777 | 393,113662 | 387,4750118 | 432,775293 | 239,5563 | 332,073262 | 309,585898 | 226,343228 | 84,08157 | 57,1001185 | 76,70923 | 112,3058 | 105,7393117 | 61,21553 | 105,0302578 | 72,24305 | 0 | 36,31485 | 129,220197 |
| Downregulated | ENSMUSG00000005320 | 2348,652757 | 3129,59349 | 2560,75384 | 3062,21727 | 4901,83476 | 5234,893567 | 4820,73056 | 3615,34457 | 3751,03978 | 4507,87271 | 3018,64579 | 1654,48 | 768,347208 | 1085,155 | 1072,082 | 1366,40714 | 436,7811 | 1947,023176 | 907,791 | 14,66644 | 318,8621 | 1528,2083 |
| Downregulated | ENSMUSG00000018470 | 28,55183451 | 27,6578623 | 40,8486078 | 16,3806462 | 45,79965 | 47,77089187 | 38,6203674 | 26,8889725 | 41,6426277 | 57,170915 | 20,9781529 | 2,627549 | 10,0175646 | 0 | 1,59299 | 18,23091581 | 22,33539 | 20,80788127 | 7,604532 | 8,666531 | 17,71456 | 8,0762623 |
| Downregulated | ENSMUSG00000086746 | 6,344852114 | 11,7014033 | 16,3394431 | 4,81783712 | 29,2608875 | 26,53938437 | 18,1742905 | 14,6667123 | 13,8808759 | 18,3378407 | 13,2493597 | 4,379248 | 0 | 3,741914 | 3,18598 | 9,115457906 | 1,654474 | 13,87192084 | 4,752832 | 0 | 0 | 0 |
| Downregulated | ENSMUSG00000029641 | 78,25317607 | 131,906728 | 200,612052 | 177,296406 | 242,992587 | 291,9332281 | 265,798999 | 182,111677 | 245,584728 | 242,706715 | 131,389484 | 148,8944 | 80,1405172 | 111,3219 | 162,485 | 100,270037 | 62,87 | 71,34130719 | 65,58909 | 162,6641 | 43,40067 | 89,7362477 |
| Downregulated | ENSMUSG00000038793 | 8,459802819 | 4,25505574 | 19,0626836 | 17,3442136 | 39,4385875 | 38,48210734 | 36,3485811 | 34,2223286 | 27,7617518 | 4,31478604 | 8,83290647 | 0 | 0 | 0 | 3,982475 | 1,823091581 | 0 | 8,917663399 | 1,901133 | 0 | 7,085824 | 5,38417486 |
| Downregulated | ENSMUSG00000024232 | 579,4964931 | 822,289521 | 694,426333 | 770,853939 | 1180,6132 | 954,0908682 | 1106,35994 | 1059,66996 | 1081,64056 | 698,995338 | 663,572099 | 627,1084 | 460,807974 | 677,2864 | 464,3566 | 544,192837 | 239,0714 | 416,1576253 | 391,6334 | 409,3269 | 265,7184 | 501,625625 |
| Downregulated | ENSMUSG00000059970 | 1019,40624 | 1161,63022 | 1217,28851 | 1123,51962 | 1861,24689 | 2099,265304 | 2356,9783 | 1921,33931 | 1879,25705 | 1669,8222 | 1252,06449 | 1227,941 | 632,108329 | 769,8987 | 1311,031 | 1087,474128 | 527,7771 | 918,5193301 | 561,7848 | 484,6591 | 412,7493 | 716,992619 |
| Downregulated | ENSMUSG00000040037 | 219,9548733 | 393,592656 | 302,279698 | 356,519947 | 525,423762 | 697,985809 | 483,890485 | 611,113011 | 582,996788 | 430,399907 | 354,420372 | 253,9964 | 225,395205 | 195,515 | 172,8394 | 175,0167918 | 22,33539 | 157,5453867 | 95,05665 | 2,666625 | 58,45805 | 174,088321 |
| Downregulated | ENSMUSG00000007872 | 11123,58323 | 11895,0083 | 10565,2655 | 13127,6426 | 16050,2329 | 17723,00088 | 15824,1276 | 13422,4862 | 14565,3099 | 12153,6736 | 10325,6677 | 6260,574 | 8246,45922 | 7232,183 | 10073,27 | 8974,168309 | 4619,291 | 8861,184864 | 5659,673 | 2293,964 | 4037,148 | 6331,78964 |
| Downregulated | ENSMUSG00000000142 | 60,27609508 | 43,6143213 | 88,0514435 | 125,263765 | 104,321425 | 141,9857064 | 148,802004 | 80,6669174 | 89,6918136 | 81,9809347 | 85,0167248 | 32,40644 | 22,0386422 | 52,32227 | 62,12661 | 46,48883532 | 95,13224 | 74,31386166 | 37,07209 | 5,33325 | 15,05738 | 47,5602113 |
| Downregulated | ENSMUSG00000071847 | 1042,670697 | 1278,64425 | 877,791194 | 2435,89845 | 1192,06311 | 2465,508808 | 1864,00067 | 1470,3379 | 1673,17943 | 2209,17045 | 1778,72654 | 412,5252 | 155,272252 | 1329,315 | 627,638 | 441,1881627 | 400,3826 | 956,1716867 | 326,9949 | 74,6655 | 265,7184 | 605,719672 |
| Downregulated | ENSMUSG00000020866 | 733,8878945 | 534,009495 | 817,879903 | 1049,32492 | 713,711212 | 1499,475217 | 1540,27112 | 1085,33671 | 896,918136 | 1406,62025 | 813,731509 | 312,6783 | 254,446142 | 632,3834 | 543,2096 | 544,192837 | 526,1226 | 725,3032898 | 483,8383 | 254,6627 | 184,2314 | 420,863002 |
| Downregulated | ENSMUSG00000060780 | 1,057475352 | 3,1912918 | 12,7084558 | 33,7248598 | 41,9830125 | 38,48210734 | 36,3485811 | 12,2222602 | 6,40655811 | 11,8656616 | 4,41645324 | 0 | 0 | 0 | 3,18598 | 0 | 0 | 0,990851489 | 0,950566 | 1,333313 | 0 | 13,4604372 |
| Downregulated | ENSMUSG00000023047 | 87,77045424 | 159,56459 | 86,2359498 | 212,948401 | 357,491712 | 248,1432439 | 286,245076 | 172,333869 | 128,131162 | 141,309243 | 175,554016 | 39,41324 | 46,0807974 | 86,99949 | 87,61444 | 120,3240444 | 44,67079 | 103,0485548 | 119,7714 | 8,666531 | 49,60077 | 187,548758 |
| Downregulated | ENSMUSG00000044550 | 16,91960564 | 13,8289311 | 17,24719 | 28,9070227 | 40,7108 | 46,44392265 | 32,9409016 | 28,1111985 | 13,8808759 | 35,5969848 | 49,6850989 | 4,379248 | 5,00878232 | 6,548349 | 6,37196 | 10,0270037 | 9,099605 | 12,88106935 | 4,752832 | 15,33309 | 13,28592 | 11,6657122 |
| Downregulated | ENSMUSG00000041789 | 4,229901409 | 22,3390426 | 13,6162026 | 40,4698318 | 52,1607124 | 50,42483031 | 39,7562605 | 19,5556163 | 24,5584728 | 29,1248057 | 36,4357392 | 7,006798 | 0 | 1,870957 | 1,59299 | 18,23091581 | 6,617895 | 9,908514888 | 0,950566 | 0 | 6,200096 | 10,7683497 |
| Downregulated | ENSMUSG00000081540 | 58,16114438 | 78,7185311 | 47,2028357 | 83,8303659 | 90,3270874 | 118,1002605 | 136,307179 | 66,0002052 | 55,5235037 | 88,4531138 | 87,2249514 | 33,28229 | 29,0509375 | 16,83861 | 50,17918 | 48,3119269 | 45,49803 | 48,55172295 | 25,6653 | 33,99947 | 46,94358 | 43,0733989 |
| Downregulated | ENSMUSG00000026167 | 1,057475352 | 12,7651672 | 8,16972156 | 52,0326409 | 5,08884999 | 10,61575375 | 1,13589316 | 4,88890409 | 14,9486356 | 64,7217906 | 40,8521924 | 3,503399 | 2,00351293 | 0 | 3,982475 | 3,646183163 | 0 | 4,954257444 | 3,802266 | 0 | 6,200096 | 8,0762623 |
| Downregulated | ENSMUSG00000032649 | 25,37940846 | 10,6376393 | 42,6641015 | 46,2512364 | 43,255225 | 9,28878453 | 15,9025042 | 20,7778424 | 7,4743178 | 66,8791836 | 39,7480791 | 12,2619 | 11,0193211 | 0 | 7,964949 | 5,469274744 | 3,308947 | 7,92681191 | 3,802266 | 3,999938 | 4,42864 | 7,17889982 |
| Downregulated | ENSMUSG00000040102 | 102,5751092 | 117,014033 | 122,545823 | 185,968513 | 69,9716874 | 139,331768 | 51,1151921 | 55,000171 | 54,455744 | 159,647083 | 89,433178 | 26,27549 | 37,0649892 | 10,29026 | 72,48104 | 67,45438851 | 20,68092 | 47,56087146 | 99,80948 | 74,6655 | 72,6297 | 79,8652605 |
| Downregulated | ENSMUSG00000044122 | 50,75881691 | 53,1881967 | 56,2803041 | 65,5225848 | 30,5331 | 75,63724546 | 23,8537563 | 35,4445546 | 36,3038293 | 73,3513626 | 46,372759 | 12,2619 | 18,0316164 | 18,70957 | 15,1334 | 14,58473265 | 40,53461 | 33,68895062 | 25,6653 | 29,99953 | 30,11475 | 27,8182368 |
| Downregulated | ENSMUSG00000034438 | 10,57475352 | 13,8289311 | 13,6162026 | 19,2713485 | 12,722125 | 54,40573796 | 7,95125211 | 15,8889383 | 12,8131162 | 34,5182883 | 11,0411331 | 1,751699 | 2,00351293 | 1,870957 | 5,575465 | 4,557728953 | 2,481711 | 9,908514888 | 0 | 0 | 1,771456 | 11,6657122 |
| Downregulated | ENSMUSG00000068606 | 7,402327466 | 12,7651672 | 16,3394431 | 29,8705901 | 53,43291249 | 210,9881058 | 35,2126879 | 77,0002394 | 88,6240539 | 174,748834 | 56,3097787 | 0,87585 | 15,026347 | 4,677392 | 33,45279 | 20,96555318 | 12,40855 | 4,954257444 | 0 | 6,666563 | 14,17165 | 5,38417486 |
| Downregulated | ENSMUSG00000027111 | 19,03455634 | 29,7853902 | 37,2176204 | 65,5225848 | 111,9547 | 157,909337 | 47,7075126 | 86,7780475 | 7,4743178 | 107,869651 | 81,7043849 | 72,69552 | 18,0316164 | 32,74174 | 28,67382 | 23,70019056 | 13,23579 | 41,61576253 | 24,71473 | 3,999938 | 38,0863 | 40,3813115 |
| Downregulated | ENSMUSG00000100779 | 11,63222888 | 22,3390426 | 11,8007089 | 11,5628091 | 29,2608875 | 43,78998421 | 17,0383974 | 34,2223286 | 6,40655811 | 43,1478604 | 20,9781529 | 14,88944 | 10,0175646 | 15,90313 | 15,9299 | 15,49627844 | 2,481711 | 11,89021787 | 0,950566 | 11,33316 | 3,542912 | 12,5630747 |
| Downregulated | ENSMUSG00000042514 | 0 | 64,8896 | 0 | 26,9798879 | 41,9830125 | 83,59906077 | 28,397329 | 0 | 59,7945424 | 72,7226661 | 44,1645324 | 0 | 0 | 4,677392 | 2,389485 | 0 | 0 | 0 | 2,851699 | 0 | 1,771456 | 8,0762623 |
| Downregulated | ENSMUSG00000021763 | 77,19570072 | 336,149403 | 82,6049624 | 141,644411 | 227,726037 | 230,892644 | 210,140234 | 145,444897 | 207,145379 | 527,482593 | 327,921653 | 183,9284 | 58,1018749 | 74,83827 | 23,09835 | 19,1424616 | 8,272368 | 83,23152506 | 42,77549 | 43,99931 | 39,85776 | 62,8153734 |
| Downregulated | ENSMUSG00000043460 | 8,459802819 | 27,6578623 | 17,24719 | 26,0163204 | 24,1720375 | 31,84726125 | 23,8537563 | 18,3333903 | 12,8131162 | 62,5643975 | 27,6028327 | 4,379248 | 0 | 15,90313 | 10,35443 | 1,823091581 | 0 | 3,963405955 | 0 | 0 | 0 | 3,58944991 |
| Downregulated | ENSMUSG00000020000 | 64,50599649 | 465,928603 | 277,770533 | 706,294922 | 283,703387 | 786,8927466 | 350,990986 | 191,889485 | 191,128984 | 909,341757 | 831,397322 | 483,469 | 121,212532 | 355,4818 | 666,6663 | 369,1760452 | 162,9657 | 110,9753667 | 34,22039 | 8,666531 | 19,48602 | 118,451847 |
| Downregulated | ENSMUSG00000029149 | 2,114950705 | 17,0202229 | 14,5239494 | 14,4535114 | 15,26655 | 27,86635359 | 11,3589316 | 6,11113011 | 7,4743178 | 19,4165372 | 22,0822662 | 14,0136 | 6,01053879 | 4,677392 | 3,18598 | 4,557728953 | 9,099605 | 0 | 1,901133 | 0,666656 | 3,542912 | 1,79472495 |
| Downregulated | ENSMUSG00000074852 | 1,057475352 | 4,25505574 | 6,35422788 | 3,8542697 | 39,4385875 | 19,90453828 | 10,2230384 | 4,88890409 | 14,9486356 | 90,6105068 | 4,41645324 | 14,88944 | 11,0193211 | 0 | 5,575465 | 4,557728953 | 2,481711 | 40,62491104 | 0 | 1,333313 | 4,42864 | 3,58944991 |
| Downregulated | ENSMUSG00000026778 | 22,2069824 | 14,8926951 | 6,35422788 | 20,2349159 | 31,8053125 | 37,15513812 | 28,397329 | 19,5556163 | 64,0655811 | 72,7226661 | 28,706946 | 15,76529 | 20,0351293 | 4,677392 | 7,168455 | 0 | 11,58132 | 43,59746551 | 10,45623 | 4,666594 | 5,314368 | 24,2287869 |
| Downregulated | ENSMUSG00000027004 | 12,68970423 | 12,7651672 | 6,35422788 | 33,7248598 | 26,7164625 | 21,2315075 | 61,3382305 | 45,2223628 | 58,7267827 | 111,10574 | 38,6439658 | 45,54418 | 5,00878232 | 23,38696 | 25,48784 | 14,58473265 | 54,59763 | 45,57916849 | 43,72606 | 17,99972 | 7,971552 | 23,3314244 |
| Downregulated | ENSMUSG00000040289 | 23,26445775 | 18,0839869 | 31,7711394 | 51,0690735 | 29,2608875 | 35,8281689 | 56,7946579 | 46,4445888 | 66,2011005 | 85,2170242 | 20,9781529 | 27,15134 | 7,01229525 | 43,03201 | 25,48784 | 35,55028583 | 28,12605 | 39,63405955 | 13,30793 | 7,999875 | 7,085824 | 23,3314244 |
| Downregulated | ENSMUSG00000074758 | 4,229901409 | 6,3825836 | 5,44648104 | 9,63567424 | 7,63327499 | 17,25059984 | 3,40767947 | 14,6667123 | 3,20327906 | 45,3052534 | 1,10411331 | 0 | 0 | 0 | 1,59299 | 0 | 12,40855 | 4,954257444 | 0,950566 | 5,999906 | 2,657184 | 5,38417486 |
| Downregulated | ENSMUSG00000034295 | 7,402327466 | 9,57387541 | 17,24719 | 25,052753 | 10,1777 | 22,55847672 | 21,58197 | 24,4445204 | 11,7453565 | 76,5874522 | 20,9781529 | 4,379248 | 3,00526939 | 2,806435 | 3,982475 | 0,911545791 | 15,7175 | 26,7529902 | 1,901133 | 15,33309 | 2,657184 | 19,7419745 |
| Downregulated | ENSMUSG00000026100 | 5,287376762 | 30,8491541 | 3,63098736 | 20,2349159 | 2,544425 | 0 | 13,6307179 | 14,6667123 | 2,13551937 | 104,633561 | 17,6658129 | 0 | 0 | 0 | 0 | 3,646183163 | 1,654474 | 0 | 0 | 0 | 0 | 11,6657122 |
| Downregulated | ENSMUSG00000020788 | 20,09203169 | 27,6578623 | 7,26197472 | 37,5791295 | 24,1720375 | 18,57756906 | 29,5332221 | 41,5556847 | 33,1005503 | 99,2400788 | 20,9781529 | 8,758497 | 11,0193211 | 0,935478 | 0 | 5,469274744 | 14,06303 | 6,935960422 | 7,604532 | 5,33325 | 24,80038 | 23,3314244 |
| Downregulated | ENSMUSG00000085977 | 3,172426057 | 3,1912918 | 4,5387342 | 3,8542697 | 2,544425 | 22,55847672 | 3,40767947 | 26,8889725 | 10,6775969 | 70,1152731 | 8,83290647 | 0,87585 | 11,0193211 | 2,806435 | 7,964949 | 7,292366325 | 0 | 0,990851489 | 0 | 1,333313 | 7,085824 | 0 |
| Downregulated | ENSMUSG00000037605 | 39,12658804 | 10,6376393 | 30,8633926 | 51,0690735 | 36,8941625 | 6,634846093 | 26,1255426 | 56,222397 | 44,8459068 | 113,263133 | 12,1452464 | 16,64114 | 0 | 4,677392 | 10,35443 | 7,292366325 | 2,481711 | 33,68895062 | 4,752832 | 3,999938 | 10,62874 | 22,4340619 |
| Downregulated | ENSMUSG00000034648 | 29,60930986 | 28,7216262 | 93,4979245 | 50,105506 | 41,9830125 | 13,26969219 | 24,9896495 | 108,778116 | 30,9650309 | 227,604963 | 46,372759 | 25,39964 | 12,0210776 | 2,806435 | 27,87732 | 27,34637372 | 14,06303 | 33,68895062 | 14,2585 | 0 | 13,28592 | 26,0235118 |
| Downregulated | ENSMUSG00000056222 | 20,09203169 | 15,956459 | 39,940861 | 47,2148038 | 11,4499125 | 2,653938437 | 11,3589316 | 52,5557189 | 30,9650309 | 122,971402 | 9,93701978 | 15,76529 | 4,00702586 | 23,38696 | 2,389485 | 25,52328214 | 0 | 24,77128722 | 5,703399 | 0 | 12,40019 | 8,97362477 |
| Downregulated | ENSMUSG00000020758 | 142,7591726 | 101,057574 | 223,305723 | 387,354104 | 202,281787 | 196,3914444 | 235,129884 | 309,223184 | 200,738821 | 633,194851 | 255,050174 | 275,8927 | 172,302112 | 46,77392 | 45,40021 | 215,1248066 | 52,11592 | 187,2709314 | 80,79815 | 42,666 | 84,14416 | 137,296459 |
| Downregulated | ENSMUSG00000039628 | 21,14950705 | 14,8926951 | 30,8633926 | 71,3039894 | 44,5274375 | 33,17423047 | 68,1535895 | 42,7779108 | 49,1169455 | 176,906228 | 39,7480791 | 4,379248 | 1,00175646 | 12,16122 | 21,50536 | 9,115457906 | 0 | 11,89021787 | 0,950566 | 0 | 12,40019 | 8,97362477 |
| Downregulated | ENSMUSG00000040624 | 187,1731374 | 259,5584 | 346,759293 | 333,394329 | 221,364975 | 416,6683346 | 366,89349 | 350,778868 | 314,989107 | 823,045437 | 355,524485 | 205,8247 | 159,279278 | 114,1284 | 92,39341 | 139,466506 | 62,04276 | 205,1062582 | 141,6344 | 86,66531 | 107,1731 | 252,158856 |
| Downregulated | ENSMUSG00000078190 | 1421,246874 | 880,796537 | 2028,81419 | 831,558687 | 871,465562 | 952,763899 | 1455,07914 | 1289,44845 | 622,503897 | 1816,52492 | 1418,7856 | 1224,438 | 454,797435 | 224,5148 | 1252,09 | 765,6984641 | 1814,13 | 897,7114489 | 933,4563 | 251,9961 | 62,88669 | 567,133086 |
| Downregulated | ENSMUSG00000057751 | 2216,468338 | 783,994019 | 2745,93419 | 2499,4939 | 583,945537 | 307,8568587 | 1425,54591 | 1025,44763 | 400,409882 | 4375,19304 | 1489,44885 | 987,0826 | 136,238879 | 332,0948 | 274,7908 | 800,3372042 | 1641,238 | 2545,497475 | 1148,284 | 563,3245 | 294,0617 | 1388,21975 |
| Downregulated | ENSMUSG00000025789 | 166,0236303 | 104,248886 | 282,309267 | 212,948401 | 133,582312 | 53,07876874 | 299,875794 | 103,889212 | 153,757395 | 380,779868 | 65,1426852 | 169,9148 | 39,0685021 | 1,870957 | 23,09835 | 43,75419795 | 175,3742 | 117,9113272 | 96,00722 | 19,33303 | 54,91514 | 75,3784481 |
| Downregulated | ENSMUSG00000027676 | 21,14950705 | 26,5940983 | 32,6788862 | 30,8341576 | 21,6276125 | 25,21241515 | 38,6203674 | 13,4444862 | 13,8808759 | 65,8004871 | 17,6658129 | 11,38605 | 5,00878232 | 2,806435 | 7,168455 | 9,115457906 | 26,47158 | 9,908514888 | 5,703399 | 27,99956 | 7,085824 | 16,1525246 |
| Downregulated | ENSMUSG00000000416 | 87,77045424 | 98,9300459 | 127,084558 | 158,988625 | 87,7826624 | 136,6778295 | 154,48147 | 156,444931 | 83,2852555 | 221,132784 | 93,8496312 | 44,66833 | 21,0368858 | 19,64505 | 54,95815 | 54,69274744 | 28,12605 | 72,33215868 | 32,31926 | 127,3313 | 16,82883 | 52,9443862 |
| Downregulated | ENSMUSG00000045871 | 13,74717958 | 3,1912918 | 49,0183294 | 30,8341576 | 49,6162874 | 26,53938437 | 35,2126879 | 31,7778766 | 13,8808759 | 104,633561 | 3,31233993 | 0 | 0 | 0 | 7,168455 | 3,646183163 | 5,790658 | 9,908514888 | 1,901133 | 7,999875 | 15,9431 | 8,0762623 |
| Downregulated | ENSMUSG00000024008 | 30,66678522 | 15,956459 | 69,8965067 | 56,850478 | 67,6272624 | 66,34846093 | 99,9585979 | 88,0002736 | 69,4043796 | 160,72578 | 23,1863795 | 56,05438 | 2,00351293 | 11,22574 | 13,54041 | 30,99255688 | 16,54474 | 32,69809913 | 18,06076 | 6,666563 | 16,82883 | 20,639337 |
| Downregulated | ENSMUSG00000029120 | 365,8864719 | 236,155593 | 325,881116 | 321,83152 | 251,898075 | 204,3532597 | 398,698499 | 317,778766 | 272,27872 | 617,014403 | 161,200543 | 89,33667 | 107,187942 | 188,0312 | 74,07403 | 144,0242349 | 57,07934 | 94,13089144 | 74,14419 | 0 | 47,82931 | 177,677771 |
| Downregulated | ENSMUSG00000000120 | 28,55183451 | 12,7651672 | 52,6493167 | 47,2148038 | 40,7108 | 29,19332281 | 55,6587648 | 34,2223286 | 21,3551937 | 63,643094 | 35,3316259 | 16,64114 | 15,026347 | 7,483827 | 7,168455 | 20,05400739 | 9,099605 | 14,86277233 | 4,752832 | 1,999969 | 20,37174 | 32,3050492 |

| | | | | | | | | | | | | | | | | | | | | | | | |
|---|---|---|---|---|---|---|---|---|---|---|---|---|---|---|---|---|---|---|---|---|---|---|---|
| Downregulated | ENSMUSG00000020081 | 31,72426057 | 4,25505574 | 1,81549368 | 2,89070227 | 5,08884999 | 17,25059984 | 9,08714527 | 9,77780817 | 7,4743178 | 36,6756813 | 6,62467985 | 0 | 0 | 0 | 0,796495 | 0 | 0 | 0,990851489 | 1,901133 | 0 | 0 | 8,0762623 |
| Downregulated | ENSMUSG00000117980 | 38,06911268 | 9,57387541 | 9,98521524 | 11,5628091 | 10,1777 | 17,25059984 | 18,1742905 | 9,77780817 | 7,4743178 | 28,0461092 | 6,62467985 | 4,379248 | 0 | 3,741914 | 0,796495 | 1,823091581 | 0 | 0,990851489 | 5,703399 | 0 | 0 | 4,48681239 |
| Downregulated | ENSMUSG00000025475 | 35,95416198 | 7,44634754 | 8,16972156 | 10,5992417 | 0 | 0 | 22,7178632 | 26,8889725 | 5,33879843 | 49,6200394 | 32,019286 | 1,751699 | 0 | 0 | 4,77897 | 1,823091581 | 0 | 3,963405955 | 2,851699 | 3,333281 | 8,85728 | 8,0762623 |
| Downregulated | ENSMUSG00000118502 | 26,43688381 | 3,1912918 | 9,98521524 | 9,63567424 | 6,36106249 | 6,634846093 | 12,4948247 | 20,7778424 | 2,13551937 | 20,4952337 | 2,20822662 | 1,751699 | 3,00526939 | 1,870957 | 2,389485 | 5,469274744 | 6,617895 | 4,954257444 | 1,901133 | 4,666594 | 3,542912 | 6,28153734 |
| Downregulated | ENSMUSG00000034255 | 100,4601585 | 59,5707803 | 90,774684 | 56,850478 | 50,8884999 | 61,04058406 | 71,561269 | 144,222671 | 44,8459068 | 151,017511 | 47,4768723 | 71,81967 | 46,0807974 | 7,483827 | 26,28433 | 25,52328214 | 55,42487 | 18,82617829 | 60,83626 | 6,666563 | 25,68611 | 50,2522987 |
| Downregulated | ENSMUSG00000019906 | 8,459802819 | 3,1912918 | 19,0626836 | 16,3806462 | 15,26655 | 27,86635359 | 21,58197 | 15,8889383 | 19,2196743 | 34,5182883 | 24,2904928 | 25,39964 | 12,0210776 | 16,83861 | 15,9299 | 17,31937002 | 10,75408 | 14,86277233 | 26,61586 | 5,33325 | 5,314368 | 8,97362477 |
| Downregulated | ENSMUSG00000021314 | 13,74717958 | 18,0839869 | 91,6824308 | 53,9597757 | 86,5104499 | 95,54178374 | 56,7946579 | 45,2223628 | 83,2852555 | 85,2170242 | 17,6658129 | 63,93703 | 48,0843103 | 28,99983 | 48,58619 | 67,45438851 | 54,59763 | 34,67980211 | 45,62719 | 5,999906 | 5,314368 | 31,4076867 |
| Downregulated | ENSMUSG00000097002 | 21,14950705 | 36,1679738 | 18,1549368 | 45,2876689 | 57,2495624 | 23,88544594 | 43,16394 | 41,5556847 | 27,7617518 | 58,2496115 | 23,1863795 | 23,64794 | 9,01580818 | 41,16105 | 23,09835 | 21,87709898 | 10,75408 | 14,86277233 | 19,01133 | 5,999906 | 0,885728 | 3,58944991 |
| Downregulated | ENSMUSG00000021469 | 9,517278171 | 23,4028065 | 11,8007089 | 29,8705901 | 39,4385875 | 21,2315075 | 27,2614358 | 40,3334587 | 27,7617518 | 25,8887162 | 30,9151726 | 22,77209 | 25,0439116 | 37,41914 | 19,11588 | 35,55028583 | 18,19921 | 10,89936638 | 23,76416 | 1,333313 | 3,542912 | 11,6657122 |
| Downregulated | ENSMUSG00000041695 | 135,3568451 | 362,743501 | 202,427545 | 249,563963 | 468,1742 | 208,3341673 | 331,680802 | 476,668148 | 404,680921 | 450,895141 | 382,023205 | 262,7549 | 38,0667457 | 43,03201 | 194,3448 | 219,6825355 | 172,0653 | 300,2280011 | 170,1514 | 50,66588 | 59,34378 | 265,619293 |
| Downregulated | ENSMUSG00000029217 | 283,4033944 | 352,105862 | 423,010027 | 395,062644 | 549,595799 | 335,7232123 | 564,5389 | 596,446299 | 525,337765 | 525,3252 | 576,347147 | 493,9792 | 210,368858 | 309,6433 | 340,8998 | 360,9721331 | 251,48 | 351,7522785 | 355,5119 | 21,333 | 122,2305 | 363,431803 |
| Downregulated | ENSMUSG00000036295 | 163,9086796 | 385,082544 | 253,261368 | 222,584075 | 403,291362 | 322,4535201 | 418,008682 | 386,223423 | 232,771611 | 358,127241 | 399,689018 | 298,6647 | 58,1018749 | 157,1604 | 288,3312 | 333,6257594 | 170,4108 | 195,1977433 | 193,9156 | 24,66628 | 28,3433 | 209,085457 |
| Downregulated | ENSMUSG00000033644 | 179,7708099 | 382,955016 | 136,162026 | 229,329047 | 385,480387 | 317,1456432 | 306,691153 | 337,334382 | 277,617518 | 290,169361 | 186,595149 | 237,3553 | 109,191455 | 104,7736 | 215,0536 | 261,6136419 | 221,6995 | 197,1794463 | 221,482 | 33,99947 | 42,51494 | 191,138208 |
| Downregulated | ENSMUSG00000001741 | 731,7729438 | 694,637849 | 568,249522 | 674,497197 | 880,371049 | 621,0215943 | 897,355595 | 713,779997 | 592,606626 | 847,855456 | 633,761039 | 559,668 | 275,483028 | 397,5783 | 575,0694 | 446,6574374 | 672,5436 | 474,6178631 | 678,7045 | 209,9967 | 120,459 | 384,968503 |
| Downregulated | ENSMUSG00000001870 | 23560,55085 | 40397,4992 | 23607,7721 | 28797,176 | 29806,6666 | 19769,18742 | 32603,5413 | 32382,8784 | 28083,1475 | 36492,3029 | 23459,0955 | 30225,57 | 11138,5301 | 14967,65 | 18692,14 | 24111,29771 | 23486,91 | 28136,21888 | 21986,6 | 2187,966 | 6305,498 | 19206,2491 |
| Downregulated | ENSMUSG00000030606 | 23,26445775 | 23,4028065 | 25,4169115 | 25,052753 | 15,26655 | 13,26969219 | 88,5996663 | 113,66702 | 99,3016508 | 113,263133 | 41,9563057 | 28,90304 | 0 | 5,61287 | 5,575465 | 18,23091581 | 16,54474 | 41,61576253 | 61,78682 | 16,66641 | 33,65766 | 25,1261494 |
| Downregulated | ENSMUSG00000055692 | 8,459802819 | 14,8926951 | 11,8007089 | 15,4170788 | 15,26655 | 6,634846093 | 44,2998332 | 26,8889725 | 25,6262325 | 26,9674127 | 26,4987194 | 13,13775 | 9,01580818 | 9,354784 | 8,761444 | 13,67318686 | 6,617895 | 24,77128722 | 24,71473 | 4,666594 | 9,743008 | 4,48681239 |
| Downregulated | ENSMUSG00000101970 | 76,13822537 | 56,3794885 | 71,7120004 | 54,9233432 | 80,1493874 | 76,96421468 | 97,6868116 | 105,111438 | 146,283077 | 159,647083 | 165,616996 | 115,6122 | 134,235366 | 87,93497 | 76,46352 | 37,37337742 | 52,94316 | 73,32301017 | 72,24305 | 27,99956 | 32,77194 | 56,5338361 |
| Downregulated | ENSMUSG00000116298 | 8,459802819 | 0 | 9,0774684 | 3,8542697 | 13,9943375 | 22,55847672 | 35,2126879 | 23,2222944 | 36,3038293 | 21,5739302 | 30,9151726 | 21,89624 | 26,0456681 | 4,677392 | 6,37196 | 1,823091581 | 9,926842 | 12,88106935 | 12,35736 | 0 | 4,42864 | 7,17889982 |
| Downregulated | ENSMUSG00000030862 | 1212,924229 | 3061,5126 | 1377,05196 | 2584,28783 | 2825,58396 | 2484,086377 | 3198,67513 | 3529,78875 | 4234,73491 | 3644,9155 | 3080,47613 | 1881,325 | 3659,41637 | 1999,117 | 1203,504 | 1889,634424 | 992,6842 | 2133,303255 | 1520,906 | 555,9913 | 753,7545 | 2318,78464 |
| Downregulated | ENSMUSG00000005968 | 345,7944402 | 413,80417 | 379,438179 | 346,884273 | 992,325749 | 826,7018232 | 2186,59433 | 867,780475 | 1797,03955 | 1671,97959 | 1168,15188 | 502,7377 | 976,712553 | 596,8352 | 517,7217 | 514,1118259 | 368,9476 | 492,4531899 | 230,0371 | 67,99894 | 322,405 | 872,236328 |
| Downregulated | ENSMUSG00000081355 | 17,97708099 | 39,3592656 | 39,940861 | 23,1256182 | 86,5104499 | 50,42483031 | 147,666111 | 67,2224312 | 135,60548 | 93,8465963 | 91,6414046 | 25,39964 | 59,1036314 | 42,09653 | 39,02825 | 41,93110637 | 33,91671 | 35,6706536 | 15,20906 | 9,999844 | 14,17165 | 59,2259235 |
| Downregulated | ENSMUSG00000029298 | 212,5525458 | 224,45419 | 356,744508 | 228,365479 | 253,170287 | 423,3031807 | 166,976294 | 293,334245 | 353,428456 | 439,029479 | 576,347147 | 218,9624 | 73,1282219 | 32,74174 | 174,4324 | 160,4320592 | 181,1649 | 177,3624165 | 117,8702 | 37,99941 | 102,7444 | 125,630747 |
| Downregulated | ENSMUSG00000046731 | 2611,96412 | 1977,53715 | 1700,20983 | 1978,20392 | 2783,60095 | 3565,56629 | 2260,42738 | 3467,45522 | 2838,10524 | 4085,02368 | 3882,06239 | 2003,068 | 520,913362 | 649,222 | 1214,655 | 1574,23958 | 848,745 | 1324,768441 | 571,2905 | 731,3219 | 1150,561 | 1536,28456 |
| Downregulated | ENSMUSG00000090799 | 74,02327466 | 42,5505574 | 29,0478989 | 21,1984833 | 100,504787 | 112,7923836 | 53,3869784 | 72,1113353 | 106,775969 | 103,554865 | 152,367637 | 31,53059 | 12,0210776 | 0 | 20,70887 | 12,76164107 | 23,98987 | 29,72554466 | 34,22039 | 27,99956 | 21,25747 | 44,8681239 |
| Downregulated | ENSMUSG00000028825 | 5,287376762 | 6,3825836 | 10,8929621 | 13,4899439 | 21,6276125 | 33,17423047 | 15,9025042 | 51,3334929 | 11,7453565 | 16,1804476 | 58,5180054 | 0,87585 | 0 | 0 | 5,575465 | 1,823091581 | 1,654474 | 23,78043573 | 2,851699 | 3,333281 | 3,542912 | 8,0762623 |
| Downregulated | ENSMUSG00000034586 | 26,43688381 | 26,5940983 | 9,0774684 | 32,7612924 | 114,499125 | 82,27209155 | 59,0664442 | 77,0002394 | 76,8786974 | 85,2170242 | 168,929336 | 41,16494 | 42,0737715 | 14,03218 | 59,73712 | 19,1424616 | 6,617895 | 28,73469318 | 21,86303 | 1,333313 | 20,37174 | 30,5103242 |
| Downregulated | ENSMUSG00000040714 | 20,09203169 | 51,0606688 | 21,7859242 | 55,8869106 | 97,9603624 | 57,0596764 | 46,5716195 | 66,0002052 | 68,3366199 | 64,7217906 | 131,389484 | 18,39284 | 14,0245905 | 19,64505 | 46,9932 | 35,55028583 | 36,39842 | 42,60661402 | 20,91246 | 1,999969 | 10,62874 | 26,9208743 |
| Downregulated | ENSMUSG00000039652 | 63,44852114 | 48,933141 | 59,9112914 | 63,59545 | 61,0661999 | 55,73270718 | 60,2023374 | 113,66702 | 100,36941 | 117,57792 | 98,2660845 | 39,41324 | 21,0368858 | 32,74174 | 15,1334 | 14,58473265 | 27,29882 | 30,71639615 | 45,62719 | 25,99959 | 38,0863 | 23,3314244 |
| Downregulated | ENSMUSG00000052861 | 48,64386621 | 37,2317377 | 41,7563546 | 20,2349159 | 45,79965 | 50,42483031 | 65,8818032 | 77,0002394 | 91,827333 | 59,328308 | 66,2467985 | 25,39964 | 37,0649892 | 4,677392 | 7,964949 | 7,292366325 | 15,7175 | 14,86277233 | 18,06076 | 23,33297 | 7,085824 | 20,639337 |
| Downregulated | ENSMUSG00000113302 | 465,289155 | 431,888157 | 352,205774 | 337,248598 | 573,767837 | 659,5037017 | 611,110519 | 953,336297 | 877,698462 | 792,841934 | 1012,4719 | 81,45402 | 407,714881 | 193,644 | 134,6076 | 127,6164107 | 205,1547 | 239,7860603 | 261,4058 | 114,6649 | 147,9166 | 343,689829 |
| Downregulated | ENSMUSG00000036306 | 0 | 7,44634754 | 4,5387342 | 14,4535114 | 10,1777 | 0 | 7,95125211 | 40,3334587 | 18,1519147 | 26,9674127 | 29,8110593 | 0 | 3,00526939 | 0 | 3,18598 | 2,734637372 | 2,481711 | 1,981702978 | 6,653965 | 9,999844 | 0 | 1,79472495 |
| Downregulated | ENSMUSG00000052581 | 0 | 3,1912918 | 2,72324052 | 36,6155621 | 24,1720375 | 30,52029203 | 11,3589316 | 48,8890409 | 36,3038293 | 48,5413429 | 44,1645324 | 0 | 7,01229525 | 7,483827 | 1,59299 | 5,469274744 | 0 | 7,92681191 | 4,752832 | 1,333313 | 0 | 15,2551621 |
| Downregulated | ENSMUSG00000049313 | 2,114950705 | 4,25505574 | 12,7084558 | 29,8705901 | 20,3554 | 18,57756906 | 20,4460768 | 79,4446914 | 40,5748681 | 71,1939696 | 70,6632518 | 22,77209 | 14,0245905 | 14,96765 | 2,389485 | 10,93854949 | 3,308947 | 22,78958424 | 4,752832 | 2,666625 | 9,743008 | 32,3050492 |
| Downregulated | ENSMUSG00000026344 | 0 | 4,25505574 | 0 | 1,92713485 | 13,9943375 | 5,307876874 | 6,81535895 | 11,0000342 | 18,1519147 | 31,2821988 | 25,3946061 | 3,503399 | 1,00175646 | 0 | 1,59299 | 3,646183163 | 0 | 0 | 0 | 0 | 3,542912 | 1,79472495 |
| Downregulated | ENSMUSG00000046215 | 35,95416198 | 49,9969049 | 31,7711394 | 35,6519947 | 80,1493874 | 38,48210734 | 54,5228716 | 102,666986 | 38,4393487 | 103,554865 | 114,827784 | 33,28229 | 68,1194396 | 33,67722 | 11,94742 | 32,81564846 | 31,435 | 17,8353268 | 30,41813 | 37,99941 | 22,1432 | 34,9971366 |
| Downregulated | ENSMUSG00000058145 | 53,93124297 | 41,4867934 | 77,1584814 | 79,9760962 | 125,949037 | 184,4487214 | 143,122538 | 178,444999 | 84,3530152 | 306,349809 | 191,011602 | 16,64114 | 20,0351293 | 11,22574 | 41,41774 | 23,70019056 | 2,481711 | 15,85362382 | 48,47889 | 9,999844 | 39,85776 | 98,7098725 |
| Downregulated | ENSMUSG00000114822 | 13,74717958 | 11,7014033 | 7,26197472 | 8,67210682 | 22,899825 | 39,80907656 | 23,8537563 | 31,7778766 | 23,4907131 | 57,170915 | 23,1863795 | 1,751699 | 4,00702586 | 18,70957 | 6,37196 | 10,0270037 | 4,136184 | 0,990851489 | 11,4068 | 0 | 7,085824 | 31,4076867 |
| Downregulated | ENSMUSG00000031618 | 116,3222888 | 125,524144 | 67,1732662 | 64,5590174 | 139,943375 | 126,0620758 | 88,5996663 | 152,778253 | 123,860124 | 223,290177 | 87,2249514 | 28,02719 | 12,0210776 | 39,29009 | 27,08083 | 20,96555318 | 9,926842 | 53,5059804 | 43,72606 | 29,33288 | 35,42912 | 73,5837231 |
| Downregulated | ENSMUSG00000007279 | 14,80465493 | 32,976682 | 15,4316963 | 15,4170788 | 33,077525 | 14,5966614 | 17,0383974 | 44,0001368 | 26,6939921 | 62,5643975 | 12,1452464 | 0 | 1,00175646 | 0,935478 | 1,59299 | 0,911545791 | 0 | 2,972554466 | 0 | 17,99972 | 11,51446 | 15,2551621 |
| Downregulated | ENSMUSG00000047146 | 29,60930986 | 77,6547672 | 39,940861 | 118,518793 | 104,321425 | 156,5823678 | 178,335226 | 195,556163 | 164,434992 | 266,438038 | 86,1208381 | 3,503399 | 5,00878232 | 33,67722 | 15,1334 | 2,734637372 | 33,08947 | 28,73469318 | 36,12153 | 53,99916 | 31,88621 | 54,7391111 |
| Downregulated | ENSMUSG00000032532 | 79,31065142 | 25,5303344 | 50,833823 | 129,118035 | 73,7883249 | 193,7375059 | 223,770952 | 261,556369 | 163,367232 | 338,710704 | 103,786651 | 42,04079 | 6,01053879 | 0 | 15,9299 | 58,3389306 | 53,77039 | 98,09429739 | 42,77549 | 43,99931 | 31,88621 | 55,6364736 |
| Downregulated | ENSMUSG00000019932 | 277,0585423 | 237,219357 | 740,721421 | 591,630398 | 405,835787 | 379,5131965 | 723,563942 | 1333,44859 | 538,150882 | 2024,71335 | 952,849785 | 91,96422 | 35,0614763 | 101,9671 | 66,10908 | 164,9897881 | 59,56105 | 416,1576253 | 165,3986 | 1,999969 | 125,7734 | 272,798193 |
| Downregulated | ENSMUSG00000020620 | 281,2884437 | 394,656419 | 399,40861 | 544,415595 | 352,402862 | 309,1838279 | 515,695494 | 798,113592 | 421,765076 | 1074,38172 | 523,349708 | 262,7549 | 130,22834 | 72,03184 | 90,00393 | 175,0167918 | 121,6038 | 358,6882389 | 161,5963 | 112,6649 | 253,3182 | 319,461042 |
| Downregulated | ENSMUSG00000031517 | 149,1040247 | 251,048288 | 216,951495 | 397,953346 | 362,580562 | 399,4177348 | 565,674793 | 590,335169 | 335,276541 | 851,091546 | 437,22887 | 102,4744 | 137,240636 | 46,77392 | 43,80722 | 73,83520904 | 52,11592 | 251,6762782 | 85,55098 | 0 | 77,94407 | 248,569406 |
| Downregulated | ENSMUSG00000040441 | 39,12658804 | 32,976682 | 68,081013 | 56,850478 | 59,7939874 | 69,00239937 | 87,4637732 | 143,000445 | 84,3530152 | 158,568387 | 98,2660845 | 17,51699 | 30,0526939 | 20,58052 | 25,48784 | 35,55028583 | 7,445132 | 64,40534677 | 45,62719 | 3,333281 | 32,77194 | 56,5338361 |
| Downregulated | ENSMUSG00000059498 | 58,16114438 | 121,269088 | 132,531039 | 161,879327 | 367,669412 | 135,3508603 | 429,367614 | 91,6669516 | 153,757395 | 340,868097 | 241,800815 | 60,43363 | 31,0544504 | 56,1287 | 38,23176 | 72,01211746 | 29,78053 | 51,52427742 | 16,15963 | 3,333281 | 37,20058 | 185,754033 |
| Downregulated | ENSMUSG00000057606 | 223,1272993 | 535,073259 | 176,102887 | 308,341576 | 1178,06877 | 703,2936859 | 1244,9389 | 750,446777 | 656,672207 | 1477,81422 | 652,530966 | 41,16494 | 98,1721335 | 62,67705 | 3,982475 | 21,87709898 | 3,308947 | 183,3075254 | 75,09475 | 0,666656 | 86,80135 | 374,200153 |
| Downregulated | ENSMUSG00000040258 | 88,82792959 | 152,118243 | 98,9444056 | 106,955984 | 393,113662 | 318,4726125 | 319,185977 | 202,88952 | 129,198922 | 416,376853 | 258,362514 | 16,64114 | 45,0790409 | 44,90296 | 18,31938 | 37,37337742 | 9,926842 | 72,33215868 | 39,92379 | 19,99969 | 35,42912 | 72,6863607 |
| Downregulated | ENSMUSG00000021313 | 15,86213028 | 62,7620721 | 8,16972156 | 30,8341576 | 164,115412 | 144,6396448 | 164,704508 | 122,222602 | 82,2174958 | 154,253601 | 115,931897 | 14,0136 | 17,0298599 | 46,77392 | 19,11588 | 25,52328214 | 10,75408 | 60,44194082 | 16,15963 | 0,666656 | 12,40019 | 43,9707614 |
| Downregulated | ENSMUSG00000014846 | 5854,18355 | 4057,19564 | 5168,71051 | 4380,37751 | 10387,6151 | 9984,116401 | 17096,3279 | 8883,13873 | 9436,8601 | 10889,4413 | 9051,52091 | 1684,259 | 3200,6119 | 2501,469 | 1898,844 | 2342,672682 | 1170,54 | 1846,947175 | 2045,619 | 1028,651 | 5259,453 | 3387,54335 |
| Downregulated | ENSMUSG00000033470 | 24,3219331 | 25,5303344 | 30,8633926 | 17,3442136 | 52,1607124 | 31,84726125 | 74,9689484 | 57,444623 | 43,7781471 | 63,643094 | 40,8521924 | 8,758497 | 10,0175646 | 16,83861 | 15,1334 | 9,115457906 | 9,926842 | 23,78043573 | 14,2585 | 12,66647 | 17,71456 | 18,844612 |
| Downregulated | ENSMUSG00000020712 | 8,459802819 | 34,0404459 | 14,5239494 | 28,9070227 | 45,79965 | 43,78998421 | 68,1535895 | 46,4445888 | 17,084155 | 69,0365766 | 32,019286 | 8,758497 | 7,01229525 | 4,677392 | 7,964949 | 25,52328214 | 0 | 6,935960422 | 8,555098 | 17,33306 | 10,62874 | 10,7683497 |
| Downregulated | ENSMUSG00000068748 | 2,114950705 | 15,956459 | 9,98521524 | 9,63567424 | 41,9830125 | 35,8281689 | 86,32788 | 69,6668832 | 23,4907131 | 127,286188 | 16,5616996 | 0 | 0 | 9,354784 | 3,18598 | 22,78864477 | 0 | 12,88106935 | 0,950566 | 2,666625 | 0 | 13,4604372 |
| Downregulated | ENSMUSG00000070564 | 45,47144015 | 42,5505574 | 49,0183294 | 67,4497197 | 69,9716874 | 76,96421468 | 102,230384 | 61,1113011 | 25,6262325 | 138,073153 | 64,0385719 | 60,43363 | 6,01053879 | 0 | 13,54041 | 49,22347269 | 10,75408 | 36,66150509 | 20,91246 | 0 | 6,200096 | 19,7419745 |
| Downregulated | ENSMUSG00000021994 | 2235,502895 | 3622,1162 | 3293,30554 | 2870,46736 | 4325,5225 | 3329,36577 | 4230,06612 | 4150,67957 | 3945,37204 | 7028,78645 | 2840,88354 | 2680,976 | 604,059148 | 1102,929 | 2469,134 | 2023,631655 | 1005,093 | 2543,515772 | 2052,273 | 1122,649 | 343,6625 | 1755,24101 |
| Downregulated | ENSMUSG00000040260 | 69,79337325 | 85,1011147 | 93,4979245 | 131,04517 | 159,026562 | 153,9284294 | 161,296828 | 173,556095 | 125,995643 | 237,313232 | 153,47175 | 87,58497 | 27,0474245 | 66,41897 | 76,46352 | 83,86221274 | 39,70737 | 100,0760004 | 38,02266 | 3,333281 | 20,37174 | 72,6863607 |
| Downregulated | ENSMUSG00000028132 | 17,97708099 | 68,0808918 | 32,6788862 | 27,9434553 | 61,0661999 | 21,2315075 | 49,979299 | 122,222602 | 81,1497361 | 166,119262 | 103,786651 | 51,67513 | 55,0966056 | 20,58052 | 3,18598 | 17,31937002 | 7,445132 | 35,6706536 | 8,555098 | 3,999938 | 10,62874 | 42,1760364 |
| Downregulated | ENSMUSG00000051515 | 6,344852114 | 10,6376393 | 7,26197472 | 14,4535114 | 12,722125 | 1,326969219 | 20,4460768 | 25,6667465 | 14,9486356 | 44,2265569 | 16,5616996 | 9,634347 | 7,01229525 | 12,16122 | 0 | 10,0270037 | 6,617895 | 8,917663399 | 2,851699 | 3,333281 | 1,771456 | 5,38417486 |
| Downregulated | ENSMUSG00000061356 | 12,68970423 | 27,6578623 | 22,693671 | 69,3768545 | 30,5331 | 61,04058406 | 39,7562605 | 70,8891093 | 54,455744 | 135,91576 | 101,578424 | 94,59177 | 2,00351293 | 19,64505 | 7,964949 | 51,95811007 | 15,7175 | 38,64320806 | 29,46756 | 17,99972 | 17,71456 | 10,7683497 |
| Downregulated | ENSMUSG00000015452 | 20,09203169 | 20,2115147 | 26,3246584 | 26,9798879 | 15,26655 | 47,77089187 | 39,7562605 | 59,8890751 | 25,6262325 | 84,1383277 | 78,3920449 | 42,91663 | 21,0368858 | 18,70957 | 0 | 26,43482793 | 0 | 17,8353268 | 2,851699 | 7,333219 | 11,51446 | 17,9472495 |
| Downregulated | ENSMUSG00000022842 | 8,459802819 | 14,8926951 | 19,9704305 | 28,9070227 | 13,9943375 | 31,84726125 | 48,8434058 | 37,8890067 | 26,6939921 | 32,3608953 | 30,9151726 | 10,5102 | 8,01405172 | 8,419306 | 3,18598 | 12,76164107 | 4,963421 | 5,945108933 | 6,653965 | 10,6665 | 5,314368 | 7,17889982 |
| Downregulated | ENSMUSG00000073598 | 1,057475352 | 0 | 1,81549368 | 16,3806462 | 3,8166375 | 10,61575375 | 21,58197 | 24,4445204 | 11,7453565 | 19,4165372 | 33,1233993 | 0 | 0 | 0,935478 | 0,796495 | 0 | 0 | 0 | 3,802266 | 1,999969 | 0,885728 | 3,58944991 |
| Downregulated | ENSMUSG00000074570 | 5,287376762 | 49,9969049 | 5,44648104 | 36,6155621 | 6,36106249 | 45,11695343 | 61,3382305 | 19,5556163 | 28,8295115 | 64,7217906 | 88,3290647 | 2,627549 | 2,00351293 | 0 | 15,1334 | 0 | 0,827237 | 1,981702978 | 7,604532 | 4,666594 | 7,971552 | 10,7683497 |
| Downregulated | ENSMUSG00000026768 | 167,0811057 | 256,367108 | 247,814887 | 352,665677 | 108,138062 | 236,2005209 | 311,234725 | 238,334074 | 202,87434 | 461,682106 | 368,773845 | 98,97102 | 48,0843103 | 0 | 60,53362 | 30,08101109 | 143,9392 | 262,5756445 | 97,90835 | 22,66631 | 40,74349 | 147,167446 |
| Downregulated | ENSMUSG00000027524 | 16,91960564 | 139,353075 | 47,2028357 | 207,166996 | 77,6049624 | 102,1766298 | 207,868448 | 232,222944 | 89,6918136 | 345,182883 | 293,69414 | 78,82647 | 17,0298599 | 0 | 22,30186 | 20,05400739 | 14,89026 | 84,22237655 | 32,31926 | 0 | 15,9431 | 205,496007 |
| Downregulated | ENSMUSG00000026073 | 24581,01456 | 20637,0203 | 19785,2501 | 12440,619 | 14841,631 | 15222,99088 | 17438,2318 | 27033,1951 | 29579,0788 | 11629,4271 | 10055,1599 | 5163,134 | 13626,8932 | 8222,855 | 4434,087 | 4085,548234 | 869,4259 | 4202,201164 | 3910,631 | 933,3188 | 5216,938 | 8804,02327 |
| Downregulated | ENSMUSG00000083674 | 25,37940846 | 17,0202229 | 17,24719 | 11,5628091 | 6,36106249 | 7,961815312 | 7,95125211 | 15,8889383 | 26,6939921 | 12,9443581 | 5,52056654 | 1,751699 | 1,00175646 | 1,870957 | 3,982475 | 4,557728953 | 1,654474 | 1,981702978 | 1,901133 | 3,333281 | 1,771456 | 2,69208743 |
| Downregulated | ENSMUSG00000003477 | 119,4947148 | 58,5070164 | 30,8633926 | 55,8869106 | 33,077525 | 25,21241515 | 0 | 89,2224996 | 185,790185 | 22,6526267 | 16,5616996 | 0 | 33,0579633 | 4,677392 | 0 | 0 | 0 | 0 | 0,950566 | 3,333281 | 7,085824 | 22,4340619 |
| Downregulated | ENSMUSG00000048826 | 10,57475352 | 5,31881967 | 4,5387342 | 3,8542697 | 11,4499125 | 19,90453828 | 1,13589316 | 7,33335613 | 19,2196743 | 20,4952337 | 2,20822662 | 0 | 0 | 6,548349 | 1,59299 | 1,823091581 | 0 | 0 | 0 | 1,333313 | 5,314368 | 3,58944991 |
| Downregulated | ENSMUSG00000028782 | 68,7358979 | 18,0839869 | 28,140152 | 22,1620508 | 61,0661999 | 72,98330702 | 37,4844742 | 42,7779108 | 65,1333408 | 43,1478604 | 22,0822662 | 11,38605 | 14,0245905 | 9,354784 | 3,982475 | 20,05400739 | 0,827237 | 19,81702978 | 25,6653 | 11,99981 | 15,05738 | 5,38417486 |
| Downregulated | ENSMUSG00000006362 | 23,26445775 | 2,12752787 | 9,0774684 | 26,9798879 | 35,62195 | 25,21241515 | 13,6307179 | 17,1111643 | 39,5071084 | 25,8887162 | 7,72879316 | 12,2619 | 4,00702586 | 0,935478 | 2,389485 | 5,469274744 | 1,654474 | 1,981702978 | 2,851699 | 0 | 0,885728 | 6,28153734 |
| Downregulated | ENSMUSG00000097790 | 51,81629226 | 121,269088 | 47,2028357 | 93,4660401 | 115,771337 | 132,6969219 | 92,0073458 | 634,335305 | 876,630702 | 63,643094 | 99,3701978 | 63,93703 | 79,1387607 | 79,51566 | 69,29506 | 58,3389306 | 62,04276 | 68,36875273 | 76,04532 | 158,6642 | 67,31533 | 66,4048233 |
| Downregulated | ENSMUSG00000097899 | 10,57475352 | 22,3390426 | 9,98521524 | 7,70853939 | 11,4499125 | 34,50119968 | 2,27178632 | 42,7779108 | 93,9628523 | 2,15739302 | 29,8110593 | 3,503399 | 2,00351293 | 0 | 1,59299 | 0 | 1,654474 | 3,963405955 | 0 | 7,333219 | 4,42864 | 5,38417486 |
| Downregulated | ENSMUSG00000007653 | 0 | 4,25505574 | 2,72324052 | 4,81783712 | 12,722125 | 25,21241515 | 4,54357263 | 51,3334929 | 65,1333408 | 2,15739302 | 28,706946 | 6,130948 | 3,00526939 | 0 | 2,389485 | 0 | 6,617895 | 10,89936638 | 5,703399 | 0,666656 | 7,971552 | 2,69208743 |
| Downregulated | ENSMUSG00000032355 | 27,49435916 | 47,869377 | 8,16972156 | 39,5062644 | 83,9660249 | 42,463015 | 70,4253758 | 213,889554 | 300,040472 | 97,0826858 | 105,994878 | 6,130948 | 12,0210776 | 0 | 7,168455 | 12,76164107 | 9,926842 | 20,80788127 | 2,851699 | 0 | 15,05738 | 39,483949 |
| Downregulated | ENSMUSG00000115453 | 14,80465493 | 27,6578623 | 10,8929621 | 16,3806462 | 35,62195 | 38,48210734 | 27,2614358 | 171,111643 | 158,028433 | 66,8791836 | 15,4575863 | 2,627549 | 5,00878232 | 5,61287 | 0 | 3,646183163 | 1,654474 | 1,981702978 | 8,555098 | 0,666656 | 7,085824 | 10,7683497 |
| Downregulated | ENSMUSG00000028270 | 2178,399226 | 2950,88115 | 3124,46462 | 2188,26162 | 4965,44538 | 5720,564301 | 3616,68382 | 12047,4819 | 12785,3545 | 4863,84256 | 3973,7038 | 1266,479 | 1452,54687 | 856,8982 | 818,7968 | 1296,218114 | 1422,847 | 1800,377155 | 1586,495 | 807,3207 | 2316,179 | 3150,63966 |
| Downregulated | ENSMUSG00000053702 | 0 | 2,12752787 | 2,72324052 | 9,63567424 | 15,26655 | 13,26969219 | 19,3101837 | 25,6667465 | 54,455744 | 23,7313232 | 4,41645324 | 0 | 11,0193211 | 0 | 1,59299 | 1,823091581 | 0 | 0,990851489 | 0 | 0 | 0 | 0,89736248 |
| Downregulated | ENSMUSG00000031302 | 15,86213028 | 27,6578623 | 51,7415699 | 35,6519947 | 35,62195 | 50,42483031 | 23,8537563 | 66,0002052 | 85,4207749 | 36,6756813 | 17,6658129 | 5,255098 | 4,00702586 | 7,483827 | 21,50536 | 8,203912116 | 39,70737 | 46,57001997 | 44,67663 | 15,99975 | 8,85728 | 10,7683497 |
| Downregulated | ENSMUSG00000010086 | 26,43688381 | 24,4665705 | 33,5866331 | 66,4861523 | 39,4385875 | 87,57996843 | 64,74591 | 157,667157 | 142,012038 | 90,6105068 | 14,353473 | 0 | 2,00351293 | 2,806435 | 0 | 0 | 0 | 5,945108933 | 19,01133 | 3,333281 | 9,743008 | 31,4076867 |
| Downregulated | ENSMUSG00000024039 | 32,78173592 | 18,0839869 | 59,0035446 | 49,1419386 | 36,8941625 | 71,65633781 | 87,4637732 | 99,0003078 | 111,047007 | 74,4300591 | 26,4987194 | 12,2619 | 25,0439116 | 13,0967 | 25,48784 | 0 | 31,435 | 13,87192084 | 20,91246 | 23,99963 | 0 | 22,4340619 |
| Downregulated | ENSMUSG00000060441 | 16,91960564 | 34,0404459 | 37,2176204 | 24,0891856 | 45,79965 | 9,28878453 | 6,81535895 | 36,6667807 | 43,7781471 | 35,5969848 | 7,72879316 | 1,751699 | 4,00702586 | 0 | 4,77897 | 16,40782423 | 21,50816 | 11,89021787 | 10,45623 | 13,99978 | 4,42864 | 5,38417486 |
| Downregulated | ENSMUSG00000031112 | 30,66678522 | 89,3561705 | 65,3577725 | 70,340422 | 64,8828374 | 67,67543015 | 36,3485811 | 103,889212 | 113,182527 | 111,10574 | 56,3097787 | 29,77889 | 9,01580818 | 7,483827 | 46,19671 | 38,28492321 | 52,94316 | 63,41449528 | 24,71473 | 35,33278 | 13,28592 | 24,2287869 |
| Downregulated | ENSMUSG00000039899 | 60538,34897 | 81894,9302 | 59894,952 | 40305,0618 | 90277,4711 | 74797,27395 | 71950,8803 | 132376,856 | 142473,31 | 70857,4163 | 68023,3168 | 40319,74 | 82005,7877 | 25665,79 | 23862,19 | 35506,53164 | 11940,34 | 41878,33817 | 26099,7 | 6549,898 | 24670,18 | 52396,9951 |
| Downregulated | ENSMUSG00000029321 | 6507,703318 | 6020,90387 | 3608,29369 | 4184,77332 | 7509,87038 | 6633,519124 | 5806,68582 | 8369,8038 | 10913,5717 | 6159,35707 | 6251,48955 | 3205,61 | 2658,66166 | 2229,245 | 1999,202 | 2632,544243 | 2820,878 | 5005,781721 | 2943,904 | 1791,972 | 2010,603 | 3539,19761 |
| Downregulated | ENSMUSG00000105504 | 219,9548733 | 154,24577 | 126,176811 | 119,482361 | 147,57665 | 151,2744909 | 206,732555 | 476,668148 | 381,190208 | 308,507202 | 168,929336 | 35,03399 | 72,1264654 | 7,483827 | 37,43526 | 57,42738481 | 117,4676 | 80,25897059 | 66,53965 | 19,99969 | 46,94358 | 138,193822 |

| | | | | | | | | | | | | | | | | | | | | | | | |
|---|---|---|---|---|---|---|---|---|---|---|---|---|---|---|---|---|---|---|---|---|---|---|---|
| Downregulated | ENSMUSG00000002565 | 632,3702607 | 704,211724 | 681,717877 | 434,568908 | 674,272624 | 1065,556283 | 830,337899 | 1669,56075 | 1320,81873 | 1087,32608 | 826,980868 | 174,2941 | 253,444386 | 63,61253 | 174,4324 | 164,9897881 | 114,1587 | 207,0879612 | 204,3718 | 20,66634 | 193,9744 | 260,235118 |
| Downregulated | ENSMUSG00000043631 | 3509,760694 | 4791,19276 | 3613,74017 | 3742,49587 | 4451,47153 | 5216,315998 | 5420,48215 | 9604,25208 | 7490,33419 | 7030,94385 | 5649,7478 | 1517,848 | 1977,46726 | 1323,702 | 1085,623 | 1212,355902 | 501,3055 | 1614,097075 | 1206,269 | 305,9952 | 1538,51 | 2883,22564 |
| Downregulated | ENSMUSG00000040987 | 297,150574 | 285,088734 | 383,976913 | 228,365479 | 298,969937 | 341,0310892 | 314,642405 | 474,223696 | 503,982572 | 332,238525 | 279,340667 | 44,66833 | 69,1211961 | 63,61253 | 26,28433 | 46,48883532 | 86,03263 | 137,7283569 | 118,8208 | 39,99938 | 138,1736 | 123,836022 |
| Downregulated | ENSMUSG00000020467 | 27701,62433 | 23281,5375 | 33261,6597 | 15769,7445 | 26773,712 | 30276,12969 | 30856,5376 | 54569,9474 | 46476,3758 | 28929,5617 | 26308,8119 | 18531,23 | 11917,8967 | 10861,84 | 12849,85 | 11718,83268 | 10532,38 | 16511,54921 | 16549,36 | 9655,183 | 17943,08 | 13448,7714 |
| Downregulated | ENSMUSG00000059852 | 7,402327466 | 4,25505574 | 1,81549368 | 9,63567424 | 21,6276125 | 3,980907656 | 6,81535895 | 24,4445204 | 21,3551937 | 4,31478604 | 7,72879316 | 0 | 0 | 0 | 0,796495 | 0,911545791 | 1,654474 | 0 | 0 | 0 | 0 | 8,97362477 |
| Downregulated | ENSMUSG00000115737 | 16,91960564 | 11,7014033 | 13,6162026 | 9,63567424 | 31,8053125 | 38,48210734 | 26,1255426 | 63,5557531 | 43,7781471 | 23,7313232 | 8,83290647 | 2,627549 | 3,00526939 | 0 | 7,964949 | 3,646183163 | 12,40855 | 2,972554466 | 0,950566 | 1,333313 | 4,42864 | 19,7419745 |
| Downregulated | ENSMUSG00000037627 | 49,70134156 | 82,9735868 | 95,3134182 | 96,3567424 | 109,410275 | 123,4081373 | 78,3766279 | 262,778595 | 207,145379 | 50,6987359 | 80,6002715 | 50,79928 | 28,049181 | 77,64471 | 67,70207 | 49,22347269 | 86,03263 | 61,43279231 | 75,09475 | 22,66631 | 20,37174 | 50,2522987 |
| Downregulated | ENSMUSG00000050875 | 37,01163733 | 98,9300459 | 78,9739751 | 161,879327 | 146,304437 | 108,8114759 | 84,0560937 | 67,2224312 | 144,147558 | 20,4952337 | 99,3701978 | 13,13775 | 18,0316164 | 39,29009 | 0,796495 | 27,34637372 | 30,60776 | 93,14003995 | 62,73739 | 3,333281 | 13,28592 | 69,0969108 |
| Downregulated | ENSMUSG00000048065 | 45,47144015 | 53,1881967 | 30,8633926 | 73,2311242 | 95,4159374 | 15,92363062 | 9,08714527 | 15,8889383 | 161,231713 | 64,7217906 | 22,0822662 | 0 | 1,00175646 | 0 | 0 | 13,67318686 | 2,481711 | 46,57001997 | 27,56643 | 0 | 5,314368 | 2,69208743 |
| Downregulated | ENSMUSG00000051041 | 4016,291388 | 4424,1942 | 3030,9667 | 5270,71381 | 4493,45455 | 2661,900253 | 3199,81103 | 2833,11992 | 7184,95492 | 4558,57145 | 2022,73558 | 585,0676 | 1360,38528 | 479,9004 | 319,3945 | 938,8921644 | 1105,188 | 2560,360247 | 1980,03 | 625,9902 | 1758,17 | 1634,09707 |
| Downregulated | ENSMUSG00000079669 | 44,4139648 | 46,8056131 | 37,2176204 | 42,3969667 | 19,0831875 | 43,78998421 | 12,4948247 | 24,4445204 | 51,2524649 | 33,4395918 | 51,8933255 | 1,751699 | 7,01229525 | 10,29026 | 8,761444 | 11,85009528 | 9,099605 | 28,73469318 | 21,86303 | 15,99975 | 15,9431 | 17,9472495 |
| Downregulated | ENSMUSG00000046618 | 283,4033944 | 297,853902 | 373,991698 | 315,086548 | 104,321425 | 221,6038595 | 263,527213 | 165,000513 | 517,863448 | 173,670138 | 156,78409 | 120,8673 | 87,1528124 | 50,51583 | 74,87053 | 76,56984641 | 71,96961 | 289,3286347 | 166,3491 | 34,66613 | 83,25843 | 83,4547104 |
| Downregulated | ENSMUSG00000106062 | 16,91960564 | 15,956459 | 27,2324052 | 26,9798879 | 12,722125 | 17,25059984 | 22,7178632 | 7,33335613 | 29,8972712 | 23,7313232 | 38,6439658 | 16,64114 | 7,01229525 | 8,419306 | 12,74392 | 15,49627844 | 8,272368 | 19,81702978 | 20,91246 | 3,333281 | 7,971552 | 7,17889982 |
| Downregulated | ENSMUSG00000044534 | 818,4859227 | 840,373508 | 601,836155 | 536,707055 | 960,520437 | 1159,771097 | 817,843074 | 867,780475 | 2163,28112 | 831,675009 | 1190,23415 | 1472,303 | 1020,78984 | 1053,349 | 1365,989 | 1551,450936 | 257,2707 | 381,4778232 | 357,413 | 89,9986 | 302,919 | 591,361873 |
| Downregulated | ENSMUSG00000051316 | 188,2306127 | 193,605036 | 175,19514 | 172,478569 | 288,792237 | 371,5513812 | 244,217029 | 259,111917 | 384,393487 | 194,165372 | 281,548894 | 219,8383 | 303,532209 | 96,35427 | 208,6817 | 230,621085 | 76,93303 | 126,8289906 | 99,80948 | 26,66625 | 137,2878 | 138,193822 |
| Downregulated | ENSMUSG00000027995 | 1377,890384 | 1725,4251 | 1540,44639 | 1639,99176 | 1834,53042 | 1840,506306 | 1723,14992 | 2160,89561 | 2444,10192 | 1586,76256 | 2423,52871 | 842,5674 | 1454,55039 | 1217,057 | 1158,104 | 1996,285281 | 770,1575 | 1050,302578 | 802,2781 | 267,3292 | 545,6085 | 1439,36941 |
| Downregulated | ENSMUSG00000094162 | 1179,085018 | 1205,24454 | 1300,80122 | 1362,48434 | 1507,57181 | 2237,270103 | 1627,7349 | 2709,67509 | 3820,44416 | 1674,13698 | 3225,11498 | 903,8769 | 2021,54455 | 1308,734 | 1136,598 | 1225,117543 | 316,0045 | 1133,534103 | 393,5345 | 11,33316 | 604,0665 | 914,412365 |
| Downregulated | ENSMUSG00000091938 | 1678,213384 | 1412,6785 | 1631,22107 | 1848,12232 | 2144,95027 | 2624,745114 | 2199,08915 | 4048,01258 | 5311,03668 | 1849,96451 | 3789,31688 | 1208,673 | 2330,08554 | 1887,795 | 1530,067 | 1668,128797 | 476,4884 | 1527,892996 | 622,6211 | 37,99941 | 792,7266 | 1084,91124 |
| Downregulated | ENSMUSG00000118633 | 951,7278171 | 1079,72039 | 1532,27667 | 1693,95153 | 1669,1428 | 3158,18674 | 1934,42605 | 2679,11944 | 3786,27585 | 2403,33582 | 4349,10232 | 853,0776 | 2606,57032 | 1680,119 | 1632,815 | 1361,849411 | 380,5289 | 1239,555213 | 469,5798 | 15,99975 | 900,7854 | 944,025326 |
| Downregulated | ENSMUSG00000095298 | 414,5303381 | 464,864839 | 805,171447 | 907,680513 | 871,465562 | 1439,761602 | 970,052757 | 1498,4491 | 2711,04184 | 1235,1075 | 2197,18548 | 559,668 | 1178,0656 | 870,9304 | 786,1405 | 753,8483689 | 197,7096 | 594,5108933 | 238,5922 | 18,66638 | 301,1475 | 456,757501 |
| Downregulated | ENSMUSG00000071005 | 1266,855472 | 1435,01755 | 1601,26543 | 2078,41493 | 2183,11665 | 3642,530505 | 2569,39032 | 3259,6768 | 6157,77011 | 2742,04653 | 5511,73364 | 1231,445 | 2962,19387 | 2094,536 | 1719,633 | 1580,620401 | 421,0636 | 1655,712838 | 519,0093 | 22,66631 | 838,7844 | 1066,96399 |
| Downregulated | ENSMUSG00000086104 | 8,459802819 | 8,51011147 | 6,35422788 | 0 | 5,08884999 | 25,21241515 | 5,67946579 | 17,1111643 | 17,084155 | 18,3378407 | 13,2493597 | 7,882647 | 6,01053879 | 14,96765 | 0,796495 | 6,380820534 | 0 | 1,981702978 | 0 | 0 | 0 | 7,17889982 |
| Downregulated | ENSMUSG00000020019 | 214,6674965 | 274,451095 | 168,840912 | 141,644411 | 264,6202 | 329,0883662 | 145,394324 | 276,223081 | 414,290758 | 406,668584 | 511,204462 | 174,2941 | 80,1405172 | 564,0935 | 220,6291 | 143,1126891 | 57,90658 | 144,6643174 | 60,83626 | 75,33216 | 181,5742 | 101,40196 |
| Downregulated | ENSMUSG00000036131 | 16,91960564 | 51,0606688 | 36,3098736 | 28,9070227 | 63,6106249 | 54,40573796 | 74,9689484 | 47,6668148 | 122,792364 | 61,485701 | 126,973031 | 1,751699 | 3,00526939 | 2,806435 | 5,575465 | 16,40782423 | 4,963421 | 15,85362382 | 7,604532 | 3,999938 | 7,085824 | 34,9971366 |
| Downregulated | ENSMUSG00000046314 | 237,9319543 | 265,940983 | 305,910685 | 243,782558 | 337,136312 | 506,9022415 | 412,329216 | 393,556779 | 565,912633 | 376,465082 | 573,034807 | 103,3503 | 96,1686206 | 110,3864 | 92,39341 | 154,0512386 | 91,82329 | 159,5270897 | 83,64985 | 105,3317 | 259,5183 | 180,369858 |
| Downregulated | ENSMUSG00000019843 | 510,7605952 | 691,446557 | 454,781167 | 539,597757 | 792,588387 | 1167,732912 | 786,038066 | 765,11349 | 933,221965 | 658,004871 | 1530,30105 | 729,5828 | 412,723663 | 697,8669 | 275,5873 | 298,0754735 | 240,7259 | 359,6790904 | 258,5541 | 251,9961 | 491,579 | 400,223665 |
| Downregulated | ENSMUSG00000025795 | 38,06911268 | 24,4665705 | 42,6641015 | 38,542697 | 69,9716874 | 61,04058406 | 49,979299 | 46,4445888 | 88,6240539 | 33,4395918 | 98,2660845 | 33,28229 | 24,0421551 | 46,77392 | 35,04578 | 9,115457906 | 6,617895 | 20,80788127 | 24,71473 | 27,99956 | 11,51446 | 16,1525246 |
| Downregulated | ENSMUSG00000032181 | 80,36812678 | 242,538177 | 159,763444 | 173,442136 | 167,93205 | 420,6492423 | 253,304174 | 262,778595 | 492,237215 | 429,321211 | 439,437097 | 123,4948 | 57,1001185 | 49,58035 | 185,5833 | 134,908777 | 23,16263 | 64,40534677 | 75,09475 | 25,99959 | 69,08679 | 86,1467978 |
| Downregulated | ENSMUSG00000079362 | 40,18406339 | 30,8491541 | 73,527494 | 46,2512364 | 41,9830125 | 108,8114759 | 37,4844742 | 55,000171 | 123,860124 | 98,1613823 | 71,7673651 | 31,53059 | 15,026347 | 22,45148 | 35,84227 | 14,58473265 | 23,16263 | 23,78043573 | 21,86303 | 5,999906 | 26,57184 | 10,7683497 |
| Downregulated | ENSMUSG00000020340 | 12,68970423 | 0 | 0 | 13,4899439 | 11,4499125 | 31,84726125 | 20,4460768 | 8,55558215 | 33,1005503 | 12,9443581 | 27,6028327 | 3,503399 | 5,00878232 | 0 | 0 | 0 | 0,827237 | 0 | 0 | 0 | 0,885728 | 0 |
| Downregulated | ENSMUSG00000024334 | 0 | 2,12752787 | 0,90774684 | 5,78140454 | 17,810975 | 22,55847672 | 10,2230384 | 1,22222602 | 49,1169455 | 6,47217906 | 30,9151726 | 0,87585 | 2,00351293 | 3,741914 | 4,77897 | 1,823091581 | 1,654474 | 3,963405955 | 2,851699 | 0 | 2,657184 | 0 |
| Downregulated | ENSMUSG00000107667 | 22,2069824 | 17,0202229 | 25,4169115 | 9,63567424 | 103,049212 | 159,2363062 | 98,8227048 | 112,444794 | 280,820797 | 43,1478604 | 96,0578579 | 0 | 9,01580818 | 0 | 0 | 0 | 0 | 20,80788127 | 0,950566 | 11,99981 | 38,0863 | 17,9472495 |
| Downregulated | ENSMUSG00000075010 | 356,3691937 | 479,757534 | 526,493167 | 693,768545 | 586,489962 | 1319,007403 | 675,856429 | 1112,22568 | 2314,903 | 627,801368 | 1241,02336 | 231,2243 | 358,628814 | 405,9976 | 262,0468 | 248,8520008 | 167,1018 | 282,3926743 | 173,9537 | 145,9977 | 431,3495 | 348,176641 |
| Downregulated | ENSMUSG00000033377 | 250,6216585 | 403,166531 | 208,781773 | 248,600395 | 432,55225 | 318,4726125 | 197,64541 | 232,222944 | 27,7617518 | 209,267123 | 550,952541 | 185,6801 | 101,177403 | 147,8056 | 393,4685 | 137,6434144 | 63,69724 | 129,801545 | 100,76 | 8,666531 | 47,82931 | 69,0969108 |
| Downregulated | ENSMUSG00000085084 | 10,57475352 | 28,7216262 | 9,0774684 | 2,89070227 | 15,26655 | 14,5966614 | 14,7666111 | 18,3333903 | 2,13551937 | 15,1017511 | 40,8521924 | 15,76529 | 5,00878232 | 4,677392 | 11,94742 | 3,646183163 | 5,790658 | 15,85362382 | 2,851699 | 0 | 0,885728 | 1,79472495 |
| Downregulated | ENSMUSG00000019856 | 22,2069824 | 56,3794885 | 22,693671 | 22,1620508 | 41,9830125 | 30,52029203 | 34,0767947 | 28,1111985 | 37,371589 | 29,1248057 | 72,8714784 | 19,26869 | 14,0245905 | 9,354784 | 38,23176 | 3,646183163 | 10,75408 | 41,61576253 | 20,91246 | 1,333313 | 7,085824 | 15,2551621 |
| Downregulated | ENSMUSG00000042873 | 59,21861973 | 78,7185311 | 46,2950888 | 62,6318826 | 63,6106249 | 39,80907656 | 74,9689484 | 68,4446572 | 29,8972712 | 65,8004871 | 73,9755917 | 25,39964 | 36,0632327 | 4,677392 | 61,33011 | 49,22347269 | 10,75408 | 33,68895062 | 18,06076 | 8,666531 | 11,51446 | 12,5630747 |
| Downregulated | ENSMUSG00000081972 | 15,86213028 | 23,4028065 | 19,0626836 | 9,63567424 | 17,810975 | 15,92363062 | 13,6307179 | 19,5556163 | 12,8131162 | 14,0230546 | 20,9781529 | 0 | 10,0175646 | 0,935478 | 13,54041 | 7,292366325 | 6,617895 | 24,77128722 | 5,703399 | 5,33325 | 0,885728 | 3,58944991 |
| Downregulated | ENSMUSG00000037499 | 1280,602652 | 1578,62568 | 1432,42451 | 1591,81338 | 1328,18985 | 1177,021697 | 1130,21369 | 1084,11448 | 1732,97397 | 1194,11704 | 1713,58386 | 827,678 | 1703,98775 | 1015,93 | 421,3458 | 695,5094383 | 539,3584 | 646,0351707 | 1115,965 | 465,9927 | 473,8645 | 494,446725 |
| Downregulated | ENSMUSG00000057280 | 197,7478909 | 235,091829 | 173,379646 | 131,04517 | 286,247812 | 210,9881058 | 180,607012 | 232,222944 | 202,87434 | 196,322765 | 249,529608 | 122,619 | 309,542748 | 68,28992 | 39,82475 | 83,86221274 | 41,36184 | 105,0302578 | 70,34192 | 117,3315 | 161,2025 | 162,422608 |
| Downregulated | ENSMUSG00000055675 | 26,43688381 | 31,912918 | 9,98521524 | 13,4899439 | 26,7164625 | 18,57756906 | 18,1742905 | 13,4444862 | 22,4229534 | 28,0461092 | 39,7480791 | 9,634347 | 25,0439116 | 6,548349 | 0,796495 | 5,469274744 | 0,827237 | 1,981702978 | 7,604532 | 5,33325 | 16,82883 | 23,3314244 |
| Downregulated | ENSMUSG00000020672 | 247,4492324 | 294,66261 | 159,763444 | 66,4861523 | 284,9756 | 214,9690134 | 166,976294 | 235,889622 | 253,059045 | 148,860118 | 340,066899 | 162,0322 | 155,272252 | 99,16071 | 146,5551 | 147,6704181 | 55,42487 | 133,764951 | 153,0412 | 29,99953 | 72,6297 | 92,4283352 |
| Downregulated | ENSMUSG00000044197 | 1356,740877 | 1128,65353 | 426,641015 | 259,199637 | 1175,52435 | 918,2626993 | 523,646746 | 633,113079 | 696,179315 | 730,277537 | 1075,40636 | 638,4944 | 738,294514 | 445,2877 | 414,9739 | 473,0922653 | 209,2909 | 410,2125164 | 356,4624 | 55,33247 | 157,6596 | 461,244313 |
| Downregulated | ENSMUSG00000041132 | 171,3110071 | 281,897442 | 96,221165 | 45,2876689 | 254,4425 | 139,331768 | 211,276127 | 244,445204 | 242,381449 | 107,869651 | 333,442219 | 143,6393 | 73,1282219 | 156,2249 | 66,10908 | 75,65830062 | 69,48789 | 124,8472876 | 91,25438 | 43,33266 | 54,91514 | 148,962171 |
| Downregulated | ENSMUSG00000041112 | 1885,478553 | 1982,85597 | 983,089828 | 872,992086 | 2683,09616 | 1293,794988 | 1747,00368 | 1465,449 | 1714,82206 | 1495,07336 | 2214,8513 | 951,1728 | 819,436788 | 567,8354 | 623,6555 | 597,0624929 | 379,7017 | 639,0992103 | 586,4995 | 288,6622 | 533,2083 | 1154,90551 |
| Downregulated | ENSMUSG00000027698 | 2886,907712 | 3478,50806 | 1987,96558 | 1482,93027 | 4451,47153 | 2774,692636 | 2618,23373 | 3109,343 | 3399,74684 | 2536,01549 | 3077,16379 | 1681,631 | 1201,106 | 1116,961 | 1529,27 | 1518,635287 | 1345,914 | 2021,337037 | 1615,963 | 267,3292 | 674,9247 | 2044,19172 |
| Downregulated | ENSMUSG00000022123 | 63,44852114 | 154,24577 | 38,1253673 | 9,63567424 | 109,410275 | 65,02149171 | 109,045743 | 64,7779792 | 76,8786974 | 74,4300591 | 61,8303453 | 25,39964 | 24,0421551 | 9,354784 | 7,168455 | 5,469274744 | 22,33539 | 70,35045571 | 18,06076 | 10,6665 | 7,971552 | 43,9707614 |
| Downregulated | ENSMUSG00000102758 | 604,8759015 | 1254,17768 | 455,688914 | 452,876689 | 848,565737 | 587,8473638 | 553,179968 | 570,779552 | 575,522471 | 766,953218 | 342,275126 | 333,6987 | 211,370614 | 199,2569 | 123,4567 | 195,982345 | 133,1851 | 421,1118827 | 442,0134 | 89,9986 | 182,46 | 423,555089 |
| Downregulated | ENSMUSG00000050097 | 96,23025706 | 225,517954 | 48,1105825 | 146,462248 | 96,6881499 | 75,63724546 | 109,045743 | 55,000171 | 91,827333 | 99,2400788 | 150,15941 | 75,32307 | 33,0579633 | 63,61253 | 19,91237 | 29,1694653 | 3,308947 | 39,63405955 | 15,20906 | 11,33316 | 28,3433 | 53,8417486 |
| Downregulated | ENSMUSG00000073535 | 330,9897853 | 391,465128 | 59,0035446 | 217,766238 | 296,425512 | 319,7995817 | 222,635059 | 155,222705 | 127,063403 | 227,604963 | 266,091307 | 162,908 | 113,19848 | 89,80593 | 177,6184 | 123,9702275 | 35,57118 | 111,9662182 | 40,87436 | 15,33309 | 42,51494 | 169,601508 |
| Downregulated | ENSMUSG00000097767 | 222,069824 | 109,567685 | 186,995849 | 100,211012 | 559,773499 | 700,6397474 | 1143,84441 | 598,890751 | 345,954138 | 502,672573 | 859,000154 | 89,33667 | 26,0456681 | 83,25758 | 42,21423 | 32,81564846 | 28,95329 | 92,14918846 | 58,93512 | 39,99938 | 70,85824 | 221,648532 |
| Downregulated | ENSMUSG00000097391 | 280,2309684 | 96,802518 | 206,058533 | 198,494889 | 435,096675 | 877,1266535 | 917,801672 | 542,668354 | 303,243751 | 420,691639 | 936,288086 | 117,3639 | 68,1194396 | 85,12853 | 54,16166 | 52,86965586 | 38,88013 | 221,9507335 | 69,39135 | 84,66535 | 106,2874 | 244,979956 |
| Downregulated | ENSMUSG00000097451 | 636,6001621 | 253,175816 | 512,876965 | 453,840257 | 597,939874 | 1092,095667 | 1360,8 | 854,335989 | 503,982572 | 785,291059 | 870,041287 | 169,9148 | 203,356562 | 216,0955 | 230,187 | 175,0167918 | 107,5408 | 410,2125164 | 151,1401 | 442,6598 | 297,6046 | 445,091789 |
| Downregulated | ENSMUSG00000097961 | 424,0476163 | 186,158688 | 337,681824 | 175,369271 | 704,805724 | 877,1266535 | 1263,11319 | 798,113592 | 269,075441 | 378,622475 | 819,252075 | 135,7567 | 143,251174 | 84,19306 | 56,55114 | 41,01956058 | 60,38829 | 305,1822586 | 102,6612 | 268,6625 | 195,7459 | 427,144539 |
| Downregulated | ENSMUSG00000021268 | 2249,250074 | 1116,95213 | 1600,35768 | 1418,37125 | 3594,00031 | 5723,21824 | 6163,35628 | 3916,01217 | 1712,68654 | 2521,99244 | 4784,12297 | 657,7631 | 1168,04804 | 596,8352 | 442,0547 | 299,8985651 | 368,1204 | 1626,978145 | 604,5603 | 1444,644 | 1124,875 | 1969,71064 |
| Downregulated | ENSMUSG00000037553 | 381,7486022 | 317,001652 | 420,286787 | 365,192054 | 913,448574 | 1027,074175 | 790,581638 | 597,668525 | 717,534509 | 728,120144 | 1017,99247 | 373,9878 | 394,692047 | 543,5129 | 379,1316 | 411,1071516 | 129,8762 | 427,0569917 | 270,9115 | 109,9983 | 182,46 | 443,297064 |
| Downregulated | ENSMUSG00000053825 | 12,68970423 | 10,6376393 | 17,24719 | 19,2713485 | 45,79965 | 55,73270718 | 53,3869784 | 47,6668148 | 29,8972712 | 22,6526267 | 66,2467985 | 27,15134 | 12,0210776 | 26,19339 | 7,168455 | 19,1424616 | 2,481711 | 36,66150509 | 4,752832 | 1,333313 | 7,085824 | 17,0498871 |
| Downregulated | ENSMUSG00000034059 | 60,27609508 | 56,3794885 | 112,560608 | 54,9233432 | 217,548337 | 334,3962431 | 356,670452 | 213,889554 | 214,619697 | 159,647083 | 244,009041 | 45,54418 | 19,0333728 | 37,41914 | 66,10908 | 29,1694653 | 59,56105 | 26,7529902 | 64,63852 | 25,99959 | 62,00096 | 70,8916357 |
| Downregulated | ENSMUSG00000027215 | 1167,452789 | 1710,53241 | 1774,64507 | 3222,16947 | 5448,88613 | 5593,175256 | 7638,88149 | 3979,56793 | 4457,89669 | 4258,69382 | 4859,20267 | 2123,936 | 821,440301 | 1789,57 | 2261,249 | 1717,35227 | 1662,746 | 1731,017551 | 1768,054 | 276,6623 | 813,984 | 2793,48939 |
| Downregulated | ENSMUSG00000022096 | 596,4160987 | 588,261455 | 777,031295 | 661,97082 | 1090,28611 | 1081,479913 | 1421,00234 | 1174,55921 | 836,055834 | 1027,99777 | 1568,94501 | 698,9281 | 222,389935 | 652,9639 | 546,3955 | 793,9563836 | 746,9949 | 649,9985767 | 619,7694 | 275,9957 | 293,176 | 646,100984 |
| Downregulated | ENSMUSG00000111229 | 119,4947148 | 59,5707803 | 87,1436966 | 62,6318826 | 100,504787 | 82,27209155 | 110,181636 | 97,7780817 | 93,9628523 | 131,600974 | 158,992316 | 44,66833 | 38,0667457 | 39,29009 | 60,53362 | 52,86965586 | 33,08947 | 82,24067357 | 47,52832 | 15,33309 | 30,11475 | 59,2259235 |
| Downregulated | ENSMUSG00000037465 | 2618,308972 | 2042,42675 | 2324,73966 | 1590,84982 | 2839,5783 | 1961,260505 | 4270,95827 | 3042,12057 | 3314,32606 | 3230,69604 | 4954,15642 | 1469,676 | 2101,68506 | 1312,476 | 1573,874 | 1472,146452 | 1212,729 | 1724,081591 | 1684,404 | 684,656 | 743,1258 | 2254,17454 |
| Downregulated | ENSMUSG00000091191 | 44,4139648 | 36,1679738 | 36,3098736 | 41,4333992 | 43,255225 | 124,7351066 | 97,6868116 | 63,5557531 | 43,7781471 | 73,3513626 | 173,345789 | 85,83327 | 21,0368858 | 2,806435 | 40,62124 | 6,380820534 | 17,37197 | 57,46938635 | 30,41813 | 7,999875 | 11,51446 | 40,3813115 |
| Downregulated | ENSMUSG00000084126 | 19,03455634 | 14,8926951 | 15,4316963 | 7,70853939 | 22,899825 | 29,19332281 | 31,8050084 | 19,5556163 | 10,6775969 | 34,5182883 | 37,5398525 | 14,88944 | 2,00351293 | 0 | 11,15093 | 3,646183163 | 10,75408 | 16,84447531 | 14,2585 | 1,333313 | 0,885728 | 11,6657122 |
| Downregulated | ENSMUSG00000072573 | 33,83921127 | 24,4665705 | 24,5091647 | 26,0163204 | 161,570987 | 99,5226914 | 163,568615 | 51,3334929 | 54,455744 | 94,9252928 | 270,507761 | 26,27549 | 25,0439116 | 28,99983 | 29,47031 | 38,28492321 | 5,790658 | 45,57916849 | 26,61586 | 12,66647 | 54,91514 | 64,6100984 |
| Downregulated | ENSMUSG00000076396 | 14,80465493 | 1,06376393 | 7,26197472 | 10,5992417 | 6,36106249 | 25,21241515 | 44,2998332 | 2,44445204 | 12,8131162 | 22,6526267 | 49,6850989 | 5,255098 | 1,00175646 | 0,935478 | 3,982475 | 0 | 0,827237 | 4,954257444 | 0,950566 | 2,666625 | 3,542912 | 8,97362477 |
| Downregulated | ENSMUSG00000039391 | 6,344852114 | 3,1912918 | 7,26197472 | 7,70853939 | 11,4499125 | 17,25059984 | 40,8921537 | 15,8889383 | 10,6775969 | 28,0461092 | 43,060419 | 0,87585 | 3,00526939 | 0 | 1,59299 | 1,823091581 | 0 | 3,963405955 | 2,851699 | 7,333219 | 2,657184 | 10,7683497 |
| Downregulated | ENSMUSG00000017400 | 340,5070634 | 245,729469 | 341,312812 | 134,899439 | 334,591887 | 570,596764 | 266,934892 | 388,667875 | 325,666704 | 329,002435 | 500,163329 | 192,6869 | 170,298599 | 301,224 | 270,8083 | 286,2253783 | 336,6854 | 225,9141394 | 281,3677 | 83,99869 | 77,05834 | 225,237982 |
| Downregulated | ENSMUSG00000025997 | 39,12658804 | 67,0171278 | 24,5091647 | 40,4698318 | 43,255225 | 99,5226914 | 88,5996663 | 74,5557873 | 65,1333408 | 60,4070045 | 80,6002715 | 17,51699 | 40,0702586 | 68,28992 | 66,90558 | 26,43482793 | 36,39842 | 36,66150509 | 20,91246 | 14,66644 | 9,743008 | 37,6892241 |
| Downregulated | ENSMUSG00000066278 | 158,6213028 | 168,074702 | 96,221165 | 90,5753379 | 242,992587 | 185,7756906 | 239,673456 | 149,111575 | 199,671061 | 182,29971 | 367,669732 | 232,1002 | 163,286304 | 198,3214 | 157,706 | 142,2011433 | 73,62408 | 142,6826144 | 96,95778 | 39,33272 | 54,91514 | 130,117559 |
| Downregulated | ENSMUSG00000024210 | 156,5063521 | 254,23958 | 63,5422788 | 49,1419386 | 203,554 | 291,9332281 | 188,558264 | 84,3335955 | 193,264503 | 118,656616 | 404,105471 | 55,17853 | 93,1633512 | 84,19306 | 125,0497 | 78,39293799 | 9,926842 | 63,41449528 | 59,88569 | 3,333281 | 23,91466 | 113,067672 |
| Downregulated | ENSMUSG00000043969 | 62,39104579 | 97,8662819 | 70,8042535 | 61,6683151 | 89,0548749 | 139,331768 | 110,181636 | 97,7780817 | 144,147558 | 146,702725 | 311,359953 | 61,30948 | 84,147543 | 77,64471 | 59,73712 | 39,196469 | 43,01632 | 81,24982208 | 54,18229 | 18,66638 | 52,25795 | 77,1731731 |
| Downregulated | ENSMUSG00000061825 | 64,50599649 | 159,56459 | 40,8486078 | 141,644411 | 291,336662 | 298,5680742 | 273,750251 | 74,5557873 | 224,229534 | 272,910217 | 904,2688 | 45,54418 | 72,1264654 | 113,1929 | 121,8637 | 93,88921644 | 31,435 | 49,54257444 | 25,6653 | 4,666594 | 27,45757 | 50,2522987 |
| Downregulated | ENSMUSG00000031884 | 15,86213028 | 91,4836983 | 13,6162026 | 64,5590174 | 146,304437 | 177,8138753 | 147,666111 | 45,2223628 | 114,250286 | 89,5318103 | 462,623476 | 11,38605 | 47,0825538 | 64,54801 | 43,80722 | 74,74675483 | 6,617895 | 39,63405955 | 3,802266 | 1,999969 | 16,82883 | 27,8182368 |
| Downregulated | ENSMUSG00000046329 | 60,27609508 | 39,3592656 | 27,2324052 | 39,5062644 | 58,5217749 | 98,19572218 | 56,7946579 | 46,4445888 | 33,1005503 | 35,5969848 | 93,8496312 | 38,53739 | 49,0860668 | 33,67722 | 18,31938 | 14,58473265 | 13,23579 | 29,72554466 | 26,61586 | 16,66641 | 31,00048 | 37,6892241 |
| Downregulated | ENSMUSG00000036863 | 264,3688381 | 319,12918 | 139,793013 | 143,571546 | 366,3972 | 432,5919653 | 397,562605 | 315,334314 | 162,299472 | 423,927728 | 560,889561 | 161,1563 | 105,184429 | 151,5475 | 99,56187 | 84,77375853 | 33,08947 | 92,14918846 | 120,7219 | 53,99916 | 45,17213 | 121,143934 |
| Downregulated | ENSMUSG00000057716 | 16,91960564 | 27,6578623 | 9,0774684 | 10,5992417 | 36,8941625 | 42,463015 | 19,3101837 | 35,4445546 | 8,54207749 | 36,6756813 | 51,8933255 | 13,13775 | 16,0281034 | 1,870957 | 4,77897 | 14,58473265 | 0 | 20,80788127 | 7,604532 | 0,666656 | 1,771456 | 7,17889982 |
| Downregulated | ENSMUSG00000040264 | 1258,395669 | 1719,04252 | 1761,93662 | 524,180679 | 3038,04345 | 3325,384862 | 1754,95493 | 97,7780817 | 74,743178 | 3061,34069 | 2400,34233 | 630,6118 | 380,667457 | 224,5148 | 218,2396 | 361,8836789 | 488,897 | 1168,213905 | 961,0227 | 14,66644 | 666,9532 | 946,717414 |
| Downregulated | ENSMUSG00000022199 | 484,3237114 | 455,290964 | 565,526281 | 364,228486 | 788,771749 | 605,0979637 | 662,225711 | 246,889656 | 311,785828 | 824,124133 | 766,254636 | 228,5968 | 123,216045 | 128,1605 | 98,76537 | 118,5009528 | 85,20539 | 235,8226543 | 232,8888 | 143,3311 | 111,6017 | 255,748306 |
| Downregulated | ENSMUSG00000039835 | 547,7722325 | 586,133928 | 502,891749 | 424,933234 | 624,656337 | 745,7567009 | 629,28481 | 411,890169 | 415,358518 | 884,531138 | 882,186534 | 259,2515 | 144,252931 | 246,9663 | 166,4674 | 209,6555318 | 57,07934 | 231,8592484 | 234,7899 | 325,9949 | 315,3192 | 269,208743 |
| Downregulated | ENSMUSG00000062995 | 103,6325845 | 118,077797 | 117,099342 | 129,118035 | 136,126737 | 141,9857064 | 165,840401 | 83,1113695 | 42,7103874 | 172,591441 | 182,178696 | 57,80608 | 21,0368858 | 64,54801 | 77,26001 | 14,58473265 | 35,57118 | 56,47853486 | 49,42946 | 21,99966 | 16,82883 | 64,6100984 |
| Downregulated | ENSMUSG00000026207 | 58,16114438 | 79,782295 | 87,1436966 | 52,9962083 | 96,6881499 | 111,4654144 | 109,045743 | 48,8890409 | 55,5235037 | 110,027044 | 90,5372913 | 35,90984 | 19,0333728 | 71,09636 | 51,77217 | 22,78864477 | 14,06303 | 26,7529902 | 34,22039 | 27,99956 | 19,48602 | 24,2287869 |
| Downregulated | ENSMUSG00000028919 | 394,4383064 | 399,975239 | 393,962129 | 371,937026 | 675,544837 | 367,5704736 | 396,426712 | 310,44541 | 237,04265 | 600,833956 | 431,708304 | 209,3281 | 42,0737715 | 172,128 | 86,81795 | 82,03912116 | 56,25211 | 172,4081591 | 199,619 | 25,99959 | 86,80135 | 235,108969 |
| Downregulated | ENSMUSG00000032184 | 1001,429159 | 1190,35184 | 924,99403 | 1031,98071 | 1094,10275 | 1145,174436 | 1151,79566 | 636,779757 | 784,803369 | 1527,43426 | 1319,4154 | 434,4214 | 616,080226 | 476,1585 | 424,5318 | 453,9498037 | 241,5532 | 490,471487 | 399,2379 | 279,9956 | 545,6085 | 393,044765 |
| Downregulated | ENSMUSG00000006931 | 2363,457412 | 3593,39457 | 3386,80346 | 3650,95697 | 4232,65098 | 2977,718927 | 3916,55961 | 1210,00376 | 2414,20465 | 4011,67232 | 3513,28855 | 1273,485 | 2278,99596 | 1565,991 | 1101,553 | 1701,855991 | 572,4479 | 1725,072442 | 1500,944 | 533,325 | 1739,57 | 1785,75133 |
| Downregulated | ENSMUSG00000102692 | 8,459802819 | 14,8926951 | 24,5091647 | 26,9798879 | 44,5274375 | 54,40573796 | 57,9305511 | 7,33335613 | 17,084155 | 65,8004871 | 55,2056654 | 27,15134 | 21,0368858 | 47,7094 | 19,91237 | 6,380820534 | 1,654474 | 2,972554466 | 4,752832 | 5,999906 | 3,542912 | 28,7155993 |
| Downregulated | ENSMUSG00000015843 | 10,57475352 | 27,6578623 | 52,6493167 | 95,393175 | 207,370637 | 168,5250908 | 178,335226 | 67,2224312 | 117,453565 | 217,896695 | 294,798253 | 9,634347 | 13,022834 | 0 | 15,9299 | 12,76164107 | 12,40855 | 30,71639615 | 7,604532 | 5,999906 | 31,88621 | 52,9443862 |

| | | | | | | | | | | | | | | | | | | | | | | | |
|---|---|---|---|---|---|---|---|---|---|---|---|---|---|---|---|---|---|---|---|---|---|---|---|
| Downregulated | ENSMUSG00000034164 | 928,4633593 | 1653,08915 | 996,70603 | 3091,1243 | 3179,25903 | 2904,73562 | 2937,41971 | 2123,0066 | 2134,45161 | 4772,15336 | 4014,55599 | 242,6104 | 309,542748 | 258,192 | 168,0604 | 390,1415984 | 75,27855 | 1453,579134 | 487,6406 | 33,33281 | 581,0376 | 1800,10913 |
| Downregulated | ENSMUSG00000079363 | 58,16114438 | 143,608131 | 151,593722 | 498,164358 | 564,862349 | 737,7948856 | 122,676461 | 202,88952 | 320,327906 | 752,930163 | 718,777764 | 66,56458 | 21,0368858 | 84,19306 | 81,24248 | 35,55028583 | 38,88013 | 171,4173076 | 86,50155 | 10,6665 | 15,05738 | 93,3256977 |
| Downregulated | ENSMUSG00000058806 | 31,72426057 | 40,4230295 | 29,9556457 | 54,9233432 | 78,8771749 | 175,1599369 | 76,1048416 | 64,7779792 | 88,6240539 | 149,938815 | 143,53473 | 7,882647 | 13,022834 | 14,03218 | 1,59299 | 10,0270037 | 0 | 28,73469318 | 4,752832 | 0 | 0,885728 | 14,3577996 |
| Downregulated | ENSMUSG00000004035 | 42,29901409 | 52,1244328 | 37,2176204 | 77,0853939 | 77,6049624 | 122,0811681 | 60,2023374 | 45,2223628 | 77,9464571 | 144,545332 | 125,868917 | 17,51699 | 37,0649892 | 30,87079 | 7,96196 | 0,911545791 | 40,53461 | 28,73469318 | 21,86303 | 5,999906 | 23,02893 | 56,5338361 |
| Downregulated | ENSMUSG00000022510 | 66,6209472 | 52,1244328 | 137,97752 | 93,4660401 | 123,404612 | 175,1599369 | 115,861102 | 97,7780817 | 88,6240539 | 204,952337 | 112,619557 | 29,77889 | 74,1299784 | 57,06418 | 61,33011 | 35,55028583 | 18,19921 | 17,8353268 | 31,36869 | 11,33316 | 14,17165 | 56,5338361 |
| Downregulated | ENSMUSG00000028545 | 5,287376762 | 14,8926951 | 34,4943799 | 27,9434553 | 19,0831875 | 41,13604578 | 22,7178632 | 20,7778424 | 14,9486356 | 45,3052534 | 45,2686457 | 12,2619 | 11,0193211 | 3,741914 | 11,15093 | 13,67318686 | 1,654474 | 3,963405955 | 10,45623 | 5,999906 | 7,971552 | 8,97362477 |
| Downregulated | ENSMUSG00000028020 | 96,23025706 | 356,360918 | 139,793013 | 194,64062 | 268,436837 | 80,94512234 | 168,112187 | 110,000342 | 183,654666 | 347,340276 | 368,773845 | 79,70232 | 74,1299784 | 56,1287 | 9,557939 | 25,52328214 | 5,790658 | 48,55172295 | 16,15963 | 16,66641 | 14,17165 | 53,8417486 |
| Downregulated | ENSMUSG00000037624 | 127,9545176 | 399,975239 | 191,534583 | 478,89301 | 442,72995 | 399,4177348 | 354,398665 | 250,556334 | 224,229534 | 679,578801 | 732,027124 | 238,2311 | 77,1352478 | 107,58 | 121,8637 | 41,01956058 | 90,99605 | 151,6002778 | 97,90835 | 64,66566 | 60,22951 | 145,372721 |
| Downregulated | ENSMUSG00000063296 | 64,50599649 | 128,715436 | 19,9704305 | 91,5389053 | 89,0548749 | 43,78998421 | 93,143239 | 69,6668832 | 60,8623021 | 170,434048 | 178,866356 | 88,46082 | 42,0737715 | 65,48349 | 50,97568 | 54,69274744 | 33,08947 | 58,46023784 | 71,29249 | 12,66647 | 7,085824 | 57,4311986 |
| Downregulated | ENSMUSG00000020393 | 2360,284986 | 2233,90426 | 1944,39373 | 1395,24563 | 2769,60661 | 2007,704428 | 2543,26478 | 1658,56071 | 2223,07567 | 3606,08243 | 4038,84648 | 3262,54 | 1364,3923 | 2203,987 | 954,2009 | 979,9117249 | 842,1271 | 1061,201945 | 1032,315 | 505,9921 | 744,8973 | 989,790813 |
| Downregulated | ENSMUSG00000025813 | 63,44852114 | 94,6749901 | 84,4204561 | 47,2148038 | 85,2382374 | 99,5226914 | 99,9585979 | 59,8890751 | 105,708209 | 175,827531 | 136,91005 | 120,8673 | 55,0966056 | 46,77392 | 50,97568 | 47,40038111 | 9,099605 | 26,7529902 | 50,38002 | 8,666531 | 18,60029 | 44,8681239 |
| Downregulated | ENSMUSG00000035877 | 2403,641476 | 2517,92923 | 1998,85854 | 1216,02209 | 2438,83136 | 2665,88116 | 2473,9753 | 2182,89567 | 2230,54998 | 3723,66035 | 3623,69988 | 2259,692 | 1254,19909 | 840,9951 | 1076,861 | 1347,264679 | 789,1839 | 1184,067529 | 936,308 | 533,325 | 673,1533 | 1544,36082 |
| Downregulated | ENSMUSG00000026840 | 4575,695849 | 7945,25282 | 3112,66391 | 5410,43109 | 8855,8712 | 4471,886267 | 7237,9112 | 4285,12443 | 3895,18733 | 8502,28589 | 5955,58719 | 6528,584 | 1764,09313 | 1873,763 | 2992,432 | 3642,536979 | 1832,33 | 2271,031612 | 2176,797 | 38,66606 | 618,2382 | 3287,93612 |
| Downregulated | ENSMUSG00000051650 | 72,96579931 | 96,802518 | 96,221165 | 53,9597757 | 110,682487 | 62,36755328 | 101,094491 | 63,5557531 | 59,7945424 | 86,2957207 | 124,764804 | 77,07477 | 56,098362 | 49,58035 | 35,84227 | 39,196469 | 31,435 | 25,76213871 | 38,02266 | 19,33303 | 22,1432 | 60,123286 |
| Downregulated | ENSMUSG00000028838 | 29,60930986 | 36,1679738 | 13,6162026 | 22,1620508 | 26,7164625 | 11,94272297 | 45,4357263 | 24,4445204 | 18,1519147 | 58,2496115 | 48,5809856 | 9,634347 | 5,00878232 | 0 | 11,15093 | 7,292366325 | 26,47158 | 10,89936638 | 39,92379 | 0 | 3,542912 | 9,87098725 |
| Downregulated | ENSMUSG00000083289 | 28,55183451 | 30,8491541 | 16,3394431 | 15,4170788 | 16,5387625 | 23,88544594 | 30,6691153 | 11,0000342 | 18,1519147 | 21,5739302 | 37,5398525 | 9,634347 | 8,01405172 | 7,483827 | 4,77897 | 6,380820534 | 16,54474 | 8,917663399 | 15,20906 | 1,333313 | 4,42864 | 8,0762623 |
| Downregulated | ENSMUSG00000053025 | 453,6569261 | 757,399921 | 179,733874 | 289,070227 | 421,102337 | 355,6277506 | 295,332221 | 341,00106 | 249,855766 | 514,538235 | 722,090104 | 160,2805 | 38,0667457 | 47,7094 | 24,69134 | 76,56984641 | 8,272368 | 64,40534677 | 39,92379 | 1,333313 | 44,2864 | 266,516656 |
| Downregulated | ENSMUSG00000040848 | 950,6703417 | 1049,935 | 504,707243 | 582,958292 | 889,276537 | 731,1600395 | 546,364609 | 713,779997 | 571,251432 | 1133,71003 | 966,099145 | 340,7055 | 327,574364 | 175,8699 | 321,784 | 293,5177446 | 176,2014 | 341,8437636 | 225,2843 | 199,9969 | 302,919 | 432,528714 |
| Downregulated | ENSMUSG00000031137 | 242,1618557 | 327,639292 | 226,028963 | 230,292614 | 385,480387 | 252,1241515 | 203,324875 | 272,556403 | 256,262325 | 357,048545 | 388,647885 | 135,7567 | 76,1334913 | 40,22557 | 90,00393 | 93,88921644 | 53,77039 | 204,1154067 | 86,50155 | 12,66647 | 35,42912 | 174,985683 |
| Downregulated | ENSMUSG00000022883 | 1144,188331 | 1805,2074 | 1043,90887 | 979,94807 | 1412,15587 | 1015,131452 | 1334,67446 | 1405,55993 | 1304,80234 | 1700,0257 | 1807,43349 | 882,8565 | 349,613006 | 447,1587 | 689,7646 | 775,7254678 | 629,5272 | 757,0105375 | 577,9444 | 76,66547 | 148,8023 | 623,666922 |
| Downregulated | ENSMUSG00000054434 | 555,17456 | 491,458937 | 464,766382 | 370,973458 | 470,718625 | 313,1647356 | 385,067781 | 513,334929 | 530,676564 | 676,342711 | 602,845867 | 393,2565 | 329,577877 | 281,579 | 169,6534 | 231,5326308 | 161,3112 | 201,1428522 | 193,9156 | 201,3302 | 150,5738 | 223,443257 |
| Downregulated | ENSMUSG00000074909 | 109,9774366 | 88,2924065 | 61,7267851 | 75,1582591 | 83,9660249 | 86,25299921 | 76,1048416 | 57,444623 | 84,3530152 | 100,318775 | 111,515444 | 38,53739 | 67,1176831 | 16,83861 | 23,09835 | 21,87709898 | 39,70737 | 36,66150509 | 20,91246 | 25,99959 | 18,60029 | 36,7918616 |
| Downregulated | ENSMUSG00000007908 | 0 | 27,6578623 | 24,5091647 | 45,2876689 | 39,4385875 | 33,17423047 | 11,3589316 | 20,7778424 | 18,1519147 | 34,5182883 | 39,7480791 | 0 | 0 | 3,741914 | 3,18598 | 2,734637372 | 0 | 19,81702978 | 4,752832 | 23,99963 | 0,885728 | 13,4604372 |
| Downregulated | ENSMUSG00000039959 | 13,74717958 | 17,0202229 | 9,98521524 | 23,1256182 | 40,7108 | 15,92363062 | 7,95125211 | 1,22222602 | 9,60983717 | 15,1017511 | 24,2904928 | 1,751699 | 2,00351293 | 10,29026 | 4,77897 | 1,823091581 | 9,926842 | 12,88106935 | 0 | 6,666563 | 7,085824 | 18,844612 |
| Downregulated | ENSMUSG00000042834 | 1940,467272 | 3259,37269 | 1184,60963 | 6363,39927 | 2745,43457 | 1787,427537 | 3907,47246 | 3652,01135 | 1029,32034 | 3368,7692 | 2570,37578 | 536,02 | 680,192639 | 861,5756 | 989,2467 | 566,9814818 | 618,7732 | 747,1020226 | 1140,68 | 779,3212 | 2193,948 | 2361,85804 |
| Downregulated | ENSMUSG00000113913 | 14,80465493 | 78,7185311 | 34,4943799 | 158,025058 | 75,0605374 | 72,98330702 | 136,307179 | 111,222568 | 68,3366199 | 89,5318103 | 91,6414046 | 7,006798 | 27,0474245 | 38,35461 | 9,557939 | 17,31937002 | 8,272368 | 36,66150509 | 27,56643 | 7,333219 | 61,11523 | 73,5837231 |
| Downregulated | ENSMUSG00000025038 | 6,344852114 | 14,8926951 | 4,5387342 | 25,052753 | 12,722125 | 2,653938437 | 34,0767947 | 13,4444862 | 14,9486356 | 11,8656616 | 14,353473 | 0,87585 | 1,00175646 | 0 | 0,796495 | 5,469274744 | 3,308947 | 7,92681191 | 9,505665 | 1,999969 | 9,743008 | 17,0498871 |
| Downregulated | ENSMUSG00000074575 | 44,4139648 | 46,8056131 | 14,5239494 | 126,227333 | 99,2325749 | 37,15513812 | 114,725209 | 75,7780133 | 26,6939921 | 156,410994 | 8,83290647 | 19,26869 | 1,00175646 | 12,16122 | 12,74392 | 16,40782423 | 28,12605 | 3,963405955 | 23,76416 | 9,999844 | 11,51446 | 31,4076867 |
| Downregulated | ENSMUSG00000040310 | 7,402327466 | 7,44634754 | 9,98521524 | 43,3605341 | 20,3554 | 10,61575375 | 31,8050084 | 12,2222602 | 4,27013874 | 24,8100197 | 13,2493597 | 12,2619 | 4,00702586 | 4,677392 | 7,964949 | 9,115457906 | 6,617895 | 6,935960422 | 0 | 0 | 2,657184 | 0 |
| Downregulated | ENSMUSG00000078850 | 27,49435916 | 11,7014033 | 20,8781773 | 21,1984833 | 22,899825 | 10,61575375 | 23,8537563 | 9,77780817 | 5,33879843 | 12,9443581 | 29,8110593 | 10,5102 | 4,00702586 | 2,806435 | 11,15093 | 3,646183163 | 0,827237 | 19,81702978 | 4,752832 | 17,33306 | 9,743008 | 18,844612 |
| Downregulated | ENSMUSG00000075592 | 604,8759015 | 273,387331 | 648,131244 | 310,268711 | 471,990837 | 269,3747514 | 316,914191 | 369,112259 | 278,685278 | 459,524713 | 407,417811 | 182,1767 | 48,0843103 | 104,7736 | 181,6008 | 160,4320592 | 203,5003 | 202,1337037 | 204,3718 | 429,3266 | 190,4315 | 244,082594 |
| Downregulated | ENSMUSG00000079495 | 48,64386621 | 24,4665705 | 49,9260762 | 21,1984833 | 30,5331 | 9,28878453 | 22,7178632 | 25,6667465 | 17,084155 | 25,8887162 | 38,6439658 | 6,130948 | 4,00702586 | 8,419306 | 6,371966 | 0,911545791 | 3,308947 | 13,87192084 | 12,35736 | 11,33281 | 11,51446 | 15,2551621 |
| Downregulated | ENSMUSG00000002346 | 210,4375951 | 117,014033 | 223,305723 | 92,5024727 | 204,826212 | 179,1408445 | 136,307179 | 162,556061 | 73,6754183 | 161,804476 | 266,091307 | 95,46762 | 44,0772844 | 28,06435 | 70,88805 | 40,10801479 | 66,17895 | 67,37790124 | 93,15552 | 101,3318 | 67,31533 | 111,272947 |
| Downregulated | ENSMUSG00000029720 | 28,55183451 | 6,3825836 | 26,3246584 | 4,81783712 | 24,1720375 | 14,5966614 | 34,0767947 | 4,88890409 | 11,7453565 | 15,1017511 | 38,6439658 | 5,255098 | 2,00351293 | 2,806435 | 3,18598 | 5,469274744 | 18,19921 | 15,85362382 | 4,752832 | 8,666531 | 7,971552 | 7,17889982 |
| Downregulated | ENSMUSG00000107068 | 33,83921127 | 5,31881967 | 23,6014178 | 7,70853939 | 19,0831875 | 15,92363062 | 34,0767947 | 7,33335613 | 6,40655811 | 23,7313232 | 14,353473 | 3,503399 | 3,00526939 | 0 | 2,389485 | 3,646183163 | 5,790658 | 13,87192084 | 12,35736 | 8,666531 | 2,657184 | 10,7683497 |
| Downregulated | ENSMUSG00000040093 | 54,98871832 | 25,5303344 | 43,5718483 | 14,4535114 | 34,3497375 | 55,73270718 | 55,6587648 | 11,0000342 | 32,0327906 | 48,5413429 | 37,5398525 | 0 | 9,01580818 | 0 | 7,964949 | 0 | 8,272368 | 12,88106935 | 13,30793 | 27,33291 | 10,62874 | 18,844612 |
| Downregulated | ENSMUSG00000000223 | 134,2993697 | 107,440157 | 106,20638 | 174,405704 | 111,9547 | 191,0835675 | 195,373623 | 173,556095 | 116,385806 | 228,68366 | 143,53473 | 55,17853 | 40,0702586 | 9,354784 | 66,90558 | 49,22347269 | 75,27855 | 44,588317 | 43,72606 | 41,33269 | 58,45805 | 113,965035 |
| Downregulated | ENSMUSG00000019990 | 69,79337325 | 84,0373508 | 90,774684 | 154,170788 | 96,6881499 | 136,6778295 | 126,084141 | 91,6669516 | 87,5562942 | 228,68366 | 114,827784 | 17,51699 | 6,01053879 | 4,677392 | 40,62124 | 10,93854949 | 42,18908 | 51,52427742 | 25,6653 | 29,99953 | 42,51494 | 44,8681239 |
| Downregulated | ENSMUSG00000003273 | 101,5176338 | 143,608131 | 167,025419 | 224,51121 | 175,565325 | 207,0071981 | 232,858097 | 129,555958 | 163,367232 | 230,841053 | 158,992316 | 14,0136 | 34,0597198 | 42,09653 | 34,24928 | 30,99255688 | 15,7175 | 53,5059804 | 35,17096 | 15,33309 | 39,85776 | 38,5865865 |
| Downregulated | ENSMUSG00000040364 | 133,2418944 | 253,175816 | 178,826127 | 307,378008 | 213,7317 | 398,0907656 | 412,329216 | 122,222602 | 124,927883 | 396,960315 | 235,176135 | 53,42683 | 30,0526939 | 75,77375 | 29,47031 | 15,49627844 | 19,02645 | 96,11259441 | 38,02266 | 22,66631 | 10,62874 | 78,967898 |
| Downregulated | ENSMUSG00000039385 | 295,0356233 | 554,22101 | 347,66704 | 506,836465 | 427,4634 | 493,6325493 | 723,563942 | 204,111746 | 171,909309 | 1112,1361 | 421,771284 | 179,5492 | 75,1317348 | 130,0315 | 169,6534 | 134,908777 | 68,66066 | 140,7009114 | 143,5355 | 40,66603 | 40,74349 | 264,721931 |
| Downregulated | ENSMUSG00000117916 | 32,78173592 | 75,5272393 | 27,2324052 | 59,7411803 | 61,0661999 | 38,48210734 | 72,6971621 | 36,6667807 | 11,7453565 | 110,027044 | 32,019286 | 0 | 0 | 0 | 0 | 0 | 0 | 10,89936638 | 5,703399 | 0 | 9,743008 | 29,6129618 |
| Downregulated | ENSMUSG00000062296 | 106,8050106 | 195,732564 | 97,1289119 | 192,713485 | 253,170287 | 258,7589976 | 295,332221 | 110,000342 | 133,469961 | 475,705161 | 196,532169 | 75,32307 | 42,0737715 | 8,419306 | 41,41774 | 29,1694653 | 56,25211 | 75,30471315 | 42,77549 | 0 | 65,54387 | 185,754033 |
| Downregulated | ENSMUSG00000060548 | 440,9672219 | 680,808918 | 574,60375 | 829,631552 | 1061,02522 | 804,1433465 | 1159,74691 | 723,557805 | 609,690781 | 1630,98912 | 847,959021 | 432,6697 | 93,1633512 | 120,6767 | 271,6048 | 302,6332025 | 157,175 | 458,7642393 | 174,9042 | 9,333188 | 89,45853 | 554,570011 |
| Downregulated | ENSMUSG00000037166 | 233,7020529 | 534,009495 | 421,194534 | 1432,82476 | 1086,46947 | 1319,007403 | 1470,98164 | 915,44729 | 635,317013 | 814,415864 | 588,492394 | 49,04758 | 71,124709 | 86,99949 | 45,40021 | 56,51583902 | 76,10579 | 298,2462981 | 207,2235 | 1,333313 | 161,2025 | 420,863002 |
| Downregulated | ENSMUSG00000085129 | 6,344852114 | 5,31881967 | 5,44648104 | 23,1256182 | 19,0831875 | 23,88544594 | 22,7178632 | 3,66667807 | 13,8808759 | 19,4165372 | 6,62467985 | 0 | 0 | 1,870957 | 0 | 1,823091581 | 0,827237 | 2,972554466 | 0,950566 | 0 | 0 | 7,17889982 |
| Downregulated | ENSMUSG00000040860 | 37,01163733 | 45,7418492 | 58,0957978 | 114,664523 | 94,1437249 | 91,56087608 | 79,5125211 | 56,222397 | 83,2852555 | 80,9022382 | 59,6221187 | 30,65474 | 6,01053879 | 20,58052 | 8,761444 | 4,557728953 | 20,68092 | 13,87192084 | 4,752832 | 19,33303 | 15,05738 | 18,844612 |
| Downregulated | ENSMUSG00000045875 | 113,1498627 | 139,353075 | 43,5718483 | 107,919551 | 141,215587 | 116,7732912 | 223,770952 | 105,111438 | 107,843728 | 226,526267 | 114,827784 | 43,79248 | 9,01580818 | 5,61287 | 4,77897 | 10,0270037 | 10,75408 | 51,52427742 | 30,41813 | 0 | 22,1432 | 55,6364736 |
| Downregulated | ENSMUSG00000073910 | 98,34520777 | 135,09802 | 76,2507346 | 128,154467 | 255,714712 | 149,9475217 | 235,129884 | 102,666986 | 114,250286 | 241,628018 | 150,15941 | 18,39284 | 20,0351293 | 29,93531 | 28,67382 | 19,1424616 | 65,35171 | 50,53342593 | 24,71473 | 11,99981 | 58,45805 | 72,6863607 |
| Downregulated | ENSMUSG00000020785 | 705,33606 | 658,469875 | 684,441117 | 589,703263 | 1260,76259 | 769,6421468 | 1343,76161 | 837,224825 | 812,565121 | 1250,20925 | 817,043849 | 300,4164 | 208,365345 | 193,644 | 284,3487 | 269,8717554 | 248,9983 | 276,4475654 | 363,1164 | 212,6633 | 403,0062 | 658,664058 |
| Downregulated | ENSMUSG00000019102 | 8440,768262 | 14838,4431 | 6806,28581 | 10275,483 | 17331,3509 | 15252,1842 | 17579,0825 | 10676,1443 | 11564,9052 | 10200,1542 | 6445,8135 | 2455,883 | 2102,68682 | 1412,572 | 1719,633 | 3011,747292 | 567,4845 | 3862,339103 | 1434,405 | 73,99885 | 995,5583 | 3372,28819 |
| Downregulated | ENSMUSG00000044555 | 1223,498983 | 1957,32564 | 1154,65398 | 1504,12875 | 2101,69505 | 1798,043291 | 2120,71253 | 1389,67099 | 1703,0767 | 1602,94301 | 1232,19045 | 258,3757 | 431,757036 | 272,2242 | 270,8083 | 450,3036206 | 274,6426 | 855,1048348 | 606,4614 | 21,99966 | 263,0612 | 952,998951 |
| Downregulated | ENSMUSG00000036972 | 56,04619367 | 96,802518 | 61,7267851 | 67,4497197 | 106,86585 | 84,92602999 | 116,996995 | 75,7780133 | 79,0142167 | 98,1613823 | 40,8521924 | 58,68193 | 19,0333728 | 24,32244 | 23,09835 | 21,87709898 | 29,78053 | 44,588317 | 33,26983 | 28,66622 | 15,9431 | 28,7155993 |
| Downregulated | ENSMUSG00000047182 | 26,43688381 | 29,7853902 | 33,5866331 | 37,5791295 | 31,8053125 | 27,86635359 | 35,2126879 | 8,55558215 | 38,4393487 | 32,3608953 | 24,2904928 | 1,751699 | 14,0245905 | 0,935478 | 18,31938 | 6,380820534 | 7,445132 | 11,89021787 | 8,555098 | 0 | 22,1432 | 26,9208743 |
| Downregulated | ENSMUSG00000035642 | 417,7027642 | 442,525797 | 389,423394 | 504,90933 | 400,746937 | 453,8234728 | 339,632054 | 481,557053 | 365,173812 | 483,256036 | 342,275126 | 86,70912 | 166,291573 | 136,5798 | 140,9796 | 167,7244255 | 140,6303 | 280,4109713 | 228,136 | 189,3304 | 350,7483 | 258,440394 |
| Downregulated | ENSMUSG00000078640 | 17,97708099 | 29,7853902 | 24,5091647 | 34,6884273 | 16,5387625 | 39,80970576 | 12,4948247 | 26,8889725 | 19,2196743 | 28,0461092 | 29,8110593 | 0 | 3,00526939 | 0 | 0 | 0 | 2,481711 | 11,89021787 | 4,752832 | 5,999906 | 15,05738 | 8,97362477 |
| Downregulated | ENSMUSG00000044328 | 1190,717247 | 1149,92881 | 1234,5357 | 1627,46538 | 1558,46031 | 1522,033694 | 1601,60935 | 1290,67068 | 1395,56191 | 1501,54554 | 1175,88067 | 349,464 | 492,864181 | 428,4491 | 348,8648 | 401,0801479 | 673,3708 | 904,6474093 | 663,4954 | 667,9896 | 1017,701 | 741,221406 |
| Downregulated | ENSMUSG00000096727 | 543,5423311 | 468,056131 | 538,293876 | 716,894163 | 480,896325 | 679,4082399 | 512,287814 | 387,445649 | 519,998967 | 473,547768 | 442,749437 | 67,44043 | 222,389935 | 140,3218 | 169,6534 | 162,2551507 | 261,4068 | 330,9443973 | 231,9382 | 137,3312 | 341,891 | 232,416882 |
| Downregulated | ENSMUSG00000036983 | 54,98871832 | 46,8056131 | 34,4943799 | 64,5590174 | 48,344075 | 74,31027624 | 67,0176963 | 31,7778766 | 33,1005503 | 48,5413429 | 43,060419 | 9,634347 | 32,0578232 | 22,45148 | 29,47031 | 10,93854949 | 24,81711 | 21,79873275 | 10,45623 | 33,33281 | 30,11475 | 18,844612 |
| Downregulated | ENSMUSG00000044345 | 958,0726692 | 571,241233 | 501,984002 | 729,42054 | 690,811387 | 1003,188729 | 696,302506 | 548,779484 | 652,401168 | 607,306135 | 701,111951 | 120,8673 | 104,182672 | 253,5146 | 229,3905 | 177,7514292 | 112,5042 | 260,5939416 | 186,311 | 175,3306 | 379,9773 | 420,863002 |
| Downregulated | ENSMUSG00000013419 | 521,3353487 | 390,401364 | 344,036052 | 594,521101 | 505,068362 | 832,0097001 | 698,574292 | 638,001983 | 565,912633 | 506,987359 | 444,957663 | 177,7975 | 194,340754 | 266,6113 | 273,1978 | 170,4590628 | 130,7034 | 294,2828922 | 210,0752 | 243,3295 | 380,863 | 370,610703 |
| Downregulated | ENSMUSG00000027339 | 1336,648845 | 714,849364 | 1361,62026 | 976,093801 | 1045,75867 | 873,1457459 | 1156,33924 | 1366,44869 | 718,602268 | 1116,45089 | 794,961582 | 308,2991 | 399,700829 | 214,2246 | 192,7518 | 243,3827261 | 162,1384 | 389,4046351 | 423,9527 | 523,3252 | 675,8105 | 715,197895 |
| Downregulated | ENSMUSG00000097325 | 148,0465493 | 102,121338 | 172,4719 | 87,6846356 | 106,86585 | 148,6205525 | 146,530217 | 129,555958 | 143,079798 | 108,948347 | 80,6002715 | 14,88944 | 45,0790409 | 16,83861 | 22,30186 | 26,43482793 | 9,926842 | 38,64320806 | 25,6653 | 23,33297 | 81,48698 | 65,5074609 |
| Downregulated | ENSMUSG00000039706 | 435,6798452 | 376,572433 | 643,59251 | 299,669469 | 412,19685 | 444,5346882 | 391,88314 | 455,890306 | 443,12027 | 517,774324 | 276,028327 | 86,70912 | 228,400474 | 152,483 | 78,05651 | 80,21602958 | 87,68711 | 247,7128722 | 187,2616 | 301,9953 | 341,0053 | 266,516656 |
| Downregulated | ENSMUSG00000025262 | 121,6096655 | 104,248866 | 109,837368 | 81,903231 | 155,209925 | 75,63724546 | 111,31753 | 90,4447256 | 76,8786974 | 118,656616 | 50,7892122 | 26,27549 | 21,0368858 | 18,70957 | 26,28433 | 17,31937002 | 42,18908 | 32,69809913 | 41,82493 | 47,99925 | 48,71504 | 45,7654863 |
| Downregulated | ENSMUSG00000040990 | 2064,191888 | 2271,136 | 2234,87272 | 1560,01566 | 2102,96726 | 1681,27 | 2168,42004 | 1994,67287 | 2094,9445 | 2360,18796 | 1538,02984 | 659,5148 | 894,568523 | 499,5455 | 657,9048 | 577,0084855 | 776,7754 | 912,5742212 | 1053,228 | 1081,983 | 1053,131 | 1197,97891 |
| Downregulated | ENSMUSG00000043251 | 29,60930986 | 14,8926951 | 21,7859242 | 15,4170788 | 24,1720375 | 15,92363062 | 23,8537563 | 22,0000684 | 22,4229534 | 22,6526267 | 7,72879316 | 2,627549 | 11,0193211 | 1,870957 | 8,761444 | 0,911545791 | 6,617895 | 15,85362382 | 11,4068 | 17,33306 | 9,743008 | 9,87098725 |
| Downregulated | ENSMUSG00000026447 | 80,36812678 | 76,5910032 | 194,257824 | 117,555226 | 223,9094 | 128,7160142 | 173,791653 | 193,111711 | 238,11041 | 245,942804 | 117,036011 | 30,65474 | 16,0281034 | 53,32227 | 16,72639 | 10,93854949 | 31,435 | 51,52427742 | 20,91246 | 29,33288 | 77,94407 | 70,8916357 |
| Downregulated | ENSMUSG00000061535 | 1506,902377 | 2126,4641 | 2318,38543 | 1730,56709 | 3227,60311 | 3205,957632 | 4686,69517 | 3548,12214 | 4802,78307 | 3583,4298 | 2473,21381 | 179,5492 | 518,909849 | 311,5143 | 352,0508 | 226,0633561 | 406,1733 | 643,0626162 | 594,1041 | 277,329 | 762,6118 | 874,031053 |
| Downregulated | ENSMUSG00000026185 | 11298,06666 | 9507,92204 | 18121,3502 | 16362,3384 | 13812,4111 | 19291,4785 | 25833,6181 | 29307,7578 | 26752,7189 | 26904,8483 | 17624,9607 | 2201,01 | 2199,8572 | 2709,145 | 1675,029 | 3068,263131 | 1314,479 | 4276,515026 | 3380,214 | 3522,612 | 3251,508 | 6172,05912 |
| Downregulated | ENSMUSG00000079481 | 274,9435916 | 280,833679 | 234,198685 | 206,203429 | 239,17595 | 283,9714128 | 372,572956 | 248,111882 | 317,124627 | 372,150296 | 178,866356 | 80,57817 | 59,1036314 | 110,3864 | 68,49857 | 81,12757537 | 55,42487 | 135,746654 | 135,931 | 146,6644 | 89,45853 | 195,62502 |
| Downregulated | ENSMUSG00000085055 | 30,66678522 | 17,0202229 | 19,9704305 | 24,0891656 | 27,988675 | 30,52029203 | 36,3485811 | 30,5556505 | 32,0327906 | 32,3608953 | 18,7691862 | 2,627549 | 7,01229525 | 0,935478 | 4,77897 | 0 | 2,481711 | 17,8353268 | 0,950566 | 15,33309 | 6,200096 | 8,97362477 |
| Downregulated | ENSMUSG00000036564 | 4663,466304 | 5961,33309 | 7089,50282 | 4971,04434 | 6041,73716 | 6008,516622 | 8578,26513 | 9494,25174 | 6095,84005 | 7855,06798 | 6024,04221 | 2067,005 | 1099,9286 | 1123,51 | 1260,055 | 1328,122217 | 1935,734 | 2984,444684 | 3469,568 | 3459,279 | 6828,077 | 4878,06243 |
| Downregulated | ENSMUSG00000038074 | 1442,396381 | 2192,41747 | 2420,96082 | 3039,09166 | 2634,75208 | 1888,277198 | 3355,42839 | 4386,56919 | 3033,50527 | 3763,57212 | 2644,35137 | 479,0898 | 929,629999 | 597,7707 | 476,304 | 510,4656428 | 954,6313 | 1629,950699 | 1202,467 | 1705,307 | 2821,044 | 2477,6178 |
| Downregulated | ENSMUSG00000029185 | 3802,681367 | 6058,1356 | 5040,7182 | 5297,6937 | 6310,17399 | 5176,506922 | 5711,2708 | 7984,8026 | 6104,38212 | 6729,98752 | 5284,2863 | 1368,953 | 2572,5106 | 1384,508 | 908,8007 | 1077,447125 | 2277,383 | 3371,867616 | 2860,255 | 4132,602 | 5652,716 | 4919,3411 |
| Downregulated | ENSMUSG00000035000 | 4160,108036 | 7605,91213 | 5330,28944 | 3778,14787 | 7927,15608 | 5749,757624 | 8724,79535 | 7780,69085 | 6497,31769 | 9076,15243 | 6064,89441 | 1339,174 | 1527,67861 | 621,1577 | 761,4492 | 976,2655418 | 1738,025 | 2400,833157 | 2863,106 | 3763,941 | 4962,734 | 5116,76085 |
| Downregulated | ENSMUSG00000053460 | 2343,365381 | 2864,71627 | 3453,06898 | 2680,64457 | 2811,58962 | 2293,00281 | 3612,14024 | 3362,34379 | 2662,99266 | 4504,63662 | 2909,33857 | 994,9652 | 1125,97427 | 636,1253 | 829,9477 | 808,5411163 | 1075,408 | 1583,380679 | 1241,44 | 1460,644 | 2073,489 | 1809,98012 |
| Downregulated | ENSMUSG00000018906 | 7173,91279 | 6830,42822 | 8408,45898 | 7812,60467 | 9587,39339 | 8310,808216 | 11798,5222 | 11107,5901 | 8322,11899 | 11202,2632 | 7760,81245 | 1654,48 | 1942,40578 | 1512,669 | 1181,202 | 1451,180899 | 2021,767 | 3551,211736 | 2908,733 | 2971,287 | 6020,293 | 5241,49423 |
| Downregulated | ENSMUSG00000046020 | 946,4404403 | 893,561705 | 1078,40325 | 901,899109 | 1166,61886 | 777,6039621 | 1449,39967 | 1313,89297 | 1023,98154 | 1336,50497 | 845,750795 | 394,1324 | 327,574364 | 311,5143 | 311,4295 | 317,2179351 | 541,0129 | 493,4440414 | 577,9444 | 940,652 | 849,4132 | 606,617035 |
| Downregulated | ENSMUSG00000052369 | 730,7154685 | 889,306649 | 823,326384 | 939,478238 | 1049,57531 | 800,1624388 | 1013,2167 | 906,891708 | 896,918136 | 1036,50735 | 856,791928 | 218,9624 | 510,895797 | 251,6437 | 229,3905 | 260,7020961 | 514,5413 | 726,2941413 | 557,032 | 751,9883 | 677,5819 | 641,614171 |
| Downregulated | ENSMUSG00000040537 | 213,6100212 | 573,36876 | 252,353622 | 453,840257 | 620,839699 | 370,224412 | 688,351254 | 645,335339 | 526,405525 | 580,338722 | 322,401086 | 86,70912 | 73,1282219 | 10,29026 | 43,80722 | 44,66574374 | 150,5571 | 347,7888726 | 291,8239 | 146,6644 | 302,0333 | 358,047628 |
| Downregulated | ENSMUSG00000031659 | 2204,83611 | 3122,14715 | 1820,03241 | 3287,69205 | 3685,59961 | 3428,888461 | 3531,49183 | 3270,67683 | 2709,97408 | 3969,60315 | 2249,07881 | 945,9177 | 661,159267 | 577,1902 | 1297,49 | 1092,943403 | 1267,327 | 1930,1787 | 1470,526 | 1334,646 | 2808,644 | 2142,00423 |
| Downregulated | ENSMUSG00000027217 | 1530,166835 | 1822,22762 | 1509,58299 | 1903,04566 | 2325,60445 | 1699,847569 | 2726,14358 | 1972,6728 | 2578,63964 | 3154,10859 | 1594,33962 | 555,2887 | 470,825538 | 398,5138 | 297,8891 | 426,60343 | 280,4333 | 1228,655846 | 923,0001 | 313,9951 | 1436,651 | 1924,84251 |
| Downregulated | ENSMUSG00000002475 | 193,5179895 | 186,158688 | 210,597267 | 170,551434 | 183,1986 | 103,5035991 | 166,976294 | 226,111814 | 182,586906 | 170,434048 | 139,118277 | 27,15134 | 61,1071443 | 52,38679 | 32,65629 | 52,86965586 | 105,0591 | 125,8381391 | 110,2657 | 74,6655 | 89,45853 | 128,322834 |
| Downregulated | ENSMUSG00000062646 | 282,3459191 | 261,685928 | 248,722634 | 230,292614 | 198,46515 | 225,5847672 | 197,64541 | 349,556642 | 287,227355 | 254,572376 | 189,907489 | 95,46762 | 62,1089008 | 55,19322 | 75,66702 | 118,5009528 | 158,8295 | 137,7283569 | 181,5582 | 153,3309 | 169,1741 | 186,651395 |
| Downregulated | ENSMUSG00000041828 | 1732,144627 | 2630,68821 | 1735,61196 | 2178,62595 | 1507,57181 | 1354,835572 | 1299,46177 | 2795,23091 | 1838,68218 | 2564,0616 | 1621,94245 | 587,6951 | 497,872963 | 164,6442 | 273,9943 | 374,64532 | 361,5025 | 794,662894 | 518,0587 | 566,6578 | 997,3297 | 799,549967 |
| Downregulated | ENSMUSG00000041797 | 1687,730662 | 2487,08008 | 1816,40143 | 2062,99785 | 1666,59837 | 1422,511002 | 1711,79099 | 2218,34023 | 1487,38924 | 2619,07512 | 1570,04913 | 773,3753 | 668,171562 | 285,3209 | 264,4363 | 403,8147853 | 593,1288 | 1425,835292 | 604,5603 | 914,6524 | 1025,673 | 1241,05231 |

| | | | | | | | | | | | | | | | | | | | | | | | |
|---|---|---|---|---|---|---|---|---|---|---|---|---|---|---|---|---|---|---|---|---|---|---|---|
| Downregulated | ENSMUSG00000041762 | 325,7024085 | 440,398269 | 265,969824 | 244,746126 | 320,59755 | 173,8329676 | 313,506512 | 333,667704 | 395,071084 | 404,511191 | 337,858672 | 86,70912 | 75,1317348 | 64,54801 | 98,76537 | 116,6778612 | 129,0489 | 212,0422186 | 123,5736 | 109,9983 | 148,8023 | 161,525246 |
| Downregulated | ENSMUSG00000070436 | 17451,51574 | 14299,1148 | 37853,951 | 22306,5859 | 15920,4672 | 18606,76238 | 27406,8301 | 17970,3892 | 11446,3838 | 14785,6931 | 20505,5924 | 3576,094 | 9921,39603 | 3641,817 | 7109,514 | 7679,773286 | 7702,402 | 9542,890689 | 5786,098 | 11465,82 | 7254,998 | 7677,83336 |
| Downregulated | ENSMUSG00000042116 | 380,6911268 | 297,853902 | 553,725572 | 556,941971 | 394,385875 | 398,0907656 | 308,962939 | 255,445239 | 277,617518 | 358,127241 | 319,088746 | 157,6529 | 258,453168 | 55,19322 | 86,81795 | 87,5083959 | 128,2217 | 166,4630501 | 137,8321 | 129,3313 | 305,5762 | 139,091184 |
| Downregulated | ENSMUSG00000022512 | 133,2418944 | 140,416839 | 317,711394 | 346,884273 | 118,315762 | 232,2196133 | 180,607012 | 191,889485 | 139,876519 | 187,693193 | 144,638843 | 52,55098 | 112,196724 | 54,25775 | 54,16166 | 32,81564846 | 30,60776 | 120,8838816 | 30,41813 | 14,66644 | 92,11571 | 68,1995483 |
| Downregulated | ENSMUSG00000032719 | 27,49435916 | 26,5940983 | 52,6493167 | 51,0690735 | 22,899825 | 27,86635359 | 42,0280469 | 18,3333903 | 13,8808759 | 47,4626464 | 26,4987194 | 17,51699 | 18,0316164 | 0 | 7,168455 | 14,58473265 | 11,58132 | 5,945108933 | 19,9619 | 6,666563 | 7,971552 | 11,6657122 |
| Downregulated | ENSMUSG00000014837 | 82,48307748 | 36,1679738 | 75,3429877 | 78,0489613 | 48,344075 | 66,34846093 | 94,2791321 | 77,0002394 | 25,6262325 | 131,600974 | 96,0578579 | 21,02039 | 20,0351293 | 9,354784 | 25,48784 | 15,49627844 | 48,80697 | 39,63405955 | 61,78682 | 29,99953 | 49,60077 | 49,3549363 |
| Downregulated | ENSMUSG00000022099 | 99,40268312 | 60,6345442 | 103,48314 | 78,0489613 | 58,5217749 | 131,3699526 | 113,589316 | 83,1113695 | 80,0819764 | 161,804476 | 114,827784 | 17,51699 | 23,0403987 | 4,677392 | 20,70887 | 26,43482793 | 52,94316 | 34,67980211 | 51,33059 | 69,33225 | 69,08679 | 83,4547104 |
| Downregulated | ENSMUSG00000046275 | 203,0352676 | 97,8662819 | 211,505014 | 196,567754 | 108,138062 | 188,429629 | 199,917196 | 140,555993 | 158,028433 | 228,68366 | 283,75712 | 53,42683 | 20,0351293 | 34,6127 | 85,22496 | 75,65830062 | 86,85987 | 65,39619826 | 38,97323 | 28,66622 | 26,57184 | 158,833159 |
| Downregulated | ENSMUSG00000038119 | 902,0264755 | 849,947383 | 1121,06735 | 1809,57962 | 999,959024 | 1080,152944 | 1133,62137 | 1329,78191 | 1107,26679 | 1828,39058 | 977,140278 | 400,2633 | 233,409256 | 524,8034 | 360,8122 | 339,0950341 | 600,5739 | 810,5165178 | 672,0505 | 528,6584 | 833,4701 | 761,860743 |
| Downregulated | ENSMUSG00000043496 | 763,4972044 | 430,824393 | 831,496105 | 903,826244 | 632,289612 | 630,3103788 | 673,584643 | 958,225201 | 628,910455 | 1303,06538 | 490,226309 | 52,55098 | 77,1352478 | 50,51583 | 86,02145 | 65,63129693 | 40,53461 | 214,0239216 | 280,4171 | 241,9962 | 434,8925 | 483,678375 |
| Downregulated | ENSMUSG00000049872 | 122,6671409 | 159,56459 | 178,826127 | 289,070227 | 134,854525 | 94,21481452 | 228,314525 | 235,889622 | 197,535542 | 335,474614 | 172,241676 | 35,90984 | 38,0667457 | 50,51583 | 60,53362 | 72,92366325 | 49,63421 | 81,24982208 | 111,2163 | 111,3316 | 84,14416 | 114,862397 |
| Downregulated | ENSMUSG00000034881 | 265,4263134 | 192,541272 | 322,250128 | 365,192054 | 138,671162 | 177,8138753 | 353,262772 | 283,556437 | 201,806581 | 310,664595 | 161,200543 | 10,5102 | 4,00702586 | 22,45148 | 9,557939 | 28,25791951 | 62,04276 | 117,9113272 | 138,7827 | 133,9979 | 44,2864 | 85,2494354 |
| Downregulated | ENSMUSG00000026080 | 204,092743 | 165,947174 | 161,578938 | 254,3818 | 145,032225 | 199,0453828 | 202,188982 | 213,889554 | 96,0983717 | 224,368874 | 75,079705 | 55,17853 | 44,0772844 | 43,03201 | 95,57939 | 97,5353996 | 74,45132 | 71,34130719 | 116,9197 | 69,33225 | 63,77242 | 104,094047 |
| Downregulated | ENSMUSG00000032593 | 222,069824 | 188,286216 | 348,574787 | 372,900593 | 253,170287 | 406,0525809 | 354,398665 | 374,001163 | 179,383627 | 439,029479 | 128,077144 | 59,55778 | 38,0667457 | 57,06418 | 46,9932 | 46,48883532 | 52,11592 | 54,49683188 | 59,88569 | 33,33281 | 54,91514 | 95,1204226 |
| Downregulated | ENSMUSG00000028226 | 820,6008734 | 903,13558 | 1127,42158 | 1100,394 | 675,544837 | 1019,11236 | 1417,59466 | 1196,55928 | 698,314834 | 1348,37064 | 765,150523 | 412,5252 | 241,423308 | 334,9013 | 502,5883 | 605,266405 | 360,6753 | 491,4623385 | 367,8692 | 296,662 | 193,0887 | 482,781013 |
| Downregulated | ENSMUSG00000049336 | 10,57475352 | 46,8056131 | 66,2655193 | 64,5590174 | 41,9830125 | 27,86635359 | 27,2614358 | 50,1112669 | 41,6426277 | 11,8656616 | 76,1838183 | 18,39284 | 21,0368858 | 6,548349 | 21,50536 | 2,734637372 | 38,05289 | 19,81702978 | 29,46756 | 1,333313 | 28,3433 | 34,9971366 |
| Downregulated | ENSMUSG00000029417 | 20,09203169 | 31,912918 | 77,1584814 | 101,17458 | 24,1720375 | 53,07876874 | 61,3382305 | 35,4445546 | 48,0491859 | 32,3608953 | 86,1208381 | 8,758497 | 13,022834 | 63,61253 | 0 | 16,40782423 | 18,19921 | 11,89021787 | 8,555098 | 5,33325 | 68,20106 | 27,8182368 |
| Downregulated | ENSMUSG00000026839 | 13,74717958 | 25,5303344 | 30,8633926 | 15,4170788 | 10,1777 | 17,25059984 | 10,2230384 | 8,55558215 | 8,54207749 | 22,6526267 | 43,060419 | 10,5102 | 6,01053879 | 1,870957 | 0,796495 | 5,469274744 | 4,136184 | 20,80788127 | 10,45623 | 4,666594 | 13,28592 | 11,6657122 |
| Downregulated | ENSMUSG00000026697 | 152,2764507 | 781,866491 | 311,357166 | 354,592812 | 115,771337 | 252,1241515 | 103,366277 | 176,000547 | 222,094015 | 206,031033 | 527,766162 | 106,8537 | 284,498836 | 30,87079 | 54,16166 | 49,22347269 | 28,95329 | 97,1034459 | 169,2008 | 11,99981 | 221,432 | 300,61643 |
| Downregulated | ENSMUSG00000063558 | 203,0352676 | 253,175816 | 142,516254 | 163,806462 | 173,0209 | 256,1050592 | 251,032388 | 125,88928 | 106,775969 | 188,771889 | 241,800815 | 59,55778 | 74,1299784 | 97,28975 | 41,41774 | 30,99255688 | 110,8497 | 175,3807135 | 157,794 | 55,99913 | 145,2594 | 289,84808 |
| Downregulated | ENSMUSG00000036264 | 3,172426057 | 35,1042098 | 18,1549368 | 9,63567424 | 19,0831875 | 2,653938437 | 39,7562605 | 9,77780817 | 0 | 9,70826858 | 29,8110593 | 0 | 1,00175646 | 0 | 0 | 0 | 0 | 1,981702978 | 0 | 0 | 1,771456 | 10,7683497 |
| Downregulated | ENSMUSG00000047904 | 15,86213028 | 44,6780852 | 27,2324052 | 13,4899439 | 48,344075 | 38,48210734 | 55,6587648 | 44,0001368 | 11,7453565 | 26,9674127 | 69,5591385 | 0 | 35,0614763 | 11,22574 | 3,982475 | 8,203912116 | 4,963421 | 14,86277233 | 6,653965 | 1,333313 | 13,28592 | 29,6129618 |
| Downregulated | ENSMUSG00000026986 | 71,90832396 | 153,182006 | 92,5901777 | 45,2876689 | 86,5104499 | 35,8281689 | 82,9202006 | 133,222636 | 166,570511 | 118,656616 | 110,411331 | 22,77209 | 46,0807974 | 19,64505 | 11,94742 | 3,646183163 | 17,37197 | 68,36875273 | 51,33059 | 67,99894 | 120,459 | 83,4547104 |
| Downregulated | ENSMUSG00000045625 | 10,57475352 | 148,926951 | 55,3725572 | 67,4497197 | 162,8432 | 71,65633781 | 89,7355595 | 136,889314 | 143,079798 | 86,2957207 | 100,474311 | 12,2619 | 28,049181 | 7,483827 | 9,557939 | 5,469274744 | 7,445132 | 47,56087146 | 14,2585 | 27,33291 | 94,7729 | 74,4810856 |
| Downregulated | ENSMUSG00000089797 | 34,89668663 | 65,9533639 | 32,6788862 | 44,3241015 | 69,9716874 | 47,77089187 | 48,8434058 | 63,5557531 | 90,7595733 | 22,6526267 | 44,1645324 | 6,130948 | 37,0649892 | 16,83861 | 2,389485 | 4,557728953 | 4,963421 | 30,71639615 | 3,802266 | 17,99972 | 58,45805 | 56,5338361 |
| Downregulated | ENSMUSG00000044749 | 517,1054473 | 788,249075 | 627,253066 | 219,693373 | 798,949449 | 525,4798106 | 634,964275 | 727,224483 | 722,873307 | 537,190862 | 473,664609 | 189,1835 | 223,391692 | 86,99949 | 59,73712 | 62,89665955 | 157,175 | 284,3743773 | 212,9269 | 237,9963 | 617,3524 | 648,793071 |
| Downregulated | ENSMUSG00000021066 | 43,35648945 | 67,0171278 | 59,0035446 | 45,2876689 | 61,0661999 | 57,0596764 | 52,2510853 | 81,8891435 | 68,3366199 | 55,013522 | 51,8933255 | 13,13775 | 8,01405172 | 3,741914 | 15,1334 | 8,203912116 | 13,23579 | 27,74384169 | 19,9619 | 32,66616 | 46,94358 | 72,6863607 |
| Downregulated | ENSMUSG00000101660 | 12,68970423 | 22,3390426 | 16,3394431 | 9,63567424 | 33,077525 | 7,961815312 | 10,2230384 | 6,11113011 | 19,2196743 | 15,1017511 | 9,93701978 | 1,751699 | 1,00175646 | 4,677392 | 8,761444 | 2,734637372 | 6,617895 | 2,972554466 | 9,505665 | 7,999875 | 11,51446 | 19,7419745 |
| Downregulated | ENSMUSG00000069830 | 12,68970423 | 11,7014033 | 4,5387342 | 37,5791295 | 21,6276125 | 51,75179953 | 60,2023374 | 17,1111643 | 76,8786974 | 52,8561289 | 1,10411331 | 1,751699 | 0 | 0 | 0 | 0 | 0,827237 | 0 | 0 | 0 | 0 | 7,17889982 |
| Downregulated | ENSMUSG00000056856 | 4,229901409 | 11,7014033 | 0 | 0 | 10,1777 | 22,55847672 | 19,3101837 | 6,11113011 | 23,4907131 | 12,9443581 | 11,0411331 | 0,87585 | 0 | 0 | 1,59299 | 5,469274744 | 9,099605 | 0,990851489 | 1,901133 | 0,666656 | 0 | 0 |
| Downregulated | ENSMUSG00000034353 | 378,5761761 | 859,521259 | 441,164964 | 715,930596 | 992,325749 | 1479,570679 | 1184,73656 | 1236,89273 | 1304,80234 | 1213,53357 | 802,690376 | 35,90984 | 324,569095 | 48,64488 | 36,63877 | 39,196469 | 23,98987 | 236,8135058 | 238,5922 | 15,33309 | 335,6909 | 769,937006 |
| Downregulated | ENSMUSG00000067001 | 15,86213028 | 47,869377 | 22,693671 | 57,8140454 | 94,1437249 | 107,4845067 | 105,638064 | 128,333732 | 138,808759 | 139,15185 | 92,7455179 | 4,379248 | 6,01053879 | 10,29026 | 3,982475 | 21,87709898 | 4,136184 | 16,84447531 | 47,52832 | 5,33325 | 77,05834 | 76,2758106 |
| Downregulated | ENSMUSG00000071984 | 4831,604885 | 9037,73838 | 3799,82827 | 5686,01137 | 17445,85 | 19909,84616 | 15875,2428 | 11909,3704 | 18756,2666 | 16881,6004 | 14494,7995 | 5836,662 | 3821,70091 | 3078,659 | 3221,822 | 2897,804068 | 1515,498 | 4628,267304 | 3646,373 | 5753,244 | 15638,41 | 8651,47164 |
| Downregulated | ENSMUSG00000003665 | 86006,58535 | 74008,1844 | 73254,2622 | 55645,0552 | 93683,184 | 91879,3487 | 117997,717 | 96321,1883 | 134058,296 | 79688,7046 | 77224,9972 | 53899,79 | 50547,6294 | 30400,24 | 59207,45 | 63763,5396 | 18962,75 | 40149,30233 | 25863,01 | 40517,37 | 74266,52 | 49345,9626 |
| Downregulated | ENSMUSG00000086825 | 182,943236 | 82,9735868 | 171,564153 | 85,7575007 | 128,493462 | 258,7589976 | 296,468114 | 189,445033 | 262,668883 | 154,253601 | 183,282809 | 31,53059 | 52,0913362 | 82,3221 | 31,8598 | 35,55028583 | 21,50816 | 49,54257444 | 24,71473 | 105,3317 | 188,6601 | 148,962171 |
| Downregulated | ENSMUSG00000030513 | 289,7482465 | 178,712341 | 233,290938 | 225,474777 | 260,803562 | 530,7876874 | 347,583306 | 364,223354 | 732,483144 | 353,812455 | 292,590027 | 38,53739 | 72,1264654 | 127,2251 | 50,17918 | 112,1201322 | 50,46145 | 97,1034459 | 69,39135 | 226,6631 | 378,2059 | 203,701282 |
| Downregulated | ENSMUSG00000021097 | 29,60930986 | 12,7651672 | 9,0774684 | 10,5992417 | 49,6162874 | 70,32936859 | 48,8434058 | 57,444623 | 54,455744 | 23,7313232 | 41,9563057 | 14,0136 | 12,0210776 | 26,19339 | 7,168455 | 18,23091581 | 3,308947 | 5,945108933 | 14,2585 | 18,66638 | 26,57184 | 8,97362477 |
| Downregulated | ENSMUSG00000107546 | 82,48307748 | 82,9735868 | 24,5091647 | 46,2512364 | 96,6881499 | 92,8878453 | 129,49182 | 111,222568 | 132,402201 | 78,7448452 | 98,2660845 | 30,65474 | 33,0579633 | 68,28992 | 18,31938 | 47,40038111 | 14,89026 | 25,76213871 | 48,47889 | 49,99922 | 16,82883 | 33,2024117 |
| Downregulated | ENSMUSG00000024018 | 50,75881691 | 49,9969049 | 27,2324052 | 48,1783712 | 31,8053125 | 45,11695343 | 55,6587648 | 51,3334929 | 41,6426277 | 24,8100197 | 51,8933255 | 20,14454 | 11,0193211 | 13,0967 | 15,1334 | 6,380820534 | 16,54474 | 31,70724764 | 18,06076 | 26,66625 | 36,31485 | 34,0997741 |
| Downregulated | ENSMUSG00000022758 | 14,80465493 | 27,6578623 | 7,26197472 | 32,7612924 | 26,7164625 | 35,8281689 | 36,3485811 | 23,2222944 | 16,0163953 | 14,0230546 | 36,4357392 | 12,2619 | 4,00702586 | 0 | 0,796495 | 0 | 4,963421 | 10,89936638 | 5,703399 | 17,33306 | 30,11475 | 28,7155993 |
| Downregulated | ENSMUSG00000060572 | 14890,31044 | 12869,4161 | 14072,7993 | 24299,2433 | 18629,0076 | 18135,68831 | 23411,8939 | 24358,9646 | 17539,0206 | 19798,3957 | 18479,5444 | 3764,402 | 8027,07455 | 3582,882 | 1909,198 | 2810,295673 | 3727,529 | 11501,80408 | 7533,239 | 4884,59 | 15207,06 | 16271,8738 |
| Downregulated | ENSMUSG00000035041 | 790,9915635 | 1500,97091 | 1410,63859 | 1992,65743 | 2334,50994 | 1993,107766 | 2736,36662 | 2559,34129 | 1996,71061 | 2464,82152 | 1830,61987 | 123,4948 | 453,795678 | 215,16 | 63,7196 | 142,2011433 | 258,9251 | 859,0682408 | 521,861 | 693,3225 | 1154,104 | 1356,81207 |
| Downregulated | ENSMUSG00000025140 | 511,8180705 | 1088,2305 | 1110,17439 | 1328,75948 | 1622,07094 | 1346,873757 | 1824,24441 | 1753,89434 | 1249,27883 | 1605,10041 | 1509,32289 | 98,97102 | 296,519914 | 159,0313 | 70,09156 | 100,270037 | 268,0247 | 641,0809133 | 409,6942 | 517,9919 | 1187,761 | 898,25984 |
| Downregulated | ENSMUSG00000039787 | 2652,148184 | 1787,12341 | 2002,48953 | 3828,25338 | 4478,188 | 3769,91955 | 4729,85911 | 4918,23751 | 3111,45172 | 4776,46814 | 4332,54062 | 902,1252 | 727,275193 | 546,3194 | 494,6234 | 783,0178342 | 680,8159 | 1484,29553 | 2044,669 | 1810,638 | 3883,032 | 3733,92527 |
| Downregulated | ENSMUSG00000020098 | 119,4947148 | 312,746597 | 167,933165 | 417,224695 | 431,280037 | 421,9762115 | 444,134225 | 524,334963 | 361,970533 | 423,927728 | 380,919092 | 87,58497 | 92,1615947 | 85,12853 | 43,80722 | 84,77375853 | 32,26224 | 184,2983769 | 161,5963 | 103,9984 | 147,9166 | 305,103242 |
| Downregulated | ENSMUSG00000026828 | 60,27609508 | 95,7387541 | 88,9591903 | 175,369271 | 128,493462 | 147,2935833 | 146,530217 | 172,333869 | 154,825154 | 143,466636 | 161,200543 | 7,882647 | 6,01053879 | 1,870957 | 6,37196 | 10,93854949 | 4,136184 | 58,46023784 | 30,41813 | 73,33219 | 54,02941 | 133,707009 |
| Downregulated | ENSMUSG00000006958 | 701,1061586 | 726,550767 | 446,611445 | 645,590174 | 952,887162 | 663,4846093 | 841,69683 | 602,557429 | 657,739966 | 917,970729 | 686,758478 | 164,6597 | 292,512888 | 244,1599 | 92,39341 | 122,1471359 | 105,8863 | 202,1337037 | 212,9269 | 232,663 | 741,3543 | 547,391111 |
| Downregulated | ENSMUSG00000030881 | 856,5550354 | 935,048498 | 983,089828 | 1003,07369 | 1130,99691 | 1024,420237 | 1193,82371 | 1173,33698 | 969,525795 | 1099,19174 | 835,813775 | 311,8025 | 577,011724 | 342,3851 | 362,4052 | 283,4907409 | 350,7484 | 325,9901398 | 451,5191 | 726,6553 | 953,0433 | 895,567752 |
| Downregulated | ENSMUSG00000031595 | 7550,374016 | 8706,9078 | 6370,56732 | 7551,4779 | 11496,9844 | 9051,25704 | 11173,781 | 13082,7073 | 10140,5137 | 9089,09679 | 11008,0097 | 2974,386 | 7656,42466 | 4173,169 | 1848,665 | 2745,575921 | 1426,984 | 5628,036456 | 3947,703 | 2489,961 | 9502,09 | 10667,8451 |
| Downregulated | ENSMUSG00000010830 | 4421,304448 | 5999,62859 | 5751,48398 | 7763,46274 | 7551,85339 | 7281,080103 | 8618,02139 | 8892,91653 | 8208,93646 | 7201,3779 | 6898,49995 | 1459,166 | 3773,6166 | 2332,148 | 1350,059 | 1482,173456 | 2643,022 | 4891,8338 | 3182,497 | 5034,588 | 8364,815 | 6566,00125 |
| Downregulated | ENSMUSG00000019539 | 5992,712822 | 8626,06174 | 8721,63164 | 11698,6721 | 17186,3186 | 16480,9577 | 17858,5122 | 17052,4975 | 14889,9088 | 13798,6857 | 15829,6725 | 2350,781 | 5871,29464 | 4162,879 | 2194,344 | 3516,74366 | 3744,901 | 8552,0392 | 5802,258 | 9074,525 | 13854,56 | 10746,813 |
| Downregulated | ENSMUSG00000025784 | 6330,047459 | 4676,30625 | 4461,57572 | 5363,21628 | 5807,65006 | 9008,794025 | 6447,32957 | 7217,24466 | 7912,09927 | 4327,73039 | 8775,49258 | 789,1406 | 3929,89061 | 1460,282 | 630,824 | 1040,985293 | 274,6426 | 1473,396164 | 1424,899 | 1283,313 | 8231,07 | 3640,59957 |
| Downregulated | ENSMUSG00000027224 | 3630,312885 | 3613,60608 | 2240,3192 | 1941,58836 | 3106,74292 | 3466,043599 | 3513,31754 | 2732,89738 | 4390,62783 | 2090,51383 | 2315,32561 | 1210,424 | 2063,61832 | 975,704 | 690,5611 | 887,8456001 | 775,1209 | 1492,222342 | 1003,798 | 1047,317 | 4312,61 | 1806,39067 |
| Downregulated | ENSMUSG00000048706 | 389,1509297 | 351,042098 | 279,586027 | 339,175733 | 390,569237 | 449,8425651 | 407,785644 | 321,445444 | 235,974891 | 226,526267 | 261,674854 | 78,82647 | 102,179159 | 88,87045 | 108,3233 | 67,45438851 | 34,74395 | 76,29556464 | 105,5129 | 151,331 | 430,4638 | 209,98282 |
| Downregulated | ENSMUSG00000023046 | 17315,10142 | 17417,0069 | 14754,5171 | 13169,076 | 18653,1797 | 15879,84064 | 19674,8054 | 13726,8205 | 13446,2977 | 10244,3807 | 11195,709 | 5030,005 | 9924,40129 | 6613,832 | 3893,267 | 6755,465854 | 3025,205 | 6424,681053 | 5364,047 | 2196,632 | 12864,31 | 8200,09832 |
| Downregulated | ENSMUSG00000016494 | 34031,67179 | 37442,3629 | 39194,6931 | 32049,2161 | 45415,4418 | 42636,84796 | 52772,4602 | 59112,9615 | 58424,6067 | 46659,0175 | 35300,7107 | 9171,898 | 19301,8436 | 8656,917 | 6553,56 | 9111,811723 | 5425,846 | 16757,28038 | 11018,02 | 10981,16 | 39610,64 | 28070,3957 |
| Downregulated | ENSMUSG00000022032 | 38539,68922 | 32764,9929 | 40790,512 | 22385,5984 | 40209,5482 | 41634,9862 | 51017,5053 | 59287,7399 | 47465,1213 | 44145,6546 | 35063,3263 | 14299,12 | 18611,6334 | 9283,688 | 8708,876 | 7520,252773 | 2868,03 | 7903,031475 | 7044,648 | 3256,616 | 18006,85 | 19914,2681 |
| Downregulated | ENSMUSG00000030711 | 138,5292712 | 117,014033 | 85,328203 | 80,9396636 | 113,226912 | 171,1790292 | 162,432722 | 152,778253 | 125,995643 | 99,2400788 | 123,660691 | 25,39964 | 57,1001185 | 12,16122 | 3,18598 | 4,557728953 | 9,099605 | 25,76213871 | 18,06076 | 7,999875 | 81,48698 | 127,425472 |
| Downregulated | ENSMUSG00000025085 | 5789,677554 | 5532,63622 | 3628,26412 | 3041,98236 | 6436,12303 | 5602,464041 | 6680,18766 | 5716,3511 | 4941,59183 | 5610,30054 | 4914,40834 | 2024,964 | 1896,32499 | 1336,799 | 1120,668 | 1215,090539 | 659,3078 | 1394,128045 | 1566,534 | 1197,315 | 2494,21 | 3086,92692 |
| Downregulated | ENSMUSG00000053007 | 2089,571296 | 2136,03798 | 1725,62674 | 1450,16897 | 2199,65541 | 3195,341878 | 2562,57496 | 2187,78458 | 1945,45815 | 2304,09574 | 2360,59425 | 522,0064 | 741,299784 | 634,2543 | 428,5143 | 469,4460822 | 332,5492 | 715,3947749 | 573,1916 | 487,3257 | 1488,023 | 1164,7765 |
| Downregulated | ENSMUSG00000038233 | 1182,257444 | 711,658072 | 730,736206 | 621,500988 | 933,803974 | 843,952423 | 736,058767 | 957,002975 | 939,628523 | 872,665476 | 564,201901 | 129,6258 | 264,463707 | 56,1287 | 92,39341 | 92,97767064 | 65,35171 | 392,3771896 | 250,9496 | 90,66525 | 373,7772 | 743,016131 |
| Downregulated | ENSMUSG00000020101 | 701,1061586 | 420,186754 | 411,209318 | 382,536267 | 573,767837 | 472,4010418 | 695,166613 | 570,779552 | 616,097339 | 706,546214 | 730,92301 | 183,0526 | 175,307381 | 168,3861 | 212,6642 | 161,3436049 | 135,6668 | 154,5728323 | 244,2956 | 202,6635 | 369,3486 | 395,736853 |
| Downregulated | ENSMUSG00000054072 | 2918,631972 | 4570,99362 | 4030,39597 | 3220,24233 | 3141,09266 | 4782,397064 | 1712,92688 | 3765,67837 | 3114,655 | 3122,82639 | 2959,02367 | 564,9231 | 1033,81267 | 453,707 | 1147,749 | 1035,516018 | 1276,426 | 1068,137905 | 689,1607 | 177,9972 | 504,865 | 1084,91124 |
| Downregulated | ENSMUSG00000063388 | 42,29901409 | 57,4432524 | 44,4795952 | 17,3442136 | 47,0718625 | 76,96421468 | 18,1742905 | 52,5557189 | 25,6262325 | 40,9904673 | 41,9563057 | 0,87585 | 3,00526939 | 6,548349 | 3,18598 | 17,31937002 | 10,75408 | 2,972554466 | 10,45623 | 1,333313 | 11,51446 | 21,5366995 |
| Downregulated | ENSMUSG00000038425 | 134,2993697 | 159,56459 | 100,759899 | 97,3203098 | 160,298775 | 191,0835675 | 74,9689484 | 177,222773 | 73,6754183 | 103,554865 | 91,6414046 | 12,2619 | 42,0737715 | 17,77409 | 26,28433 | 10,0270037 | 50,46145 | 62,4236438 | 28,51699 | 75,99881 | 54,02941 | 84,3520729 |
| Downregulated | ENSMUSG00000069874 | 968,6474227 | 1241,41251 | 893,222891 | 441,31388 | 1228,95727 | 1366,778295 | 617,925878 | 1367,67092 | 995,152027 | 1395,83328 | 1026,82538 | 189,1835 | 132,231853 | 97,28975 | 231,78 | 235,178814 | 203,5003 | 293,2920407 | 200,5695 | 116,6648 | 325,0622 | 452,270689 |
| Downregulated | ENSMUSG00000036944 | 67,67842255 | 145,735659 | 51,7415699 | 66,4861523 | 117,04355 | 115,446322 | 69,2894827 | 145,444897 | 96,0983717 | 116,499223 | 87,2249514 | 27,15134 | 12,0210776 | 21,516 | 11,15093 | 18,23091581 | 8,272368 | 26,7529902 | 0,950566 | 34,66613 | 34,54339 | 50,2522987 |
| Downregulated | ENSMUSG00000004360 | 80,36812678 | 196,796328 | 98,9444056 | 166,697164 | 80,1493874 | 156,5823678 | 112,453423 | 215,11178 | 74,743178 | 176,906228 | 166,72111 | 59,55778 | 17,0298599 | 28,06435 | 34,24928 | 2,734637372 | 29,78053 | 24,77128722 | 27,56643 | 10,6665 | 17,71456 | 56,5338361 |
| Downregulated | ENSMUSG00000029134 | 14,80465493 | 92,5474623 | 10,8929621 | 43,3605341 | 53,4329249 | 103,5035991 | 24,9896495 | 94,1114037 | 71,5398989 | 66,8791836 | 75,079705 | 15,76529 | 28,049181 | 12,16122 | 0 | 4,557728953 | 0,827237 | 30,71639615 | 12,35736 | 0 | 6,200096 | 29,6129618 |
| Downregulated | ENSMUSG00000050640 | 169,1960564 | 312,746597 | 291,386736 | 566,577645 | 634,834037 | 832,0097001 | 592,936229 | 578,112908 | 616,097339 | 554,450006 | 592,908847 | 119,9914 | 165,289817 | 75,77375 | 122,6602 | 175,0167918 | 81,06921 | 380,4869717 | 173,9537 | 5,33325 | 87,68707 | 329,332029 |
| Downregulated | ENSMUSG00000040605 | 664,0945213 | 957,387541 | 1051,17084 | 1546,52572 | 1494,84969 | 2011,685335 | 1809,4778 | 2015,45071 | 1582,41985 | 2168,17998 | 1882,51319 | 505,3653 | 551,967812 | 452,7715 | 387,0965 | 431,161159 | 541,0129 | 1181,094975 | 707,2215 | 262,6626 | 557,1229 | 1208,74726 |
| Downregulated | ENSMUSG00000061080 | 716,9682889 | 1449,91024 | 1029,38492 | 1573,5056 | 1401,97817 | 2177,556488 | 1586,84274 | 1621,89393 | 2232,6855 | 1785,24272 | 1616,42188 | 582,44 | 401,704342 | 361,0947 | 444,4442 | 716,4749914 | 448,3624 | 878,8852706 | 450,5685 | 9,999844 | 202,8317 | 725,068882 |
| Downregulated | ENSMUSG00000036446 | 90651,0171 | 137619,14 | 137439,226 | 151534,467 | 177812,052 | 182345,4752 | 181270,374 | 256300,797 | 289921,313 | 207009,411 | 210858,039 | 81256,08 | 72748,5562 | 87757,23 | 83060,88 | 99497,04614 | 75300,89 | 161036,1565 | 93154,57 | 29258,88 | 61539,5 | 135446,098 |
| Downregulated | ENSMUSG00000029090 | 92,00035565 | 248,920761 | 187,903596 | 284,25239 | 395,658087 | 354,3007814 | 326,001336 | 474,223696 | 459,136665 | 382,937261 | 310,25584 | 73,57137 | 84,147543 | 296,5466 | 167,2639 | 106,6508575 | 45,49803 | 206,0971097 | 117,8702 | 131,3313 | 105,4016 | 157,038434 |
| Downregulated | ENSMUSG00000085338 | 3,172426057 | 19,1477508 | 7,26197472 | 41,4333992 | 40,7108 | 49,09786109 | 12,4948247 | 35,4445546 | 43,7781471 | 59,328308 | 33,1233993 | 5,255098 | 6,01053879 | 17,77409 | 7,964949 | 1,823091581 | 5,790658 | 8,917663399 | 2,851699 | 1,333313 | 12,40019 | 11,6657122 |
| Downregulated | ENSMUSG00000036617 | 114,2073381 | 129,7792 | 177,010634 | 121,409495 | 133,582312 | 152,6014601 | 174,927546 | 166,222739 | 127,063403 | 174,748834 | 232,967908 | 55,17853 | 27,0474245 | 37,41914 | 19,91237 | 52,86965586 | 59,56105 | 66,38704975 | 102,6612 | 19,99969 | 43,40067 | 100,504597 |
| Downregulated | ENSMUSG00000050821 | 97,28773241 | 92,5474623 | 91,6824308 | 105,028849 | 134,854525 | 103,5035991 | 140,850752 | 133,222636 | 85,4207749 | 88,4531138 | 124,764804 | 41,16494 | 20,0351293 | 10,29026 | 13,54041 | 20,96555318 | 30,60776 | 36,66150509 | 36,12153 | 33,99947 | 34,54339 | 56,5338361 |
| Downregulated | ENSMUSG00000025978 | 849,1527079 | 951,004957 | 889,591903 | 683,169304 | 1095,37496 | 927,5514838 | 1004,12955 | 1078,00335 | 808,294082 | 949,252928 | 1119,5709 | 287,2787 | 323,567338 | 230,1277 | 214,2571 | 292,6061988 | 349,0939 | 518,2153286 | 449,618 | 441,9931 | 580,1519 | 702,63482 |
| Downregulated | ENSMUSG00000024299 | 2179,456701 | 1842,43913 | 2158,62199 | 1523,4001 | 2432,4703 | 2693,747514 | 3451,97931 | 2585,00804 | 1346,44496 | 2924,34624 | 2731,57633 | 641,122 | 631,106573 | 479,9004 | 690,5611 | 535,0773791 | 679,9887 | 1215,774777 | 890,6808 | 991,9845 | 823,7271 | 1416,03799 |
| Downregulated | ENSMUSG00000050505 | 688,4164544 | 1330,76868 | 706,227041 | 682,205736 | 901,998662 | 789,5466851 | 1469,84575 | 1178,22588 | 580,861269 | 1720,52093 | 715,465424 | 231,2243 | 137,240636 | 136,5798 | 133,8112 | 147,6704181 | 118,2949 | 268,5207535 | 216,7292 | 66,66563 | 300,2618 | 481,88365 |
| Downregulated | ENSMUSG00000027015 | 580,5539684 | 619,11061 | 765,230586 | 516,472139 | 709,894574 | 558,654041 | 1445,99199 | 1136,6702 | 452,730107 | 1546,85079 | 946,225106 | 77,07477 | 47,0825538 | 19,64505 | 53,36516 | 68,3659343 | 17,37197 | 149,6185748 | 37,07209 | 19,33303 | 77,05834 | 288,053355 |
| Downregulated | ENSMUSG00000021991 | 103,6325845 | 190,413744 | 211,505014 | 220,65694 | 267,164625 | 238,8544594 | 350,990986 | 282,334211 | 132,402201 | 354,891152 | 134,701824 | 21,89624 | 11,0193211 | 41,16105 | 7,168455 | 41,93110637 | 7,445132 | 84,22237655 | 76,04532 | 3,999938 | 29,22902 | 113,067672 |
| Downregulated | ENSMUSG00000097040 | 14,80465493 | 24,4665705 | 15,4316963 | 26,9798879 | 29,2608875 | 27,86635359 | 39,7562605 | 28,1111985 | 7,4743178 | 36,6756813 | 25,3946061 | 8,758497 | 0 | 1,870957 | 3,982475 | 0 | 0,827237 | 10,89936638 | 6,653965 | 0 | 6,200096 | 23,3314244 |
| Downregulated | ENSMUSG00000004473 | 5027,237825 | 8048,43792 | 7916,46019 | 9545,0989 | 9834,20261 | 10390,16898 | 14434,9303 | 14703,379 | 9110,12564 | 10946,6122 | 9993,32956 | 1708,783 | 3186,58731 | 2060,859 | 1448,824 | 2786,595482 | 1509,707 | 4220,036491 | 2115,961 | 633,3234 | 2695,27 | 5359,04872 |
| Downregulated | ENSMUSG00000028339 | 2376,147117 | 3128,52973 | 3365,92528 | 2855,05028 | 3975,66406 | 4304,688145 | 4765,0718 | 4766,68148 | 2920,32274 | 4390,29479 | 4119,44676 | 963,4347 | 876,536906 | 829,7693 | 755,8737 | 1134,874509 | 419,4091 | 1272,253312 | 821,2895 | 124,6647 | 834,3558 | 1971,50536 |
| Downregulated | ENSMUSG00000014361 | 580,5539684 | 901,008052 | 537,386129 | 925,024727 | 1020,31442 | 1027,074175 | 1364,20768 | 1134,22575 | 809,361842 | 1093,79826 | 999,222544 | 322,3127 | 145,254687 | 169,3216 | 241,338 | 185,0437955 | 113,3314 | 399,31315 | 273,7632 | 89,33194 | 157,6596 | 577,901435 |
| Downregulated | ENSMUSG00000006800 | 11894,48276 | 13421,5096 | 11348,651 | 15321,6856 | 17609,9654 | 17926,02717 | 21831,8665 | 18581,5022 | 14724,4061 | 20715,2878 | 15359,3202 | 3499,02 | 4065,12773 | 4161,008 | 2711,269 | 4056,378768 | 1809,994 | 6683,293292 | 5254,732 | 963,985 | 4558,842 | 9791,12199 |

| | | | | | | | | | | | | | | | | | | | | | | | |
|---|---|---|---|---|---|---|---|---|---|---|---|---|---|---|---|---|---|---|---|---|---|---|---|
| Downregulated | ENSMUSG00000031398 | 187,1731374 | 235,091829 | 287,755748 | 270,762446 | 278,614537 | 348,9929045 | 403,242071 | 413,112395 | 296,837193 | 428,242514 | 194,323942 | 79,70232 | 52,0913362 | 56,1287 | 83,63197 | 39,196469 | 76,10579 | 127,8198421 | 67,49022 | 85,332 | 85,91562 | 170,498871 |
| Downregulated | ENSMUSG00000023393 | 290,8057219 | 318,065416 | 378,530432 | 409,516155 | 463,08535 | 545,3843489 | 609,974626 | 602,557429 | 368,377092 | 690,365766 | 474,768723 | 60,43363 | 71,124709 | 67,35444 | 42,21423 | 55,60429323 | 80,24197 | 233,8409514 | 176,8054 | 111,3316 | 123,1162 | 328,434667 |
| Downregulated | ENSMUSG00000007097 | 296,0930986 | 414,867934 | 479,290331 | 364,228486 | 765,871924 | 1023,093268 | 997,314193 | 1135,44797 | 755,973857 | 1121,84437 | 594,01296 | 68,31628 | 103,180916 | 32,74174 | 23,89485 | 76,56984641 | 32,26224 | 193,2160403 | 141,6344 | 21,333 | 93,88717 | 429,836627 |
| Downregulated | ENSMUSG00000041193 | 1544,97149 | 3000,87806 | 1345,28082 | 2181,51665 | 3885,33697 | 3420,926646 | 3440,62038 | 2831,89769 | 2482,54127 | 2402,25713 | 3763,92227 | 897,7459 | 1130,98305 | 1433,153 | 1514,933 | 1404,692063 | 347,4395 | 1382,237827 | 730,0351 | 22,66631 | 343,6625 | 1434,8826 |
| Downregulated | ENSMUSG00000036596 | 20639,80393 | 27999,3305 | 15692,2196 | 26595,4245 | 44902,7401 | 42662,06038 | 35434,1871 | 48742,3737 | 29059,0798 | 32040,5224 | 37659,0967 | 13476,7 | 19833,7762 | 17764,73 | 11381,12 | 16698,60734 | 3852,442 | 16302,47955 | 11267,06 | 1209,314 | 9161,085 | 24884,7589 |
| Downregulated | ENSMUSG00000018593 | 602987,2505 | 810530,674 | 506265,844 | 833512,802 | 1152822,99 | 885609,9677 | 977103,246 | 1022223,4 | 664836,297 | 786397,801 | 1011333,56 | 302233,8 | 606396,246 | 518060,4 | 299584,1 | 456046,3591 | 152486,2 | 507433,8736 | 312305,8 | 154652,3 | 509335,2 | 760656,483 |
| Downregulated | ENSMUSG00000083282 | 3820,658448 | 3880,61083 | 3076,35404 | 3296,36416 | 5417,08082 | 4102,988824 | 4465,19601 | 4172,67964 | 3385,86596 | 4621,13585 | 4423,07792 | 1360,195 | 2066,62359 | 1285,347 | 967,7414 | 1516,812196 | 726,3139 | 1987,648087 | 1539,918 | 593,9907 | 2248,863 | 2902,96761 |
| Downregulated | ENSMUSG00000001506 | 2140733,011 | 2132896,68 | 1671894,48 | 2251680,74 | 3447611,91 | 2874977,008 | 3301905,1 | 2726772,81 | 1781363,77 | 2612378,58 | 3128490,71 | 987497,8 | 1234100,85 | 826923,6 | 816608,8 | 1248454,026 | 419909,6 | 1314421,969 | 908231,1 | 432597,9 | 976325,6 | 2079433,84 |
| Downregulated | ENSMUSG00000029661 | 1904071,085 | 2162426,77 | 1314806,85 | 2040099,64 | 3274496,86 | 2446619,401 | 2699868,1 | 2649964,46 | 1867491,4 | 2527319,04 | 2509809,65 | 842598,9 | 1285330,68 | 993040,2 | 692301,5 | 1029465,177 | 464034,4 | 1203068,097 | 968423,8 | 354626,5 | 971936,8 | 2079381,79 |
| Downregulated | ENSMUSG00000042129 | 306,6678522 | 323,384236 | 369,452964 | 415,29756 | 782,410687 | 759,026393 | 538,413357 | 771,22462 | 492,237215 | 780,976273 | 772,879316 | 0,87585 | 27,0474245 | 33,67722 | 11,94742 | 7,292366325 | 4,963421 | 97,1034459 | 101,7106 | 46,66594 | 63,77242 | 257,543031 |
| Downregulated | ENSMUSG00000048834 | 265,4263134 | 538,264551 | 359,467749 | 449,02242 | 659,006074 | 617,0406867 | 578,169618 | 680,779894 | 541,354161 | 654,768781 | 868,937174 | 194,4386 | 193,338998 | 217,9665 | 115,4918 | 152,228147 | 64,52447 | 244,7403177 | 332,6983 | 62,66569 | 98,31581 | 375,994878 |
| Downregulated | ENSMUSG00000031428 | 238,9894296 | 224,45419 | 164,302178 | 157,06149 | 470,718625 | 441,8807498 | 346,447413 | 73,3335613 | 347,021898 | 276,146306 | 226,343228 | 36,78569 | 71,124709 | 101,9671 | 29,47031 | 68,3659343 | 21,50816 | 159,5270897 | 98,85892 | 5,999906 | 25,68611 | 108,58086 |
| Downregulated | ENSMUSG00000038508 | 678,8991762 | 748,889809 | 806,986941 | 479,856577 | 1424,878 | 1369,432234 | 1514,14558 | 854,335989 | 1548,25154 | 951,410321 | 893,227667 | 487,8483 | 632,108329 | 482,7068 | 653,9224 | 371,9106826 | 608,0191 | 716,3856264 | 734,7879 | 227,3298 | 255,9754 | 475,602113 |
| Downregulated | ENSMUSG00000061013 | 902,0264755 | 1048,87124 | 852,374283 | 547,306297 | 1802,72511 | 829,3557616 | 1405,09984 | 1206,33708 | 1272,76955 | 1193,03834 | 630,448699 | 201,4454 | 154,270496 | 188,0312 | 470,7285 | 272,5521914 | 350,7484 | 668,8247549 | 465,7776 | 63,33234 | 140,8308 | 723,274157 |
| Downregulated | ENSMUSG00000020774 | 85,65550354 | 177,648577 | 106,20638 | 124,300198 | 262,075775 | 230,892644 | 205,596662 | 172,333869 | 191,128984 | 159,647083 | 155,679977 | 19,26869 | 41,072015 | 28,06435 | 16,72639 | 31,90410267 | 73,62408 | 105,0302578 | 40,87436 | 10,6665 | 60,22951 | 97,81251 |
| Downregulated | ENSMUSG00000024697 | 93,057831 | 268,068511 | 182,457115 | 141,644411 | 525,423762 | 245,4893054 | 313,506512 | 286,000889 | 404,680921 | 306,349809 | 255,050174 | 56,93023 | 24,0421551 | 17,77409 | 46,19671 | 36,46183163 | 126,5672 | 216,996476 | 93,15552 | 33,33281 | 119,5733 | 174,985683 |
| Downregulated | ENSMUSG00000028121 | 87,77045424 | 103,185102 | 95,3134182 | 171,515001 | 390,569237 | 244,1623362 | 243,081136 | 198,000616 | 240,245929 | 257,808466 | 165,616996 | 33,28229 | 53,0930926 | 65,48349 | 66,10908 | 62,89665955 | 23,98987 | 124,8472876 | 82,69928 | 49,99922 | 44,2864 | 110,375585 |
| Downregulated | ENSMUSG00000003746 | 12420,04801 | 13931,0525 | 13784,1358 | 9468,01351 | 17023,4754 | 16534,03646 | 15141,4558 | 12525,3723 | 13372,6223 | 14728,5221 | 18082,0637 | 10392,83 | 8108,21682 | 7947,824 | 7195,535 | 8552,122608 | 3765,582 | 7417,514245 | 5922,98 | 3983,938 | 5473,799 | 6500,49379 |
| Downregulated | ENSMUSG00000034037 | 813,1985459 | 836,118452 | 755,245371 | 540,561325 | 1049,57531 | 932,8593607 | 1006,40134 | 625,779723 | 755,973857 | 1348,37064 | 883,290647 | 567,5506 | 409,718394 | 354,5463 | 502,5883 | 472,1807195 | 196,0551 | 250,6854267 | 310,8352 | 91,33191 | 209,9175 | 448,681239 |
| Downregulated | ENSMUSG00000028871 | 1967,961631 | 2228,58544 | 1745,59717 | 1514,72799 | 2525,34181 | 2277,079179 | 2764,76395 | 1719,67201 | 2189,97512 | 3035,45198 | 2206,01839 | 1193,783 | 875,53515 | 1295,638 | 627,638 | 906,9880617 | 209,2909 | 624,236438 | 493,344 | 17,99972 | 136,4021 | 616,488022 |
| Downregulated | ENSMUSG00000025357 | 776,1869086 | 765,910032 | 692,610839 | 615,719584 | 997,414599 | 834,6636385 | 857,599334 | 827,447017 | 669,485323 | 925,521605 | 894,33178 | 509,7445 | 371,651648 | 295,6112 | 467,5425 | 348,210492 | 236,5897 | 342,8346151 | 311,7858 | 253,3294 | 317,0906 | 505,215075 |
| Downregulated | ENSMUSG00000032010 | 3112,149962 | 2948,75362 | 2438,20801 | 2344,35954 | 4661,3866 | 3316,096077 | 4546,98031 | 3312,23252 | 3413,62772 | 4714,98244 | 4214,4005 | 2220,279 | 1581,77346 | 1388,25 | 2022,301 | 1822,180035 | 1062,999 | 1877,663571 | 1398,283 | 378,6608 | 945,9575 | 1963,4291 |
| Downregulated | ENSMUSG00000043456 | 359,5416198 | 458,482256 | 226,028963 | 267,871744 | 477,079687 | 602,4440253 | 405,513857 | 320,223218 | 326,734464 | 471,390375 | 396,376678 | 63,93703 | 26,0456681 | 42,09653 | 69,29506 | 81,12757537 | 36,39842 | 165,4721986 | 59,88569 | 0,666656 | 21,25747 | 148,962171 |
| Downregulated | ENSMUSG00000026170 | 1357,798352 | 1472,24928 | 872,344713 | 804,578799 | 1614,43766 | 2095,284396 | 1436,90485 | 1235,67051 | 1218,3138 | 1686,00264 | 1779,83065 | 529,8891 | 439,771088 | 157,1604 | 201,5132 | 302,6332025 | 141,4575 | 428,0478432 | 335,55 | 39,99938 | 331,2623 | 569,825173 |
| Downregulated | ENSMUSG00000021408 | 5802,367258 | 6330,45917 | 4900,01744 | 4013,25832 | 7441,1709 | 6796,736338 | 5143,32422 | 4463,56943 | 7093,12759 | 6327,63372 | 5437,75805 | 4064,818 | 3203,61717 | 2767,145 | 2464,355 | 2841,288229 | 1264,845 | 2851,670585 | 2778,506 | 1128,649 | 1416,279 | 3227,81283 |
| Downregulated | ENSMUSG00000052062 | 1362,028254 | 1833,92902 | 1050,26309 | 1006,92796 | 1543,19376 | 1512,744909 | 1341,48982 | 1359,11534 | 1267,43075 | 1532,82774 | 1425,41028 | 690,1696 | 807,41571 | 478,0295 | 307,4471 | 425,6918842 | 262,2341 | 620,273032 | 330,7971 | 77,33213 | 390,6061 | 656,869333 |
| Downregulated | ENSMUSG00000028476 | 3867,187363 | 5620,92863 | 2950,17723 | 3499,67688 | 5298,76506 | 4641,738327 | 4364,10151 | 3623,90015 | 4890,33936 | 6131,31096 | 4431,91082 | 2405,083 | 1277,23949 | 1477,12 | 1170,848 | 1947,973355 | 972,0033 | 1917,297631 | 1904,935 | 538,6583 | 1203,704 | 1527,31094 |
| Downregulated | ENSMUSG00000024219 | 906,2563769 | 552,093482 | 616,360104 | 519,362842 | 1118,27479 | 942,1481452 | 1027,98331 | 706,446641 | 741,025222 | 841,383277 | 746,380597 | 472,083 | 168,295086 | 113,1929 | 226,2046 | 229,7095392 | 158,8295 | 315,0907734 | 223,3831 | 173,9973 | 253,3182 | 307,79533 |
| Downregulated | ENSMUSG00000030768 | 657,7496691 | 582,942636 | 533,755142 | 481,783712 | 989,781324 | 967,3605604 | 834,881471 | 671,002086 | 870,224144 | 682,81489 | 871,145401 | 255,7481 | 151,265226 | 188,9666 | 298,6856 | 247,0289093 | 241,5532 | 265,548199 | 309,8847 | 404,6604 | 397,6919 | 457,654863 |
| Downregulated | ENSMUSG00000033102 | 395,4957818 | 354,23339 | 289,571242 | 136,826574 | 530,512612 | 456,4774112 | 490,705844 | 328,7788 | 502,914812 | 316,058077 | 441,645324 | 202,3213 | 118,207263 | 102,9026 | 203,9027 | 216,0363524 | 125,74 | 216,0056246 | 162,5469 | 169,9973 | 150,5738 | 228,827432 |
| Downregulated | ENSMUSG00000019647 | 539,3124297 | 521,244328 | 580,050231 | 318,940817 | 781,138474 | 812,1051618 | 998,450086 | 825,002565 | 805,090803 | 1059,27997 | 642,593946 | 326,6919 | 227,398717 | 217,031 | 364,7947 | 342,7412173 | 246,5166 | 387,4229321 | 329,8466 | 63,33234 | 150,5738 | 439,707614 |
| Downregulated | ENSMUSG00000025648 | 1299,637208 | 1096,74062 | 1195,50259 | 659,080118 | 2068,61752 | 2072,725919 | 2130,93556 | 2025,22852 | 1711,61878 | 2094,82862 | 2112,16876 | 1552,006 | 883,549202 | 752,1246 | 772,6001 | 968,9731754 | 583,202 | 799,6171515 | 602,6592 | 225,9965 | 306,4619 | 956,588401 |
| Downregulated | ENSMUSG00000026510 | 2533,710944 | 2421,12671 | 2422,77632 | 2126,5933 | 3274,67497 | 3211,265509 | 3821,14458 | 3251,12122 | 2925,66154 | 3059,1833 | 3038,51983 | 2339,395 | 1537,69617 | 1689,474 | 1870,967 | 1913,334615 | 1165,577 | 1412,954223 | 1165,395 | 673,3228 | 674,039 | 1776,77771 |
| Downregulated | ENSMUSG00000001552 | 1690,903088 | 1174,39538 | 1222,73499 | 849,866468 | 2544,425 | 2412,430039 | 2756,8127 | 2178,00677 | 2813,54677 | 1747,48834 | 2247,9747 | 947,6694 | 846,484213 | 1016,865 | 925,5271 | 916,1035196 | 612,9825 | 674,7698639 | 826,0423 | 275,329 | 539,4084 | 1205,15781 |
| Downregulated | ENSMUSG00000074170 | 1800,880525 | 1698,831 | 1299,89347 | 1261,30976 | 2854,84485 | 2437,642455 | 2895,39166 | 2156,0067 | 2580,77516 | 1921,15848 | 2275,57753 | 658,639 | 1320,31502 | 860,6401 | 727,1999 | 868,7031385 | 540,1857 | 1174,159014 | 735,7385 | 411,9936 | 703,268 | 1363,99097 |
| Downregulated | ENSMUSG00000038332 | 485,3811867 | 372,317377 | 487,460053 | 375,791295 | 632,289612 | 590,5013023 | 756,504843 | 798,113592 | 749,567299 | 474,626464 | 748,588823 | 257,4998 | 230,403987 | 258,192 | 168,0604 | 289,8715614 | 136,4941 | 224,923288 | 173,0031 | 135,9979 | 149,688 | 393,044765 |
| Downregulated | ENSMUSG00000098188 | 3510,81817 | 3327,45359 | 3212,51607 | 2466,73261 | 5667,70668 | 4929,690647 | 5581,77898 | 6524,2425 | 6497,31769 | 4220,93944 | 5542,64881 | 3356,256 | 2682,70381 | 1662,345 | 2930,305 | 2562,355217 | 1389,758 | 3049,840883 | 2033,262 | 625,3236 | 1388,822 | 3320,24117 |
| Downregulated | ENSMUSG00000090439 | 70,85084861 | 221,262898 | 79,8817219 | 162,842895 | 366,3972 | 216,2959826 | 336,224375 | 179,667225 | 191,128984 | 303,113719 | 411,834264 | 22,77209 | 39,0685021 | 107,58 | 129,8287 | 141,2895975 | 11,58132 | 27,74384169 | 33,26983 | 7,333219 | 50,4865 | 134,604372 |
| Downregulated | ENSMUSG00000030717 | 16971,42193 | 28835,449 | 13614,3871 | 16073,2682 | 42433,3757 | 25915,70884 | 34749,2435 | 24218,4086 | 26652,3495 | 30479,6486 | 38919,9941 | 13110,59 | 20987,7997 | 18516,86 | 16348,06 | 17765,11591 | 6229,093 | 13583,58306 | 11497,1 | 4309,266 | 13786,36 | 27991,4278 |
| Downregulated | ENSMUSG00000031274 | 336,277162 | 1088,2305 | 423,917774 | 557,905539 | 1391,80047 | 780,2579005 | 1038,20635 | 946,002941 | 855,275508 | 1172,54311 | 1504,90644 | 279,3961 | 374,656918 | 264,7404 | 252,4889 | 274,375283 | 55,42487 | 312,118219 | 334,5994 | 217,3299 | 476,5217 | 791,473705 |
| Downregulated | ENSMUSG00000049690 | 84,59802819 | 149,990715 | 30,8633926 | 79,0125288 | 232,814887 | 173,8329676 | 96,5509185 | 51,3334929 | 85,4207749 | 158,568387 | 166,72111 | 57,80608 | 84,147543 | 59,87062 | 62,12661 | 38,28492321 | 33,08947 | 66,38704975 | 37,07209 | 17,33306 | 69,97251 | 134,604372 |
| Downregulated | ENSMUSG00000037490 | 85,65550354 | 127,651672 | 110,745114 | 79,9760962 | 281,158962 | 205,6802289 | 111,31753 | 127,111506 | 134,53772 | 171,512745 | 183,282809 | 46,42003 | 51,0895797 | 28,99983 | 44,60372 | 25,52328214 | 27,29882 | 59,45108933 | 48,47889 | 29,33288 | 64,65815 | 113,067672 |
| Downregulated | ENSMUSG00000039316 | 1378,947859 | 1964,77199 | 1247,24416 | 1238,18414 | 2954,07742 | 2007,704428 | 1927,61069 | 1096,33674 | 1966,81334 | 2720,4726 | 2224,78832 | 717,3209 | 675,183857 | 678,2218 | 816,4073 | 1091,120311 | 550,1125 | 953,1991322 | 668,2482 | 329,3282 | 666,0675 | 1091,19277 |
| Downregulated | ENSMUSG00000015966 | 45,47144015 | 152,118243 | 35,4021268 | 36,6155621 | 185,743025 | 116,7732912 | 94,2791321 | 77,0002394 | 81,1497361 | 125,128795 | 118,140124 | 42,91663 | 37,0649892 | 21,516 | 27,08083 | 24,61173635 | 6,617895 | 39,63405955 | 24,71473 | 2,666625 | 21,25747 | 65,5074609 |
| Downregulated | ENSMUSG00000055555 | 23,26445775 | 275,514859 | 62,634532 | 138,753709 | 718,800062 | 461,7852881 | 354,398665 | 528,001641 | 265,872162 | 171,512745 | 285,965347 | 141,8877 | 117,205506 | 231,0632 | 82,03898 | 66,54284272 | 112,5042 | 190,2434859 | 75,09475 | 8,666531 | 49,60077 | 227,032707 |
| Downregulated | ENSMUSG00000030043 | 159,6787782 | 392,528892 | 88,0514435 | 63,59545 | 600,484299 | 410,0334886 | 227,178632 | 336,112156 | 180,451387 | 256,729769 | 215,302095 | 14,88944 | 20,0351293 | 56,1287 | 129,8287 | 108,4739491 | 26,47158 | 74,31386166 | 38,02266 | 0 | 13,28592 | 101,40196 |
| Downregulated | ENSMUSG00000037860 | 200,9203169 | 272,323567 | 150,685975 | 212,948401 | 550,868012 | 502,9213339 | 304,419366 | 336,112156 | 298,972712 | 217,896695 | 287,06946 | 131,3775 | 57,1001185 | 82,3221 | 115,4918 | 198,7169824 | 69,48789 | 213,0330701 | 85,55098 | 1,999969 | 9,743008 | 169,601508 |
| Downregulated | ENSMUSG00000087385 | 49,70134156 | 42,5505574 | 42,6641015 | 36,6155621 | 66,1550499 | 61,04058406 | 51,1151921 | 61,1113011 | 40,5748681 | 34,5182883 | 45,2686457 | 14,88944 | 6,01053879 | 27,12887 | 11,15093 | 9,115457906 | 21,50816 | 24,77128722 | 45,62719 | 15,33309 | 35,42912 | 42,1760364 |
| Downregulated | ENSMUSG00000049598 | 185,0581867 | 143,608131 | 138,885267 | 102,138147 | 326,958612 | 189,7565983 | 245,352922 | 158,889383 | 119,589085 | 101,397472 | 200,948622 | 51,67513 | 58,1018749 | 85,12853 | 43,80722 | 67,45438851 | 20,68092 | 54,49683188 | 93,15552 | 12,66647 | 85,91562 | 147,167446 |
| Downregulated | ENSMUSG00000040280 | 3649,347441 | 6182,59598 | 3032,78219 | 4672,33844 | 9929,61855 | 10083,63909 | 10553,5833 | 6532,79809 | 5572,6378 | 7238,05358 | 6768,21458 | 807,5334 | 2116,71141 | 1483,669 | 361,6087 | 708,2710793 | 358,1936 | 2016,38278 | 989,5397 | 527,9918 | 3554,427 | 4150,30146 |
| Downregulated | ENSMUSG00000074240 | 179,7708099 | 240,410649 | 161,578938 | 290,033795 | 643,739524 | 822,7209155 | 605,431053 | 372,778937 | 342,750859 | 343,02549 | 515,620915 | 14,88944 | 102,179159 | 53,32227 | 7,964949 | 15,49627844 | 3,308947 | 160,5179412 | 68,44079 | 3,333281 | 106,2874 | 309,590055 |
| Downregulated | ENSMUSG00000015970 | 119,4947148 | 174,457285 | 16,3394431 | 57,8140454 | 227,726037 | 241,5083978 | 258,98364 | 147,889349 | 68,3366199 | 144,545332 | 149,055297 | 15,76529 | 8,01405172 | 13,0967 | 44,60372 | 23,70019056 | 7,445132 | 30,71639615 | 21,86303 | 2,666625 | 31,88621 | 152,551621 |
| Downregulated | ENSMUSG00000040420 | 3,172426057 | 22,3390426 | 0,90774684 | 8,67210682 | 64,8828374 | 59,71361484 | 27,2614358 | 45,2223628 | 10,6775969 | 29,1248057 | 30,9151726 | 0 | 3,00526939 | 0 | 0 | 0 | 0 | 10,89936638 | 3,802266 | 1,333313 | 7,971552 | 17,9472495 |
| Downregulated | ENSMUSG00000017491 | 42,29901409 | 63,825836 | 25,4169115 | 67,4497197 | 227,726037 | 149,9475217 | 115,861102 | 118,555924 | 61,9300618 | 126,207492 | 125,868917 | 7,006798 | 43,075528 | 10,29026 | 20,70887 | 11,85009528 | 25,64434 | 46,57001997 | 59,88569 | 28,66622 | 91,22999 | 88,8388853 |
| Downregulated | ENSMUSG00000081189 | 206,2076937 | 71,2721836 | 252,353622 | 52,9962083 | 57,2495624 | 181,794783 | 105,638064 | 386,223423 | 77,9464571 | 220,054088 | 181,074583 | 35,03399 | 43,075528 | 46,77392 | 97,17238 | 23,70019056 | 39,70737 | 13,87192084 | 64,63852 | 107,3317 | 39,85776 | 117,554485 |
| Downregulated | ENSMUSG00000037813 | 87,77045424 | 29,7853902 | 110,745114 | 58,7776129 | 44,5274375 | 74,31027624 | 69,2894827 | 146,667123 | 68,3366199 | 119,735313 | 96,0578579 | 15,76529 | 17,0298599 | 0 | 12,74392 | 23,70019056 | 9,926842 | 97,1034459 | 38,02266 | 2,666625 | 37,20058 | 76,2758106 |
| Downregulated | ENSMUSG00000031748 | 138,5292712 | 174,457285 | 170,656406 | 114,664523 | 222,637187 | 305,2029203 | 240,80935 | 419,223525 | 139,876519 | 209,267123 | 211,989755 | 42,04079 | 40,0702586 | 60,8061 | 53,36516 | 38,28492321 | 57,90658 | 75,30471315 | 71,29249 | 62,66569 | 80,60125 | 137,296459 |
| Downregulated | ENSMUSG00000090098 | 75,08075001 | 60,6345442 | 49,9260762 | 28,9070227 | 57,2495624 | 70,32936859 | 49,979299 | 108,778116 | 18,1519147 | 63,643094 | 32,019286 | 2,627549 | 13,022834 | 33,67722 | 12,74392 | 1,823091581 | 5,790658 | 27,74384169 | 28,51699 | 9,999844 | 7,085824 | 23,3314244 |
| Downregulated | ENSMUSG00000051262 | 149,1040247 | 203,178911 | 164,302178 | 75,1582591 | 145,032225 | 181,794783 | 223,770952 | 251,77856 | 37,371589 | 175,827531 | 140,22239 | 27,15134 | 44,0772844 | 45,83844 | 35,84227 | 12,76164107 | 60,38829 | 94,13089144 | 122,6231 | 55,33247 | 22,1432 | 79,8652605 |
| Downregulated | ENSMUSG00000044139 | 20,09203169 | 15,956459 | 24,5091647 | 10,5992417 | 16,5387625 | 27,86635359 | 27,2614358 | 26,8889725 | 11,7453565 | 18,3378407 | 16,5616996 | 5,255098 | 12,0210776 | 14,03218 | 1,59299 | 18,23091581 | 1,654474 | 10,89936638 | 1,901133 | 3,333281 | 1,771456 | 11,6657122 |
| Downregulated | ENSMUSG00000070031 | 82,48307748 | 57,4432524 | 100,759899 | 85,7575007 | 41,9830125 | 159,2363062 | 51,1151921 | 135,667088 | 29,8972712 | 50,6987359 | 69,5591385 | 55,17853 | 45,0790409 | 20,58052 | 37,43526 | 48,3119269 | 33,08947 | 22,78958424 | 31,36869 | 40,66603 | 71,74397 | 31,4076867 |
| Downregulated | ENSMUSG00000028441 | 24,3219331 | 6,3825836 | 28,140152 | 13,4899439 | 17,810975 | 31,84726125 | 9,08714527 | 29,3334245 | 8,54207749 | 4,31478604 | 14,353473 | 21,89624 | 4,00702586 | 9,354784 | 7,964949 | 2,734637372 | 7,445132 | 3,963405955 | 0,950566 | 0,666656 | 0,885728 | 1,79472495 |
| Downregulated | ENSMUSG00000078921 | 65,56347184 | 59,5707803 | 91,6824308 | 60,7047477 | 54,7051374 | 104,8305683 | 24,9896495 | 108,778116 | 65,1333408 | 61,485701 | 47,4768723 | 11,38605 | 9,01580818 | 13,0967 | 3,18598 | 8,203912116 | 14,89026 | 20,80788127 | 1,901133 | 3,999938 | 16,82883 | 20,639337 |
| Downregulated | ENSMUSG00000078922 | 194,5754648 | 165,947174 | 285,940255 | 169,587867 | 176,837537 | 236,2005209 | 38,6203674 | 194,333937 | 180,451387 | 212,503212 | 78,3920449 | 18,39284 | 24,0421551 | 42,09653 | 23,09835 | 11,85009528 | 24,81711 | 51,52427742 | 7,604532 | 3,999938 | 52,25795 | 52,9443862 |
| Downregulated | ENSMUSG00000025766 | 30,66678522 | 45,7418492 | 75,3429877 | 46,2512364 | 27,988675 | 75,63724546 | 14,7666111 | 75,7780133 | 33,1005503 | 40,9904673 | 22,0822662 | 4,379248 | 20,0351293 | 25,25792 | 9,557939 | 11,85009528 | 13,23579 | 20,80788127 | 17,1102 | 21,333 | 23,91466 | 26,9208743 |
| Downregulated | ENSMUSG00000072769 | 21,14950705 | 2,12752787 | 25,4169115 | 17,3442136 | 11,4499125 | 41,13604578 | 11,3589316 | 22,0000684 | 16,0163953 | 21,5739302 | 6,62467985 | 0 | 3,00526939 | 8,419306 | 3,982475 | 1,823091581 | 0 | 13,87192084 | 0,950566 | 0 | 7,971552 | 5,38417486 |
| Downregulated | ENSMUSG00000105096 | 189,2880881 | 109,567685 | 155,22471 | 99,2474447 | 117,04355 | 226,9117364 | 97,6868116 | 185,778355 | 198,603302 | 169,355352 | 138,014164 | 53,42683 | 35,0614763 | 18,70957 | 45,40021 | 20,96555318 | 46,32526 | 48,55172295 | 34,22039 | 9,333188 | 16,82883 | 62,8153734 |
| Downregulated | ENSMUSG00000078920 | 767,7271058 | 747,826046 | 837,850333 | 677,387899 | 793,860599 | 1454,358264 | 472,531554 | 1342,00417 | 1349,64824 | 1059,27997 | 891,01944 | 117,3639 | 234,411013 | 94,48332 | 324,9699 | 212,3901692 | 239,0714 | 319,0541794 | 169,2008 | 73,99885 | 424,2637 | 439,707614 |
| Downregulated | ENSMUSG00000090942 | 168,138581 | 158,500826 | 187,903596 | 161,879327 | 157,75435 | 395,4368271 | 101,094491 | 337,334382 | 237,04265 | 252,414983 | 177,762243 | 28,02719 | 132,231853 | 57,06418 | 285,9417 | 49,22347269 | 69,48789 | 77,28641613 | 36,12153 | 14,66644 | 81,48698 | 92,4283352 |
| Downregulated | ENSMUSG00000043664 | 59,21861973 | 55,3157246 | 42,6641015 | 26,0163204 | 27,988675 | 94,21481452 | 20,4460768 | 17,1111643 | 8,54207749 | 14,0230546 | 7,72879316 | 12,2619 | 6,01053879 | 17,77409 | 0 | 9,115457906 | 8,272368 | 6,935960422 | 13,30793 | 1,333313 | 10,62874 | 0 |
| Downregulated | ENSMUSG00000073555 | 271,7711655 | 276,578623 | 249,630381 | 211,984833 | 145,032225 | 322,4535201 | 67,0176963 | 96,5558557 | 87,5562942 | 118,656616 | 86,1208381 | 25,39964 | 79,1387607 | 74,83827 | 41,41774 | 41,93110637 | 84,37816 | 80,25897059 | 89,35325 | 25,99959 | 45,17213 | 53,8417486 |
| Downregulated | ENSMUSG00000108004 | 15,86213028 | 21,2752787 | 11,8007089 | 20,2349159 | 3,8166375 | 34,50119968 | 7,95125211 | 6,11113011 | 6,40655811 | 12,9443581 | 0 | 4,379248 | 1,00175646 | 1,870957 | 0,796495 | 5,469274744 | 6,617895 | 2,972554466 | 0 | 0 | 7,085824 | 0 |
| Downregulated | ENSMUSG00000063767 | 27,49435916 | 15,956459 | 15,4316963 | 1,92713485 | 7,63327499 | 43,78998421 | 23,8537563 | 7,33335613 | 7,4743178 | 33,4395918 | 3,31233993 | 0,87585 | 0 | 0 | 0 | 0 | 11,58132 | 0,990851489 | 0 | 0,666656 | 0 | 2,69208743 |
| Downregulated | ENSMUSG00000039699 | 253,7940846 | 346,787042 | 239,645166 | 221,620508 | 264,6202 | 423,3031807 | 221,499166 | 135,667088 | 215,687457 | 407,74728 | 204,260962 | 70,06798 | 70,1229525 | 106,6445 | 139,3866 | 190,5130702 | 172,0653 | 122,8655846 | 105,5129 | 35,33278 | 16,82883 | 107,683497 |
| Downregulated | ENSMUSG00000048388 | 1894,995831 | 2920,032 | 1892,65216 | 1264,20046 | 1413,42809 | 951,4369297 | 711,069117 | 1191,67037 | 873,427423 | 1816,52492 | 1571,15324 | 569,3023 | 925,622973 | 371,3849 | 214,2571 | 456,6844411 | 136,4941 | 698,5502996 | 522,8116 | 458,6595 | 544,7227 | 763,655468 |
| Downregulated | ENSMUSG00000043257 | 452,5994508 | 427,633101 | 335,866331 | 252,454665 | 231,542675 | 276,0095975 | 235,129884 | 146,667123 | 123,860124 | 322,530256 | 175,554016 | 117,3639 | 115,201993 | 98,22523 | 136,9971 | 173,1937002 | 107,5408 | 168,4447531 | 153,9918 | 137,3312 | 130,202 | 160,627883 |
| Downregulated | ENSMUSG00000043342 | 87,77045424 | 98,9300459 | 74,4352409 | 37,5791295 | 35,62195 | 46,44392265 | 29,5332221 | 14,6667123 | 25,6262325 | 76,5874522 | 47,4768723 | 35,90984 | 22,0386422 | 18,70957 | 13,54041 | 15,49627844 | 26,47158 | 16,84447531 | 17,1102 | 4,666594 | 0,885728 | 9,87098725 |
| Downregulated | ENSMUSG00000062563 | 371,1738487 | 432,951921 | 216,043748 | 270,762446 | 283,703387 | 179,1408445 | 159,025042 | 127,111506 | 162,299472 | 230,841053 | 282,653007 | 95,46762 | 58,1018749 | 61,74157 | 97,17238 | 84,77375853 | 34,74395 | 130,7923965 | 69,39135 | 12,66647 | 34,54339 | 62,8153734 |
| Downregulated | ENSMUSG00000064105 | 253,7940846 | 235,091829 | 177,918381 | 211,984833 | 204,826212 | 143,3126756 | 139,714858 | 78,2224654 | 119,589085 | 234,077142 | 224,135002 | 114,7363 | 85,1492995 | 103,8381 | 80,44599 | 122,1471359 | 29,78053 | 68,36875273 | 112,1668 | 41,99934 | 42,51494 | 87,9415228 |
| Downregulated | ENSMUSG00000031750 | 1202,349476 | 1753,08296 | 1247,24416 | 1856,79443 | 937,620612 | 859,8760537 | 868,958266 | 711,335545 | 546,692959 | 1089,48347 | 1116,25856 | 424,7871 | 194,340754 | 260,9985 | 461,1706 | 619,8511376 | 506,2689 | 770,8824583 | 435,3595 | 117,9982 | 180,6885 | 465,731126 |
| Downregulated | ENSMUSG00000039611 | 268,5987395 | 292,535082 | 225,121216 | 309,305143 | 208,64285 | 201,6993212 | 128,355927 | 221,22291 | 143,079798 | 269,674127 | 202,052736 | 110,3571 | 92,1615947 | 139,3863 | 152,927 | 155,8743302 | 71,14237 | 159,5270897 | 129,277 | 20,66634 | 43,40067 | 120,246572 |
| Downregulated | ENSMUSG00000030107 | 1129,383676 | 1190,35184 | 1259,04487 | 777,598911 | 839,660249 | 1285,833173 | 831,473792 | 1018,11428 | 696,179315 | 1027,99777 | 908,685253 | 120,8673 | 142,249418 | 79,51566 | 786,1405 | 377,3799573 | 605,5374 | 290,3194862 | 370,7209 | 70,66556 | 492,4648 | 513,291337 |
| Downregulated | ENSMUSG00000046879 | 2661,665462 | 3996,5611 | 3433,09855 | 2493,71249 | 2662,74076 | 4704,10588 | 1761,77029 | 2487,22995 | 1427,5947 | 4189,65724 | 2056,96309 | 895,1184 | 401,704342 | 242,2889 | 913,5797 | 821,3027574 | 547,6308 | 774,8458643 | 575,0927 | 160,6642 | 672,2676 | 1119,90837 |
| Downregulated | ENSMUSG00000049502 | 581,6114438 | 748,889809 | 637,238282 | 381,5727 | 519,062699 | 663,4846093 | 480,482806 | 466,89034 | 358,767254 | 892,082013 | 430,60419 | 237,3553 | 94,1651077 | 78,58018 | 317,005 | 295,3408362 | 371,4293 | 173,3990105 | 295,6262 | 198,6636 | 232,0607 | 363,431803 |
| Downregulated | ENSMUSG00000043065 | 351,081817 | 337,213167 | 297,740964 | 164,77003 | 242,992587 | 262,7399053 | 184,014692 | 206,556198 | 280,820797 | 297,720237 | 220,822662 | 109,4812 | 59,1036314 | 108,5155 | 147,3516 | 63,80820534 | 127,3945 | 106,0211093 | 112,1668 | 134,6646 | 147,0309 | 116,657122 |
| Downregulated | ENSMUSG00000040253 | 1474,120641 | 1342,47008 | 1604,89641 | 975,130233 | 1101,73602 | 1178,348666 | 757,640737 | 1129,33684 | 1543,98051 | 1540,37862 | 913,101706 | 270,6376 | 343,602467 | 296,5466 | 635,603 | 464,8883532 | 741,2042 | 462,7276453 | 389,7323 | 281,3289 | 395,0347 | 522,264962 |
| Downregulated | ENSMUSG00000104713 | 979,2221762 | 960,578832 | 1394,29915 | 539,597757 | 446,546587 | 1057,594467 | 739,466446 | 858,002667 | 679,09516 | 1465,94856 | 777,295769 | 350,3399 | 336,590172 | 316,1917 | 332,1384 | 359,1490415 | 342,4761 | 225,9141394 | 209,1246 | 67,99894 | 215,2319 | 383,173778 |

| | | | | | | | | | | | | | | | | | | | | | | | |
|---|---|---|---|---|---|---|---|---|---|---|---|---|---|---|---|---|---|---|---|---|---|---|---|
| Downregulated | ENSMUSG00000016496 | 256,9665106 | 491,458937 | 594,57418 | 383,499835 | 372,758262 | 457,8043804 | 400,970285 | 322,66767 | 394,003324 | 797,15672 | 474,768723 | 269,7617 | 138,242392 | 152,483 | 171,2464 | 274,375283 | 104,2318 | 135,746654 | 124,5242 | 32,66616 | 135,5164 | 163,319971 |
| Downregulated | ENSMUSG00000117406 | 58,16114438 | 104,248866 | 108,929621 | 77,0853939 | 49,6162874 | 123,4081373 | 74,9689484 | 69,6668832 | 74,743178 | 120,814009 | 56,3097787 | 66,56458 | 24,0421551 | 28,06435 | 18,31938 | 28,25791951 | 27,29882 | 36,66150509 | 14,2585 | 8,666531 | 16,82883 | 28,7155993 |
| Downregulated | ENSMUSG00000004187 | 209,3801198 | 85,1011147 | 148,870482 | 43,3605341 | 104,321425 | 180,4678137 | 185,150585 | 144,222671 | 64,0655811 | 98,1613823 | 114,827784 | 33,28229 | 39,0685021 | 4,677392 | 13,54041 | 13,67318686 | 9,926842 | 51,52427742 | 70,34192 | 28,66622 | 16,82883 | 47,5602113 |
| Downregulated | ENSMUSG00000032131 | 330,9897853 | 249,984524 | 241,460659 | 246,673261 | 251,898075 | 387,4750118 | 441,862439 | 267,667499 | 219,958495 | 347,340276 | 141,326504 | 84,08157 | 31,0544504 | 52,38679 | 57,34764 | 76,56984641 | 80,24197 | 117,9113272 | 131,1782 | 36,66609 | 44,2864 | 87,0441603 |
| Downregulated | ENSMUSG00000026413 | 271,7711655 | 242,538177 | 143,424001 | 143,571546 | 129,765675 | 233,5465825 | 400,970285 | 133,222636 | 133,469961 | 229,762356 | 99,3701978 | 35,03399 | 15,026347 | 16,83861 | 6,37196 | 19,1424616 | 7,445132 | 18,82617829 | 38,97323 | 0 | 16,82883 | 76,2758106 |
| Downregulated | ENSMUSG00000055980 | 489,6110881 | 451,035908 | 264,15433 | 336,285031 | 477,079687 | 509,5561799 | 487,298165 | 268,889725 | 222,094015 | 449,816444 | 278,236554 | 198,8179 | 97,1703771 | 86,99949 | 156,9095 | 108,4739491 | 230,7991 | 206,0971097 | 258,5541 | 202,6635 | 149,688 | 323,947854 |
| Downregulated | ENSMUSG00000021453 | 12274,11641 | 9703,65461 | 6797,20834 | 8812,78766 | 13933,2713 | 18589,51178 | 13058,2277 | 4629,79217 | 7031,19753 | 9206,67471 | 9510,83204 | 4117,369 | 2460,31388 | 3212,433 | 4079,647 | 2660,802163 | 3423,933 | 4977,047028 | 3936,296 | 3377,281 | 2723,614 | 5881,31368 |
| Downregulated | ENSMUSG00000029708 | 1925,662617 | 1445,65519 | 1359,80477 | 1177,47939 | 1446,50561 | 2251,866764 | 1352,84875 | 1411,67106 | 1047,47225 | 1662,27132 | 665,780325 | 513,2479 | 376,660431 | 288,1273 | 552,7675 | 568,8045734 | 503,7872 | 618,291329 | 618,8188 | 240,6629 | 172,717 | 477,396838 |
| Downregulated | ENSMUSG00000069893 | 105,7475352 | 56,3794885 | 69,8965067 | 79,0125288 | 67,4272624 | 111,4654144 | 97,6868116 | 74,5557873 | 72,6076586 | 91,6892033 | 41,9563057 | 12,2619 | 5,00878232 | 12,16122 | 41,41774 | 28,25791951 | 37,22566 | 47,56087146 | 23,76416 | 13,99978 | 36,31485 | 54,7391111 |
| Downregulated | ENSMUSG00000030956 | 593,2436726 | 369,126085 | 457,504407 | 363,264919 | 526,695974 | 302,5489818 | 314,642405 | 237,111848 | 281,888557 | 407,74728 | 404,105471 | 124,3707 | 102,179159 | 68,28992 | 270,8083 | 149,4935097 | 142,2847 | 139,7100599 | 200,5695 | 158,6642 | 186,0029 | 272,798193 |
| Downregulated | ENSMUSG00000035863 | 1775,501117 | 1176,52291 | 1315,32517 | 874,919221 | 1095,37496 | 1235,408343 | 1067,73957 | 688,11325 | 867,020865 | 1102,42783 | 1206,79585 | 456,3177 | 468,822025 | 276,9016 | 473,9145 | 288,0484698 | 363,157 | 442,9106155 | 516,1576 | 531,325 | 886,6137 | 544,699024 |
| Downregulated | ENSMUSG00000038967 | 790,9915635 | 499,969049 | 544,648104 | 405,661886 | 530,512612 | 468,4201342 | 421,416362 | 425,334656 | 329,937743 | 568,47306 | 474,768723 | 111,2329 | 121,212532 | 149,6765 | 210,2747 | 194,1592534 | 217,5633 | 284,3743773 | 288,0216 | 309,3285 | 361,377 | 322,153129 |
| Downregulated | ENSMUSG00000086448 | 25,37940846 | 19,1477508 | 22,693671 | 6,74497197 | 16,5387625 | 13,26969219 | 17,0383974 | 4,88890409 | 13,8808759 | 28,0461092 | 16,5616996 | 3,503399 | 3,00526939 | 3,741914 | 3,982475 | 0 | 0,827237 | 1,981702978 | 0,950566 | 0 | 7,971552 | 7,17889982 |
| Downregulated | ENSMUSG00000027188 | 3779,416909 | 2608,34917 | 2047,87687 | 1378,86498 | 2954,07742 | 2291,675841 | 2394,46278 | 518,223833 | 1661,43407 | 3259,82085 | 2170,68677 | 1041,385 | 482,846616 | 149,6765 | 428,5143 | 441,1881627 | 445,0534 | 741,1569136 | 932,5057 | 221,3299 | 2014,146 | 1540,77137 |
| Downregulated | ENSMUSG00000034614 | 582,6689191 | 296,790138 | 401,224103 | 214,875536 | 323,141975 | 390,1289503 | 319,185977 | 308,000957 | 418,561797 | 317,136774 | 433,91653 | 160,2805 | 75,1317348 | 110,3864 | 107,5268 | 121,2355902 | 83,55092 | 218,978179 | 236,6911 | 159,3308 | 309,1191 | 192,932933 |
| Downregulated | ENSMUSG00000030731 | 211,4950705 | 153,182006 | 146,147241 | 123,33663 | 104,321425 | 91,56087608 | 78,3766279 | 79,4446914 | 93,9628523 | 181,221014 | 121,452464 | 54,30268 | 21,0368858 | 43,03201 | 27,08083 | 10,93854949 | 4,963421 | 62,4236438 | 66,53965 | 14,66644 | 15,9431 | 43,9707614 |
| Downregulated | ENSMUSG00000051373 | 214,6674965 | 163,819646 | 149,778229 | 187,895648 | 111,9547 | 139,331768 | 156,753256 | 129,555958 | 153,757395 | 198,480158 | 158,992316 | 65,68873 | 61,1071443 | 43,96748 | 47,7897 | 63,80820534 | 74,45132 | 62,4236438 | 75,09475 | 53,3325 | 67,31533 | 87,9415228 |
| Downregulated | ENSMUSG00000036599 | 4006,77411 | 3113,63703 | 3020,98148 | 3198,08028 | 2773,42325 | 2578,301192 | 2565,98264 | 2334,4517 | 1839,74994 | 2833,73573 | 2503,02487 | 808,4093 | 836,466648 | 765,2213 | 1166,069 | 778,4601052 | 542,6674 | 940,3180629 | 1145,433 | 999,9844 | 1446,394 | 1107,3453 |
| Downregulated | ENSMUSG00000057778 | 398,6682078 | 280,833679 | 291,386736 | 290,997362 | 189,559662 | 205,6802289 | 235,129884 | 221,22291 | 177,248108 | 318,21547 | 193,219829 | 81,45402 | 84,147543 | 107,58 | 83,63197 | 59,25047639 | 65,35171 | 102,0577033 | 134,0299 | 121,3314 | 164,7454 | 111,272947 |
| Downregulated | ENSMUSG00000027490 | 853,3826093 | 688,255265 | 688,072105 | 569,468348 | 473,26305 | 553,3461642 | 592,936229 | 622,113045 | 443,12027 | 752,930163 | 420,667171 | 126,9982 | 231,405743 | 101,9671 | 178,4149 | 192,3361618 | 179,5104 | 342,8346151 | 319,3903 | 337,9947 | 400,3491 | 324,845217 |
| Downregulated | ENSMUSG00000071176 | 1137,843479 | 1302,04706 | 1117,43636 | 771,817507 | 1039,39761 | 1105,365359 | 991,634727 | 869,002701 | 755,973857 | 1067,90954 | 994,806091 | 409,8977 | 264,463707 | 286,2564 | 318,598 | 455,7728953 | 378,0472 | 597,4834478 | 494,2946 | 425,3267 | 713,8968 | 438,810251 |
| Downregulated | ENSMUSG00000027412 | 1176,970067 | 1193,54313 | 1047,53985 | 907,680513 | 1043,21425 | 1301,756803 | 1138,16494 | 1173,33698 | 748,49954 | 1855,358 | 994,806091 | 211,9556 | 256,449655 | 174,9345 | 187,9728 | 189,6015245 | 218,3905 | 331,9352488 | 410,6447 | 473,3259 | 615,581 | 607,514397 |
| Downregulated | ENSMUSG00000040183 | 395,4957818 | 371,253613 | 310,449419 | 229,329047 | 333,319675 | 297,241105 | 248,760602 | 253,000787 | 257,330084 | 484,334733 | 319,088746 | 124,3707 | 96,1686206 | 82,3221 | 99,56187 | 123,9702275 | 76,93303 | 96,11259441 | 149,2389 | 34,66613 | 46,05786 | 109,478222 |
| Downregulated | ENSMUSG00000078853 | 1726,85725 | 1588,19955 | 1522,29145 | 1045,47066 | 1190,7909 | 1694,539692 | 861,007014 | 1269,89284 | 1179,87445 | 1872,61714 | 1414,36915 | 353,8433 | 436,765819 | 298,4176 | 614,8941 | 702,8018046 | 337,5126 | 402,2857045 | 237,6416 | 87,99863 | 346,3197 | 536,622762 |
| Downregulated | ENSMUSG00000110123 | 45,47144015 | 42,5505574 | 29,9556457 | 33,7248598 | 52,1607124 | 33,17423047 | 30,6691153 | 41,5556847 | 34,1683099 | 36,6756813 | 17,6658129 | 13,13775 | 11,0193211 | 15,90313 | 11,94742 | 12,76164107 | 18,19921 | 7,92681191 | 10,45623 | 5,999906 | 14,17165 | 13,4604372 |
| Downregulated | ENSMUSG00000038742 | 58,16114438 | 37,2317377 | 31,7711394 | 47,2148038 | 45,79965 | 33,17423047 | 27,2614358 | 55,000171 | 38,4393487 | 39,9117708 | 20,9781529 | 10,5102 | 8,01405172 | 17,77409 | 11,94742 | 9,115457906 | 19,85368 | 14,86277233 | 13,30793 | 17,99972 | 22,1432 | 18,844612 |
| Downregulated | ENSMUSG00000046532 | 1607,362536 | 1582,88073 | 814,248915 | 1517,61869 | 1297,65675 | 1362,797388 | 1043,88581 | 1036,44767 | 1142,50286 | 731,356233 | 506,788009 | 313,5542 | 477,837834 | 490,1907 | 395,858 | 226,0633561 | 197,7096 | 435,9746551 | 394,4851 | 225,9965 | 449,9498 | 487,267825 |
| Downregulated | ENSMUSG00000022604 | 186,115662 | 161,692118 | 92,5901777 | 147,425816 | 198,46515 | 167,1981215 | 105,638064 | 111,222568 | 107,843728 | 87,3744172 | 115,931897 | 15,76529 | 16,0281034 | 28,06435 | 52,56867 | 72,01211746 | 47,1525 | 65,39619826 | 36,12153 | 85,332 | 35,42912 | 64,6100984 |
| Downregulated | ENSMUSG00000002578 | 470,5765318 | 527,626911 | 245,091647 | 303,523739 | 604,300937 | 550,6922257 | 423,688148 | 447,334724 | 294,701673 | 334,395918 | 352,212146 | 193,5628 | 85,1492995 | 169,3216 | 221,4256 | 92,97767064 | 80,24197 | 140,7009114 | 120,7219 | 80,66541 | 64,65815 | 172,293596 |
| Downregulated | ENSMUSG00000071669 | 130,0694683 | 104,248866 | 60,8190383 | 77,0853939 | 151,393287 | 130,0429834 | 144,258431 | 116,111472 | 62,9978215 | 87,3744172 | 94,9537446 | 32,40644 | 24,0421551 | 42,09653 | 33,45279 | 34,63874004 | 31,435 | 35,6706536 | 24,71473 | 60,66572 | 37,20058 | 38,5865865 |
| Downregulated | ENSMUSG00000074063 | 301,3804754 | 363,807265 | 210,597267 | 358,447082 | 405,835787 | 356,9547198 | 238,537563 | 152,778253 | 229,568332 | 232,998446 | 207,573302 | 312,6783 | 202,354806 | 154,3539 | 423,7353 | 242,4711803 | 95,13224 | 112,9570697 | 107,414 | 51,99919 | 72,6297 | 244,082594 |
| Downregulated | ENSMUSG00000087088 | 13,74717958 | 46,8056131 | 14,5239494 | 41,4333992 | 25,44425 | 25,21241515 | 18,1742905 | 9,77780817 | 2,13551937 | 10,7869651 | 6,62467985 | 8,758497 | 14,0245905 | 4,677392 | 39,02825 | 9,115457906 | 4,963421 | 7,92681191 | 5,703399 | 7,999875 | 3,542912 | 14,3577996 |
| Downregulated | ENSMUSG00000021411 | 5938,781579 | 6936,80461 | 5689,75719 | 3698,17177 | 5714,77854 | 2189,499211 | 5293,26212 | 1533,89366 | 1786,36195 | 1269,62579 | 2505,2331 | 1406,615 | 689,208448 | 4453,813 | 1305,455 | 1583,355038 | 1482,408 | 1383,228678 | 2525,655 | 378,6608 | 1715,655 | 2306,22157 |
| Downregulated | ENSMUSG00000021559 | 13706,99552 | 12477,9509 | 8332,20824 | 6851,92795 | 13083,4333 | 8558,95146 | 8121,63608 | 8669,24917 | 6451,40402 | 6797,9454 | 5591,2298 | 4842,573 | 3645,39177 | 1956,085 | 2994,821 | 2846,757504 | 1949,797 | 3257,919695 | 3576,982 | 1614,641 | 2157,633 | 3990,57094 |
| Downregulated | ENSMUSG00000039457 | 1874,9038 | 1924,34896 | 1416,99282 | 1208,31355 | 1945,21291 | 1125,269897 | 1209,72621 | 750,446777 | 694,043796 | 1126,15916 | 547,640201 | 681,4111 | 460,807974 | 278,7726 | 672,2417 | 397,4339647 | 119,1221 | 216,0056246 | 495,2451 | 61,33238 | 383,5202 | 594,951323 |
| Downregulated | ENSMUSG00000110874 | 1076,509909 | 1433,95378 | 675,363649 | 1018,49077 | 1362,53959 | 906,3199763 | 920,073458 | 801,78027 | 334,208782 | 747,536681 | 693,383158 | 235,6036 | 597,046853 | 205,8052 | 182,3973 | 281,6676493 | 143,9392 | 622,254735 | 411,5953 | 38,66606 | 203,7174 | 620,077472 |
| Downregulated | ENSMUSG00000019768 | 2828,746567 | 3864,65437 | 2728,687 | 2496,6032 | 4454,01596 | 2201,441934 | 2982,85543 | 2281,89598 | 2093,87674 | 3523,0228 | 1911,22014 | 854,8293 | 1616,83493 | 658,5768 | 843,4882 | 767,5215557 | 335,8582 | 1372,329312 | 904,9393 | 305,9952 | 805,1268 | 1711,27024 |
| Downregulated | ENSMUSG00000118010 | 52,87376762 | 60,6345442 | 17,24719 | 46,2512364 | 64,8828374 | 37,15513812 | 64,74591 | 24,4445204 | 20,287434 | 29,1248057 | 28,706946 | 2,627549 | 13,022834 | 8,419306 | 3,982475 | 3,646183163 | 0 | 12,88106935 | 2,851699 | 1,333313 | 17,71456 | 17,0498871 |
| Downregulated | ENSMUSG00000063887 | 41,24153874 | 77,6547672 | 9,98521524 | 26,0163204 | 96,6881499 | 19,90453828 | 98,8227048 | 31,7778766 | 10,6775969 | 31,2821988 | 29,8110593 | 10,5102 | 27,0474245 | 0,935478 | 0 | 0 | 0 | 13,87192084 | 6,653965 | 0 | 3,542912 | 15,2551621 |
| Downregulated | ENSMUSG00000074865 | 20,09203169 | 93,6112262 | 49,0183294 | 27,9434553 | 29,2608875 | 46,44392265 | 52,2510853 | 52,5557189 | 36,3038293 | 33,4395918 | 18,7699262 | 21,02039 | 7,01229525 | 0 | 17,52289 | 6,380820534 | 33,91671 | 8,917663399 | 54,18229 | 43,33266 | 22,1432 | 38,5865865 |
| Downregulated | ENSMUSG00000107278 | 14,80465493 | 26,5940983 | 9,0774684 | 4,81783712 | 11,4499125 | 15,92363062 | 24,9896495 | 34,2223286 | 3,20327906 | 8,62957207 | 5,52056654 | 0 | 7,01229525 | 0,935478 | 3,18598 | 0 | 0,827237 | 10,89936638 | 28,51699 | 2,666625 | 1,771456 | 7,17889982 |
| Downregulated | ENSMUSG00000045394 | 48,64386621 | 47,869377 | 16,3394431 | 29,8705901 | 6,36106249 | 53,07876874 | 47,7075126 | 31,7778766 | 44,8459068 | 10,7869651 | 16,5616996 | 13,13775 | 11,0193211 | 0 | 5,575465 | 14,58473265 | 34,74395 | 18,82617829 | 31,36869 | 0,666656 | 15,9431 | 13,4604372 |
| Downregulated | ENSMUSG00000034379 | 46,5289155 | 30,8491541 | 29,9556457 | 39,5062644 | 3,8166375 | 30,52029203 | 92,0073458 | 22,0000684 | 18,1519147 | 28,0461092 | 32,019286 | 21,02039 | 6,01053879 | 3,741914 | 20,70887 | 0 | 30,60776 | 15,85362382 | 29,46756 | 25,33294 | 6,200096 | 13,4604372 |
| Downregulated | ENSMUSG00000052974 | 25011,40703 | 33856,4147 | 26389,1084 | 28404,0405 | 42563,1414 | 16910,89572 | 90888,4911 | 31703,3208 | 24406,8509 | 32891,614 | 19185,0729 | 9315,537 | 17000,809 | 13438,15 | 2569,493 | 4868,566068 | 6198,486 | 20647,36332 | 12781,32 | 2850,622 | 6130,124 | 22985,0425 |
| Downregulated | ENSMUSG00000042501 | 23,26445775 | 80,846059 | 50,833823 | 94,4296076 | 86,5104499 | 59,71361484 | 202,188982 | 64,7779792 | 44,8459068 | 70,1152731 | 87,2249514 | 19,26869 | 14,0245905 | 9,354784 | 5,575465 | 11,85009528 | 76,10579 | 32,69809913 | 38,97323 | 8,666531 | 9,743008 | 90,6336102 |
| Downregulated | ENSMUSG00000056755 | 0 | 3,1912918 | 9,0774684 | 0 | 27,988675 | 21,2315075 | 109,045743 | 12,2222602 | 4,27103874 | 4,31478604 | 12,1452464 | 0 | 2,00351293 | 0 | 3,982475 | 0 | 1,654474 | 0 | 0 | 1,333313 | 0 | 30,5103242 |
| Downregulated | ENSMUSG00000075289 | 28,55183451 | 61,6983082 | 47,2028357 | 37,5791295 | 55,9773499 | 47,77089187 | 76,1048416 | 50,1112669 | 43,7781471 | 47,4626464 | 46,372759 | 58,68193 | 15,026347 | 1,870957 | 21,50536 | 45,57728953 | 31,435 | 14,86277233 | 15,20906 | 23,99963 | 38,0863 | 23,3314244 |
| Downregulated | ENSMUSG00000000791 | 111,034912 | 109,567685 | 130,715545 | 84,7939333 | 80,1493874 | 233,5465825 | 273,750251 | 23,2222944 | 12,8131162 | 98,1613823 | 158,992316 | 87,58497 | 16,0281034 | 34,6127 | 5,575465 | 58,3389306 | 37,22566 | 28,73469318 | 112,1668 | 11,33316 | 57,57232 | 32,3050492 |
| Downregulated | ENSMUSG00000112058 | 45,47144015 | 71,2721836 | 52,6493167 | 6,74497197 | 66,1550499 | 115,446322 | 120,404675 | 34,2223286 | 25,6262325 | 46,3839499 | 88,3290647 | 32,40644 | 17,0298599 | 11,22574 | 42,21423 | 47,40038111 | 42,18908 | 12,88106935 | 49,42946 | 6,666563 | 28,3433 | 37,6892241 |
| Downregulated | ENSMUSG00000024948 | 89,88540495 | 107,440157 | 81,6972156 | 99,2474447 | 62,3384124 | 86,25299921 | 84,0560937 | 58,666849 | 29,8972712 | 132,679671 | 113,7236711 | 60,43363 | 44,0772844 | 63,61253 | 23,09835 | 30,99255688 | 34,74395 | 54,49683188 | 39,92379 | 37,99941 | 18,60029 | 19,7419745 |
| Downregulated | ENSMUSG00000034201 | 3852,382708 | 3377,45049 | 3093,60123 | 1837,52308 | 4259,36745 | 3256,382462 | 2855,6354 | 1299,22626 | 2355,47787 | 6033,14958 | 6243,76076 | 2600,398 | 2329,08378 | 1591,249 | 970,1308 | 1381,903419 | 408,655 | 760,9739434 | 1293,721 | 431,9933 | 914,957 | 1234,77077 |
| Downregulated | ENSMUSG00000032997 | 10688,96086 | 11033,3595 | 8491,06394 | 5957,73738 | 12491,8545 | 4129,528208 | 7246,99835 | 2998,12043 | 4115,14583 | 14248,5022 | 15282,0323 | 4470,337 | 7050,362 | 3100,175 | 1237,753 | 2591,524683 | 494,6876 | 2402,81486 | 2089,345 | 1415,978 | 2487,124 | 3408,18269 |
| Downregulated | ENSMUSG00000038822 | 435,6798452 | 534,009495 | 458,412154 | 355,556379 | 457,9965 | 262,7399053 | 218,091486 | 195,556163 | 324,598944 | 625,643975 | 496,850989 | 292,5338 | 262,460194 | 286,2564 | 251,6924 | 257,9674588 | 72,79684 | 196,1885948 | 202,4707 | 96,66516 | 122,2305 | 171,396233 |
| Downregulated | ENSMUSG00000006651 | 389,1509297 | 603,154151 | 449,334686 | 238,001154 | 398,202512 | 180,4678137 | 233,993991 | 101,44476 | 201,806581 | 680,657497 | 619,407566 | 409,0218 | 216,379396 | 221,7084 | 109,1198 | 151,3166012 | 59,56105 | 100,0760004 | 112,1668 | 22,66631 | 12,40019 | 61,0206485 |
| Downregulated | ENSMUSG00000021464 | 1157,935511 | 1589,26332 | 1105,63565 | 770,853939 | 912,176362 | 719,2173165 | 632,692489 | 352,001094 | 519,998967 | 1816,52492 | 1321,62363 | 826,8021 | 314,55153 | 411,6105 | 375,1491 | 459,4190785 | 105,0591 | 348,7797241 | 402,0896 | 127,998 | 137,2878 | 263,824568 |
| Downregulated | ENSMUSG00000040913 | 93,057831 | 58,5070164 | 50,833823 | 37,5791295 | 50,8884999 | 38,48210734 | 24,9896495 | 12,2222602 | 28,8295115 | 96,0039893 | 66,2467985 | 49,92343 | 14,0245905 | 24,32244 | 11,94742 | 12,76164107 | 4,963421 | 4,954257444 | 10,45623 | 10,6665 | 15,9431 | 19,7419745 |
| Downregulated | ENSMUSG00000024190 | 93679,62651 | 72196,5944 | 76528,5051 | 58853,7347 | 72924,4926 | 62696,64164 | 56993,4392 | 26446,5267 | 49580,3532 | 128096,289 | 91607,1771 | 69009,95 | 49197,2617 | 44300,51 | 39252,07 | 39002,30974 | 13359,05 | 36147,25316 | 30185,24 | 8191,872 | 11076,03 | 29163,3832 |
| Downregulated | ENSMUSG00000010054 | 305,6103768 | 181,903633 | 187,903596 | 155,134355 | 111,9547 | 159,2363062 | 71,561269 | 113,66702 | 119,589085 | 244,864108 | 181,074583 | 161,1563 | 162,284547 | 162,7732 | 105,9338 | 79,30448379 | 45,49803 | 116,9204757 | 82,69928 | 55,99913 | 42,51494 | 67,3021858 |
| Downregulated | ENSMUSG00000037685 | 150,1615 | 112,758977 | 50,833823 | 49,1419386 | 29,2608875 | 72,98330702 | 36,3485811 | 44,0001368 | 28,8295115 | 143,466636 | 80,6002715 | 80,57817 | 69,1211961 | 2,806435 | 7,964949 | 0,911545791 | 14,89026 | 17,8353268 | 19,01133 | 15,33309 | 13,28592 | 36,7918616 |
| Downregulated | ENSMUSG00000087408 | 84,59802819 | 69,1446557 | 84,4204561 | 34,6884273 | 57,2495624 | 23,88544594 | 10,2230384 | 24,4445204 | 19,2196743 | 111,10574 | 66,2467985 | 35,03399 | 17,0298599 | 6,548349 | 22,30186 | 20,96555318 | 5,790658 | 32,69809913 | 29,46756 | 10,6665 | 12,40019 | 27,8182368 |
| Downregulated | ENSMUSG00000002504 | 290,8057219 | 199,98762 | 288,663495 | 107,919551 | 117,04355 | 103,5035991 | 38,6203674 | 33,0001026 | 61,9300618 | 251,336287 | 156,78409 | 73,57137 | 56,098362 | 44,90296 | 25,48784 | 27,34637372 | 23,16263 | 24,77128722 | 29,46756 | 30,66619 | 24,80038 | 29,6129618 |
| Downregulated | ENSMUSG00000032492 | 2111,778279 | 1667,98185 | 2475,42563 | 1088,83119 | 727,705549 | 602,4440253 | 489,569951 | 407,001265 | 558,438316 | 2356,95187 | 1763,26895 | 971,3173 | 700,227769 | 448,0941 | 278,7732 | 404,726331 | 150,5571 | 474,6178631 | 367,8692 | 159,3308 | 227,6321 | 295,232255 |
| Downregulated | ENSMUSG00000037852 | 13973,47931 | 12069,4656 | 14572,9678 | 5506,78783 | 5317,84824 | 3164,821586 | 3426,98966 | 2550,78571 | 2998,2692 | 16811,4851 | 13958,2004 | 4660,396 | 3361,8947 | 1926,15 | 1716,447 | 1876,872783 | 868,5987 | 3143,971774 | 3154,93 | 151,9976 | 766,1547 | 2256,86663 |
| Downregulated | ENSMUSG00000027860 | 1123,038824 | 887,179121 | 1243,61317 | 502,982195 | 530,512612 | 493,6325493 | 371,437063 | 388,667875 | 439,91699 | 1213,53357 | 941,808652 | 410,7735 | 342,600711 | 275,0306 | 215,8501 | 255,2328214 | 149,7299 | 317,0724764 | 354,5613 | 258,6626 | 291,4045 | 221,648532 |
| Downregulated | ENSMUSG00000035936 | 33,83921127 | 22,3390426 | 35,4021268 | 14,4535114 | 5,08884999 | 1,326969219 | 0 | 3,66667807 | 8,54207749 | 43,1478604 | 30,9151726 | 16,64114 | 5,00878232 | 3,741914 | 0 | 1,823091581 | 0 | 3,963405955 | 10,45623 | 16,66641 | 7,085824 | 5,38417486 |
| Downregulated | ENSMUSG00000028445 | 34,89668663 | 22,3390426 | 22,693671 | 16,3806462 | 6,36106249 | 2,653938437 | 6,81535895 | 12,2222602 | 9,60983717 | 36,6756813 | 23,1863795 | 18,39284 | 0 | 0,935478 | 1,59299 | 5,469274744 | 0,827237 | 2,972554466 | 1,901133 | 6,666563 | 2,657184 | 3,58944991 |
| Downregulated | ENSMUSG00000006205 | 20443,11351 | 23487,9077 | 20122,9319 | 6875,05357 | 10933,3942 | 8499,237845 | 4475,41904 | 3654,45581 | 5708,24328 | 31288,6709 | 26780,2683 | 8760,249 | 8450,81754 | 3330,303 | 1586,618 | 2015,427743 | 328,413 | 2212,571375 | 1580,792 | 188,6637 | 1413,622 | 1988,55525 |
| Downregulated | ENSMUSG00000020814 | 1959,501828 | 2423,25424 | 1492,3358 | 763,1454 | 659,006074 | 574,5776717 | 269,206679 | 260,334143 | 419,629556 | 2969,65149 | 2785,67788 | 1093,06 | 646,13292 | 219,8374 | 138,5901 | 144,9357807 | 76,93303 | 295,2737437 | 146,3872 | 111,3316 | 268,3756 | 191,138208 |
| Downregulated | ENSMUSG00000036622 | 609,1058029 | 509,542924 | 327,696609 | 125,263765 | 201,009575 | 163,2172139 | 132,8995 | 68,4446572 | 177,248108 | 783,133666 | 547,640201 | 219,8383 | 142,249418 | 75,77375 | 41,41774 | 21,87709898 | 25,64434 | 43,59746551 | 53,23172 | 48,66591 | 81,48698 | 38,5865865 |
| Downregulated | ENSMUSG00000003070 | 111,034912 | 84,0373508 | 85,328203 | 35,6519947 | 53,4329249 | 27,86635359 | 22,7178632 | 14,6667123 | 19,2196743 | 146,702725 | 103,786651 | 50,79928 | 28,049181 | 27,12887 | 19,11588 | 25,52328214 | 4,963421 | 11,89021787 | 11,4068 | 11,99981 | 3,542912 | 11,6657122 |
| Downregulated | ENSMUSG00000028329 | 129,011993 | 99,9938098 | 127,084558 | 44,3241015 | 57,2495624 | 45,11695343 | 29,5332221 | 26,8889725 | 39,5071084 | 162,883173 | 130,28537 | 71,81967 | 71,124709 | 60,8061 | 43,01073 | 34,63874004 | 12,40855 | 16,84447531 | 41,82493 | 18,66638 | 14,17165 | 20,639337 |
| Downregulated | ENSMUSG00000036339 | 564,6918381 | 474,438715 | 501,076256 | 137,790142 | 192,104087 | 80,94512234 | 103,366277 | 78,2224654 | 130,266682 | 606,227438 | 430,60419 | 217,2107 | 224,393448 | 95,4188 | 72,48104 | 84,77375853 | 24,81711 | 126,8289906 | 92,20495 | 71,33222 | 75,28688 | 72,6863607 |
| Downregulated | ENSMUSG00000021973 | 826,9457255 | 851,011147 | 862,359498 | 269,798879 | 211,187275 | 264,0668745 | 94,2791321 | 107,55589 | 203,9421 | 949,252928 | 618,303453 | 430,918 | 448,786896 | 304,966 | 166,4674 | 163,1666965 | 63,69724 | 135,746654 | 158,7446 | 145,3311 | 121,3447 | 95,1204226 |
| Downregulated | ENSMUSG00000071253 | 347,9093909 | 496,777757 | 446,611445 | 188,859215 | 218,82055 | 144,6396448 | 127,220034 | 122,222602 | 123,860124 | 541,505648 | 535,494955 | 289,0304 | 285,500592 | 185,2247 | 60,53362 | 87,5083959 | 38,88013 | 68,36875273 | 116,9197 | 81,99872 | 67,31533 | 69,0969108 |
| Downregulated | ENSMUSG00000032431 | 4439,281529 | 4689,07142 | 4413,46514 | 1991,69387 | 2508,80305 | 1405,260403 | 1144,9803 | 641,668661 | 1230,05916 | 5980,29345 | 5711,57815 | 2941,979 | 3742,56215 | 1673,571 | 595,7782 | 884,1994169 | 330,0675 | 1167,223054 | 1139,729 | 599,324 | 626,2097 | 638,922084 |
| Downregulated | ENSMUSG00000047187 | 6463,289353 | 7010,20432 | 6894,33725 | 3741,53231 | 4287,35612 | 2673,842976 | 2296,77597 | 1837,00571 | 2931,00034 | 8527,09591 | 7641,56821 | 4903,882 | 5761,10143 | 3116,079 | 1801,672 | 2009,046923 | 777,6026 | 1915,315928 | 1866,913 | 1208,648 | 1755,513 | 1612,56037 |
| Downregulated | ENSMUSG00000030104 | 3989,854504 | 3925,28892 | 3281,50483 | 748,691888 | 1314,19551 | 729,8330702 | 654,274459 | 573,224004 | 935,357485 | 3626,57766 | 3723,07008 | 2710,755 | 1756,07908 | 977,5749 | 568,6974 | 589,7701265 | 461,5982 | 609,3736656 | 593,1535 | 365,3276 | 466,7787 | 437,015527 |
| Downregulated | ENSMUSG00000019189 | 4637,02942 | 4535,88941 | 3850,6621 | 1723,82212 | 2508,80305 | 1348,200726 | 1371,02304 | 1208,78154 | 1771,41332 | 4695,5659 | 5158,41738 | 3883,518 | 2403,21376 | 1915,86 | 1234,567 | 1413,807521 | 748,6493 | 1339,631213 | 1359,31 | 715,9888 | 868,0135 | 1234,77077 |
| Downregulated | ENSMUSG00000020253 | 927,405884 | 1049,935 | 798,817219 | 240,891856 | 260,803562 | 240,1814286 | 224,906845 | 81,8891435 | 207,145379 | 1132,63133 | 1087,55161 | 614,8465 | 565,992402 | 267,5468 | 99,56187 | 96,62385381 | 47,1525 | 112,9570697 | 121,6725 | 59,99906 | 101,8587 | 71,7889982 |
| Downregulated | ENSMUSG00000014470 | 593,2436726 | 523,371855 | 480,198078 | 154,170788 | 175,565325 | 124,7351066 | 130,627713 | 55,000171 | 81,1497361 | 594,361777 | 642,593946 | 387,1256 | 250,439116 | 134,7089 | 54,95815 | 71,10057167 | 81,89645 | 127,8198421 | 97,90835 | 68,6656 | 127,5448 | 74,4810856 |
| Downregulated | ENSMUSG00000026833 | 1233,016261 | 931,857206 | 793,370738 | 471,18447 | 276,070112 | 388,8019811 | 335,088482 | 253,000787 | 379,054688 | 1152,04787 | 1322,7274 | 680,5352 | 242,425064 | 565,0289 | 262,8433 | 324,5103015 | 199,3641 | 221,9507335 | 199,619 | 112,6649 | 105,4016 | 218,956444 |
| Downregulated | ENSMUSG00000050947 | 308,7828029 | 210,625259 | 192,44233 | 171,515001 | 73,7883249 | 115,446322 | 78,3766279 | 96,5558557 | 101,43717 | 275,06761 | 263,883081 | 189,1835 | 79,1387607 | 107,58 | 53,36516 | 72,92366325 | 28,12605 | 60,44194082 | 15,20906 | 6,666563 | 40,74349 | 43,0733989 |
| Downregulated | ENSMUSG00000038777 | 344,7369649 | 176,584813 | 214,228254 | 18,3077811 | 41,9830125 | 30,52029203 | 48,8434058 | 33,0001026 | 38,4393487 | 293,40545 | 350,003919 | 96,34347 | 18,0316164 | 46,77392 | 11,15093 | 3,646183163 | 13,23579 | 13,87192084 | 19,01133 | 11,33316 | 10,62874 | 10,7683497 |
| Downregulated | ENSMUSG00000038072 | 1120,923873 | 778,6752 | 941,333473 | 336,285031 | 363,852775 | 323,7804893 | 407,785644 | 409,445717 | 578,72575 | 1282,57015 | 1077,61459 | 401,1392 | 586,027532 | 540,7065 | 246,1169 | 242,4711803 | 143,112 | 153,5819808 | 293,725 | 120,6648 | 257,7469 | 382,276415 |
| Downregulated | ENSMUSG00000043673 | 80,36812678 | 84,0373508 | 99,8521524 | 55,8869106 | 67,4272624 | 86,25299921 | 36,3485811 | 47,6668148 | 104,640449 | 201,716247 | 85,0167248 | 105,9778 | 77,1352478 | 55,19322 | 51,77217 | 60,16202218 | 6,617895 | 46,57001997 | 35,17096 | 13,99978 | 7,085824 | 34,0997741 |
| Downregulated | ENSMUSG00000091337 | 618,6230811 | 731,869587 | 677,179143 | 562,723376 | 527,968187 | 565,2888871 | 304,419366 | 367,890033 | 682,298439 | 957,8825 | 700,007838 | 492,2275 | 436,765819 | 479,9004 | 261,2503 | 215,1248066 | 151,3843 | 268,5207535 | 265,2081 | 274,6624 | 178,9171 | 207,290732 |
| Downregulated | ENSMUSG00000036186 | 75,08075001 | 104,248866 | 118,914836 | 47,2148038 | 45,79965 | 57,0596764 | 15,9025042 | 30,5556505 | 36,3038293 | 181,221014 | 105,994878 | 67,44043 | 50,0878232 | 43,96748 | 11,15093 | 23,70019056 | 14,89026 | 23,78043573 | 21,86303 | 27,33291 | 4,42864 | 17,0498871 |

| | | | | | | | | | | | | | | | | | | | | | | | |
|---|---|---|---|---|---|---|---|---|---|---|---|---|---|---|---|---|---|---|---|---|---|---|---|
| Downregulated | ENSMUSG00000019857 | 129,011993 | 188,286216 | 229,659951 | 97,3203098 | 80,1493874 | 63,69452249 | 34,0767947 | 64,7779792 | 119,589085 | 201,716247 | 244,009041 | 218,0866 | 220,386422 | 165,5797 | 54,95815 | 91,15457906 | 33,91671 | 62,4236438 | 94,10608 | 33,99947 | 22,1432 | 37,6892241 |
| Downregulated | ENSMUSG00000110808 | 21,14950705 | 13,8289311 | 24,5091647 | 4,81783712 | 6,36106249 | 5,307876874 | 1,13589316 | 1,22222602 | 10,6775969 | 28,0461092 | 35,3316259 | 16,64114 | 31,0544504 | 11,22574 | 5,575465 | 0 | 4,136184 | 1,981702978 | 1,901133 | 0 | 4,42864 | 1,79472495 |
| Downregulated | ENSMUSG00000113132 | 66,6209472 | 64,8896 | 72,6197472 | 18,3077811 | 39,4385875 | 50,42483031 | 12,4948247 | 17,1111643 | 21,3551937 | 90,6105068 | 132,493597 | 89,33667 | 93,1633512 | 39,29009 | 15,1334 | 26,43482793 | 8,272368 | 17,8353268 | 15,20906 | 10,6665 | 7,085824 | 9,87098725 |
| Downregulated | ENSMUSG00000028669 | 298,2080494 | 291,471318 | 216,951495 | 119,482361 | 206,098425 | 159,2363062 | 152,209683 | 41,5556847 | 148,418596 | 393,724226 | 478,081063 | 359,9742 | 325,570851 | 152,483 | 92,39341 | 104,8277659 | 65,35171 | 89,17663399 | 49,42946 | 52,66584 | 32,77194 | 87,0441603 |
| Downregulated | ENSMUSG00000024761 | 200,9203169 | 177,648577 | 129,807798 | 163,806462 | 137,39895 | 57,0596764 | 80,6484142 | 46,4445888 | 65,1333408 | 179,063621 | 194,323942 | 113,8605 | 159,279278 | 93,54784 | 40,62124 | 72,92366325 | 19,02645 | 58,46023784 | 66,53965 | 23,33297 | 42,51494 | 56,5338361 |
| Downregulated | ENSMUSG00000062762 | 842,8078558 | 927,60215 | 807,894688 | 449,985987 | 562,317924 | 383,4941042 | 404,377964 | 366,667807 | 501,847052 | 1142,3396 | 971,619712 | 732,2103 | 707,240064 | 629,577 | 549,5815 | 597,0624929 | 148,0754 | 359,6790904 | 412,5459 | 169,3307 | 196,6316 | 258,440394 |
| Downregulated | ENSMUSG00000024790 | 308,7828029 | 278,706151 | 255,076862 | 203,312726 | 234,0871 | 120,7541989 | 198,781303 | 110,000342 | 197,535542 | 336,553311 | 263,883081 | 207,5764 | 264,463707 | 192,7085 | 157,706 | 154,0512386 | 67,83342 | 121,8747331 | 140,6838 | 65,99897 | 62,88669 | 105,888772 |
| Downregulated | ENSMUSG00000029346 | 102,5751092 | 147,863187 | 88,9591903 | 31,797725 | 39,4385875 | 17,25059984 | 9,08714527 | 15,8889383 | 22,4229534 | 84,1383277 | 98,2660845 | 102,4744 | 60,1053879 | 29,93531 | 21,50536 | 12,76164107 | 7,445132 | 26,7529902 | 12,35736 | 7,999875 | 1,771456 | 5,38417486 |
| Downregulated | ENSMUSG00000022663 | 2406,813902 | 3817,84876 | 2593,43272 | 1441,49687 | 1445,2334 | 760,3533623 | 711,069117 | 686,891024 | 1179,87445 | 2589,95032 | 2486,46317 | 2099,412 | 2243,93448 | 1437,83 | 841,8952 | 857,764589 | 325,9313 | 774,8458643 | 800,377 | 501,9922 | 651,8958 | 607,514397 |
| Downregulated | ENSMUSG00000064181 | 136,4143204 | 246,793233 | 97,1289119 | 88,648203 | 119,587975 | 67,67543015 | 49,979299 | 58,666849 | 60,8623021 | 173,670138 | 179,970469 | 134,005 | 130,22834 | 76,70923 | 39,02825 | 35,55028583 | 27,29882 | 34,67980211 | 48,47889 | 19,33303 | 31,88621 | 27,8182368 |
| Downregulated | ENSMUSG00000084094 | 58,16114438 | 95,7387541 | 59,0035446 | 24,0891856 | 52,1607124 | 27,86635359 | 19,3101837 | 20,7778424 | 28,8295115 | 62,5643975 | 61,8303453 | 70,06798 | 69,1211961 | 42,09653 | 15,9299 | 22,78864477 | 10,75408 | 17,8353268 | 17,1102 | 5,999906 | 15,9431 | 16,1525246 |
| Downregulated | ENSMUSG00000049940 | 1023,636141 | 982,917875 | 1101,09692 | 665,82509 | 893,093174 | 650,2149171 | 446,406011 | 299,445375 | 537,083122 | 899,632889 | 1123,98735 | 849,5742 | 842,477187 | 552,8677 | 446,0372 | 555,1313865 | 177,8559 | 472,6361602 | 469,5798 | 195,9969 | 185,1172 | 290,745443 |
| Downregulated | ENSMUSG00000001157 | 580,5539684 | 661,661167 | 763,415092 | 315,086548 | 343,497375 | 176,4869061 | 97,6868116 | 105,111438 | 156,960674 | 508,066056 | 533,286728 | 476,4622 | 371,651648 | 235,7406 | 160,892 | 200,5400739 | 55,42487 | 137,7283569 | 194,8661 | 74,6655 | 81,48698 | 113,067672 |
| Downregulated | ENSMUSG00000024072 | 1935,179895 | 2939,17975 | 2619,75738 | 1607,23046 | 1666,59837 | 1032,382052 | 829,202006 | 981,447495 | 1476,71165 | 2628,78339 | 2425,73694 | 1537,116 | 1961,43916 | 1183,38 | 681,0032 | 824,9489405 | 211,7726 | 825,3792902 | 783,2668 | 285,3289 | 440,2068 | 527,649137 |
| Downregulated | ENSMUSG00000022498 | 1642,259222 | 1810,52622 | 1253,59839 | 847,939333 | 999,959024 | 732,4870087 | 499,79299 | 371,556711 | 597,945424 | 1704,34048 | 1235,50279 | 1181,521 | 1015,78106 | 519,1905 | 395,858 | 407,4609684 | 215,0816 | 417,1484768 | 465,7776 | 277,9957 | 345,4339 | 322,153129 |
| Downregulated | ENSMUSG00000007570 | 270,7136902 | 284,02497 | 243,276153 | 152,243653 | 153,937712 | 141,9857064 | 77,2407348 | 37,8890067 | 89,6918136 | 329,002435 | 236,280248 | 189,1835 | 227,398717 | 160,9023 | 74,87053 | 61,98511376 | 38,05289 | 76,29556464 | 83,64985 | 51,33253 | 45,17213 | 35,8944991 |
| Downregulated | ENSMUSG00000049109 | 0 | 3,1912918 | 0 | 2,89070227 | 2,544425 | 0 | 0 | 8,55558215 | 9,60983717 | 19,4165372 | 34,2275126 | 14,0136 | 0 | 0 | 0,796495 | 0,911545791 | 9,926842 | 3,963405955 | 0 | 0,666656 | 1,771456 | 0 |
| Downregulated | ENSMUSG00000031994 | 79,31065142 | 285,088734 | 111,652861 | 352,665677 | 165,387625 | 100,8496606 | 86,32788 | 94,1114037 | 208,213139 | 722,726661 | 932,975746 | 465,0762 | 58,1018749 | 325,5465 | 141,7761 | 165,9013339 | 91,82329 | 178,353268 | 122,6231 | 5,999906 | 38,97203 | 81,6599854 |
| Downregulated | ENSMUSG00000070570 | 4,229901409 | 14,8926951 | 8,16972156 | 8,67210682 | 1,2722125 | 0 | 0 | 4,88890409 | 0 | 32,3608953 | 64,0385719 | 26,27549 | 7,01229525 | 6,548349 | 1,59299 | 14,58473265 | 0 | 3,963405955 | 3,802266 | 0 | 0,885728 | 1,79472495 |
| Downregulated | ENSMUSG00000061947 | 23,26445775 | 5,31881967 | 57,1880509 | 26,0163204 | 39,4385875 | 10,61575375 | 49,979299 | 25,6667465 | 30,9650309 | 81,9809347 | 183,282809 | 37,66154 | 4,00702586 | 30,87079 | 32,65629 | 21,87709898 | 22,33539 | 41,61576253 | 5,703399 | 1,333313 | 0 | 28,7155993 |
| Downregulated | ENSMUSG00000037031 | 307,7253275 | 120,205325 | 182,457115 | 90,5753379 | 323,141975 | 136,6778295 | 251,032388 | 242,000752 | 235,974891 | 618,0931 | 677,925572 | 214,5832 | 78,1370042 | 83,25758 | 140,1831 | 189,6015245 | 52,11592 | 151,6002778 | 96,95778 | 5,999906 | 33,65766 | 182,164583 |
| Downregulated | ENSMUSG00000043795 | 19,03455634 | 3,1912918 | 18,1549368 | 6,74497197 | 24,1720375 | 11,94272297 | 24,9896495 | 12,2222602 | 16,0163953 | 44,2265569 | 73,9755917 | 7,882647 | 11,0193211 | 0 | 7,964949 | 0 | 2,481711 | 15,85362382 | 18,06076 | 4,666594 | 7,085824 | 11,6657122 |
| Downregulated | ENSMUSG00000030223 | 31,72426057 | 82,9735868 | 27,2324052 | 36,6155621 | 86,5104499 | 62,36755328 | 26,1255426 | 78,2224654 | 95,030612 | 206,031033 | 230,759682 | 9,634347 | 19,0333728 | 10,29026 | 9,557939 | 8,203912116 | 14,06303 | 6,935960422 | 18,06076 | 8,666531 | 13,28592 | 61,9180109 |
| Downregulated | ENSMUSG00000032807 | 23,26445775 | 43,6143213 | 9,98521524 | 15,4170788 | 21,6276125 | 5,307876874 | 11,3589316 | 25,6667465 | 10,6775969 | 89,5318103 | 101,578424 | 11,38605 | 4,00702586 | 13,0967 | 12,74392 | 0,911545791 | 4,963421 | 0 | 4,752832 | 0 | 0 | 10,7683497 |
| Downregulated | ENSMUSG00000024041 | 93,057831 | 84,0373508 | 101,667646 | 90,5753379 | 19,0831875 | 98,19572218 | 113,589316 | 81,8891435 | 105,708209 | 170,434048 | 273,820101 | 44,66833 | 15,026347 | 32,74174 | 3,18598 | 17,31937002 | 10,75408 | 49,54257444 | 27,56643 | 2,666625 | 14,17165 | 67,3021858 |
| Downregulated | ENSMUSG00000040234 | 3444,197222 | 4164,6358 | 2683,29966 | 2925,3907 | 2884,10573 | 2565,0315 | 2412,63707 | 2905,23125 | 2758,02327 | 5494,88002 | 6760,48579 | 2067,881 | 1547,71374 | 1302,186 | 861,011 | 1005,435007 | 1160,613 | 1852,892284 | 1409,69 | 1044,65 | 1550,91 | 2089,95721 |
| Downregulated | ENSMUSG00000024601 | 405,0130599 | 502,096577 | 403,039597 | 310,268711 | 351,13065 | 367,5704736 | 260,119533 | 319,000992 | 343,818619 | 798,235417 | 1093,07218 | 357,3467 | 235,412769 | 270,3533 | 145,7586 | 144,0242349 | 102,5774 | 309,1456645 | 155,8929 | 191,997 | 239,1466 | 197,419745 |
| Downregulated | ENSMUSG00000038042 | 311,9552289 | 253,175816 | 239,645166 | 156,097923 | 174,293112 | 181,794783 | 123,812354 | 123,444828 | 184,722426 | 310,664595 | 357,732712 | 218,9624 | 57,1001185 | 55,19322 | 27,87732 | 16,40782423 | 53,77039 | 113,9479212 | 127,3759 | 124,6647 | 113,3732 | 96,9151476 |
| Downregulated | ENSMUSG00000032558 | 205,1502183 | 139,353075 | 186,088102 | 79,9760962 | 73,7883249 | 96,86875296 | 42,0280469 | 94,1114037 | 95,030612 | 211,424516 | 245,113155 | 116,488 | 32,0562069 | 56,1287 | 30,26681 | 20,96555318 | 18,19921 | 41,61576253 | 33,26983 | 57,99909 | 62,00096 | 50,2522987 |
| Downregulated | ENSMUSG00002076083 | 137,4717958 | 60,6345442 | 81,6972156 | 38,542697 | 35,62195 | 37,15513812 | 28,397329 | 40,3334587 | 32,0327906 | 148,860118 | 184,386923 | 102,4744 | 28,049181 | 42,09653 | 12,74392 | 5,469274744 | 15,7175 | 16,84447531 | 53,23172 | 20,66634 | 24,80038 | 14,3577996 |
| Downregulated | ENSMUSG00000105867 | 38,06911268 | 11,7014033 | 18,1549368 | 12,5263765 | 6,36106249 | 7,961815312 | 0 | 2,44445204 | 10,6775969 | 35,5969848 | 52,9974388 | 21,02039 | 3,00526939 | 3,741914 | 2,389485 | 0 | 7,445132 | 2,972554466 | 0,950566 | 0 | 0,885728 | 6,28153734 |
| Downregulated | ENSMUSG00000045790 | 289,7482465 | 276,578623 | 186,088102 | 187,895648 | 235,359312 | 183,1217522 | 120,404675 | 196,77839 | 158,028433 | 410,98337 | 460,41525 | 185,6801 | 93,1633512 | 241,3534 | 170,4499 | 89,33148748 | 142,2847 | 123,8564361 | 99,80948 | 34,66613 | 100,0873 | 87,0441603 |
| Downregulated | ENSMUSG00000039621 | 850,2101833 | 1008,44821 | 676,271396 | 560,796241 | 645,011737 | 911,6278532 | 706,525544 | 506,001573 | 673,756362 | 2139,05518 | 2270,05696 | 1257,72 | 343,602467 | 618,3512 | 347,2718 | 319,9525725 | 205,1547 | 326,9809913 | 222,4326 | 93,33188 | 378,2059 | 306,897967 |
| Downregulated | ENSMUSG00000015957 | 387,0359789 | 395,720183 | 416,6558 | 160,91576 | 338,408525 | 513,5370876 | 329,409016 | 156,444931 | 470,882021 | 1201,66791 | 1780,93477 | 674,4043 | 275,483028 | 219,8374 | 132,2182 | 157,6974218 | 6,617895 | 77,28641613 | 79,84759 | 41,99934 | 52,25795 | 93,3256977 |
| Downregulated | ENSMUSG00000033152 | 172,3684824 | 169,138465 | 111,652861 | 36,6155621 | 89,0548749 | 19,90453828 | 28,397329 | 19,5556163 | 41,6426277 | 355,969848 | 408,521924 | 86,70912 | 155,272252 | 34,6127 | 5,575465 | 19,1424616 | 0 | 28,73469318 | 19,9619 | 7,333219 | 7,085824 | 18,844612 |
| Downregulated | ENSMUSG00000097915 | 27,49435916 | 13,8289311 | 14,5239494 | 9,63567424 | 21,6276125 | 19,90453828 | 7,95125211 | 7,33335613 | 1,06775969 | 56,0922185 | 55,2056654 | 21,02039 | 19,0333728 | 8,419306 | 1,59299 | 0,911545791 | 4,136184 | 12,88106935 | 12,35736 | 2,666625 | 1,771456 | 1,79472495 |
| Downregulated | ENSMUSG00000079055 | 27,49435916 | 31,912918 | 30,8633926 | 15,4170788 | 20,3554 | 53,07876874 | 9,08714527 | 14,6667123 | 3,20327906 | 57,170915 | 79,4961582 | 19,26869 | 4,00702586 | 6,548349 | 3,18598 | 2,734637372 | 0,827237 | 8,917663399 | 0 | 0 | 4,42864 | 0 |
| Downregulated | ENSMUSG00000040841 | 141,7016972 | 142,544367 | 190,626836 | 113,700956 | 83,9660249 | 144,6396448 | 79,5125211 | 95,3336297 | 61,9300618 | 317,136774 | 291,485914 | 97,21932 | 66,1159267 | 51,45131 | 27,08083 | 50,13501849 | 43,84355 | 61,43279231 | 75,09475 | 71,99888 | 67,31533 | 29,6129618 |
| Downregulated | ENSMUSG00000054364 | 1705,707743 | 919,092039 | 1879,94371 | 960,676722 | 1029,21991 | 1541,938232 | 857,599334 | 421,667978 | 489,033936 | 2101,3008 | 2475,42204 | 1327,788 | 898,575549 | 1078,607 | 692,1541 | 517,7580091 | 188,61 | 433,0021006 | 351,7096 | 200,6635 | 266,6041 | 313,179505 |
| Downregulated | ENSMUSG00000038037 | 3249,621758 | 3295,54067 | 2696,00811 | 1299,85245 | 2894,28343 | 2501,336977 | 1144,9803 | 332,445478 | 1026,11706 | 3547,83282 | 5491,8596 | 2425,228 | 1253,19734 | 1166,542 | 739,1473 | 831,3297611 | 127,3945 | 775,8367157 | 574,1422 | 188,6637 | 228,5178 | 362,534441 |
| Downregulated | ENSMUSG00000031983 | 734,9453699 | 553,157246 | 531,939648 | 295,815199 | 333,319675 | 388,8019811 | 206,732555 | 233,44517 | 270,1432 | 620,250493 | 1054,42821 | 481,7173 | 353,620032 | 302,1595 | 201,5132 | 264,3482793 | 153,0388 | 194,2068918 | 222,4326 | 179,3305 | 177,1456 | 170,498871 |
| Downregulated | ENSMUSG00000035745 | 15,86213028 | 10,6376393 | 18,1549368 | 12,5263765 | 6,36106249 | 2,653938437 | 2,27178632 | 2,44445204 | 17,084155 | 118,656616 | 52,9974388 | 4,379248 | 7,01229525 | 1,870957 | 0,796495 | 5,469274744 | 5,790658 | 13,87192084 | 0 | 3,333281 | 2,657184 | 4,48681239 |
| Downregulated | ENSMUSG00000042988 | 19,03455634 | 31,912918 | 58,0957978 | 42,3969667 | 24,1720375 | 37,15513812 | 21,58197 | 14,6667123 | 38,4393487 | 148,860118 | 101,578424 | 33,28229 | 14,0245905 | 8,419306 | 23,09835 | 4,557728953 | 0 | 7,92681191 | 3,802266 | 0,666656 | 0,885728 | 5,38417486 |
| Downregulated | ENSMUSG00000039323 | 577,3815424 | 299,981429 | 309,541672 | 68,4132871 | 7,63327499 | 6,634846093 | 2,27178632 | 216,334006 | 131,334441 | 1213,53357 | 493,538649 | 35,90984 | 25,0439116 | 4,677392 | 5,575465 | 0 | 0,827237 | 11,89021787 | 8,555098 | 0 | 4,42864 | 10,7683497 |
| Downregulated | ENSMUSG00000031661 | 571,0366903 | 810,588118 | 587,312205 | 376,754863 | 525,423762 | 287,9523204 | 295,332221 | 180,889451 | 333,141022 | 2168,17998 | 935,183973 | 368,7327 | 111,194968 | 113,1929 | 43,80722 | 78,39293799 | 27,29882 | 187,2709314 | 75,09475 | 13,33313 | 20,37174 | 58,328561 |
| Downregulated | ENSMUSG00000034324 | 2411,043803 | 3306,17831 | 2461,80943 | 1226,62133 | 1094,10275 | 1330,950126 | 1445,99199 | 689,335476 | 983,40667 | 6610,25221 | 3653,51094 | 1303,264 | 451,792166 | 339,5787 | 169,6534 | 157,6974218 | 72,79684 | 393,3680411 | 187,2616 | 0,666656 | 148,8023 | 256,645669 |
| Downregulated | ENSMUSG00000017417 | 2609,84917 | 2405,17025 | 2209,45581 | 1555,19782 | 1373,9895 | 805,4703157 | 811,027715 | 774,891298 | 989,813229 | 4905,91172 | 2116,58521 | 1686,887 | 314,55153 | 306,8369 | 524,0937 | 715,5634456 | 220,8722 | 1157,314539 | 623,5716 | 21,99966 | 143,4879 | 533,033312 |
| Downregulated | ENSMUSG00000055725 | 34,89668663 | 14,8926951 | 33,5866331 | 14,4535114 | 7,63327499 | 5,307876874 | 15,9025042 | 9,77780817 | 17,084155 | 73,3513626 | 33,1233993 | 21,89624 | 23,0403987 | 9,354784 | 7,168455 | 2,734637372 | 13,23579 | 18,82617829 | 21,86303 | 1,999969 | 7,085824 | 4,48681239 |
| Downregulated | ENSMUSG00000019831 | 61,33357043 | 47,869377 | 101,667646 | 45,2876689 | 17,810975 | 38,48210734 | 20,4460768 | 37,8890067 | 38,4393487 | 126,207492 | 118,140124 | 60,43363 | 21,0368858 | 32,74174 | 43,80722 | 9,115457906 | 22,33539 | 33,68895062 | 16,15963 | 13,33313 | 18,60029 | 28,7155993 |
| Downregulated | ENSMUSG00000003949 | 28,55183451 | 34,0404459 | 25,4169115 | 28,9070227 | 5,08884999 | 23,88544594 | 17,0383974 | 13,4444862 | 12,8131162 | 72,2726661 | 57,4138921 | 38,53739 | 25,0439116 | 8,419306 | 37,43526 | 22,78864477 | 16,54474 | 38,64320806 | 8,555098 | 4,666594 | 15,9431 | 5,38417486 |
| Downregulated | ENSMUSG00000092268 | 9,517278171 | 3,1912918 | 9,98521524 | 0,96356742 | 10,1777 | 6,634846093 | 17,0383974 | 12,2222602 | 12,8131162 | 15,1017511 | 27,6028327 | 28,90304 | 21,0368858 | 3,741914 | 12,74392 | 18,23091581 | 4,136184 | 6,935960422 | 8,555098 | 5,33325 | 4,42864 | 0,89736248 |
| Downregulated | ENSMUSG00000087259 | 27,49435916 | 48,933141 | 29,9556457 | 26,9798879 | 66,1550499 | 34,50119968 | 68,1535895 | 68,4446572 | 102,50493 | 56,0922185 | 103,786651 | 93,71592 | 42,0737715 | 28,06435 | 42,21423 | 41,93110637 | 31,435 | 24,77128722 | 34,22039 | 13,99978 | 14,17165 | 12,5630747 |
| Downregulated | ENSMUSG00000048616 | 32,78173592 | 23,4028065 | 10,8929621 | 19,2713485 | 16,5387625 | 6,634846093 | 14,7666111 | 22,0000684 | 59,7945424 | 52,8561289 | 69,5591385 | 69,19213 | 15,026347 | 28,06435 | 35,04578 | 31,90410267 | 9,926842 | 27,74384169 | 6,653965 | 3,999938 | 3,542912 | 6,28153734 |
| Downregulated | ENSMUSG00000097090 | 60,27609508 | 38,2955016 | 39,940861 | 39,5062644 | 73,7883249 | 50,42483031 | 120,404675 | 62,3335271 | 39,5071084 | 78,7448452 | 115,931897 | 142,7635 | 49,0860668 | 5,61287 | 43,80722 | 52,86965586 | 73,62408 | 99,08514888 | 76,99589 | 16,66641 | 13,28592 | 43,0733989 |
| Downregulated | ENSMUSG00000004661 | 45,47144015 | 36,1679738 | 29,0478989 | 19,2713485 | 21,6276125 | 51,75179953 | 48,8434058 | 41,5556847 | 54,455744 | 74,4300591 | 67,3509118 | 80,57817 | 29,0509375 | 18,70957 | 40,62124 | 26,43482793 | 40,53461 | 12,88106935 | 32,31926 | 3,333281 | 7,971552 | 34,0997741 |
| Downregulated | ENSMUSG00000109378 | 105,7475352 | 48,933141 | 51,7415699 | 26,0163204 | 50,8884999 | 55,73270718 | 73,8330553 | 67,2224312 | 39,5071084 | 124,050099 | 101,578424 | 83,20572 | 28,049181 | 36,48366 | 30,26681 | 37,37337742 | 55,42487 | 58,46023784 | 50,38002 | 19,33303 | 2,657184 | 29,6129618 |
| Downregulated | ENSMUSG00000024186 | 70,85084861 | 56,3794885 | 34,4943799 | 31,797725 | 49,6162874 | 74,31027624 | 112,453423 | 59,8890751 | 45,9136665 | 153,174904 | 96,0578579 | 113,8605 | 22,0386422 | 53,32227 | 76,46352 | 69,27748009 | 23,98987 | 55,48768337 | 77,94645 | 17,99972 | 15,05738 | 43,0733989 |
| Downregulated | ENSMUSG00000061576 | 26,43688381 | 19,1477508 | 37,2176204 | 34,6884273 | 33,077525 | 49,09786109 | 55,6587648 | 31,7778766 | 21,3551937 | 38,8330743 | 69,5591385 | 57,80608 | 26,0456681 | 18,70957 | 13,54041 | 45,57728953 | 10,75408 | 17,8353268 | 11,4068 | 29,99953 | 29,22902 | 46,6628488 |
| Downregulated | ENSMUSG00000040390 | 27,49435916 | 22,3390426 | 46,2950888 | 31,797725 | 38,166375 | 37,15513812 | 38,6203674 | 35,4445546 | 33,1005503 | 48,5413429 | 72,8714784 | 59,55778 | 31,0544504 | 24,32244 | 23,09835 | 31,90410267 | 8,272368 | 13,87192084 | 15,20906 | 19,99969 | 45,17213 | 12,5630747 |
| Downregulated | ENSMUSG00000031737 | 118,4372395 | 107,440157 | 68,081013 | 157,06149 | 78,8771749 | 153,9284294 | 122,676461 | 95,3336297 | 125,995643 | 112,184437 | 197,636282 | 111,2329 | 88,1545689 | 138,4508 | 98,76537 | 69,27748009 | 37,22566 | 84,22237655 | 64,63852 | 25,33294 | 34,54339 | 69,9942732 |
| Downregulated | ENSMUSG00000003378 | 20,09203169 | 15,956459 | 4,5387342 | 34,6884273 | 24,1720375 | 13,26969219 | 26,1255426 | 24,4445204 | 19,2196743 | 24,8100197 | 56,3097787 | 35,03399 | 7,01229525 | 20,58052 | 2,389485 | 3,646183163 | 8,272368 | 8,917663399 | 3,802266 | 4,666594 | 12,40019 | 9,87098725 |
| Downregulated | ENSMUSG00000020882 | 161,7937289 | 98,9300459 | 65,3577725 | 87,6846356 | 115,771337 | 183,1217522 | 130,627713 | 68,4446572 | 56,5912633 | 105,712258 | 250,633721 | 149,7703 | 56,098362 | 71,09636 | 86,02145 | 133,0856854 | 67,00618 | 83,23152506 | 67,49022 | 29,99953 | 33,65766 | 56,5338361 |
| Downregulated | ENSMUSG00000052040 | 1644,374173 | 1550,96782 | 1991,59657 | 1173,62512 | 1352,36189 | 1630,84517 | 1915,11586 | 982,669721 | 1018,64274 | 2214,56393 | 3464,70756 | 1742,065 | 999,752952 | 806,3824 | 1337,315 | 1524,104562 | 1052,245 | 1148,396876 | 858,3615 | 427,3267 | 360,4913 | 894,67039 |
| Downregulated | ENSMUSG00000039377 | 31,72426057 | 41,4867934 | 59,9112914 | 19,2713485 | 48,344075 | 47,77089187 | 42,0280469 | 12,2222602 | 16,0163953 | 35,5969848 | 107,098991 | 67,44043 | 30,0526939 | 11,22574 | 28,67382 | 48,3119269 | 6,617895 | 15,85362382 | 26,61586 | 5,999906 | 1,771456 | 16,1525246 |
| Downregulated | ENSMUSG00000003068 | 204,092743 | 105,312629 | 92,5901777 | 139,717276 | 129,765675 | 260,0859668 | 152,209683 | 128,333732 | 249,855766 | 172,591441 | 584,07594 | 275,0168 | 138,242392 | 345,1915 | 146,5551 | 90,24303327 | 74,45132 | 145,6551689 | 107,414 | 103,3317 | 117,8018 | 67,3021858 |
| Downregulated | ENSMUSG00000052632 | 60,27609508 | 19,1477508 | 30,8633926 | 24,0891856 | 59,7939874 | 74,31027624 | 64,74591 | 102,666986 | 106,775969 | 91,6892033 | 240,696701 | 131,3775 | 52,0913362 | 67,35444 | 45,40021 | 40,10801479 | 10,75408 | 60,44194082 | 31,36869 | 27,33291 | 37,20058 | 50,2522987 |
| Downregulated | ENSMUSG00000042364 | 439,9097466 | 253,175816 | 327,696609 | 275,580283 | 306,603212 | 456,4774112 | 403,242071 | 468,112566 | 665,214284 | 448,737748 | 1214,52464 | 754,9824 | 445,781627 | 523,8679 | 272,4013 | 349,1220378 | 124,9128 | 323,0175854 | 210,0752 | 176,6639 | 270,147 | 271,003468 |
| Downregulated | ENSMUSG00000050910 | 931,6357854 | 780,802728 | 784,29327 | 419,151829 | 609,389787 | 927,5514838 | 648,594993 | 557,335066 | 612,89406 | 1338,66237 | 2913,75502 | 848,6984 | 428,751767 | 277,8371 | 141,7761 | 94,80076223 | 167,9291 | 267,529902 | 178,7065 | 599,324 | 627,0954 | 304,20588 |
| Downregulated | ENSMUSG00000042804 | 1073,337483 | 656,342347 | 758,876358 | 1144,7181 | 617,023062 | 1226,119558 | 813,299501 | 826,224791 | 968,458035 | 1637,4613 | 3524,32968 | 1233,196 | 397,697316 | 1042,123 | 422,9388 | 307,1909314 | 305,2504 | 446,8740215 | 314,6375 | 354,6611 | 591,6663 | 434,323439 |
| Downregulated | ENSMUSG00000042812 | 32,78173592 | 21,2752787 | 29,0478989 | 16,3806462 | 38,166375 | 43,78998421 | 34,0767947 | 26,8889725 | 43,7781471 | 51,7774324 | 157,888203 | 54,30268 | 9,01580818 | 33,67722 | 25,48784 | 31,90410267 | 23,98987 | 36,66150509 | 12,35736 | 24,66628 | 5,314368 | 13,4604372 |
| Downregulated | ENSMUSG00000015981 | 17,97708099 | 17,0202229 | 14,5239494 | 9,63567424 | 25,44425 | 9,28878453 | 17,0383974 | 25,6667465 | 25,6262325 | 40,9904673 | 145,742957 | 70,06798 | 13,022834 | 14,96765 | 7,964949 | 7,292366325 | 9,926842 | 10,89936638 | 4,752832 | 0,666656 | 0 | 6,28153734 |
| Downregulated | ENSMUSG00000012017 | 256,9665106 | 272,323567 | 177,010634 | 255,345367 | 169,204262 | 542,7304104 | 249,896495 | 209,00065 | 297,904952 | 445,501658 | 1827,30753 | 1089,557 | 137,240636 | 246,0308 | 136,9971 | 189,6015245 | 112,5042 | 202,1337037 | 143,5355 | 146,6644 | 131,9735 | 96,9151476 |
| Downregulated | ENSMUSG00000031154 | 392,3233557 | 249,984524 | 327,696609 | 221,620508 | 176,837537 | 333,0692739 | 241,945243 | 244,445204 | 194,332263 | 330,081132 | 836,917888 | 794,3957 | 288,505862 | 357,3527 | 272,4013 | 218,7709898 | 159,6567 | 174,389862 | 153,0412 | 99,33178 | 150,5738 | 138,193822 |
| Downregulated | ENSMUSG00000042594 | 929,5208347 | 787,185311 | 599,112914 | 382,536267 | 610,661999 | 697,985809 | 547,500502 | 614,779689 | 813,63288 | 997,794271 | 2385,98886 | 2559,233 | 700,227769 | 691,3185 | 478,6935 | 434,8073421 | 349,9212 | 607,3919626 | 335,55 | 122,6648 | 270,147 | 383,173778 |
| Downregulated | ENSMUSG00000008305 | 240,046905 | 195,732564 | 127,992304 | 177,296406 | 206,098425 | 279,9905051 | 168,112187 | 194,333937 | 132,402201 | 305,271112 | 937,392199 | 786,513 | 189,331972 | 230,1277 | 128,2357 | 104,8277659 | 67,00618 | 147,6368718 | 57,98456 | 34,66613 | 115,1446 | 119,349209 |
| Downregulated | ENSMUSG00000000531 | 127,9545176 | 104,248866 | 127,084558 | 57,8140454 | 114,499125 | 94,21481452 | 101,094491 | 42,7779108 | 80,0819764 | 158,568387 | 581,867714 | 536,8959 | 99,17389 | 90,7414 | 52,56867 | 29,1694653 | 45,49803 | 94,13089144 | 67,49022 | 40,66603 | 33,65766 | 51,1496612 |
| Downregulated | ENSMUSG00000020525 | 72,96579931 | 52,1244328 | 46,2950888 | 27,9434553 | 55,9773499 | 49,09786109 | 14,7666111 | 25,6667465 | 74,743178 | 116,499223 | 207,573302 | 208,4522 | 50,0878232 | 79,51566 | 42,21423 | 19,1424616 | 23,16263 | 34,67980211 | 36,12153 | 7,333219 | 14,17165 | 36,7918616 |
| Downregulated | ENSMUSG00000002910 | 83,54055283 | 84,0373508 | 59,9112914 | 20,2349159 | 67,4272624 | 80,94512234 | 40,8921537 | 48,8890409 | 35,2360696 | 100,318775 | 238,488475 | 177,7975 | 50,0878232 | 84,19306 | 37,43526 | 22,78864477 | 37,22566 | 23,78043573 | 30,41813 | 37,99941 | 14,17165 | 23,3314244 |
| Downregulated | ENSMUSG00000037573 | 1072,280007 | 1077,59287 | 1569,49429 | 604,156775 | 725,161124 | 717,8903473 | 579,305511 | 620,890819 | 904,392454 | 1684,92395 | 3498,93508 | 2074,012 | 807,41571 | 1133,8 | 810,8319 | 409,28406 | 202,673 | 332,9261002 | 200,5695 | 297,3287 | 285,2044 | 389,455315 |
| Downregulated | ENSMUSG00000027552 | 214,6674965 | 144,671895 | 206,96628 | 67,4497197 | 80,1493874 | 106,1575375 | 67,0176963 | 58,666849 | 105,708209 | 245,942804 | 538,807295 | 448,435 | 249,43736 | 172,128 | 101,9514 | 100,270037 | 48,80697 | 94,13089144 | 70,34192 | 25,33294 | 43,40067 | 60,123286 |
| Downregulated | ENSMUSG00000052681 | 6639,887737 | 9395,16306 | 7522,49806 | 3560,38163 | 3975,66406 | 3665,088982 | 3495,14325 | 3708,23375 | 5833,17116 | 9667,27812 | 17410,7628 | 14917,47 | 7895,84445 | 6433,285 | 3158,899 | 3381,834883 | 1516,325 | 3065,694506 | 2131,17 | 1201,315 | 2720,956 | 3136,28186 |
| Downregulated | ENSMUSG00000008429 | 825,8882502 | 708,46678 | 879,606688 | 325,685789 | 431,280037 | 338,3771507 | 251,032388 | 297,000923 | 458,068905 | 1077,61781 | 1789,76767 | 1461,793 | 790,385851 | 632,3834 | 285,9417 | 289,8715614 | 126,5672 | 228,8866939 | 207,2235 | 152,6643 | 290,5188 | 244,979956 |
| Downregulated | ENSMUSG00000036478 | 8687,160019 | 8509,04771 | 7332,77897 | 2714,36943 | 3225,05868 | 3760,630766 | 2819,28682 | 2796,45314 | 4329,76553 | 11202,2632 | 25185,9287 | 22600,43 | 7176,58331 | 7897,309 | 3195,538 | 3539,532305 | 1583,331 | 3198,468606 | 1869,764 | 1256,647 | 2165,605 | 2413,90506 |
| Downregulated | ENSMUSG00000063049 | 105,7475352 | 104,248866 | 81,6972156 | 24,0891856 | 25,44425 | 69,00239937 | 42,0280469 | 37,8890067 | 54,455744 | 139,15185 | 261,674854 | 200,5696 | 72,1264654 | 74,83827 | 26,28433 | 32,81564846 | 17,37197 | 34,67980211 | 21,86303 | 24,66628 | 27,45757 | 26,0235118 |
| Downregulated | ENSMUSG00000082838 | 99,40268312 | 56,3794885 | 46,2950888 | 4,81783712 | 19,0831875 | 2,653938437 | 9,08714527 | 8,55558215 | 5,33879843 | 45,3052534 | 209,781529 | 151,522 | 45,0790409 | 26,19339 | 4,77897 | 18,23091581 | 2,481711 | 6,935960422 | 9,505665 | 5,33325 | 9,743008 | 7,17889982 |
| Downregulated | ENSMUSG00000068263 | 221,0123486 | 165,947174 | 156,132456 | 73,2311242 | 72,5161124 | 59,71361484 | 106,773957 | 47,6668148 | 69,4043796 | 224,368874 | 538,807295 | 473,8347 | 76,1334913 | 74,83827 | 29,47031 | 36,46183163 | 32,26224 | 38,64320806 | 19,9619 | 15,33309 | 37,20058 | 43,0733989 |
| Downregulated | ENSMUSG00000053175 | 1391,637564 | 989,300459 | 729,828459 | 432,641773 | 433,824462 | 765,6612391 | 385,067781 | 310,44541 | 536,015362 | 1552,24428 | 3700,98781 | 2873,663 | 539,946734 | 354,5463 | 312,226 | 275,2868288 | 156,3478 | 279,4201198 | 214,828 | 176,6639 | 155,0024 | 227,930069 |

| | | | | | | | | | | | | | | | | | | | | | | | |
|---|---|---|---|---|---|---|---|---|---|---|---|---|---|---|---|---|---|---|---|---|---|---|---|
| Downregulated | ENSMUSG00000097025 | 21,14950705 | 11,7014033 | 12,7084558 | 2,89070227 | 8,90548749 | 5,307876874 | 0 | 4,88890409 | 3,20327906 | 25,8887162 | 60,726232 | 48,17173 | 9,01580818 | 2,806435 | 2,389485 | 6,380820534 | 0 | 0,990851489 | 0 | 1,999969 | 2,657184 | 0 |
| Downregulated | ENSMUSG00000020105 | 633,427736 | 363,807265 | 334,958584 | 274,616716 | 188,28745 | 261,4129361 | 214,683807 | 215,11178 | 217,822976 | 418,534246 | 859,000154 | 702,4315 | 164,28806 | 388,2235 | 209,4782 | 209,6555318 | 77,76026 | 179,3441195 | 128,3265 | 70,66556 | 130,202 | 104,094047 |
| Downregulated | ENSMUSG00000078619 | 488,5536128 | 312,746597 | 287,755748 | 196,567754 | 111,9547 | 314,4917048 | 136,307179 | 105,111438 | 156,960674 | 448,737748 | 719,881877 | 626,2325 | 171,300355 | 237,6115 | 88,41094 | 52,86965586 | 61,21553 | 139,7100599 | 82,69928 | 81,33206 | 135,5164 | 76,2758106 |
| Downregulated | ENSMUSG00000003154 | 489,6110881 | 288,280026 | 286,848001 | 167,660732 | 147,57665 | 205,6802289 | 156,753256 | 128,333732 | 186,857945 | 371,071599 | 612,782886 | 536,8959 | 208,365345 | 185,2247 | 149,7411 | 120,3240444 | 120,7766 | 112,9570697 | 104,5623 | 105,3317 | 123,1162 | 110,375585 |
| Downregulated | ENSMUSG00000049686 | 132,184419 | 159,56459 | 92,5901777 | 84,7939333 | 133,582312 | 57,0596764 | 60,2023374 | 31,7778766 | 91,827333 | 196,322765 | 358,836825 | 253,9964 | 74,1299784 | 65,48349 | 76,46352 | 75,65830062 | 21,50816 | 55,48768337 | 39,92379 | 17,33306 | 57,57232 | 43,9707614 |
| Downregulated | ENSMUSG00000042680 | 65,56347184 | 158,500826 | 36,3098736 | 32,7612924 | 40,7108 | 50,42483031 | 51,1151921 | 6,11113011 | 44,8459068 | 120,814009 | 206,469189 | 175,1699 | 17,0298599 | 21,516 | 43,80722 | 13,67318686 | 28,95329 | 21,79873275 | 12,35736 | 22,66631 | 23,02893 | 13,4604372 |
| Downregulated | ENSMUSG00000036158 | 365,8864719 | 382,955016 | 312,264913 | 408,552588 | 208,64285 | 368,8974428 | 251,032388 | 216,334006 | 218,890736 | 481,098643 | 704,424291 | 437,9248 | 164,28806 | 144,0637 | 203,1062 | 128,5279565 | 122,4311 | 155,5636837 | 115,0185 | 167,3307 | 90,34426 | 139,091184 |
| Downregulated | ENSMUSG00000023307 | 1006,716535 | 1081,84792 | 773,400308 | 576,21332 | 851,110162 | 720,5442857 | 748,553591 | 887,336092 | 987,677709 | 1335,42628 | 2082,3577 | 1418,001 | 889,559741 | 704,4152 | 500,1988 | 515,9349175 | 362,3297 | 558,8402397 | 479,0855 | 407,327 | 403,892 | 532,135949 |
| Downregulated | ENSMUSG00000057614 | 250,6216585 | 330,830583 | 279,586027 | 172,478569 | 147,57665 | 159,2363062 | 143,122538 | 149,111575 | 201,806581 | 359,205938 | 556,473108 | 314,43 | 221,388179 | 143,1282 | 107,5268 | 81,12757537 | 60,38829 | 101,0668519 | 97,90835 | 94,66519 | 69,97251 | 126,528109 |
| Downregulated | ENSMUSG00000031016 | 323,5874578 | 239,346885 | 206,058533 | 127,1909 | 91,5992999 | 119,4272297 | 120,404675 | 122,222602 | 84,3530152 | 620,250493 | 759,629956 | 408,146 | 170,298599 | 166,5152 | 118,6777 | 76,56984641 | 25,64434 | 82,24067357 | 78,89702 | 88,66528 | 57,57232 | 116,657122 |
| Downregulated | ENSMUSG00000035172 | 138,5292712 | 99,9938098 | 89,8669372 | 21,1984833 | 22,899825 | 15,92363062 | 7,95125211 | 30,5556505 | 9,60983717 | 142,387939 | 316,88052 | 144,5152 | 12,0210776 | 3,741914 | 11,94742 | 3,646183163 | 5,790658 | 4,954257444 | 8,555098 | 6,666563 | 3,542912 | 12,5630747 |
| Downregulated | ENSMUSG00000091811 | 195,6329402 | 221,262898 | 147,054988 | 41,4333992 | 62,3384124 | 75,63724546 | 49,979299 | 44,0001368 | 48,0491859 | 297,720237 | 459,311136 | 211,9556 | 93,1633512 | 108,5155 | 23,89485 | 56,51583902 | 28,12605 | 41,61576253 | 51,33059 | 28,66622 | 59,34378 | 50,2522987 |
| Downregulated | ENSMUSG00000050445 | 32,78173592 | 21,2752787 | 19,0626836 | 2,89070227 | 50,8884999 | 15,92363062 | 27,2614358 | 8,55558215 | 35,2360696 | 21,5739302 | 48,5809856 | 67,44043 | 28,049181 | 83,25758 | 32,65629 | 38,28492321 | 8,272368 | 8,917663399 | 16,15963 | 0,666656 | 0 | 7,17889982 |
| Downregulated | ENSMUSG00000031609 | 118,4372395 | 138,289311 | 121,638077 | 58,7776129 | 80,1493874 | 59,71361484 | 65,8818032 | 24,4445204 | 98,2338911 | 55,013522 | 165,616996 | 225,9692 | 222,389935 | 195,515 | 77,26001 | 151,3166012 | 15,7175 | 70,35045571 | 79,84759 | 19,33303 | 23,91466 | 23,3314244 |
| Downregulated | ENSMUSG00000020656 | 29,60930986 | 18,0839869 | 9,98521524 | 7,70853939 | 30,5331 | 18,57756906 | 6,81535895 | 4,88890409 | 19,2196743 | 3,23608953 | 41,9563057 | 26,27549 | 32,0562069 | 12,16122 | 29,47031 | 13,67318686 | 5,790658 | 0,990851489 | 0,950566 | 4,666594 | 12,40019 | 9,87098725 |
| Downregulated | ENSMUSG00000040010 | 1570,350898 | 990,364222 | 1047,53985 | 832,522254 | 3451,51251 | 386,1480426 | 382,795994 | 234,667396 | 2447,3052 | 2310,56792 | 2580,3128 | 4567,556 | 2289,01352 | 1075,8 | 2373,555 | 575,1853939 | 695,7062 | 1625,987293 | 245,2462 | 175,9973 | 183,3457 | 929,667527 |
| Downregulated | ENSMUSG00000022742 | 267,5412641 | 311,682833 | 174,287393 | 146,462248 | 101,777 | 141,9857064 | 74,9689484 | 108,778116 | 103,57269 | 239,470625 | 433,91653 | 253,9964 | 73,1282219 | 119,7412 | 106,7303 | 114,8547696 | 62,87 | 110,9753667 | 71,29249 | 75,99881 | 93,88717 | 61,9180109 |
| Downregulated | ENSMUSG00000016984 | 26,43688381 | 46,8056131 | 25,4169115 | 14,4535114 | 1,2722125 | 14,5966614 | 0 | 9,77780817 | 16,0163953 | 24,8100197 | 45,2686457 | 31,53059 | 9,01580818 | 11,22574 | 11,94742 | 4,557728953 | 12,40855 | 7,92681191 | 1,901133 | 2,666625 | 0,885728 | 9,87098725 |
| Downregulated | ENSMUSG00000062232 | 380,6911268 | 273,387331 | 194,257824 | 164,77003 | 160,298775 | 191,0835675 | 189,694157 | 129,555958 | 202,87434 | 266,438038 | 422,875397 | 317,9334 | 104,182672 | 298,4176 | 160,0955 | 124,8817733 | 57,07934 | 109,9845153 | 90,30382 | 73,33219 | 88,5728 | 83,4547104 |
| Downregulated | ENSMUSG00000040488 | 8258,882502 | 8771,7974 | 6460,43426 | 4162,61127 | 4815,32431 | 5597,156164 | 3395,18465 | 3814,56741 | 4947,99838 | 9926,16528 | 8177,06317 | 7260,794 | 3631,36718 | 3109,53 | 1661,488 | 3201,348817 | 837,9909 | 1399,082302 | 1561,781 | 331,3282 | 962,7864 | 1152,21342 |
| Downregulated | ENSMUSG00000037174 | 1830,489835 | 2201,99134 | 1610,34289 | 1025,23574 | 1342,18419 | 1600,324878 | 1222,22104 | 1035,22544 | 1636,8756 | 2088,35644 | 2530,6277 | 1662,363 | 1518,6628 | 1392,927 | 930,3061 | 958,034626 | 435,9538 | 766,9190523 | 653,9897 | 406,6603 | 456,1499 | 805,831505 |
| Downregulated | ENSMUSG00000030878 | 840,6929051 | 946,749901 | 749,79889 | 468,293768 | 473,26305 | 646,2340095 | 571,354259 | 458,334758 | 706,856912 | 1213,53357 | 1316,10306 | 856,581 | 451,792166 | 567,8354 | 183,1938 | 273,4637372 | 125,74 | 294,2828922 | 191,0639 | 113,3316 | 251,5468 | 243,185231 |
| Downregulated | ENSMUSG00000042042 | 1136,786004 | 1631,81387 | 1155,56173 | 803,615232 | 889,276537 | 824,0478848 | 638,371955 | 784,669106 | 1077,36952 | 1626,67434 | 1754,43605 | 1317,278 | 1025,79862 | 1011,252 | 531,2621 | 410,1956058 | 263,0613 | 630,1815469 | 365,0175 | 229,9964 | 588,1234 | 464,833763 |
| Downregulated | ENSMUSG00000041037 | 850,2101833 | 804,205534 | 689,887598 | 453,840257 | 728,977762 | 508,2292107 | 270,342572 | 282,334211 | 383,325727 | 1188,72355 | 1106,32154 | 1071,164 | 410,72015 | 558,4806 | 347,2718 | 446,6574374 | 190,2645 | 294,2828922 | 400,1885 | 155,9976 | 193,0887 | 269,208743 |
| Downregulated | ENSMUSG00000024201 | 1037,383321 | 888,242885 | 816,972156 | 365,192054 | 686,994749 | 375,5322889 | 415,736896 | 235,889622 | 383,325727 | 1132,63133 | 962,786805 | 848,6984 | 517,908092 | 412,546 | 238,9485 | 432,9842506 | 156,3478 | 280,4109713 | 333,6488 | 145,9977 | 157,6596 | 313,179505 |
| Downregulated | ENSMUSG00000000957 | 60005,38139 | 44835,5223 | 48568,9947 | 25384,2202 | 31833,3011 | 25085,02611 | 17587,0338 | 12531,4834 | 28467,541 | 53100,9931 | 57848,9127 | 55462,31 | 29881,3936 | 29957,76 | 16498,6 | 24150,49418 | 7569,217 | 18460,55409 | 19016,08 | 5469,915 | 10959,11 | 12374,6286 |
| Downregulated | ENSMUSG00000035354 | 1429,706676 | 1383,95688 | 1106,5434 | 719,784866 | 1156,44116 | 850,5872691 | 615,654092 | 407,001265 | 728,212106 | 1349,44933 | 1407,74447 | 1296,258 | 738,294514 | 896,1883 | 574,2729 | 601,6202218 | 242,3804 | 540,0140614 | 640,6818 | 264,6625 | 300,2618 | 398,42894 |
| Downregulated | ENSMUSG00000027367 | 907,3138523 | 779,738964 | 530,124155 | 383,499835 | 623,384124 | 411,3604578 | 307,827046 | 360,556676 | 470,882021 | 967,590769 | 1021,30481 | 1101,819 | 578,01348 | 425,6427 | 269,2153 | 308,1024772 | 156,3478 | 222,941585 | 279,4665 | 142,6644 | 201,946 | 247,672044 |
| Downregulated | ENSMUSG00000028911 | 397,6107325 | 327,639292 | 249,630381 | 79,0125288 | 143,760012 | 127,389045 | 97,6868116 | 55,000171 | 98,2338911 | 327,923739 | 357,732712 | 399,3875 | 230,403987 | 105,7091 | 52,56867 | 92,97767064 | 42,18908 | 60,44194082 | 63,68796 | 23,99963 | 38,0863 | 53,8417486 |
| Downregulated | ENSMUSG00000011158 | 817,4284473 | 584,0064 | 472,028357 | 215,839103 | 413,469062 | 281,3174743 | 199,917196 | 129,555958 | 223,161774 | 723,805358 | 676,821458 | 812,7885 | 494,867693 | 296,5466 | 221,4256 | 185,0437955 | 93,47776 | 192,2251888 | 191,0639 | 87,99863 | 109,8303 | 168,704146 |
| Downregulated | ENSMUSG00000028862 | 1588,327979 | 1091,4218 | 744,352409 | 533,816353 | 852,382374 | 690,0239937 | 722,428049 | 364,223354 | 540,286401 | 1244,81577 | 1447,49255 | 1723,672 | 828,452596 | 687,5766 | 599,7607 | 900,6072411 | 191,0917 | 512,2702197 | 565,5871 | 200,6635 | 178,9171 | 436,118164 |
| Downregulated | ENSMUSG00000028826 | 826,9457255 | 829,735868 | 677,179143 | 399,880481 | 664,094924 | 554,6731334 | 432,775293 | 277,445307 | 440,98475 | 829,517616 | 944,016879 | 1185,025 | 624,094277 | 634,2543 | 371,1666 | 404,726331 | 223,3539 | 413,1850708 | 373,5726 | 156,6642 | 207,2604 | 296,129618 |
| Downregulated | ENSMUSG00000052934 | 143,8166479 | 141,480603 | 84,4204561 | 65,5225848 | 50,8884999 | 70,32936859 | 31,8050084 | 51,3334929 | 82,2174958 | 157,48969 | 175,554016 | 127,8741 | 59,1036314 | 35,54818 | 23,09835 | 28,25791951 | 37,22566 | 32,69809913 | 60,83626 | 25,33294 | 32,77194 | 49,3549363 |
| Downregulated | ENSMUSG00000032855 | 7323,016815 | 7577,1905 | 3901,49592 | 2220,05934 | 2866,29476 | 2522,568485 | 2603,46712 | 2070,45088 | 2697,16097 | 8343,7175 | 8083,21353 | 6548,728 | 2119,71668 | 2269,471 | 1375,547 | 1772,045017 | 871,9076 | 1539,783214 | 1186,307 | 1006,651 | 1388,822 | 1549,745 |
| Downregulated | ENSMUSG00000029178 | 3220,012448 | 3531,69626 | 2001,58178 | 581,031157 | 1558,46031 | 1423,837972 | 1510,7379 | 1081,67003 | 1168,1291 | 3868,20568 | 5193,749 | 4329,325 | 922,617704 | 1223,606 | 889,6849 | 912,4573364 | 743,6859 | 831,3243991 | 718,6283 | 611,9904 | 738,6972 | 1088,50069 |
| Downregulated | ENSMUSG00000021303 | 1331,361469 | 1657,34421 | 1135,5913 | 328,576492 | 1092,83054 | 603,7709945 | 566,810686 | 279,889759 | 303,243751 | 2488,55285 | 1806,32937 | 1815,636 | 912,600139 | 368,5785 | 277,9767 | 427,5149758 | 175,3742 | 357,6973875 | 327,9454 | 84,66535 | 416,2922 | 390,352678 |
| Downregulated | ENSMUSG00000041120 | 26014,95114 | 30299,1881 | 18603,3637 | 12670,9116 | 17626,5042 | 9862,035233 | 8444,22974 | 4887,68186 | 11570,244 | 33346,8239 | 37070,6043 | 23233,66 | 15951,9699 | 9305,204 | 4203,104 | 6333,420153 | 2162,397 | 5347,625485 | 4170,135 | 950,6518 | 2388,808 | 3451,25609 |
| Downregulated | ENSMUSG00000034707 | 17394,41207 | 18124,4099 | 13493,6568 | 5552,0755 | 8919,48183 | 6207,562005 | 5094,48081 | 3412,45505 | 6076,62037 | 26430,2219 | 23397,2651 | 17553,78 | 9877,31874 | 6347,221 | 3106,33 | 3087,405593 | 1835,639 | 4567,825363 | 4231,922 | 1871,304 | 3433,968 | 3067,18495 |
| Downregulated | ENSMUSG00000034761 | 1145,245807 | 1671,17314 | 1019,3997 | 389,281239 | 541,962524 | 359,6086582 | 345,31152 | 361,778902 | 485,830657 | 1636,3826 | 1435,3473 | 1271,734 | 688,206691 | 635,1898 | 298,6856 | 352,768221 | 105,8863 | 349,7705756 | 313,6869 | 355,9944 | 267,4899 | 218,059082 |
| Downregulated | ENSMUSG00000018415 | 1068,050106 | 1347,7889 | 874,160207 | 446,131717 | 676,817049 | 386,1480426 | 322,593657 | 344,667738 | 357,699495 | 1393,67589 | 1232,19045 | 907,3803 | 747,310323 | 417,2234 | 443,6477 | 370,9991368 | 305,2504 | 323,0175854 | 431,5572 | 255,996 | 236,4894 | 343,689829 |
| Downregulated | ENSMUSG00000050730 | 1075,452433 | 1338,21503 | 1078,40325 | 428,787504 | 901,998662 | 817,4130387 | 512,287814 | 432,668012 | 731,415385 | 1429,27287 | 1557,90388 | 1206,921 | 895,570279 | 610,8674 | 505,7743 | 621,6742292 | 196,0551 | 370,5784568 | 398,2874 | 234,663 | 244,4609 | 384,07114 |
| Downregulated | ENSMUSG00000030557 | 2970,448265 | 4444,40572 | 2802,21449 | 1629,39251 | 3088,93195 | 2177,556488 | 2204,76862 | 1885,89475 | 2441,9664 | 4227,41162 | 4881,28494 | 4296,919 | 2464,3209 | 2360,212 | 1757,864 | 2077,412857 | 837,9909 | 1643,82262 | 1470,526 | 474,6593 | 749,3259 | 1774,08562 |
| Downregulated | ENSMUSG00000030849 | 3619,738131 | 3675,30439 | 4224,65379 | 1444,38757 | 2791,23422 | 1633,499108 | 1892,398 | 1741,67208 | 2321,30956 | 5369,75122 | 5821,98948 | 4190,065 | 2253,95205 | 1545,41 | 1148,546 | 1336,326129 | 605,5374 | 1451,597431 | 1441,059 | 221,3299 | 412,7493 | 1217,72088 |
| Downregulated | ENSMUSG00000017831 | 2173,111849 | 2931,7334 | 2750,47293 | 1257,45549 | 1769,64759 | 1159,771097 | 1438,04074 | 1520,44917 | 1665,70511 | 3336,4083 | 3588,36825 | 3413,186 | 1993,49536 | 1954,214 | 966,1484 | 1204,151989 | 558,3849 | 1062,192796 | 970,5284 | 654,6564 | 800,6981 | 1221,31033 |
| Downregulated | ENSMUSG00000074622 | 1451,913659 | 704,211724 | 1185,51737 | 202,349159 | 187,015237 | 346,3389661 | 116,996995 | 235,889622 | 282,956317 | 917,970729 | 1337,08122 | 1671,121 | 330,579633 | 254,4501 | 341,6963 | 239,7365429 | 140,6303 | 258,6122386 | 155,8929 | 93,33188 | 162,0882 | 153,448984 |
| Downregulated | ENSMUSG00000029208 | 238,9894296 | 249,984524 | 212,412761 | 77,0853939 | 58,5217749 | 58,38664562 | 30,6691153 | 37,8890067 | 79,0142167 | 242,706715 | 380,919092 | 289,9062 | 213,374127 | 102,9026 | 72,48104 | 55,60429323 | 77,76026 | 71,34130719 | 42,77549 | 56,66578 | 57,57232 | 46,6628488 |
| Downregulated | ENSMUSG00000015337 | 123,7246162 | 80,846059 | 100,759899 | 7,70853939 | 33,077525 | 14,5966614 | 5,67946579 | 9,77780817 | 14,9486356 | 114,34183 | 173,345789 | 140,136 | 108,189698 | 42,09653 | 15,1334 | 17,31937002 | 12,40855 | 7,92681191 | 11,4068 | 1,999969 | 13,28592 | 2,69208743 |
| Downregulated | ENSMUSG00000039208 | 6620,853181 | 3963,58442 | 4906,37167 | 640,772337 | 1460,49995 | 1230,100466 | 959,829719 | 631,890853 | 1278,10834 | 5267,27505 | 9994,43367 | 6069,638 | 4029,0645 | 1672,635 | 508,9603 | 785,7524715 | 285,3967 | 983,9155284 | 589,3512 | 411,3269 | 996,444 | 563,543636 |
| Downregulated | ENSMUSG00000031681 | 1727,914726 | 2364,74723 | 1232,72021 | 466,366633 | 858,743437 | 678,0812707 | 449,813691 | 470,557018 | 662,011005 | 2510,12678 | 3532,05847 | 2480,406 | 1438,52228 | 762,4149 | 332,9349 | 386,4954152 | 198,5368 | 399,31315 | 298,4779 | 189,997 | 272,8042 | 311,38478 |
| Downregulated | ENSMUSG00000028756 | 2531,595993 | 3320,00724 | 1847,26482 | 613,792449 | 1066,11407 | 695,3318706 | 435,04708 | 290,889793 | 591,538866 | 3190,78427 | 3671,17675 | 2723,893 | 1829,2073 | 651,093 | 322,5805 | 448,480529 | 205,982 | 398,3222985 | 333,6488 | 142,6644 | 268,3756 | 328,434667 |
| Downregulated | ENSMUSG00000027602 | 1650,719025 | 2215,82027 | 1534,99991 | 818,068743 | 837,115824 | 897,0311918 | 513,423708 | 576,890682 | 715,398989 | 1810,05274 | 2565,95933 | 1940,007 | 1351,36947 | 737,157 | 373,5561 | 445,7458916 | 466,5616 | 583,6115269 | 505,7014 | 479,9925 | 400,3491 | 438,810251 |
| Downregulated | ENSMUSG00000037514 | 483,266236 | 511,670452 | 335,866331 | 79,0125288 | 218,82055 | 128,7160142 | 115,861102 | 118,555924 | 129,198922 | 477,862554 | 704,424291 | 453,6901 | 293,514644 | 158,0958 | 121,8637 | 146,7588723 | 97,61395 | 134,7558025 | 145,4367 | 92,66522 | 113,3732 | 133,707009 |
| Downregulated | ENSMUSG00000049339 | 1798,765574 | 1507,35349 | 1046,63211 | 439,386745 | 550,868012 | 493,6325493 | 405,513857 | 342,223286 | 534,947603 | 1277,17667 | 2103,33585 | 1799,871 | 750,315592 | 655,7704 | 480,2865 | 441,1881627 | 305,2504 | 397,331447 | 395,4357 | 243,9962 | 356,0627 | 318,563679 |
| Downregulated | ENSMUSG00000029030 | 1579,868176 | 1576,49815 | 1074,77226 | 379,645565 | 555,956862 | 443,207719 | 326,001336 | 277,445307 | 428,171634 | 1734,54399 | 2751,45037 | 1750,824 | 752,319105 | 597,7707 | 217,4431 | 254,3212756 | 151,3843 | 282,3926743 | 160,6457 | 151,9976 | 259,5183 | 234,211607 |
| Downregulated | ENSMUSG00000020087 | 261,196412 | 207,433967 | 180,641621 | 52,9962083 | 27,988675 | 67,67543015 | 28,397329 | 18,3333903 | 23,4907131 | 249,178894 | 420,667171 | 284,6511 | 99,17389 | 56,1287 | 29,47031 | 29,1694653 | 11,58132 | 37,65235657 | 23,76416 | 21,99966 | 30,11475 | 24,2287869 |
| Downregulated | ENSMUSG00000027829 | 8612,079269 | 6777,24002 | 7151,22961 | 2437,82558 | 3618,17235 | 4036,640363 | 2426,26779 | 2103,45098 | 2813,54677 | 11761,028 | 13272,5461 | 14298,25 | 3929,89061 | 2901,854 | 2487,454 | 1648,986335 | 1087,816 | 2251,214583 | 1244,292 | 785,3211 | 744,8973 | 1319,12284 |
| Downregulated | ENSMUSG00000020134 | 2309,526169 | 3033,85474 | 2862,12579 | 504,90933 | 917,265212 | 834,6636385 | 561,13122 | 498,668217 | 807,226322 | 4362,24868 | 4621,81831 | 4610,473 | 1968,45145 | 1152,509 | 464,3566 | 649,9321487 | 295,3236 | 446,8740215 | 382,1277 | 232,663 | 318,8621 | 324,845217 |
| Downregulated | ENSMUSG00000032370 | 212,5525458 | 147,863187 | 211,505014 | 54,9233432 | 43,255225 | 46,44392265 | 48,8434058 | 37,8890067 | 59,7945424 | 266,438038 | 324,609313 | 186,556 | 115,201993 | 87,93497 | 63,7196 | 35,55028583 | 22,33539 | 33,68895062 | 62,73739 | 29,33288 | 36,31485 | 52,9443862 |
| Downregulated | ENSMUSG00000025158 | 663,0370459 | 442,525797 | 584,588965 | 183,077811 | 213,7317 | 261,4129361 | 148,802004 | 108,778116 | 145,215317 | 746,457984 | 739,755917 | 531,6408 | 294,516401 | 144,9992 | 99,56187 | 85,68530432 | 73,62408 | 122,8655846 | 79,84759 | 119,3315 | 77,94407 | 94,2230601 |
| Downregulated | ENSMUSG00000021476 | 1216,096655 | 1105,25073 | 881,422182 | 156,097923 | 325,6864 | 282,6444436 | 71,561269 | 83,1113695 | 181,519147 | 1232,95011 | 1540,23807 | 890,7391 | 517,908092 | 238,547 | 103,5443 | 119,4124986 | 36,39842 | 124,8472876 | 91,25438 | 85,99866 | 110,716 | 98,7098725 |
| Downregulated | ENSMUSG00000034789 | 514,9904966 | 436,143213 | 443,888205 | 82,8667985 | 108,138062 | 102,1766298 | 92,0073458 | 28,1111985 | 79,0142167 | 512,380842 | 653,635079 | 472,083 | 268,470733 | 126,2896 | 86,81795 | 41,93110637 | 52,11592 | 93,14003995 | 56,08342 | 33,99947 | 51,37222 | 46,6628488 |
| Downregulated | ENSMUSG00000030055 | 169,1960564 | 128,715436 | 88,9591903 | 32,7612924 | 64,8828374 | 88,90693765 | 62,4741237 | 40,3334587 | 49,1169455 | 125,128795 | 211,989755 | 210,2039 | 123,216045 | 62,67705 | 46,9932 | 66,54284272 | 52,11592 | 37,65235657 | 72,24305 | 23,33297 | 23,91466 | 43,0733989 |
| Downregulated | ENSMUSG00000078676 | 576,324067 | 324,448 | 402,13185 | 154,170788 | 208,64285 | 189,7565983 | 151,07379 | 79,4446914 | 154,825154 | 434,714693 | 824,772642 | 799,6508 | 303,532209 | 342,3851 | 181,6008 | 160,4320592 | 89,34158 | 123,8564361 | 129,277 | 110,6649 | 69,08679 | 120,246572 |
| Downregulated | ENSMUSG00000031734 | 199,8628416 | 178,712341 | 143,424001 | 56,850478 | 77,6049624 | 72,98330702 | 72,6971621 | 26,8889725 | 66,2011005 | 215,739302 | 407,417811 | 292,5338 | 109,191455 | 115,9993 | 58,94063 | 90,24303327 | 22,33539 | 26,7529902 | 16,15963 | 5,999906 | 21,25747 | 19,7419745 |
| Downregulated | ENSMUSG00000017692 | 377,5187008 | 341,468223 | 200,612052 | 80,9396636 | 117,04355 | 111,4654144 | 51,1151921 | 64,7779792 | 139,876519 | 308,507202 | 662,467985 | 442,3041 | 247,433847 | 160,9023 | 96,37589 | 82,03912116 | 27,29882 | 101,0668519 | 51,33059 | 21,99966 | 38,0863 | 56,5338361 |
| Downregulated | ENSMUSG00000036087 | 1111,406595 | 1379,70182 | 809,710181 | 315,086548 | 366,3972 | 371,5513812 | 241,945243 | 273,778629 | 499,711533 | 1236,1862 | 1500,48999 | 1699,148 | 534,937952 | 584,674 | 351,2543 | 266,1713709 | 190,2645 | 231,8592484 | 219,5809 | 159,3308 | 246,2324 | 199,21447 |
| Downregulated | ENSMUSG00000004591 | 1107,176694 | 1702,02229 | 1070,23352 | 476,965875 | 463,08535 | 492,3055801 | 449,813691 | 476,668148 | 680,16292 | 1383,96762 | 1952,07233 | 1926,869 | 857,503534 | 934,5429 | 410,9914 | 636,2589619 | 234,108 | 400,3040015 | 321,2915 | 303,3286 | 358,7198 | 389,455315 |
| Downregulated | ENSMUSG00000015214 | 853,3826093 | 940,367318 | 636,330535 | 209,094131 | 352,402862 | 216,2959826 | 271,478465 | 246,889656 | 374,78365 | 983,771216 | 1417,68149 | 1474,931 | 804,410441 | 608,061 | 204,6992 | 302,6332025 | 90,99605 | 235,8226543 | 197,7178 | 73,99885 | 180,6885 | 256,645669 |
| Downregulated | ENSMUSG00000025858 | 709,5659614 | 765,910032 | 681,717877 | 322,795087 | 454,179862 | 347,6659353 | 245,352922 | 161,333835 | 268,007681 | 859,721118 | 1333,76888 | 1271,734 | 793,39112 | 660,4477 | 190,3623 | 226,9749019 | 108,368 | 240,7769118 | 257,6035 | 160,6642 | 162,0882 | 192,932933 |
| Downregulated | ENSMUSG00000026608 | 510,7605952 | 524,435619 | 387,607901 | 185,968513 | 264,6202 | 204,3532597 | 110,181636 | 133,222636 | 216,755216 | 615,935707 | 676,821458 | 673,5284 | 441,774601 | 428,4491 | 164,078 | 106,6508575 | 71,14237 | 114,9387727 | 101,7106 | 98,66513 | 113,3732 | 136,399097 |
| Downregulated | ENSMUSG00000100153 | 2989,482821 | 3469,99795 | 2707,80882 | 848,902901 | 1826,89715 | 959,3987451 | 537,277464 | 503,557121 | 1153,18046 | 3332,09352 | 4718,98028 | 4035,915 | 2748,81974 | 1605,281 | 629,231 | 807,6295705 | 222,5267 | 639,0992103 | 560,8342 | 183,3305 | 285,2044 | 447,783876 |
| Downregulated | ENSMUSG00000004455 | 2849,896074 | 3454,04149 | 2339,26361 | 1155,31734 | 1947,75734 | 1356,162541 | 820,11486 | 772,446846 | 1566,40346 | 3017,11414 | 4428,59848 | 3690,831 | 2697,73016 | 2132,891 | 1061,728 | 1195,948077 | 466,5616 | 1103,808559 | 780,4151 | 447,993 | 681,1248 | 766,347556 |
| Downregulated | ENSMUSG00000017299 | 501,243317 | 603,154151 | 473,84385 | 180,187108 | 245,537012 | 199,0453828 | 105,638064 | 46,4445888 | 133,469961 | 498,357787 | 653,635079 | 514,1238 | 528,927413 | 240,4179 | 152,1305 | 122,1471359 | 51,28868 | 109,9845153 | 106,4634 | 69,33225 | 62,88669 | 81,6599854 |
| Downregulated | ENSMUSG00000071637 | 12892,7395 | 14098,0634 | 8750,67954 | 3160,50115 | 5929,78246 | 4357,766914 | 1709,5192 | 1427,55999 | 3446,72827 | 10435,31 | 12250,1372 | 8390,64 | 8013,04996 | 3473,431 | 1431,301 | 2234,198733 | 470,6978 | 1279,189272 | 1173,95 | 217,3299 | 265,7184 | 670,329771 |
| Downregulated | ENSMUSG00000104923 | 358,4841444 | 356,360918 | 320,434634 | 179,223541 | 170,476475 | 152,6014601 | 110,181636 | 113,66702 | 137,740999 | 367,83551 | 390,856111 | 275,8927 | 312,548017 | 116,9348 | 86,81795 | 80,21602958 | 44,67079 | 81,24982208 | 86,50155 | 39,99938 | 83,25843 | 88,8388853 |
| Downregulated | ENSMUSG00000029416 | 764,5546797 | 786,121547 | 537,386129 | 130,081602 | 267,164625 | 176,4869061 | 86,32788 | 74,5557873 | 202,87434 | 625,643975 | 739,755917 | 702,4315 | 432,758793 | 257,2566 | 121,0672 | 176,8398834 | 62,87 | 111,9662182 | 117,8702 | 37,33275 | 53,14368 | 61,9180109 |
| Downregulated | ENSMUSG00000036615 | 301,3804754 | 338,276931 | 170,656406 | 50,105506 | 75,0605374 | 43,78998421 | 26,1255426 | 35,4445546 | 40,5748681 | 250,25759 | 346,691579 | 308,2991 | 142,249418 | 65,48349 | 43,80722 | 25,52328214 | 4,136184 | 13,87192084 | 19,9619 | 8,666531 | 13,28592 | 25,1261494 |
| Downregulated | ENSMUSG00000032285 | 453,6569261 | 468,056131 | 250,538128 | 70,340422 | 131,037887 | 63,69452249 | 65,8818032 | 41,5556847 | 66,2011005 | 320,372863 | 511,204462 | 437,049 | 281,493567 | 86,06401 | 61,33011 | 57,42738481 | 27,29882 | 26,7529902 | 57,03399 | 16,66641 | 23,91466 | 17,9472495 |
| Downregulated | ENSMUSG00000000738 | 1096,60194 | 878,669009 | 742,536915 | 242,818991 | 255,714712 | 258,7589976 | 162,432722 | 147,889349 | 136,67324 | 791,763238 | 998,118431 | 827,678 | 582,020506 | 286,2564 | 136,2006 | 125,7933191 | 89,34158 | 127,8198421 | 109,3151 | 130,6646 | 129,3163 | 116,657122 |
| Downregulated | ENSMUSG00000031823 | 872,4171657 | 552,093482 | 562,803041 | 124,300198 | 207,370637 | 86,25299921 | 60,2023374 | 67,2224312 | 129,198922 | 473,547768 | 716,569537 | 544,7785 | 359,630571 | 218,9019 | 86,02145 | 112,1201322 | 47,97974 | 95,12174293 | 79,84759 | 49,99922 | 143,4879 | 57,4311986 |
| Downregulated | ENSMUSG00000022175 | 7781,961118 | 6062,39066 | 5246,77673 | 1902,0821 | 2074,97859 | 1378,721018 | 1142,70852 | 576,890682 | 1280,24386 | 5708,46193 | 6138,87 | 7664,561 | 5202,12132 | 3604,398 | 2132,217 | 2762,895291 | 462,4254 | 1039,403212 | 1443,911 | 527,9918 | 725,4112 | 760,066018 |
| Downregulated | ENSMUSG00000003135 | 253,7940846 | 206,370203 | 197,888811 | 80,9396636 | 142,4878 | 114,1193528 | 60,2023374 | 26,8889725 | 61,9300618 | 156,410994 | 287,06946 | 277,6444 | 208,365345 | 123,4831 | 101,1549 | 70,18902588 | 14,06303 | 60,44194082 | 50,38002 | 29,99953 | 26,57184 | 36,7918616 |
| Downregulated | ENSMUSG00000022200 | 5928,206825 | 6397,4763 | 6432,29411 | 2764,47494 | 2923,54432 | 1981,165043 | 1784,48815 | 1882,22807 | 2700,36425 | 6847,56544 | 7289,35606 | 6548,728 | 5505,65353 | 3473,431 | 2046,992 | 2117,520872 | 978,6212 | 2321,565038 | 1601,705 | 969,3182 | 1204,59 | 1599,09993 |
| Downregulated | ENSMUSG00000035953 | 1175,912592 | 1089,29427 | 1161,00821 | 317,013683 | 515,246062 | 379,5131965 | 246,488815 | 210,222876 | 430,307153 | 1195,19573 | 1281,87555 | 1282,244 | 755,324374 | 536,9646 | 324,1734 | 307,1909314 | 162,1384 | 269,511605 | 255,7024 | 147,9977 | 170,0598 | 183,061945 |
| Downregulated | ENSMUSG00000041740 | 3345,852015 | 3335,9637 | 3219,77804 | 1172,66156 | 1647,51519 | 1206,21502 | 890,540236 | 721,113353 | 1078,43728 | 3340,72309 | 4112,82208 | 3858,994 | 2941,15698 | 1768,99 | 1095,181 | 1268,871741 | 545,9763 | 879,8761221 | 864,0649 | 547,3248 | 594,3235 | 593,156598 |
| Downregulated | ENSMUSG00000049969 | 458,9443029 | 347,850806 | 398,500863 | 108,883119 | 139,943375 | 82,27209155 | 93,143239 | 57,444623 | 172,977069 | 503,75127 | 581,867714 | 478,2139 | 349,613006 | 289,9983 | 116,2883 | 142,2011433 | 39,70737 | 99,08514888 | 78,89702 | 33,33281 | 46,05786 | 70,8916357 |
| Downregulated | ENSMUSG00000026499 | 1930,949993 | 1727,55263 | 1438,77874 | 554,051269 | 847,293524 | 559,9810103 | 492,977631 | 408,223491 | 854,207749 | 2177,88825 | 2331,88731 | 1981,172 | 1338,34664 | 1000,962 | 397,451 | 459,4190785 | 266,3703 | 527,132992 | 405,8919 | 418,6601 | 430,4638 | 497,138812 |

| | | | | | | | | | | | | | | | | | | | | | | | |
|---|---|---|---|---|---|---|---|---|---|---|---|---|---|---|---|---|---|---|---|---|---|---|---|
| Downregulated | ENSMUSG00000090036 | 40,18406339 | 34,0404459 | 39,940861 | 4,81783712 | 8,90548749 | 1,326969219 | 3,40767947 | 0 | 0 | 47,4626464 | 46,372759 | 57,80608 | 35,0614763 | 2,806435 | 2,389485 | 2,734637372 | 0 | 0,990851489 | 0 | 0 | 0,885728 | 1,79472495 |
| Downregulated | ENSMUSG00000032199 | 2430,07836 | 2417,93542 | 1849,98806 | 528,998516 | 459,268712 | 566,6158564 | 370,30117 | 348,334416 | 545,625199 | 2754,99088 | 2806,65603 | 2847,387 | 1796,14934 | 854,0918 | 384,7071 | 346,3874004 | 210,1182 | 352,74313 | 342,2039 | 291,3288 | 348,0911 | 279,07973 |
| Downregulated | ENSMUSG00000061130 | 737,0603206 | 802,078006 | 570,972762 | 178,259973 | 227,726037 | 210,9881058 | 131,763606 | 119,77815 | 229,568332 | 755,087556 | 834,709661 | 936,2833 | 579,015237 | 427,5136 | 175,2289 | 195,982345 | 108,368 | 173,3990105 | 123,5736 | 102,6651 | 102,7444 | 122,041297 |
| Downregulated | ENSMUSG00000100347 | 175,5409085 | 177,648577 | 114,376102 | 27,9434553 | 48,344075 | 41,13604578 | 17,0383974 | 25,6667465 | 30,9650309 | 160,72578 | 185,491036 | 206,7005 | 97,1703771 | 66,41897 | 22,30186 | 25,52328214 | 18,19921 | 25,76213871 | 8,555098 | 15,99975 | 23,02893 | 12,5630747 |
| Downregulated | ENSMUSG00000019158 | 491,7260388 | 523,371855 | 361,283242 | 88,648203 | 100,504787 | 110,1384451 | 53,3869784 | 35,4445546 | 62,9978215 | 435,79339 | 628,240473 | 646,3771 | 452,793922 | 153,4185 | 37,43526 | 54,69274744 | 72,79684 | 71,34130719 | 71,29249 | 58,66575 | 74,40115 | 49,3549363 |
| Downregulated | ENSMUSG00000029221 | 771,9570072 | 863,776314 | 673,548155 | 259,199637 | 372,758262 | 323,7804893 | 160,160935 | 156,444931 | 284,024076 | 923,364212 | 1127,29969 | 1182,397 | 733,285732 | 384,4816 | 247,7099 | 153,1396928 | 167,9291 | 231,8592484 | 141,6344 | 167,9974 | 222,3177 | 191,138208 |
| Downregulated | ENSMUSG00000039662 | 1494,212673 | 1148,86505 | 1192,77935 | 443,241015 | 297,697725 | 317,1456432 | 198,781303 | 149,111575 | 321,395665 | 1447,61072 | 1031,24183 | 1518,723 | 584,024019 | 368,5785 | 259,6574 | 331,8026678 | 216,7361 | 250,6854267 | 232,8888 | 247,9961 | 276,3471 | 175,883046 |
| Downregulated | ENSMUSG00000019861 | 733,8878945 | 618,046846 | 462,043142 | 199,458457 | 201,009575 | 130,0429834 | 77,2407348 | 121,000376 | 146,283077 | 652,611388 | 647,010399 | 640,2461 | 267,468976 | 261,9339 | 149,7411 | 129,4395023 | 134,8396 | 134,7558025 | 91,25438 | 117,3315 | 135,5164 | 100,504597 |
| Downregulated | ENSMUSG00000036955 | 635,5426867 | 684,00021 | 436,62623 | 132,972305 | 197,192937 | 124,7351066 | 57,9305511 | 74,5557873 | 116,385806 | 690,365766 | 586,284167 | 605,2121 | 362,63584 | 184,2892 | 74,87053 | 60,16202218 | 34,74395 | 62,4236438 | 92,20495 | 73,99885 | 69,08679 | 62,8153734 |
| Downregulated | ENSMUSG00000006526 | 574,2091163 | 577,623816 | 384,88466 | 158,025058 | 165,387625 | 123,4081373 | 48,8434058 | 66,0002052 | 89,6918136 | 615,935707 | 520,037368 | 430,0422 | 201,353049 | 90,7414 | 61,33011 | 88,41994169 | 61,21553 | 62,4236438 | 78,89702 | 72,66553 | 109,8303 | 84,3520729 |
| Downregulated | ENSMUSG00000025907 | 1302,809634 | 1763,7206 | 1302,61672 | 460,585229 | 917,265212 | 638,2721942 | 514,559601 | 601,335203 | 763,448175 | 1706,49788 | 1406,64036 | 1810,381 | 1031,80916 | 880,2852 | 468,339 | 520,4926465 | 260,5796 | 486,508081 | 450,5685 | 267,9958 | 353,4055 | 361,637078 |
| Downregulated | ENSMUSG00000083964 | 49,70134156 | 70,2084196 | 58,0957978 | 8,67210682 | 26,7164625 | 19,90453828 | 4,54357263 | 4,88890409 | 18,1519147 | 70,1152731 | 54,1015521 | 56,93023 | 36,0632327 | 28,99983 | 5,575465 | 10,0270037 | 5,790658 | 0,990851489 | 5,703399 | 1,999969 | 2,657184 | 0,89736248 |
| Downregulated | ENSMUSG00000021171 | 3232,702152 | 3924,22515 | 3403,1429 | 1320,08737 | 1859,97467 | 1039,016898 | 996,1783 | 936,225133 | 1713,7543 | 4201,5229 | 3540,89138 | 3544,564 | 2283,00298 | 1932,698 | 1115,093 | 1178,628707 | 636,9724 | 1010,668519 | 979,0835 | 579,9909 | 683,782 | 718,787344 |
| Downregulated | ENSMUSG00000052253 | 842,8078558 | 840,373508 | 1019,3997 | 531,889218 | 660,278287 | 509,5561799 | 383,931887 | 168,667191 | 327,802223 | 1083,0113 | 836,917888 | 992,3377 | 685,201422 | 448,0941 | 342,4928 | 420,2226095 | 111,677 | 274,4658624 | 284,2194 | 116,6648 | 140,8308 | 163,319971 |
| Downregulated | ENSMUSG00000019777 | 1244,64849 | 1310,55717 | 1464,19565 | 673,533629 | 748,060949 | 378,1862273 | 286,245076 | 305,556505 | 486,898417 | 1527,43426 | 1067,67757 | 1318,154 | 1112,95143 | 652,9639 | 492,2339 | 424,7803384 | 103,4046 | 339,8620607 | 467,6787 | 189,3304 | 245,3467 | 325,742579 |
| Downregulated | ENSMUSG00000047443 | 42,29901409 | 67,0171278 | 48,1105825 | 10,5992417 | 47,0718625 | 17,25059984 | 28,397329 | 19,5556163 | 7,4743178 | 45,3052534 | 128,077144 | 77,07477 | 53,0930926 | 28,06435 | 39,82475 | 3,646183163 | 7,445132 | 2,972554466 | 8,555098 | 17,33306 | 23,91466 | 31,4076867 |
| Downregulated | ENSMUSG00000071537 | 109,9774366 | 85,1011147 | 133,438785 | 16,3806462 | 89,0548749 | 46,44392265 | 40,8921537 | 62,3335271 | 30,9650309 | 90,6105068 | 185,491036 | 84,95742 | 80,1405172 | 65,48349 | 36,63877 | 27,34637372 | 6,617895 | 21,79873275 | 34,22039 | 1,999969 | 24,80038 | 36,7918616 |
| Downregulated | ENSMUSG00000113728 | 8,459802819 | 8,51011147 | 17,24719 | 3,8542697 | 7,63327499 | 0 | 2,27178632 | 1,22222602 | 8,54207749 | 7,55087556 | 32,019286 | 14,88944 | 12,0210776 | 7,483827 | 1,59299 | 5,469274744 | 2,481711 | 0,990851489 | 0 | 1,999969 | 0,885728 | 0 |
| Downregulated | ENSMUSG00000037035 | 793,1065142 | 584,0064 | 592,758686 | 171,515001 | 194,648512 | 179,1408445 | 191,965944 | 119,77815 | 227,432813 | 593,28308 | 881,08242 | 429,1663 | 356,625301 | 610,8674 | 451,6126 | 326,333393 | 202,673 | 456,7825363 | 269,9609 | 119,9981 | 101,8587 | 166,012058 |
| Downregulated | ENSMUSG00000042622 | 242,1618557 | 323,384236 | 417,563546 | 225,474777 | 262,075775 | 416,6683346 | 264,663106 | 262,778595 | 311,785828 | 499,436484 | 852,375474 | 504,4894 | 343,602467 | 559,4161 | 271,6048 | 297,1639277 | 154,6933 | 178,353268 | 153,0412 | 108,665 | 187,7743 | 122,938659 |
| Downregulated | ENSMUSG00000032870 | 247,4492324 | 140,416839 | 299,556457 | 151,280086 | 203,554 | 144,6396448 | 90,8714527 | 86,7780475 | 113,182527 | 288,011968 | 329,025766 | 253,1206 | 189,331972 | 248,8373 | 135,4041 | 197,8054366 | 64,52447 | 105,0302578 | 105,5129 | 83,99869 | 50,4865 | 82,5573479 |
| Downregulated | ENSMUSG00000020038 | 1163,222888 | 874,413954 | 1220,01175 | 465,403066 | 615,750849 | 335,7232123 | 344,175627 | 194,333937 | 491,169455 | 941,702053 | 800,482149 | 1509,965 | 1118,96197 | 732,4796 | 422,1423 | 583,389306 | 71,14237 | 369,5876053 | 259,5047 | 103,9984 | 153,2309 | 183,959308 |
| Downregulated | ENSMUSG00000026820 | 518,1629226 | 441,462033 | 396,685369 | 157,06149 | 180,654175 | 87,57996843 | 63,6100169 | 44,0001368 | 61,9300618 | 506,987359 | 303,63116 | 502,7377 | 455,799191 | 199,2569 | 87,61444 | 144,0242349 | 42,18908 | 59,45108933 | 116,9197 | 53,3325 | 59,34378 | 41,278674 |
| Downregulated | ENSMUSG00000054716 | 216,7824472 | 197,860092 | 214,228254 | 55,8869106 | 96,6881499 | 70,32936859 | 36,3485811 | 15,8889383 | 28,8295115 | 200,637551 | 152,367637 | 247,8655 | 255,447898 | 94,48332 | 45,40021 | 70,18902588 | 21,50816 | 34,67980211 | 48,47889 | 21,99966 | 17,71456 | 26,0235118 |
| Downregulated | ENSMUSG00000041846 | 835,4055283 | 992,49175 | 866,898232 | 310,268711 | 412,19685 | 352,9738122 | 218,091486 | 246,889656 | 427,103874 | 1019,3682 | 1148,27784 | 1490,696 | 766,343695 | 571,5773 | 340,1033 | 334,5373052 | 167,1018 | 317,0724764 | 268,0598 | 158,6642 | 213,4605 | 250,364131 |
| Downregulated | ENSMUSG00000035529 | 775,1294332 | 696,765377 | 648,131244 | 190,78635 | 203,554 | 193,7375059 | 121,540568 | 59,8890751 | 163,367232 | 834,911098 | 803,794489 | 1173,639 | 427,75001 | 275,0306 | 150,5375 | 124,8817733 | 59,56105 | 133,764951 | 115,9691 | 77,33213 | 90,34426 | 97,81251 |
| Downregulated | ENSMUSG00000031671 | 306,6678522 | 325,511764 | 303,187445 | 62,6318826 | 92,8715124 | 80,94512234 | 37,4844742 | 47,6668148 | 35,2360696 | 338,710704 | 378,710865 | 725,2035 | 284,498836 | 163,7087 | 83,63197 | 95,71230802 | 50,46145 | 60,44194082 | 79,84759 | 30,66619 | 38,0863 | 38,5865865 |
| Downregulated | ENSMUSG00000030374 | 2289,434138 | 1974,34586 | 1634,85206 | 686,060006 | 985,964686 | 642,2531018 | 574,761938 | 403,334587 | 581,929029 | 2216,72133 | 2335,19965 | 3229,258 | 1918,36363 | 1200,219 | 625,2485 | 752,9368231 | 229,9718 | 573,703012 | 581,7467 | 333,3281 | 290,5188 | 421,760364 |
| Downregulated | ENSMUSG00000039148 | 923,1759826 | 778,6752 | 639,961522 | 307,378008 | 358,763925 | 244,1623362 | 209,004341 | 95,3336297 | 225,297294 | 964,354679 | 826,980868 | 1525,73 | 838,470161 | 585,6095 | 265,2328 | 369,1760452 | 147,2482 | 215,0147731 | 247,1473 | 123,9981 | 114,2589 | 188,44612 |
| Downregulated | ENSMUSG00000024843 | 1238,303638 | 1039,29736 | 1743,78168 | 519,362842 | 815,488212 | 930,2054223 | 639,507848 | 383,778971 | 560,573835 | 1470,26334 | 1723,52088 | 2315,747 | 1203,10951 | 859,7046 | 583,8308 | 491,3231812 | 140,6303 | 362,6516449 | 224,3337 | 177,3306 | 198,4031 | 233,314244 |
| Downregulated | ENSMUSG00000017781 | 2587,642187 | 2686,00393 | 2782,24406 | 1030,05358 | 1769,64759 | 1154,46322 | 951,878467 | 651,44647 | 1157,4515 | 3455,06492 | 3772,75518 | 5138,61 | 3563,24774 | 2416,341 | 1275,985 | 1370,964869 | 431,8176 | 894,7388944 | 925,8518 | 397,3271 | 596,9807 | 838,136554 |
| Downregulated | ENSMUSG00000024970 | 376,4612254 | 305,300249 | 419,37904 | 134,899439 | 241,720375 | 161,8902447 | 134,035393 | 79,4446914 | 136,67324 | 346,261579 | 527,766162 | 776,8787 | 423,742985 | 274,0952 | 196,7343 | 197,8054366 | 74,45132 | 108,0028123 | 131,1782 | 61,33238 | 63,77242 | 91,5309727 |
| Downregulated | ENSMUSG00000027519 | 1652,833976 | 1661,59926 | 1759,21338 | 702,440652 | 1248,04046 | 825,374854 | 537,277464 | 515,779381 | 903,324694 | 1786,32142 | 2064,69189 | 2425,228 | 1620,84196 | 1379,831 | 939,0675 | 995,4080034 | 355,7118 | 755,0288345 | 942,0114 | 265,3292 | 446,4069 | 550,083199 |
| Downregulated | ENSMUSG00000056216 | 877,7045424 | 797,82295 | 997,613777 | 344,957138 | 764,599712 | 577,2316101 | 402,106178 | 360,556676 | 436,713711 | 867,271993 | 1257,58506 | 1449,531 | 1058,85658 | 811,9952 | 416,5669 | 623,4973208 | 282,0878 | 404,2674074 | 455,3214 | 181,9972 | 218,7748 | 340,100379 |
| Downregulated | ENSMUSG00000061979 | 259,0814613 | 327,639292 | 302,279698 | 175,369271 | 132,3101 | 134,0238911 | 82,9202006 | 56,222397 | 111,047007 | 281,539789 | 341,171012 | 571,054 | 450,790409 | 310,5788 | 136,9971 | 194,1592534 | 63,69724 | 86,20407953 | 76,04532 | 60,66572 | 45,17213 | 85,2494354 |
| Downregulated | ENSMUSG00000024527 | 1029,980993 | 876,541482 | 843,296814 | 347,84784 | 404,563575 | 315,818674 | 206,732555 | 169,889417 | 259,465604 | 899,632889 | 882,186534 | 2273,706 | 1480,59605 | 754,9311 | 398,2475 | 379,2030489 | 167,9291 | 303,2005556 | 359,3141 | 211,9967 | 160,3168 | 203,701282 |
| Downregulated | ENSMUSG00000027637 | 345,7944402 | 375,508669 | 367,63747 | 61,6683151 | 108,138062 | 39,80907656 | 28,397329 | 15,8889383 | 34,1683099 | 282,618485 | 417,354831 | 509,7445 | 553,971325 | 159,9668 | 54,16166 | 72,01211746 | 4,963421 | 37,65235657 | 32,31926 | 7,333219 | 6,200096 | 16,1525246 |
| Downregulated | ENSMUSG00000006442 | 1853,754293 | 1826,48267 | 1767,3831 | 937,551104 | 802,766087 | 663,4846093 | 563,403006 | 567,112874 | 769,854733 | 2445,40499 | 2626,68556 | 2924,462 | 2785,88473 | 1089,832 | 462,7636 | 616,2049545 | 453,3258 | 589,5566358 | 590,3018 | 477,3259 | 680,2391 | 513,291337 |
| Downregulated | ENSMUSG00000024091 | 2847,781124 | 3618,9249 | 3062,73784 | 1081,12265 | 1166,61886 | 717,8903473 | 581,577297 | 600,112977 | 910,799012 | 3338,5657 | 4003,51486 | 5444,282 | 3681,45501 | 1785,828 | 969,3344 | 894,2264206 | 450,8441 | 808,5348149 | 607,412 | 448,6597 | 491,579 | 477,396838 |
| Downregulated | ENSMUSG00000059422 | 131,1269437 | 209,561495 | 134,346532 | 55,8869106 | 64,8828374 | 25,21241515 | 31,8050084 | 42,7779108 | 41,6426277 | 194,165372 | 257,258401 | 303,9198 | 225,395205 | 107,58 | 48,58619 | 73,83520904 | 30,60776 | 65,39619826 | 33,26983 | 36,66609 | 23,02893 | 23,3314244 |
| Downregulated | ENSMUSG00000021102 | 425,1050916 | 432,951921 | 285,032508 | 87,6846356 | 113,226912 | 95,54178374 | 43,16394 | 46,4445888 | 71,5398989 | 389,40944 | 753,005277 | 723,4518 | 501,879989 | 173,999 | 126,6427 | 95,71230802 | 68,66066 | 76,29556464 | 105,5129 | 73,99885 | 74,40115 | 50,2522987 |
| Downregulated | ENSMUSG00000105081 | 143,8166479 | 136,161784 | 128,900051 | 57,8140454 | 82,6938124 | 80,94512234 | 46,5716195 | 34,2223286 | 62,9978215 | 154,253601 | 266,091307 | 267,1342 | 225,395205 | 126,2896 | 43,80722 | 51,95811007 | 16,54474 | 47,56087146 | 38,97323 | 24,66628 | 38,97203 | 50,2522987 |
| Downregulated | ENSMUSG00000028850 | 174,4834331 | 238,283121 | 154,316963 | 48,1783712 | 80,1493874 | 79,61815312 | 86,32788 | 12,2222602 | 76,8786974 | 144,545332 | 156,78409 | 319,6851 | 149,261713 | 80,45114 | 83,63197 | 192,3361618 | 16,54474 | 60,44194082 | 69,39135 | 19,33303 | 16,82883 | 43,0733989 |
| Downregulated | ENSMUSG00000016239 | 661,9795706 | 515,925508 | 463,858635 | 289,070227 | 380,391537 | 321,1265509 | 343,039734 | 151,556027 | 201,806581 | 357,048545 | 443,85355 | 740,093 | 306,537478 | 387,2881 | 289,1277 | 277,1099204 | 87,68711 | 254,6488326 | 169,2008 | 55,33247 | 51,37222 | 108,58086 |
| Downregulated | ENSMUSG00000021846 | 501,243317 | 444,653324 | 339,497318 | 136,826574 | 174,293112 | 242,835367 | 149,937897 | 167,444965 | 181,519147 | 344,104186 | 384,231431 | 388,0014 | 226,396961 | 224,5148 | 185,5833 | 196,8938908 | 67,00618 | 178,353268 | 116,9197 | 55,99913 | 103,6302 | 104,99141 |
| Downregulated | ENSMUSG00000033697 | 277,0585423 | 208,497731 | 170,656406 | 103,101714 | 100,504787 | 123,4081373 | 77,2407348 | 45,2223628 | 66,2011005 | 182,29971 | 117,036011 | 215,459 | 121,212532 | 105,7091 | 106,7303 | 63,80820534 | 23,98987 | 47,56087146 | 77,94645 | 45,99928 | 27,45757 | 34,0997741 |
| Downregulated | ENSMUSG00000081884 | 9,517278171 | 11,7014033 | 23,6014178 | 10,5992417 | 15,26655 | 11,94272297 | 1,13589316 | 11,0000342 | 3,20327906 | 44,2265569 | 32,019286 | 39,41324 | 27,0474245 | 7,483827 | 8,761444 | 16,40782423 | 12,40855 | 30,71639615 | 6,653965 | 3,999938 | 4,42864 | 0,89736248 |
| Downregulated | ENSMUSG00000029330 | 54,98871832 | 61,6983082 | 46,2950888 | 29,8705901 | 47,0718625 | 91,56087608 | 45,4357263 | 42,7779108 | 40,5748681 | 58,2496115 | 100,474311 | 146,2669 | 34,0597198 | 68,28992 | 55,75465 | 58,3389306 | 25,64434 | 45,57916849 | 14,2585 | 7,999875 | 17,71456 | 26,9208743 |
| Downregulated | ENSMUSG00000041570 | 730,7154685 | 855,266203 | 708,042535 | 404,698318 | 722,616699 | 828,0287924 | 624,741237 | 640,446435 | 1045,33673 | 1141,26091 | 1521,46814 | 1687,762 | 770,350721 | 910,2205 | 491,4374 | 515,9349175 | 281,2605 | 548,9317248 | 418,2493 | 247,9961 | 343,6625 | 376,892241 |
| Downregulated | ENSMUSG00000054843 | 1249,935866 | 1364,80913 | 826,957371 | 889,372732 | 815,488212 | 1184,983512 | 813,299501 | 1037,66989 | 1048,54001 | 2168,17998 | 2350,65723 | 2688,859 | 795,394633 | 1205,832 | 735,9613 | 690,0401635 | 506,2689 | 663,8704975 | 442,964 | 312,6618 | 501,3221 | 554,570011 |
| Downregulated | ENSMUSG00000046791 | 44,4139648 | 56,3794885 | 59,9112914 | 8,67210682 | 24,1720375 | 19,90453828 | 18,1742905 | 17,1111643 | 22,4229534 | 160,72578 | 239,592588 | 336,3263 | 94,1651077 | 32,74174 | 27,87732 | 24,61173635 | 5,790658 | 37,65235657 | 4,752832 | 3,333281 | 32,77194 | 8,97362477 |
| Downregulated | ENSMUSG00000033016 | 4396,982515 | 6298,54625 | 4758,40893 | 1262,27333 | 2971,8884 | 1766,19603 | 2037,79233 | 1400,67102 | 1980,69422 | 9885,17481 | 13337,6888 | 13079,94 | 4987,74544 | 3598,785 | 1976,9 | 2230,55255 | 1086,162 | 1407,999966 | 1245,242 | 755,9882 | 1069,959 | 1638,58388 |
| Downregulated | ENSMUSG00000039481 | 11,63222888 | 10,6376393 | 7,26197472 | 2,89070227 | 11,4499125 | 6,634846093 | 3,40767947 | 4,88890409 | 4,27103874 | 28,0461092 | 28,706946 | 35,03399 | 12,0210776 | 3,741914 | 2,389485 | 2,734637372 | 2,481711 | 0 | 3,802266 | 0 | 0 | 1,79472495 |
| Downregulated | ENSMUSG00000058152 | 149,1040247 | 149,990715 | 201,519798 | 58,7776129 | 115,771337 | 63,69452249 | 64,74591 | 70,8891093 | 65,1333408 | 277,225003 | 312,464066 | 468,5796 | 300,526939 | 188,9666 | 105,1373 | 111,2085865 | 34,74395 | 66,38704975 | 87,45212 | 57,33244 | 43,40067 | 38,5865865 |
| Downregulated | ENSMUSG00000035671 | 1532,281786 | 1630,75011 | 2104,15717 | 588,739696 | 1594,08226 | 1041,670837 | 836,017364 | 668,557634 | 1318,68321 | 2552,19594 | 3480,16515 | 4610,473 | 2267,97664 | 1922,408 | 901,6323 | 1349,999316 | 483,9336 | 891,7663399 | 1234,786 | 451,9929 | 820,1841 | 824,676117 |
| Downregulated | ENSMUSG00000039244 | 304,5529015 | 280,833679 | 191,534583 | 88,648203 | 150,121075 | 145,966614 | 47,7075126 | 66,0002052 | 69,4043796 | 349,497669 | 512,308575 | 592,0744 | 250,439116 | 277,8371 | 146,5551 | 192,3361618 | 47,97974 | 80,25897059 | 78,89702 | 38,66606 | 68,20106 | 95,1204226 |
| Downregulated | ENSMUSG00000032024 | 3631,37036 | 3702,96225 | 3168,03647 | 1263,23689 | 1711,12581 | 2050,167443 | 1628,87079 | 1144,00356 | 1829,07234 | 4694,48721 | 6929,41513 | 8179,56 | 3833,72199 | 3034,692 | 1562,723 | 1630,755419 | 1173,022 | 1374,311015 | 1252,847 | 666,6563 | 885,728 | 1129,77936 |
| Downregulated | ENSMUSG00000060708 | 175,5409085 | 207,433967 | 125,269064 | 36,6155621 | 24,1720375 | 33,17423047 | 11,3589316 | 15,8889383 | 24,5584728 | 249,178894 | 333,442219 | 396,7599 | 218,382909 | 105,7091 | 24,69134 | 68,3659343 | 25,64434 | 12,88106935 | 30,41813 | 19,99969 | 18,60029 | 17,0498871 |
| Downregulated | ENSMUSG00000085281 | 151,2189754 | 163,819646 | 129,807798 | 52,9962083 | 91,5992999 | 83,59906077 | 71,561269 | 50,1112669 | 52,3202246 | 175,827531 | 250,633721 | 260,1274 | 160,281034 | 101,9671 | 63,7196 | 79,30448379 | 59,56105 | 35,6706536 | 42,77549 | 37,99941 | 45,17213 | 47,5602113 |
| Downregulated | ENSMUSG00000035891 | 615,450655 | 809,524354 | 466,581876 | 331,467194 | 510,157212 | 435,2459037 | 411,193323 | 254,223013 | 601,148703 | 655,847478 | 1344,81001 | 1850,67 | 559,981864 | 702,5443 | 441,2582 | 410,1956058 | 184,4738 | 342,8346151 | 229,0865 | 109,9983 | 231,175 | 240,493144 |
| Downregulated | ENSMUSG00000020482 | 209,3801198 | 257,430872 | 121,638077 | 102,138147 | 92,8715124 | 69,00239937 | 74,9689484 | 124,667054 | 133,469961 | 210,345819 | 501,267442 | 668,2733 | 199,349536 | 194,5795 | 96,37589 | 144,9357807 | 100,9229 | 102,0577033 | 104,5623 | 88,66528 | 61,11523 | 98,7098725 |
| Downregulated | ENSMUSG00000040606 | 768,7845811 | 769,101324 | 565,526281 | 239,928289 | 526,695974 | 585,1934254 | 413,46511 | 378,890067 | 422,832836 | 1147,73309 | 2072,42068 | 2801,843 | 895,570279 | 712,8345 | 427,7178 | 523,2272838 | 363,157 | 464,7093483 | 332,6983 | 91,33191 | 176,2599 | 321,255767 |
| Downregulated | ENSMUSG00000021681 | 302,4379508 | 327,639292 | 254,169115 | 145,498681 | 183,1986 | 220,2768903 | 90,8714527 | 169,889417 | 136,67324 | 407,74728 | 804,898602 | 1291,002 | 271,476002 | 331,1593 | 183,1938 | 234,2672682 | 99,26842 | 133,764951 | 77,94645 | 114,6649 | 127,5448 | 141,783271 |
| Downregulated | ENSMUSG00000040396 | 319,3575564 | 317,001652 | 199,704305 | 52,9962083 | 44,5274375 | 50,42483031 | 77,2407348 | 81,8891435 | 60,8623021 | 347,340276 | 508,996235 | 805,7817 | 325,570851 | 188,9666 | 123,4567 | 115,7663154 | 47,1525 | 52,51512891 | 49,42946 | 49,99922 | 84,14416 | 78,0705355 |
| Downregulated | ENSMUSG00000040028 | 870,302215 | 913,773219 | 688,979852 | 343,030003 | 265,892412 | 279,9905051 | 124,948247 | 248,111882 | 356,631735 | 982,69252 | 1842,76511 | 2753,671 | 834,463135 | 552,8677 | 314,6155 | 347,2989462 | 231,6263 | 282,3926743 | 192,965 | 163,3308 | 267,4899 | 262,029843 |
| Downregulated | ENSMUSG00000020358 | 1481,522969 | 1757,33802 | 1185,51737 | 251,491098 | 333,319675 | 287,9523204 | 157,889149 | 176,000547 | 296,837193 | 1774,45576 | 2931,42083 | 4896 | 1636,87006 | 882,1561 | 333,7314 | 345,4758547 | 231,6263 | 303,2005556 | 202,4707 | 221,3299 | 277,2329 | 174,985683 |
| Downregulated | ENSMUSG00000005312 | 1483,637919 | 1557,3504 | 1003,06026 | 519,362842 | 353,675075 | 347,6659353 | 272,614358 | 309,223184 | 379,054688 | 1605,10041 | 2416,90403 | 3974,606 | 1065,86888 | 896,1883 | 437,2757 | 463,0652616 | 325,9313 | 371,5693083 | 312,7364 | 365,9943 | 448,1784 | 252,158856 |
| Downregulated | ENSMUSG00000054321 | 998,2567326 | 1400,9771 | 1206,39555 | 336,285031 | 431,280037 | 674,1003631 | 486,162272 | 613,557463 | 564,844874 | 2170,33738 | 2426,84105 | 3709,223 | 851,492995 | 901,8012 | 365,5912 | 461,2421701 | 206,8092 | 372,5601598 | 185,3605 | 169,3307 | 341,891 | 282,66918 |
| Downregulated | ENSMUSG00000056708 | 4671,926107 | 4653,96721 | 3467,59293 | 1514,72799 | 1245,49604 | 2497,356069 | 1064,33189 | 1514,33804 | 1566,40346 | 4696,6446 | 8086,52587 | 11598 | 2895,07618 | 3931,816 | 1419,354 | 1092,031857 | 769,3303 | 1070,119608 | 757,6015 | 1098,65 | 1395,907 | 746,605581 |
| Downregulated | ENSMUSG00000070407 | 473,7489578 | 317,001652 | 184,272609 | 137,790142 | 264,6202 | 226,9117364 | 26,1255426 | 116,111472 | 248,788007 | 440,108176 | 801,586262 | 1084,302 | 139,244149 | 149,6765 | 250,0994 | 338,1834883 | 71,96961 | 72,33215868 | 21,86303 | 100,6651 | 147,9166 | 77,1731731 |
| Downregulated | ENSMUSG00000002083 | 227,3572007 | 109,567685 | 246,90714 | 91,5389053 | 95,4159374 | 163,2172139 | 129,49182 | 67,2224312 | 93,9628523 | 258,887162 | 406,313698 | 528,1374 | 115,201993 | 167,4506 | 138,5901 | 136,7318686 | 83,55092 | 79,2681191 | 89,35325 | 28,66622 | 19,48602 | 67,3021858 |
| Downregulated | ENSMUSG00000042608 | 3029,666884 | 2771,10505 | 2650,62077 | 1018,49077 | 1928,67415 | 1812,639953 | 1831,05977 | 1491,11575 | 2008,45597 | 3708,5586 | 5840,7594 | 6754,553 | 2864,02173 | 1797,054 | 846,6741 | 1813,064578 | 665,0984 | 1472,405312 | 962,9239 | 405,327 | 751,9831 | 1446,54831 |
| Downregulated | ENSMUSG00000031728 | 123,7246162 | 119,141561 | 111,652861 | 67,4497197 | 85,2382374 | 70,32936859 | 57,9305511 | 40,3334587 | 101,43717 | 115,420526 | 269,403647 | 301,2923 | 87,1528124 | 72,96731 | 42,21423 | 33,72719425 | 16,54474 | 43,59746551 | 35,17096 | 25,33294 | 42,51494 | 61,0206485 |
| Downregulated | ENSMUSG00000021794 | 3132,241994 | 3691,26085 | 2770,44336 | 1446,3147 | 1619,52651 | 2433,661547 | 1403,96394 | 1449,56006 | 2320,2418 | 4426,97047 | 6201,80446 | 6476,908 | 1389,43622 | 2414,47 | 1353,245 | 1440,242349 | 838,8182 | 1459,524243 | 977,1824 | 719,3221 | 1320,62 | 1217,72088 |
| Downregulated | ENSMUSG00000108799 | 188,2306127 | 218,071606 | 141,608507 | 72,2675568 | 75,0605374 | 106,1575375 | 62,4741237 | 67,2224312 | 106,775969 | 235,155839 | 336,754559 | 423,9113 | 73,1282219 | 114,1284 | 54,16166 | 57,42738481 | 46,32526 | 71,34130719 | 60,83626 | 43,33266 | 61,11523 | 56,5338361 |
| Downregulated | ENSMUSG00000048429 | 397,6107325 | 343,595751 | 162,486684 | 137,790142 | 100,504787 | 84,92602999 | 94,2791321 | 72,1113353 | 128,131162 | 465,996892 | 778,399883 | 915,2629 | 216,379396 | 267,5468 | 121,0672 | 96,62385381 | 65,35171 | 103,0485548 | 89,35325 | 71,99888 | 85,91562 | 86,1467978 |
| Downregulated | ENSMUSG00000031065 | 2036,697529 | 1778,6133 | 1637,5753 | 490,455819 | 463,08535 | 597,1361484 | 440,726545 | 502,334895 | 483,695138 | 2280,36442 | 4009,03542 | 4055,184 | 1254,19909 | 806,3824 | 469,1355 | 439,3650711 | 351,5757 | 514,2519227 | 405,8919 | 550,6581 | 604,0665 | 430,733989 |
| Downregulated | ENSMUSG00000025165 | 34330,93731 | 25298,4339 | 27718,9575 | 27408,6754 | 7756,6796 | 6865,738737 | 13239,9707 | 16570,9404 | 8939,28409 | 15485,7671 | 13476,807 | 10263,21 | 7183,59561 | 28697,67 | 18113,89 | 17292,02365 | 14348,42 | 17761,01294 | 14578,84 | 2385,296 | 1292,277 | 4214,01419 |
| Downregulated | ENSMUSG00000067199 | 9,517278171 | 15,956459 | 28,140152 | 22,1620508 | 15,26655 | 13,26969219 | 9,08714527 | 15,8889383 | 6,40655811 | 33,4395918 | 11,0411331 | 14,88944 | 5,00878232 | 27,12887 | 19,11588 | 20,96555318 | 16,54474 | 7,92681191 | 11,4068 | 2,666625 | 0,885728 | 3,58944991 |
| Downregulated | ENSMUSG00000049892 | 527,6802008 | 518,053036 | 583,681218 | 501,05506 | 250,625862 | 437,8998421 | 763,320202 | 789,55801 | 880,901741 | 257,808466 | 289,277687 | 331,947 | 526,9239 | 723,1248 | 1138,988 | 342,7412173 | 342,4761 | 352,74313 | 294,6756 | 179,3305 | 62,00096 | 172,293596 |
| Downregulated | ENSMUSG00000042745 | 127,9545176 | 157,437062 | 147,054988 | 256,308935 | 180,654175 | 253,4511208 | 193,101837 | 251,77856 | 219,958495 | 141,309243 | 196,532169 | 189,1835 | 64,1124137 | 207,6762 | 305,0576 | 370,9991368 | 119,9493 | 227,8958424 | 208,1741 | 47,99925 | 44,2864 | 63,7127359 |
| Downregulated | ENSMUSG00000048251 | 503,3582677 | 319,12918 | 379,438179 | 615,719584 | 442,72995 | 380,8401657 | 480,482806 | 471,779244 | 484,762897 | 377,543778 | 513,412689 | 263,6308 | 190,333728 | 389,159 | 309,04 | 421,1341553 | 165,4474 | 422,1027342 | 239,5428 | 39,99938 | 82,37271 | 208,188095 |
| Downregulated | ENSMUSG00000029406 | 75,08075001 | 73,3997114 | 52,6493167 | 84,7939333 | 55,9773499 | 102,1766298 | 40,8921537 | 70,8891093 | 68,3366199 | 69,0365766 | 67,3509118 | 34,15814 | 22,0386422 | 60,8061 | 69,29506 | 67,45438851 | 26,47158 | 38,64320806 | 37,07209 | 14,66644 | 23,02893 | 32,3050492 |
| Downregulated | ENSMUSG00000069378 | 25,37940846 | 13,8289311 | 71,7120004 | 22,1620508 | 11,4499125 | 41,13604578 | 15,9025042 | 33,0001026 | 17,084155 | 21,5739302 | 67,3509118 | 21,89624 | 17,0298599 | 5,61287 | 10,35443 | 10,93854949 | 0,827237 | 1,981702978 | 0,950566 | 6,666563 | 0,885728 | 4,48681239 |
| Downregulated | ENSMUSG00000067860 | 38,06911268 | 27,6578623 | 67,1732662 | 25,052753 | 35,62195 | 25,21241515 | 32,9409016 | 51,3334929 | 32,0327906 | 38,8330743 | 49,6850989 | 24,52379 | 31,0544504 | 40,22557 | 28,67382 | 20,96555318 | 22,33539 | 18,82617829 | 15,20906 | 7,333219 | 4,42864 | 7,17889982 |
| Downregulated | ENSMUSG00000057722 | 786,7616621 | 687,191501 | 2185,85439 | 792,052423 | 741,699887 | 621,0215943 | 1408,50752 | 2024,00629 | 1698,80566 | 1404,46285 | 1095,2804 | 1451,283 | 292,512888 | 364,8366 | 491,4374 | 875,9955048 | 1209,42 | 1598,243451 | 987,6386 | 163,3308 | 386,1774 | 709,81372 |
| Downregulated | ENSMUSG00000021187 | 0 | 1,06376393 | 17,24719 | 17,3442136 | 7,63327499 | 7,961815312 | 13,6307179 | 37,8890067 | 6,40655811 | 15,1017511 | 22,0822662 | 35,90984 | 0 | 0 | 0 | 2,734637372 | 0 | 20,80788127 | 0 | 1,333313 | 1,771456 | 0,89736248 |

| | | | | | | | | | | | | | | | | | | | | | | | |
|---|---|---|---|---|---|---|---|---|---|---|---|---|---|---|---|---|---|---|---|---|---|---|---|
| Downregulated | ENSMUSG00000047777 | 1248,878391 | 796,759187 | 1200,04132 | 807,469501 | 1274,75692 | 1017,785391 | 1031,39099 | 508,446025 | 1162,7903 | 1157,44135 | 1302,8537 | 1008,979 | 915,605409 | 1091,703 | 512,9427 | 659,9591524 | 191,0917 | 651,9802796 | 593,1535 | 175,9973 | 191,3173 | 605,719672 |
| Downregulated | ENSMUSG00000040616 | 1182,257444 | 699,956669 | 1024,84618 | 759,29113 | 1203,51302 | 1413,222218 | 1142,70852 | 754,113455 | 842,462392 | 1143,4183 | 1422,09794 | 1131,598 | 1193,09195 | 1196,477 | 906,4113 | 1017,285102 | 408,655 | 763,9464979 | 557,032 | 107,9983 | 264,8327 | 699,942732 |
| Downregulated | ENSMUSG00000032352 | 39,12658804 | 31,912918 | 40,8486078 | 26,9798879 | 31,8053125 | 42,463015 | 21,58197 | 8,55558215 | 25,6262325 | 40,9904673 | 41,9563057 | 54,30268 | 31,0544504 | 19,64505 | 19,91237 | 26,43482793 | 5,790658 | 6,935960422 | 2,851699 | 13,33313 | 4,42864 | 14,3577996 |
| Downregulated | ENSMUSG00000036561 | 172,3684824 | 238,283121 | 216,951495 | 118,518793 | 189,559662 | 210,9881058 | 101,094491 | 73,3335613 | 90,7595733 | 215,739302 | 259,466628 | 217,2107 | 177,310894 | 155,2894 | 82,83547 | 73,83520904 | 59,56105 | 106,0211093 | 77,94645 | 47,33259 | 46,94358 | 60,123286 |
| Downregulated | ENSMUSG00000119926 | 22,2069824 | 22,3390426 | 28,140152 | 19,2713485 | 29,2608875 | 14,5966614 | 15,9025042 | 25,6667465 | 18,1519147 | 18,3378407 | 18,7699262 | 33,28229 | 18,0316164 | 16,83861 | 16,72639 | 12,76164107 | 6,617895 | 20,80788127 | 14,2585 | 3,999938 | 5,314368 | 2,69208743 |
| Downregulated | ENSMUSG00000022895 | 14685,16022 | 15391,6004 | 13236,7644 | 10035,5547 | 18388,5595 | 15423,36323 | 15482,2237 | 13108,3741 | 18949,5311 | 16672,3332 | 16638,9876 | 18447,15 | 12281,5343 | 13255,73 | 15244,12 | 17333,04321 | 9431,327 | 11515,676 | 11568,39 | 2886,622 | 3881,26 | 10339,4105 |
| Downregulated | ENSMUSG00000035954 | 845,9802819 | 1233,96616 | 913,193321 | 614,756017 | 1344,72861 | 993,8999447 | 850,783976 | 998,55866 | 1187,34877 | 1097,03435 | 1139,44493 | 1135,101 | 636,115355 | 645,4801 | 580,6448 | 834,0643984 | 547,6308 | 770,8824583 | 759,5026 | 345,9946 | 218,7748 | 585,080335 |
| Downregulated | ENSMUSG00000034839 | 1722,627349 | 1864,77818 | 1466,91889 | 1139,90026 | 2086,4285 | 2076,706827 | 1669,76294 | 1274,78174 | 2037,28548 | 1904,97804 | 2469,90147 | 1595,798 | 1713,00355 | 1221,735 | 1000,398 | 1520,458379 | 823,9279 | 1183,076678 | 1069,387 | 220,6632 | 496,8934 | 977,227738 |
| Downregulated | ENSMUSG00000028525 | 6290,920871 | 8801,58279 | 5922,14038 | 6376,88921 | 9326,58983 | 11431,83982 | 6723,3516 | 6828,57678 | 7556,5353 | 6762,34842 | 5396,90585 | 6777,325 | 2908,09902 | 6006,707 | 5093,585 | 6080,010424 | 5601,221 | 6484,132143 | 3577,932 | 2308,631 | 1495,109 | 2545,81735 |
| Downregulated | ENSMUSG00000115223 | 35,95416198 | 35,1042098 | 24,5091647 | 31,797725 | 43,255225 | 34,50119968 | 29,5332221 | 22,0000684 | 33,1005503 | 33,4395918 | 24,2904928 | 35,90984 | 18,0316164 | 22,45148 | 23,89485 | 26,43482793 | 16,54474 | 16,84447531 | 13,30793 | 11,33316 | 7,085824 | 9,87098725 |
| Downregulated | ENSMUSG00000048120 | 2339,135479 | 4452,91583 | 3641,88032 | 4685,82838 | 3882,79255 | 3115,723725 | 4636,71587 | 4541,7919 | 6116,12748 | 6590,83567 | 7177,84062 | 3717,982 | 1587,784 | 3189,981 | 6250,892 | 3455,670092 | 2142,543 | 2381,016128 | 1634,024 | 128,6647 | 861,8134 | 2017,27085 |
| Downregulated | ENSMUSG00000043953 | 1031,038469 | 1137,16365 | 897,761625 | 1038,72568 | 1029,21991 | 846,6063615 | 1175,64942 | 812,780304 | 822,174958 | 1284,72754 | 1778,72654 | 1031,751 | 938,645807 | 849,4144 | 1373,157 | 1151,282334 | 409,4822 | 721,3398839 | 429,6561 | 114,6649 | 453,4927 | 622,769559 |
| Downregulated | ENSMUSG00000020893 | 330,9897853 | 225,517954 | 357,652255 | 283,288823 | 309,147637 | 363,5895659 | 478,21102 | 268,889725 | 235,974891 | 308,507202 | 464,831703 | 387,1256 | 204,358319 | 302,1595 | 454,7986 | 350,0335836 | 306,9049 | 347,7888726 | 155,8929 | 101,3318 | 72,6297 | 129,220197 |
| Downregulated | ENSMUSG00000015396 | 74,02327466 | 154,24577 | 217,859242 | 151,280086 | 215,003912 | 209,6611365 | 277,157931 | 166,222739 | 180,451387 | 159,647083 | 320,19286 | 155,0254 | 204,358319 | 136,5798 | 250,0994 | 433,8957963 | 41,36184 | 109,9845153 | 28,51699 | 5,33325 | 36,31485 | 115,75976 |
| Downregulated | ENSMUSG00000030905 | 25,37940846 | 25,5303344 | 48,1105825 | 36,6155621 | 38,166375 | 63,69452249 | 52,2510853 | 56,222397 | 48,0491859 | 30,2035023 | 120,348351 | 67,44043 | 21,0368858 | 29,93531 | 67,70207 | 113,031678 | 14,06303 | 55,48768337 | 23,76416 | 9,999844 | 11,51446 | 7,17889982 |
| Downregulated | ENSMUSG00000027253 | 930,57831 | 1047,80747 | 993,98279 | 870,101384 | 673,000412 | 622,3485635 | 764,456095 | 657,5576 | 654,536687 | 694,680552 | 760,73407 | 1232,321 | 646,13292 | 616,4803 | 980,4853 | 1146,724605 | 474,0067 | 733,2301017 | 826,9928 | 91,99856 | 155,8881 | 395,736853 |
| Downregulated | ENSMUSG00000073565 | 748,6925494 | 769,101324 | 979,45884 | 665,82509 | 496,162874 | 671,4464246 | 414,601003 | 332,445478 | 626,774935 | 691,444462 | 849,063134 | 922,2697 | 568,997672 | 852,2208 | 960,5729 | 824,0373947 | 497,9966 | 545,9591703 | 517,1082 | 100,6651 | 174,4884 | 346,381916 |
| Downregulated | ENSMUSG00000049107 | 85,65550354 | 98,9300459 | 158,855697 | 100,211012 | 96,6881499 | 107,4845067 | 113,589316 | 69,6668832 | 112,114767 | 193,086675 | 114,827784 | 104,2261 | 14,0245905 | 15,90313 | 114,6953 | 95,71230802 | 26,47158 | 45,57916849 | 57,03399 | 3,999938 | 15,9431 | 39,483949 |
| Downregulated | ENSMUSG00000027765 | 155,4488768 | 92,5474623 | 190,626836 | 189,822783 | 194,648512 | 143,3126756 | 85,1919869 | 70,8891093 | 113,182527 | 198,480158 | 146,84707 | 166,4114 | 105,184429 | 79,51566 | 113,1023 | 171,3706086 | 52,11592 | 84,22237655 | 49,42946 | 11,99981 | 42,51494 | 56,5338361 |
| Downregulated | ENSMUSG00000056004 | 50,75881691 | 151,054479 | 98,9444056 | 90,5753379 | 150,121075 | 167,1981215 | 79,5125211 | 149,111575 | 185,790185 | 189,850586 | 147,951183 | 112,1088 | 57,1001185 | 98,22523 | 109,1198 | 164,9897881 | 88,51434 | 143,6734659 | 68,44079 | 37,33275 | 21,25747 | 114,862397 |
| Downregulated | ENSMUSG00000045776 | 31,72426057 | 138,289311 | 55,3725572 | 31,797725 | 123,404612 | 230,892644 | 24,9896495 | 33,0001026 | 72,6076586 | 141,309243 | 75,079705 | 102,4744 | 10,0175646 | 9,354784 | 13,54041 | 143,1126891 | 19,85368 | 117,9113272 | 36,12153 | 0 | 3,542912 | 33,2024117 |
| Downregulated | ENSMUSG00000031778 | 403,9555846 | 631,875777 | 436,62623 | 950,07748 | 940,165037 | 800,1624388 | 845,10451 | 689,335476 | 806,158563 | 1001,03036 | 671,300892 | 341,5814 | 258,453168 | 339,5787 | 460,3741 | 687,3055261 | 595,6105 | 872,9401616 | 516,1576 | 228,6631 | 161,2025 | 406,505202 |
| Downregulated | ENSMUSG00000036528 | 223,1272993 | 269,132275 | 164,302178 | 238,964721 | 561,045712 | 229,5656748 | 190,830051 | 193,111711 | 379,054688 | 308,507202 | 228,551455 | 145,391 | 65,1141702 | 104,7736 | 171,2464 | 247,9404551 | 258,9251 | 282,3926743 | 194,8661 | 30,66619 | 66,4296 | 145,372721 |
| Downregulated | ENSMUSG00000029001 | 32,78173592 | 71,2721836 | 33,5866331 | 53,9597757 | 86,5104499 | 53,07876874 | 74,9689484 | 36,6667807 | 51,2524649 | 59,328308 | 75,079705 | 29,77889 | 27,0474245 | 71,09636 | 45,40021 | 84,77375853 | 28,95329 | 68,36875273 | 30,41813 | 16,66641 | 9,743008 | 23,3314244 |
| Downregulated | ENSMUSG00000039328 | 7602,190308 | 7104,87931 | 6034,70099 | 7315,40388 | 9247,71265 | 10753,75855 | 7904,68049 | 8956,47229 | 8542,07749 | 7774,16574 | 9145,37054 | 6732,657 | 6248,95683 | 7900,115 | 7765,826 | 7755,431587 | 5992,504 | 9739,079284 | 5172,032 | 1294,646 | 1641,254 | 5272,90192 |
| Downregulated | ENSMUSG00000038679 | 1271,085373 | 1537,13888 | 1170,99342 | 1489,67524 | 1625,88757 | 1385,355864 | 1493,6995 | 1431,22667 | 1396,62967 | 2103,45819 | 1036,7624 | 1235,824 | 980,719579 | 1419,121 | 1221,027 | 1590,647405 | 1031,564 | 1629,950699 | 794,6736 | 177,9972 | 254,2039 | 1119,90837 |
| Downregulated | ENSMUSG00000029361 | 9,517278171 | 77,6547672 | 5,44648104 | 22,1620508 | 133,582312 | 34,50119968 | 0 | 73,3335613 | 38,4393487 | 32,3608953 | 18,7699262 | 11,38605 | 36,0632327 | 77,64471 | 38,23176 | 61,98511376 | 40,53461 | 121,8747331 | 88,40268 | 3,333281 | 0,885728 | 11,6657122 |
| Downregulated | ENSMUSG00000039329 | 0 | 3,1912918 | 0,90774684 | 2,89070227 | 21,6276125 | 11,94272297 | 0 | 19,5556163 | 11,7453565 | 0 | 0 | 5,255098 | 4,00702586 | 0 | 3,18598 | 36,46183163 | 0,827237 | 0 | 0 | 0 | 0,885728 | 8,0762623 |
| Downregulated | ENSMUSG00000034171 | 14,80465493 | 13,8289311 | 28,140152 | 52,9962083 | 75,0605374 | 30,52029203 | 19,3101837 | 56,222397 | 71,5398989 | 31,2821988 | 22,0822662 | 33,28229 | 14,0245905 | 18,70957 | 35,84227 | 47,40038111 | 19,85368 | 25,76213871 | 6,653965 | 8,666531 | 11,51446 | 6,28153734 |
| Downregulated | ENSMUSG00000096862 | 3,172426057 | 3,1912918 | 9,0774684 | 8,67210682 | 26,7164625 | 18,57756906 | 11,3589316 | 18,3333903 | 7,4743178 | 30,2035023 | 8,83290647 | 18,39284 | 32,0562069 | 14,03218 | 17,52289 | 8,203912116 | 4,136184 | 12,88106935 | 2,851699 | 3,333281 | 5,314368 | 11,6657122 |
| Downregulated | ENSMUSG00000024176 | 37,01163733 | 51,0606688 | 63,5422788 | 65,5225848 | 133,582312 | 135,3508603 | 115,861102 | 89,2224996 | 91,827333 | 107,869651 | 80,6002715 | 97,21932 | 49,0860668 | 89,80593 | 148,1481 | 60,16202218 | 62,04276 | 42,60661402 | 29,46756 | 27,99956 | 34,54339 | 45,7654863 |
| Downregulated | ENSMUSG00000109438 | 8,459802819 | 7,44634754 | 10,8929621 | 7,70853939 | 43,255225 | 62,36755328 | 10,2230384 | 24,4445204 | 12,8131162 | 5,39348255 | 11,0411331 | 29,77889 | 19,0333728 | 32,74174 | 32,65629 | 15,49627844 | 15,7175 | 23,78043573 | 5,703399 | 7,999875 | 2,657184 | 19,7419745 |
| Downregulated | ENSMUSG00000105161 | 25,37940846 | 9,57387541 | 6,35422788 | 13,4899439 | 43,255225 | 59,71361484 | 20,4460768 | 25,6667465 | 14,9486356 | 18,3378407 | 13,2493597 | 46,42003 | 22,0386422 | 26,19339 | 38,23176 | 30,08101109 | 8,272368 | 18,82617829 | 7,604532 | 21,99966 | 10,62874 | 20,639337 |
| Downregulated | ENSMUSG00000012819 | 17,97708099 | 26,5940983 | 12,7084558 | 22,1620508 | 64,8828374 | 29,19332281 | 49,979299 | 34,2223286 | 44,8459068 | 30,2035023 | 28,706946 | 13,13775 | 9,01580818 | 16,83861 | 21,50536 | 45,57728953 | 2,481711 | 5,945108933 | 14,2585 | 15,99975 | 7,085824 | 42,1760364 |
| Downregulated | ENSMUSG00000073405 | 394,4383064 | 392,528892 | 385,792407 | 418,188262 | 2181,84444 | 1352,181634 | 1347,16929 | 755,335681 | 1563,20018 | 434,714693 | 1121,77912 | 286,4028 | 220,386422 | 233,8696 | 547,9885 | 494,9693643 | 312,6955 | 490,471487 | 194,8661 | 285,9955 | 682,8963 | 262,029843 |
| Downregulated | ENSMUSG00000073406 | 314,0701796 | 238,283121 | 283,217014 | 313,159413 | 1180,6132 | 826,7018232 | 963,237398 | 271,334177 | 511,456889 | 267,516734 | 723,194217 | 366,981 | 181,31792 | 294,6757 | 490,6409 | 544,192837 | 432,6449 | 399,31315 | 150,1895 | 84,66535 | 503,0935 | 146,270084 |
| Downregulated | ENSMUSG00000092457 | 6,344852114 | 1,06376393 | 4,5387342 | 6,74497197 | 58,5217749 | 23,88544594 | 28,397329 | 2,44445204 | 24,5584728 | 4,31478604 | 26,4987194 | 12,2619 | 1,00175646 | 3,741914 | 23,89485 | 18,23091581 | 9,099605 | 10,89936638 | 0 | 0,666656 | 10,62874 | 0,89736248 |
| Downregulated | ENSMUSG00000054034 | 9,517278171 | 18,0839869 | 11,8007089 | 25,052753 | 40,7108 | 26,53938437 | 42,0280469 | 17,1111643 | 22,4229534 | 11,8656616 | 44,1645324 | 2,627549 | 15,026347 | 15,90313 | 18,31938 | 6,380820534 | 8,272368 | 9,908514888 | 10,45623 | 7,333219 | 7,971552 | 17,0498871 |
| Downregulated | ENSMUSG00000028528 | 20,09203169 | 4,25505574 | 9,0774684 | 20,2349159 | 41,9830125 | 19,90453828 | 39,7562605 | 12,2222602 | 19,2196743 | 14,0230546 | 39,7480791 | 2,627549 | 0 | 8,419306 | 26,28433 | 13,67318686 | 0 | 5,945108933 | 9,505665 | 1,999969 | 1,771456 | 9,87098725 |
| Downregulated | ENSMUSG00000051043 | 45,47144015 | 104,248866 | 22,693671 | 41,4333992 | 127,22125 | 96,86875296 | 97,6868116 | 56,222397 | 12,8131162 | 26,9674127 | 8,83290647 | 49,04758 | 18,0316164 | 26,19339 | 23,09835 | 41,01956058 | 48,80697 | 38,64320806 | 22,8136 | 9,333188 | 12,40019 | 88,8388853 |
| Downregulated | ENSMUSG00000108801 | 29,60930986 | 69,1446557 | 41,7563546 | 65,5225848 | 64,8828374 | 70,32936859 | 104,502171 | 45,2223628 | 70,4721393 | 57,170915 | 45,2686457 | 56,93023 | 33,0579633 | 27,12887 | 22,30186 | 49,22347269 | 9,926842 | 34,67980211 | 30,41813 | 16,66641 | 31,00048 | 29,6129618 |
| Downregulated | ENSMUSG00000039313 | 10,57475352 | 29,7853902 | 7,26197472 | 9,63567424 | 31,8053125 | 45,11695343 | 56,7946579 | 23,2222944 | 29,8972712 | 17,2591441 | 6,62467985 | 14,88944 | 0 | 1,870957 | 1,59299 | 0 | 0 | 6,935960422 | 23,76416 | 3,999938 | 0,885728 | 1,79472495 |
| Downregulated | ENSMUSG00000030428 | 51,81629226 | 121,269088 | 59,0035446 | 33,7248598 | 166,659837 | 130,0429834 | 224,906845 | 30,5556505 | 53,3879843 | 69,0365766 | 72,8714784 | 23,64794 | 124,217802 | 78,58018 | 23,09835 | 17,31937002 | 23,16263 | 17,8353268 | 45,62719 | 11,33316 | 7,085824 | 56,5338361 |
| Downregulated | ENSMUSG00000041556 | 74,02327466 | 69,1446557 | 62,634532 | 84,7939333 | 142,4878 | 107,4845067 | 182,878798 | 34,2223286 | 37,371589 | 88,4531138 | 66,2467985 | 41,16494 | 47,0825538 | 28,06435 | 69,29506 | 64,71975114 | 31,435 | 39,63405955 | 40,87436 | 9,999844 | 18,60029 | 42,1760364 |
| Downregulated | ENSMUSG00000025496 | 5,287376762 | 10,6376393 | 4,5387342 | 18,3077811 | 17,810975 | 35,8281689 | 14,7666111 | 31,7778766 | 20,287434 | 33,4395918 | 36,4357392 | 2,627549 | 2,00351293 | 1,870957 | 1,59299 | 2,734637372 | 0 | 1,981702978 | 1,901133 | 23,99963 | 23,91466 | 12,5630747 |
| Downregulated | ENSMUSG00000085584 | 15,86213028 | 5,31881967 | 21,7859242 | 19,2713485 | 12,722125 | 83,59906077 | 31,8050084 | 57,444623 | 43,7781471 | 77,6661487 | 124,764804 | 7,006798 | 5,00878232 | 0 | 0 | 4,557728953 | 0 | 3,963405955 | 18,06076 | 36,66609 | 18,60029 | 19,7419745 |
| Downregulated | ENSMUSG00000031636 | 3,172426057 | 51,0606688 | 9,0774684 | 35,6519947 | 39,4385875 | 104,8305683 | 17,0383974 | 28,1111985 | 42,7103874 | 46,3839499 | 111,515444 | 0 | 13,022834 | 0 | 3,982475 | 0 | 0 | 3,963405955 | 6,653965 | 11,99981 | 35,42912 | 8,0762623 |
| Downregulated | ENSMUSG00000020607 | 29,60930986 | 52,1244328 | 22,693671 | 37,5791295 | 47,0718625 | 157,909337 | 78,3766279 | 75,7780133 | 59,7945424 | 81,9809347 | 99,3701978 | 26,27549 | 62,1089008 | 13,0967 | 21,50536 | 10,0270037 | 19,02645 | 18,82617829 | 25,6653 | 54,66581 | 26,57184 | 43,0733989 |
| Downregulated | ENSMUSG00000061723 | 2,114950705 | 20,2115147 | 9,0774684 | 10,5992417 | 13,9943375 | 45,11695343 | 20,4460768 | 26,8889725 | 12,8131162 | 23,7313232 | 35,3316259 | 7,882647 | 11,0193211 | 7,483827 | 5,575465 | 0,911545791 | 3,308947 | 3,963405955 | 3,802266 | 27,33291 | 21,25747 | 10,7683497 |
| Downregulated | ENSMUSG00000034664 | 322,5299825 | 182,967397 | 344,943799 | 167,660732 | 597,939874 | 749,7376085 | 770,135561 | 480,334827 | 549,896238 | 482,17734 | 701,111951 | 345,0848 | 497,872963 | 357,3527 | 367,9807 | 437,5419795 | 151,3843 | 237,8043573 | 345,0556 | 207,3301 | 382,6345 | 463,936401 |
| Downregulated | ENSMUSG00000087691 | 7,402327466 | 6,3825836 | 11,8007089 | 1,92713485 | 13,9943375 | 34,50119968 | 23,8537563 | 28,1111985 | 20,287434 | 15,1017511 | 34,2275126 | 3,503399 | 17,0298599 | 0,935478 | 3,982475 | 3,646183163 | 3,308947 | 4,954257444 | 5,703399 | 10,6665 | 15,05738 | 29,6129618 |
| Downregulated | ENSMUSG00000112080 | 15,86213028 | 29,7853902 | 25,4169115 | 10,5992417 | 59,7939874 | 95,54178374 | 63,6100169 | 26,8889725 | 33,1005503 | 45,3052534 | 41,9563057 | 16,64114 | 8,01405172 | 13,0967 | 14,33691 | 50,13501849 | 31,435 | 9,908514888 | 22,8136 | 10,6665 | 13,28592 | 34,9971366 |
| Downregulated | ENSMUSG00000034209 | 32,78173592 | 19,1477508 | 53,5570636 | 10,5992417 | 41,9830125 | 88,90693765 | 64,74591 | 30,5556505 | 53,3879843 | 53,9348255 | 66,2467985 | 46,42003 | 27,0474245 | 17,77409 | 16,72639 | 30,99255688 | 9,099605 | 13,87192084 | 20,91246 | 30,66619 | 31,00048 | 21,5366995 |
| Downregulated | ENSMUSG00000052373 | 8,459802819 | 1,06376393 | 24,5091647 | 2,89070227 | 22,899825 | 30,52029203 | 12,4948247 | 19,5556163 | 20,287434 | 35,5969848 | 19,8740396 | 7,882647 | 3,00526939 | 0 | 0 | 15,49627844 | 0 | 0,990851489 | 1,901133 | 5,999906 | 7,085824 | 6,28153734 |
| Downregulated | ENSMUSG00000038055 | 24,3219331 | 23,4028065 | 10,8929621 | 31,797725 | 47,0718625 | 66,34846093 | 51,1151921 | 28,1111985 | 53,3879843 | 8,62957207 | 43,060419 | 16,64114 | 12,0210776 | 26,19339 | 32,65629 | 15,49627844 | 34,74395 | 11,89021787 | 5,703399 | 31,33284 | 31,88621 | 21,5366995 |
| Downregulated | ENSMUSG00000062859 | 27,49435916 | 10,6376393 | 13,6162026 | 10,5992417 | 10,1777 | 70,32936859 | 52,2510853 | 36,6667807 | 16,0163953 | 22,6526267 | 23,1863795 | 11,38605 | 6,01053879 | 22,45148 | 6,37196 | 5,469274744 | 1,654474 | 4,954257444 | 2,851699 | 10,6665 | 27,45757 | 20,639337 |
| Downregulated | ENSMUSG00000104639 | 10,57475352 | 7,44634754 | 3,63098736 | 7,70853939 | 8,90548749 | 26,53938437 | 22,7178632 | 12,2222602 | 8,54207749 | 6,47217906 | 5,52056654 | 11,38605 | 7,01229525 | 5,61287 | 7,964949 | 6,380820534 | 0,827237 | 4,954257444 | 1,901133 | 3,333281 | 17,71456 | 7,17889982 |
| Downregulated | ENSMUSG00000074466 | 82,48307748 | 129,7792 | 76,2507346 | 90,5753379 | 175,565325 | 272,0286898 | 161,296828 | 172,333869 | 152,689635 | 78,7448452 | 122,556577 | 44,66833 | 49,0860668 | 45,83844 | 32,65629 | 31,90410267 | 29,78053 | 45,57916849 | 71,29249 | 47,99925 | 198,4031 | 107,683497 |
| Downregulated | ENSMUSG00000025427 | 46,5289155 | 55,3157246 | 47,2028357 | 57,8140454 | 71,2438999 | 139,331768 | 53,3869784 | 46,4445888 | 50,1847052 | 30,2035023 | 40,8521924 | 17,51699 | 37,0649892 | 56,1287 | 24,69134 | 16,40782423 | 11,58132 | 39,63405955 | 30,41813 | 19,99969 | 92,11571 | 58,328561 |
| Downregulated | ENSMUSG00000034245 | 21,14950705 | 31,912918 | 35,4021268 | 45,2876689 | 87,7826624 | 130,0429834 | 48,8434058 | 58,666849 | 74,743178 | 65,8004871 | 56,3097787 | 20,14454 | 8,01405172 | 0,935478 | 14,33691 | 4,557728953 | 14,89026 | 54,49683188 | 26,61586 | 62,66569 | 18,60029 | 40,3813115 |
| Downregulated | ENSMUSG00000052951 | 3,172426057 | 3,1912918 | 4,5387342 | 18,3077811 | 22,899825 | 50,42483031 | 17,0383974 | 35,4445546 | 10,6775969 | 6,47217906 | 11,0411331 | 0 | 0 | 0 | 0 | 0 | 0 | 0,990851489 | 1,901133 | 14,66644 | 10,62874 | 0 |
| Downregulated | ENSMUSG00000072591 | 15,86213028 | 21,2752787 | 9,98521524 | 19,2713485 | 20,3554 | 50,42483031 | 17,0383974 | 31,7778766 | 26,6939921 | 35,5969848 | 14,353473 | 0 | 0 | 0 | 4,77897 | 0 | 3,308947 | 0,990851489 | 4,752832 | 1,333313 | 6,200096 | 3,58944991 |
| Downregulated | ENSMUSG00000039492 | 10,57475352 | 10,6376393 | 13,6162026 | 5,78140454 | 8,90548749 | 66,34846093 | 10,2230384 | 36,6667807 | 29,8972712 | 21,5739302 | 5,52056654 | 0 | 1,00175646 | 0 | 0 | 0 | 0 | 0 | 0 | 0 | 2,657184 | 1,79472495 |
| Downregulated | ENSMUSG00000050463 | 19,03455634 | 9,57387541 | 5,44648104 | 9,63567424 | 6,36106249 | 46,44392265 | 5,67946579 | 24,4445204 | 35,2360696 | 22,6526267 | 0 | 0 | 0 | 0,935478 | 0 | 0 | 0 | 0 | 0 | 0 | 0,885728 | 0,89736248 |
| Downregulated | ENSMUSG00000059146 | 75,08075001 | 45,7418492 | 66,2655193 | 61,6683151 | 90,3270874 | 205,6802289 | 78,3766279 | 135,667088 | 187,925705 | 110,027044 | 76,1838183 | 1,751699 | 25,0439116 | 7,483827 | 16,72639 | 5,469274744 | 19,85368 | 46,57001997 | 26,61586 | 0 | 23,91466 | 77,1731731 |
| Downregulated | ENSMUSG00000025739 | 45,47144015 | 51,0606688 | 19,0626836 | 41,4333992 | 49,6162874 | 115,446322 | 44,2998332 | 59,8890751 | 74,743178 | 35,5969848 | 16,5616996 | 4,379248 | 23,0403987 | 11,22574 | 3,982475 | 5,469274744 | 5,790658 | 14,86277233 | 13,30793 | 0 | 24,80038 | 32,3050492 |
| Downregulated | ENSMUSG00000029348 | 4,229901409 | 10,6376393 | 13,6162026 | 26,0163204 | 10,1777 | 55,73270718 | 18,1742905 | 28,1111985 | 51,2524649 | 31,2821988 | 8,83290647 | 8,758497 | 2,00351293 | 0 | 10,35443 | 3,646183163 | 0 | 4,954257444 | 0 | 27,99956 | 19,48602 | 2,69208743 |
| Downregulated | ENSMUSG00000053199 | 66,6209472 | 28,7216262 | 68,9887598 | 38,542697 | 63,6106249 | 222,9308287 | 54,5228716 | 89,2224996 | 137,740999 | 92,7678998 | 87,2249514 | 40,28909 | 35,0614763 | 18,70957 | 14,33691 | 24,61173635 | 16,54474 | 36,66150509 | 38,97323 | 73,99885 | 126,6591 | 59,2259235 |
| Downregulated | ENSMUSG00000097589 | 434,6223698 | 385,082544 | 457,504407 | 264,981042 | 502,523937 | 911,6278532 | 491,841738 | 701,557736 | 657,739966 | 309,585898 | 500,163329 | 336,3263 | 495,86945 | 286,2564 | 190,3623 | 230,621085 | 225,0084 | 363,6424964 | 228,136 | 123,9981 | 123,1162 | 240,493144 |
| Downregulated | ENSMUSG00000090222 | 169,1960564 | 135,09802 | 159,763444 | 105,028849 | 276,070112 | 420,6492423 | 163,568615 | 352,001094 | 289,362875 | 195,244068 | 280,44478 | 150,6461 | 132,231853 | 119,7412 | 78,853 | 71,10057167 | 62,87 | 120,8838816 | 58,93512 | 51,99919 | 131,9735 | 147,167446 |
| Downregulated | ENSMUSG00000031144 | 10,57475352 | 18,0839869 | 19,9704305 | 43,3605341 | 92,8715124 | 151,2744909 | 60,2023374 | 116,111472 | 67,2688602 | 34,5182883 | 51,8933255 | 10,5102 | 20,0351293 | 1,870957 | 15,1334 | 9,115457906 | 9,099605 | 15,85362382 | 14,2585 | 7,333219 | 16,82883 | 45,7654863 |
| Downregulated | ENSMUSG00000025892 | 118,4372395 | 197,860092 | 254,169115 | 196,567754 | 543,234737 | 686,043086 | 233,993991 | 628,224175 | 518,931207 | 273,988913 | 266,091307 | 56,93023 | 68,1194396 | 98,22523 | 98,76537 | 157,6974218 | 88,51434 | 188,2617829 | 116,9197 | 41,99934 | 32,77194 | 147,167446 |
| Downregulated | ENSMUSG00000114894 | 10,57475352 | 11,7014033 | 4,5387342 | 12,5263765 | 22,899825 | 39,80907656 | 7,95125211 | 12,2222602 | 22,4229534 | 4,31478604 | 25,3946061 | 1,751699 | 18,0316164 | 5,61287 | 3,982475 | 6,380820534 | 9,099605 | 10,89936638 | 7,604532 | 1,999969 | 4,42864 | 2,69208743 |
| Downregulated | ENSMUSG00000024610 | 23,26445775 | 73,3997114 | 19,0626836 | 67,4497197 | 95,4159374 | 191,0835675 | 145,394324 | 96,5558557 | 140,944279 | 59,328308 | 101,578424 | 9,634347 | 39,0685021 | 29,93531 | 16,72639 | 34,63874004 | 14,06303 | 21,79873275 | 12,35736 | 12,66647 | 47,82931 | 11,6657122 |
| Downregulated | ENSMUSG00000073421 | 25,37940846 | 68,0808918 | 34,4943799 | 48,1783712 | 69,9716874 | 94,21481452 | 51,1151921 | 55,000171 | 93,9628523 | 30,2035023 | 72,8714784 | 9,634347 | 41,072015 | 11,22574 | 5,575465 | 36,46183163 | 8,272368 | 15,85362382 | 15,20906 | 3,333281 | 52,25795 | 14,3577996 |
| Downregulated | ENSMUSG00000052605 | 68,7358979 | 104,248866 | 49,0183294 | 72,2675568 | 71,2438999 | 126,0620758 | 59,0664442 | 88,0002736 | 59,7945424 | 50,6987359 | 32,019286 | 6,130948 | 32,0562069 | 13,0967 | 0 | 16,40782423 | 9,926842 | 19,81702978 | 31,36869 | 31,33284 | 37,20058 | 31,4076867 |
| Downregulated | ENSMUSG00000030433 | 146,989074 | 319,12918 | 125,269064 | 116,591658 | 228,99825 | 345,0119968 | 135,171286 | 257,889691 | 217,822976 | 133,758367 | 103,786651 | 37,66154 | 103,180916 | 14,96765 | 40,62124 | 44,66574374 | 14,06303 | 64,40534677 | 35,17096 | 6,666563 | 13,28592 | 42,1760364 |
| Downregulated | ENSMUSG00000044991 | 41,24153874 | 46,8056131 | 34,4943799 | 27,9434553 | 49,6162874 | 50,42483031 | 44,2998332 | 47,6668148 | 45,9136665 | 25,8887162 | 23,1863795 | 10,5102 | 23,0403987 | 21,516 | 15,1334 | 9,115457906 | 8,272368 | 25,76213871 | 16,15963 | 29,33288 | 18,60029 | 29,6129618 |
| Downregulated | ENSMUSG00000071001 | 96,23025706 | 136,161784 | 139,793013 | 144,535114 | 142,4878 | 338,3771507 | 176,06344 | 96,5558557 | 113,182527 | 116,499223 | 83,9126115 | 21,89624 | 15,026347 | 66,41897 | 59,73712 | 39,196469 | 38,05289 | 76,29556464 | 91,25438 | 56,66578 | 57,57232 | 110,375585 |
| Downregulated | ENSMUSG00000081769 | 40,18406339 | 41,4867934 | 31,7711394 | 45,2876689 | 96,6881499 | 108,8114759 | 73,8330553 | 42,7779108 | 40,5748681 | 22,6526267 | 34,2275126 | 7,006798 | 20,0351293 | 13,0967 | 3,982475 | 7,292366325 | 12,40855 | 16,84447531 | 15,20906 | 3,999938 | 26,57184 | 19,7419745 |
| Downregulated | ENSMUSG00000037493 | 51,81629226 | 72,3359475 | 32,6788862 | 72,2675568 | 127,22125 | 151,2744909 | 86,32788 | 59,8890751 | 65,1333408 | 87,3744172 | 65,1426852 | 32,40644 | 35,0614763 | 45,83844 | 21,50536 | 45,57728953 | 34,74395 | 44,588317 | 45,62719 | 24,66628 | 23,91466 | 76,2758106 |
| Downregulated | ENSMUSG00000036095 | 31,72426057 | 56,3794885 | 25,4169115 | 32,7612924 | 54,7051374 | 96,86875296 | 51,1151921 | 73,3335613 | 40,5748681 | 38,8330743 | 22,0822662 | 5,255098 | 7,01229525 | 1,870957 | 0 | 13,67318686 | 4,136184 | 58,46023784 | 20,91246 | 0 | 3,542912 | 7,17889982 |
| Downregulated | ENSMUSG00000057897 | 47,58639085 | 203,178911 | 51,7415699 | 60,7047477 | 189,559662 | 279,9905051 | 124,948247 | 201,667294 | 125,995643 | 223,290177 | 129,181257 | 1,751699 | 23,0403987 | 18,70957 | 8,761444 | 0 | 0 | 49,54257444 | 16,15963 | 2,666625 | 35,42912 | 105,888772 |
| Downregulated | ENSMUSG00000041617 | 72,96579931 | 111,695213 | 93,4979245 | 68,4132871 | 211,187275 | 236,2005209 | 143,122538 | 168,667191 | 168,70603 | 181,221014 | 138,014164 | 27,15134 | 21,0368858 | 43,96748 | 35,84227 | 30,99255688 | 31,435 | 74,31386166 | 67,49022 | 54,66581 | 84,14416 | 79,8652605 |
| Downregulated | ENSMUSG00000035578 | 99,40268312 | 86,1648786 | 52,6493167 | 58,7776129 | 100,504787 | 171,1790292 | 94,2791321 | 77,0002394 | 13,8808759 | 76,5874522 | 47,4768723 | 34,15814 | 94,1651077 | 75,77375 | 74,87053 | 69,27748009 | 19,02645 | 40,62491104 | 55,13286 | 1,333313 | 14,17165 | 84,3520729 |
| Downregulated | ENSMUSG00000102854 | 65,56347184 | 35,1042098 | 19,9704305 | 22,1620508 | 30,5331 | 94,21481452 | 65,8818032 | 46,4445888 | 33,1005503 | 20,4952337 | 24,2904928 | 7,882647 | 29,0509375 | 24,32244 | 11,94742 | 8,203912116 | 3,308947 | 24,77128722 | 15,20906 | 8,666531 | 22,1432 | 20,639337 |
| Downregulated | ENSMUSG00000102964 | 48,64386621 | 28,7216262 | 29,9556457 | 14,4535114 | 40,7108 | 82,27209155 | 78,3766279 | 51,3334929 | 43,7781471 | 43,1478604 | 35,3316259 | 19,26869 | 32,0562069 | 26,19339 | 31,8598 | 17,31937002 | 6,617895 | 39,63405955 | 29,46756 | 10,6665 | 4,42864 | 22,4340619 |

| | | | | | | | | | | | | | | | | | | | | | | | |
|---|---|---|---|---|---|---|---|---|---|---|---|---|---|---|---|---|---|---|---|---|---|---|---|
| Downregulated | ENSMUSG00000003032 | 9167,253829 | 6881,48889 | 8816,94506 | 11915,4748 | 11308,6969 | 19737,34016 | 13690,9202 | 9248,58431 | 10575,0919 | 10416,9722 | 12243,5125 | 5900,599 | 5068,88771 | 13353,02 | 8071,68 | 6669,78055 | 3000,388 | 7158,902007 | 3599,795 | 3998,604 | 8325,843 | 5724,27524 |
| Downregulated | ENSMUSG00000038539 | 332,0472606 | 254,23958 | 264,15433 | 249,563963 | 351,13065 | 603,7709945 | 277,157931 | 382,556745 | 464,475463 | 224,368874 | 323,505199 | 148,0186 | 219,384666 | 382,6107 | 276,3837 | 238,8249971 | 116,6404 | 171,4173076 | 110,2657 | 95,9985 | 234,7179 | 134,604372 |
| Downregulated | ENSMUSG00000045038 | 125,8395669 | 110,631449 | 175,19514 | 135,863007 | 265,892412 | 311,8377664 | 280,56561 | 204,111746 | 245,584728 | 126,207492 | 133,59771 | 141,0118 | 150,26347 | 218,9019 | 157,706 | 168,6359713 | 65,35171 | 123,8564361 | 87,45212 | 49,99922 | 98,31581 | 89,7362477 |
| Downregulated | ENSMUSG00000037872 | 818,4859227 | 373,381141 | 371,268458 | 358,447082 | 1016,49779 | 1438,434633 | 1020,03206 | 770,002394 | 800,819764 | 485,413429 | 948,433332 | 706,8107 | 641,124137 | 1018,736 | 589,4063 | 602,5317676 | 124,0855 | 573,703012 | 237,6416 | 105,3317 | 531,4368 | 511,496612 |
| Downregulated | ENSMUSG00000005338 | 2044,099856 | 1132,90859 | 1251,78289 | 1502,20161 | 2702,17935 | 3399,695138 | 3110,07547 | 2383,34074 | 2440,89864 | 1443,29593 | 2998,77175 | 1594,922 | 2141,75532 | 3566,979 | 2070,887 | 2112,051597 | 420,2363 | 1785,514383 | 1038,969 | 325,9949 | 1395,907 | 1422,31953 |
| Downregulated | ENSMUSG00002076173 | 21594,70417 | 16092,6208 | 18024,2213 | 33083,1239 | 65249,2346 | 104604,9835 | 50797,142 | 35816,1113 | 42412,4825 | 16345,4882 | 24439,5481 | 13869,08 | 35683,567 | 43360,36 | 24481,07 | 37883,84306 | 14666,08 | 29586,82546 | 26567,38 | 15901,75 | 25449,62 | 31389,7395 |
| Downregulated | ENSMUSG00002075551 | 3223,184874 | 1820,10009 | 2341,0791 | 3993,02341 | 7424,63214 | 14048,62312 | 7367,40302 | 3807,23406 | 6434,31987 | 2144,44866 | 3901,93643 | 1642,218 | 4322,57914 | 5801,837 | 2967,74 | 4987,067021 | 2700,101 | 3559,138548 | 3095,995 | 2151,3 | 2837,873 | 3896,34788 |
| Downregulated | ENSMUSG00002075501 | 542,4848557 | 405,294059 | 447,519192 | 1166,88015 | 1684,40935 | 3261,690339 | 1587,97864 | 1117,11458 | 1409,44279 | 384,015957 | 765,150523 | 338,9538 | 1028,80389 | 1488,346 | 728,7929 | 1146,724605 | 530,2588 | 835,2878051 | 655,8909 | 466,6594 | 534,9797 | 750,195031 |
| Downregulated | ENSMUSG00000063903 | 180,8282852 | 314,874124 | 54,4648104 | 178,259973 | 376,5749 | 540,076472 | 296,468114 | 376,445615 | 359,835014 | 224,368874 | 319,088746 | 84,95742 | 336,590172 | 292,8047 | 122,6602 | 204,1862571 | 46,32526 | 116,9204757 | 136,8816 | 3,333281 | 100,973 | 299,719067 |
| Downregulated | ENSMUSG00000074676 | 26,43688381 | 21,2752787 | 14,5239494 | 26,9798879 | 43,255225 | 72,98330702 | 52,2510853 | 33,0001026 | 34,1683099 | 26,9674127 | 51,8933255 | 6,130948 | 44,0772844 | 37,41914 | 14,33691 | 11,85009528 | 4,136184 | 15,85362382 | 8,555098 | 0 | 23,02893 | 39,483949 |
| Downregulated | ENSMUSG00000049676 | 20,09203169 | 5,31881967 | 21,7859242 | 29,8705901 | 6,36106249 | 35,8281689 | 14,7666111 | 7,33335613 | 20,287434 | 5,39348255 | 26,4987194 | 4,379248 | 7,01229525 | 0 | 8,761444 | 4,557728953 | 5,790658 | 13,87192084 | 3,802266 | 15,99975 | 0 | 8,0762623 |
| Downregulated | ENSMUSG00000055148 | 319,3575564 | 146,799423 | 330,41985 | 549,233432 | 278,614537 | 842,6254538 | 368,029383 | 213,889554 | 231,703852 | 453,052534 | 707,736631 | 403,7667 | 106,186185 | 377,9333 | 291,5172 | 126,7048649 | 93,47776 | 205,1062582 | 102,6612 | 149,9977 | 147,9166 | 130,117559 |
| Downregulated | ENSMUSG00000038910 | 46,5289155 | 17,0202229 | 27,2324052 | 33,7248598 | 25,44425 | 62,36755328 | 21,58197 | 19,5556163 | 27,7617518 | 45,3052534 | 58,5180054 | 12,2619 | 17,0298599 | 14,96765 | 11,15093 | 16,40782423 | 4,136184 | 11,89021787 | 4,752832 | 22,66631 | 9,743008 | 4,48681239 |
| Downregulated | ENSMUSG00000046207 | 146,989074 | 146,799423 | 59,9112914 | 89,6117704 | 263,347987 | 339,70412 | 162,432722 | 95,3336297 | 195,400022 | 127,286188 | 165,616996 | 30,65474 | 30,0526939 | 2,806435 | 18,31938 | 9,115457906 | 0,827237 | 23,78043573 | 13,30793 | 0,666656 | 10,62874 | 13,4604372 |
| Downregulated | ENSMUSG00000059857 | 108,9199613 | 175,521049 | 100,759899 | 29,8705901 | 223,9094 | 432,5919653 | 179,471119 | 108,778116 | 227,432813 | 108,948347 | 88,3290647 | 18,39284 | 12,0210776 | 43,96748 | 15,1334 | 18,23091581 | 12,40855 | 67,37790124 | 19,01133 | 1,333313 | 8,85728 | 10,7683497 |
| Downregulated | ENSMUSG00000036594 | 21,14950705 | 24,4665705 | 12,7084558 | 10,5992417 | 57,2495624 | 100,8496606 | 32,9409016 | 25,6667465 | 61,9300618 | 32,3608953 | 28,706946 | 14,0136 | 10,0175646 | 4,677392 | 4,77897 | 1,823091581 | 0 | 14,86277233 | 3,802266 | 0 | 14,17165 | 8,0762623 |
| Downregulated | ENSMUSG00000016386 | 37,01163733 | 57,4432524 | 34,4943799 | 41,4333992 | 54,7051374 | 98,19572218 | 51,1151921 | 35,4445546 | 58,7267827 | 17,2591441 | 50,7892122 | 1,751699 | 17,0298599 | 5,61287 | 16,72639 | 17,31937002 | 3,308947 | 10,89936638 | 14,2585 | 7,999875 | 7,085824 | 12,5630747 |
| Downregulated | ENSMUSG00000022528 | 517,1054473 | 432,951921 | 544,648104 | 664,861523 | 647,556162 | 1011,150545 | 697,438399 | 455,890306 | 797,616485 | 517,774324 | 497,955102 | 261,8791 | 239,419795 | 181,4828 | 407,8054 | 204,1862571 | 267,1975 | 314,099922 | 236,6911 | 353,9945 | 247,1181 | 310,487417 |
| Downregulated | ENSMUSG00000040270 | 47,58639085 | 31,912918 | 26,3246584 | 35,6519947 | 52,1607124 | 95,54178374 | 37,4844742 | 46,4445888 | 40,5748681 | 49,6200394 | 67,3509118 | 3,503399 | 11,0193211 | 4,677392 | 10,35443 | 1,823091581 | 2,481711 | 13,87192084 | 0,950566 | 28,66622 | 11,51446 | 17,0498871 |
| Downregulated | ENSMUSG00000030401 | 240,046905 | 108,503921 | 163,394431 | 123,33663 | 179,381962 | 375,5322889 | 134,035393 | 133,222636 | 113,182527 | 182,29971 | 175,554016 | 45,54418 | 97,1703771 | 23,38696 | 53,36516 | 22,78864477 | 87,68711 | 105,0302578 | 64,63852 | 103,3317 | 106,2874 | 113,067672 |
| Downregulated | ENSMUSG00000092021 | 25,37940846 | 6,3825836 | 29,0478989 | 22,1620508 | 48,344075 | 106,1575375 | 11,3589316 | 24,4445204 | 44,8459068 | 45,3052534 | 59,6221187 | 8,758497 | 10,0175646 | 6,548349 | 18,31938 | 4,557728953 | 2,481711 | 13,87192084 | 8,555098 | 0,666656 | 7,085824 | 25,1261494 |
| Downregulated | ENSMUSG00000087291 | 35,95416198 | 4,25505574 | 28,140152 | 20,2349159 | 20,3554 | 115,446322 | 39,7562605 | 42,7779108 | 16,0163953 | 44,2265569 | 35,3316259 | 22,77209 | 29,0509375 | 5,61287 | 24,69134 | 2,734637372 | 9,926842 | 18,82617829 | 19,01133 | 32,66616 | 5,314368 | 17,0498871 |
| Downregulated | ENSMUSG00000054672 | 0 | 0 | 7,26197472 | 0 | 30,5331 | 95,54178374 | 10,2230384 | 26,8889725 | 2,13551937 | 1,07869651 | 38,6439658 | 14,0136 | 0 | 0 | 17,52289 | 4,557728953 | 0 | 0 | 0 | 0 | 0 | 2,69208743 |
| Downregulated | ENSMUSG00000106678 | 19,03455634 | 14,8926951 | 10,8929621 | 20,2349159 | 50,8884999 | 107,4845067 | 30,6691153 | 39,1112327 | 21,3551937 | 42,0691639 | 30,9151726 | 22,77209 | 14,0245905 | 16,83861 | 24,69134 | 28,25791951 | 19,02645 | 11,89021787 | 12,35736 | 24,66628 | 2,657184 | 20,639337 |
| Downregulated | ENSMUSG00000084399 | 30,66678522 | 36,1679738 | 5,44648104 | 38,542697 | 19,0831875 | 115,446322 | 39,7562605 | 13,4444862 | 17,084155 | 16,1804476 | 26,4987194 | 15,76529 | 36,0632327 | 33,67722 | 20,70887 | 23,70019056 | 2,481711 | 25,76213871 | 7,604532 | 0,666656 | 17,71456 | 29,6129618 |
| Downregulated | ENSMUSG00000049321 | 37,01163733 | 36,1679738 | 21,7859242 | 28,9070227 | 38,166375 | 87,57996843 | 15,9025042 | 24,4445204 | 29,8972712 | 21,5739302 | 33,1233993 | 18,39284 | 45,0790409 | 26,19339 | 33,45279 | 25,52328214 | 11,58132 | 19,81702978 | 9,505665 | 35,99944 | 16,82883 | 19,7419745 |
| Downregulated | ENSMUSG00000085517 | 146,989074 | 101,057574 | 102,575393 | 56,850478 | 292,608875 | 84,92602999 | 86,32788 | 147,889349 | 88,6240539 | 53,9348255 | 168,929336 | 38,53739 | 29,0509375 | 19,64505 | 29,47031 | 10,0270037 | 53,77039 | 95,12174293 | 97,90835 | 71,33222 | 66,4296 | 102,299322 |
| Downregulated | ENSMUSG00000062310 | 9,517278171 | 7,44634754 | 10,8929621 | 20,2349159 | 53,4329249 | 17,25059984 | 18,1742905 | 18,3333903 | 11,7453565 | 23,7313232 | 28,706946 | 6,130948 | 13,022834 | 5,61287 | 5,575465 | 12,76164107 | 3,308947 | 8,917663399 | 9,505665 | 4,666594 | 3,542912 | 8,97362477 |
| Downregulated | ENSMUSG00000025213 | 1008,831486 | 2950,88115 | 1789,16902 | 2316,41609 | 5815,28333 | 4428,096283 | 5009,28883 | 7560,69017 | 4087,38408 | 4103,36152 | 4254,14858 | 315,3059 | 1232,16045 | 627,706 | 240,5415 | 405,6378768 | 73,62408 | 1237,57351 | 501,8991 | 47,99925 | 1024,787 | 2183,28291 |
| Downregulated | ENSMUSG00000020251 | 2469,204948 | 3100,87187 | 1972,53388 | 2431,08061 | 5226,24894 | 3666,415951 | 4517,44709 | 5833,6848 | 4372,47591 | 3645,9942 | 3952,72565 | 720,8243 | 1311,29921 | 921,4462 | 558,343 | 486,7654522 | 132,3579 | 1171,18646 | 780,4151 | 111,3316 | 978,7295 | 2658,88502 |
| Downregulated | ENSMUSG00000022132 | 1414,902021 | 3885,92965 | 1557,69358 | 4260,89515 | 6780,89262 | 4735,953141 | 5012,69651 | 7005,79956 | 6275,22367 | 3555,38369 | 4356,83112 | 516,7513 | 1867,27405 | 932,672 | 187,1763 | 268,9060082 | 173,7197 | 1839,020363 | 842,2019 | 187,3304 | 1510,166 | 2827,58917 |
| Downregulated | ENSMUSG00000024076 | 182,943236 | 226,581718 | 281,40152 | 322,795087 | 507,612787 | 258,7589976 | 578,169618 | 502,334895 | 384,393487 | 362,442027 | 215,302095 | 77,95062 | 171,300355 | 53,32227 | 46,9932 | 42,84265216 | 75,27855 | 170,4264561 | 100,76 | 71,33222 | 134,6307 | 300,61643 |
| Downregulated | ENSMUSG00000025551 | 96,23025706 | 203,178911 | 116,191596 | 245,709693 | 384,208175 | 132,6969219 | 281,701503 | 294,556471 | 237,04265 | 255,651073 | 168,929336 | 56,93023 | 71,124709 | 46,77392 | 50,17918 | 60,16202218 | 44,67079 | 141,6917629 | 74,14419 | 11,33316 | 35,42912 | 157,935796 |
| Downregulated | ENSMUSG00000028399 | 150,1615 | 238,283121 | 127,084558 | 173,442136 | 461,813137 | 315,818674 | 444,134225 | 305,556505 | 333,141022 | 221,132784 | 210,885642 | 53,42683 | 96,1686206 | 154,3539 | 40,62124 | 42,84265216 | 80,24197 | 138,7192084 | 98,85892 | 81,99872 | 140,8308 | 195,62502 |
| Downregulated | ENSMUSG00000071648 | 194,5754648 | 370,189849 | 310,449419 | 382,536267 | 731,522187 | 498,9404262 | 717,884476 | 537,77945 | 497,576014 | 540,426951 | 433,91653 | 70,94383 | 177,310894 | 134,7089 | 173,6359 | 154,9627844 | 109,1953 | 206,0971097 | 163,4974 | 127,998 | 234,7179 | 259,337756 |
| Downregulated | ENSMUSG00000023960 | 673,6117994 | 1230,77487 | 801,54046 | 1014,6365 | 3716,13271 | 2395,17944 | 2910,15827 | 2867,34225 | 2128,04505 | 1375,33805 | 1856,01447 | 634,1152 | 749,313835 | 419,0943 | 624,452 | 746,5560025 | 661,7895 | 1403,045708 | 670,1494 | 383,994 | 1201,933 | 1559,61599 |
| Downregulated | ENSMUSG00000024784 | 248,5067078 | 1258,43273 | 337,681824 | 1013,67293 | 3226,3309 | 1990,453828 | 3606,46078 | 2935,7869 | 1033,59138 | 1739,93747 | 1724,62499 | 151,522 | 263,46195 | 532,2872 | 140,1831 | 360,0605873 | 33,91671 | 778,8092702 | 272,8126 | 6,666563 | 377,3201 | 1382,83558 |
| Downregulated | ENSMUSG00000026950 | 11,63222888 | 8,51011147 | 0,90774684 | 11,5628091 | 207,370637 | 110,1384451 | 118,132888 | 243,222978 | 106,775969 | 207,10973 | 68,4550251 | 9,634347 | 5,00878232 | 0,935478 | 1,59299 | 9,115457906 | 14,06303 | 26,7529902 | 57,03399 | 59,33241 | 77,94407 | 180,369858 |
| Downregulated | ENSMUSG00000016942 | 1,057475352 | 12,7651672 | 6,35422788 | 10,5992417 | 36,8941625 | 17,25059984 | 18,1742905 | 37,8890067 | 11,7453565 | 33,4395918 | 16,5616996 | 1,751699 | 0 | 9,354784 | 0 | 0 | 1,654474 | 0,990851489 | 4,752832 | 3,999938 | 24,80038 | 17,0498871 |
| Downregulated | ENSMUSG00000024617 | 163,9086796 | 299,981429 | 157,040203 | 324,722222 | 683,178112 | 506,9022415 | 736,058767 | 832,335921 | 455,933386 | 709,782303 | 811,523282 | 26,27549 | 63,1106573 | 101,0317 | 46,9932 | 30,99255688 | 18,19921 | 186,2800799 | 87,45212 | 37,99941 | 149,688 | 448,681239 |
| Downregulated | ENSMUSG00000057969 | 4154,820659 | 6603,8465 | 3491,19435 | 5952,91955 | 12649,6089 | 10654,23586 | 12944,6384 | 11216,3682 | 9220,10489 | 11900,1799 | 11015,7385 | 1527,482 | 3304,79458 | 1677,313 | 911,9867 | 1399,222789 | 861,9808 | 4296,332055 | 2632,119 | 1103,983 | 6224,896 | 7812,43773 |
| Downregulated | ENSMUSG00000019852 | 99,40268312 | 397,847711 | 84,4204561 | 290,997362 | 596,667662 | 214,9690134 | 487,298165 | 694,22438 | 369,444851 | 559,843488 | 293,69414 | 26,27549 | 75,1317348 | 55,19322 | 35,04578 | 38,28492321 | 36,39842 | 113,9479212 | 149,2389 | 12,66647 | 89,45853 | 431,631352 |
| Downregulated | ENSMUSG00000025429 | 39,12658804 | 51,0606688 | 28,140152 | 45,2876689 | 148,848862 | 53,07876874 | 103,366277 | 162,556061 | 121,724604 | 120,814009 | 118,140124 | 51,67513 | 38,0667457 | 28,99983 | 20,70887 | 24,61173635 | 12,40855 | 51,52427742 | 39,92379 | 34,66613 | 54,02941 | 97,81251 |
| Downregulated | ENSMUSG00000074505 | 6,344852114 | 36,1679738 | 6,35422788 | 52,0326409 | 38,166375 | 54,40573796 | 40,8921537 | 90,4447256 | 34,1683099 | 35,5969848 | 58,5180054 | 0 | 3,00526939 | 0,935478 | 7,168455 | 1,823091581 | 0 | 22,78958424 | 25,6653 | 0 | 8,85728 | 44,8681239 |
| Downregulated | ENSMUSG00000049744 | 188,2306127 | 411,676642 | 387,607901 | 465,403066 | 659,006074 | 575,9046409 | 587,256763 | 1076,78113 | 778,396811 | 464,918195 | 379,814978 | 124,3707 | 194,340754 | 86,99949 | 30,26681 | 68,3659343 | 95,95947 | 461,7367938 | 249,0484 | 119,3315 | 243,5752 | 521,367599 |
| Downregulated | ENSMUSG00000027499 | 501,243317 | 1043,55242 | 484,736813 | 805,542366 | 770,960774 | 748,4106393 | 606,566947 | 1334,67082 | 1051,74329 | 723,805358 | 720,985991 | 229,4726 | 324,569095 | 140,3218 | 141,7761 | 241,5596345 | 193,5734 | 391,3863381 | 364,067 | 220,6632 | 544,7227 | 698,148007 |
| Downregulated | ENSMUSG00000030249 | 455,7718768 | 981,854111 | 569,157269 | 450,949554 | 1066,11407 | 879,7805919 | 763,320202 | 1278,44842 | 1037,86241 | 554,450006 | 483,601629 | 207,5764 | 415,728933 | 142,1927 | 86,02145 | 79,30448379 | 281,2605 | 233,8409514 | 769,9589 | 242,6629 | 819,2984 | 677,50867 |
| Downregulated | ENSMUSG00000021223 | 12,68970423 | 48,933141 | 45,387342 | 11,5628091 | 55,9773499 | 92,8878453 | 71,561269 | 154,000479 | 84,3530152 | 70,1152731 | 51,8933255 | 10,5102 | 35,0614763 | 24,32244 | 15,9299 | 6,380820534 | 21,50816 | 7,92681191 | 32,31926 | 31,33284 | 83,25843 | 66,4048233 |
| Downregulated | ENSMUSG00000038173 | 7,402327466 | 3,1912918 | 2,72324052 | 5,78140454 | 26,7164625 | 13,26969219 | 32,9409016 | 40,3334587 | 8,54207749 | 23,7313232 | 26,4987194 | 6,130948 | 0 | 0 | 3,18598 | 0 | 5,790658 | 5,945108933 | 7,604532 | 19,33303 | 28,3433 | 4,48681239 |
| Downregulated | ENSMUSG00000023092 | 2181,571652 | 1916,90261 | 2526,25946 | 2286,5455 | 4464,19366 | 4871,304002 | 8152,3052 | 6065,90775 | 3764,92065 | 5710,61932 | 4286,16786 | 502,7377 | 1706,99302 | 655,7704 | 635,603 | 635,3474161 | 328,413 | 571,721309 | 711,9743 | 1812,638 | 4894,533 | 3333,7016 |
| Downregulated | ENSMUSG00000029830 | 0 | 0 | 2,72324052 | 2,89070227 | 24,1720375 | 26,53938437 | 48,8434058 | 46,4445888 | 23,4907131 | 12,9443581 | 27,6028327 | 1,751699 | 4,00702586 | 0,935478 | 0,796495 | 1,823091581 | 3,308947 | 6,935960422 | 0 | 1,333313 | 16,82883 | 9,87098725 |
| Downregulated | ENSMUSG00000047261 | 21,14950705 | 260,622164 | 191,534583 | 435,532476 | 361,30835 | 111,4654144 | 762,184309 | 810,335852 | 601,148703 | 134,837064 | 448,270003 | 148,8944 | 195,342511 | 400,3848 | 270,8083 | 70,18902588 | 91,82329 | 95,12174293 | 118,8208 | 195,3303 | 256,8611 | 125,630747 |
| Downregulated | ENSMUSG00000026208 | 7,402327466 | 69,1446557 | 16,3394431 | 99,2474447 | 128,493462 | 135,3508603 | 121,540568 | 228,556266 | 172,977069 | 78,7448452 | 240,696701 | 27,15134 | 23,0403987 | 50,51583 | 51,77217 | 33,72719425 | 8,272368 | 40,62491104 | 15,20906 | 79,3321 | 181,5742 | 70,8916357 |
| Downregulated | ENSMUSG00000036699 | 0 | 12,7651672 | 9,98521524 | 17,3442136 | 47,0718625 | 37,15513812 | 32,9409016 | 57,444623 | 50,1847052 | 15,1017511 | 78,3920449 | 7,882647 | 3,00526939 | 5,61287 | 3,982475 | 2,734637372 | 0 | 2,972554466 | 6,653965 | 15,99975 | 10,62874 | 13,4604372 |
| Downregulated | ENSMUSG00000117786 | 1,057475352 | 1,06376393 | 0 | 0 | 133,582312 | 71,65633781 | 77,2407348 | 195,556163 | 176,180348 | 0 | 41,9563057 | 49,04758 | 0 | 33,67722 | 58,14413 | 35,55028583 | 31,435 | 35,6706536 | 0 | 97,99847 | 25,68611 | 0 |
| Downregulated | ENSMUSG00000067235 | 415,5878135 | 278,706151 | 333,14309 | 297,742334 | 895,637599 | 797,5085004 | 1064,33189 | 1203,89263 | 950,30612 | 421,770335 | 687,862591 | 402,8909 | 187,328459 | 247,9018 | 507,3673 | 541,4581996 | 330,0675 | 429,0386947 | 208,1741 | 1302,646 | 307,3476 | 221,648532 |
| Downregulated | ENSMUSG00000043541 | 22,2069824 | 17,0202229 | 17,24719 | 7,70853939 | 33,077525 | 30,52029203 | 24,9896495 | 48,8890409 | 12,8131162 | 3,23608953 | 11,0411331 | 17,51699 | 27,0474245 | 28,99983 | 23,89485 | 14,58473265 | 7,445132 | 8,917663399 | 3,802266 | 33,99947 | 25,68611 | 15,2551621 |
| Downregulated | ENSMUSG00000090397 | 20,09203169 | 8,51011147 | 37,2176204 | 5,78140454 | 35,62195 | 49,09786109 | 34,0767947 | 53,777945 | 5,33879843 | 7,55087556 | 11,0411331 | 12,2619 | 9,01580818 | 30,87079 | 7,964949 | 11,85009528 | 16,54474 | 6,935960422 | 14,2585 | 17,99972 | 9,743008 | 17,9472495 |
| Downregulated | ENSMUSG00000070323 | 1845,29449 | 5037,98599 | 3986,82412 | 3826,32624 | 2080,06744 | 2273,098271 | 5194,43941 | 9297,47335 | 8443,84359 | 1881,24671 | 4468,34656 | 1518,723 | 1201,106 | 857,8337 | 865,79 | 555,1313865 | 1087,816 | 2837,798664 | 1736,685 | 1398,645 | 1697,941 | 3852,37712 |
| Downregulated | ENSMUSG00000050447 | 2,114950705 | 17,0202229 | 41,7563546 | 47,2148038 | 20,3554 | 46,44392265 | 68,1535895 | 255,445239 | 58,7267827 | 1,07869651 | 61,8303453 | 18,39284 | 10,0175646 | 29,93531 | 9,557939 | 26,43482793 | 7,445132 | 15,85362382 | 9,505665 | 17,99972 | 10,62874 | 24,2287869 |
| Downregulated | ENSMUSG00000038738 | 11,63222888 | 15,956459 | 30,8633926 | 16,3806462 | 50,8884999 | 46,44392265 | 88,5996663 | 199,222842 | 89,6918136 | 50,6987359 | 51,8933255 | 8,758497 | 16,0281034 | 20,58052 | 9,557939 | 11,85009528 | 16,54474 | 32,69809913 | 12,35736 | 19,33303 | 10,62874 | 39,483949 |
| Downregulated | ENSMUSG00000067242 | 336,277162 | 303,172721 | 317,711394 | 288,10666 | 306,603212 | 376,8592581 | 372,572956 | 994,891982 | 548,828478 | 335,474614 | 219,718548 | 117,3639 | 124,217802 | 62,67705 | 60,53362 | 82,03912116 | 79,41474 | 52,51512891 | 148,2884 | 42,666 | 85,91562 | 128,322834 |
| Downregulated | ENSMUSG00000116093 | 126,8970423 | 124,46038 | 100,759899 | 109,846686 | 81,4215999 | 131,3699526 | 148,802004 | 380,112293 | 232,771611 | 141,309243 | 96,0578579 | 43,79248 | 60,1053879 | 16,83861 | 31,8598 | 17,31937002 | 23,98987 | 31,70724764 | 57,03399 | 32,66616 | 48,71504 | 88,8388853 |
| Downregulated | ENSMUSG00000067203 | 1613,707388 | 1397,78581 | 1366,15899 | 1622,64754 | 659,006074 | 1215,503804 | 1472,11753 | 4324,23566 | 2628,82435 | 1434,66636 | 1236,60691 | 750,6032 | 1109,94616 | 615,5448 | 400,637 | 988,1156371 | 478,9701 | 859,0682408 | 797,5253 | 1022,651 | 236,4894 | 902,746652 |
| Downregulated | ENSMUSG00000034488 | 25,37940846 | 251,048288 | 35,4021268 | 73,2311242 | 162,8432 | 62,36755328 | 62,4741237 | 640,446435 | 295,769433 | 163,961869 | 133,59771 | 11,38605 | 14,0245905 | 10,29026 | 0 | 6,380820534 | 4,136184 | 94,13089144 | 30,41813 | 6,666563 | 7,971552 | 127,425472 |
| Downregulated | ENSMUSG00000092224 | 196,6904155 | 169,138465 | 136,162026 | 141,644411 | 286,247812 | 250,7971823 | 272,614358 | 2209,78465 | 1358,19032 | 155,332297 | 204,260962 | 112,1088 | 73,1282219 | 124,4186 | 130,6252 | 127,6164107 | 107,5408 | 108,9936638 | 65,58909 | 486,6591 | 96,54435 | 55,6364736 |
| Downregulated | ENSMUSG00000091705 | 731,7729438 | 622,301901 | 604,559395 | 608,974612 | 886,732112 | 869,1648382 | 950,742573 | 3319,56588 | 2177,162 | 524,246503 | 864,520721 | 493,9792 | 443,778114 | 407,8686 | 539,2271 | 678,1900682 | 425,1997 | 530,1055465 | 308,9341 | 763,9881 | 347,2054 | 306,000605 |
| Downregulated | ENSMUSG00000063929 | 29,60930986 | 14,8926951 | 23,6014178 | 33,7248598 | 17,810975 | 41,13604578 | 15,9025042 | 58,666849 | 42,7103874 | 12,9443581 | 12,1452464 | 32,40644 | 8,01405172 | 24,32244 | 7,964949 | 20,96555318 | 14,89026 | 19,81702978 | 14,2585 | 5,33325 | 0 | 7,17889982 |
| Downregulated | ENSMUSG00000006638 | 40,18406339 | 12,7651672 | 15,4316963 | 17,3442136 | 13,9943375 | 43,78998421 | 36,3485811 | 80,6669174 | 68,3366199 | 16,1804476 | 69,5591385 | 31,53059 | 22,0386422 | 22,45148 | 7,964949 | 24,61173635 | 12,40855 | 20,80788127 | 16,15963 | 5,33325 | 7,971552 | 25,1261494 |
| Downregulated | ENSMUSG00000082902 | 1962,674254 | 1042,48866 | 1215,47302 | 1310,4517 | 1421,06136 | 879,7805919 | 1555,03773 | 4668,9034 | 4945,86286 | 636,43094 | 2484,25494 | 796,1474 | 693,215473 | 1500,507 | 532,0586 | 1236,967638 | 496,3421 | 944,2814688 | 722,4305 | 34,66613 | 175,3741 | 741,221406 |
| Downregulated | ENSMUSG00000118433 | 2474,492324 | 1605,21978 | 1741,05844 | 1973,38608 | 1929,94636 | 1616,248508 | 2157,06111 | 5820,24032 | 6808,03576 | 1325,71801 | 3769,44284 | 1153,494 | 1403,46081 | 2066,472 | 1011,549 | 1592,470496 | 648,5537 | 1555,636837 | 865,0155 | 45,99928 | 376,4344 | 1119,90837 |
| Downregulated | ENSMUSG00000067279 | 90,9428803 | 149,990715 | 14,5239494 | 22,1620508 | 145,032225 | 79,61815312 | 21,58197 | 224,889588 | 72,6076586 | 58,2496115 | 64,0385719 | 30,65474 | 128,224827 | 26,19339 | 21,50536 | 10,93854949 | 10,75408 | 18,82617829 | 15,20906 | 15,33309 | 23,91466 | 51,1496612 |
| Downregulated | ENSMUSG00000063713 | 5,287376762 | 8,51011147 | 1,81549368 | 5,78140454 | 21,6276125 | 6,634846093 | 4,54357263 | 25,6667465 | 22,4229534 | 14,0230546 | 11,0411331 | 0 | 17,0298599 | 16,83861 | 2,389485 | 0,911545791 | 1,654474 | 5,945108933 | 3,802266 | 0 | 4,42864 | 1,79472495 |
| Downregulated | ENSMUSG00000019368 | 19,03455634 | 5,31881967 | 3,63098736 | 0 | 11,4499125 | 25,21241515 | 9,08714527 | 20,7778424 | 14,9486356 | 8,62957207 | 7,72879316 | 4,379248 | 33,0579633 | 0 | 1,59299 | 0 | 2,481711 | 0 | 0 | 0 | 0 | 8,97362477 |
| Downregulated | ENSMUSG00000021340 | 94,11530636 | 40,4230295 | 76,2507346 | 29,8705901 | 100,504787 | 102,1766298 | 40,8921537 | 144,222671 | 87,5562942 | 158,568387 | 109,307218 | 38,53739 | 120,210776 | 42,09653 | 37,43526 | 25,52328214 | 25,64434 | 55,48768337 | 9,505665 | 6,666563 | 52,25795 | 62,8153734 |
| Upregulated | ENSMUSG00000039116 | 12,68970423 | 5,31881967 | 29,0478989 | 15,4170788 | 5,08884999 | 0 | 0 | 7,33335613 | 14,9486356 | 0 | 4,41645324 | 1,751699 | 5,00878232 | 9,354784 | 0,796495 | 10,93854949 | 0 | 3,963405955 | 11,4068 | 41,99934 | 48,71504 | 19,7419745 |
| Upregulated | ENSMUSG00000025964 | 372,231324 | 141,480603 | 274,139546 | 168,624299 | 58,5217749 | 143,3126756 | 59,0664442 | 149,111575 | 129,198922 | 99,2400788 | 99,3701978 | 32,40644 | 132,231853 | 105,7091 | 47,7897 | 41,01956058 | 103,4046 | 149,6185748 | 225,2843 | 456,6595 | 511,0651 | 189,343483 |
| Upregulated | ENSMUSG00000008136 | 226,2997254 | 94,6749901 | 264,15433 | 117,555226 | 7,63327499 | 17,25059984 | 20,4460768 | 53,777945 | 22,4229534 | 38,8330743 | 73,9755917 | 107,7295 | 95,1668641 | 129,096 | 61,33011 | 60,16202218 | 189,4372 | 144,6643174 | 183,4593 | 445,3264 | 480,0646 | 224,340619 |
| Upregulated | ENSMUSG00000024866 | 68,7358979 | 32,976682 | 54,4648104 | 34,6884273 | 13,9943375 | 38,48210734 | 29,5332221 | 15,8889383 | 18,1519147 | 17,2591441 | 11,0411331 | 23,64794 | 20,0351293 | 32,74174 | 23,09835 | 38,28492321 | 33,08947 | 42,60661402 | 53,23172 | 85,99866 | 124,8877 | 44,8681239 |
| Upregulated | ENSMUSG00000087365 | 6,344852114 | 5,31881967 | 20,8781773 | 8,67210682 | 2,544425 | 5,307876874 | 2,27178632 | 0 | 2,13551937 | 4,31478604 | 0 | 2,627549 | 0 | 0 | 2,389485 | 2,734637372 | 4,963421 | 2,972554466 | 1,901133 | 19,99969 | 24,80038 | 3,58944991 |
| Upregulated | ENSMUSG00000078452 | 19,03455634 | 20,2115147 | 88,9591903 | 25,052753 | 3,8166375 | 7,961815312 | 12,4948247 | 17,1111643 | 2,13551937 | 44,2265569 | 32,019286 | 1,751699 | 18,0316164 | 6,548349 | 14,33691 | 13,67318686 | 15,7175 | 56,47853486 | 7,604532 | 96,66516 | 103,6302 | 35,8944991 |
| Upregulated | ENSMUSG00000022844 | 964,4175213 | 1002,06563 | 1301,70897 | 1397,17276 | 926,170699 | 711,2555012 | 1299,46177 | 1016,89205 | 681,230679 | 1054,96519 | 776,191656 | 323,1885 | 509,89404 | 246,0308 | 394,265 | 412,9302432 | 1551,896 | 1637,877511 | 1169,197 | 2591,293 | 2615,555 | 1650,2496 |
| Upregulated | ENSMUSG00000070858 | 27,49435916 | 41,4867934 | 75,3429877 | 66,4861523 | 41,9830125 | 54,40573796 | 65,8818032 | 41,5556847 | 29,8972712 | 40,9904673 | 51,8933255 | 37,66154 | 25,0439116 | 28,06435 | 11,94742 | 13,67318686 | 66,17895 | 73,32301017 | 58,93512 | 137,3312 | 167,4026 | 52,0470237 |
| Upregulated | ENSMUSG00000044433 | 31,72426057 | 4,25505574 | 49,0183294 | 24,0891856 | 0 | 14,5966614 | 1,13589316 | 22,0000684 | 7,4743178 | 25,8887162 | 25,3946061 | 0 | 9,01580818 | 20,58052 | 4,77897 | 11,85009528 | 52,94316 | 34,67980211 | 51,33059 | 101,3318 | 81,48698 | 35,8944991 |
| Upregulated | ENSMUSG00000068794 | 40,18406339 | 24,4665705 | 74,4352409 | 56,850478 | 34,3497375 | 18,57756906 | 9,08714527 | 31,7778766 | 23,4907131 | 46,3839499 | 25,3946061 | 3,503399 | 31,0544504 | 0 | 34,24928 | 33,72719425 | 26,47158 | 112,9570697 | 75,09475 | 145,9977 | 147,0309 | 78,967898 |
| Upregulated | ENSMUSG00000015829 | 0 | 1,06376393 | 0,90774684 | 4,81783712 | 1,2722125 | 0 | 0 | 9,77780817 | 5,33879843 | 11,8656616 | 6,62467985 | 1,751699 | 3,00526939 | 10,29026 | 0,796495 | 4,557728953 | 6,617895 | 0 | 13,30793 | 41,99934 | 32,77194 | 16,1525246 |
| Upregulated | ENSMUSG00000067049 | 0 | 2,12752787 | 0 | 0,96356742 | 0 | 0 | 0 | 0 | 5,33879843 | 0 | 0 | 0 | 3,00526939 | 0 | 1,59299 | 0 | 17,37197 | 0,990851489 | 13,30793 | 50,66588 | 46,05786 | 9,87098725 |

| | | | | | | | | | | | | | | | | | | | | | | | |
|---|---|---|---|---|---|---|---|---|---|---|---|---|---|---|---|---|---|---|---|---|---|---|---|
| Upregulated | ENSMUSG00000056133 | 0 | 0 | 0 | 0 | 0 | 0 | 0 | 0 | 0 | 0 | 0 | 0 | 1,00175646 | 0,935478 | 0,796495 | 4,557728953 | 15,7175 | 1,981702978 | 11,4068 | 34,66613 | 34,54339 | 14,3577996 |
| Upregulated | ENSMUSG00000033453 | 974,9922748 | 296,790138 | 785,201017 | 614,756017 | 202,281787 | 240,1814286 | 207,868448 | 480,334827 | 162,299472 | 709,782303 | 356,628599 | 909,132 | 644,129407 | 381,6752 | 436,4792 | 217,859444 | 1808,34 | 930,409548 | 1999,992 | 5067,254 | 4836,075 | 2666,06392 |
| Upregulated | ENSMUSG00000029669 | 29,60930986 | 34,0404459 | 71,7120004 | 43,3605341 | 13,9943375 | 27,86635359 | 17,0383974 | 63,5557531 | 30,9650309 | 44,2265569 | 32,019286 | 52,55098 | 43,075528 | 48,64488 | 68,49857 | 59,25047639 | 116,6404 | 51,52427742 | 92,20495 | 251,9961 | 209,9175 | 127,425472 |
| Upregulated | ENSMUSG00000030162 | 0 | 6,3825836 | 10,8929621 | 0 | 0 | 0 | 0 | 6,11113011 | 6,40655811 | 0 | 6,62467985 | 36,78569 | 51,0895797 | 31,80627 | 6,37196 | 8,203912116 | 19,02645 | 37,65235657 | 22,8136 | 307,9952 | 248,0038 | 116,657122 |
| Upregulated | ENSMUSG00000017713 | 15,86213028 | 8,51011147 | 19,0626836 | 3,8542697 | 25,44425 | 10,61575375 | 52,2510853 | 8,55558215 | 14,9486356 | 20,4952337 | 16,5616996 | 37,66154 | 41,072015 | 38,35461 | 11,94742 | 8,203912116 | 38,88013 | 28,73469318 | 38,97323 | 193,997 | 200,1745 | 70,8916357 |
| Upregulated | ENSMUSG00000001864 | 111,034912 | 65,9533639 | 143,424001 | 46,2512364 | 67,4272624 | 82,27209155 | 38,6203674 | 74,5557873 | 82,2174958 | 96,0039893 | 91,6414046 | 178,6733 | 177,310894 | 42,09653 | 96,37589 | 151,3166012 | 151,3843 | 105,0302578 | 143,5355 | 573,991 | 652,7815 | 158,833159 |
| Upregulated | ENSMUSG00000036585 | 10,57475352 | 15,956459 | 29,0478989 | 11,5628091 | 12,722125 | 0 | 0 | 12,2222602 | 10,6775969 | 21,5739302 | 0 | 34,15814 | 9,01580818 | 1,870957 | 9,557939 | 2,734637372 | 8,272368 | 4,954257444 | 9,505665 | 130,6646 | 153,2309 | 61,9180109 |
| Upregulated | ENSMUSG00000025572 | 111,034912 | 77,6547672 | 152,501469 | 62,6318826 | 106,86585 | 98,19572218 | 106,773957 | 90,4447256 | 98,2338911 | 117,57792 | 112,619557 | 100,7227 | 137,240636 | 47,7094 | 50,97568 | 92,06612485 | 88,51434 | 117,9113272 | 198,6684 | 633,9901 | 653,6673 | 258,440394 |
| Upregulated | ENSMUSG00000027221 | 2,114950705 | 34,0404459 | 6,35422788 | 11,5628091 | 0 | 2,653938437 | 0 | 4,88890409 | 14,9486356 | 16,1804476 | 4,41645324 | 17,51699 | 9,01580818 | 0 | 3,18598 | 11,85009528 | 4,136184 | 14,86277233 | 21,86303 | 229,3298 | 193,0887 | 42,1760364 |
| Upregulated | ENSMUSG00000025278 | 527,6802008 | 554,22101 | 802,448207 | 510,690735 | 346,0418 | 447,1886267 | 477,075126 | 695,446606 | 513,592409 | 787,448452 | 455,998797 | 750,6032 | 574,006454 | 345,1915 | 422,9388 | 675,4554309 | 967,0399 | 662,879646 | 1117,866 | 4295,933 | 3661,6 | 1359,50415 |
| Upregulated | ENSMUSG00000019936 | 0 | 1,06376393 | 13,6162026 | 14,4535114 | 0 | 0 | 0 | 13,4444862 | 1,06775969 | 6,47217906 | 3,31233993 | 0 | 12,0210776 | 0 | 0 | 0 | 0 | 7,92681191 | 17,1102 | 45,99928 | 54,91514 | 22,4340619 |
| Upregulated | ENSMUSG00000090124 | 117,3797641 | 22,3390426 | 77,1584814 | 18,3077811 | 13,9943375 | 18,57756906 | 18,1742905 | 64,7779792 | 48,0491859 | 20,4952337 | 18,7699262 | 25,39964 | 40,0702586 | 14,96765 | 3,18598 | 11,85009528 | 27,29882 | 54,49683188 | 62,73739 | 312,6618 | 358,7198 | 104,094047 |
| Upregulated | ENSMUSG00000047686 | 42,29901409 | 1,06376393 | 128,900051 | 13,4899439 | 0 | 13,26969219 | 6,81535895 | 24,4445204 | 11,7453565 | 25,8887162 | 27,6028327 | 0 | 21,0368858 | 0 | 0 | 9,115457906 | 0 | 3,963405955 | 30,41813 | 303,3286 | 394,149 | 80,762623 |
| Upregulated | ENSMUSG00000025791 | 502,3007924 | 638,25836 | 679,902383 | 674,497197 | 575,040049 | 456,4774112 | 687,215361 | 545,112806 | 493,304975 | 708,703607 | 486,913969 | 472,9588 | 442,776357 | 512,6422 | 525,6867 | 387,406961 | 845,4361 | 738,1843592 | 799,4264 | 2099,301 | 2333,893 | 990,688175 |
| Upregulated | ENSMUSG00000067377 | 272,8286409 | 295,726374 | 494,722028 | 321,83152 | 155,209925 | 161,8902447 | 214,683807 | 288,445341 | 238,11041 | 303,113719 | 280,44478 | 158,5288 | 328,57612 | 202,0633 | 207,0887 | 168,6359713 | 386,3196 | 343,8254666 | 495,2451 | 1129,982 | 1093,874 | 400,223665 |
| Upregulated | ENSMUSG00000074280 | 42,29901409 | 19,1477508 | 57,1880509 | 59,7411803 | 68,6994749 | 120,7541989 | 118,132888 | 66,0002052 | 75,8109377 | 107,869651 | 124,764804 | 65,68873 | 231,405743 | 97,28975 | 95,57939 | 131,2625939 | 183,6466 | 107,0119608 | 179,6571 | 444,6597 | 423,378 | 169,601508 |
| Upregulated | ENSMUSG00000029084 | 2,114950705 | 1,06376393 | 9,98521524 | 26,9798879 | 1,2722125 | 0 | 1,13589316 | 17,1111643 | 8,54207749 | 4,31478604 | 11,0411331 | 36,78569 | 132,231853 | 12,16122 | 21,50536 | 0 | 29,78053 | 10,89936638 | 75,09475 | 235,9963 | 207,2604 | 40,3813115 |
| Upregulated | ENSMUSG00000035413 | 274,9435916 | 132,970492 | 298,64871 | 254,3818 | 161,570987 | 195,0644751 | 136,307179 | 127,111506 | 158,028433 | 204,952337 | 162,304656 | 226,8451 | 531,932683 | 234,8051 | 144,1656 | 92,06612485 | 366,4659 | 232,8500999 | 476,2338 | 863,3199 | 873,3278 | 271,003468 |
| Upregulated | ENSMUSG00000031963 | 673,6117994 | 157,437062 | 990,351802 | 292,924497 | 41,9830125 | 53,07876874 | 54,5228716 | 297,000923 | 127,063403 | 134,837064 | 207,573302 | 341,5814 | 1482,59957 | 226,3858 | 73,27754 | 128,5279565 | 450,0168 | 369,5876053 | 1197,714 | 4701,927 | 3232,022 | 756,476568 |
| Upregulated | ENSMUSG00000067586 | 595,3586234 | 159,56459 | 716,212257 | 387,354104 | 78,8771749 | 153,9284294 | 144,258431 | 344,667738 | 124,927883 | 496,200394 | 575,243034 | 1535,365 | 2191,84314 | 322,74 | 333,7314 | 568,8045734 | 1475,791 | 760,9739434 | 2771,852 | 7521,883 | 6457,843 | 1727,42277 |
| Upregulated | ENSMUSG00000016918 | 778,3018593 | 829,735868 | 759,784105 | 700,513517 | 388,024812 | 342,3580584 | 188,558264 | 691,779928 | 277,617518 | 594,361777 | 450,47823 | 1170,135 | 2266,97488 | 1020,607 | 657,9048 | 704,6248962 | 1804,204 | 1083,991529 | 2633,069 | 9640,516 | 8657,106 | 2535,94636 |
| Upregulated | ENSMUSG00000028487 | 46,5289155 | 43,6143213 | 74,4352409 | 23,1256182 | 3,8166375 | 13,26969219 | 19,3101837 | 23,2222944 | 23,4907131 | 65,8004871 | 45,2686457 | 74,44722 | 177,310894 | 16,83861 | 20,70887 | 0 | 125,74 | 82,24067357 | 156,8435 | 623,9903 | 493,3505 | 147,167446 |
| Upregulated | ENSMUSG00000024885 | 124,7820916 | 105,312629 | 143,424001 | 71,3039894 | 8,90548749 | 17,25059984 | 36,3485811 | 55,000171 | 22,4229534 | 131,600974 | 59,6221187 | 73,57137 | 194,340754 | 72,96731 | 62,12661 | 61,07356797 | 223,3539 | 149,6185748 | 294,6756 | 660,6564 | 595,2092 | 209,085457 |
| Upregulated | ENSMUSG00000031380 | 206,2076937 | 115,950269 | 226,028963 | 182,114243 | 39,4385875 | 49,09786109 | 105,638064 | 57,444623 | 92,8950927 | 174,748834 | 71,7673651 | 180,425 | 300,526939 | 145,9346 | 78,853 | 115,7663154 | 313,5228 | 347,7888726 | 530,4161 | 1152,649 | 980,5009 | 279,977093 |
| Upregulated | ENSMUSG00000023456 | 2374,032166 | 2371,12981 | 2637,91232 | 2664,26393 | 2124,59487 | 2087,322581 | 2621,64141 | 2154,78448 | 2257,24398 | 2496,10372 | 1766,58129 | 1983,8 | 2426,25416 | 3356,496 | 2996,414 | 2691,79472 | 3667,141 | 2644,582624 | 3612,153 | 7959,209 | 7996,353 | 2857,20213 |
| Upregulated | ENSMUSG00000106755 | 106,8050106 | 130,842964 | 149,778229 | 129,118035 | 156,482137 | 155,2553986 | 145,394324 | 70,8891093 | 104,640449 | 133,758367 | 117,036011 | 125,2465 | 119,209019 | 173,999 | 167,2639 | 185,9553413 | 248,1711 | 168,4447531 | 242,3945 | 468,6594 | 494,2362 | 131,014922 |
| Upregulated | ENSMUSG00000024909 | 3004,287476 | 2148,80315 | 3883,34098 | 3094,97857 | 1866,33574 | 2645,976622 | 2897,66345 | 2961,45365 | 2880,81563 | 3101,25246 | 3558,55719 | 4023,653 | 5137,00715 | 4113,298 | 3255,275 | 3751,922474 | 9107,878 | 6146,251785 | 7480,958 | 17397,73 | 22173,32 | 5740,42777 |
| Upregulated | ENSMUSG00000028480 | 684,1865529 | 277,642387 | 881,422182 | 425,896801 | 146,304437 | 303,8759511 | 251,032388 | 592,779621 | 508,25361 | 256,729769 | 432,812417 | 828,5538 | 1662,91573 | 944,8332 | 527,2797 | 935,2459812 | 2252,566 | 836,2786566 | 2081,741 | 4654,594 | 4455,212 | 931,462252 |
| Upregulated | ENSMUSG00000110063 | 0 | 2,12752787 | 11,8007089 | 0 | 0 | 9,28878453 | 0 | 2,44445204 | 0 | 0 | 0 | 8,758497 | 8,01405172 | 5,61287 | 2,389485 | 3,646183163 | 21,50816 | 6,935960422 | 14,2585 | 43,99931 | 46,05786 | 8,97362477 |
| Upregulated | ENSMUSG00000046623 | 26,43688381 | 2,12752787 | 3,63098736 | 18,3077811 | 0 | 5,307876874 | 0 | 0 | 17,084155 | 12,9443581 | 2,20822662 | 0,87585 | 23,0403987 | 4,677392 | 9,557939 | 5,469274744 | 26,47158 | 28,73469318 | 16,15963 | 103,9984 | 107,1731 | 17,9472495 |
| Upregulated | ENSMUSG00000035107 | 59,21861973 | 65,9533639 | 103,48314 | 99,2474447 | 49,6162874 | 82,27209155 | 115,861102 | 69,6668832 | 107,843728 | 50,6987359 | 87,2249514 | 95,46762 | 163,286304 | 238,547 | 106,7303 | 85,68530432 | 120,7766 | 190,2434859 | 67,49022 | 499,9922 | 610,2666 | 106,786135 |
| Upregulated | ENSMUSG00000045763 | 39,12658804 | 42,5505574 | 38,1253673 | 56,850478 | 6,36106249 | 31,84726125 | 53,3869784 | 6,11113011 | 6,40655811 | 35,5969848 | 39,7480791 | 27,15134 | 41,072015 | 84,19306 | 25,48784 | 32,81564846 | 59,56105 | 52,51512891 | 67,49022 | 223,9965 | 240,918 | 67,3021858 |
| Upregulated | ENSMUSG00000015143 | 43,35648945 | 39,3592656 | 101,667646 | 77,0853939 | 27,988675 | 21,2315075 | 38,6203674 | 33,0001026 | 29,8972712 | 52,8561289 | 76,1838183 | 122,619 | 83,1457866 | 68,28992 | 64,51609 | 113,031678 | 136,4941 | 110,9753667 | 145,4367 | 498,6589 | 284,3187 | 104,99141 |
| Upregulated | ENSMUSG00000027035 | 101,5176338 | 232,964302 | 148,870482 | 260,163204 | 213,7317 | 127,389045 | 152,209683 | 166,222739 | 104,640449 | 229,762356 | 207,573302 | 413,4011 | 262,460194 | 270,3533 | 367,9807 | 271,6406456 | 494,6876 | 333,9169517 | 406,8425 | 1881,971 | 1050,473 | 533,033312 |
| Upregulated | ENSMUSG00000011256 | 266,4837888 | 56,3794885 | 285,032508 | 94,4296076 | 72,5161124 | 106,1575375 | 124,948247 | 184,556129 | 79,0142167 | 134,837064 | 227,447342 | 261,0032 | 296,519914 | 204,8698 | 49,38269 | 79,30448379 | 410,3095 | 332,9261002 | 645,4346 | 2711,958 | 1595,196 | 585,977698 |
| Upregulated | ENSMUSG00000030041 | 0 | 4,25505574 | 6,35422788 | 3,8542697 | 0 | 0 | 0 | 7,33335613 | 1,06775969 | 0 | 1,10411331 | 0,87585 | 5,00878232 | 3,741914 | 0,796495 | 0 | 9,099605 | 1,981702978 | 23,76416 | 88,66528 | 46,94358 | 17,9472495 |
| Upregulated | ENSMUSG00000022054 | 1,057475352 | 6,3825836 | 3,63098736 | 3,8542697 | 2,544425 | 1,326969219 | 2,27178632 | 4,88890409 | 1,06775969 | 0 | 13,2493597 | 7,006798 | 14,0245905 | 7,483827 | 2,389485 | 2,734637372 | 19,85368 | 11,89021787 | 9,505665 | 125,3314 | 62,00096 | 20,639337 |
| Upregulated | ENSMUSG00000059654 | 11,63222888 | 0 | 3,63098736 | 3,8542697 | 1,2722125 | 0 | 0 | 2,44445204 | 0 | 0 | 0 | 9,634347 | 75,1317348 | 0 | 12,74392 | 0 | 23,98987 | 56,47853486 | 64,63852 | 668,6562 | 310,8905 | 19,7419745 |
| Upregulated | ENSMUSG00000024304 | 5,287376762 | 9,57387541 | 12,7084558 | 2,89070227 | 0 | 1,326969219 | 0 | 14,6667123 | 9,60983717 | 1,07869651 | 7,72879316 | 2,627549 | 11,0193211 | 0 | 8,761444 | 0,911545791 | 7,445132 | 30,71639615 | 29,46756 | 431,3266 | 201,946 | 20,639337 |
| Upregulated | ENSMUSG00000015652 | 28,55183451 | 13,8289311 | 24,5091647 | 17,3442136 | 12,722125 | 9,28878453 | 2,27178632 | 24,4445204 | 14,9486356 | 7,55087556 | 20,9781529 | 155,9012 | 246,43209 | 302,1595 | 65,31259 | 131,2625939 | 320,1407 | 149,6185748 | 331,7477 | 1870,637 | 899,0139 | 158,833159 |
| Upregulated | ENSMUSG00000000693 | 356,3691937 | 327,639292 | 676,271396 | 478,89301 | 124,676825 | 169,85206 | 132,8995 | 238,334074 | 142,012038 | 319,294167 | 248,425494 | 498,3585 | 1046,83551 | 398,5138 | 219,0361 | 104,8277659 | 1053,073 | 662,879646 | 2039,916 | 8341,87 | 3556,198 | 804,03678 |
| Upregulated | ENSMUSG00000022871 | 13,74717958 | 5,31881967 | 18,1549368 | 7,70853939 | 2,544425 | 0 | 2,27178632 | 0 | 10,6775969 | 12,9443581 | 2,20822662 | 11,38605 | 22,0386422 | 10,29026 | 19,91237 | 19,1424616 | 35,57118 | 12,88106935 | 9,505665 | 123,9981 | 62,88669 | 15,2551621 |
| Upregulated | ENSMUSG00000034786 | 15,86213028 | 8,51011147 | 18,1549368 | 10,5992417 | 1,2722125 | 7,961815312 | 2,27178632 | 1,22222602 | 2,13551937 | 3,23608953 | 16,5616996 | 4,379248 | 18,0316164 | 6,548349 | 9,557939 | 4,557728953 | 19,85368 | 12,88106935 | 11,4068 | 90,66525 | 47,82931 | 17,0498871 |
| Upregulated | ENSMUSG00000041696 | 11,63222888 | 1,06376393 | 0 | 27,9434553 | 0 | 3,980907656 | 2,27178632 | 2,44445204 | 2,13551937 | 22,6526267 | 7,72879316 | 16,64114 | 18,0316164 | 13,0967 | 4,77897 | 8,203912116 | 38,88013 | 25,76213871 | 17,1102 | 208,6634 | 75,28688 | 16,1525246 |
| Upregulated | ENSMUSG00000031289 | 2,114950705 | 0 | 8,16972156 | 0 | 2,544425 | 19,90453828 | 0 | 0 | 5,33879843 | 6,47217906 | 4,41645324 | 3,503399 | 2,00351293 | 12,16122 | 7,168455 | 2,734637372 | 100,0957 | 13,87192084 | 65,58909 | 413,3269 | 141,7165 | 17,9472495 |
| Upregulated | ENSMUSG00000074743 | 1459,315986 | 676,553862 | 1839,0951 | 624,391691 | 552,140224 | 642,2531018 | 556,587648 | 878,78051 | 1104,06351 | 1077,61781 | 1029,0336 | 2042,481 | 2310,05041 | 961,6718 | 1197,928 | 1498,58128 | 1464,209 | 1236,582658 | 3250,937 | 13330,46 | 4983,106 | 3222,42866 |
| Upregulated | ENSMUSG00000042684 | 0 | 0 | 1,81549368 | 12,5263765 | 0 | 0 | 0 | 2,44445204 | 7,4743178 | 0 | 0 | 1,751699 | 7,01229525 | 0 | 0 | 7,292366325 | 9,926842 | 10,89936638 | 14,2585 | 94,66519 | 43,40067 | 23,3314244 |
| Upregulated | ENSMUSG00000030761 | 13,74717958 | 15,956459 | 1,81549368 | 28,9070227 | 17,810975 | 5,307876874 | 0 | 13,4444862 | 2,13551937 | 23,7313232 | 15,4575863 | 2,627549 | 4,00702586 | 15,90313 | 28,67382 | 34,63874004 | 37,22566 | 36,66150509 | 47,52832 | 398,6604 | 124,8877 | 71,7889982 |
| Upregulated | ENSMUSG00000019853 | 19,03455634 | 34,0404459 | 65,3577725 | 37,5791295 | 5,08884999 | 34,50119968 | 4,54357263 | 66,0002052 | 61,9300618 | 44,2265569 | 55,2056654 | 8,758497 | 30,0526939 | 13,0967 | 0 | 11,85009528 | 29,78053 | 19,81702978 | 82,69928 | 280,6623 | 202,8317 | 57,4311986 |
| Upregulated | ENSMUSG00000021087 | 0 | 4,25505574 | 10,8929621 | 1,92713485 | 5,08884999 | 0 | 2,27178632 | 8,55558215 | 10,6775969 | 7,55087556 | 9,93701978 | 1,751699 | 16,0281034 | 4,677392 | 0,796495 | 0 | 8,272368 | 6,935960422 | 11,4068 | 61,33238 | 44,2864 | 8,0762623 |
| Upregulated | ENSMUSG00000029082 | 3,172426057 | 0 | 8,16972156 | 4,81783712 | 0 | 3,980907656 | 0 | 3,66667807 | 21,3551937 | 26,9674127 | 9,93701978 | 10,5102 | 31,0544504 | 19,64505 | 2,389485 | 5,469274744 | 19,85368 | 1,981702978 | 14,2585 | 116,6648 | 74,40115 | 23,3314244 |
| Upregulated | ENSMUSG00000026051 | 84,59802819 | 82,9735868 | 87,1436966 | 52,9962083 | 24,1720375 | 21,2315075 | 26,1255426 | 124,667054 | 20,287434 | 77,6661487 | 44,1645324 | 10,5102 | 190,333728 | 26,19339 | 3,982475 | 17,31937002 | 26,47158 | 36,66150509 | 125,4748 | 1071,317 | 683,782 | 301,513792 |
| Upregulated | ENSMUSG00000009614 | 28,55183451 | 37,2317377 | 73,527494 | 85,7575007 | 41,9830125 | 94,21481452 | 17,0383974 | 83,1113695 | 48,0491859 | 60,4070045 | 86,1208381 | 29,77889 | 93,1633512 | 13,0967 | 23,09835 | 19,1424616 | 98,44118 | 84,22237655 | 89,35325 | 713,3222 | 542,9513 | 173,190958 |
| Upregulated | ENSMUSG00000032374 | 33,83921127 | 39,3592656 | 69,8965067 | 53,9597757 | 6,36106249 | 3,980907656 | 10,2230384 | 39,1112327 | 23,4907131 | 31,2821988 | 43,060419 | 110,3571 | 127,223071 | 43,96748 | 32,65629 | 43,75419795 | 64,52447 | 100,0760004 | 96,00722 | 1499,31 | 1069,959 | 166,909421 |
| Upregulated | ENSMUSG00000022464 | 7,402327466 | 8,51011147 | 9,0774684 | 6,74497197 | 2,544425 | 17,25059984 | 0 | 13,4444862 | 5,33879843 | 0 | 1,10411331 | 15,76529 | 8,01405172 | 5,61287 | 4,77897 | 5,469274744 | 3,308947 | 8,917663399 | 17,1102 | 405,327 | 253,3182 | 34,9971366 |
| Upregulated | ENSMUSG00000020865 | 3,172426057 | 0 | 0,90774684 | 10,5992417 | 1,2722125 | 0 | 0 | 4,88890409 | 4,27103874 | 0 | 0 | 0,87585 | 32,0562069 | 7,483827 | 1,59299 | 9,115457906 | 25,64434 | 10,89936638 | 35,17096 | 272,6624 | 195,7459 | 29,6129618 |
| Upregulated | ENSMUSG00000039683 | 30,66678522 | 20,2115147 | 37,2176204 | 37,5791295 | 8,90548749 | 37,15513812 | 13,6307179 | 35,4445546 | 18,1519147 | 29,1248057 | 24,2904928 | 24,52379 | 48,0843103 | 63,61253 | 23,89485 | 6,380820534 | 36,39842 | 23,78043573 | 58,93512 | 509,3254 | 423,378 | 87,9415228 |
| Upregulated | ENSMUSG00000051323 | 41,24153874 | 24,4665705 | 108,929621 | 62,6318826 | 10,1777 | 18,57756906 | 26,1255426 | 56,222397 | 89,6918136 | 62,5643975 | 41,9563057 | 34,15814 | 71,124709 | 83,25758 | 14,33691 | 12,76164107 | 38,05289 | 95,12174293 | 47,52832 | 607,3239 | 326,8336 | 75,3784481 |
| Upregulated | ENSMUSG00000027684 | 1,057475352 | 2,12752787 | 2,72324052 | 11,5628091 | 0 | 0 | 0 | 6,11113011 | 1,06775969 | 0 | 0 | 0 | 20,0351293 | 0,935478 | 3,982475 | 0 | 0 | 1,981702978 | 6,653965 | 100,6651 | 58,45805 | 6,28153734 |
| Upregulated | ENSMUSG00000029814 | 2,114950705 | 11,7014033 | 4,5387342 | 8,67210682 | 2,544425 | 0 | 6,81535895 | 9,77780817 | 13,8808759 | 8,62957207 | 6,62467985 | 7,006798 | 12,0210776 | 0,935478 | 10,35443 | 0,911545791 | 14,89026 | 3,963405955 | 8,555098 | 105,3317 | 67,31533 | 1,79472495 |
| Upregulated | ENSMUSG00000031562 | 16,91960564 | 35,1042098 | 68,081013 | 32,7612924 | 10,1777 | 0 | 7,95125211 | 31,7778766 | 22,4229534 | 29,1248057 | 18,7699262 | 20,14454 | 64,1124137 | 66,41897 | 59,73712 | 37,37337742 | 43,01632 | 65,39619826 | 53,23172 | 179,9972 | 111,6017 | 29,6129618 |
| Upregulated | ENSMUSG00000020399 | 12,68970423 | 11,7014033 | 14,5239494 | 5,78140454 | 7,63327499 | 0 | 0 | 0 | 0 | 0 | 0 | 4,379248 | 9,01580818 | 0 | 0 | 0,911545791 | 15,7175 | 40,62491104 | 16,15963 | 73,33219 | 62,88669 | 18,844612 |
| Upregulated | ENSMUSG00000031304 | 6,344852114 | 2,12752787 | 9,0774684 | 4,81783712 | 3,8166375 | 0 | 5,67946579 | 2,44445204 | 10,6775969 | 3,23608953 | 1,10411331 | 13,13775 | 1,00175646 | 9,354784 | 1,59299 | 4,557728953 | 2,481711 | 33,68895062 | 11,4068 | 57,99909 | 47,82931 | 10,7683497 |
| Upregulated | ENSMUSG00000042228 | 100,4601585 | 77,6547672 | 96,221165 | 69,3768545 | 48,344075 | 41,13604578 | 61,3382305 | 67,2224312 | 83,2852555 | 35,5969848 | 60,726232 | 33,28229 | 64,1124137 | 28,06435 | 45,40021 | 41,93110637 | 82,72368 | 118,9021787 | 124,5242 | 305,9952 | 212,5747 | 81,6599854 |
| Upregulated | ENSMUSG00000033068 | 207,2651691 | 55,3157246 | 102,575393 | 108,883119 | 122,1324 | 61,04058406 | 115,861102 | 101,44476 | 79,0142167 | 111,10574 | 72,8714784 | 92,84007 | 122,214289 | 177,7409 | 113,1023 | 91,15457906 | 198,5368 | 201,1428522 | 249,0484 | 651,3232 | 441,9783 | 176,780408 |
| Upregulated | ENSMUSG00000076437 | 164,966155 | 177,648577 | 256,892356 | 274,616716 | 199,737362 | 281,3174743 | 228,314525 | 199,222842 | 206,077619 | 158,568387 | 147,951183 | 110,3571 | 189,331972 | 228,2567 | 197,5307 | 129,4395023 | 349,0939 | 408,2308134 | 394,4851 | 1059,983 | 707,6967 | 323,050492 |
| Upregulated | ENSMUSG00000020911 | 5,287376762 | 0 | 2,72324052 | 20,2349159 | 0 | 0 | 0 | 1,22222602 | 0 | 0 | 1,10411331 | 5,255098 | 6,01053879 | 0 | 1,59299 | 2,734637372 | 0 | 0 | 0,950566 | 34,66613 | 27,45757 | 4,48681239 |
| Upregulated | ENSMUSG00000021108 | 4,229901409 | 20,2115147 | 21,7859242 | 40,4698318 | 13,9943375 | 19,90453828 | 32,9409016 | 20,7778424 | 14,9486356 | 23,7313232 | 1,10411331 | 25,39964 | 27,0474245 | 37,41914 | 19,91237 | 21,87709898 | 11,58132 | 9,908514888 | 14,2585 | 125,3314 | 71,74397 | 31,4076867 |
| Upregulated | ENSMUSG00000024479 | 2,114950705 | 2,12752787 | 13,6162026 | 23,1256182 | 6,36106249 | 9,28878453 | 10,2230384 | 7,33335613 | 12,8131162 | 8,62957207 | 3,31233993 | 2,627549 | 10,0175646 | 6,548349 | 0,796495 | 14,58473265 | 1,654474 | 12,88106935 | 0,950566 | 63,33234 | 30,11475 | 10,7683497 |
| Upregulated | ENSMUSG00000028182 | 8,459802819 | 1,06376393 | 6,35422788 | 7,70853939 | 2,544425 | 18,57756906 | 6,81535895 | 3,66667807 | 8,54207749 | 1,07869651 | 5,52056654 | 10,5102 | 3,00526939 | 0 | 3,18598 | 9,115457906 | 0 | 5,945108933 | 5,703399 | 35,99944 | 27,45757 | 7,17889982 |
| Upregulated | ENSMUSG00000047250 | 80,36812678 | 124,46038 | 291,386736 | 132,972305 | 161,570987 | 383,4941042 | 63,6100169 | 143,000445 | 269,075441 | 257,808466 | 291,485914 | 301,2923 | 83,1457866 | 49,58035 | 246,9134 | 166,8128797 | 114,9859 | 76,29556464 | 210,0752 | 1552,642 | 1231,162 | 327,537304 |
| Upregulated | ENSMUSG00000068699 | 649,2898663 | 276,578623 | 665,378434 | 408,552588 | 267,164625 | 484,3437648 | 290,788649 | 420,445751 | 188,993464 | 621,329189 | 568,618354 | 390,629 | 284,498836 | 72,03184 | 368,7772 | 361,8836789 | 245,6893 | 263,566496 | 506,6519 | 2093,301 | 1407,422 | 428,041902 |
| Upregulated | ENSMUSG00000037762 | 47,58639085 | 21,2752787 | 43,5718483 | 38,542697 | 7,63327499 | 7,961815312 | 7,95125211 | 117,333698 | 37,371589 | 64,7217906 | 38,6439658 | 0 | 11,0193211 | 7,483827 | 7,964949 | 2,734637372 | 3,308947 | 5,945108933 | 17,1102 | 146,6644 | 180,6885 | 40,3813115 |
| Upregulated | ENSMUSG00000014158 | 47,58639085 | 17,0202229 | 27,2324052 | 20,2349159 | 17,810975 | 14,5966614 | 5,67946579 | 157,667157 | 35,2360696 | 34,5182883 | 28,706946 | 37,66154 | 49,0860668 | 29,93531 | 38,23176 | 35,55028583 | 72,79684 | 38,64320806 | 80,79815 | 237,3296 | 226,7464 | 73,5837231 |
| Upregulated | ENSMUSG00000021701 | 1787,133345 | 1573,30686 | 3249,73369 | 1610,12117 | 2648,74642 | 2245,231918 | 2102,53824 | 1860,22801 | 1121,14767 | 2190,83261 | 2542,77295 | 3210,865 | 1542,70496 | 1451,862 | 2634,009 | 2412,861708 | 1819,921 | 2122,403889 | 1352,656 | 6655,229 | 8112,383 | 2113,28863 |
| Upregulated | ENSMUSG00000037185 | 25,37940846 | 44,6780852 | 18,1549368 | 24,0891856 | 44,5274375 | 54,40573796 | 60,2023374 | 51,3334929 | 36,3038293 | 64,7217906 | 36,4357392 | 45,54418 | 28,049181 | 7,483827 | 10,35443 | 7,292366325 | 25,64434 | 35,6706536 | 38,02266 | 186,6638 | 244,4609 | 77,1731731 |
| Upregulated | ENSMUSG00000116114 | 0 | 15,956459 | 9,0774684 | 4,81783712 | 2,544425 | 18,57756906 | 13,6307179 | 22,0000684 | 20,287434 | 14,0230546 | 11,0411331 | 3,503399 | 2,00351293 | 0,935478 | 0,796495 | 0 | 12,40855 | 0,990851489 | 8,555098 | 66,66563 | 93,88717 | 18,844612 |
| Upregulated | ENSMUSG00000031442 | 10,57475352 | 40,4230295 | 9,0774684 | 10,5992417 | 22,899825 | 13,26969219 | 12,4948247 | 13,4444862 | 9,60983717 | 23,7313232 | 40,8521924 | 2,627549 | 16,0281034 | 4,677392 | 11,94742 | 3,646183163 | 19,02645 | 1,981702978 | 22,8136 | 149,331 | 191,3173 | 35,8944991 |
| Upregulated | ENSMUSG00000030674 | 0 | 14,8926951 | 18,1549368 | 6,74497197 | 0 | 5,307876874 | 6,81535895 | 8,55558215 | 9,60983717 | 5,39348255 | 1,10411331 | 3,503399 | 12,0210776 | 0 | 6,37196 | 0,911545791 | 5,790658 | 4,954257444 | 6,653965 | 72,66553 | 93,00144 | 19,7419745 |
| Upregulated | ENSMUSG00000034675 | 288,6907712 | 153,182006 | 367,63747 | 238,964721 | 62,3384124 | 107,4845067 | 103,366277 | 182,111677 | 93,9628523 | 251,336287 | 170,03345 | 211,0798 | 267,468976 | 182,4183 | 89,20743 | 81,12757537 | 220,8722 | 187,2709314 | 390,6828 | 2067,301 | 2416,266 | 580,593523 |
| Upregulated | ENSMUSG00000045392 | 3,172426057 | 0 | 4,5387342 | 0,96356742 | 0 | 1,326969219 | 1,13589316 | 3,66667807 | 6,40655811 | 1,07869651 | 0 | 15,76529 | 23,0403987 | 6,548349 | 2,389485 | 8,203912116 | 6,617895 | 8,917663399 | 9,505665 | 133,9979 | 156,7739 | 20,639337 |
| Upregulated | ENSMUSG00000062345 | 68,7358979 | 9,57387541 | 87,1436966 | 338,212166 | 20,3554 | 41,13604578 | 17,0383974 | 146,667123 | 97,1661314 | 1,07869651 | 0 | 191,8111 | 1436,51877 | 138,4508 | 43,01073 | 47,40038111 | 309,3866 | 269,511605 | 1112,163 | 25743,6 | 32138,64 | 1079,52706 |
| Upregulated | ENSMUSG00000021702 | 260,1389367 | 370,189849 | 595,481927 | 876,846356 | 225,181612 | 119,4272297 | 89,7355595 | 548,779484 | 100,36941 | 228,68366 | 323,505199 | 305,6715 | 726,273437 | 405,0621 | 104,3408 | 137,6434144 | 540,1857 | 616,309626 | 719,5788 | 7203,888 | 9065,426 | 884,799403 |
| Upregulated | ENSMUSG00000086712 | 27,49435916 | 29,7853902 | 68,081013 | 33,7248598 | 8,90548749 | 47,77089187 | 23,8537563 | 35,4445546 | 21,3551937 | 77,6661487 | 25,3946061 | 0,87585 | 5,00878232 | 11,22574 | 10,35443 | 15,49627844 | 38,88013 | 17,8353268 | 51,33059 | 219,9966 | 347,2054 | 60,123286 |
| Upregulated | ENSMUSG00000038903 | 4,229901409 | 14,8926951 | 11,8007089 | 10,5992417 | 0 | 29,19332281 | 11,3589316 | 9,77780817 | 0 | 14,0230546 | 2,20822662 | 4,379248 | 0 | 0 | 0 | 0 | 0 | 3,963405955 | 10,45623 | 73,99885 | 129,3163 | 5,38417486 |
| Upregulated | ENSMUSG00000029378 | 6,344852114 | 2,12752787 | 6,35422788 | 9,63567424 | 3,8166375 | 17,25059984 | 0 | 0 | 0 | 11,8656616 | 7,72879316 | 0 | 2,00351293 | 1,870957 | 5,575465 | 15,49627844 | 8,272368 | 0 | 0 | 57,99909 | 78,82979 | 9,87098725 |
| Upregulated | ENSMUSG00000029136 | 26,43688381 | 23,4028065 | 19,0626836 | 47,2148038 | 26,7164625 | 5,307876874 | 9,08714527 | 6,11113011 | 18,1519147 | 12,9443581 | 0 | 7,006798 | 25,0439116 | 37,41914 | 21,50536 | 16,40782423 | 17,37197 | 24,77128722 | 32,31926 | 35,33278 | 61,11523 | 38,5865865 |

| | | | | | | | | | | | | | | | | | | | | | | | |
|---|---|---|---|---|---|---|---|---|---|---|---|---|---|---|---|---|---|---|---|---|---|---|---|
| Upregulated | ENSMUSG00000089665 | 39,12658804 | 29,7853902 | 11,8007089 | 30,8341576 | 6,36106249 | 7,961815312 | 10,2230384 | 13,4444862 | 11,7453565 | 6,47217906 | 6,62467985 | 18,39284 | 55,0966056 | 73,90279 | 28,67382 | 33,72719425 | 31,435 | 28,73469318 | 32,31926 | 81,33206 | 68,20106 | 81,6599854 |
| Upregulated | ENSMUSG00000092344 | 7,402327466 | 10,6376393 | 17,24719 | 19,2713485 | 5,08884999 | 3,980907656 | 6,81535895 | 4,88890409 | 10,6775969 | 2,15739302 | 2,20822662 | 8,758497 | 20,0351293 | 14,96765 | 11,94742 | 13,67318686 | 15,7175 | 13,87192084 | 8,555098 | 22,66631 | 23,91466 | 17,0498871 |
| Upregulated | ENSMUSG00000045506 | 0 | 0 | 0,90774684 | 1,92713485 | 0 | 7,961815312 | 5,67946579 | 1,22222602 | 0 | 4,31478604 | 2,20822662 | 7,006798 | 7,01229525 | 15,90313 | 1,59299 | 2,734637372 | 5,790658 | 0 | 1,901133 | 17,33306 | 20,37174 | 1,79472495 |
| Upregulated | ENSMUSG00000110834 | 24,3219331 | 31,912918 | 19,9704305 | 33,7248598 | 24,1720375 | 18,57756906 | 36,3485811 | 23,2222944 | 27,7617518 | 15,1017511 | 13,2493597 | 39,41324 | 30,0526939 | 62,67705 | 36,63877 | 19,1424616 | 29,78053 | 28,73469318 | 41,82493 | 89,33194 | 79,71552 | 34,9971366 |
| Upregulated | ENSMUSG00000026166 | 2,114950705 | 0 | 0 | 21,1984833 | 0 | 0 | 0 | 0 | 0 | 0 | 0 | 0 | 14,0245905 | 27,12887 | 0,796495 | 20,05400739 | 6,617895 | 1,981702978 | 10,45623 | 39,33272 | 13,28592 | 7,17889982 |
| Upregulated | ENSMUSG00000097075 | 11,63222888 | 2,12752787 | 6,35422788 | 11,5628091 | 3,8166375 | 0 | 4,54357263 | 2,44445204 | 4,27103874 | 2,15739302 | 3,31233993 | 1,751699 | 1,00175646 | 20,58052 | 0,796495 | 8,203912116 | 2,481711 | 3,963405955 | 7,604532 | 23,33297 | 17,71456 | 8,97362477 |
| Upregulated | ENSMUSG00000039079 | 7,402327466 | 6,3825836 | 14,5239494 | 4,81783712 | 0 | 0 | 18,1742905 | 2,44445204 | 3,20327906 | 1,07869651 | 2,20822662 | 0,87585 | 0 | 0 | 1,59299 | 2,734637372 | 23,16263 | 3,963405955 | 13,30793 | 17,99972 | 30,11475 | 11,6657122 |
| Upregulated | ENSMUSG00000118097 | 12,68970423 | 18,0839869 | 18,1549368 | 14,4535114 | 11,4499125 | 14,5966614 | 30,6691153 | 4,88890409 | 8,54207749 | 14,0230546 | 5,52056654 | 20,14454 | 13,022834 | 14,03218 | 12,74392 | 20,96555318 | 44,67079 | 13,87192084 | 26,61586 | 43,33266 | 35,42912 | 13,4604372 |
| Upregulated | ENSMUSG00000069272 | 1,057475352 | 14,8926951 | 2,72324052 | 2,89070227 | 3,8166375 | 0 | 10,2230384 | 0 | 5,33879843 | 0 | 0 | 0 | 0 | 0,935478 | 0 | 0 | 8,272368 | 0 | 14,2585 | 17,99972 | 15,05738 | 8,0762623 |
| Upregulated | ENSMUSG00000028125 | 24,3219331 | 44,6780852 | 7,26197472 | 4,81783712 | 27,988675 | 10,61575375 | 40,8921537 | 7,33335613 | 16,0163953 | 8,62957207 | 5,52056654 | 21,89624 | 11,0193211 | 0 | 3,18598 | 11,85009528 | 28,95329 | 20,80788127 | 3,802266 | 51,99919 | 81,48698 | 26,9208743 |
| Upregulated | ENSMUSG00000032221 | 11,63222888 | 5,31881967 | 3,63098736 | 1,92713485 | 12,722125 | 6,634846093 | 5,67946579 | 0 | 12,8131162 | 5,39348255 | 3,31233993 | 0,87585 | 0 | 0 | 0,796495 | 0 | 9,099605 | 14,86277233 | 19,9619 | 18,66638 | 18,60029 | 2,69208743 |
| Upregulated | ENSMUSG00000104876 | 0 | 1,06376393 | 0 | 0 | 0 | 1,326969219 | 0 | 0 | 3,20327906 | 0 | 0 | 5,255098 | 0 | 0 | 11,15093 | 3,646183163 | 1,654474 | 19,81702978 | 40,87436 | 14,66644 | 19,48602 | 11,6657122 |
| Upregulated | ENSMUSG00000021697 | 2,114950705 | 6,3825836 | 16,3394431 | 6,74497197 | 11,4499125 | 0 | 14,7666111 | 3,66667807 | 2,13551937 | 1,07869651 | 2,20822662 | 1,751699 | 0 | 13,0967 | 7,168455 | 0 | 2,481711 | 36,66150509 | 34,22039 | 47,99925 | 23,02893 | 18,844612 |
| Upregulated | ENSMUSG00000034883 | 1,057475352 | 4,25505574 | 3,63098736 | 3,8542697 | 0 | 0 | 0 | 2,44445204 | 5,33879843 | 5,39348255 | 0 | 3,503399 | 6,01053879 | 2,806435 | 1,59299 | 0 | 13,23579 | 24,77128722 | 23,76416 | 27,33291 | 11,51446 | 12,5630747 |
| Upregulated | ENSMUSG00000044229 | 44,4139648 | 41,4867934 | 40,8486078 | 70,340422 | 53,4329249 | 66,34846093 | 74,9689484 | 75,7780133 | 87,5562942 | 30,2035023 | 55,2056654 | 64,81288 | 83,1457866 | 61,74157 | 67,70207 | 80,21602958 | 103,4046 | 193,2160403 | 194,8661 | 239,9963 | 209,9175 | 160,627883 |
| Upregulated | ENSMUSG00000060639 | 13,74717958 | 7,44634754 | 17,24719 | 8,67210682 | 12,722125 | 1,326969219 | 5,67946579 | 3,66667807 | 4,27103874 | 2,15739302 | 11,0411331 | 3,503399 | 7,01229525 | 24,32244 | 12,74392 | 14,58473265 | 9,926842 | 20,80788127 | 53,23172 | 51,99919 | 52,25795 | 3,58944991 |
| Upregulated | ENSMUSG00000042367 | 14,80465493 | 10,6376393 | 5,44648104 | 5,78140454 | 0 | 0 | 0 | 0 | 1,06775969 | 8,62957207 | 3,31233993 | 21,89624 | 13,022834 | 7,483827 | 2,389485 | 7,292366325 | 18,19921 | 8,917663399 | 89,35325 | 76,66547 | 75,28688 | 16,1525246 |
| Upregulated | ENSMUSG00000062380 | 49,70134156 | 74,4634754 | 73,527494 | 69,3768545 | 49,6162874 | 76,96421468 | 29,5332221 | 57,444623 | 83,2852555 | 20,4952337 | 36,4357392 | 83,20572 | 68,1194396 | 104,7736 | 19,11588 | 38,28492321 | 38,88013 | 83,23152506 | 217,6797 | 280,6623 | 232,9465 | 37,6892241 |
| Upregulated | ENSMUSG00000063171 | 215,7249719 | 210,625259 | 286,848001 | 293,888064 | 152,6655 | 359,6086582 | 470,259768 | 253,000787 | 182,586906 | 173,670138 | 155,679977 | 144,5152 | 301,528696 | 205,8052 | 263,6398 | 274,375283 | 377,22 | 266,5390505 | 460,0742 | 1001,318 | 680,2391 | 249,466769 |
| Upregulated | ENSMUSG00000098019 | 5,287376762 | 7,44634754 | 2,72324052 | 7,70853939 | 10,1777 | 9,28878453 | 20,4460768 | 3,66667807 | 5,33879843 | 4,31478604 | 8,83290647 | 4,379248 | 18,0316164 | 19,64505 | 8,761444 | 10,0270037 | 24,81711 | 2,972554466 | 21,86303 | 37,99941 | 31,00048 | 7,17889982 |
| Upregulated | ENSMUSG00000031371 | 231,5871022 | 312,746597 | 209,68952 | 387,354104 | 231,542675 | 298,5680742 | 268,070785 | 178,444999 | 213,551937 | 270,752824 | 187,699262 | 481,7173 | 414,727176 | 516,3841 | 456,3916 | 456,6844411 | 791,6657 | 659,9070915 | 591,2524 | 929,9855 | 627,9812 | 394,83949 |
| Upregulated | ENSMUSG00000081600 | 28,55183451 | 39,3592656 | 44,4795952 | 76,1218265 | 63,6106249 | 55,73270718 | 70,4253758 | 39,1112327 | 58,7267827 | 56,0922185 | 35,3316259 | 84,95742 | 66,1159267 | 95,4188 | 86,02145 | 81,12757537 | 133,1851 | 97,1034459 | 97,90835 | 205,3301 | 131,0877 | 74,4810856 |
| Upregulated | ENSMUSG00000113611 | 8,459802819 | 20,2115147 | 10,8929621 | 21,1984833 | 8,90548749 | 17,25059984 | 4,54357263 | 7,33335613 | 17,084155 | 9,70826858 | 6,62467985 | 28,90304 | 14,0245905 | 26,19339 | 15,9299 | 27,34637372 | 46,32526 | 25,76213871 | 29,46756 | 47,99925 | 29,22902 | 10,7683497 |
| Upregulated | ENSMUSG00000083044 | 17,97708099 | 37,2317377 | 15,4316963 | 46,2512364 | 26,7164625 | 37,15513812 | 28,397329 | 12,2222602 | 24,5584728 | 14,0230546 | 17,6658129 | 43,79248 | 27,0474245 | 50,51583 | 31,0633 | 33,72719425 | 98,44118 | 34,67980211 | 61,78682 | 84,66535 | 84,14416 | 28,7155993 |
| Upregulated | ENSMUSG00000097989 | 14,80465493 | 31,912918 | 21,7859242 | 33,7248598 | 39,4385875 | 46,44392265 | 38,6203674 | 26,8889725 | 29,8972712 | 17,2591441 | 19,8740396 | 55,17853 | 26,0456681 | 64,54801 | 52,56867 | 54,69274744 | 100,0957 | 28,73469318 | 74,14419 | 111,3316 | 88,5728 | 21,5366995 |
| Upregulated | ENSMUSG00000059658 | 5,287376762 | 29,7853902 | 8,16972156 | 18,3077811 | 10,1777 | 19,90453828 | 29,5332221 | 8,55558215 | 13,8808759 | 7,55087556 | 12,1452464 | 12,2619 | 21,0368858 | 24,32244 | 23,09835 | 57,42738481 | 52,94316 | 16,84447531 | 50,38002 | 50,66588 | 68,20106 | 8,97362477 |
| Upregulated | ENSMUSG00000098113 | 5,287376762 | 15,956459 | 9,98521524 | 17,3442136 | 24,1720375 | 23,88544594 | 28,397329 | 18,3333903 | 10,6775969 | 4,31478604 | 12,1452464 | 16,64114 | 15,026347 | 21,516 | 19,91237 | 27,34637372 | 47,1525 | 15,85362382 | 35,17096 | 59,99906 | 41,62922 | 20,639337 |
| Upregulated | ENSMUSG00000081721 | 19,03455634 | 28,7216262 | 22,693671 | 25,052753 | 34,3497375 | 49,09786109 | 32,9409016 | 23,2222944 | 29,8972712 | 12,9443581 | 27,6028327 | 28,90304 | 24,0421551 | 37,41914 | 51,77217 | 67,45438851 | 88,51434 | 38,64320806 | 46,57776 | 92,66522 | 79,71552 | 27,8182368 |
| Upregulated | ENSMUSG00000069376 | 44,4139648 | 67,0171278 | 68,081013 | 98,2838772 | 66,1550499 | 71,65633781 | 74,9689484 | 63,5557531 | 54,455744 | 38,8330743 | 48,5809856 | 74,44722 | 92,1615947 | 99,16071 | 94,7829 | 85,68530432 | 123,2583 | 62,4236438 | 120,7219 | 191,997 | 171,8312 | 85,2494354 |
| Upregulated | ENSMUSG00000070443 | 50,75881691 | 70,2084196 | 59,0035446 | 99,2474447 | 95,4159374 | 54,40573796 | 92,0073458 | 48,8890409 | 72,6076586 | 55,013522 | 51,8933255 | 77,07477 | 106,186185 | 128,1605 | 123,4567 | 129,4395023 | 167,1018 | 76,29556464 | 182,5088 | 219,3299 | 211,689 | 64,6100984 |
| Upregulated | ENSMUSG00000105558 | 125,8395669 | 129,7792 | 107,114127 | 118,518793 | 96,6881499 | 116,7732912 | 132,8995 | 80,6669174 | 60,8623021 | 81,9809347 | 72,8714784 | 102,4744 | 102,179159 | 119,7412 | 136,2006 | 138,5549602 | 275,4699 | 122,8655846 | 230,0371 | 371,9942 | 275,4614 | 97,81251 |
| Upregulated | ENSMUSG00000063524 | 1488,925296 | 1754,14673 | 1613,97388 | 1900,15496 | 1279,84577 | 1252,658942 | 1542,54291 | 1106,11455 | 1224,72036 | 1347,29194 | 1016,88836 | 1572,15 | 1478,59254 | 1981,343 | 2036,638 | 2340,84959 | 3314,738 | 2078,806424 | 2798,468 | 5447,248 | 3814,831 | 1672,68366 |
| Upregulated | ENSMUSG00000059040 | 3631,37036 | 3765,72433 | 3847,03111 | 4317,74563 | 2936,26645 | 2924,640158 | 3572,38398 | 2644,89711 | 2802,86917 | 3345,03787 | 2310,90916 | 3940,448 | 3645,39177 | 4881,326 | 5013,936 | 5806,546686 | 7723,083 | 5221,787346 | 7302,252 | 12551,14 | 9605,72 | 4415,02339 |
| Upregulated | ENSMUSG00000064193 | 4024,751191 | 4379,51612 | 4372,61653 | 4936,35591 | 3490,9511 | 3351,924246 | 4345,92722 | 3128,89862 | 3142,41675 | 3890,85831 | 2603,49918 | 4260,133 | 4073,14178 | 5214,357 | 5618,475 | 6558,571964 | 8614,844 | 5622,091348 | 8054,15 | 14093,78 | 10965,31 | 5019,8457 |
| Upregulated | ENSMUSG00000070025 | 75,08075001 | 74,4634754 | 68,9887598 | 98,2838772 | 99,2325749 | 69,00239937 | 102,230384 | 72,1113353 | 79,0142167 | 70,1152731 | 49,6850989 | 91,08837 | 118,207263 | 143,1282 | 150,5375 | 175,0167918 | 265,543 | 95,12174293 | 158,7446 | 328,6615 | 202,8317 | 98,7098725 |
| Upregulated | ENSMUSG00000070490 | 86,71297889 | 112,758977 | 105,298633 | 125,263765 | 123,404612 | 131,3699526 | 130,627713 | 101,44476 | 107,843728 | 75,5087556 | 71,7673651 | 141,8877 | 128,224827 | 204,8698 | 172,8394 | 193,2477076 | 275,4699 | 168,4447531 | 253,8013 | 396,6605 | 303,8047 | 152,551621 |
| Upregulated | ENSMUSG00000097979 | 83,54055283 | 91,4836983 | 68,9887598 | 80,9396636 | 108,138062 | 84,92602999 | 104,502171 | 77,0002394 | 89,6918136 | 58,2496115 | 79,4961582 | 90,21252 | 110,193211 | 188,9666 | 146,5551 | 164,9897881 | 241,5532 | 137,7283569 | 195,8167 | 309,3285 | 232,9465 | 109,478222 |
| Upregulated | ENSMUSG00000007682 | 24,3219331 | 31,912918 | 52,6493167 | 28,9070227 | 53,4329249 | 55,73270718 | 35,2126879 | 37,8890067 | 117,453565 | 62,5643975 | 98,2660845 | 129,6258 | 238,418039 | 64,54801 | 260,4538 | 219,6825355 | 198,5368 | 134,7558025 | 88,40268 | 49,99922 | 270,147 | 142,680634 |
| Upregulated | ENSMUSG00000024737 | 6,344852114 | 3,1912918 | 2,72324052 | 0 | 2,544425 | 18,57756906 | 2,27178632 | 3,66667807 | 19,2196743 | 0 | 18,7699262 | 10,5102 | 10,0175646 | 11,22574 | 23,89485 | 9,115457906 | 18,19921 | 7,92681191 | 2,851699 | 17,99972 | 26,57184 | 13,4604372 |
| Upregulated | ENSMUSG00000105651 | 1,057475352 | 0 | 0 | 0 | 5,08884999 | 7,961815312 | 1,13589316 | 1,22222602 | 0 | 5,39348255 | 2,20822662 | 15,76529 | 4,00702586 | 7,483827 | 11,15093 | 16,40782423 | 14,06303 | 6,935960422 | 20,91246 | 35,99944 | 19,48602 | 6,28153734 |
| Upregulated | ENSMUSG00000029860 | 1857,984194 | 823,353285 | 1510,49074 | 1184,22436 | 600,484299 | 1199,580174 | 998,450086 | 800,558044 | 1016,50722 | 960,039893 | 1037,86651 | 3161,817 | 1705,99126 | 3626,85 | 2220,628 | 3278,830209 | 3198,098 | 2498,927455 | 2648,278 | 7557,215 | 4469,384 | 1545,25819 |
| Upregulated | ENSMUSG00000053897 | 142,7591726 | 120,205325 | 85,328203 | 79,9760962 | 190,831875 | 155,2553986 | 94,2791321 | 178,444999 | 124,927883 | 153,174904 | 73,9755917 | 330,1953 | 315,553286 | 381,6752 | 147,3516 | 243,3827261 | 244,8621 | 191,2343373 | 295,6262 | 736,6552 | 482,7218 | 253,056219 |
| Upregulated | ENSMUSG00000106874 | 263,3113627 | 94,6749901 | 183,364862 | 122,373063 | 151,393287 | 220,2768903 | 244,217029 | 179,667225 | 185,790185 | 159,647083 | 295,902367 | 993,2136 | 826,449083 | 733,4151 | 600,5572 | 585,2123976 | 745,3404 | 455,7916849 | 703,4192 | 1087,316 | 874,2136 | 588,669785 |
| Upregulated | ENSMUSG00000005686 | 424,0476163 | 365,934793 | 285,940255 | 167,660732 | 235,359312 | 301,2220126 | 173,791653 | 308,000957 | 245,584728 | 376,465082 | 632,656926 | 1311,147 | 1262,21315 | 1202,09 | 899,2428 | 697,3325298 | 927,3325 | 591,5383388 | 1305,128 | 1867,304 | 1774,999 | 939,538514 |
| Upregulated | ENSMUSG00000086320 | 132,184419 | 104,248866 | 54,4648104 | 54,9233432 | 97,9603624 | 79,61815312 | 109,045743 | 74,5557873 | 139,876519 | 74,4300591 | 161,200543 | 329,3195 | 578,01348 | 400,3848 | 274,7908 | 556,9544781 | 378,8745 | 246,7220207 | 351,7096 | 761,3215 | 505,7507 | 386,763228 |
| Upregulated | ENSMUSG00000028896 | 240,046905 | 382,955016 | 416,6558 | 334,357896 | 293,881087 | 257,4320284 | 274,886144 | 295,778697 | 293,633914 | 324,687649 | 403,001358 | 1039,634 | 970,702014 | 406,9331 | 710,4735 | 605,266405 | 833,8547 | 759,9830919 | 785,1679 | 1337,979 | 1131,96 | 753,784481 |
| Upregulated | ENSMUSG00000033581 | 299,2655247 | 185,094925 | 362,190989 | 134,899439 | 123,404612 | 240,1814286 | 141,986645 | 204,111746 | 155,892914 | 141,309243 | 273,820101 | 528,1374 | 501,879989 | 238,547 | 302,6681 | 341,8296715 | 464,9071 | 298,2462981 | 504,7508 | 800,6542 | 601,4093 | 413,684102 |
| Upregulated | ENSMUSG00000047330 | 1437,109004 | 740,379698 | 1495,96679 | 587,776129 | 234,0871 | 203,0262904 | 277,157931 | 1096,33674 | 782,66785 | 440,108176 | 700,007838 | 1665,866 | 2694,72489 | 1522,959 | 988,4502 | 1110,262773 | 2743,117 | 2038,181512 | 2008,547 | 3811,274 | 3279,851 | 2455,18374 |
| Upregulated | ENSMUSG00000060459 | 44,4139648 | 6,3825836 | 9,0774684 | 1,92713485 | 1,2722125 | 0 | 0 | 0 | 3,20327906 | 0 | 2,20822662 | 54,30268 | 286,502349 | 141,2572 | 17,52289 | 11,85009528 | 304,4232 | 89,17663399 | 237,6416 | 269,3291 | 375,5487 | 113,965035 |
| Upregulated | ENSMUSG00000022875 | 3,172426057 | 0 | 0 | 0 | 0 | 0 | 0 | 0 | 0 | 0 | 0 | 8,758497 | 18,0316164 | 3,741914 | 1,59299 | 0,911545791 | 28,95329 | 11,89021787 | 16,15963 | 31,33284 | 36,31485 | 17,0498871 |
| Upregulated | ENSMUSG00000074802 | 238,9894296 | 239,346885 | 403,039597 | 125,263765 | 212,459487 | 216,2959826 | 210,140234 | 389,890101 | 274,414239 | 297,720237 | 292,590027 | 736,5896 | 780,368286 | 535,0936 | 497,8093 | 829,5066695 | 1048,936 | 636,1266558 | 663,4954 | 1199,981 | 1418,051 | 743,016131 |
| Upregulated | ENSMUSG00000021950 | 380,6911268 | 621,238137 | 290,478989 | 173,442136 | 479,624112 | 69,00239937 | 72,6971621 | 260,334143 | 451,662347 | 59,328308 | 270,507761 | 749,7273 | 807,41571 | 986,9297 | 561,5289 | 1380,080327 | 1619,73 | 458,7642393 | 1066,536 | 2104,634 | 2737,785 | 1770,49617 |
| Upregulated | ENSMUSG00000046079 | 94,11530636 | 72,3359475 | 66,2655193 | 46,2512364 | 111,9547 | 33,17423047 | 38,6203674 | 37,8890067 | 19,2196743 | 21,5739302 | 43,060419 | 119,1156 | 145,254687 | 29,93531 | 121,0672 | 100,270037 | 200,1913 | 35,6706536 | 168,2503 | 259,3293 | 279,0043 | 168,704146 |
| Upregulated | ENSMUSG00000108814 | 0 | 0 | 0 | 2,89070227 | 2,544425 | 0 | 1,13589316 | 0 | 0 | 0 | 4,41645324 | 2,627549 | 15,026347 | 6,548349 | 10,35443 | 6,380820534 | 14,06303 | 4,954257444 | 5,703399 | 32,66616 | 23,02893 | 11,6657122 |
| Upregulated | ENSMUSG00000019139 | 659,8646198 | 293,598846 | 491,09104 | 440,350313 | 381,66375 | 403,3986425 | 336,224375 | 319,000992 | 453,797866 | 256,729769 | 451,582343 | 980,9517 | 1811,17569 | 922,3817 | 1061,728 | 802,1602958 | 1394,721 | 663,8704975 | 935,3574 | 2116,634 | 2285,178 | 1337,07009 |
| Upregulated | ENSMUSG00000068922 | 222,069824 | 186,158688 | 207,874026 | 234,146884 | 192,104087 | 155,2553986 | 186,286478 | 154,000479 | 121,724604 | 182,29971 | 144,638843 | 294,2855 | 290,509375 | 232,9341 | 313,0225 | 302,6332025 | 253,1345 | 393,3680411 | 313,6869 | 590,6574 | 518,1509 | 359,842353 |
| Upregulated | ENSMUSG00000073514 | 19,03455634 | 5,31881967 | 3,63098736 | 3,8542697 | 6,36106249 | 3,980907656 | 3,40767947 | 0 | 3,20327906 | 0 | 0 | 8,758497 | 30,0526939 | 0,935478 | 20,70887 | 10,0270037 | 7,445132 | 21,79873275 | 14,2585 | 50,66588 | 58,45805 | 9,87098725 |
| Upregulated | ENSMUSG00000022894 | 2073,709166 | 1388,21193 | 2716,88629 | 1507,01945 | 1180,6132 | 2476,124562 | 1508,46611 | 1920,11708 | 1900,61224 | 1586,76256 | 1453,01311 | 2814,105 | 4290,52294 | 5208,744 | 2561,528 | 3479,370283 | 3007,833 | 4014,930233 | 3654,928 | 8823,196 | 8737,707 | 3200,89196 |
| Upregulated | ENSMUSG00000033307 | 867,1297889 | 1022,27714 | 1004,87575 | 1507,01945 | 935,076187 | 927,5514838 | 1135,89316 | 817,669209 | 1149,97718 | 865,1146 | 853,479588 | 1503,834 | 2135,74478 | 2776,5 | 1997,609 | 2431,092624 | 2604,142 | 2090,696641 | 2174,896 | 3872,606 | 3905,175 | 1530,00302 |
| Upregulated | ENSMUSG00000037362 | 1744,834331 | 988,236695 | 1476,90411 | 2069,74283 | 402,01915 | 601,117056 | 807,620036 | 541,446128 | 918,27333 | 846,77676 | 908,685253 | 3346,622 | 2371,15755 | 1643,636 | 5175,624 | 4267,857392 | 4792,183 | 3170,724764 | 1896,38 | 10461,17 | 13610,1 | 2657,98766 |
| Upregulated | ENSMUSG00000000295 | 62,39104579 | 130,842964 | 118,007089 | 147,425816 | 114,499125 | 104,8305683 | 88,5996663 | 133,222636 | 115,318046 | 74,4300591 | 79,4961582 | 157,6529 | 144,252931 | 146,8701 | 173,6359 | 198,7169824 | 184,4738 | 190,2434859 | 205,3224 | 304,6619 | 356,9484 | 193,830295 |
| Upregulated | ENSMUSG00000083344 | 8,459802819 | 7,44634754 | 20,8781773 | 8,67210682 | 12,722125 | 10,61575375 | 10,2230384 | 11,0000342 | 5,33879843 | 4,31478604 | 7,72879316 | 17,51699 | 21,0368858 | 34,6127 | 30,26681 | 15,49627844 | 34,74395 | 13,87192084 | 22,8136 | 25,33294 | 49,60077 | 10,7683497 |
| Upregulated | ENSMUSG00000026271 | 23,26445775 | 9,57387541 | 33,5866331 | 0 | 7,63327499 | 0 | 3,40767947 | 7,33335613 | 0 | 0 | 0 | 39,41324 | 62,1089008 | 26,19339 | 65,31259 | 43,75419795 | 76,93303 | 20,80788127 | 49,42946 | 81,33206 | 93,00144 | 32,3050492 |
| Upregulated | ENSMUSG00000009687 | 1860,099145 | 2043,49052 | 2115,05014 | 2122,73904 | 1597,8989 | 1200,907143 | 1750,41136 | 787,113558 | 1186,28101 | 1461,63377 | 1363,57994 | 3267,795 | 3883,80981 | 3299,432 | 4028,671 | 3946,081728 | 4832,718 | 2652,509436 | 3382,116 | 2907,288 | 5701,431 | 2703,75314 |
| Upregulated | ENSMUSG00000032548 | 1311,269437 | 524,435619 | 678,086889 | 481,783712 | 164,115412 | 273,355659 | 257,847747 | 402,112361 | 255,194565 | 206,031033 | 352,212146 | 2477,779 | 1839,22487 | 773,6406 | 692,9506 | 583,389306 | 1997,777 | 602,4377052 | 1735,734 | 1705,973 | 3082,333 | 1410,65381 |
| Upregulated | ENSMUSG00000030790 | 942,2105389 | 1158,43892 | 1132,86806 | 1595,66765 | 820,577062 | 1973,203228 | 1118,85476 | 1201,44818 | 872,359663 | 793,920631 | 1343,7059 | 3819,581 | 2990,24305 | 1939,247 | 1643,169 | 996,3195492 | 2840,731 | 1720,118185 | 5442,944 | 3979,938 | 4309,953 | 2185,975 |
| Upregulated | ENSMUSG00000006014 | 3708,566061 | 1119,07966 | 1925,33105 | 1999,4024 | 192,104087 | 99,5226914 | 260,119533 | 457,112532 | 227,432813 | 215,739302 | 338,962786 | 7836,227 | 19061,422 | 17295,12 | 5753,083 | 5822,954511 | 11920,48 | 3557,156845 | 9403,004 | 10480,5 | 16737,6 | 4687,82158 |
| Upregulated | ENSMUSG00000023367 | 256,9665106 | 71,2721836 | 315,8959 | 372,900593 | 64,8828374 | 196,3914444 | 147,666111 | 195,556163 | 123,860124 | 133,758367 | 447,16589 | 879,3531 | 2765,8496 | 2137,568 | 638,7889 | 867,7915927 | 1926,635 | 2054,035136 | 1454,367 | 2571,293 | 3249,736 | 1217,72088 |
| Upregulated | ENSMUSG00000029810 | 731,7729438 | 175,521049 | 618,175598 | 650,408011 | 189,559662 | 538,7495028 | 372,572956 | 309,223184 | 319,260146 | 276,146306 | 921,934613 | 1873,442 | 6052,61256 | 3997,299 | 1287,136 | 1638,047786 | 3514,929 | 3350,068884 | 2403,983 | 3849,273 | 4677,53 | 1867,41132 |
| Upregulated | ENSMUSG00000024810 | 366,9439473 | 559,539829 | 1179,16315 | 687,023573 | 215,003912 | 139,331768 | 237,40167 | 304,334279 | 300,040472 | 393,724226 | 814,835622 | 7717,987 | 34159,8954 | 9572,75 | 5898,045 | 6127,410805 | 12911,51 | 8540,148982 | 8848,823 | 17609,06 | 19636,59 | 7526,1791 |
| Upregulated | ENSMUSG00000038067 | 103,6325845 | 15,956459 | 117,099342 | 43,3605341 | 16,5387625 | 5,307876874 | 11,3589316 | 52,5557189 | 40,5748681 | 0 | 24,2904928 | 3574,343 | 8991,76603 | 3702,623 | 696,1366 | 1453,915536 | 2270,765 | 845,19632 | 2202,463 | 6017,239 | 8385,187 | 964,664663 |
| Upregulated | ENSMUSG00000029380 | 21063,85154 | 17510,6181 | 41674,6574 | 17283,5089 | 7411,91002 | 8199,342802 | 6457,5526 | 14137,4884 | 13600,0551 | 11450,3634 | 21943,1479 | 85600,29 | 142718,24 | 106913 | 69980,84 | 61548,48333 | 63350,62 | 68435,13978 | 66718,36 | 169028 | 239014,6 | 55640,9604 |
| Upregulated | ENSMUSG00000047907 | 178,7133345 | 113,822741 | 177,010634 | 175,369271 | 27,988675 | 173,8329676 | 68,1535895 | 201,667294 | 147,350837 | 146,702725 | 110,411331 | 292,5338 | 648,136433 | 489,2552 | 350,4578 | 354,5913126 | 274,6426 | 174,389862 | 484,7889 | 671,3229 | 843,2131 | 191,138208 |
| Upregulated | ENSMUSG00000017210 | 744,462648 | 465,928603 | 722,566485 | 730,384107 | 681,905899 | 740,448824 | 901,899168 | 597,668525 | 537,083122 | 883,452441 | 542,119635 | 1200,79 | 2032,56387 | 1138,477 | 810,8319 | 794,8679294 | 1598,222 | 729,2666958 | 1279,463 | 2460,628 | 2224,063 | 952,998951 |
| Upregulated | ENSMUSG00000030587 | 11,63222888 | 22,3390426 | 7,26197472 | 7,70853939 | 5,08884999 | 9,28878453 | 15,9025042 | 9,77780817 | 2,13551937 | 9,70826858 | 12,1452464 | 15,76529 | 51,0895797 | 34,6127 | 14,33691 | 12,76164107 | 28,95329 | 14,86277233 | 34,22039 | 56,66578 | 64,65815 | 22,4340619 |
| Upregulated | ENSMUSG00000020553 | 2,114950705 | 4,25505574 | 5,44648104 | 0,96356742 | 15,26655 | 22,55847672 | 0 | 4,88890409 | 1,06775969 | 2,15739302 | 0 | 0 | 21,0368858 | 7,483827 | 0 | 0 | 11,58132 | 6,935960422 | 24,71473 | 23,99963 | 20,37174 | 24,2287869 |
| Upregulated | ENSMUSG00000019178 | 4,229901409 | 0 | 2,72324052 | 1,92713485 | 12,722125 | 30,52029203 | 4,54357263 | 2,44445204 | 10,6775969 | 3,23608953 | 3,31233993 | 0 | 8,01405172 | 2,806435 | 1,59299 | 0 | 15,7175 | 16,84447531 | 12,35736 | 20,66634 | 28,3433 | 17,9472495 |
| Upregulated | ENSMUSG00000075225 | 10,57475352 | 14,8926951 | 12,7084558 | 9,63567424 | 17,810975 | 26,53938437 | 3,40767947 | 6,11113011 | 4,27103874 | 1,07869651 | 7,72879316 | 0 | 13,022834 | 14,03218 | 1,59299 | 3,646183163 | 4,963421 | 2,972554466 | 10,45623 | 15,99975 | 36,31485 | 29,6129618 |
| Upregulated | ENSMUSG00000074899 | 2,114950705 | 9,57387541 | 0,90774684 | 2,89070227 | 7,63327499 | 15,92363062 | 10,2230384 | 6,11113011 | 2,13551937 | 0 | 11,0411331 | 0,87585 | 6,01053879 | 0,935478 | 1,59299 | 0 | 16,54474 | 8,917663399 | 5,703399 | 21,99966 | 40,74349 | 42,1760364 |
| Upregulated | ENSMUSG00000034738 | 10,57475352 | 10,6376393 | 9,98521524 | 4,81783712 | 11,4499125 | 37,15513812 | 15,9025042 | 8,55558215 | 21,3551937 | 12,9443581 | 26,4987194 | 0,87585 | 22,0386422 | 0 | 8,761444 | 18,23091581 | 17,37197 | 11,89021787 | 0 | 41,99934 | 62,88669 | 49,3549363 |
| Upregulated | ENSMUSG00000031461 | 12,68970423 | 9,57387541 | 1,81549368 | 18,3077811 | 0 | 2,653938437 | 10,2230384 | 8,55558215 | 4,27103874 | 0 | 23,1863795 | 45,54418 | 7,01229525 | 6,548349 | 18,31938 | 6,380820534 | 8,272368 | 19,81702978 | 7,604532 | 34,66613 | 85,02989 | 34,9971366 |
| Upregulated | ENSMUSG00000022126 | 12,68970423 | 48,933141 | 7,26197472 | 0 | 12,722125 | 9,28878453 | 0 | 7,33335613 | 13,8808759 | 30,2035023 | 19,8740396 | 35,03399 | 28,049181 | 10,29026 | 5,575465 | 5,469274744 | 33,91671 | 32,69809913 | 45,62719 | 34,66613 | 133,7449 | 65,5074609 |
| Upregulated | ENSMUSG00000026875 | 53,93124297 | 32,976682 | 59,0035446 | 27,9434553 | 19,0831875 | 47,77089187 | 35,2126879 | 51,3334929 | 30,9650309 | 16,1804476 | 70,6632518 | 77,95062 | 42,0737715 | 29,93531 | 26,28433 | 48,3119269 | 71,96961 | 71,34130719 | 93,15552 | 82,66538 | 183,3457 | 93,3256977 |
| Upregulated | ENSMUSG00000097028 | 9,517278171 | 9,57387541 | 5,44648104 | 0 | 0 | 3,980907656 | 12,4948247 | 24,4445204 | 14,9486356 | 8,62957207 | 23,1863795 | 46,42003 | 9,01580818 | 2,806435 | 19,11588 | 13,67318686 | 55,42487 | 47,56087146 | 53,23172 | 54,66581 | 155,8881 | 30,5103242 |
| Upregulated | ENSMUSG00000090231 | 309,8402782 | 305,300249 | 531,939648 | 233,183317 | 146,304437 | 281,3174743 | 355,534559 | 239,5563 | 395,071084 | 503,75127 | 435,020644 | 818,9195 | 964,691475 | 512,6422 | 366,3877 | 596,1509471 | 1354,187 | 460,7459423 | 1039,92 | 1601,308 | 3525,198 | 909,02819 |
| Upregulated | ENSMUSG00000015745 | 6,344852114 | 3,1912918 | 11,8007089 | 8,67210682 | 6,36106249 | 7,961815312 | 0 | 2,44445204 | 6,40655811 | 1,07869651 | 18,7699262 | 7,006798 | 9,01580818 | 3,741914 | 5,575465 | 1,823091581 | 34,74395 | 13,87192084 | 28,51699 | 39,33272 | 100,973 | 29,6129618 |
| Upregulated | ENSMUSG00000033576 | 4,229901409 | 5,31881967 | 14,5239494 | 22,1620508 | 3,8166375 | 25,21241515 | 5,67946579 | 17,1111643 | 0 | 18,3378407 | 2,20822662 | 7,006798 | 16,0281034 | 14,96765 | 4,77897 | 4,557728953 | 20,68092 | 6,935960422 | 27,56643 | 70,66556 | 115,1446 | 37,6892241 |

| | | | | | | | | | | | | | | | | | | | | | | | |
|---|---|---|---|---|---|---|---|---|---|---|---|---|---|---|---|---|---|---|---|---|---|---|---|
| Upregulated | ENSMUSG00000031538 | 3126,954617 | 2020,08771 | 4426,17359 | 2081,30564 | 957,976012 | 1260,620758 | 1271,06444 | 2212,2291 | 2547,67461 | 3610,39722 | 3855,56367 | 2489,165 | 3382,93158 | 1833,538 | 1458,382 | 1661,747976 | 4536,567 | 3473,92532 | 5987,618 | 19146,37 | 29700,23 | 7632,06787 |
| Upregulated | ENSMUSG00000009621 | 61,33357043 | 89,3561705 | 89,8669372 | 56,850478 | 39,4385875 | 69,00239937 | 101,094491 | 94,1114037 | 76,8786974 | 115,420526 | 104,890764 | 100,7227 | 93,1633512 | 75,77375 | 66,90558 | 87,5083959 | 79,41474 | 92,14918846 | 109,3151 | 293,3288 | 429,5781 | 201,009195 |
| Upregulated | ENSMUSG00000023064 | 38,06911268 | 31,912918 | 41,7563546 | 62,6318826 | 5,08884999 | 18,57756906 | 0 | 9,77780817 | 16,0163953 | 4,31478604 | 1,10411331 | 5,255098 | 17,0298599 | 0 | 7,168455 | 27,34637372 | 63,69724 | 20,80788127 | 11,4068 | 150,6643 | 290,5188 | 113,067672 |
| Upregulated | ENSMUSG00000090667 | 0 | 0 | 8,16972156 | 0 | 0 | 0 | 0 | 2,44445204 | 0 | 2,15739302 | 0 | 0,87585 | 8,01405172 | 0 | 0 | 0 | 0 | 0 | 6,653965 | 57,99909 | 120,459 | 68,1995483 |
| Upregulated | ENSMUSG00000015134 | 1,057475352 | 1,06376393 | 25,4169115 | 26,0163204 | 0 | 0 | 17,0383974 | 62,3335271 | 9,60983717 | 23,7313232 | 1,10411331 | 116,488 | 166,291573 | 154,3539 | 37,43526 | 36,46183163 | 71,96961 | 70,35045571 | 58,93512 | 575,991 | 1226,733 | 466,628488 |
| Upregulated | ENSMUSG00000057729 | 4,229901409 | 1,06376393 | 1,81549368 | 4,81783712 | 0 | 1,326969219 | 0 | 0 | 0 | 0 | 0 | 0,87585 | 13,022834 | 9,354784 | 5,575465 | 0 | 7,445132 | 1,981702978 | 12,35736 | 31,9995 | 72,6297 | 34,9971366 |
| Upregulated | ENSMUSG00000021069 | 781,4742854 | 711,658072 | 321,342381 | 449,02242 | 582,673324 | 291,9332281 | 449,813691 | 398,445683 | 168,70603 | 209,267123 | 284,861234 | 452,8143 | 499,876476 | 309,6433 | 179,2114 | 197,8054366 | 375,5655 | 315,0907734 | 469,5798 | 1101,983 | 2090,318 | 868,646878 |
| Upregulated | ENSMUSG00000020810 | 2088,513821 | 1062,70017 | 1921,70006 | 1467,51319 | 218,82055 | 287,9523204 | 127,220034 | 1345,67085 | 634,249253 | 720,569268 | 1031,24183 | 784,7613 | 2585,53344 | 527,6098 | 253,2854 | 226,9749019 | 1977,923 | 1555,636837 | 2929,646 | 3079,952 | 8995,454 | 4548,7304 |
| Upregulated | ENSMUSG00000054404 | 1184,372395 | 706,339252 | 944,96446 | 606,08391 | 325,6864 | 457,8043804 | 398,698499 | 740,668969 | 336,344301 | 560,922185 | 423,979511 | 357,3467 | 1098,92684 | 275,0306 | 497,0128 | 288,0484698 | 709,7692 | 476,5995661 | 1354,557 | 1699,307 | 3503,054 | 1711,27024 |
| Upregulated | ENSMUSG00000027931 | 113,1498627 | 61,6983082 | 142,516254 | 86,7210682 | 55,9773499 | 75,63724546 | 36,3485811 | 59,8890751 | 109,979248 | 87,3744172 | 60,726232 | 107,7295 | 167,29333 | 29,93531 | 21,50536 | 9,115457906 | 46,32526 | 49,54257444 | 136,8816 | 239,3296 | 476,5217 | 175,883046 |
| Upregulated | ENSMUSG00000054196 | 3031,781835 | 2662,60113 | 2648,80528 | 2806,87191 | 1937,57964 | 1492,840371 | 964,373291 | 3480,89971 | 1428,66246 | 2043,05119 | 2719,43108 | 1273,485 | 6648,65766 | 1194,606 | 474,711 | 365,529862 | 1186,258 | 2654,491139 | 2902,08 | 7762,546 | 20827,89 | 8128,30932 |
| Upregulated | ENSMUSG00000071656 | 3644,060064 | 3311,49713 | 3358,66331 | 2040,8358 | 1826,89715 | 2652,611468 | 2354,70652 | 3366,01046 | 4189,88901 | 2519,83505 | 3062,81032 | 3319,47 | 4589,04636 | 1112,284 | 884,9059 | 852,2953142 | 1171,367 | 1993,593195 | 4597,89 | 8573,866 | 20909,38 | 8183,94579 |
| Upregulated | ENSMUSG00000012428 | 4962,731828 | 4271,01219 | 6248,92925 | 3370,55885 | 3020,23247 | 1812,639953 | 2953,32221 | 4528,34741 | 3696,58403 | 3150,8725 | 9611,30635 | 8077,086 | 12081,183 | 8114,34 | 5181,2 | 4278,795941 | 10357,83 | 4336,956967 | 10706,23 | 12828,47 | 25434,57 | 14705,9763 |
| Upregulated | ENSMUSG00000041540 | 99,40268312 | 77,6547672 | 66,2655193 | 60,7047477 | 94,1437249 | 59,71361484 | 47,7075126 | 55,000171 | 86,4885345 | 56,0922185 | 94,9537446 | 110,3571 | 126,221315 | 189,9021 | 80,44599 | 99,35849118 | 134,0124 | 58,46023784 | 157,794 | 181,3305 | 344,5482 | 156,141071 |
| Upregulated | ENSMUSG00000024998 | 67,67842255 | 59,5707803 | 50,833823 | 65,5225848 | 82,6938124 | 69,00239937 | 51,1151921 | 113,66702 | 83,2852555 | 61,485701 | 27,6028327 | 69,19213 | 130,22834 | 96,35427 | 44,60372 | 49,22347269 | 46,32526 | 52,51512891 | 71,29249 | 165,9974 | 319,7478 | 178,575133 |
| Upregulated | ENSMUSG00000016024 | 5209,123586 | 1490,33327 | 3443,99151 | 2783,74629 | 349,858437 | 241,5083978 | 859,871121 | 1614,56057 | 1225,78812 | 737,828412 | 1400,01568 | 6522,453 | 18745,8687 | 14842,3 | 3390,679 | 6878,524536 | 5592,121 | 5194,043504 | 10359,27 | 10625,83 | 49837,26 | 18945,1166 |
| Upregulated | ENSMUSG00000026272 | 2,114950705 | 0 | 0 | 0 | 0 | 0 | 0 | 0 | 0 | 0 | 0 | 2,627549 | 20,0351293 | 18,70957 | 3,982475 | 1,823091581 | 0 | 1,981702978 | 2,851699 | 20,66634 | 55,80086 | 6,28153734 |
| Upregulated | ENSMUSG00000039252 | 51,81629226 | 29,7853902 | 41,7563546 | 54,9233432 | 8,90548749 | 13,26969219 | 0 | 74,5557873 | 43,7781471 | 15,1017511 | 54,1015521 | 82,32987 | 327,574364 | 224,5148 | 27,87732 | 13,67318686 | 54,59763 | 72,33215868 | 173,9537 | 320,6617 | 641,2671 | 151,654259 |
| Upregulated | ENSMUSG00000027276 | 72,96579931 | 32,976682 | 67,1732662 | 58,7776129 | 43,255225 | 54,40573796 | 29,5332221 | 57,444623 | 71,5398989 | 72,2726661 | 91,6414046 | 72,69552 | 93,1633512 | 191,7731 | 86,81795 | 72,92366325 | 105,8863 | 116,9204757 | 89,35325 | 170,664 | 441,9783 | 157,038434 |
| Upregulated | ENSMUSG00000017390 | 0 | 0 | 0 | 0 | 0 | 0 | 0 | 0 | 2,13551937 | 0 | 0 | 0 | 0 | 16,83861 | 0 | 0 | 1,654474 | 0 | 4,752832 | 13,99978 | 41,62922 | 0 |
| Upregulated | ENSMUSG00000020826 | 68,7358979 | 42,5505574 | 36,3098736 | 26,9798879 | 33,077525 | 55,73270718 | 60,2023374 | 61,1113011 | 69,4043796 | 66,8791836 | 264,987194 | 32,40644 | 99,17389 | 313,3853 | 117,0848 | 35,55028583 | 66,17895 | 27,74384169 | 15,20906 | 293,9954 | 1022,13 | 77,1731731 |
| Upregulated | ENSMUSG00000042286 | 0 | 7,44634754 | 3,63098736 | 7,70853939 | 0 | 0 | 4,54357263 | 3,66667807 | 3,20327906 | 14,0230546 | 15,4575863 | 39,41324 | 32,0562069 | 8,419306 | 3,18598 | 1,823091581 | 36,39842 | 4,954257444 | 7,604532 | 59,33241 | 110,716 | 9,87098725 |
| Upregulated | ENSMUSG00000054855 | 26,43688381 | 63,825836 | 22,693671 | 34,6884273 | 95,4159374 | 63,69452249 | 104,502171 | 162,556061 | 96,0983717 | 51,7774324 | 44,1645324 | 465,0762 | 1219,13762 | 278,7726 | 91,59692 | 88,41994169 | 273,8154 | 97,1034459 | 279,4665 | 1403,978 | 2298,464 | 406,505202 |
| Upregulated | ENSMUSG00000048636 | 0 | 0 | 8,16972156 | 0,96356742 | 1,2722125 | 1,326969219 | 2,27178632 | 3,66667807 | 1,06775969 | 4,31478604 | 2,20822662 | 11,38605 | 35,0614763 | 0 | 1,59299 | 2,734637372 | 14,06303 | 6,935960422 | 5,703399 | 25,99959 | 73,51543 | 13,4604372 |
| Upregulated | ENSMUSG00000026255 | 7,402327466 | 3,1912918 | 9,0774684 | 14,4535114 | 24,1720375 | 25,21241515 | 19,3101837 | 64,7779792 | 21,3551937 | 33,4395918 | 17,6658129 | 13,13775 | 13,022834 | 0 | 0 | 9,115457906 | 9,099605 | 9,908514888 | 13,30793 | 218,6633 | 432,2353 | 89,7362477 |
| Upregulated | ENSMUSG00000030116 | 2370,85974 | 2683,87641 | 3798,92053 | 2464,80547 | 989,781324 | 1665,346369 | 1547,08648 | 2532,45232 | 2905,3741 | 2486,39545 | 1780,93477 | 1602,805 | 1878,29337 | 1013,123 | 1153,325 | 938,8921644 | 1262,363 | 1321,795886 | 2083,642 | 16067,08 | 27754,29 | 5293,54125 |
| Upregulated | ENSMUSG00000037161 | 2,114950705 | 7,44634754 | 5,44648104 | 3,8542697 | 7,63327499 | 0 | 3,40767947 | 4,88890409 | 3,20327906 | 1,07869651 | 16,5616996 | 7,006798 | 4,00702586 | 2,806435 | 5,575465 | 0 | 0,827237 | 0 | 0,950566 | 75,33216 | 126,6591 | 18,844612 |
| Upregulated | ENSMUSG00000026109 | 410,3004367 | 329,76682 | 439,349471 | 380,609132 | 128,493462 | 179,1408445 | 134,035393 | 361,778902 | 240,245929 | 263,201948 | 368,773845 | 200,5696 | 540,948491 | 167,4506 | 54,16166 | 76,56984641 | 300,287 | 222,941585 | 558,9331 | 1661,307 | 3348,938 | 773,526456 |
| Upregulated | ENSMUSG00000029484 | 176,5983838 | 325,511764 | 223,305723 | 233,183317 | 50,8884999 | 27,86635359 | 233,993991 | 301,889827 | 207,145379 | 218,975391 | 80,6002715 | 323,1885 | 770,350721 | 159,0313 | 100,3584 | 186,8668871 | 336,6854 | 374,5418628 | 766,1566 | 2679,292 | 5006,135 | 1242,84703 |
| Upregulated | ENSMUSG00000115439 | 3,172426057 | 6,3825836 | 19,9704305 | 7,70853939 | 10,1777 | 0 | 4,54357263 | 3,66667807 | 5,33879843 | 3,23608953 | 0 | 14,0136 | 51,0895797 | 17,77409 | 1,59299 | 0 | 20,68092 | 8,917663399 | 28,51699 | 133,3313 | 302,919 | 60,123286 |
| Upregulated | ENSMUSG00000017969 | 23,26445775 | 62,7620721 | 85,328203 | 53,9597757 | 8,90548749 | 2,653938437 | 0 | 64,7779792 | 19,2196743 | 51,7774324 | 11,0411331 | 272,3893 | 706,238308 | 260,9985 | 49,38269 | 20,96555318 | 171,238 | 104,0394063 | 435,3595 | 2695,958 | 4768,76 | 570,722536 |
| Upregulated | ENSMUSG00000045608 | 10,57475352 | 8,51011147 | 13,6162026 | 1,92713485 | 5,08884999 | 11,94272297 | 0 | 6,11113011 | 1,06775969 | 2,15739302 | 5,52056654 | 0,87585 | 33,0579633 | 4,677392 | 7,964949 | 2,734637372 | 8,272368 | 13,87192084 | 16,15963 | 107,9983 | 187,7743 | 15,2551621 |
| Upregulated | ENSMUSG00000042357 | 6,344852114 | 2,12752787 | 15,4316963 | 10,5992417 | 0 | 11,94272297 | 11,3589316 | 0 | 3,20327906 | 19,4165372 | 7,72879316 | 0 | 18,0316164 | 3,741914 | 4,77897 | 3,646183163 | 5,790658 | 20,80788127 | 22,8136 | 113,9982 | 262,1755 | 36,7918616 |
| Upregulated | ENSMUSG00000029379 | 83,54055283 | 38,2955016 | 183,364862 | 54,9233432 | 17,810975 | 10,61575375 | 43,16394 | 61,1113011 | 33,1005503 | 18,3378407 | 134,701824 | 578,0608 | 878,540419 | 555,6742 | 223,0186 | 165,9013339 | 255,6162 | 573,703012 | 499,0474 | 3176,617 | 7784,664 | 524,957049 |
| Upregulated | ENSMUSG00000115759 | 115,2648134 | 10,6376393 | 200,612052 | 13,4899439 | 1,2722125 | 0 | 3,40767947 | 14,6667123 | 2,13551937 | 2,15739302 | 15,4575863 | 152,3978 | 296,519914 | 65,48349 | 10,35443 | 6,380820534 | 62,04276 | 59,45108933 | 139,7333 | 1009,318 | 2570,383 | 308,692692 |
| Upregulated | ENSMUSG00000115414 | 58,16114438 | 3,1912918 | 77,1584814 | 4,81783712 | 0 | 2,653938437 | 1,13589316 | 17,1111643 | 0 | 4,31478604 | 4,41645324 | 77,07477 | 106,186185 | 36,48366 | 7,964949 | 12,76164107 | 21,50816 | 20,80788127 | 38,97323 | 452,6596 | 1033,645 | 96,9151476 |
| Upregulated | ENSMUSG00000046997 | 32,78173592 | 9,57387541 | 5,44648104 | 21,1984833 | 11,4499125 | 10,61575375 | 15,9025042 | 22,0000684 | 4,27103874 | 30,2035023 | 11,0411331 | 14,0136 | 17,0298599 | 16,83861 | 8,761444 | 1,823091581 | 51,28868 | 5,945108933 | 31,36869 | 112,6649 | 232,0607 | 11,6657122 |
| Upregulated | ENSMUSG00000104266 | 16,91960564 | 13,8289311 | 2,72324052 | 2,89070227 | 1,2722125 | 2,653938437 | 12,4948247 | 13,4444862 | 10,6775969 | 6,47217906 | 1,10411331 | 15,76529 | 11,0193211 | 0,935478 | 3,18598 | 5,469274744 | 6,617895 | 3,963405955 | 25,6653 | 84,66535 | 119,5733 | 17,9472495 |
| Upregulated | ENSMUSG00000026576 | 6,344852114 | 0 | 3,63098736 | 0,96356742 | 0 | 0 | 2,27178632 | 8,55558215 | 5,33879843 | 0 | 1,10411331 | 7,006798 | 3,00526939 | 0 | 0 | 0 | 4,136184 | 1,981702978 | 13,30793 | 33,99947 | 69,97251 | 6,28153734 |
| Upregulated | ENSMUSG00000097754 | 2,114950705 | 1,06376393 | 0,90774684 | 13,4899439 | 3,8166375 | 1,326969219 | 0 | 3,66667807 | 5,33879843 | 0 | 3,31233993 | 18,39284 | 20,0351293 | 23,38696 | 22,30186 | 7,292366325 | 9,099605 | 14,86277233 | 19,01133 | 54,66581 | 131,0877 | 14,3577996 |
| Upregulated | ENSMUSG00000027859 | 108,9199613 | 210,625259 | 272,324052 | 146,462248 | 287,520025 | 266,7208129 | 455,493156 | 254,223013 | 279,753038 | 333,317221 | 394,168451 | 518,503 | 773,355991 | 703,4797 | 323,3769 | 608,0010424 | 570,7934 | 606,4011112 | 714,826 | 1647,308 | 3454,339 | 572,517261 |
| Upregulated | ENSMUSG00000087008 | 5,287376762 | 18,0839869 | 9,0774684 | 8,67210682 | 5,08884999 | 9,28878453 | 6,81535895 | 7,33335613 | 17,084155 | 16,1804476 | 16,5616996 | 24,52379 | 34,0597198 | 16,83861 | 10,35443 | 7,292366325 | 32,26224 | 16,84447531 | 35,17096 | 75,33216 | 157,6596 | 28,7155993 |
| Upregulated | ENSMUSG00000029223 | 5,287376762 | 9,57387541 | 5,44648104 | 0 | 0 | 0 | 2,27178632 | 0 | 2,13551937 | 0 | 0 | 21,89624 | 7,01229525 | 4,677392 | 7,964949 | 11,85009528 | 13,23579 | 0 | 1,901133 | 42,666 | 88,5728 | 17,9472495 |
| Upregulated | ENSMUSG00000024059 | 11,63222888 | 5,31881967 | 39,0331141 | 17,3442136 | 2,544425 | 10,61575375 | 6,81535895 | 13,4444862 | 19,2196743 | 3,23608953 | 8,83290647 | 31,53059 | 15,026347 | 9,354784 | 1,59299 | 6,380820534 | 7,445132 | 10,89936638 | 32,31926 | 84,66535 | 194,8602 | 45,7654863 |
| Upregulated | ENSMUSG00000057967 | 2,114950705 | 0 | 3,63098736 | 0,96356742 | 0 | 2,653938437 | 0 | 1,22222602 | 7,4743178 | 1,07869651 | 3,31233993 | 15,76529 | 7,01229525 | 0,935478 | 0,796495 | 0 | 3,308947 | 2,972554466 | 4,752832 | 27,33291 | 52,25795 | 9,87098725 |
| Upregulated | ENSMUSG00000021943 | 1431,821627 | 630,812013 | 1703,84082 | 1146,64523 | 87,7826624 | 155,2553986 | 76,1048416 | 1060,89219 | 429,239394 | 831,675009 | 927,455179 | 162,0322 | 1046,83551 | 144,9992 | 46,19671 | 30,08101109 | 508,7507 | 630,1815469 | 655,8909 | 1811,305 | 3866,203 | 1680,75992 |
| Upregulated | ENSMUSG00000028681 | 305,6103768 | 15,956459 | 318,619141 | 107,919551 | 64,8828374 | 23,88544594 | 72,6971621 | 161,333835 | 133,469961 | 65,8004871 | 109,307218 | 112,9846 | 139,244149 | 52,38679 | 37,43526 | 15,49627844 | 181,1649 | 143,6734659 | 170,1514 | 391,3272 | 647,4672 | 342,792466 |
| Upregulated | ENSMUSG00000031283 | 1083,912236 | 534,009495 | 1298,07798 | 761,218265 | 40,7108 | 53,07876874 | 23,8537563 | 971,669687 | 201,806581 | 418,534246 | 495,746876 | 261,0032 | 1341,35191 | 345,1915 | 43,80722 | 16,40782423 | 417,7546 | 405,2582589 | 868,8178 | 1321,313 | 2309,093 | 1779,46979 |
| Upregulated | ENSMUSG00000052504 | 197,7478909 | 238,283121 | 268,693065 | 191,749917 | 99,2325749 | 75,63724546 | 29,5332221 | 287,223115 | 170,84155 | 96,0039893 | 132,493597 | 27,15134 | 319,560312 | 52,38679 | 54,95815 | 79,30448379 | 213,4271 | 217,9873275 | 200,5695 | 445,993 | 632,4098 | 544,699024 |
| Upregulated | ENSMUSG00000040254 | 121,6096655 | 357,424682 | 125,269064 | 185,004945 | 99,2325749 | 70,32936859 | 111,31753 | 248,111882 | 192,196743 | 161,804476 | 112,619557 | 21,02039 | 118,207263 | 44,90296 | 6,37196 | 20,05400739 | 93,47776 | 130,7923965 | 385,93 | 543,3249 | 760,8404 | 409,19729 |
| Upregulated | ENSMUSG00000011171 | 47,58639085 | 61,6983082 | 34,4943799 | 53,9597757 | 50,8884999 | 27,86635359 | 17,0383974 | 63,5557531 | 54,455744 | 59,328308 | 61,8303453 | 21,89624 | 22,0386422 | 2,806435 | 3,18598 | 3,646183163 | 30,60776 | 63,41449528 | 113,1174 | 138,6645 | 211,689 | 62,8153734 |
| Upregulated | ENSMUSG00000026893 | 0 | 11,7014033 | 9,0774684 | 10,5992417 | 10,1777 | 0 | 0 | 14,6667123 | 18,1519147 | 4,31478604 | 3,31233993 | 6,130948 | 15,026347 | 0 | 12,74392 | 4,557728953 | 5,790658 | 9,908514888 | 33,26983 | 36,66609 | 74,40115 | 24,2287869 |
| Upregulated | ENSMUSG00000033715 | 21,14950705 | 14,8926951 | 19,0626836 | 21,1984833 | 1,2722125 | 1,326969219 | 20,4460768 | 26,8889725 | 44,8459068 | 17,2591441 | 9,93701978 | 0 | 10,0175646 | 3,741914 | 0,796495 | 2,734637372 | 0 | 36,66150509 | 8,555098 | 53,99916 | 108,0588 | 31,4076867 |
| Upregulated | ENSMUSG00000037206 | 548,8297079 | 511,670452 | 837,850333 | 706,294922 | 310,41985 | 459,1313496 | 529,326212 | 1372,55982 | 972,729074 | 902,868978 | 654,739192 | 126,1224 | 286,502349 | 105,7091 | 84,42846 | 168,6359713 | 157,175 | 426,0661402 | 368,8198 | 1770,639 | 4690,816 | 1072,34816 |
| Upregulated | ENSMUSG00000000392 | 201,9777923 | 431,888157 | 514,692458 | 331,467194 | 176,837537 | 144,6396448 | 184,014692 | 315,334314 | 354,496216 | 281,539789 | 337,858672 | 122,619 | 209,367101 | 138,4508 | 60,53362 | 97,5353996 | 87,68711 | 199,1611493 | 285,1699 | 656,6564 | 1432,222 | 524,059687 |
| Upregulated | ENSMUSG00000030111 | 11,63222888 | 21,2752787 | 11,8007089 | 6,74497197 | 0 | 0 | 0 | 31,7778766 | 0 | 0 | 33,1233993 | 0,87585 | 21,0368858 | 0 | 0,796495 | 0 | 6,617895 | 13,87192084 | 1,901133 | 73,33219 | 189,5458 | 31,4076867 |
| Upregulated | ENSMUSG00000075122 | 0 | 9,57387541 | 11,8007089 | 8,67210682 | 1,2722125 | 0 | 0 | 3,66667807 | 8,54207749 | 9,70826858 | 0 | 0 | 3,00526939 | 0 | 3,982475 | 0 | 0 | 2,972554466 | 0 | 47,99925 | 115,1446 | 30,5103242 |
| Upregulated | ENSMUSG00000074968 | 0 | 11,7014033 | 19,0626836 | 4,81783712 | 0 | 0 | 0 | 3,66667807 | 0 | 8,62957207 | 1,10411331 | 0 | 3,00526939 | 0 | 0 | 0,911545791 | 0 | 4,954257444 | 0 | 39,99938 | 81,48698 | 20,639337 |
| Upregulated | ENSMUSG00000022371 | 2934,494103 | 1946,688 | 3305,10624 | 1805,72535 | 605,573149 | 958,0717758 | 586,12087 | 2132,78441 | 1250,34659 | 2674,08865 | 1931,09418 | 945,0418 | 1269,22544 | 510,7712 | 231,78 | 393,7877816 | 947,1862 | 1351,521431 | 3390,671 | 6613,897 | 17873,11 | 5379,68805 |
| Upregulated | ENSMUSG00000006386 | 159,6787782 | 103,185102 | 151,593722 | 71,3039894 | 5,08884999 | 5,307876874 | 17,0383974 | 185,778355 | 65,1333408 | 116,499223 | 98,2660845 | 75,32307 | 166,291573 | 4,677392 | 3,982475 | 1,823091581 | 26,47158 | 30,71639615 | 165,3986 | 583,9909 | 1144,361 | 424,452452 |
| Upregulated | ENSMUSG00000043719 | 124,7820916 | 106,376393 | 141,608507 | 79,9760962 | 27,988675 | 11,94272297 | 3,40767947 | 213,889554 | 19,2196743 | 97,0826858 | 117,036011 | 44,66833 | 133,23361 | 16,83861 | 5,575465 | 0 | 29,78053 | 44,588317 | 178,7065 | 503,9921 | 1030,102 | 289,84808 |
| Upregulated | ENSMUSG00000040829 | 2,114950705 | 4,25505574 | 1,81549368 | 8,67210682 | 7,63327499 | 9,28878453 | 1,13589316 | 9,77780817 | 2,13551937 | 8,62957207 | 16,5616996 | 11,38605 | 22,0386422 | 0 | 0 | 12,76164107 | 14,89026 | 11,89021787 | 26,61586 | 41,33269 | 75,28688 | 45,7654863 |
| Upregulated | ENSMUSG00000026249 | 1331,361469 | 1691,38465 | 1780,9993 | 1403,91774 | 1553,37146 | 769,6421468 | 1658,40401 | 1624,33838 | 2124,84177 | 1347,29194 | 1292,91668 | 2794,836 | 2832,96728 | 1496,765 | 2966,147 | 2287,979934 | 3172,453 | 2755,55799 | 3642,571 | 6279,902 | 6757,219 | 6048,2231 |
| Upregulated | ENSMUSG00000031722 | 850,2101833 | 24,4665705 | 234,198685 | 266,908176 | 24,1720375 | 33,17423047 | 43,16394 | 55,000171 | 32,0327906 | 14,0230546 | 41,9563057 | 1002,848 | 5888,3245 | 1378,895 | 290,7207 | 90,24303327 | 10455,45 | 965,0893501 | 5077,926 | 15217,76 | 18076,82 | 17328,9668 |
| Upregulated | ENSMUSG00000086058 | 2,114950705 | 0 | 3,63098736 | 14,4535114 | 0 | 0 | 0 | 2,44445204 | 0 | 0 | 0 | 2,627549 | 12,0210776 | 7,483827 | 0 | 3,646183163 | 10,75408 | 0 | 17,1102 | 33,33281 | 48,71504 | 40,3813115 |
| Upregulated | ENSMUSG00000025092 | 61,33357043 | 76,5910032 | 87,1436966 | 113,700956 | 26,7164625 | 76,96421468 | 77,2407348 | 100,222534 | 66,2011005 | 100,318775 | 104,890764 | 94,59177 | 130,22834 | 54,25775 | 54,95815 | 63,80820534 | 137,3213 | 61,43279231 | 125,4748 | 297,9953 | 400,3491 | 294,334893 |
| Upregulated | ENSMUSG00000035112 | 4,229901409 | 0 | 9,98521524 | 0 | 5,08884999 | 0 | 6,81535895 | 6,11113011 | 5,33879843 | 2,15739302 | 7,72879316 | 0,87585 | 8,01405172 | 7,483827 | 0 | 7,292366325 | 2,481711 | 5,945108933 | 11,4068 | 24,66628 | 24,80038 | 22,4340619 |
| Upregulated | ENSMUSG00000098318 | 29,60930986 | 13,8289311 | 57,1880509 | 15,4170788 | 7,63327499 | 0 | 12,4948247 | 14,6667123 | 17,084155 | 9,70826858 | 14,353473 | 0,87585 | 9,01580818 | 1,870957 | 3,982475 | 0,911545791 | 9,926842 | 41,61576253 | 21,86303 | 86,66531 | 94,7729 | 69,0969108 |
| Upregulated | ENSMUSG00000038591 | 0 | 0 | 16,3394431 | 2,89070227 | 0 | 2,653938437 | 7,95125211 | 2,44445204 | 0 | 4,31478604 | 0 | 0,87585 | 8,01405172 | 0 | 2,389485 | 4,557728953 | 0 | 10,89936638 | 4,752832 | 47,99925 | 35,42912 | 21,5366995 |
| Upregulated | ENSMUSG00000027459 | 75,08075001 | 28,7216262 | 113,468355 | 51,0690735 | 41,9830125 | 46,44392265 | 52,2510853 | 61,1113011 | 57,659023 | 32,3608953 | 48,5809856 | 65,68873 | 131,230097 | 94,48332 | 40,62124 | 82,03912116 | 95,13224 | 61,43279231 | 96,00722 | 152,6643 | 213,4605 | 61,0206485 |
| Upregulated | ENSMUSG00000031480 | 14,80465493 | 6,3825836 | 34,4943799 | 12,5263765 | 11,4499125 | 7,961815312 | 10,2230384 | 25,6667465 | 9,60983717 | 11,8656616 | 24,2904928 | 27,15134 | 40,0702586 | 18,70957 | 15,9299 | 20,05400739 | 28,95329 | 17,8353268 | 34,22039 | 49,99922 | 90,34426 | 26,0235118 |
| Upregulated | ENSMUSG00000099406 | 22,2069824 | 0 | 6,35422788 | 0,96356742 | 5,08884999 | 10,61575375 | 4,54357263 | 9,77780817 | 3,20327906 | 4,31478604 | 3,31233993 | 26,27549 | 28,049181 | 14,03218 | 4,77897 | 11,85009528 | 14,89026 | 6,935960422 | 16,15963 | 44,66597 | 65,54387 | 10,7683497 |
| Upregulated | ENSMUSG00000024827 | 276,001067 | 74,4634754 | 167,933165 | 79,9760962 | 34,3497375 | 2,653938437 | 26,1255426 | 31,7778766 | 11,7453565 | 11,8656616 | 32,019286 | 140,136 | 487,855398 | 227,3212 | 93,9864 | 86,59685011 | 153,8661 | 183,3075254 | 339,3522 | 748,655 | 814,8698 | 255,748306 |
| Upregulated | ENSMUSG00000039960 | 116,3222888 | 20,2115147 | 88,9591903 | 53,9597757 | 41,9830125 | 69,00239937 | 65,8818032 | 57,444623 | 80,0819764 | 46,3839499 | 82,8084982 | 62,18533 | 180,316164 | 122,5477 | 54,95815 | 81,12757537 | 105,0591 | 102,0577033 | 99,80948 | 227,9964 | 317,9764 | 131,014922 |
| Upregulated | ENSMUSG00000029596 | 37,01163733 | 15,956459 | 22,693671 | 23,1256182 | 12,722125 | 5,307876874 | 22,7178632 | 8,55558215 | 0 | 16,1804476 | 2,20822662 | 7,882647 | 51,0895797 | 16,83861 | 3,982475 | 13,67318686 | 14,06303 | 16,84447531 | 71,29249 | 61,33238 | 108,9445 | 52,0470237 |
| Upregulated | ENSMUSG00000026686 | 44,4139648 | 11,7014033 | 51,7415699 | 22,1620508 | 5,08884999 | 1,326969219 | 0 | 7,33335613 | 9,60983717 | 0 | 30,9151726 | 58,68193 | 76,1334913 | 5,61287 | 14,33691 | 6,380820534 | 4,963421 | 18,82617829 | 63,68796 | 103,3317 | 142,6022 | 74,4810856 |
| Upregulated | ENSMUSG00000027074 | 3779,416909 | 1439,2726 | 2453,63971 | 1009,81866 | 393,113662 | 1214,176835 | 555,451754 | 2228,11804 | 2617,07899 | 1018,2895 | 1359,16348 | 2688,859 | 5406,47964 | 2177,794 | 1213,858 | 1115,732048 | 2465,166 | 1363,411649 | 2899,228 | 6389,234 | 11487,01 | 3945,70281 |
| Upregulated | ENSMUSG00000018339 | 34390,15593 | 16860,6584 | 16476,5129 | 19311,8183 | 3759,38793 | 2616,783299 | 1367,61536 | 23560,851 | 8306,10259 | 10070,7106 | 15861,6918 | 18302,63 | 70767,0819 | 13129,44 | 2399,839 | 2163,098161 | 18703 | 10080,92305 | 44328,72 | 44251,98 | 137324,2 | 56917,9072 |
| Upregulated | ENSMUSG00000031497 | 47,58639085 | 9,57387541 | 25,4169115 | 19,2713485 | 5,08884999 | 3,980907656 | 18,1742905 | 29,3334245 | 16,0163953 | 10,7869651 | 15,4575863 | 16,64114 | 87,1528124 | 28,99983 | 3,982475 | 0,911545791 | 21,50816 | 40,62491104 | 52,28116 | 67,33228 | 155,0024 | 60,123286 |
| Upregulated | ENSMUSG00000030523 | 7,402327466 | 22,3390426 | 17,24719 | 17,3442136 | 19,0831875 | 14,5966614 | 3,40767947 | 4,88890409 | 3,20327906 | 3,23608953 | 1,10411331 | 12,2619 | 16,0281034 | 14,03218 | 14,33691 | 26,43482793 | 13,23579 | 25,76213871 | 21,86303 | 33,99947 | 67,31533 | 31,4076867 |
| Upregulated | ENSMUSG00000017493 | 4197,119673 | 1371,19171 | 7191,17047 | 3281,91065 | 917,265212 | 1127,923836 | 1154,06745 | 2717,00845 | 1765,00676 | 2732,33826 | 1250,96038 | 2897,311 | 8602,08276 | 1586,571 | 1652,727 | 2225,994821 | 6953,753 | 4794,730354 | 4508,537 | 11185,16 | 16556,91 | 7177,10509 |
| Upregulated | ENSMUSG00000072620 | 131,1269437 | 134,034256 | 229,659951 | 15,4170788 | 5,08884999 | 37,15513812 | 18,1742905 | 40,3334587 | 10,6775969 | 36,6756813 | 107,098991 | 107,7295 | 201,353049 | 40,22557 | 97,96888 | 180,4860665 | 309,3866 | 218,978179 | 326,0443 | 477,9925 | 685,5535 | 236,903694 |
| Upregulated | ENSMUSG00000026479 | 617,5656058 | 393,592656 | 802,448207 | 379,645565 | 129,765675 | 204,3532597 | 155,617363 | 253,000787 | 254,126805 | 209,267123 | 300,31882 | 254,8723 | 417,732446 | 138,4508 | 179,2114 | 133,0856854 | 301,1142 | 237,8043573 | 408,7436 | 1044,65 | 1794,485 | 713,40317 |
| Upregulated | ENSMUSG00000000567 | 372,231324 | 169,138465 | 578,234737 | 208,130564 | 92,8715124 | 111,4654144 | 118,132888 | 112,444794 | 69,4043796 | 266,438038 | 310,25584 | 244,3621 | 399,700829 | 184,2892 | 146,5551 | 174,105246 | 153,0388 | 297,2554466 | 346,0062 | 947,3185 | 1486,252 | 450,475964 |
| Upregulated | ENSMUSG00000019122 | 86,71297889 | 78,7185311 | 302,279698 | 143,571546 | 55,9773499 | 72,98330702 | 61,3382305 | 30,5556505 | 45,9136665 | 87,3744172 | 102,682538 | 28,02719 | 169,296843 | 55,19322 | 136,9971 | 61,98511376 | 102,5774 | 176,371565 | 112,1668 | 356,6611 | 629,7526 | 150,756896 |
| Upregulated | ENSMUSG00000075410 | 9,517278171 | 3,1912918 | 19,9704305 | 8,67210682 | 3,8166375 | 0 | 0 | 6,11113011 | 3,20327906 | 7,55087556 | 4,41645324 | 4,379248 | 21,0368858 | 4,677392 | 2,389485 | 0,911545791 | 3,308947 | 15,85362382 | 19,01133 | 26,66625 | 53,14368 | 13,4604372 |
| Upregulated | ENSMUSG00000038600 | 1,057475352 | 0 | 1,81549368 | 0 | 0 | 1,326969219 | 5,67946579 | 6,11113011 | 1,06775969 | 0 | 1,10411331 | 2,627549 | 3,00526939 | 0 | 0 | 0 | 0 | 21,79873275 | 14,2585 | 11,99981 | 25,68611 | 16,1525246 |

| | | | | | | | | | | | | | | | | | | | | | | | |
|---|---|---|---|---|---|---|---|---|---|---|---|---|---|---|---|---|---|---|---|---|---|---|---|
| Upregulated | ENSMUSG00000027460 | 199,8628416 | 30,8491541 | 205,150786 | 753,509726 | 71,2438999 | 136,6778295 | 202,188982 | 243,222978 | 74,743178 | 625,643975 | 470,35227 | 70,94383 | 198,34778 | 943,8977 | 171,2464 | 560,6006612 | 2826,668 | 1572,481313 | 3150,177 | 1766,639 | 3716,515 | 3142,5634 |
| Upregulated | ENSMUSG00000038578 | 7,402327466 | 12,7651672 | 24,5091647 | 13,4899439 | 13,9943375 | 14,5966614 | 2,27178632 | 6,11113011 | 6,40655811 | 9,70826858 | 8,83290647 | 4,379248 | 28,049181 | 25,25792 | 31,0633 | 9,115457906 | 30,60776 | 36,66150509 | 57,98456 | 49,99922 | 57,57232 | 47,5602113 |
| Upregulated | ENSMUSG00000009378 | 29,60930986 | 21,2752787 | 29,9556457 | 88,648203 | 38,166375 | 18,57756906 | 20,4460768 | 12,2222602 | 28,8295115 | 22,6526267 | 100,474311 | 43,79248 | 102,179159 | 26,19339 | 71,68455 | 97,5353996 | 50,46145 | 51,52427742 | 92,20495 | 85,99866 | 145,2594 | 179,472495 |
| Upregulated | ENSMUSG00000020178 | 2,114950705 | 2,12752787 | 4,5387342 | 0 | 2,544425 | 0 | 2,27178632 | 12,2222602 | 14,9486356 | 2,15739302 | 25,3946061 | 25,39964 | 60,1053879 | 6,548349 | 20,70887 | 7,292366325 | 35,57118 | 13,87192084 | 33,26983 | 33,99947 | 63,77242 | 33,2024117 |
| Upregulated | ENSMUSG00000070594 | 15,86213028 | 9,57387541 | 2,72324052 | 8,67210682 | 2,544425 | 1,326969219 | 1,13589316 | 9,77780817 | 11,7453565 | 5,39348255 | 11,0411331 | 18,39284 | 46,0807974 | 10,29026 | 5,575465 | 4,557728953 | 45,49803 | 9,908514888 | 29,46756 | 23,99963 | 69,08679 | 73,5837231 |
| Upregulated | ENSMUSG00000033898 | 2,114950705 | 9,57387541 | 19,0626836 | 24,0891856 | 3,8166375 | 5,307876874 | 1,13589316 | 31,7778766 | 23,4907131 | 5,39348255 | 18,7699262 | 35,03399 | 70,1229525 | 10,29026 | 5,575465 | 12,76164107 | 93,47776 | 16,84447531 | 50,38002 | 64,66566 | 145,2594 | 107,683497 |
| Upregulated | ENSMUSG00000026365 | 1237,246162 | 714,849364 | 1056,61732 | 1119,66535 | 137,39895 | 173,8329676 | 172,65576 | 1701,33862 | 1120,07991 | 807,943685 | 872,249514 | 2026,716 | 4953,68572 | 1079,542 | 755,8737 | 690,9517093 | 4367,811 | 1418,899332 | 2674,894 | 3166,617 | 10045,04 | 6034,76266 |
| Upregulated | ENSMUSG00000089991 | 143,8166479 | 107,440157 | 84,4204561 | 99,2474447 | 8,90548749 | 3,980907656 | 24,9896495 | 179,667225 | 92,8950927 | 69,0365766 | 103,786651 | 231,2243 | 472,829051 | 76,70923 | 92,39341 | 70,18902588 | 417,7546 | 133,764951 | 298,4779 | 318,6617 | 869,7849 | 658,664058 |
| Upregulated | ENSMUSG00000069763 | 334,1622113 | 257,430872 | 814,248915 | 567,541213 | 501,251724 | 451,1695343 | 553,179968 | 898,336126 | 697,247075 | 763,717128 | 224,135002 | 37,66154 | 180,316164 | 29,93531 | 26,28433 | 30,99255688 | 49,63421 | 114,9387727 | 88,40268 | 534,6583 | 1767,027 | 558,159461 |
| Upregulated | ENSMUSG00000078350 | 205,1502183 | 168,074702 | 197,888811 | 221,620508 | 249,35365 | 281,3174743 | 232,858097 | 356,889998 | 207,145379 | 249,178894 | 207,573302 | 63,06118 | 99,17389 | 64,54801 | 46,9932 | 51,04656428 | 71,96961 | 128,8106935 | 136,8816 | 303,3286 | 552,6943 | 236,903694 |
| Upregulated | ENSMUSG00000001604 | 98,34520777 | 88,2924065 | 157,94795 | 141,644411 | 178,10975 | 205,6802289 | 126,084141 | 163,778287 | 176,180348 | 143,466636 | 146,84707 | 19,26869 | 72,1264654 | 39,29009 | 31,8598 | 27,34637372 | 40,53461 | 158,5362382 | 105,5129 | 172,664 | 295,8332 | 230,622157 |
| Upregulated | ENSMUSG00000033578 | 0 | 4,25505574 | 15,4316963 | 4,81783712 | 1,2722125 | 18,57756906 | 4,54357263 | 6,11113011 | 18,1519147 | 10,7869651 | 5,52056654 | 0 | 0 | 0 | 0,796495 | 0 | 0 | 1,981702978 | 0 | 24,66628 | 60,22951 | 4,48681239 |
| Upregulated | ENSMUSG00000035314 | 43,35648945 | 64,8896 | 97,1289119 | 95,393175 | 52,1607124 | 72,98330702 | 44,2998332 | 77,0002394 | 64,0655811 | 85,2170242 | 37,5398525 | 33,28229 | 43,075528 | 14,03218 | 11,94742 | 13,67318686 | 61,21553 | 56,47853486 | 73,19362 | 153,3309 | 225,8606 | 88,8388853 |
| Upregulated | ENSMUSG00000026193 | 29151,42304 | 34042,5734 | 38161,6772 | 18985,169 | 21195,0602 | 21292,54808 | 24418,2952 | 37747,2285 | 26936,3736 | 43581,4964 | 21596,4563 | 9310,282 | 9464,59508 | 5798,095 | 6269,212 | 7673,392466 | 4667,27 | 13168,41629 | 19056,96 | 63009,68 | 92203,4 | 31901,2361 |
| Upregulated | ENSMUSG00000004098 | 6835,520677 | 5678,37188 | 9521,3566 | 4549,96538 | 7345,75497 | 9385,653283 | 7805,85778 | 5441,35025 | 3835,39279 | 10797,7521 | 6051,64505 | 2031,971 | 2253,95205 | 1189,929 | 874,5515 | 1365,495594 | 3406,561 | 2542,52492 | 4722,414 | 16975,07 | 26591,33 | 7097,23983 |
| Upregulated | ENSMUSG00000045954 | 211,4950705 | 186,158688 | 292,294482 | 245,709693 | 239,17595 | 400,744704 | 424,824041 | 188,222807 | 131,334441 | 325,766346 | 315,776406 | 81,45402 | 135,237123 | 80,45114 | 25,48784 | 51,04656428 | 78,5875 | 108,9936638 | 249,999 | 507,3254 | 759,9546 | 334,716204 |
| Upregulated | ENSMUSG00000030409 | 379,6336515 | 252,112052 | 395,777622 | 284,25239 | 288,792237 | 530,7876874 | 358,942238 | 371,556711 | 216,755216 | 433,635997 | 444,957663 | 119,9914 | 126,221315 | 58,93514 | 111,5093 | 80,21602958 | 161,3112 | 193,2160403 | 301,3296 | 637,3234 | 848,5274 | 452,270689 |
| Upregulated | ENSMUSG00000032179 | 26,43688381 | 1,06376393 | 10,8929621 | 31,797725 | 41,9830125 | 51,75179953 | 32,9409016 | 17,1111643 | 9,60983717 | 23,7313232 | 5,52056654 | 0 | 49,0860668 | 3,741914 | 0 | 0 | 0 | 2,972554466 | 11,4068 | 17,99972 | 65,54387 | 9,87098725 |
| Upregulated | ENSMUSG00000071713 | 24,3219331 | 20,2115147 | 39,0331141 | 79,0125288 | 142,4878 | 208,3341673 | 60,2023374 | 106,333664 | 87,5562942 | 83,0596312 | 155,679977 | 133,1292 | 282,495323 | 118,8058 | 139,3866 | 34,63874004 | 19,85368 | 102,0577033 | 62,73739 | 125,3314 | 521,6938 | 145,372721 |
| Upregulated | ENSMUSG00000042842 | 1,057475352 | 6,3825836 | 4,5387342 | 19,2713485 | 11,4499125 | 39,80907656 | 46,5716195 | 12,2222602 | 22,4229534 | 20,4952337 | 7,72879316 | 1,751699 | 31,0544504 | 0 | 0 | 0 | 6,617895 | 1,981702978 | 4,752832 | 42,666 | 92,11571 | 8,0762623 |
| Upregulated | ENSMUSG00000078612 | 15,86213028 | 46,8056131 | 16,3394431 | 24,0891856 | 24,1720375 | 46,44392265 | 57,9305511 | 40,3334587 | 23,4907131 | 35,5969848 | 49,6850989 | 27,15134 | 59,1036314 | 12,16122 | 5,575465 | 5,469274744 | 0,827237 | 17,8353268 | 11,4068 | 51,99919 | 99,20154 | 43,0733989 |
| Upregulated | ENSMUSG00000004791 | 68,7358979 | 341,468223 | 126,176811 | 110,810254 | 265,892412 | 114,1193528 | 306,691153 | 190,667259 | 219,958495 | 224,368874 | 150,15941 | 126,1224 | 85,1492995 | 175,8699 | 129,8287 | 90,24303327 | 42,18908 | 46,57001997 | 71,29249 | 275,9957 | 584,5805 | 157,038434 |
| Upregulated | ENSMUSG00000068452 | 166,0236303 | 159,56459 | 98,9444056 | 87,6846356 | 157,75435 | 155,2553986 | 210,140234 | 125,88928 | 146,283077 | 165,040566 | 139,118277 | 79,70232 | 97,1703771 | 97,28975 | 56,55114 | 66,54284272 | 35,57118 | 57,46938635 | 73,19362 | 79,3321 | 374,663 | 111,272947 |
| Upregulated | ENSMUSG00000102252 | 70,85084861 | 89,3561705 | 69,8965067 | 98,2838772 | 33,077525 | 38,48210734 | 71,561269 | 63,5557531 | 45,9136665 | 89,5318103 | 56,3097787 | 12,2619 | 36,0632327 | 12,16122 | 11,94742 | 24,61173635 | 24,81711 | 33,68895062 | 28,51699 | 47,99925 | 155,0024 | 81,6599854 |
| Upregulated | ENSMUSG00000020096 | 11,63222888 | 21,2752787 | 19,9704305 | 20,2349159 | 3,8166375 | 14,5966614 | 45,4357263 | 17,1111643 | 6,40655811 | 63,643094 | 47,4768723 | 6,130948 | 5,00878232 | 1,870957 | 6,37196 | 0,911545791 | 5,790658 | 17,8353268 | 7,604532 | 9,999844 | 116,0304 | 55,6364736 |
| Upregulated | ENSMUSG00000111954 | 4,229901409 | 22,3390426 | 15,4316963 | 6,74497197 | 6,36106249 | 3,980907656 | 20,4460768 | 14,6667123 | 3,20327906 | 3,23608953 | 3,31233993 | 0 | 3,00526939 | 1,870957 | 0,796495 | 3,646183163 | 1,654474 | 0 | 7,604532 | 1,999969 | 23,02893 | 17,9472495 |
| Upregulated | ENSMUSG00000004885 | 7,402327466 | 9,57387541 | 16,3394431 | 25,052753 | 19,0831875 | 22,55847672 | 35,2126879 | 25,6667465 | 19,2196743 | 20,4952337 | 12,1452464 | 12,2619 | 14,0245905 | 2,806435 | 7,964949 | 33,72719425 | 9,926842 | 3,963405955 | 4,752832 | 28,66622 | 38,97203 | 16,1525246 |
| Upregulated | ENSMUSG00000023078 | 4608,477585 | 301,045193 | 1269,03008 | 483,710847 | 12,722125 | 9,28878453 | 23,8537563 | 209,00065 | 32,0327906 | 43,1478604 | 57,4138921 | 378,3671 | 3765,60255 | 511,7067 | 91,59692 | 142,2011433 | 317,6589 | 58,46023784 | 267,1092 | 377,3274 | 6445,443 | 2944,24629 |
| Upregulated | ENSMUSG00000027562 | 15,86213028 | 2,12752787 | 0,90774684 | 6,74497197 | 0 | 0 | 0 | 0 | 0 | 0 | 0 | 0,87585 | 21,0368858 | 0 | 0 | 0 | 1,654474 | 2,972554466 | 11,4068 | 5,33325 | 31,00048 | 2,69208743 |
| Upregulated | ENSMUSG00000006344 | 1380,005335 | 252,112052 | 822,418637 | 408,552588 | 86,5104499 | 53,07876874 | 160,160935 | 262,778595 | 103,57269 | 118,656616 | 203,156849 | 913,5112 | 2160,78869 | 609,9319 | 186,3798 | 247,9404551 | 852,0539 | 527,132992 | 1244,292 | 724,6554 | 1498,652 | 707,121632 |
| Upregulated | ENSMUSG00000022181 | 1428,649201 | 525,499383 | 551,910079 | 476,965875 | 111,9547 | 19,90453828 | 48,8434058 | 575,668456 | 338,47982 | 157,48969 | 524,453822 | 684,0386 | 2423,24889 | 485,5133 | 31,0633 | 115,7663154 | 332,5492 | 368,5967538 | 1120,718 | 51,33253 | 1516,366 | 1077,73234 |
| Upregulated | ENSMUSG00000031981 | 263,3113627 | 52,1244328 | 84,4204561 | 44,3241015 | 34,3497375 | 31,84726125 | 19,3101837 | 34,2223286 | 26,6939921 | 39,9117708 | 67,3509118 | 76,19892 | 216,379396 | 37,41914 | 3,18598 | 20,05400739 | 47,97974 | 49,54257444 | 164,448 | 36,66609 | 201,946 | 172,293596 |
| Upregulated | ENSMUSG00000032281 | 238,9894296 | 26,5940983 | 167,025419 | 133,935872 | 13,9943375 | 53,07876874 | 0 | 141,778219 | 64,0655811 | 43,1478604 | 66,2467985 | 6,130948 | 402,706099 | 57,99966 | 2,389485 | 4,557728953 | 78,5875 | 62,4236438 | 61,78682 | 120,6648 | 346,3197 | 369,713341 |
| Upregulated | ENSMUSG00000031980 | 1189,659771 | 491,458937 | 539,201623 | 567,541213 | 151,393287 | 72,98330702 | 22,7178632 | 837,224825 | 198,603302 | 140,230546 | 542,119635 | 628,8601 | 3718,52 | 706,2862 | 73,27754 | 82,95066695 | 694,0517 | 335,8986547 | 1363,112 | 505,3254 | 4171,779 | 2697,47161 |
| Upregulated | ENSMUSG00000026582 | 9,517278171 | 61,6983082 | 49,9260762 | 17,3442136 | 0 | 1,326969219 | 0 | 24,4445204 | 33,1005503 | 31,2821988 | 71,7673651 | 141,8877 | 480,843103 | 82,3221 | 0 | 25,52328214 | 39,70737 | 50,53342593 | 59,88569 | 76,66547 | 814,8698 | 275,490281 |
| Upregulated | ENSMUSG00000029322 | 429,334993 | 314,874124 | 404,855091 | 298,705901 | 3,8166375 | 15,92363062 | 4,54357263 | 140,555993 | 42,7103874 | 52,8561289 | 50,7892122 | 235,6036 | 2944,16225 | 449,9651 | 142,5726 | 44,66574374 | 391,283 | 97,1034459 | 548,4769 | 841,3202 | 3774,973 | 934,154339 |
| Upregulated | ENSMUSG00000033174 | 320,4150318 | 168,074702 | 380,345926 | 94,4296076 | 48,344075 | 33,17423047 | 44,2998332 | 217,556232 | 133,469961 | 74,4300591 | 81,7043849 | 225,9692 | 910,596626 | 286,2564 | 98,76537 | 113,9432238 | 214,2543 | 167,4539016 | 345,0556 | 313,9951 | 1411,85 | 597,64341 |
| Upregulated | ENSMUSG00000008734 | 70,85084861 | 0 | 31,7711394 | 17,3442136 | 7,63327499 | 43,78998421 | 9,08714527 | 12,2222602 | 8,54207749 | 11,8656616 | 23,1863795 | 57,80608 | 143,251174 | 27,12887 | 11,15093 | 19,1424616 | 7,445132 | 21,79873275 | 40,87436 | 86,66531 | 158,5453 | 42,1760364 |
| Upregulated | ENSMUSG00000047180 | 0 | 0 | 0 | 0,96356742 | 0 | 0 | 0 | 0 | 1,06775969 | 0 | 0 | 0 | 24,0421551 | 0,935478 | 0 | 0 | 0 | 4,954257444 | 0 | 11,99981 | 20,37174 | 10,7683497 |
| Upregulated | ENSMUSG00000051048 | 249,5641831 | 152,118243 | 357,652255 | 82,8667985 | 11,4499125 | 13,26969219 | 5,67946579 | 4,88890409 | 3,20327906 | 45,3052534 | 83,9126115 | 95,46762 | 204,358319 | 113,1929 | 97,17238 | 24,61173635 | 116,6404 | 147,6368718 | 178,7065 | 320,6617 | 471,2073 | 87,9415228 |
| Upregulated | ENSMUSG00000074736 | 22,2069824 | 8,51011147 | 27,2324052 | 2,89070227 | 0 | 0 | 0 | 0 | 0 | 7,55087556 | 5,52056654 | 14,88944 | 20,0351293 | 13,0967 | 16,72639 | 0,911545791 | 10,75408 | 5,945108933 | 16,15963 | 29,99953 | 24,80038 | 0,89736248 |
| Upregulated | ENSMUSG00000045868 | 153,3339261 | 58,5070164 | 141,608507 | 45,2876689 | 10,1777 | 38,48210734 | 15,9025042 | 23,2222944 | 10,6775969 | 50,6987359 | 39,7480791 | 21,02039 | 141,247662 | 78,58018 | 84,42846 | 96,62385381 | 74,45132 | 12,88106935 | 39,92379 | 45,33263 | 147,0309 | 102,299322 |
| Upregulated | ENSMUSG00000032661 | 67,67842255 | 18,0839869 | 59,9112914 | 6,74497197 | 0 | 10,61575375 | 0 | 0 | 0 | 47,4626464 | 0 | 0 | 9,01580818 | 0 | 14,33691 | 2,734637372 | 34,74395 | 2,972554466 | 17,1102 | 1,333313 | 111,6017 | 7,17889982 |
| Upregulated | ENSMUSG00000052776 | 289,7482465 | 168,074702 | 320,434634 | 74,1946916 | 12,722125 | 27,86635359 | 9,08714527 | 15,8889383 | 5,33879843 | 51,7774324 | 33,1233993 | 28,02719 | 93,1633512 | 26,19339 | 82,83547 | 164,9897881 | 104,2318 | 23,78043573 | 82,69928 | 107,3317 | 503,0935 | 82,5573479 |
| Upregulated | ENSMUSG00000041827 | 51,81629226 | 12,7651672 | 103,48314 | 23,1256182 | 1,2722125 | 14,5966614 | 9,08714527 | 0 | 6,40655811 | 10,7869651 | 6,62467985 | 10,5102 | 26,0456681 | 17,77409 | 58,94063 | 13,67318686 | 27,29882 | 4,954257444 | 20,91246 | 29,33288 | 153,2309 | 42,1760364 |
| Upregulated | ENSMUSG00000034853 | 49,70134156 | 73,3997114 | 49,0183294 | 35,6519947 | 30,5331 | 41,13604578 | 27,2614358 | 25,6667465 | 46,9814262 | 65,8004871 | 48,5809856 | 16,64114 | 2,00351293 | 11,22574 | 30,26681 | 26,43482793 | 77,76026 | 79,2681191 | 74,14419 | 77,99878 | 120,459 | 89,7362477 |
| Upregulated | ENSMUSG00000035473 | 493,8409895 | 544,647134 | 283,217014 | 310,268711 | 339,680737 | 330,4153354 | 241,945243 | 304,334279 | 264,804402 | 320,372863 | 357,732712 | 169,9148 | 187,328459 | 95,4188 | 126,6427 | 125,7933191 | 279,6061 | 382,4686747 | 333,6488 | 321,3283 | 868,8992 | 421,760364 |
| Upregulated | ENSMUSG00000048965 | 82,48307748 | 95,7387541 | 93,4979245 | 150,316518 | 44,5274375 | 10,61575375 | 37,4844742 | 62,3335271 | 39,5071084 | 102,476168 | 34,2275126 | 63,06118 | 14,0245905 | 0 | 50,17918 | 20,96555318 | 99,26842 | 92,14918846 | 76,04532 | 101,9984 | 113,3732 | 97,81251 |
| Upregulated | ENSMUSG00000027656 | 6574,324265 | 7148,49364 | 3605,57045 | 15868,0283 | 2092,78956 | 1756,907245 | 3587,15059 | 2118,1177 | 837,123594 | 4962,00394 | 1311,68661 | 2001,317 | 3235,67338 | 1328,379 | 1085,623 | 1542,335478 | 7160,562 | 9083,135598 | 8347,875 | 8040,541 | 12625,17 | 9530,88687 |
| Upregulated | ENSMUSG00000030562 | 683,1290776 | 698,892905 | 401,224103 | 719,784866 | 231,542675 | 286,6253512 | 177,199333 | 259,111917 | 199,671061 | 382,937261 | 501,267442 | 477,3381 | 462,811487 | 654,8349 | 559,9359 | 473,0922653 | 1024,119 | 859,0682408 | 635,929 | 884,6529 | 1309,992 | 879,415228 |
| Upregulated | ENSMUSG00000062960 | 31,72426057 | 36,1679738 | 19,0626836 | 29,8705901 | 0 | 5,307876874 | 0 | 9,77780817 | 8,54207749 | 0 | 2,20822662 | 41,16494 | 17,0298599 | 14,96765 | 4,77897 | 5,469274744 | 44,67079 | 10,89936638 | 14,2585 | 24,66628 | 65,54387 | 44,8681239 |
| Upregulated | ENSMUSG00000064294 | 208,3226444 | 103,185102 | 125,269064 | 65,5225848 | 27,988675 | 13,26969219 | 10,2230384 | 74,5557873 | 35,2360696 | 72,2726661 | 71,7673651 | 123,4948 | 173,303868 | 55,19322 | 3,18598 | 10,93854949 | 14,06303 | 34,67980211 | 195,8167 | 17,99972 | 348,0911 | 233,314244 |
| Upregulated | ENSMUSG00000118590 | 27,49435916 | 7,44634754 | 27,2324052 | 7,70853939 | 3,8166375 | 3,980907656 | 5,67946579 | 0 | 4,27103874 | 6,47217906 | 3,31233993 | 2,627549 | 5,00878232 | 0 | 0 | 0,911545791 | 11,58132 | 4,954257444 | 7,604532 | 0 | 49,60077 | 30,5103242 |
| Upregulated | ENSMUSG00000025370 | 56,04619367 | 27,6578623 | 39,940861 | 17,3442136 | 0 | 10,61575375 | 0 | 26,8889725 | 4,27103874 | 48,5413429 | 11,0411331 | 13,13775 | 35,0614763 | 1,870957 | 0 | 0 | 37,22566 | 22,78958424 | 51,33059 | 17,99972 | 155,0024 | 64,6100984 |
| Upregulated | ENSMUSG00000091345 | 435,6798452 | 127,651672 | 170,656406 | 365,192054 | 31,8053125 | 41,13604578 | 2,27178632 | 154,000479 | 36,3038293 | 309,585898 | 260,570741 | 176,9216 | 306,537478 | 72,96731 | 35,84227 | 22,78864477 | 136,4941 | 169,4356046 | 419,1998 | 55,33247 | 2333,008 | 1543,46346 |
| Upregulated | ENSMUSG00000032380 | 13,74717958 | 13,8289311 | 22,693671 | 8,67210682 | 13,9943375 | 3,980907656 | 3,40767947 | 40,3334587 | 28,8295115 | 14,0230546 | 13,2493597 | 13,13775 | 5,00878232 | 2,806435 | 21,50536 | 0 | 13,23579 | 10,89936638 | 21,86303 | 23,99963 | 128,4306 | 92,4283352 |
| Upregulated | ENSMUSG00000041957 | 15,86213028 | 5,31881967 | 0 | 4,81783712 | 3,8166375 | 0 | 0 | 2,44445204 | 1,06775969 | 9,70826858 | 0 | 4,379248 | 3,00526939 | 0 | 2,389485 | 0 | 0 | 0 | 2,851699 | 8,666531 | 25,68611 | 41,278674 |
| Upregulated | ENSMUSG00000032561 | 27,49435916 | 6,3825836 | 0,90774684 | 16,3806462 | 17,810975 | 6,634846093 | 3,40767947 | 13,4444862 | 5,33879843 | 31,2821988 | 36,4357392 | 21,89624 | 15,026347 | 0 | 5,575465 | 9,115457906 | 6,617895 | 0 | 11,4068 | 27,99956 | 102,7444 | 104,094047 |
| Upregulated | ENSMUSG00000048455 | 7,402327466 | 0 | 16,3394431 | 22,1620508 | 0 | 3,980907656 | 0 | 0 | 5,33879843 | 0 | 0 | 0 | 0 | 0 | 0 | 0 | 0 | 0 | 0 | 13,33313 | 42,51494 | 1,79472495 |
| Upregulated | ENSMUSG00000030559 | 6,344852114 | 9,57387541 | 17,24719 | 36,6155621 | 1,2722125 | 6,634846093 | 0 | 17,1111643 | 1,06775969 | 4,31478604 | 11,0411331 | 0 | 14,0245905 | 5,61287 | 5,575465 | 0 | 19,02645 | 1,981702978 | 12,35736 | 19,33303 | 68,20106 | 28,7155993 |
| Upregulated | ENSMUSG00000021319 | 964,4175213 | 2270,07224 | 2536,24467 | 2077,45137 | 141,215587 | 58,38664562 | 36,3485811 | 570,779552 | 359,835014 | 283,697182 | 380,919092 | 1188,528 | 2762,84433 | 226,3858 | 152,927 | 54,69274744 | 1173,022 | 781,7818247 | 2018,053 | 981,318 | 8092,897 | 1631,40498 |
| Upregulated | ENSMUSG00000040612 | 34,89668663 | 74,4634754 | 94,4056714 | 113,700956 | 10,1777 | 18,57756906 | 4,54357263 | 34,2223286 | 26,6939921 | 19,4165372 | 30,9151726 | 46,42003 | 81,1422736 | 48,64488 | 19,91237 | 14,58473265 | 57,90658 | 36,66150509 | 113,1174 | 109,9983 | 363,1485 | 146,270084 |
| Upregulated | ENSMUSG00000025780 | 678,8991762 | 458,482256 | 1102,91241 | 681,242169 | 269,70905 | 337,0501815 | 346,447413 | 950,891845 | 863,817586 | 1348,37064 | 882,186534 | 296,913 | 388,681508 | 198,3214 | 123,4567 | 252,498184 | 167,1018 | 457,7733878 | 481,9372 | 761,3215 | 3640,342 | 1432,19051 |
| Upregulated | ENSMUSG00000079017 | 541,4273804 | 715,913128 | 1742,87393 | 288,10666 | 40,7108 | 102,1766298 | 82,9202006 | 264,000821 | 106,775969 | 409,904673 | 461,519363 | 42,04079 | 290,509375 | 36,48366 | 51,77217 | 75,65830062 | 177,8559 | 142,6826144 | 437,2606 | 475,3259 | 3849,374 | 560,851548 |
| Upregulated | ENSMUSG00000055632 | 103,6325845 | 381,891252 | 132,531039 | 164,77003 | 40,7108 | 172,5059984 | 37,4844742 | 259,111917 | 142,012038 | 126,207492 | 333,442219 | 95,46762 | 231,405743 | 117,8703 | 43,01073 | 41,93110637 | 122,4311 | 75,30471315 | 272,8126 | 330,6615 | 1637,711 | 656,869333 |
| Upregulated | ENSMUSG00000079343 | 1470,948215 | 959,515068 | 1563,14006 | 1039,68925 | 600,484299 | 985,9381294 | 1160,88281 | 1269,89284 | 1148,90942 | 1330,0328 | 1376,8293 | 1267,355 | 2069,62886 | 865,3175 | 677,8172 | 804,8949331 | 1504,744 | 1173,168163 | 2557,974 | 1217,981 | 7102,653 | 2665,16656 |
| Upregulated | ENSMUSG00000018845 | 4,229901409 | 2,12752787 | 24,5091647 | 5,78140454 | 0 | 0 | 3,40767947 | 4,88890409 | 7,4743178 | 7,55087556 | 11,0411331 | 20,14454 | 17,0298599 | 3,741914 | 0 | 2,734637372 | 19,85368 | 7,92681191 | 39,92379 | 49,99922 | 149,688 | 59,2259235 |
| Upregulated | ENSMUSG00000024653 | 13,74717958 | 0 | 29,0478989 | 1,92713485 | 2,544425 | 1,326969219 | 7,95125211 | 0 | 0 | 2,15739302 | 2,20822662 | 3,503399 | 14,0245905 | 0 | 0,796495 | 1,823091581 | 1,654474 | 2,972554466 | 19,01133 | 15,99975 | 108,9445 | 46,6628488 |
| Upregulated | ENSMUSG00000058952 | 0 | 0 | 0 | 0 | 0 | 0 | 0 | 19,5556163 | 5,33879843 | 0 | 0 | 0 | 47,0825538 | 0 | 0 | 0 | 16,54474 | 0 | 3,802266 | 25,99959 | 162,0882 | 68,1995483 |
| Upregulated | ENSMUSG00000079434 | 53,93124297 | 80,846059 | 159,763444 | 30,8341576 | 2,544425 | 0 | 1,13589316 | 2,44445204 | 0 | 4,31478604 | 19,8740396 | 90,21252 | 561,985377 | 15,90313 | 1,59299 | 10,93854949 | 53,77039 | 83,23152506 | 30,41813 | 399,9938 | 2245,321 | 299,719067 |
| Upregulated | ENSMUSG00000033066 | 151,2189754 | 101,057574 | 137,97752 | 106,955984 | 151,393287 | 130,0429834 | 136,307179 | 107,55589 | 105,708209 | 133,758367 | 196,532169 | 169,039 | 389,683265 | 136,5798 | 91,59692 | 116,6778612 | 138,1486 | 214,0239216 | 194,8661 | 385,3273 | 1009,73 | 297,924343 |
| Upregulated | ENSMUSG00000032068 | 31,72426057 | 7,44634754 | 47,2028357 | 29,8705901 | 10,1777 | 10,61575375 | 12,4948247 | 2,44445204 | 43,7781471 | 16,1804476 | 9,93701978 | 8,758497 | 82,1440301 | 48,64488 | 18,31938 | 27,34637372 | 18,19921 | 17,8353268 | 24,71473 | 34,66613 | 499,5506 | 56,5338361 |
| Upregulated | ENSMUSG00000017344 | 33,83921127 | 44,6780852 | 66,2655193 | 101,17458 | 44,5274375 | 53,07876874 | 26,1255426 | 57,444623 | 79,0142167 | 83,0596312 | 87,2249514 | 51,67513 | 191,335485 | 62,67705 | 31,0633 | 38,28492321 | 54,59763 | 45,57916849 | 98,85892 | 89,9986 | 687,3249 | 148,962171 |
| Upregulated | ENSMUSG00000005864 | 0 | 0 | 0 | 0 | 0 | 0 | 0 | 7,33335613 | 0 | 0 | 1,10411331 | 19,26869 | 7,01229525 | 4,677392 | 0 | 0 | 0 | 0,990851489 | 3,802266 | 14,66644 | 74,40115 | 12,5630747 |
| Upregulated | ENSMUSG00000032726 | 2,114950705 | 0 | 0,90774684 | 3,8542697 | 7,63327499 | 2,653938437 | 0 | 8,55558215 | 0 | 4,31478604 | 8,83290647 | 21,02039 | 17,0298599 | 0 | 0,796495 | 0 | 0,827237 | 0,990851489 | 0,950566 | 10,6665 | 113,3732 | 34,9971366 |
| Upregulated | ENSMUSG00000091649 | 19,03455634 | 21,2752787 | 20,8781773 | 14,4535114 | 1,2722125 | 25,21241515 | 5,67946579 | 9,77780817 | 4,27103874 | 6,47217906 | 6,62467985 | 2,627549 | 6,01053879 | 3,741914 | 18,31938 | 14,58473265 | 36,39842 | 3,963405955 | 4,752832 | 35,99944 | 118,6876 | 17,0498871 |
| Upregulated | ENSMUSG00000001281 | 68,7358979 | 85,1011147 | 42,6641015 | 57,8140454 | 66,1550499 | 78,2911839 | 39,7562605 | 33,0001026 | 24,5584728 | 26,9674127 | 26,4987194 | 57,80608 | 15,026347 | 26,19339 | 36,63877 | 35,55028583 | 38,05289 | 85,21322804 | 51,33059 | 111,9983 | 449,0641 | 134,604372 |
| Upregulated | ENSMUSG00000024302 | 30,66678522 | 22,3390426 | 7,26197472 | 10,5992417 | 22,899825 | 19,90453828 | 18,1742905 | 14,6667123 | 21,3551937 | 18,3378407 | 9,93701978 | 28,02719 | 19,0333728 | 14,96765 | 11,15093 | 13,67318686 | 0 | 14,86277233 | 14,2585 | 25,99959 | 126,6591 | 22,4340619 |
| Upregulated | ENSMUSG00000024511 | 2,114950705 | 5,31881967 | 10,8929621 | 5,78140454 | 17,810975 | 19,90453828 | 18,1742905 | 20,7778424 | 7,4743178 | 3,23608953 | 9,93701978 | 0 | 0 | 0 | 4,77897 | 13,67318686 | 1,654474 | 8,917663399 | 12,35736 | 58,66575 | 146,1451 | 22,4340619 |
| Upregulated | ENSMUSG00000056671 | 4,229901409 | 19,1477508 | 19,9704305 | 19,2713485 | 0 | 23,88544594 | 6,81535895 | 6,11113011 | 14,9486356 | 14,0230546 | 20,9781529 | 0 | 5,00878232 | 6,548349 | 3,982475 | 4,557728953 | 9,926842 | 23,78043573 | 5,703399 | 51,99919 | 182,46 | 19,7419745 |
| Upregulated | ENSMUSG00000079654 | 6,344852114 | 11,7014033 | 4,5387342 | 15,4170788 | 2,544425 | 11,94272297 | 19,3101837 | 3,66667807 | 11,7453565 | 6,47217906 | 1,10411331 | 1,751699 | 0 | 1,870957 | 0 | 5,469274744 | 3,308947 | 10,89936638 | 14,2585 | 40,66603 | 143,4879 | 15,2551621 |
| Upregulated | ENSMUSG00000021403 | 7,402327466 | 0 | 8,16972156 | 25,052753 | 1,2722125 | 15,92363062 | 3,40767947 | 2,44445204 | 0 | 0 | 0 | 1,751699 | 0 | 0 | 0 | 1,823091581 | 9,926842 | 5,945108933 | 10,45623 | 98,66513 | 364,0342 | 74,4810856 |
| Upregulated | ENSMUSG00000023031 | 2,114950705 | 1,06376393 | 3,63098736 | 3,8542697 | 0 | 0 | 0 | 11,0000342 | 0 | 0 | 0 | 0 | 7,01229525 | 1,870957 | 1,59299 | 0 | 8,272368 | 5,945108933 | 9,505665 | 63,999 | 215,2319 | 23,3314244 |
| Upregulated | ENSMUSG00000046056 | 35,95416198 | 5,31881967 | 53,5570636 | 43,3605341 | 0 | 34,50119968 | 3,40767947 | 4,88890409 | 22,4229534 | 1,07869651 | 5,52056654 | 7,882647 | 41,072015 | 6,548349 | 12,74392 | 10,93854949 | 28,95329 | 5,945108933 | 19,01133 | 365,3276 | 1425,136 | 178,575133 |
| Upregulated | ENSMUSG00000016756 | 88,82792959 | 86,1648786 | 81,6972156 | 67,4497197 | 242,992587 | 261,4129361 | 213,547914 | 188,222807 | 332,073262 | 241,628018 | 278,236554 | 34,15814 | 63,1106573 | 11,22574 | 2,389485 | 0 | 25,64434 | 34,67980211 | 60,83626 | 303,3286 | 2061,975 | 498,933537 |
| Upregulated | ENSMUSG00000043629 | 48,64386621 | 95,7387541 | 31,7711394 | 33,7248598 | 58,5217749 | 31,84726125 | 27,2614358 | 143,000445 | 115,318046 | 72,2726661 | 72,8714784 | 36,78569 | 49,0860668 | 6,548349 | 23,09835 | 29,1694653 | 15,7175 | 35,6706536 | 57,98456 | 145,3311 | 606,7237 | 151,654259 |
| Upregulated | ENSMUSG00000073940 | 45,47144015 | 58,5070164 | 23,6014178 | 25,052753 | 33,077525 | 25,21241515 | 29,5332221 | 73,3335613 | 67,2688602 | 15,1017511 | 30,9151726 | 29,77889 | 18,0316164 | 39,29009 | 14,33691 | 45,57728953 | 33,91671 | 30,71639615 | 20,91246 | 130,6646 | 532,3225 | 50,2522987 |

| | | | | | | | | | | | | | | | | | | | | | | | |
|---|---|---|---|---|---|---|---|---|---|---|---|---|---|---|---|---|---|---|---|---|---|---|---|
| Upregulated | ENSMUSG00000052305 | 9,517278171 | 13,8289311 | 6,35422788 | 5,78140454 | 8,90548749 | 2,653938437 | 9,08714527 | 56,222397 | 25,6262325 | 6,47217906 | 5,52056654 | 5,255098 | 6,01053879 | 10,29026 | 6,37196 | 10,93854949 | 23,16263 | 22,78958424 | 13,30793 | 121,9981 | 520,8081 | 58,328561 |
| Upregulated | ENSMUSG00000069919 | 78,25317607 | 90,4199344 | 51,7415699 | 56,850478 | 40,7108 | 27,86635359 | 32,9409016 | 152,778253 | 96,0983717 | 36,6756813 | 49,6850989 | 42,91663 | 29,0509375 | 38,35461 | 47,7897 | 51,95811007 | 124,0855 | 39,63405955 | 55,13286 | 317,3284 | 1393,25 | 154,346346 |
| Upregulated | ENSMUSG00000069917 | 44,4139648 | 42,5505574 | 22,693671 | 16,3806462 | 19,0831875 | 11,94272297 | 13,6307179 | 75,7780133 | 65,1333408 | 14,0230546 | 8,83290647 | 22,77209 | 11,0193211 | 14,96765 | 18,31938 | 17,31937002 | 47,1525 | 25,76213871 | 34,22039 | 140,6645 | 652,7815 | 82,5573479 |
| Upregulated | ENSMUSG00000035506 | 26,43688381 | 17,0202229 | 16,3394431 | 2,89070227 | 3,8166375 | 6,634846093 | 20,4460768 | 15,8889383 | 8,54207749 | 10,7869651 | 35,3316259 | 32,40644 | 13,022834 | 2,806435 | 8,761444 | 18,23091581 | 4,963421 | 9,908514888 | 16,15963 | 65,99897 | 401,2348 | 90,6336102 |
| Upregulated | ENSMUSG00000075254 | 1388,465138 | 1147,80128 | 1020,30745 | 811,323771 | 826,938124 | 1182,329574 | 1252,89015 | 1097,55897 | 1063,48865 | 1259,91752 | 1331,56065 | 811,9127 | 1217,1341 | 1258,218 | 715,2525 | 563,3352986 | 517,023 | 843,214617 | 943,9125 | 1766,639 | 7200,969 | 1732,80694 |
| Upregulated | ENSMUSG00000024379 | 10,57475352 | 18,0839869 | 29,9556457 | 13,4899439 | 40,7108 | 7,961815312 | 23,8537563 | 30,5556505 | 27,7617518 | 21,5739302 | 28,706946 | 33,28229 | 85,1492995 | 25,25792 | 21,50536 | 25,52328214 | 101,7501 | 148,6277233 | 38,97323 | 217,3299 | 787,4122 | 35,8944991 |
| Upregulated | ENSMUSG00000118000 | 3,172426057 | 7,44634754 | 9,0774684 | 3,8542697 | 6,36106249 | 3,980907656 | 9,08714527 | 3,66667807 | 8,54207749 | 3,23608953 | 4,41645324 | 7,006798 | 18,0316164 | 4,677392 | 13,54041 | 6,380820534 | 12,40855 | 9,908514888 | 11,4068 | 38,66606 | 112,4875 | 15,2551621 |
| Upregulated | ENSMUSG00000021822 | 127,9545176 | 132,970492 | 168,840912 | 142,607979 | 111,9547 | 106,1575375 | 109,045743 | 202,88952 | 146,283077 | 152,096208 | 216,406209 | 218,0866 | 334,586659 | 100,0962 | 88,41094 | 166,8128797 | 253,1345 | 330,9443973 | 448,6674 | 673,3228 | 3069,048 | 603,924947 |
| Upregulated | ENSMUSG00000032369 | 39,12658804 | 81,9098229 | 71,7120004 | 49,1419386 | 21,6276125 | 58,38664562 | 44,2998332 | 40,3334587 | 69,4043796 | 61,485701 | 81,7043849 | 153,2737 | 207,363588 | 73,90279 | 46,9932 | 59,25047639 | 122,4311 | 113,9479212 | 127,3759 | 331,9948 | 1112,474 | 271,003468 |
| Upregulated | ENSMUSG00000002847 | 95,17278171 | 93,6112262 | 151,593722 | 58,7776129 | 30,5331 | 18,57756906 | 18,1742905 | 108,778116 | 104,640449 | 262,123252 | 136,91005 | 136,6326 | 120,210776 | 164,6442 | 125,0497 | 57,42738481 | 367,2932 | 171,4173076 | 192,965 | 474,6593 | 2446,381 | 397,531578 |
| Upregulated | ENSMUSG00000041261 | 1,057475352 | 1,06376393 | 8,16972156 | 1,92713485 | 6,36106249 | 10,61575375 | 2,27178632 | 28,1111985 | 3,20327906 | 21,5739302 | 4,41645324 | 19,26869 | 29,0509375 | 18,70957 | 2,389485 | 6,380820534 | 0 | 7,92681191 | 16,15963 | 44,66597 | 276,3471 | 30,5103242 |
| Upregulated | ENSMUSG00000018916 | 9,517278171 | 7,44634754 | 9,98521524 | 3,8542697 | 5,08884999 | 0 | 15,9025042 | 6,11113011 | 12,8131162 | 3,23608953 | 36,4357392 | 88,46082 | 218,382909 | 56,1287 | 19,11588 | 21,87709898 | 40,53461 | 125,8381391 | 172,0525 | 656,6564 | 5333,854 | 336,510929 |
| Upregulated | ENSMUSG00000060962 | 75,08075001 | 1,06376393 | 6,35422788 | 92,5024727 | 0 | 65,02149171 | 5,67946579 | 24,4445204 | 69,4043796 | 0 | 0 | 0 | 90,1580818 | 0 | 0 | 9,115457906 | 0,827237 | 3,963405955 | 0,950566 | 368,6609 | 2661,613 | 302,411155 |
| Upregulated | ENSMUSG00000085721 | 10,57475352 | 2,12752787 | 1,81549368 | 3,8542697 | 2,544425 | 10,61575375 | 3,40767947 | 6,11113011 | 2,13551937 | 7,55087556 | 5,52056654 | 9,634347 | 12,0210776 | 4,677392 | 6,37196 | 4,557728953 | 14,06303 | 9,908514888 | 3,802266 | 34,66613 | 98,31581 | 27,8182368 |
| Upregulated | ENSMUSG00000071714 | 1,057475352 | 0 | 0 | 3,8542697 | 0 | 7,961815312 | 2,27178632 | 4,88890409 | 6,40655811 | 0 | 1,10411331 | 10,5102 | 38,0667457 | 5,61287 | 7,964949 | 0 | 1,654474 | 0,990851489 | 22,8136 | 77,99878 | 255,0897 | 75,3784481 |
| Upregulated | ENSMUSG00000032372 | 3,172426057 | 1,06376393 | 1,81549368 | 4,81783712 | 0 | 5,307876874 | 0 | 0 | 4,27103874 | 1,07869651 | 0 | 0 | 18,0316164 | 1,870957 | 0 | 0 | 12,40855 | 0,990851489 | 19,01133 | 38,66606 | 140,8308 | 29,6129618 |
| Upregulated | ENSMUSG00000019832 | 465,289155 | 385,082544 | 364,006483 | 310,268711 | 270,981262 | 193,7375059 | 170,383974 | 386,223423 | 253,059045 | 443,344265 | 492,434536 | 231,2243 | 374,656918 | 292,8047 | 358,4227 | 278,0214661 | 419,4091 | 319,0541794 | 554,1803 | 666,6563 | 2448,152 | 871,338966 |
| Upregulated | ENSMUSG00000010797 | 106,8050106 | 104,248866 | 140,70076 | 195,604187 | 7,63327499 | 19,90453828 | 0 | 67,2224312 | 32,0327906 | 48,5413429 | 54,1015521 | 25,39964 | 51,0895797 | 32,74174 | 8,761444 | 0 | 52,94316 | 10,89936638 | 87,45212 | 317,3284 | 1263,048 | 323,947854 |
| Upregulated | ENSMUSG00000042485 | 65,56347184 | 47,869377 | 123,45357 | 48,1783712 | 25,44425 | 25,21241515 | 13,6307179 | 40,3334587 | 39,5071084 | 32,3608953 | 55,2056654 | 46,42003 | 57,1001185 | 34,6127 | 32,65629 | 38,28492321 | 86,03263 | 69,35960422 | 69,39135 | 200,6635 | 760,8404 | 253,953581 |
| Upregulated | ENSMUSG00000027068 | 31,72426057 | 6,3825836 | 28,140152 | 20,2349159 | 0 | 2,653938437 | 0 | 6,11113011 | 0 | 2,15739302 | 0 | 28,90304 | 23,0403987 | 0 | 1,59299 | 21,87709898 | 12,40855 | 1,981702978 | 40,87436 | 105,9983 | 393,2632 | 121,143934 |
| Upregulated | ENSMUSG00000024053 | 8649,090907 | 5853,89293 | 8713,46192 | 4268,60369 | 4156,31823 | 5088,926953 | 4400,4501 | 3889,1232 | 5029,14812 | 5840,0629 | 3995,78606 | 2477,779 | 2298,02933 | 906,4786 | 1444,045 | 1558,743302 | 2235,194 | 2930,938704 | 5566,517 | 6191,237 | 13130,03 | 5260,33884 |
| Upregulated | ENSMUSG00000022053 | 318,300081 | 234,028065 | 449,334686 | 370,009891 | 166,659837 | 156,5823678 | 173,791653 | 267,667499 | 167,638271 | 376,465082 | 167,825223 | 98,97102 | 148,259957 | 67,35444 | 60,53362 | 21,87709898 | 78,5875 | 90,16748548 | 185,3605 | 429,3266 | 770,5834 | 318,563679 |
| Upregulated | ENSMUSG00000038530 | 564,6918381 | 287,216262 | 681,717877 | 584,885426 | 72,5161124 | 58,38664562 | 89,7355595 | 111,222568 | 51,2524649 | 254,572376 | 229,655568 | 185,6801 | 162,284547 | 50,51583 | 99,56187 | 96,62385381 | 204,3275 | 396,3405955 | 572,241 | 741,9884 | 1326,821 | 497,138812 |
| Upregulated | ENSMUSG00000033208 | 33,83921127 | 7,44634754 | 55,3725572 | 28,9070227 | 0 | 3,980907656 | 7,95125211 | 22,0000684 | 6,40655811 | 30,2035023 | 8,83290647 | 3,503399 | 13,022834 | 7,483827 | 1,59299 | 2,734637372 | 5,790658 | 14,86277233 | 35,17096 | 24,66628 | 75,28688 | 26,9208743 |
| Upregulated | ENSMUSG00000049929 | 84,59802819 | 58,5070164 | 210,597267 | 158,988625 | 29,2608875 | 37,15513812 | 55,6587648 | 81,8891435 | 105,708209 | 94,9252928 | 99,3701978 | 28,02719 | 57,1001185 | 20,58052 | 29,47031 | 12,76164107 | 79,41474 | 51,52427742 | 146,3872 | 189,3304 | 286,0901 | 102,299322 |
| Upregulated | ENSMUSG00000048520 | 26,43688381 | 7,44634754 | 25,4169115 | 45,2876689 | 1,2722125 | 9,28878453 | 1,13589316 | 26,8889725 | 6,40655811 | 24,8100197 | 11,0411331 | 0 | 4,00702586 | 5,61287 | 1,59299 | 6,380820534 | 19,02645 | 14,86277233 | 19,9619 | 51,99919 | 56,68659 | 26,9208743 |
| Upregulated | ENSMUSG00000041945 | 31,72426057 | 9,57387541 | 21,7859242 | 26,0163204 | 0 | 21,2315075 | 1,13589316 | 6,11113011 | 7,4743178 | 7,55087556 | 3,31233993 | 11,38605 | 9,01580818 | 14,03218 | 11,94742 | 8,203912116 | 21,50816 | 7,92681191 | 16,15963 | 30,66619 | 39,85776 | 17,0498871 |
| Upregulated | ENSMUSG00000021668 | 223,1272993 | 132,970492 | 265,062077 | 108,883119 | 94,1437249 | 110,1384451 | 209,004341 | 154,000479 | 106,775969 | 250,25759 | 77,2879316 | 42,04079 | 36,0632327 | 28,99983 | 134,6076 | 66,54284272 | 100,0957 | 79,2681191 | 119,7714 | 192,6637 | 126,6591 | 87,9415228 |
| Upregulated | ENSMUSG00000045257 | 28,55183451 | 17,0202229 | 48,1105825 | 21,1984833 | 8,90548749 | 7,961815312 | 29,5332221 | 30,5556505 | 10,6775969 | 24,8100197 | 2,20822662 | 7,006798 | 12,0210776 | 9,354784 | 5,575465 | 11,85009528 | 11,58132 | 7,92681191 | 26,61586 | 32,66616 | 21,25747 | 8,97362477 |
| Upregulated | ENSMUSG00000031842 | 10,57475352 | 2,12752787 | 31,7711394 | 0 | 2,544425 | 6,634846093 | 10,2230384 | 4,88890409 | 4,27103874 | 10,7869651 | 3,31233993 | 0 | 2,00351293 | 0 | 1,59299 | 0 | 14,89026 | 0,990851489 | 7,604532 | 33,99947 | 2,657184 | 1,79472495 |
| Upregulated | ENSMUSG00000024169 | 500,1858416 | 347,850806 | 398,500863 | 255,345367 | 279,88675 | 244,1623362 | 385,067781 | 233,44517 | 264,804402 | 319,294167 | 149,055297 | 119,9914 | 181,31792 | 45,83844 | 122,6602 | 73,83520904 | 263,8886 | 219,9690305 | 315,5881 | 520,6585 | 198,4031 | 251,261494 |
| Upregulated | ENSMUSG00000022364 | 208,3226444 | 147,863187 | 195,165571 | 59,7411803 | 90,3270874 | 95,54178374 | 101,094491 | 123,444828 | 111,047007 | 79,8235417 | 51,8933255 | 24,52379 | 33,0579633 | 43,03201 | 82,83547 | 32,81564846 | 83,55092 | 156,5545352 | 135,931 | 200,6635 | 123,1162 | 100,504597 |
| Upregulated | ENSMUSG00000035878 | 136,4143204 | 86,1648786 | 142,516254 | 55,8869106 | 76,3327499 | 21,2315075 | 80,6484142 | 112,444794 | 53,3879843 | 77,6661487 | 72,8714784 | 35,90984 | 11,0193211 | 13,0967 | 32,65629 | 1,823091581 | 35,57118 | 78,27726762 | 63,68796 | 152,6643 | 83,25843 | 75,3784481 |
| Upregulated | ENSMUSG00000031066 | 346,8519156 | 159,56459 | 426,641015 | 294,851632 | 171,748687 | 193,7375059 | 202,188982 | 237,111848 | 217,822976 | 327,923739 | 94,9537446 | 95,46762 | 76,1334913 | 113,1929 | 39,02825 | 66,54284272 | 100,0957 | 189,2526344 | 122,6231 | 265,9958 | 178,0313 | 141,783271 |
| Upregulated | ENSMUSG00000040350 | 22,2069824 | 8,51011147 | 24,5091647 | 4,81783712 | 10,1777 | 17,25059984 | 2,27178632 | 9,77780817 | 12,8131162 | 17,2591441 | 1,10411331 | 4,379248 | 4,00702586 | 0,935478 | 1,59299 | 0,911545791 | 2,481711 | 5,945108933 | 5,703399 | 20,66634 | 2,657184 | 2,69208743 |
| Upregulated | ENSMUSG00000051977 | 93,057831 | 68,0808918 | 84,4204561 | 68,4132871 | 62,3384124 | 22,55847672 | 28,397329 | 64,7779792 | 16,0163953 | 43,1478604 | 70,6632518 | 28,90304 | 14,0245905 | 12,16122 | 36,63877 | 17,31937002 | 22,33539 | 18,82617829 | 29,46756 | 75,99881 | 37,20058 | 27,8182368 |
| Upregulated | ENSMUSG00000034706 | 41,24153874 | 17,0202229 | 36,3098736 | 19,2713485 | 2,544425 | 9,28878453 | 11,3589316 | 25,6667465 | 1,06775969 | 38,8330743 | 43,060419 | 2,627549 | 4,00702586 | 0 | 4,77897 | 3,646183163 | 3,308947 | 7,92681191 | 16,15963 | 43,99931 | 0,885728 | 4,48681239 |
| Upregulated | ENSMUSG00000040170 | 1076,509909 | 520,180564 | 1108,35889 | 565,614078 | 81,4215999 | 35,8281689 | 26,1255426 | 524,334963 | 242,381449 | 155,332297 | 198,740396 | 91,96422 | 571,001185 | 144,9992 | 68,49857 | 44,66574374 | 155,5205 | 80,25897059 | 251,9001 | 1383,978 | 853,8418 | 925,180714 |
| Upregulated | ENSMUSG00000022206 | 49,70134156 | 40,4230295 | 105,298633 | 29,8705901 | 2,544425 | 1,326969219 | 6,81535895 | 23,2222944 | 2,13551937 | 19,4165372 | 16,5616996 | 4,379248 | 42,0737715 | 21,516 | 2,389485 | 5,469274744 | 0,827237 | 9,908514888 | 24,71473 | 59,99906 | 35,42912 | 18,844612 |
| Upregulated | ENSMUSG00000024253 | 148,0465493 | 138,289311 | 186,088102 | 217,766238 | 85,2382374 | 71,65633781 | 113,589316 | 114,889246 | 134,53772 | 116,499223 | 92,7455179 | 53,42683 | 73,1282219 | 54,25775 | 58,14413 | 52,86965586 | 62,04276 | 79,2681191 | 102,6612 | 183,3305 | 146,1451 | 91,5309727 |
| Upregulated | ENSMUSG00000009894 | 689,4739297 | 799,950478 | 724,381978 | 968,385261 | 419,830125 | 407,3795501 | 436,182973 | 496,223765 | 416,426277 | 610,542224 | 358,836825 | 181,3009 | 343,602467 | 384,4816 | 270,8083 | 196,8938908 | 311,8683 | 348,7797241 | 365,0175 | 758,6548 | 724,5255 | 321,255767 |
| Upregulated | ENSMUSG00000022323 | 80,36812678 | 142,544367 | 229,659951 | 123,33663 | 55,9773499 | 82,27209155 | 80,6484142 | 124,667054 | 136,67324 | 87,3744172 | 80,6002715 | 14,88944 | 67,1176831 | 25,25792 | 16,72639 | 28,25791951 | 16,54474 | 79,2681191 | 8,555098 | 179,9972 | 121,3447 | 60,123286 |
| Upregulated | ENSMUSG00000037855 | 72,96579931 | 62,7620721 | 89,8669372 | 59,7411803 | 3,8166375 | 10,61575375 | 27,2614358 | 41,5556847 | 32,0327906 | 64,7217906 | 45,2686457 | 19,26869 | 37,0649892 | 0,935478 | 2,389485 | 22,78864477 | 14,89026 | 48,55172295 | 54,18229 | 87,33197 | 44,2864 | 10,7683497 |
| Upregulated | ENSMUSG00000038122 | 172,3684824 | 130,842964 | 134,346532 | 121,409495 | 105,593637 | 86,25299921 | 193,101837 | 169,889417 | 140,944279 | 59,328308 | 88,3290647 | 40,28909 | 63,1106573 | 43,96748 | 57,34764 | 32,81564846 | 74,45132 | 108,9936638 | 109,3151 | 170,664 | 118,6876 | 92,4283352 |
| Upregulated | ENSMUSG00000054057 | 34,89668663 | 37,2317377 | 45,387342 | 43,3605341 | 24,1720375 | 22,55847672 | 43,16394 | 31,7778766 | 16,0163953 | 9,70826858 | 27,6028327 | 0 | 9,01580818 | 2,806435 | 7,964949 | 3,646183163 | 24,81711 | 14,86277233 | 22,8136 | 51,33253 | 14,17165 | 18,844612 |
| Upregulated | ENSMUSG00000097908 | 58,16114438 | 61,6983082 | 38,1253673 | 26,9798879 | 19,0831875 | 1,326969219 | 35,2126879 | 59,8890751 | 21,3551937 | 22,6526267 | 11,0411331 | 0 | 2,00351293 | 0 | 14,33691 | 4,557728953 | 9,099605 | 9,908514888 | 19,9619 | 88,66528 | 22,1432 | 9,87098725 |
| Upregulated | ENSMUSG00000027330 | 303,4954261 | 154,24577 | 285,032508 | 169,587867 | 197,192937 | 177,8138753 | 354,398665 | 233,44517 | 209,280898 | 214,660605 | 186,595149 | 51,67513 | 38,0667457 | 48,64488 | 93,18991 | 87,5083959 | 185,3011 | 217,9873275 | 159,6952 | 526,6584 | 186,0029 | 174,088321 |
| Upregulated | ENSMUSG00000058290 | 141,7016972 | 144,671895 | 177,918381 | 156,097923 | 166,659837 | 195,0644751 | 210,140234 | 113,66702 | 114,250286 | 221,132784 | 145,742957 | 16,64114 | 18,0316164 | 40,22557 | 51,77217 | 42,84265216 | 196,0551 | 229,8775454 | 214,828 | 386,6606 | 129,3163 | 149,859534 |
| Upregulated | ENSMUSG00000049811 | 100,4601585 | 58,5070164 | 64,4500256 | 45,2876689 | 36,8941625 | 23,88544594 | 89,7355595 | 45,2223628 | 20,287434 | 61,485701 | 52,9974388 | 8,758497 | 36,0632327 | 18,70957 | 19,91237 | 10,0270037 | 42,18908 | 55,48768337 | 75,09475 | 141,9978 | 86,80135 | 87,9415228 |
| Upregulated | ENSMUSG00000053219 | 11,63222888 | 1,06376393 | 22,693671 | 9,63567424 | 1,2722125 | 2,653938437 | 2,27178632 | 3,66667807 | 4,27103874 | 24,8100197 | 11,0411331 | 0 | 1,00175646 | 0 | 2,389485 | 3,646183163 | 3,308947 | 23,78043573 | 3,802266 | 39,99938 | 19,48602 | 9,87098725 |
| Upregulated | ENSMUSG00000052316 | 808,9686445 | 314,874124 | 1543,16963 | 1003,07369 | 170,476475 | 249,4702131 | 475,939233 | 707,668867 | 401,477642 | 2241,53135 | 696,695498 | 313,5542 | 52,0913362 | 80,45114 | 106,7303 | 144,0242349 | 746,1676 | 1198,930301 | 659,6931 | 3563,278 | 961,0149 | 882,107315 |
| Upregulated | ENSMUSG00000045930 | 89,88540495 | 37,2317377 | 129,807798 | 116,591658 | 8,90548749 | 51,75179953 | 29,5332221 | 30,5556505 | 19,2196743 | 67,9578801 | 54,1015521 | 42,91663 | 16,0281034 | 9,354784 | 50,97568 | 35,55028583 | 65,35171 | 108,0028123 | 42,77549 | 121,9981 | 91,22999 | 78,967898 |
| Upregulated | ENSMUSG00000047003 | 61,33357043 | 25,5303344 | 57,1880509 | 89,6117704 | 21,6276125 | 13,26969219 | 14,7666111 | 22,0000684 | 34,1683099 | 84,1383277 | 7,72879316 | 20,14454 | 12,0210776 | 4,677392 | 13,54041 | 1,823091581 | 44,67079 | 14,86277233 | 38,97323 | 78,66544 | 15,05738 | 36,7918616 |
| Upregulated | ENSMUSG00000022178 | 60,27609508 | 68,0808918 | 62,634532 | 78,0489613 | 39,4385875 | 33,17423047 | 11,3589316 | 19,5556163 | 19,2196743 | 32,3608953 | 35,3316259 | 28,02719 | 1,00175646 | 23,38696 | 27,87732 | 10,93854949 | 8,272368 | 17,8353268 | 24,71473 | 128,6647 | 15,9431 | 30,5103242 |
| Upregulated | ENSMUSG00000034413 | 14,80465493 | 15,956459 | 23,6014178 | 26,0163204 | 7,63327499 | 5,307876874 | 2,27178632 | 25,6667465 | 0 | 19,4165372 | 11,0411331 | 0 | 0 | 2,806435 | 0 | 0 | 19,02645 | 15,85362382 | 0,950566 | 21,333 | 9,743008 | 13,4604372 |
| Upregulated | ENSMUSG00000111375 | 15,86213028 | 29,7853902 | 29,0478989 | 34,6884273 | 21,6276125 | 10,61575375 | 23,8537563 | 24,4445204 | 23,4907131 | 33,4395918 | 32,019286 | 3,503399 | 4,00702586 | 0,935478 | 0,796495 | 5,469274744 | 28,12605 | 41,61576253 | 6,653965 | 37,99941 | 14,17165 | 24,2287869 |
| Upregulated | ENSMUSG00000007646 | 31,72426057 | 25,5303344 | 22,693671 | 31,797725 | 0 | 5,307876874 | 5,67946579 | 8,55558215 | 11,7453565 | 2,15739302 | 9,93701978 | 8,758497 | 0 | 1,870957 | 9,557939 | 0,911545791 | 28,12605 | 24,77128722 | 21,86303 | 48,66591 | 11,51446 | 6,28153734 |
| Upregulated | ENSMUSG00000027353 | 67,67842255 | 71,2721836 | 28,140152 | 49,1419386 | 1,2722125 | 19,90453828 | 11,3589316 | 31,7778766 | 22,4229534 | 35,5969848 | 29,8110593 | 28,90304 | 20,0351293 | 1,870957 | 12,74392 | 34,63874004 | 61,21553 | 53,5059804 | 22,8136 | 67,99894 | 36,31485 | 18,844612 |
| Upregulated | ENSMUSG00000047415 | 21,14950705 | 3,1912918 | 29,9556457 | 2,89070227 | 5,08884999 | 11,94272297 | 5,67946579 | 2,44445204 | 12,8131162 | 35,5969848 | 4,41645324 | 11,38605 | 2,00351293 | 0 | 15,1334 | 21,87709898 | 48,80697 | 32,69809913 | 55,13286 | 18,66638 | 60,22951 | 20,639337 |
| Upregulated | ENSMUSG00000027750 | 21694,10685 | 5857,08422 | 19200,6612 | 9849,58621 | 2469,36446 | 1358,81648 | 4072,17697 | 4901,12635 | 3621,84085 | 10779,4142 | 6423,73123 | 7196,857 | 5069,88947 | 6659,671 | 3427,318 | 3826,669229 | 19385,47 | 13222,91312 | 14102,6 | 17657,72 | 23671,08 | 10599,6456 |
| Upregulated | ENSMUSG00000019734 | 33,83921127 | 12,7651672 | 18,1549368 | 26,9798879 | 7,63327499 | 5,307876874 | 5,67946579 | 20,7778424 | 6,40655811 | 18,3378407 | 8,83290647 | 7,882647 | 20,0351293 | 3,741914 | 21,50536 | 13,67318686 | 28,12605 | 31,70724764 | 41,82493 | 29,99953 | 60,22951 | 24,2287869 |
| Upregulated | ENSMUSG00000026023 | 17,97708099 | 10,6376393 | 36,3098736 | 18,3077811 | 12,722125 | 13,26969219 | 36,3485811 | 24,4445204 | 16,0163953 | 32,3608953 | 29,8110593 | 0 | 2,00351293 | 0 | 1,59299 | 2,734637372 | 14,06303 | 57,46938635 | 29,46756 | 28,66622 | 36,31485 | 31,4076867 |
| Upregulated | ENSMUSG00000037994 | 59,21861973 | 18,0839869 | 15,4316963 | 6,74497197 | 5,08884999 | 15,92363062 | 18,1742905 | 0 | 1,06775969 | 59,328308 | 37,5398525 | 21,89624 | 17,0298599 | 13,0967 | 8,761444 | 3,646183163 | 26,47158 | 66,38704975 | 37,07209 | 41,99934 | 38,0863 | 43,0733989 |
| Upregulated | ENSMUSG00000042766 | 30,66678522 | 40,4230295 | 97,1289119 | 38,542697 | 30,5331 | 5,307876874 | 24,9896495 | 4,88890409 | 10,6775969 | 11,8656616 | 32,019286 | 5,255098 | 17,0298599 | 22,45148 | 23,09835 | 16,40782423 | 40,53461 | 43,59746551 | 18,06076 | 67,99894 | 31,00048 | 17,9472495 |
| Upregulated | ENSMUSG00000030610 | 71,90832396 | 71,2721836 | 106,20638 | 62,6318826 | 72,5161124 | 42,463015 | 45,4357263 | 28,1111985 | 34,1683099 | 26,9674127 | 16,5616996 | 19,26869 | 23,0403987 | 57,06418 | 59,73712 | 33,72719425 | 86,03263 | 44,588317 | 71,29249 | 116,6648 | 74,40115 | 54,7391111 |
| Upregulated | ENSMUSG00000021176 | 9,517278171 | 17,0202229 | 25,4169115 | 4,81783712 | 0 | 25,21241515 | 7,95125211 | 7,33335613 | 1,06775969 | 0 | 1,10411331 | 2,627549 | 2,00351293 | 0 | 3,18598 | 0 | 5,790658 | 27,74384169 | 33,26983 | 52,66584 | 15,9431 | 16,1525246 |
| Upregulated | ENSMUSG00000046295 | 20,09203169 | 20,2115147 | 31,7711394 | 3,8542697 | 3,8166375 | 30,52029203 | 2,27178632 | 15,8889383 | 6,40655811 | 0 | 0 | 0 | 0 | 0 | 0,796495 | 0 | 7,445132 | 44,588317 | 44,67663 | 116,6648 | 29,22902 | 17,0498871 |
| Upregulated | ENSMUSG00000019845 | 30,66678522 | 15,956459 | 37,2176204 | 19,2713485 | 8,90548749 | 14,5966614 | 14,7666111 | 33,0001026 | 14,9486356 | 5,39348255 | 15,4575863 | 3,503399 | 10,0175646 | 12,16122 | 7,964949 | 4,557728953 | 21,50816 | 50,53342593 | 33,26983 | 77,33213 | 26,57184 | 20,639337 |
| Upregulated | ENSMUSG00000046179 | 20,09203169 | 20,2115147 | 25,4169115 | 8,67210682 | 0 | 11,94272297 | 0 | 15,8889383 | 5,33879843 | 3,23608953 | 3,31233993 | 7,006798 | 5,00878232 | 0 | 0,796495 | 0 | 12,40855 | 32,69809913 | 27,56643 | 51,33253 | 6,200096 | 6,28153734 |
| Upregulated | ENSMUSG00000041064 | 16,91960564 | 19,1477508 | 21,7859242 | 0 | 0 | 5,307876874 | 0 | 8,55558215 | 5,33879843 | 3,23608953 | 0 | 0 | 0 | 0 | 0 | 0 | 0 | 15,85362382 | 15,20906 | 43,99931 | 25,68611 | 7,17889982 |
| Upregulated | ENSMUSG00000027715 | 149,1040247 | 171,265993 | 382,16142 | 113,700956 | 13,9943375 | 30,52029203 | 65,8818032 | 123,444828 | 80,0819764 | 39,9117708 | 29,8110593 | 10,5102 | 7,01229525 | 17,77409 | 11,94742 | 0 | 153,8661 | 354,724833 | 300,379 | 556,658 | 325,9479 | 240,493144 |
| Upregulated | ENSMUSG00000022033 | 82,48307748 | 114,886505 | 218,766988 | 118,518793 | 3,8166375 | 10,61575375 | 79,5125211 | 105,111438 | 58,7267827 | 55,013522 | 3,31233993 | 7,006798 | 3,00526939 | 5,61287 | 6,37196 | 0 | 236,5897 | 374,5418628 | 281,3677 | 595,324 | 249,7753 | 229,724794 |
| Upregulated | ENSMUSG00000046341 | 111,034912 | 140,416839 | 260,523343 | 114,664523 | 71,2438999 | 58,38664562 | 97,6868116 | 166,222739 | 144,147558 | 87,3744172 | 41,9563057 | 15,76529 | 25,0439116 | 27,12887 | 39,82475 | 20,05400739 | 311,8683 | 388,4137836 | 288,0216 | 513,3253 | 310,0048 | 242,287869 |
| Upregulated | ENSMUSG00000027699 | 102,5751092 | 142,544367 | 309,541672 | 95,393175 | 13,9943375 | 39,80907656 | 134,035393 | 160,111609 | 92,8950927 | 67,9578801 | 23,1863795 | 69,19213 | 12,0210776 | 21,516 | 15,1334 | 20,05400739 | 250,6528 | 285,3652288 | 350,759 | 515,3253 | 177,1456 | 270,106106 |
| Upregulated | ENSMUSG00000050107 | 0 | 11,7014033 | 24,5091647 | 2,89070227 | 1,2722125 | 0 | 2,27178632 | 3,66667807 | 3,20327906 | 3,23608953 | 0 | 1,751699 | 1,00175646 | 0 | 0,796495 | 0 | 0 | 27,74384169 | 9,505665 | 25,99959 | 11,51446 | 8,97362477 |
| Upregulated | ENSMUSG00000028832 | 111,034912 | 218,071606 | 250,538128 | 164,77003 | 63,6106249 | 79,61815312 | 112,453423 | 78,2224654 | 86,4885345 | 42,0691639 | 76,1838183 | 18,39284 | 35,0614763 | 16,83861 | 39,02825 | 29,1694653 | 373,0838 | 387,4229321 | 199,619 | 358,6611 | 448,1784 | 120,246572 |
| Upregulated | ENSMUSG00000026278 | 203,0352676 | 203,178911 | 335,866331 | 161,879327 | 45,79965 | 71,65633781 | 188,558264 | 256,667465 | 194,332263 | 163,961869 | 141,326504 | 73,57137 | 178,312651 | 46,77392 | 98,76537 | 203,2747113 | 530,2588 | 617,3004775 | 505,7014 | 755,3215 | 729,8399 | 362,534441 |
| Upregulated | ENSMUSG00000021714 | 11,63222888 | 31,912918 | 42,6641015 | 23,1256182 | 15,26655 | 1,326969219 | 22,7178632 | 9,77780817 | 24,5584728 | 8,62957207 | 5,52056654 | 5,255098 | 12,0210776 | 2,806435 | 4,77897 | 5,469274744 | 47,1525 | 55,48768337 | 45,62719 | 67,99894 | 30,11475 | 25,1261494 |
| Upregulated | ENSMUSG00000082279 | 34,89668663 | 36,1679738 | 36,3098736 | 43,3605341 | 16,5387625 | 10,61575375 | 28,397329 | 20,7778424 | 18,1519147 | 28,0461092 | 12,1452464 | 13,13775 | 41,072015 | 24,32244 | 21,50536 | 16,40782423 | 63,69724 | 77,28641613 | 55,13286 | 92,66522 | 41,62922 | 23,3314244 |
| Upregulated | ENSMUSG00000074867 | 26,43688381 | 46,8056131 | 58,0957978 | 26,9798879 | 8,90548749 | 14,5966614 | 17,0383974 | 24,4445204 | 17,084155 | 28,0461092 | 2,20822662 | 11,38605 | 3,00526939 | 2,806435 | 15,1334 | 7,292366325 | 42,18908 | 26,7529902 | 32,31926 | 69,99891 | 26,57184 | 13,4604372 |
| Upregulated | ENSMUSG00000028212 | 49,70134156 | 139,353075 | 118,914836 | 99,2474447 | 36,8941625 | 33,17423047 | 57,9305511 | 64,7779792 | 87,5562942 | 111,10574 | 20,9781529 | 17,51699 | 37,0649892 | 14,03218 | 53,36516 | 34,63874004 | 116,6404 | 99,08514888 | 137,8321 | 244,6628 | 98,31581 | 55,6364736 |
| Upregulated | ENSMUSG00000113836 | 54,98871832 | 59,5707803 | 60,8190383 | 47,2148038 | 26,7164625 | 18,57756906 | 49,979299 | 70,8891093 | 40,5748681 | 47,4626464 | 25,3946061 | 7,006798 | 5,00878232 | 5,61287 | 34,24928 | 3,646183163 | 67,83342 | 58,46023784 | 68,44079 | 93,99853 | 36,31485 | 44,8681239 |
| Upregulated | ENSMUSG00000017550 | 24,3219331 | 41,4867934 | 67,1732662 | 31,797725 | 29,2608875 | 34,50119968 | 21,58197 | 33,0001026 | 32,0327906 | 28,0461092 | 25,3946061 | 9,634347 | 5,00878232 | 4,677392 | 15,9299 | 6,380820534 | 73,62408 | 48,55172295 | 66,53965 | 76,66547 | 36,31485 | 26,9208743 |
| Upregulated | ENSMUSG00000024791 | 7,402327466 | 22,3390426 | 46,2950888 | 10,5992417 | 3,8166375 | 3,980907656 | 14,7666111 | 25,6667465 | 26,6939921 | 19,4165372 | 1,10411331 | 0 | 10,0175646 | 0 | 3,982475 | 2,734637372 | 76,93303 | 53,5059804 | 67,49022 | 79,3321 | 39,85776 | 41,278674 |
| Upregulated | ENSMUSG00000069270 | 4,229901409 | 14,8926951 | 18,1549368 | 1,92713485 | 2,544425 | 6,634846093 | 4,54357263 | 9,77780817 | 8,54207749 | 7,55087556 | 2,20822662 | 1,751699 | 0 | 0 | 4,77897 | 0,911545791 | 38,88013 | 14,86277233 | 39,92379 | 39,99938 | 20,37174 | 11,6657122 |
| Upregulated | ENSMUSG00000028718 | 12,68970423 | 53,1881967 | 109,837368 | 24,0891856 | 6,36106249 | 30,52029203 | 39,7562605 | 52,5557189 | 27,7617518 | 55,013522 | 8,83290647 | 68,31628 | 8,01405172 | 7,483827 | 7,168455 | 10,0270037 | 123,2583 | 75,30471315 | 111,2163 | 174,6639 | 63,77242 | 93,3256977 |
| Upregulated | ENSMUSG00000072980 | 33,83921127 | 41,4867934 | 37,2176204 | 7,70853939 | 5,08884999 | 9,28878453 | 20,4460768 | 11,0000342 | 4,27103874 | 2,15739302 | 6,62467985 | 11,38605 | 7,01229525 | 11,22574 | 8,761444 | 10,0270037 | 88,51434 | 41,61576253 | 74,14419 | 113,3316 | 41,62922 | 32,3050492 |

| | | | | | | | | | | | | | | | | | | | | | | | |
|---|---|---|---|---|---|---|---|---|---|---|---|---|---|---|---|---|---|---|---|---|---|---|---|
| Upregulated | ENSMUSG00000051235 | 9,517278171 | 27,6578623 | 23,6014178 | 2,89070227 | 1,2722125 | 0 | 3,40767947 | 1,22222602 | 3,20327906 | 0 | 0 | 0 | 0 | 0 | 3,18598 | 0 | 33,91671 | 27,74384169 | 31,36869 | 55,99913 | 17,71456 | 12,5630747 |
| Upregulated | ENSMUSG00000028702 | 22,2069824 | 75,5272393 | 62,634532 | 29,8705901 | 13,9943375 | 17,25059984 | 34,0767947 | 39,1112327 | 17,084155 | 11,8656616 | 30,9151726 | 28,90304 | 16,0281034 | 38,35461 | 17,52289 | 22,78864477 | 131,5307 | 131,783248 | 115,9691 | 179,9972 | 46,94358 | 43,0733989 |
| Upregulated | ENSMUSG00000069793 | 169,1960564 | 163,819646 | 280,493774 | 65,5225848 | 24,1720375 | 26,53938437 | 124,948247 | 117,333698 | 54,455744 | 61,485701 | 13,2493597 | 43,79248 | 50,0878232 | 23,38696 | 26,28433 | 27,34637372 | 369,7749 | 228,8866939 | 270,9115 | 537,9916 | 147,9166 | 88,8388853 |
| Upregulated | ENSMUSG00000022034 | 28,55183451 | 24,4665705 | 52,6493167 | 17,3442136 | 3,8166375 | 1,326969219 | 27,2614358 | 14,6667123 | 3,20327906 | 11,8656616 | 0 | 7,882647 | 0 | 3,741914 | 0 | 0 | 47,97974 | 37,65235657 | 45,62719 | 95,9985 | 25,68611 | 9,87098725 |
| Upregulated | ENSMUSG00000050410 | 205,1502183 | 294,66261 | 312,264913 | 191,749917 | 106,86585 | 156,5823678 | 214,683807 | 188,222807 | 142,012038 | 176,906228 | 52,9974388 | 65,68873 | 51,0895797 | 34,6127 | 74,87053 | 107,5624033 | 402,0371 | 334,9078032 | 450,5685 | 561,9912 | 271,0328 | 195,62502 |
| Upregulated | ENSMUSG00000032815 | 57,10366903 | 53,1881967 | 60,8190383 | 39,5062644 | 29,2608875 | 26,53938437 | 56,7946579 | 34,2223286 | 36,3038293 | 58,2496115 | 34,2275126 | 14,0136 | 10,0175646 | 9,354784 | 11,15093 | 9,115457906 | 167,1018 | 93,14003995 | 107,414 | 191,997 | 74,40115 | 56,5338361 |
| Upregulated | ENSMUSG00000028312 | 195,6329402 | 261,685928 | 354,021268 | 234,146884 | 99,2325749 | 119,4272297 | 228,314525 | 176,000547 | 181,519147 | 223,290177 | 86,1208381 | 113,8605 | 74,1299784 | 146,8701 | 196,7343 | 153,1396928 | 532,7405 | 507,3159623 | 393,5345 | 871,3197 | 309,1191 | 279,977093 |
| Upregulated | ENSMUSG00000020649 | 407,1280106 | 482,948826 | 904,115853 | 267,871744 | 111,9547 | 110,1384451 | 258,98364 | 398,445683 | 255,194565 | 311,743291 | 160,09643 | 159,4046 | 99,17389 | 150,612 | 193,5483 | 133,9972312 | 1932,425 | 1526,902144 | 1435,355 | 2602,626 | 612,9238 | 624,564284 |
| Upregulated | ENSMUSG00000023908 | 95,17278171 | 56,3794885 | 135,254279 | 80,9396636 | 13,9943375 | 41,13604578 | 61,3382305 | 52,5557189 | 61,9300618 | 74,4300591 | 44,1645324 | 35,90984 | 43,075528 | 38,35461 | 27,87732 | 51,04656428 | 272,1609 | 238,7952088 | 206,2729 | 351,3279 | 120,459 | 106,786135 |
| Upregulated | ENSMUSG00000107060 | 9,517278171 | 11,7014033 | 29,9556457 | 5,78140454 | 3,8166375 | 2,653938437 | 3,40767947 | 13,4444862 | 5,33879843 | 12,9443581 | 4,41645324 | 5,255098 | 0 | 4,677392 | 1,59299 | 0,911545791 | 2,481711 | 8,917663399 | 28,51699 | 10,6665 | 23,02893 | 9,87098725 |
| Upregulated | ENSMUSG00000030322 | 41,24153874 | 25,5303344 | 44,4795952 | 27,9434553 | 41,9830125 | 22,55847672 | 11,3589316 | 20,7778424 | 35,2360696 | 50,6987359 | 9,93701978 | 3,503399 | 6,01053879 | 24,32244 | 9,557939 | 8,203912116 | 9,099605 | 37,65235657 | 74,14419 | 38,66606 | 27,45757 | 26,9208743 |
| Upregulated | ENSMUSG00000051022 | 215,7249719 | 64,8896 | 229,659951 | 51,0690735 | 13,9943375 | 26,53938437 | 24,9896495 | 18,3333903 | 35,2360696 | 15,1017511 | 46,372759 | 60,43363 | 101,177403 | 23,38696 | 14,33691 | 40,10801479 | 67,83342 | 84,22237655 | 129,277 | 361,9944 | 229,4036 | 109,478222 |
| Upregulated | ENSMUSG00000029521 | 106,8050106 | 48,933141 | 65,3577725 | 35,6519947 | 11,4499125 | 13,26969219 | 28,397329 | 22,0000684 | 12,8131162 | 26,9674127 | 13,2493597 | 27,15134 | 35,0614763 | 8,419306 | 23,89485 | 8,203912116 | 66,17895 | 63,41449528 | 48,47889 | 124,6647 | 94,7729 | 47,5602113 |
| Upregulated | ENSMUSG00000097993 | 67,67842255 | 5,31881967 | 61,7267851 | 26,0163204 | 0 | 0 | 0 | 7,33335613 | 18,1519147 | 6,47217906 | 0 | 14,88944 | 1,00175646 | 13,0967 | 0 | 0 | 4,136184 | 17,8353268 | 25,6653 | 77,99878 | 35,42912 | 8,97362477 |
| Upregulated | ENSMUSG00000021185 | 253,7940846 | 113,822741 | 196,981064 | 91,5389053 | 48,344075 | 55,73270718 | 106,773957 | 89,2224996 | 43,7781471 | 87,3744172 | 29,8110593 | 21,02039 | 72,1264654 | 30,87079 | 31,0633 | 14,58473265 | 75,27855 | 60,44194082 | 121,6725 | 465,3261 | 88,5728 | 40,3813115 |
| Upregulated | ENSMUSG00000031803 | 150,1615 | 105,312629 | 69,8965067 | 59,7411803 | 12,722125 | 66,34846093 | 48,8434058 | 18,3333903 | 16,0163953 | 31,2821988 | 5,52056654 | 45,54418 | 36,0632327 | 28,99983 | 10,35443 | 31,90410267 | 53,77039 | 35,6706536 | 152,0906 | 231,3297 | 124,8877 | 21,5366995 |
| Upregulated | ENSMUSG00000020185 | 201,9777923 | 172,329757 | 175,19514 | 84,7939333 | 54,7051374 | 173,8329676 | 95,4150253 | 48,8890409 | 61,9300618 | 23,7313232 | 29,8110593 | 44,66833 | 13,022834 | 15,90313 | 43,80722 | 27,34637372 | 156,3478 | 111,9662182 | 143,5355 | 335,9948 | 124,0019 | 116,657122 |
| Upregulated | ENSMUSG00000059323 | 137,4717958 | 58,5070164 | 76,2507346 | 32,7612924 | 22,899825 | 21,2315075 | 36,3485811 | 74,5557873 | 39,5071084 | 74,4300591 | 39,7480791 | 40,28909 | 11,0193211 | 27,12887 | 49,38269 | 60,16202218 | 116,6404 | 66,38704975 | 119,7714 | 189,3304 | 65,54387 | 36,7918616 |
| Upregulated | ENSMUSG00000033762 | 40,18406339 | 8,51011147 | 25,4169115 | 14,4535114 | 11,4499125 | 0 | 7,95125211 | 8,55558215 | 6,40655811 | 17,2591441 | 12,1452464 | 4,379248 | 7,01229525 | 12,16122 | 30,26681 | 16,40782423 | 33,08947 | 23,78043573 | 51,33059 | 53,3325 | 27,45757 | 10,7683497 |
| Upregulated | ENSMUSG00000034206 | 49,70134156 | 25,5303344 | 36,3098736 | 32,7612924 | 0 | 19,90453828 | 9,08714527 | 26,8889725 | 6,40655811 | 22,6526267 | 13,2493597 | 5,255098 | 23,0403987 | 8,419306 | 3,982475 | 17,31937002 | 31,435 | 20,80788127 | 79,84759 | 59,33241 | 16,82883 | 38,5865865 |
| Upregulated | ENSMUSG00000015568 | 21019,43758 | 6360,24456 | 17489,5584 | 7081,257 | 3153,81478 | 4207,819392 | 4816,18699 | 6393,46432 | 4986,43773 | 5407,5056 | 4209,98405 | 2741,41 | 8936,66942 | 2112,31 | 1230,585 | 1183,186436 | 7122,509 | 8788,852706 | 16485,67 | 15303,09 | 10399,33 | 12317,1974 |
| Upregulated | ENSMUSG00000049791 | 429,334993 | 157,437062 | 782,477776 | 186,93208 | 33,077525 | 62,36755328 | 43,16394 | 239,5563 | 155,892914 | 162,883173 | 79,4961582 | 104,2261 | 288,505862 | 211,4181 | 86,81795 | 67,45438851 | 311,8683 | 156,5545352 | 623,5716 | 849,9867 | 488,0361 | 555,467374 |
| Upregulated | ENSMUSG00000111740 | 3,172426057 | 3,1912918 | 1,81549368 | 0 | 3,8166375 | 5,307876874 | 1,13589316 | 0 | 0 | 2,15739302 | 6,62467985 | 11,38605 | 14,0245905 | 17,77409 | 15,9299 | 5,469274744 | 8,272368 | 9,908514888 | 17,1102 | 15,33309 | 5,314368 | 2,69208743 |
| Upregulated | ENSMUSG00000039364 | 142,7591726 | 117,014033 | 169,748659 | 105,028849 | 59,7939874 | 87,57996843 | 78,3766279 | 68,4446572 | 83,2852555 | 42,0691639 | 73,9755917 | 423,0354 | 633,110086 | 328,3529 | 442,0547 | 356,4144041 | 699,0151 | 310,136516 | 551,3286 | 633,3234 | 52,25795 | 166,909421 |
| Upregulated | ENSMUSG00000089992 | 39,12658804 | 34,0404459 | 34,4943799 | 27,9434553 | 31,8053125 | 14,5966614 | 12,4948247 | 30,5556505 | 23,4907131 | 31,2821988 | 22,0822662 | 42,91663 | 56,098362 | 43,03201 | 50,17918 | 29,1694653 | 65,35171 | 39,63405955 | 65,58909 | 62,66569 | 34,54339 | 28,7155993 |
| Upregulated | ENSMUSG00000052482 | 5,287376762 | 2,12752787 | 8,16972156 | 7,70853939 | 2,544425 | 0 | 3,40767947 | 0 | 2,13551937 | 0 | 2,20822662 | 7,006798 | 19,0333728 | 28,06435 | 26,28433 | 13,67318686 | 31,435 | 5,945108933 | 8,555098 | 16,66641 | 6,200096 | 14,3577996 |
| Upregulated | ENSMUSG00000038459 | 17,97708099 | 13,8289311 | 24,5091647 | 25,052753 | 29,2608875 | 29,19332281 | 29,5332221 | 12,2222602 | 40,5748681 | 30,2035023 | 46,372759 | 34,15814 | 48,0843103 | 131,9025 | 105,9338 | 57,42738481 | 108,368 | 48,55172295 | 57,03399 | 90,66525 | 52,25795 | 55,6364736 |
| Upregulated | ENSMUSG00000042377 | 29,60930986 | 26,5940983 | 11,8007089 | 26,0163204 | 34,3497375 | 30,52029203 | 71,561269 | 37,8890067 | 49,1169455 | 6,47217906 | 17,6658129 | 52,55098 | 74,1299784 | 103,8381 | 117,8813 | 32,81564846 | 86,85987 | 54,49683188 | 74,14419 | 151,9976 | 47,82931 | 65,5074609 |
| Upregulated | ENSMUSG00000098123 | 8,459802819 | 28,7216262 | 13,6162026 | 13,4899439 | 11,4499125 | 3,980907656 | 15,9025042 | 18,3333903 | 27,7617518 | 11,8656616 | 13,2493597 | 19,26869 | 25,0439116 | 40,22557 | 39,82475 | 30,99255688 | 33,08947 | 33,68895062 | 35,17096 | 48,66591 | 31,00048 | 23,3314244 |
| Upregulated | ENSMUSG00000070524 | 7,402327466 | 0 | 9,98521524 | 4,81783712 | 12,722125 | 0 | 0 | 2,44445204 | 4,27103874 | 12,9443581 | 28,706946 | 27,15134 | 30,0526939 | 54,25775 | 46,9932 | 42,84265216 | 47,1525 | 21,79873275 | 22,8136 | 62,66569 | 30,11475 | 25,1261494 |
| Upregulated | ENSMUSG00000042784 | 5,287376762 | 7,44634754 | 7,26197472 | 24,0891856 | 11,4499125 | 11,94272297 | 4,54357263 | 13,4444862 | 16,0163953 | 4,31478604 | 57,4138921 | 43,79248 | 34,0597198 | 48,64488 | 61,33011 | 37,37337742 | 75,27855 | 49,54257444 | 34,22039 | 81,99872 | 38,97203 | 14,3577996 |
| Upregulated | ENSMUSG00000035778 | 85,65550354 | 65,9533639 | 129,807798 | 71,3039894 | 19,0831875 | 18,57756906 | 28,397329 | 37,8890067 | 46,9814262 | 112,184437 | 125,868917 | 84,95742 | 135,237123 | 388,2235 | 200,7167 | 239,7365429 | 301,1142 | 231,8592484 | 163,4974 | 384,6607 | 127,5448 | 106,786135 |
| Upregulated | ENSMUSG00000016477 | 49,70134156 | 58,5070164 | 59,0035446 | 57,8140454 | 31,8053125 | 19,90453828 | 36,3485811 | 69,6668832 | 74,743178 | 79,8235417 | 117,036011 | 93,71592 | 109,191455 | 241,3534 | 97,96888 | 106,6508575 | 143,9392 | 125,8381391 | 142,585 | 214,6633 | 112,4875 | 109,478222 |
| Upregulated | ENSMUSG00000058254 | 244,2768064 | 151,054479 | 238,737419 | 200,422024 | 180,654175 | 143,3126756 | 170,383974 | 232,222944 | 164,434992 | 404,511191 | 241,800815 | 395,0082 | 368,646379 | 275,0306 | 448,4267 | 502,2617306 | 779,2571 | 404,2674074 | 583,6478 | 628,6569 | 200,1745 | 340,100379 |
| Upregulated | ENSMUSG00000049871 | 16,91960564 | 7,44634754 | 0 | 9,63567424 | 1,2722125 | 2,653938437 | 6,81535895 | 3,66667807 | 14,9486356 | 16,1804476 | 13,2493597 | 34,15814 | 13,022834 | 16,83861 | 21,50536 | 38,28492321 | 57,07934 | 14,86277233 | 39,92379 | 57,33244 | 19,48602 | 25,1261494 |
| Upregulated | ENSMUSG00000028364 | 11415,44643 | 16683,0098 | 18681,43 | 4332,19914 | 6246,56337 | 1910,835675 | 5109,24743 | 2680,34167 | 2155,80681 | 9119,30029 | 11237,6653 | 122658,4 | 13125,0132 | 54753,55 | 160315,3 | 128109,557 | 266650,7 | 57574,41661 | 223862,2 | 274129,1 | 10019,36 | 106522,31 |
| Upregulated | ENSMUSG00000023008 | 169,1960564 | 269,132275 | 341,312812 | 285,215958 | 159,026562 | 134,0238911 | 211,276127 | 166,222739 | 198,603302 | 353,812455 | 223,030888 | 503,6136 | 182,319677 | 363,9011 | 434,0897 | 700,978713 | 963,7309 | 329,9535458 | 746,1947 | 1160,649 | 229,4036 | 321,255767 |
| Upregulated | ENSMUSG00000062588 | 22,2069824 | 25,5303344 | 21,7859242 | 17,3442136 | 25,44425 | 9,28878453 | 22,7178632 | 24,4445204 | 22,4229534 | 18,3378407 | 26,4987194 | 35,03399 | 31,0544504 | 28,99983 | 57,34764 | 41,01956058 | 103,4046 | 79,2681191 | 78,89702 | 142,6644 | 35,42912 | 43,0733989 |
| Upregulated | ENSMUSG00000041324 | 372,231324 | 424,44181 | 628,160813 | 578,140454 | 254,4425 | 317,1456432 | 399,834392 | 250,556334 | 438,849231 | 659,083567 | 354,420372 | 1082,55 | 489,858911 | 970,0911 | 1482,277 | 2131,194059 | 5836,983 | 1674,539016 | 5238,572 | 7131,889 | 949,5004 | 2594,27492 |
| Upregulated | ENSMUSG00000015217 | 46,5289155 | 46,8056131 | 67,1732662 | 110,810254 | 30,5331 | 35,8281689 | 49,979299 | 78,2224654 | 73,6754183 | 43,1478604 | 80,6002715 | 72,69552 | 81,1422736 | 119,7412 | 126,6427 | 119,4124986 | 264,7158 | 172,4081591 | 238,5922 | 315,9951 | 121,3447 | 131,014922 |
| Upregulated | ENSMUSG00000028614 | 799,4513664 | 723,359475 | 776,123548 | 771,817507 | 576,312262 | 400,744704 | 599,751588 | 622,113045 | 541,354161 | 784,212362 | 548,744314 | 744,4722 | 590,034558 | 588,4159 | 1319,792 | 957,1230802 | 2111,108 | 1304,951411 | 1620,716 | 1925,303 | 698,8394 | 1259,89692 |
| Upregulated | ENSMUSG00000040697 | 114,2073381 | 105,312629 | 92,5901777 | 87,6846356 | 78,8771749 | 27,86635359 | 81,7843074 | 79,4446914 | 51,2524649 | 132,679671 | 87,2249514 | 91,96422 | 77,1352478 | 128,1605 | 170,4499 | 154,9627844 | 253,1345 | 114,9387727 | 188,2122 | 251,3294 | 128,4306 | 154,346346 |
| Upregulated | ENSMUSG00000004864 | 14,80465493 | 23,4028065 | 24,5091647 | 22,1620508 | 13,9943375 | 10,61575375 | 24,9896495 | 17,1111643 | 3,20327906 | 30,2035023 | 17,6658129 | 15,76529 | 7,01229525 | 13,0967 | 48,58619 | 45,57728953 | 53,77039 | 21,79873275 | 78,89702 | 67,33228 | 4,42864 | 64,6100984 |
| Upregulated | ENSMUSG00000020899 | 103,6325845 | 88,2924065 | 133,438785 | 93,4660401 | 54,7051374 | 37,15513812 | 43,16394 | 108,778116 | 69,4043796 | 58,2496115 | 73,9755917 | 150,6461 | 181,31792 | 114,1284 | 246,1169 | 184,1322497 | 197,7096 | 152,5911293 | 355,5119 | 345,3279 | 69,97251 | 146,270084 |
| Upregulated | ENSMUSG00000039985 | 23,26445775 | 34,0404459 | 55,3725572 | 25,052753 | 12,722125 | 1,326969219 | 15,9025042 | 13,4444862 | 24,5584728 | 45,3052534 | 15,4575863 | 67,44043 | 30,0526939 | 33,67722 | 98,76537 | 39,196469 | 67,83342 | 59,45108933 | 98,85892 | 73,33219 | 23,91466 | 49,3549363 |
| Upregulated | ENSMUSG00000021259 | 24,3219331 | 5,31881967 | 23,6014178 | 4,81783712 | 0 | 14,5966614 | 11,3589316 | 1,22222602 | 5,33879843 | 28,0461092 | 23,1863795 | 37,66154 | 19,0333728 | 41,16105 | 28,67382 | 12,76164107 | 33,91671 | 21,79873275 | 58,93512 | 51,99919 | 15,05738 | 24,2287869 |
| Upregulated | ENSMUSG00000056313 | 378,5761761 | 213,816551 | 418,471293 | 90,5753379 | 137,39895 | 138,0047987 | 227,178632 | 95,3336297 | 93,9628523 | 504,829966 | 605,054093 | 356,4708 | 167,29333 | 272,2242 | 628,4345 | 375,5568657 | 635,3179 | 369,5876053 | 753,7992 | 1240,647 | 328,6051 | 553,672649 |
| Upregulated | ENSMUSG00002075252 | 29,60930986 | 11,7014033 | 26,3246584 | 11,5628091 | 16,5387625 | 19,90453828 | 20,4460768 | 3,66667807 | 2,13551937 | 30,2035023 | 49,6850989 | 82,32987 | 51,0895797 | 31,80627 | 61,33011 | 20,05400739 | 66,17895 | 67,37790124 | 76,04532 | 132,6646 | 20,37174 | 31,4076867 |
| Upregulated | ENSMUSG00000022122 | 25,37940846 | 12,7651672 | 140,70076 | 21,1984833 | 0 | 0 | 1,13589316 | 12,2222602 | 5,33879843 | 0 | 12,1452464 | 252,2447 | 11,0193211 | 47,7094 | 157,706 | 98,44694539 | 139,803 | 73,32301017 | 169,2008 | 181,9972 | 77,05834 | 210,880182 |
| Upregulated | ENSMUSG00000029819 | 3,172426057 | 10,6376393 | 9,98521524 | 2,89070227 | 5,08884999 | 0 | 21,58197 | 7,33335613 | 9,60983717 | 12,9443581 | 19,8740396 | 117,3639 | 20,0351293 | 33,67722 | 51,77217 | 50,13501849 | 80,24197 | 56,47853486 | 115,9691 | 78,66544 | 7,971552 | 85,2494354 |
| Upregulated | ENSMUSG00000020407 | 58,16114438 | 45,7418492 | 48,1105825 | 13,4899439 | 5,08884999 | 37,15513812 | 13,6307179 | 29,3334245 | 32,0327906 | 21,5739302 | 19,8740396 | 135,7567 | 48,0843103 | 47,7094 | 90,80042 | 68,3659343 | 107,5408 | 84,22237655 | 159,6952 | 148,6643 | 74,40115 | 104,094047 |
| Upregulated | ENSMUSG00000058013 | 3051,873867 | 2160,50455 | 3203,4386 | 2014,81948 | 1250,58489 | 1084,133852 | 1732,23707 | 1476,44903 | 1632,60456 | 1683,84525 | 1521,46814 | 4104,232 | 2597,55451 | 2138,504 | 3101,551 | 3632,509976 | 5043,663 | 3065,694506 | 4036,105 | 4558,596 | 2623,526 | 2706,44523 |
| Upregulated | ENSMUSG00000073821 | 51,81629226 | 8,51011147 | 42,6641015 | 2,89070227 | 8,90548749 | 6,634846093 | 6,81535895 | 11,0000342 | 1,06775969 | 24,8100197 | 48,5809856 | 94,59177 | 21,0368858 | 56,1287 | 52,56867 | 52,86965586 | 123,2583 | 77,28641613 | 57,03399 | 95,33185 | 11,51446 | 44,8681239 |
| Upregulated | ENSMUSG00000024966 | 2570,722581 | 3028,53592 | 1882,66695 | 1824,03313 | 1318,01215 | 792,2006235 | 1141,57262 | 1558,33818 | 1746,85485 | 1167,14962 | 1485,0324 | 3058,467 | 2894,07443 | 2941,144 | 3013,937 | 3001,720289 | 3629,088 | 2740,695218 | 3048,467 | 3089,285 | 2486,239 | 2395,06045 |
| Upregulated | ENSMUSG00000042246 | 50,75881691 | 22,3390426 | 49,0183294 | 13,4899439 | 13,9943375 | 3,980907656 | 19,3101837 | 26,8889725 | 10,6775969 | 12,9443581 | 46,372759 | 62,18533 | 65,1141702 | 54,25775 | 69,29506 | 131,2625939 | 117,4676 | 64,40534677 | 80,79815 | 74,6655 | 52,25795 | 48,4575738 |
| Upregulated | ENSMUSG00000040663 | 143,8166479 | 56,3794885 | 232,383191 | 87,6846356 | 232,814887 | 99,5226914 | 240,80935 | 116,111472 | 230,636092 | 127,286188 | 223,030888 | 509,7445 | 307,539235 | 695,0604 | 621,2661 | 451,2151664 | 962,0764 | 435,9746551 | 663,4954 | 759,3215 | 231,175 | 594,05396 |
| Upregulated | ENSMUSG00000062480 | 109,9774366 | 105,312629 | 120,73033 | 172,478569 | 101,777 | 79,61815312 | 89,7355595 | 127,111506 | 149,486356 | 98,1613823 | 93,8496312 | 749,7273 | 473,830808 | 827,8984 | 561,5289 | 525,0503754 | 1067,963 | 502,3617048 | 596,9558 | 605,3239 | 426,9209 | 428,041902 |
| Upregulated | ENSMUSG00000022246 | 802,6237924 | 698,892905 | 701,688307 | 495,273656 | 585,217749 | 424,63015 | 594,072122 | 444,890272 | 564,844874 | 626,722672 | 707,736631 | 1930,373 | 1782,12475 | 2781,177 | 2133,81 | 2858,607599 | 4411,654 | 1938,105512 | 2373,565 | 2028,635 | 1675,797 | 1825,23528 |
| Upregulated | ENSMUSG00000086515 | 2,114950705 | 2,12752787 | 1,81549368 | 0 | 0 | 3,980907656 | 3,40767947 | 1,22222602 | 2,13551937 | 0 | 0 | 8,758497 | 0 | 14,96765 | 16,72639 | 16,40782423 | 28,12605 | 13,87192084 | 23,76416 | 14,66644 | 6,200096 | 11,6657122 |
| Upregulated | ENSMUSG00000008398 | 233,7020529 | 299,981429 | 253,261368 | 400,844048 | 128,493462 | 203,0262904 | 188,558264 | 160,111609 | 224,229534 | 237,313232 | 305,839387 | 445,8075 | 159,279278 | 687,5766 | 659,4978 | 638,9935992 | 876,8711 | 443,901467 | 726,2328 | 643,3233 | 422,4923 | 418,170914 |
| Upregulated | ENSMUSG00000024664 | 548,8297079 | 308,491541 | 563,710788 | 463,475931 | 349,858437 | 445,8616575 | 492,977631 | 391,112327 | 326,734464 | 511,302145 | 564,201901 | 1059,778 | 564,990646 | 1208,638 | 1481,481 | 1426,569162 | 1698,317 | 903,6565578 | 1280,413 | 1374,645 | 929,1287 | 757,373931 |
| Upregulated | ENSMUSG00000112546 | 47,58639085 | 80,846059 | 66,2655193 | 80,9396636 | 86,5104499 | 50,42483031 | 62,4741237 | 64,7779792 | 42,7103874 | 81,9809347 | 38,6439658 | 136,6326 | 110,193211 | 145,9346 | 136,2006 | 208,7439861 | 313,5228 | 97,1034459 | 205,3224 | 245,9962 | 155,0024 | 89,7362477 |
| Upregulated | ENSMUSG00000116398 | 30,66678522 | 31,912918 | 32,6788862 | 41,4333992 | 40,7108 | 21,2315075 | 27,2614358 | 33,0001026 | 30,9650309 | 31,2821988 | 20,9781529 | 66,56458 | 35,0614763 | 59,87062 | 65,31259 | 72,01211746 | 123,2583 | 46,57001997 | 83,64985 | 89,33194 | 57,57232 | 27,8182368 |
| Upregulated | ENSMUSG00000021048 | 260,1389367 | 326,575528 | 470,212863 | 497,200791 | 286,247812 | 175,1599369 | 269,206679 | 387,445649 | 322,463425 | 412,062067 | 193,219829 | 818,9195 | 594,041583 | 824,1565 | 994,0257 | 680,9247056 | 1278,081 | 752,05628 | 942,0114 | 1058,65 | 600,5236 | 476,499476 |
| Upregulated | ENSMUSG00000050370 | 13,74717958 | 8,51011147 | 26,3246584 | 40,4698318 | 6,36106249 | 17,25059984 | 11,3589316 | 28,1111985 | 17,084155 | 21,5739302 | 3,31233993 | 154,1495 | 111,194968 | 154,3539 | 130,6252 | 119,4124986 | 287,8784 | 57,46938635 | 187,2616 | 293,3288 | 65,54387 | 16,1525246 |
| Upregulated | ENSMUSG00000029231 | 9463,346928 | 8064,39438 | 10487,1992 | 8038,07945 | 5906,88263 | 5774,970039 | 6524,5703 | 7447,02315 | 6836,86527 | 8618,78511 | 7120,42673 | 17347,08 | 11538,231 | 18082,8 | 17270,4 | 16076,02156 | 25634,42 | 11693,03842 | 21332,61 | 21477 | 7532,231 | 9500,37655 |
| Upregulated | ENSMUSG00000001131 | 2454,400293 | 1904,13744 | 3695,43739 | 2648,84685 | 1365,08401 | 1653,403646 | 2085,49984 | 1950,67273 | 2706,7708 | 1516,64729 | 3246,09313 | 9940,018 | 13546,7527 | 19009,86 | 12201,51 | 14727,84534 | 22054,13 | 11778,25165 | 13730,93 | 28140,23 | 19079,47 | 9827,91385 |
| Upregulated | ENSMUSG00000026749 | 364,8289965 | 272,323567 | 548,279091 | 471,18447 | 264,6202 | 233,5465825 | 332,816695 | 398,445683 | 416,426277 | 261,044555 | 749,692937 | 1508,213 | 2118,71492 | 2228,31 | 1686,18 | 1947,973355 | 3002,043 | 1650,75858 | 2142,577 | 4093,269 | 2280,75 | 1665,50476 |
| Upregulated | ENSMUSG00000029552 | 65,56347184 | 82,9735868 | 92,5901777 | 67,4497197 | 41,9830125 | 26,53938437 | 26,1255426 | 37,8890067 | 18,1519147 | 18,3378407 | 79,4961582 | 211,0798 | 148,259957 | 323,6755 | 378,3351 | 216,0363524 | 377,22 | 180,334971 | 368,8198 | 488,659 | 284,3187 | 246,774681 |
| Upregulated | ENSMUSG00000040675 | 124,7820916 | 163,819646 | 130,715545 | 184,041378 | 138,671162 | 80,94512234 | 171,519867 | 97,7780817 | 69,4043796 | 141,309243 | 143,53473 | 487,8483 | 212,37237 | 280,6435 | 404,6194 | 444,8343458 | 530,2588 | 320,0450309 | 362,1658 | 579,9909 | 435,7782 | 235,108969 |
| Upregulated | ENSMUSG00000104913 | 2365,572363 | 2778,5514 | 2358,32629 | 2836,7425 | 2251,81612 | 1734,348769 | 2525,09049 | 2129,11773 | 1568,53898 | 2167,10129 | 1667,2111 | 5018,619 | 3221,64879 | 4492,167 | 5170,845 | 5878,558804 | 8083,758 | 4289,396095 | 6942,938 | 8679,198 | 5306,397 | 3700,72286 |
| Upregulated | ENSMUSG00000032294 | 9733,003143 | 11217,3907 | 9947,99762 | 11805,6281 | 9488,16082 | 7099,28532 | 10838,6925 | 8010,46935 | 7129,43142 | 9185,10078 | 7598,50779 | 21246,36 | 13303,3258 | 18726,41 | 22441,25 | 23136,85526 | 32270,51 | 17605,44925 | 26987,53 | 34927,45 | 20038,71 | 15020,0531 |
| Upregulated | ENSMUSG00000028613 | 1,057475352 | 6,3825836 | 1,81549368 | 4,81783712 | 16,5387625 | 21,2315075 | 14,7666111 | 4,88890409 | 4,27103874 | 4,31478604 | 35,3316259 | 97,21932 | 26,0456681 | 63,61253 | 125,0497 | 54,69274744 | 148,9026 | 82,24067357 | 61,78682 | 107,9983 | 13,28592 | 24,2287869 |
| Upregulated | ENSMUSG00000029228 | 33,83921127 | 44,6780852 | 72,6197472 | 55,8869106 | 66,1550499 | 70,32936859 | 36,3485811 | 26,8889725 | 39,5071084 | 47,4626464 | 102,682538 | 219,8383 | 73,1282219 | 302,1595 | 192,7518 | 184,1322497 | 403,6916 | 226,9049909 | 145,4367 | 249,3294 | 108,0588 | 94,2230601 |
| Upregulated | ENSMUSG00000027560 | 68,7358979 | 87,2286426 | 53,5570636 | 79,9760962 | 29,2608875 | 38,48210734 | 21,58197 | 24,4445204 | 26,6939921 | 34,5182883 | 13,2493597 | 196,1903 | 78,1370042 | 144,0637 | 245,3204 | 266,1713709 | 489,7242 | 145,6551689 | 150,1895 | 203,3302 | 61,11523 | 96,9151476 |
| Upregulated | ENSMUSG00000020534 | 64,50599649 | 98,9300459 | 123,45357 | 70,340422 | 86,5104499 | 90,23390687 | 84,0560937 | 146,667123 | 104,640449 | 117,57792 | 90,5372913 | 205,8247 | 143,251174 | 121,6122 | 211,0712 | 251,5866382 | 426,8542 | 248,7037237 | 163,4974 | 332,6615 | 132,8592 | 148,962171 |
| Upregulated | ENSMUSG00000094873 | 7,402327466 | 15,956459 | 9,98521524 | 9,63567424 | 19,0831875 | 7,961815312 | 7,95125211 | 4,88890409 | 6,40655811 | 4,31478604 | 3,31233993 | 18,39284 | 8,01405172 | 14,03218 | 29,47031 | 23,70019056 | 31,435 | 11,89021787 | 19,01133 | 36,66609 | 22,1432 | 7,17889982 |
| Upregulated | ENSMUSG00000041817 | 1,057475352 | 15,956459 | 10,8929621 | 1,92713485 | 5,08884999 | 0 | 6,81535895 | 2,44445204 | 2,13551937 | 0 | 0 | 10,5102 | 7,01229525 | 10,29026 | 22,30186 | 8,203912116 | 28,95329 | 8,917663399 | 0 | 19,33303 | 11,51446 | 5,38417486 |
| Upregulated | ENSMUSG00000028347 | 0 | 7,44634754 | 1,81549368 | 0 | 0 | 17,25059984 | 5,67946579 | 0 | 4,27103874 | 2,15739302 | 4,41645324 | 8,758497 | 0 | 23,38696 | 4,77897 | 15,49627844 | 49,63421 | 10,89936638 | 20,91246 | 35,33278 | 10,62874 | 12,5630747 |
| Upregulated | ENSMUSG00000115004 | 29,60930986 | 17,0202229 | 9,98521524 | 32,7612924 | 17,810975 | 11,94272297 | 14,7666111 | 18,3333903 | 8,54207749 | 11,8656616 | 17,6658129 | 30,65474 | 27,0474245 | 38,35461 | 20,70887 | 39,196469 | 62,04276 | 24,77128722 | 45,62719 | 39,99938 | 30,11475 | 26,9208743 |
| Upregulated | ENSMUSG00000056220 | 376,4612254 | 504,224105 | 364,91423 | 269,798879 | 320,59755 | 228,2387056 | 204,460768 | 455,890306 | 553,099517 | 380,779868 | 421,771284 | 832,9331 | 539,946734 | 930,801 | 983,6713 | 1241,525367 | 2881,266 | 788,7177851 | 1734,784 | 579,9909 | 956,5863 | 1230,28396 |
| Upregulated | ENSMUSG00000047562 | 488,5536128 | 1130,78106 | 605,467142 | 643,663039 | 1596,62669 | 1049,632652 | 842,832723 | 530,446093 | 672,688602 | 825,20283 | 1299,54136 | 3253,782 | 1839,22487 | 3185,304 | 1714,057 | 3053,678399 | 6868,548 | 2317,601632 | 3858,349 | 2527,294 | 1990,231 | 2970,2698 |
| Upregulated | ENSMUSG00000023088 | 957,0151938 | 790,376603 | 727,105219 | 664,861523 | 922,354062 | 744,4297316 | 999,585979 | 1097,55897 | 947,102841 | 916,892033 | 1127,29969 | 2071,385 | 1662,91573 | 1444,379 | 1747,51 | 2009,046923 | 3667,968 | 1677,511571 | 2362,158 | 1793,305 | 1100,96 | 2150,0805 |
| Upregulated | ENSMUSG00000029762 | 399,7256832 | 547,838426 | 499,260762 | 600,302505 | 531,784824 | 475,0549803 | 499,79299 | 416,779073 | 430,307153 | 565,236971 | 468,144043 | 740,093 | 828,452596 | 853,1563 | 948,6255 | 1038,250656 | 4220,562 | 1115,698776 | 2545,617 | 2727,291 | 1456,137 | 1589,22895 |
| Upregulated | ENSMUSG00000056758 | 39,12658804 | 23,4028065 | 44,4795952 | 3,8542697 | 5,08884999 | 2,653938437 | 1,13589316 | 6,11113011 | 8,54207749 | 8,62957207 | 1,10411331 | 58,68193 | 15,026347 | 51,45131 | 60,53362 | 50,13501849 | 201,8458 | 63,41449528 | 134,9804 | 112,6649 | 13,28592 | 78,0705355 |
| Upregulated | ENSMUSG00000017724 | 12,68970423 | 24,4665705 | 28,140152 | 41,4333992 | 10,1777 | 17,25059984 | 7,95125211 | 20,7778424 | 4,27103874 | 28,0461092 | 20,9781529 | 64,81288 | 20,0351293 | 25,25792 | 38,23176 | 50,13501849 | 178,6832 | 42,60661402 | 111,2163 | 69,99891 | 20,37174 | 41,278674 |
| Upregulated | ENSMUSG00000107479 | 0 | 1,06376393 | 0 | 0 | 0 | 2,653938437 | 0 | 6,11113011 | 1,06775969 | 0 | 0 | 0 | 27,0474245 | 19,64505 | 12,74392 | 9,115457906 | 43,84355 | 13,87192084 | 20,91246 | 15,99975 | 20,37174 | 11,6657122 |

| | | | | | | | | | | | | | | | | | | | | | | | |
|---|---|---|---|---|---|---|---|---|---|---|---|---|---|---|---|---|---|---|---|---|---|---|---|
| Upregulated | ENSMUSG00000117359 | 12,68970423 | 8,51011147 | 15,4316963 | 20,2349159 | 8,90548749 | 11,94272297 | 19,3101837 | 7,33335613 | 20,287434 | 15,1017511 | 23,1863795 | 18,39284 | 29,0509375 | 36,48366 | 24,69134 | 29,1694653 | 52,11592 | 27,74384169 | 40,87436 | 38,66606 | 38,0863 | 17,9472495 |
| Upregulated | ENSMUSG00000049985 | 90,9428803 | 30,8491541 | 68,081013 | 35,6519947 | 61,0661999 | 49,09786109 | 15,9025042 | 78,2224654 | 45,9136665 | 23,7313232 | 54,1015521 | 112,9846 | 198,34778 | 236,676 | 215,8501 | 257,055913 | 389,6286 | 248,7037237 | 175,8548 | 180,6638 | 207,2604 | 145,372721 |
| Upregulated | ENSMUSG00000027408 | 14503,27446 | 11900,3271 | 16890,4455 | 16412,4439 | 16013,3387 | 15654,25587 | 16846,4314 | 22719,9595 | 14511,9219 | 20511,4141 | 20274,8327 | 56319,76 | 60936,8457 | 63852,01 | 46857 | 73352,08977 | 135302 | 74344,57806 | 92025,29 | 76634,14 | 82668,54 | 40432,4611 |
| Upregulated | ENSMUSG00000030088 | 69,79337325 | 53,1881967 | 71,7120004 | 79,0125288 | 7,63327499 | 23,88544594 | 10,2230384 | 62,3335271 | 25,6262325 | 17,2591441 | 49,6850989 | 569,3023 | 434,762306 | 676,3509 | 400,637 | 540,5466538 | 970,3488 | 595,5017448 | 702,4686 | 735,9885 | 677,5819 | 424,452452 |
| Upregulated | ENSMUSG00000023832 | 314,0701796 | 377,636197 | 452,057926 | 558,869106 | 363,852775 | 334,3962431 | 357,806345 | 554,890614 | 555,235037 | 350,576365 | 399,689018 | 1398,732 | 1212,12532 | 1498,636 | 1378,733 | 1315,360576 | 2574,361 | 1557,61854 | 1315,584 | 1425,311 | 1437,537 | 1041,83784 |
| Upregulated | ENSMUSG00000037936 | 343,6794895 | 361,679738 | 395,777622 | 534,77992 | 199,737362 | 177,8138753 | 202,188982 | 326,334348 | 292,566154 | 209,267123 | 229,655568 | 1013,358 | 899,577305 | 902,7366 | 896,0568 | 1164,95552 | 1949,797 | 999,7691522 | 1278,512 | 908,6525 | 922,9286 | 845,315454 |
| Upregulated | ENSMUSG00000030220 | 233,7020529 | 277,642387 | 377,622685 | 363,264919 | 64,8828374 | 98,19572218 | 104,502171 | 249,334108 | 170,84155 | 282,618485 | 186,595149 | 597,3295 | 1016,78281 | 923,3172 | 335,3244 | 661,782244 | 1055,554 | 1112,726222 | 722,4305 | 1019,317 | 797,1552 | 872,236328 |
| Upregulated | ENSMUSG00000021190 | 2772,700374 | 2204,11887 | 4403,47992 | 3047,76376 | 1746,74776 | 1301,756803 | 2016,21036 | 897,1139 | 1454,28869 | 1870,45975 | 1319,4154 | 7379,034 | 10672,7134 | 9103,14 | 7068,893 | 7620,52281 | 11176,8 | 8083,366446 | 9569,353 | 7316,552 | 7811,235 | 4237,34562 |
| Upregulated | ENSMUSG00000037601 | 760,3247783 | 844,628564 | 989,444056 | 1563,86993 | 784,955112 | 766,9882084 | 984,819368 | 994,891982 | 1203,36517 | 764,795825 | 863,416607 | 1969,786 | 2714,76002 | 2736,274 | 2512,145 | 2503,104741 | 2780,343 | 2550,451732 | 2235,732 | 2424,629 | 2675,784 | 1769,59881 |
| Upregulated | ENSMUSG00000021252 | 383,8635529 | 448,90838 | 407,578331 | 608,974612 | 246,809225 | 233,5465825 | 316,914191 | 299,445375 | 282,956317 | 332,238525 | 331,233993 | 429,1663 | 636,115355 | 994,4135 | 699,3226 | 564,2468444 | 1512,189 | 1327,740995 | 953,4182 | 848,6534 | 923,8143 | 443,297064 |
| Upregulated | ENSMUSG00000041939 | 180,8282852 | 155,309534 | 161,578938 | 344,957138 | 138,671162 | 176,4869061 | 101,094491 | 213,889554 | 159,096193 | 141,309243 | 140,22239 | 159,4046 | 196,344267 | 383,5461 | 266,0293 | 206,0093487 | 548,458 | 544,9683188 | 389,7323 | 359,9944 | 482,7218 | 264,721931 |
| Upregulated | ENSMUSG00000027952 | 70,85084861 | 134,034256 | 95,3134182 | 204,276294 | 134,854525 | 118,1002605 | 97,6868116 | 99,0003078 | 123,860124 | 62,5643975 | 69,5591385 | 162,0322 | 233,409256 | 351,7399 | 234,1695 | 140,3780518 | 434,2993 | 320,0450309 | 179,6571 | 223,9965 | 403,892 | 157,935796 |
| Upregulated | ENSMUSG00000031349 | 388,0934543 | 389,3376 | 353,113521 | 366,155621 | 198,46515 | 205,6802289 | 153,345576 | 295,778697 | 224,229534 | 331,159828 | 181,074583 | 381,8705 | 415,728933 | 650,1575 | 475,5075 | 347,2989462 | 1032,392 | 647,0260222 | 455,3214 | 609,3238 | 906,9855 | 401,121027 |
| Upregulated | ENSMUSG00000006517 | 171,3110071 | 176,584813 | 108,021874 | 298,705901 | 136,126737 | 50,42483031 | 127,220034 | 128,333732 | 70,4721393 | 75,5087556 | 83,9126115 | 183,9284 | 246,43209 | 397,5783 | 257,2679 | 144,0242349 | 578,2386 | 323,0175854 | 319,3903 | 352,6612 | 523,4653 | 224,340619 |
| Upregulated | ENSMUSG00000113859 | 10,57475352 | 5,31881967 | 9,0774684 | 13,4899439 | 1,2722125 | 3,980907656 | 5,67946579 | 0 | 1,06775969 | 2,15739302 | 1,10411331 | 6,130948 | 15,026347 | 14,96765 | 7,964949 | 5,469274744 | 49,63421 | 19,81702978 | 12,35736 | 26,66625 | 26,57184 | 6,28153734 |
| Upregulated | ENSMUSG00000084331 | 8,459802819 | 15,956459 | 15,4316963 | 11,5628091 | 6,36106249 | 1,326969219 | 6,81535895 | 9,77780817 | 8,54207749 | 11,8656616 | 3,31233993 | 14,88944 | 8,01405172 | 26,19339 | 19,11588 | 26,43482793 | 65,35171 | 30,71639615 | 34,22039 | 33,33281 | 40,74349 | 10,7683497 |
| Upregulated | ENSMUSG00000033105 | 138,5292712 | 135,09802 | 178,826127 | 383,499835 | 52,1607124 | 86,25299921 | 52,2510853 | 124,667054 | 112,114767 | 235,155839 | 85,0167248 | 425,663 | 368,646379 | 566,8999 | 684,9857 | 350,0335836 | 1417,057 | 672,7881609 | 538,0206 | 705,989 | 690,8679 | 327,537304 |
| Upregulated | ENSMUSG00000059743 | 15,86213028 | 20,2115147 | 22,693671 | 47,2148038 | 8,90548749 | 5,307876874 | 12,4948247 | 22,0000684 | 22,4229534 | 10,7869651 | 8,83290647 | 53,42683 | 43,075528 | 87,93497 | 82,03898 | 49,22347269 | 176,2014 | 85,21322804 | 86,50155 | 80,66541 | 147,0309 | 29,6129618 |
| Upregulated | ENSMUSG00000090610 | 240,046905 | 255,303344 | 385,792407 | 701,477085 | 124,676825 | 114,1193528 | 147,666111 | 259,111917 | 211,416418 | 194,165372 | 170,03345 | 771,6236 | 785,377068 | 1456,54 | 902,4288 | 704,6248962 | 2288,964 | 1342,603767 | 1165,395 | 926,6522 | 1672,254 | 590,46451 |
| Upregulated | ENSMUSG00000093651 | 29,60930986 | 44,6780852 | 45,387342 | 105,992417 | 10,1777 | 23,88544594 | 10,2230384 | 37,8890067 | 23,4907131 | 6,47217906 | 27,6028327 | 119,9914 | 79,1387607 | 161,8378 | 111,5093 | 85,68530432 | 339,9943 | 183,3075254 | 154,9423 | 123,9981 | 191,3173 | 81,6599854 |
| Upregulated | ENSMUSG00000073394 | 3,172426057 | 2,12752787 | 14,5239494 | 5,78140454 | 3,8166375 | 2,653938437 | 2,27178632 | 7,33335613 | 9,60983717 | 1,07869651 | 4,41645324 | 14,88944 | 12,0210776 | 9,354784 | 3,982475 | 13,67318686 | 26,47158 | 16,84447531 | 13,30793 | 19,33303 | 19,48602 | 7,17889982 |
| Upregulated | ENSMUSG00000024665 | 99,40268312 | 130,842964 | 145,239494 | 340,139301 | 72,5161124 | 106,1575375 | 92,0073458 | 194,333937 | 163,367232 | 196,322765 | 112,619557 | 553,537 | 509,89404 | 534,1582 | 650,7364 | 414,7533347 | 1623,866 | 793,6720425 | 789,9208 | 1381,978 | 1821,943 | 417,273552 |
| Upregulated | ENSMUSG00000068196 | 284,4608698 | 651,023528 | 985,813068 | 1642,88246 | 162,8432 | 108,8114759 | 185,150585 | 249,334108 | 114,250286 | 338,710704 | 279,340667 | 742,7205 | 1434,51526 | 1272,251 | 1577,856 | 1927,007801 | 7482,357 | 4623,313047 | 3916,334 | 6773,228 | 5954,749 | 1276,04944 |
| Upregulated | ENSMUSG00000028362 | 11,63222888 | 0 | 3,63098736 | 5,78140454 | 3,8166375 | 0 | 0 | 0 | 1,06775969 | 0 | 5,52056654 | 25,39964 | 159,279278 | 129,096 | 40,62124 | 102,0931286 | 314,35 | 210,0605156 | 173,9537 | 280,6623 | 270,147 | 117,554485 |
| Upregulated | ENSMUSG00000034926 | 7,402327466 | 15,956459 | 19,9704305 | 14,4535114 | 0 | 1,326969219 | 6,81535895 | 20,7778424 | 14,9486356 | 3,23608953 | 9,93701978 | 59,55778 | 84,147543 | 83,25758 | 35,84227 | 59,25047639 | 152,2116 | 92,14918846 | 68,44079 | 147,9977 | 114,2589 | 97,81251 |
| Upregulated | ENSMUSG00000028044 | 505,4732184 | 348,91457 | 554,633319 | 593,557533 | 237,903737 | 338,3771507 | 316,914191 | 465,668114 | 537,083122 | 283,697182 | 217,510322 | 441,4282 | 687,204935 | 823,221 | 553,564 | 519,5811007 | 1101,052 | 1012,650222 | 915,3955 | 1623,308 | 1201,933 | 727,760969 |
| Upregulated | ENSMUSG00000024352 | 0 | 1,06376393 | 2,72324052 | 5,78140454 | 0 | 2,653938437 | 1,13589316 | 3,66667807 | 2,13551937 | 0 | 0 | 7,882647 | 10,0175646 | 13,0967 | 5,575465 | 5,469274744 | 17,37197 | 10,89936638 | 14,2585 | 26,66625 | 15,9431 | 8,97362477 |
| Upregulated | ENSMUSG00000047298 | 0 | 0 | 0 | 0 | 0 | 5,307876874 | 0 | 0 | 0 | 5,39348255 | 1,10411331 | 14,88944 | 8,01405172 | 0 | 10,35443 | 0 | 28,95329 | 5,945108933 | 4,752832 | 11,33316 | 15,05738 | 12,5630747 |
| Upregulated | ENSMUSG00000032315 | 0 | 0 | 7,26197472 | 0,96356742 | 1,2722125 | 6,634846093 | 0 | 17,1111643 | 19,2196743 | 1,07869651 | 7,72879316 | 10,5102 | 36,0632327 | 0 | 11,94742 | 2,734637372 | 165,4474 | 24,77128722 | 65,58909 | 139,9978 | 57,57232 | 124,733384 |
| Upregulated | ENSMUSG00000052336 | 6,344852114 | 0 | 0 | 2,89070227 | 0 | 0 | 0 | 0 | 2,13551937 | 7,55087556 | 4,41645324 | 5,255098 | 14,0245905 | 1,870957 | 4,77897 | 0 | 42,18908 | 26,7529902 | 11,4068 | 29,99953 | 11,51446 | 31,4076867 |
| Upregulated | ENSMUSG00000032415 | 21,14950705 | 21,2752787 | 24,5091647 | 19,2713485 | 3,8166375 | 25,21241515 | 6,81535895 | 20,7778424 | 32,0327906 | 21,5739302 | 28,706946 | 22,77209 | 27,0474245 | 27,12887 | 53,36516 | 18,23091581 | 52,11592 | 42,60661402 | 38,97323 | 65,33231 | 60,22951 | 44,8681239 |
| Upregulated | ENSMUSG00000085196 | 10,57475352 | 4,25505574 | 3,63098736 | 2,89070227 | 10,1777 | 10,61575375 | 6,81535895 | 12,2222602 | 3,20327906 | 7,55087556 | 9,93701978 | 14,0136 | 17,0298599 | 0,935478 | 23,89485 | 6,380820534 | 22,33539 | 9,908514888 | 7,604532 | 38,66606 | 25,68611 | 23,3314244 |
| Upregulated | ENSMUSG00000019891 | 33,83921127 | 51,0606688 | 81,6972156 | 43,3605341 | 34,3497375 | 31,84726125 | 77,2407348 | 58,666849 | 67,2688602 | 65,8004871 | 100,474311 | 97,21932 | 23,0403987 | 68,28992 | 84,42846 | 72,92366325 | 205,982 | 153,5819808 | 110,2657 | 171,9973 | 31,00048 | 95,1204226 |
| Upregulated | ENSMUSG00000032291 | 1833,662261 | 1856,26806 | 3432,1908 | 4129,84998 | 1433,78349 | 1373,413141 | 2200,22505 | 2273,3404 | 1444,67885 | 3873,59916 | 1847,18157 | 3143,425 | 674,182101 | 2454,695 | 3424,928 | 3729,133829 | 13451,7 | 5661,725407 | 7493,316 | 10739,17 | 770,5834 | 4111,71487 |
| Upregulated | ENSMUSG00000002020 | 2770,585423 | 3218,94966 | 2472,70239 | 2981,27761 | 1636,06527 | 1346,873757 | 3331,57463 | 4448,90272 | 1818,39474 | 4812,06513 | 1884,72142 | 2977,013 | 611,071443 | 2051,504 | 2397,45 | 3010,835746 | 15815,94 | 5150,446039 | 4557,016 | 10390,5 | 891,0424 | 3149,7423 |
| Upregulated | ENSMUSG00000060487 | 21,14950705 | 9,57387541 | 25,4169115 | 9,63567424 | 8,90548749 | 9,28878453 | 9,08714527 | 23,2222944 | 16,0163953 | 11,8656616 | 0 | 13,13775 | 32,0562069 | 19,64505 | 11,94742 | 17,31937002 | 67,83342 | 56,47853486 | 14,2585 | 60,66572 | 9,743008 | 34,0997741 |
| Upregulated | ENSMUSG00000026196 | 0 | 6,3825836 | 9,0774684 | 0,96356742 | 0 | 0 | 3,40767947 | 13,4444862 | 1,06775969 | 16,1804476 | 3,31233993 | 0 | 0 | 0 | 7,168455 | 0 | 31,435 | 19,81702978 | 7,604532 | 43,99931 | 3,542912 | 11,6657122 |
| Upregulated | ENSMUSG00000032113 | 26,43688381 | 59,5707803 | 33,5866331 | 26,0163204 | 16,5387625 | 27,86635359 | 17,0383974 | 24,4445204 | 18,1519147 | 11,8656616 | 15,4575863 | 35,03399 | 30,0526939 | 14,96765 | 47,7897 | 4,557728953 | 110,8497 | 38,64320806 | 93,15552 | 99,99844 | 21,25747 | 25,1261494 |
| Upregulated | ENSMUSG00000041859 | 318,300081 | 457,418492 | 374,899445 | 437,459611 | 134,854525 | 183,1217522 | 164,704508 | 286,000889 | 234,907131 | 212,503212 | 175,554016 | 434,4214 | 381,669213 | 312,4498 | 442,0547 | 490,4116354 | 1034,046 | 644,0534677 | 644,4841 | 933,9854 | 294,0617 | 266,516656 |
| Upregulated | ENSMUSG00000022673 | 679,9566515 | 749,953573 | 666,286181 | 644,626607 | 334,591887 | 306,5298895 | 289,652755 | 624,557497 | 568,048153 | 535,033469 | 337,858672 | 705,9349 | 494,867693 | 427,5136 | 744,7228 | 577,0084855 | 1644,547 | 855,1048348 | 1133,075 | 1529,309 | 622,6668 | 467,525851 |
| Upregulated | ENSMUSG00000116912 | 62,39104579 | 84,0373508 | 55,3725572 | 71,3039894 | 36,8941625 | 33,17423047 | 42,0280469 | 72,1113353 | 48,0491859 | 61,485701 | 29,8110593 | 111,2329 | 81,1422736 | 57,99966 | 74,87053 | 59,25047639 | 203,5003 | 117,9113272 | 131,1782 | 212,6633 | 62,00096 | 50,2522987 |
| Upregulated | ENSMUSG00000053024 | 0 | 0 | 0 | 0 | 0 | 0 | 4,54357263 | 0 | 0 | 0 | 0 | 14,88944 | 13,022834 | 4,677392 | 14,33691 | 0 | 49,63421 | 12,88106935 | 20,91246 | 35,33278 | 2,657184 | 5,38417486 |
| Upregulated | ENSMUSG00000029730 | 364,8289965 | 331,894347 | 292,294482 | 244,746126 | 228,99825 | 180,4678137 | 257,847747 | 292,112019 | 197,535542 | 298,798933 | 255,050174 | 531,6408 | 361,634084 | 335,8367 | 413,3809 | 316,3063894 | 966,2126 | 570,7304576 | 670,1494 | 751,3216 | 322,405 | 303,308517 |
| Upregulated | ENSMUSG00000036875 | 7,402327466 | 7,44634754 | 14,5239494 | 21,1984833 | 0 | 1,326969219 | 5,67946579 | 15,8889383 | 4,27103874 | 5,39348255 | 0 | 6,130948 | 1,00175646 | 17,77409 | 1,59299 | 0,911545791 | 63,69724 | 9,908514888 | 36,12153 | 41,33269 | 20,37174 | 7,17889982 |
| Upregulated | ENSMUSG00000017499 | 13,74717958 | 35,1042098 | 17,24719 | 4,81783712 | 15,26655 | 9,28878453 | 23,8537563 | 57,444623 | 14,9486356 | 12,9443581 | 4,41645324 | 9,634347 | 14,0245905 | 0 | 11,15093 | 14,58473265 | 155,5205 | 62,4236438 | 63,68796 | 116,6648 | 45,17213 | 35,8944991 |
| Upregulated | ENSMUSG00000113996 | 1,057475352 | 8,51011147 | 13,6162026 | 2,89070227 | 2,544425 | 0 | 10,2230384 | 17,1111643 | 7,4743178 | 0 | 7,72879316 | 4,379248 | 4,00702586 | 5,61287 | 0,796495 | 1,823091581 | 62,04276 | 22,78958424 | 28,51699 | 55,33247 | 5,314368 | 8,97362477 |
| Upregulated | ENSMUSG00000033970 | 51,81629226 | 62,7620721 | 158,855697 | 91,5389053 | 22,899825 | 57,0596764 | 92,0073458 | 108,778116 | 85,4207749 | 51,7774324 | 60,726232 | 49,92343 | 52,0913362 | 111,3219 | 123,4567 | 63,80820534 | 213,4271 | 142,6826144 | 173,0031 | 276,6623 | 108,9445 | 108,58086 |
| Upregulated | ENSMUSG00000046591 | 8,459802819 | 3,1912918 | 25,4169115 | 14,4535114 | 0 | 0 | 11,3589316 | 11,0000342 | 27,7617518 | 2,15739302 | 0 | 3,503399 | 0 | 0 | 10,35443 | 0 | 76,93303 | 63,41449528 | 67,49022 | 121,3314 | 15,05738 | 26,9208743 |
| Upregulated | ENSMUSG00000051220 | 20,09203169 | 10,6376393 | 24,5091647 | 15,4170788 | 0 | 1,326969219 | 19,3101837 | 24,4445204 | 29,8972712 | 2,15739302 | 11,0411331 | 5,255098 | 5,00878232 | 17,77409 | 23,89485 | 2,734637372 | 91,82329 | 71,34130719 | 102,6612 | 123,9981 | 15,9431 | 52,0470237 |
| Upregulated | ENSMUSG00000022945 | 109,9774366 | 126,587908 | 147,962735 | 145,498681 | 99,2325749 | 61,04058406 | 74,9689484 | 105,111438 | 75,8109377 | 87,3744172 | 96,0578579 | 131,3775 | 95,1668641 | 105,7091 | 199,1237 | 121,2355902 | 716,3871 | 432,0112491 | 499,998 | 634,6568 | 237,3751 | 204,598645 |
| Upregulated | ENSMUSG00000026355 | 863,9573628 | 739,315934 | 868,713726 | 769,890372 | 249,35365 | 222,9308287 | 535,005678 | 431,445786 | 355,563975 | 540,426951 | 306,9435 | 714,6933 | 587,029288 | 439,6748 | 875,3479 | 654,4898777 | 2351,007 | 1334,676955 | 1606,457 | 2211,299 | 901,6711 | 681,995483 |
| Upregulated | ENSMUSG00000019214 | 99,40268312 | 86,1648786 | 112,560608 | 118,518793 | 47,0718625 | 41,13604578 | 39,7562605 | 69,6668832 | 45,9136665 | 36,6756813 | 13,2493597 | 47,29588 | 26,0456681 | 73,90279 | 45,40021 | 40,10801479 | 288,7057 | 195,1977433 | 266,1586 | 364,661 | 117,8018 | 64,6100984 |
| Upregulated | ENSMUSG00000005410 | 255,9090353 | 387,210072 | 360,375495 | 310,268711 | 35,62195 | 62,36755328 | 110,181636 | 199,222842 | 127,063403 | 88,4531138 | 60,726232 | 112,9846 | 135,237123 | 126,2896 | 232,5765 | 156,785876 | 1235,892 | 624,236438 | 730,0351 | 1423,311 | 373,7772 | 197,419745 |
| Upregulated | ENSMUSG00000002870 | 269,6562148 | 394,656419 | 297,740964 | 311,232278 | 62,3384124 | 14,5966614 | 182,878798 | 211,445102 | 127,063403 | 155,332297 | 76,1838183 | 227,7209 | 109,191455 | 212,3536 | 343,2893 | 195,0707992 | 1172,195 | 593,5200418 | 720,5294 | 1187,981 | 449,9498 | 178,575133 |
| Upregulated | ENSMUSG00000007080 | 35,95416198 | 81,9098229 | 96,221165 | 34,6884273 | 10,1777 | 2,653938437 | 26,1255426 | 74,5557873 | 17,084155 | 43,1478604 | 5,52056654 | 40,28909 | 18,0316164 | 33,67722 | 18,31938 | 45,57728953 | 313,5228 | 192,2251888 | 198,6684 | 364,661 | 121,3447 | 63,7127359 |
| Upregulated | ENSMUSG00000056394 | 191,4030388 | 288,280026 | 216,043748 | 173,442136 | 122,1324 | 183,1217522 | 141,986645 | 172,333869 | 177,248108 | 151,017511 | 152,367637 | 188,3077 | 90,1580818 | 147,8056 | 211,0712 | 110,2970407 | 800,7653 | 564,7853486 | 501,8991 | 843,9868 | 329,4908 | 191,138208 |
| Upregulated | ENSMUSG00000001525 | 1455,086085 | 1265,87908 | 2714,16305 | 2971,64194 | 1293,84011 | 1683,923938 | 1682,25777 | 2137,67331 | 2035,14996 | 1511,25381 | 1526,98871 | 1409,242 | 1767,0984 | 3364,916 | 2977,298 | 2452,969723 | 5754,259 | 4258,679699 | 4095,991 | 5509,247 | 2506,61 | 1924,84251 |
| Upregulated | ENSMUSG00000027454 | 13,74717958 | 9,57387541 | 27,2324052 | 14,4535114 | 12,722125 | 11,94272297 | 2,27178632 | 12,2222602 | 17,084155 | 31,2821988 | 6,62467985 | 12,2619 | 14,0245905 | 34,6127 | 38,23176 | 25,52328214 | 76,93303 | 57,46938635 | 57,98456 | 83,99869 | 20,37174 | 26,0235118 |
| Upregulated | ENSMUSG00000022978 | 135,3568451 | 129,7792 | 199,704305 | 131,04517 | 83,9660249 | 145,966614 | 86,32788 | 149,111575 | 134,53772 | 127,286188 | 141,326504 | 212,8315 | 110,193211 | 187,0957 | 254,0819 | 166,8128797 | 391,283 | 320,0450309 | 296,5767 | 320,6617 | 271,0328 | 172,293596 |
| Upregulated | ENSMUSG00000029005 | 15,86213028 | 1,06376393 | 29,0478989 | 3,8542697 | 11,4499125 | 9,28878453 | 2,27178632 | 7,33335613 | 6,40655811 | 16,1804476 | 3,31233993 | 13,13775 | 1,00175646 | 25,25792 | 7,964949 | 2,734637372 | 57,07934 | 50,53342593 | 26,61586 | 53,3325 | 14,17165 | 7,17889982 |
| Upregulated | ENSMUSG00000035783 | 70,85084861 | 167,010938 | 118,914836 | 592,593966 | 36,8941625 | 155,2553986 | 59,0664442 | 158,889383 | 144,147558 | 61,485701 | 118,140124 | 133,1292 | 81,1422736 | 202,9988 | 151,334 | 525,9619212 | 776,7754 | 568,7487546 | 292,7745 | 815,9873 | 469,4358 | 388,557953 |
| Upregulated | ENSMUSG00000044447 | 82,48307748 | 52,1244328 | 87,1436966 | 58,7776129 | 10,1777 | 26,53938437 | 22,7178632 | 40,3334587 | 10,6775969 | 50,6987359 | 19,8740396 | 56,05438 | 32,0562069 | 40,22557 | 43,80722 | 102,0931286 | 143,112 | 75,30471315 | 68,44079 | 153,9976 | 46,94358 | 35,8944991 |
| Upregulated | ENSMUSG00000046934 | 24,3219331 | 27,6578623 | 15,4316963 | 26,0163204 | 11,4499125 | 10,61575375 | 7,95125211 | 20,7778424 | 7,4743178 | 15,1017511 | 24,2904928 | 22,77209 | 26,0456681 | 21,516 | 20,70887 | 38,28492321 | 44,67079 | 18,82617829 | 34,22039 | 48,66591 | 24,80038 | 14,3577996 |
| Upregulated | ENSMUSG00000057497 | 177,6558592 | 178,712341 | 215,136001 | 258,23607 | 117,04355 | 138,0047987 | 65,8818032 | 177,222773 | 136,67324 | 124,050099 | 145,742957 | 291,6579 | 309,542748 | 238,547 | 259,6574 | 163,1666965 | 424,3725 | 271,4933079 | 289,9228 | 325,3283 | 326,8336 | 304,20588 |
| Upregulated | ENSMUSG00000040213 | 77,19570072 | 48,933141 | 86,2359498 | 113,700956 | 39,4385875 | 18,57756906 | 15,9025042 | 52,5557189 | 29,8972712 | 35,5969848 | 25,3946061 | 69,19213 | 63,1106573 | 77,64471 | 83,63197 | 18,23091581 | 151,3843 | 82,24067357 | 87,45212 | 127,998 | 90,34426 | 82,5573479 |
| Upregulated | ENSMUSG00000028937 | 543,5423311 | 359,55221 | 669,917168 | 401,807616 | 150,121075 | 192,4105367 | 311,234725 | 211,445102 | 198,603302 | 254,572376 | 277,132441 | 331,0712 | 323,567338 | 373,2559 | 468,339 | 386,4954152 | 870,2532 | 444,8923185 | 693,9135 | 717,3221 | 410,0921 | 507,0098 |
| Upregulated | ENSMUSG00000039153 | 115,2648134 | 59,5707803 | 103,48314 | 74,1946916 | 29,2608875 | 10,61575375 | 17,0383974 | 53,777945 | 23,4907131 | 39,9117708 | 25,3946061 | 123,4948 | 73,1282219 | 177,7409 | 124,2532 | 82,95066695 | 203,5003 | 134,7558025 | 117,8702 | 259,3293 | 144,3737 | 73,5837231 |
| Upregulated | ENSMUSG00000116475 | 34,89668663 | 36,1679738 | 60,8190383 | 32,7612924 | 8,90548749 | 14,5966614 | 24,9896495 | 19,5556163 | 30,9650309 | 18,3378407 | 8,83290647 | 42,04079 | 34,0597198 | 38,35461 | 53,36516 | 41,01956058 | 56,25211 | 54,49683188 | 63,68796 | 85,99866 | 41,62922 | 29,6129618 |
| Upregulated | ENSMUSG00000025089 | 207,2651691 | 78,7185311 | 238,737419 | 254,3818 | 155,209925 | 228,2387056 | 280,56561 | 160,111609 | 192,196743 | 456,288623 | 266,091307 | 1405,739 | 1401,45729 | 724,9958 | 1165,272 | 1993,550644 | 1359,15 | 575,684715 | 921,0989 | 5796,576 | 534,094 | 526,751774 |
| Upregulated | ENSMUSG00000069601 | 1,057475352 | 4,25505574 | 13,6162026 | 0,96356742 | 0 | 2,653938437 | 0 | 2,44445204 | 0 | 0 | 4,41645324 | 9,634347 | 45,0790409 | 5,61287 | 43,01073 | 45,57728953 | 32,26224 | 4,954257444 | 6,653965 | 249,3294 | 38,0863 | 33,2024117 |
| Upregulated | ENSMUSG00000021364 | 0 | 2,12752787 | 4,5387342 | 1,92713485 | 3,8166375 | 2,653938437 | 3,40767947 | 18,3333903 | 6,40655811 | 4,31478604 | 12,1452464 | 33,28229 | 9,01580818 | 0,935478 | 13,54041 | 19,1424616 | 7,445132 | 11,89021787 | 2,851699 | 69,99891 | 9,743008 | 11,6657122 |
| Upregulated | ENSMUSG00000031548 | 366,9439473 | 297,853902 | 384,88466 | 226,438345 | 131,037887 | 185,7756906 | 127,220034 | 429,001334 | 301,108231 | 224,368874 | 800,482149 | 1099,191 | 657,152241 | 1371,411 | 483,4724 | 439,3650711 | 775,9482 | 481,5538236 | 704,3698 | 2308,631 | 864,4705 | 282,66918 |
| Upregulated | ENSMUSG00000021478 | 0 | 7,44634754 | 29,9556457 | 4,81783712 | 3,8166375 | 0 | 0 | 0 | 1,06775969 | 3,23608953 | 5,52056654 | 65,68873 | 8,01405172 | 38,35461 | 4,77897 | 14,58473265 | 50,46145 | 17,8353268 | 24,71473 | 112,6649 | 15,9431 | 15,2551621 |
| Upregulated | ENSMUSG00000038357 | 1,057475352 | 1,06376393 | 9,98521524 | 6,74497197 | 0 | 2,653938437 | 0 | 0 | 2,13551937 | 0 | 2,20822662 | 29,77889 | 5,00878232 | 21,516 | 15,1334 | 13,67318686 | 17,37197 | 0,990851489 | 27,56643 | 82,66538 | 1,771456 | 6,28153734 |
| Upregulated | ENSMUSG00000040133 | 118,4372395 | 127,651672 | 132,531039 | 100,211012 | 71,2438999 | 132,6969219 | 63,6100169 | 40,3334587 | 72,6076586 | 169,355352 | 187,699262 | 284,6511 | 53,0930926 | 115,0638 | 553,564 | 427,5149758 | 851,2267 | 404,2674074 | 398,2874 | 1669,307 | 183,3457 | 255,748306 |
| Upregulated | ENSMUSG00000046687 | 114,2073381 | 84,0373508 | 293,202229 | 180,187108 | 63,6106249 | 71,65633781 | 97,6868116 | 94,1114037 | 178,315868 | 138,073153 | 113,723671 | 380,1188 | 66,1159267 | 200,1924 | 423,7353 | 428,4265216 | 536,0495 | 311,1273675 | 389,7323 | 1215,981 | 314,4334 | 204,598645 |
| Upregulated | ENSMUSG00000076441 | 26,43688381 | 34,0404459 | 57,1880509 | 36,6155621 | 27,988675 | 23,88544594 | 21,58197 | 29,3334245 | 26,6939921 | 9,70826858 | 18,7699262 | 90,21252 | 20,0351293 | 49,58035 | 121,0672 | 115,7663154 | 142,2847 | 57,46938635 | 101,7106 | 271,9958 | 69,97251 | 36,7918616 |
| Upregulated | ENSMUSG00000020181 | 190,3455634 | 60,6345442 | 133,438785 | 112,737389 | 137,39895 | 90,23390687 | 127,220034 | 92,8891777 | 118,521325 | 193,086675 | 39,7480791 | 196,1903 | 142,249418 | 225,4503 | 193,5483 | 370,087591 | 449,1896 | 193,2160403 | 273,7632 | 641,99 | 299,3761 | 289,84808 |
| Upregulated | ENSMUSG00000044583 | 0 | 0 | 0 | 0 | 0 | 0 | 0 | 0 | 0 | 0 | 0 | 3,503399 | 2,00351293 | 0 | 0 | 8,203912116 | 19,02645 | 0 | 14,2585 | 27,99956 | 9,743008 | 0 |
| Upregulated | ENSMUSG00000015709 | 0 | 4,25505574 | 7,26197472 | 0 | 1,2722125 | 0 | 0 | 0 | 0 | 0 | 0 | 65,68873 | 39,0685021 | 62,67705 | 51,77217 | 48,3119269 | 144,7664 | 43,59746551 | 61,78682 | 242,6629 | 50,4865 | 20,639337 |
| Upregulated | ENSMUSG00000052698 | 353,1967677 | 210,625259 | 204,243039 | 232,219749 | 133,582312 | 188,429629 | 179,471119 | 114,889246 | 165,502751 | 174,748834 | 138,014164 | 512,3721 | 381,669213 | 424,7072 | 445,2407 | 602,5317676 | 1006,747 | 487,4989325 | 564,6365 | 1429,311 | 450,8356 | 272,798193 |
| Upregulated | ENSMUSG00000036356 | 687,358979 | 469,119895 | 650,854484 | 559,832673 | 87,7826624 | 228,2387056 | 126,084141 | 185,778355 | 144,147558 | 228,68366 | 192,115716 | 1150,866 | 821,440301 | 1024,349 | 930,3061 | 1638,047786 | 2944,136 | 1300,988005 | 2297,519 | 4539,262 | 1745,77 | 900,951927 |
| Upregulated | ENSMUSG00000033965 | 694,7613065 | 518,053036 | 642,684763 | 783,380316 | 329,503037 | 384,8210734 | 550,908182 | 405,779039 | 397,206603 | 596,51917 | 600,63764 | 2182,617 | 822,442057 | 1447,185 | 1166,069 | 1663,571068 | 3854,924 | 1926,215294 | 2698,658 | 4569,262 | 2323,265 | 1115,42156 |
| Upregulated | ENSMUSG00000061531 | 5,287376762 | 5,31881967 | 28,140152 | 0,96356742 | 13,9943375 | 18,57756906 | 11,3589316 | 20,7778424 | 17,084155 | 0 | 14,353473 | 49,04758 | 13,022834 | 30,87079 | 42,21423 | 29,1694653 | 67,83342 | 32,69809913 | 29,46756 | 95,9985 | 65,54387 | 24,2287869 |
| Upregulated | ENSMUSG00000100303 | 6,344852114 | 0 | 5,44648104 | 9,63567424 | 0 | 0 | 0 | 3,66667807 | 2,13551937 | 1,07869651 | 8,83290647 | 21,02039 | 10,0175646 | 4,677392 | 2,389485 | 13,67318686 | 35,57118 | 14,86277233 | 13,30793 | 39,99938 | 20,37174 | 3,58944991 |
| Upregulated | ENSMUSG00000021678 | 19,03455634 | 20,2115147 | 7,26197472 | 29,8705901 | 0 | 2,653938437 | 6,81535895 | 0 | 8,54207749 | 1,07869651 | 0 | 19,26869 | 4,00702586 | 11,22574 | 41,41774 | 33,72719425 | 97,61395 | 28,73469318 | 31,36869 | 117,9982 | 50,4865 | 23,3314244 |
| Upregulated | ENSMUSG00000074221 | 61,33357043 | 36,1679738 | 67,1732662 | 61,6683151 | 36,8941625 | 65,02149171 | 57,9305511 | 39,1112327 | 153,757395 | 75,5087556 | 68,4550251 | 163,7839 | 67,1176831 | 76,70923 | 119,4742 | 119,4124986 | 304,4232 | 107,0119608 | 117,8702 | 344,6613 | 101,8587 | 92,4283352 |

| | | | | | | | | | | | | | | | | | | | | | | | |
|---|---|---|---|---|---|---|---|---|---|---|---|---|---|---|---|---|---|---|---|---|---|---|---|
| Upregulated | ENSMUSG00000021125 | 5,287376762 | 10,6376393 | 13,6162026 | 10,5992417 | 7,63327499 | 22,55847672 | 10,2230384 | 4,88890409 | 11,7453565 | 14,0230546 | 5,52056654 | 42,91663 | 26,0456681 | 28,06435 | 50,97568 | 6,380820534 | 23,98987 | 20,80788127 | 51,33059 | 129,998 | 32,77194 | 45,7654863 |
| Upregulated | ENSMUSG00000063060 | 17,97708099 | 6,3825836 | 42,6641015 | 36,6155621 | 12,722125 | 9,28878453 | 15,9025042 | 7,33335613 | 17,084155 | 1,07869651 | 2,20822662 | 79,70232 | 36,0632327 | 12,16122 | 58,94063 | 30,08101109 | 30,60776 | 7,92681191 | 50,38002 | 166,6641 | 34,54339 | 42,1760364 |
| Upregulated | ENSMUSG00000024883 | 187,1731374 | 30,8491541 | 189,71909 | 143,571546 | 49,6162874 | 75,63724546 | 76,1048416 | 30,5556505 | 29,8972712 | 56,0922185 | 23,1863795 | 197,942 | 92,1615947 | 174,9345 | 262,8433 | 219,6825355 | 263,8886 | 134,7558025 | 312,7364 | 566,6578 | 108,9445 | 152,551621 |
| Upregulated | ENSMUSG00000051232 | 95,17278171 | 119,141561 | 121,638077 | 116,591658 | 86,5104499 | 66,34846093 | 96,5509185 | 95,3336297 | 77,9464571 | 50,6987359 | 59,6221187 | 132,2533 | 123,216045 | 152,483 | 195,9378 | 112,1201322 | 173,7197 | 100,0760004 | 231,9382 | 321,3283 | 157,6596 | 129,220197 |
| Upregulated | ENSMUSG00000024713 | 2282,03181 | 1321,19481 | 3464,86969 | 3226,9873 | 665,367137 | 623,6755328 | 1401,69216 | 1546,11592 | 1198,02637 | 1196,27443 | 1161,5272 | 6437,495 | 3946,92047 | 8102,178 | 5221,821 | 4822,077232 | 9669,571 | 5635,963268 | 7805,101 | 17727,06 | 6712,933 | 2727,08457 |
| Upregulated | ENSMUSG00000025068 | 576,324067 | 658,469875 | 814,248915 | 937,551104 | 695,900237 | 764,3342699 | 692,894827 | 685,668798 | 578,72575 | 460,603409 | 781,712223 | 1682,507 | 1573,75941 | 1970,117 | 1856,63 | 1697,298262 | 2059,82 | 1395,118896 | 1909,688 | 3885,273 | 1398,565 | 1285,02307 |
| Upregulated | ENSMUSG00000099871 | 5,287376762 | 3,1912918 | 11,8007089 | 3,8542697 | 5,08884999 | 3,980907656 | 13,6307179 | 23,2222944 | 8,54207749 | 7,55087556 | 11,0411331 | 90,21252 | 43,075528 | 9,354784 | 1,59299 | 14,58473265 | 82,72368 | 49,54257444 | 23,76416 | 165,9974 | 18,60029 | 54,7391111 |
| Upregulated | ENSMUSG00000097418 | 19,03455634 | 24,4665705 | 36,3098736 | 13,4899439 | 10,1777 | 13,26969219 | 21,58197 | 29,3334245 | 8,54207749 | 46,3839499 | 27,6028327 | 138,3843 | 24,0421551 | 26,19339 | 69,29506 | 37,37337742 | 94,305 | 55,48768337 | 88,40268 | 214,6633 | 54,91514 | 73,5837231 |
| Upregulated | ENSMUSG00000106987 | 0 | 5,31881967 | 4,5387342 | 0 | 1,2722125 | 1,326969219 | 2,27178632 | 4,88890409 | 7,4743178 | 4,31478604 | 5,52056654 | 22,77209 | 8,01405172 | 4,677392 | 7,964949 | 3,646183163 | 22,33539 | 4,954257444 | 14,2585 | 48,66591 | 2,657184 | 6,28153734 |
| Upregulated | ENSMUSG00000097081 | 6,344852114 | 3,1912918 | 2,72324052 | 10,5992417 | 6,36106249 | 1,326969219 | 0 | 3,66667807 | 3,20327906 | 0 | 5,52056654 | 27,15134 | 10,0175646 | 7,483827 | 11,15093 | 2,734637372 | 21,50816 | 8,917663399 | 13,30793 | 26,66625 | 15,05738 | 11,6657122 |
| Upregulated | ENSMUSG00000070583 | 22,2069824 | 13,8289311 | 11,8007089 | 29,8705901 | 16,5387625 | 9,28878453 | 21,58197 | 22,0000684 | 7,4743178 | 21,5739302 | 8,83290647 | 55,17853 | 28,049181 | 8,419306 | 33,45279 | 13,67318686 | 33,08947 | 40,62491104 | 33,26983 | 83,33203 | 31,88621 | 30,5103242 |
| Upregulated | ENSMUSG00000039934 | 88,82792959 | 110,631449 | 159,763444 | 81,903231 | 30,5331 | 37,15513812 | 40,8921537 | 111,222568 | 117,453565 | 78,7448452 | 48,5809856 | 91,08837 | 80,1405172 | 55,19322 | 48,58619 | 99,35849118 | 218,3905 | 139,7100599 | 186,311 | 362,661 | 130,202 | 122,938659 |
| Upregulated | ENSMUSG00000042029 | 256,9665106 | 359,55221 | 367,63747 | 158,988625 | 269,70905 | 179,1408445 | 181,742905 | 473,00147 | 374,78365 | 268,595431 | 224,135002 | 179,5492 | 175,307381 | 196,4505 | 219,0361 | 272,5521914 | 782,5661 | 526,1421406 | 679,655 | 1274,647 | 379,0916 | 454,962776 |
| Upregulated | ENSMUSG00000083668 | 30,66678522 | 18,0839869 | 41,7563546 | 25,052753 | 0 | 10,61575375 | 7,95125211 | 36,6667807 | 14,9486356 | 19,4165372 | 6,62467985 | 13,13775 | 13,022834 | 11,22574 | 28,67382 | 1,823091581 | 75,27855 | 64,40534677 | 62,73739 | 169,9973 | 41,62922 | 23,3314244 |
| Upregulated | ENSMUSG00000005470 | 51,81629226 | 105,312629 | 157,040203 | 82,8667985 | 20,3554 | 21,2315075 | 80,6484142 | 51,3334929 | 42,7103874 | 35,5969848 | 15,4575863 | 18,39284 | 30,0526939 | 89,80593 | 47,7897 | 39,196469 | 352,4029 | 309,1456645 | 288,0216 | 751,9883 | 247,1181 | 75,3784481 |
| Upregulated | ENSMUSG00000045751 | 95,17278171 | 94,6749901 | 117,099342 | 94,4296076 | 29,2608875 | 29,19332281 | 82,9202006 | 66,0002052 | 60,8623021 | 73,3513626 | 54,1015521 | 64,81288 | 32,0562069 | 105,7091 | 100,3584 | 64,71975114 | 262,2341 | 193,2160403 | 175,8548 | 510,6587 | 137,2878 | 116,657122 |
| Upregulated | ENSMUSG00000042489 | 16,91960564 | 8,51011147 | 16,3394431 | 14,4535114 | 2,544425 | 1,326969219 | 11,3589316 | 11,0000342 | 5,33879843 | 8,62957207 | 0 | 2,627549 | 0 | 2,806435 | 0,796495 | 0 | 47,97974 | 33,68895062 | 34,22039 | 94,66519 | 31,88621 | 27,8182368 |
| Upregulated | ENSMUSG00000025001 | 77,19570072 | 103,185102 | 106,20638 | 108,883119 | 39,4385875 | 10,61575375 | 68,1535895 | 57,444623 | 102,50493 | 75,5087556 | 32,019286 | 32,40644 | 44,0772844 | 59,87062 | 84,42846 | 29,1694653 | 339,1671 | 236,8135058 | 286,1205 | 605,9905 | 124,0019 | 132,809647 |
| Upregulated | ENSMUSG00000001228 | 233,7020529 | 171,265993 | 249,630381 | 135,863007 | 35,62195 | 57,0596764 | 90,8714527 | 138,11154 | 107,843728 | 78,7448452 | 39,7480791 | 90,21252 | 72,1264654 | 103,8381 | 140,1831 | 67,45438851 | 671,7163 | 463,7184968 | 490,4923 | 1018,651 | 317,9764 | 176,780408 |
| Upregulated | ENSMUSG00000116864 | 14,80465493 | 7,44634754 | 10,8929621 | 11,5628091 | 0 | 0 | 7,95125211 | 1,22222602 | 6,40655811 | 6,47217906 | 0 | 0,87585 | 0 | 7,483827 | 3,982475 | 1,823091581 | 50,46145 | 35,6706536 | 44,67663 | 85,99866 | 24,80038 | 17,9472495 |
| Upregulated | ENSMUSG00000030528 | 39,12658804 | 49,9969049 | 60,8190383 | 37,5791295 | 17,810975 | 17,25059984 | 38,6203674 | 44,0001368 | 24,5584728 | 43,1478604 | 23,1863795 | 29,77889 | 11,0193211 | 57,99966 | 55,75465 | 20,96555318 | 120,7766 | 91,15833697 | 105,5129 | 199,9969 | 46,05786 | 82,5573479 |
| Upregulated | ENSMUSG00000037572 | 65,56347184 | 32,976682 | 71,7120004 | 35,6519947 | 21,6276125 | 17,25059984 | 40,8921537 | 47,6668148 | 36,3038293 | 35,5969848 | 28,706946 | 53,42683 | 9,01580818 | 23,38696 | 31,8598 | 53,78120165 | 138,1486 | 115,9296242 | 145,4367 | 223,9965 | 49,60077 | 66,4048233 |
| Upregulated | ENSMUSG00000027635 | 99,40268312 | 96,802518 | 121,638077 | 64,5590174 | 40,7108 | 57,0596764 | 45,4357263 | 84,3335955 | 45,9136665 | 50,6987359 | 44,1645324 | 114,7363 | 51,0895797 | 70,16088 | 48,58619 | 77,4813922 | 191,9189 | 168,4447531 | 187,2616 | 330,6615 | 103,6302 | 156,141071 |
| Upregulated | ENSMUSG00000026669 | 31,72426057 | 48,933141 | 30,8633926 | 34,6884273 | 15,26655 | 15,92363062 | 18,1742905 | 35,4445546 | 10,6775969 | 11,8656616 | 9,93701978 | 48,17173 | 11,0193211 | 22,45148 | 9,557939 | 23,70019056 | 89,34158 | 93,14003995 | 107,414 | 213,9967 | 46,05786 | 55,6364736 |
| Upregulated | ENSMUSG00000094777 | 24,3219331 | 51,0606688 | 113,468355 | 34,6884273 | 27,988675 | 38,48210734 | 128,355927 | 67,2224312 | 90,7595733 | 32,3608953 | 29,8110593 | 7,882647 | 12,0210776 | 13,0967 | 19,91237 | 20,96555318 | 295,3236 | 257,6213871 | 358,3636 | 459,9928 | 232,0607 | 160,627883 |
| Upregulated | ENSMUSG00000094248 | 34,89668663 | 62,7620721 | 127,992304 | 49,1419386 | 34,3497375 | 31,84726125 | 132,8995 | 83,1113695 | 75,8109377 | 32,3608953 | 29,8110593 | 12,2619 | 14,0245905 | 17,77409 | 15,1334 | 28,25791951 | 301,9414 | 296,2645952 | 385,93 | 562,6579 | 259,5183 | 163,319971 |
| Upregulated | ENSMUSG00000069301 | 9,517278171 | 13,8289311 | 40,8486078 | 16,3806462 | 8,90548749 | 18,57756906 | 27,2614358 | 14,6667123 | 6,40655811 | 6,47217906 | 7,72879316 | 3,503399 | 5,00878232 | 8,419306 | 3,18598 | 0,911545791 | 75,27855 | 61,43279231 | 118,8208 | 159,3308 | 84,14416 | 35,8944991 |
| Upregulated | ENSMUSG00000071478 | 5,287376762 | 2,12752787 | 6,35422788 | 3,8542697 | 0 | 2,653938437 | 9,08714527 | 2,44445204 | 3,20327906 | 2,15739302 | 0 | 2,627549 | 1,00175646 | 5,61287 | 0 | 3,646183163 | 23,16263 | 15,85362382 | 29,46756 | 42,666 | 18,60029 | 5,38417486 |
| Upregulated | ENSMUSG00000112222 | 27,49435916 | 29,7853902 | 26,3246584 | 37,5791295 | 13,9943375 | 14,5966614 | 28,397329 | 35,4445546 | 32,0327906 | 28,0461092 | 11,0411331 | 16,64114 | 27,0474245 | 15,90313 | 20,70887 | 28,25791951 | 74,45132 | 63,41449528 | 51,33059 | 113,3316 | 62,88669 | 49,3549363 |
| Upregulated | ENSMUSG00000031821 | 95,17278171 | 126,587908 | 102,575393 | 104,065282 | 24,1720375 | 53,07876874 | 36,3485811 | 78,2224654 | 69,4043796 | 49,6200394 | 29,8110593 | 49,04758 | 98,1721335 | 111,3219 | 90,00393 | 128,5279565 | 277,1243 | 255,6396841 | 231,9382 | 430,6599 | 265,7184 | 108,58086 |
| Upregulated | ENSMUSG00000047894 | 3,172426057 | 0 | 2,72324052 | 0 | 0 | 0 | 0 | 0 | 0 | 0 | 1,10411331 | 0 | 1,00175646 | 0 | 0 | 0 | 23,98987 | 13,87192084 | 12,35736 | 35,33278 | 15,9431 | 1,79472495 |
| Upregulated | ENSMUSG00000073705 | 10,57475352 | 30,8491541 | 47,2028357 | 26,0163204 | 19,0831875 | 29,19332281 | 13,6307179 | 12,2222602 | 11,7453565 | 15,1017511 | 12,1452464 | 2,627549 | 9,01580818 | 34,6127 | 26,28433 | 4,557728953 | 71,96961 | 48,55172295 | 43,72606 | 105,9983 | 36,31485 | 16,1525246 |
| Upregulated | ENSMUSG00000029366 | 113,1498627 | 120,205325 | 186,995849 | 114,664523 | 61,0661999 | 49,09786109 | 81,7843074 | 132,00041 | 76,8786974 | 101,397472 | 78,3920449 | 41,16494 | 19,0333728 | 39,29009 | 79,64949 | 77,4813922 | 236,5897 | 213,0330701 | 151,1401 | 318,6617 | 175,3741 | 102,299322 |
| Upregulated | ENSMUSG00000021391 | 7,402327466 | 19,1477508 | 31,7711394 | 19,2713485 | 0 | 13,26969219 | 12,4948247 | 30,5556505 | 8,54207749 | 6,47217906 | 9,93701978 | 14,0136 | 2,00351293 | 0 | 0,796495 | 0 | 50,46145 | 28,73469318 | 34,22039 | 77,99878 | 43,40067 | 11,6657122 |
| Upregulated | ENSMUSG00000021175 | 25,37940846 | 29,7853902 | 12,7084558 | 26,0163204 | 8,90548749 | 5,307876874 | 11,3589316 | 36,6667807 | 25,6262325 | 12,9443581 | 2,20822662 | 2,627549 | 9,01580818 | 0 | 4,77897 | 0 | 43,01632 | 42,60661402 | 57,03399 | 106,665 | 39,85776 | 13,4604372 |
| Upregulated | ENSMUSG00000029516 | 153,3339261 | 109,567685 | 118,007089 | 83,8303659 | 73,7883249 | 79,61815312 | 106,773957 | 42,7779108 | 52,3202246 | 89,5318103 | 68,4550251 | 13,13775 | 19,0333728 | 42,09653 | 43,80722 | 46,48883532 | 210,9454 | 140,7009114 | 166,3491 | 401,3271 | 109,8303 | 70,8916357 |
| Upregulated | ENSMUSG00000034023 | 40,18406339 | 32,976682 | 27,2324052 | 32,7612924 | 24,1720375 | 39,80907656 | 32,9409016 | 35,4445546 | 21,3551937 | 42,0691639 | 16,5616996 | 1,751699 | 5,00878232 | 8,419306 | 16,72639 | 5,469274744 | 56,25211 | 47,56087146 | 93,15552 | 169,3307 | 48,71504 | 59,2259235 |
| Upregulated | ENSMUSG00000037466 | 106,8050106 | 77,6547672 | 109,837368 | 118,518793 | 40,7108 | 65,02149171 | 122,676461 | 86,7780475 | 50,1847052 | 120,814009 | 58,5180054 | 37,66154 | 30,0526939 | 22,45148 | 28,67382 | 4,557728953 | 144,7664 | 178,353268 | 192,0144 | 425,9934 | 186,0029 | 136,399097 |
| Upregulated | ENSMUSG00000041219 | 122,6671409 | 128,715436 | 158,855697 | 58,7776129 | 36,8941625 | 43,78998421 | 77,2407348 | 70,8891093 | 52,3202246 | 85,2170242 | 17,6658129 | 27,15134 | 9,01580818 | 21,516 | 22,30186 | 18,23091581 | 167,9291 | 206,0971097 | 210,0752 | 546,6581 | 156,7739 | 116,657122 |
| Upregulated | ENSMUSG00000020897 | 119,4947148 | 80,846059 | 224,213469 | 72,2675568 | 5,08884999 | 37,15513812 | 69,2894827 | 85,5558215 | 30,9650309 | 29,1248057 | 9,93701978 | 0,87585 | 4,00702586 | 6,548349 | 4,77897 | 5,469274744 | 237,417 | 253,6579811 | 268,0598 | 707,989 | 190,4315 | 127,425472 |
| Upregulated | ENSMUSG00000020914 | 235,8170036 | 241,474413 | 466,581876 | 104,065282 | 6,36106249 | 27,86635359 | 221,499166 | 183,333903 | 137,740999 | 67,9578801 | 9,93701978 | 25,39964 | 18,0316164 | 11,22574 | 12,74392 | 13,67318686 | 454,153 | 745,1203196 | 606,4614 | 1874,637 | 357,8341 | 384,968503 |
| Upregulated | ENSMUSG00000051378 | 25,37940846 | 34,0404459 | 57,1880509 | 27,9434553 | 2,544425 | 0 | 17,0383974 | 30,5556505 | 12,8131162 | 16,1804476 | 0 | 0 | 0 | 0 | 1,59299 | 0 | 43,01632 | 109,9845153 | 133,0793 | 241,3296 | 45,17213 | 39,483949 |
| Upregulated | ENSMUSG00000030346 | 7,402327466 | 27,6578623 | 55,3725572 | 21,1984833 | 6,36106249 | 0 | 17,0383974 | 26,8889725 | 16,0163953 | 7,55087556 | 0 | 0 | 0 | 3,741914 | 12,74392 | 9,115457906 | 58,73382 | 64,40534677 | 89,35325 | 189,997 | 53,14368 | 41,278674 |
| Upregulated | ENSMUSG00000030677 | 50,75881691 | 106,376393 | 229,659951 | 63,59545 | 24,1720375 | 22,55847672 | 68,1535895 | 41,5556847 | 29,8972712 | 53,9348255 | 24,2904928 | 27,15134 | 20,0351293 | 37,41914 | 56,55114 | 40,10801479 | 177,8559 | 162,4996442 | 214,828 | 561,3246 | 216,1176 | 128,322834 |
| Upregulated | ENSMUSG00000020493 | 16,91960564 | 26,5940983 | 67,1732662 | 21,1984833 | 3,8166375 | 5,307876874 | 11,3589316 | 25,6667465 | 6,40655811 | 20,4952337 | 4,41645324 | 0,87585 | 0 | 2,806435 | 19,11588 | 5,469274744 | 34,74395 | 41,61576253 | 62,73739 | 153,9976 | 25,68611 | 17,9472495 |
| Upregulated | ENSMUSG00000038379 | 8,459802819 | 32,976682 | 68,9887598 | 24,0891856 | 16,5387625 | 17,25059984 | 12,4948247 | 23,2222944 | 7,4743178 | 28,0461092 | 13,2493597 | 4,379248 | 10,0175646 | 5,61287 | 4,77897 | 11,85009528 | 33,08947 | 62,4236438 | 79,84759 | 178,6639 | 43,40067 | 31,4076867 |
| Upregulated | ENSMUSG00000032783 | 20,09203169 | 8,51011147 | 49,0183294 | 2,89070227 | 0 | 2,653938437 | 5,67946579 | 3,66667807 | 4,27103874 | 0 | 4,41645324 | 0 | 0 | 4,677392 | 0,796495 | 0 | 48,80697 | 35,6706536 | 54,18229 | 96,66516 | 17,71456 | 18,844612 |
| Upregulated | ENSMUSG00000034906 | 75,08075001 | 87,2286426 | 190,626836 | 43,3605341 | 22,899825 | 25,21241515 | 34,0767947 | 91,6669516 | 52,3202246 | 65,8004871 | 12,1452464 | 14,88944 | 32,0562069 | 48,64488 | 44,60372 | 30,99255688 | 211,7726 | 224,923288 | 235,7405 | 531,325 | 130,202 | 110,375585 |
| Upregulated | ENSMUSG00000029414 | 45,47144015 | 68,0808918 | 126,176811 | 14,4535114 | 0 | 3,980907656 | 32,9409016 | 53,777945 | 30,9650309 | 35,5969848 | 1,10411331 | 4,379248 | 15,026347 | 22,45148 | 11,15093 | 14,58473265 | 159,6567 | 167,4539016 | 178,7065 | 393,9939 | 46,94358 | 49,3549363 |
| Upregulated | ENSMUSG00000022385 | 19,03455634 | 23,4028065 | 87,1436966 | 10,5992417 | 2,544425 | 1,326969219 | 31,8050084 | 19,5556163 | 14,9486356 | 11,8656616 | 5,52056654 | 5,255098 | 8,01405172 | 4,677392 | 18,31938 | 0,911545791 | 137,3213 | 146,6460203 | 198,6684 | 323,9949 | 70,85824 | 52,9443862 |
| Upregulated | ENSMUSG00000017861 | 52,87376762 | 27,6578623 | 54,4648104 | 41,4333992 | 8,90548749 | 6,634846093 | 13,6307179 | 19,5556163 | 21,3551937 | 26,9674127 | 7,72879316 | 5,255098 | 3,00526939 | 9,354784 | 0 | 1,823091581 | 114,1587 | 63,41449528 | 168,2503 | 278,6623 | 66,4296 | 59,2259235 |
| Upregulated | ENSMUSG00000062510 | 23,26445775 | 13,8289311 | 39,0331141 | 8,67210682 | 10,1777 | 5,307876874 | 15,9025042 | 14,6667123 | 6,40655811 | 14,0230546 | 6,62467985 | 0,87585 | 0 | 5,61287 | 2,389485 | 3,646183163 | 59,56105 | 30,71639615 | 77,94645 | 147,331 | 9,743008 | 41,278674 |
| Upregulated | ENSMUSG00000024660 | 132,184419 | 107,440157 | 214,228254 | 84,7939333 | 83,9660249 | 45,11695343 | 81,7843074 | 101,44476 | 81,1497361 | 77,6661487 | 28,706946 | 49,04758 | 51,0895797 | 84,19306 | 82,83547 | 96,62385381 | 218,3905 | 155,5636837 | 305,1318 | 598,6573 | 139,0593 | 150,756896 |
| Upregulated | ENSMUSG00000024177 | 94,11530636 | 24,4665705 | 186,088102 | 59,7411803 | 8,90548749 | 41,13604578 | 19,3101837 | 58,666849 | 81,1497361 | 31,2821988 | 54,1015521 | 43,79248 | 35,0614763 | 43,03201 | 43,01073 | 47,40038111 | 124,9128 | 88,1857825 | 151,1401 | 309,3285 | 91,22999 | 68,1995483 |
| Upregulated | ENSMUSG00000001739 | 49,70134156 | 11,7014033 | 35,4021268 | 0,96356742 | 1,2722125 | 1,326969219 | 1,13589316 | 2,44445204 | 3,20327906 | 0 | 2,20822662 | 12,2619 | 6,01053879 | 0,935478 | 1,59299 | 1,823091581 | 46,32526 | 14,86277233 | 73,19362 | 142,6644 | 43,40067 | 10,7683497 |
| Upregulated | ENSMUSG00000039748 | 20,09203169 | 8,51011147 | 15,4316963 | 9,63567424 | 5,08884999 | 0 | 4,54357263 | 20,7778424 | 4,27103874 | 0 | 0 | 9,634347 | 1,00175646 | 8,419306 | 1,59299 | 0 | 33,91671 | 19,81702978 | 28,51699 | 92,66522 | 17,71456 | 7,17889982 |
| Upregulated | ENSMUSG00000048327 | 112,0923873 | 109,567685 | 197,888811 | 23,1256182 | 19,0831875 | 0 | 36,3485811 | 79,4446914 | 22,4229534 | 24,8100197 | 9,93701978 | 0 | 1,00175646 | 8,419306 | 19,11588 | 7,292366325 | 141,4575 | 120,8838816 | 103,6117 | 385,994 | 112,4875 | 131,912284 |
| Upregulated | ENSMUSG00000038943 | 194,5754648 | 168,074702 | 448,426939 | 109,846686 | 25,44425 | 15,92363062 | 119,268782 | 112,444794 | 70,4721393 | 74,4300591 | 57,4138921 | 13,13775 | 8,01405172 | 21,516 | 18,31938 | 0 | 269,6792 | 389,4046351 | 338,4017 | 825,3205 | 340,1196 | 312,282142 |
| Upregulated | ENSMUSG00000026622 | 104,6900599 | 64,8896 | 205,150786 | 28,9070227 | 11,4499125 | 6,634846093 | 57,9305511 | 47,6668148 | 28,8295115 | 26,9674127 | 0 | 13,13775 | 7,01229525 | 2,806435 | 14,33691 | 6,380820534 | 124,9128 | 148,6277233 | 151,1401 | 385,3273 | 131,9735 | 161,525246 |
| Upregulated | ENSMUSG00000039187 | 34,89668663 | 43,6143213 | 63,5422788 | 16,3806462 | 11,4499125 | 18,57756906 | 27,2614358 | 29,3334245 | 32,0327906 | 8,62957207 | 2,20822662 | 12,2619 | 24,0421551 | 6,548349 | 10,35443 | 0 | 65,35171 | 78,27726762 | 60,83626 | 149,9977 | 44,2864 | 51,1496612 |
| Upregulated | ENSMUSG00000036223 | 9,517278171 | 15,956459 | 52,6493167 | 19,2713485 | 6,36106249 | 5,307876874 | 9,08714527 | 25,6667465 | 12,8131162 | 1,07869651 | 2,20822662 | 0 | 2,00351293 | 3,741914 | 1,59299 | 0 | 47,1525 | 53,5059804 | 40,87436 | 123,9981 | 37,20058 | 34,0997741 |
| Upregulated | ENSMUSG00000034311 | 29,60930986 | 35,1042098 | 94,4056714 | 16,3806462 | 7,63327499 | 5,307876874 | 21,58197 | 42,7779108 | 20,287434 | 3,23608953 | 4,41645324 | 2,627549 | 5,00878232 | 0 | 0 | 0 | 119,1221 | 93,14003995 | 111,2163 | 289,9955 | 64,65815 | 80,762623 |
| Upregulated | ENSMUSG00000031262 | 35,95416198 | 85,1011147 | 147,962735 | 77,0853939 | 31,8053125 | 45,11695343 | 70,4253758 | 59,8890751 | 80,0819764 | 48,5413429 | 25,3946061 | 47,29588 | 54,0948491 | 39,29009 | 32,65629 | 5,469274744 | 211,7726 | 179,3441195 | 175,8548 | 437,3265 | 130,202 | 131,014922 |
| Upregulated | ENSMUSG00000048922 | 28,55183451 | 48,933141 | 87,1436966 | 17,3442136 | 19,0831875 | 14,5966614 | 60,2023374 | 25,6667465 | 27,7617518 | 25,8887162 | 13,2493597 | 25,39964 | 28,049181 | 37,41914 | 14,33691 | 25,52328214 | 74,45132 | 101,0668519 | 69,39135 | 197,3303 | 58,45805 | 52,0470237 |
| Upregulated | ENSMUSG00000027469 | 159,6787782 | 126,587908 | 301,371951 | 78,0489613 | 10,1777 | 41,13604578 | 105,638064 | 81,8891435 | 48,0491859 | 59,328308 | 60,726232 | 30,65474 | 16,0281034 | 33,67722 | 65,31259 | 32,81564846 | 230,7991 | 284,3743773 | 244,2956 | 731,9886 | 167,4026 | 170,498871 |
| Upregulated | ENSMUSG00000037725 | 127,9545176 | 112,758977 | 306,818432 | 70,340422 | 25,44425 | 53,07876874 | 107,90985 | 96,5558557 | 80,0819764 | 50,6987359 | 67,3509118 | 128,7499 | 74,1299784 | 55,19322 | 122,6602 | 82,03912116 | 225,0084 | 297,2554466 | 227,1854 | 675,9895 | 193,9744 | 148,962171 |
| Upregulated | ENSMUSG00000118240 | 13,74717958 | 25,5303344 | 31,7711394 | 14,4535114 | 2,544425 | 3,980907656 | 5,67946579 | 22,0000684 | 13,8808759 | 11,8656616 | 1,10411331 | 2,627549 | 0 | 0 | 2,389485 | 0,911545791 | 27,29882 | 30,71639615 | 24,71473 | 117,3315 | 20,37174 | 17,0498871 |
| Upregulated | ENSMUSG00000027326 | 27,49435916 | 62,7620721 | 60,8190383 | 19,2713485 | 5,08884999 | 0 | 12,4948247 | 33,0001026 | 16,0163953 | 18,3378407 | 6,62467985 | 16,64114 | 4,00702586 | 0 | 2,389485 | 0 | 59,56105 | 57,46938635 | 65,58909 | 243,9962 | 62,00096 | 66,4048233 |
| Upregulated | ENSMUSG00000031004 | 119,4947148 | 165,947174 | 232,383191 | 85,7575007 | 20,3554 | 37,15513812 | 73,8330553 | 147,889349 | 80,0819764 | 31,2821988 | 34,2275126 | 12,2619 | 1,00175646 | 7,483827 | 25,48784 | 11,85009528 | 236,5897 | 261,584793 | 307,033 | 955,9851 | 248,8896 | 273,695556 |
| Upregulated | ENSMUSG00000028678 | 47,58639085 | 11,7014033 | 47,2028357 | 28,9070227 | 3,8166375 | 9,28878453 | 4,54357263 | 22,0000684 | 22,4229534 | 7,55087556 | 0 | 5,255098 | 0 | 0 | 1,59299 | 0,911545791 | 76,93303 | 42,60661402 | 48,47889 | 326,6616 | 43,40067 | 40,3813115 |
| Upregulated | ENSMUSG00000045328 | 46,5289155 | 56,3794885 | 104,390887 | 15,4170788 | 8,90548749 | 9,28878453 | 18,1742905 | 30,5556505 | 5,33879843 | 34,5182883 | 1,10411331 | 0,87585 | 3,00526939 | 0 | 7,168455 | 8,203912116 | 99,26842 | 105,0302578 | 46,57776 | 463,9928 | 81,48698 | 61,0206485 |
| Upregulated | ENSMUSG00000003779 | 166,0236303 | 67,0171278 | 281,40152 | 38,542697 | 15,26655 | 39,80907656 | 69,2894827 | 72,1113353 | 56,5912633 | 34,5182883 | 37,5398525 | 16,64114 | 19,0333728 | 25,25792 | 22,30186 | 25,52328214 | 209,2909 | 239,7860603 | 145,4367 | 1074,65 | 254,2039 | 203,701282 |
| Upregulated | ENSMUSG00000015880 | 34,89668663 | 36,1679738 | 97,1289119 | 23,1256182 | 20,3554 | 14,5966614 | 13,6307179 | 52,5557189 | 18,1519147 | 8,62957207 | 1,10411331 | 7,006798 | 13,022834 | 8,419306 | 7,168455 | 0 | 138,1486 | 114,9387727 | 160,6457 | 475,3259 | 79,71552 | 88,8388853 |
| Upregulated | ENSMUSG00000026779 | 42,29901409 | 51,0606688 | 117,099342 | 26,0163204 | 17,810975 | 7,961815312 | 26,1255426 | 40,3334587 | 34,1683099 | 34,5182883 | 24,2904928 | 5,255098 | 7,01229525 | 25,25792 | 20,70887 | 13,67318686 | 110,0225 | 77,28641613 | 108,3646 | 373,9942 | 78,82979 | 99,607235 |
| Upregulated | ENSMUSG00000002055 | 52,87376762 | 30,8491541 | 78,9739751 | 21,1984833 | 24,1720375 | 25,21241515 | 23,8537563 | 34,2223286 | 12,8131162 | 17,2591441 | 0 | 0 | 1,00175646 | 0,935478 | 14,33691 | 0 | 60,38829 | 77,28641613 | 75,09475 | 287,9955 | 49,60077 | 63,7127359 |
| Upregulated | ENSMUSG00000038252 | 158,6213028 | 90,4199344 | 241,460659 | 134,899439 | 83,9660249 | 123,4081373 | 127,220034 | 133,222636 | 61,9300618 | 124,050099 | 65,1426852 | 107,7295 | 50,0878232 | 89,80593 | 94,7829 | 59,25047639 | 242,3804 | 270,5024564 | 225,2843 | 850,6534 | 182,46 | 213,57227 |
| Upregulated | ENSMUSG00000039396 | 16,91960564 | 25,5303344 | 42,6641015 | 10,5992417 | 2,544425 | 0 | 22,7178632 | 29,3334245 | 14,9486356 | 37,7543778 | 0 | 0 | 0 | 0 | 0 | 0 | 76,10579 | 93,14003995 | 39,92379 | 228,6631 | 38,0863 | 40,3813115 |
| Upregulated | ENSMUSG00000029910 | 60,27609508 | 72,3359475 | 129,807798 | 85,7575007 | 27,988675 | 37,15513812 | 69,2894827 | 108,778116 | 68,3366199 | 98,1613823 | 13,2493597 | 26,27549 | 41,072015 | 51,45131 | 36,63877 | 26,43482793 | 205,1547 | 267,529902 | 161,5963 | 610,6571 | 205,4889 | 151,654259 |
| Upregulated | ENSMUSG00000024989 | 34,89668663 | 27,6578623 | 70,8042535 | 6,74497197 | 3,8166375 | 0 | 22,7178632 | 31,7778766 | 10,6775969 | 14,0230546 | 7,72879316 | 0 | 6,01053879 | 18,70957 | 0,796495 | 0 | 67,83342 | 62,4236438 | 37,07209 | 199,9969 | 70,85824 | 55,6364736 |
| Upregulated | ENSMUSG00000031756 | 32,78173592 | 43,6143213 | 86,2359498 | 48,1783712 | 3,8166375 | 9,28878453 | 42,0280469 | 72,1113353 | 26,6939921 | 21,5739302 | 12,1452464 | 27,15134 | 14,0245905 | 13,0967 | 11,94742 | 24,61173635 | 99,26842 | 104,0394063 | 75,09475 | 361,3277 | 109,8303 | 93,3256977 |
| Upregulated | ENSMUSG00000028873 | 95,17278171 | 87,2286426 | 186,088102 | 69,3768545 | 13,9943375 | 23,88544594 | 54,5228716 | 73,3335613 | 41,6426277 | 32,3608953 | 34,2275126 | 20,14454 | 10,0175646 | 9,354784 | 15,9299 | 7,292366325 | 166,2746 | 191,2343373 | 153,0412 | 560,6579 | 170,9455 | 154,346346 |
| Upregulated | ENSMUSG00000023015 | 166,0236303 | 162,755882 | 361,283242 | 148,389383 | 27,988675 | 39,80907656 | 144,258431 | 154,000479 | 99,3016508 | 152,096208 | 13,2493597 | 28,90304 | 5,00878232 | 6,548349 | 19,91237 | 14,58473265 | 290,3601 | 481,5538236 | 327,9454 | 1311,98 | 324,1765 | 368,815978 |
| Upregulated | ENSMUSG00000024056 | 11,63222888 | 37,2317377 | 88,0514435 | 24,0891856 | 33,077525 | 7,961815312 | 15,9025042 | 40,3334587 | 17,084155 | 15,1017511 | 1,10411331 | 8,758497 | 0 | 9,354784 | 0,796495 | 1,823091581 | 73,62408 | 87,19493102 | 70,34192 | 279,9956 | 75,28688 | 54,7391111 |
| Upregulated | ENSMUSG00000025758 | 141,7016972 | 187,222452 | 245,999394 | 185,004945 | 95,4159374 | 122,0811681 | 101,094491 | 172,333869 | 129,198922 | 165,040566 | 96,0578579 | 78,82647 | 101,177403 | 105,7091 | 62,9231 | 82,95066695 | 323,4496 | 288,3377832 | 229,0865 | 709,3223 | 269,2613 | 231,519519 |
| Upregulated | ENSMUSG00000023940 | 19,03455634 | 18,0839869 | 47,2028357 | 13,4899439 | 5,08884999 | 33,17423047 | 21,58197 | 26,8889725 | 13,8808759 | 20,4952337 | 30,9151726 | 25,39964 | 3,00526939 | 13,0967 | 12,74392 | 3,646183163 | 72,79684 | 57,46938635 | 45,62719 | 145,9977 | 38,0863 | 53,8417486 |
| Upregulated | ENSMUSG00000021485 | 11,63222888 | 13,8289311 | 97,1289119 | 24,0891856 | 1,2722125 | 6,634846093 | 36,3485811 | 34,2223286 | 23,4907131 | 18,3378407 | 3,31233993 | 2,627549 | 6,01053879 | 10,29026 | 9,557939 | 8,203912116 | 92,65053 | 44,588317 | 77,94645 | 215,33 | 53,14368 | 46,6628488 |
| Upregulated | ENSMUSG00000034329 | 5,287376762 | 31,912918 | 85,328203 | 6,74497197 | 3,8166375 | 0 | 28,397329 | 23,2222944 | 25,6262325 | 14,0230546 | 12,1452464 | 7,882647 | 11,0193211 | 21,516 | 13,54041 | 13,67318686 | 128,2217 | 75,30471315 | 57,98456 | 257,3293 | 46,94358 | 38,5865865 |

| | | | | | | | | | | | | | | | | | | | | | | | |
|---|---|---|---|---|---|---|---|---|---|---|---|---|---|---|---|---|---|---|---|---|---|---|---|
| Upregulated | ENSMUSG00000035842 | 38,06911268 | 44,6780852 | 59,9112914 | 34,6884273 | 13,9943375 | 23,88544594 | 24,9896495 | 37,8890067 | 17,084155 | 29,1248057 | 26,4987194 | 61,30948 | 32,0562069 | 28,06435 | 46,9932 | 20,96555318 | 96,78671 | 46,57001997 | 56,08342 | 120,6648 | 44,2864 | 45,7654863 |
| Upregulated | ENSMUSG00000055612 | 37,01163733 | 29,7853902 | 52,6493167 | 17,3442136 | 0 | 18,57756906 | 43,16394 | 22,0000684 | 17,084155 | 20,4952337 | 22,0822662 | 43,79248 | 38,0667457 | 23,38696 | 54,95815 | 20,96555318 | 97,61395 | 48,55172295 | 47,52832 | 161,9975 | 51,37222 | 47,5602113 |
| Upregulated | ENSMUSG00000026274 | 37,01163733 | 32,976682 | 49,0183294 | 24,0891856 | 10,1777 | 14,5966614 | 20,4460768 | 22,0000684 | 38,4393487 | 64,7217906 | 22,0822662 | 38,53739 | 13,022834 | 17,77409 | 10,35443 | 20,05400739 | 150,5571 | 77,28641613 | 72,24305 | 229,9964 | 44,2864 | 31,4076867 |
| Upregulated | ENSMUSG00000029409 | 17,97708099 | 4,25505574 | 8,16972156 | 0,96356742 | 6,36106249 | 9,28878453 | 11,3589316 | 0 | 1,06775969 | 5,39348255 | 0 | 20,14454 | 27,0474245 | 28,99983 | 4,77897 | 0 | 186,1283 | 27,74384169 | 101,7106 | 355,3278 | 22,1432 | 50,2522987 |
| Upregulated | ENSMUSG00000052430 | 2,114950705 | 3,1912918 | 17,24719 | 14,4535114 | 0 | 0 | 9,08714527 | 13,4444862 | 4,27103874 | 0 | 7,72879316 | 47,29588 | 36,0632327 | 22,45148 | 23,09835 | 20,05400739 | 244,8621 | 72,33215868 | 86,50155 | 309,9952 | 53,14368 | 32,3050492 |
| Upregulated | ENSMUSG00000049001 | 42,29901409 | 99,9938098 | 95,3134182 | 178,259973 | 99,2325749 | 46,44392265 | 52,2510853 | 61,1113011 | 67,2688602 | 182,29971 | 26,4987194 | 129,6258 | 214,375883 | 99,16071 | 187,1763 | 179,5745208 | 775,9482 | 343,8254666 | 423,9527 | 1257,98 | 140,8308 | 133,707009 |
| Upregulated | ENSMUSG00000037544 | 69,79337325 | 63,825836 | 149,778229 | 49,1419386 | 44,5274375 | 62,36755328 | 49,979299 | 23,2222944 | 29,8972712 | 24,8100197 | 32,019286 | 162,908 | 62,1089008 | 17,77409 | 67,70207 | 47,40038111 | 363,9842 | 111,9662182 | 237,6416 | 411,9936 | 94,7729 | 174,088321 |
| Upregulated | ENSMUSG00000037474 | 44,4139648 | 57,4432524 | 123,45357 | 55,8869106 | 17,810975 | 33,17423047 | 64,74591 | 105,111438 | 70,4721393 | 39,9117708 | 14,353473 | 65,68873 | 22,0386422 | 41,16105 | 63,7196 | 18,23091581 | 304,4232 | 128,8106935 | 262,3564 | 389,9939 | 42,51494 | 79,8652605 |
| Upregulated | ENSMUSG00000028587 | 5,287376762 | 9,57387541 | 41,7563546 | 16,3806462 | 12,722125 | 1,326969219 | 14,7666111 | 18,3333903 | 29,8972712 | 25,8887162 | 18,7699262 | 16,64114 | 22,0386422 | 19,64505 | 21,50536 | 18,23091581 | 100,0957 | 33,68895062 | 75,09475 | 146,6644 | 21,25747 | 35,8944991 |
| Upregulated | ENSMUSG00000026429 | 28,55183451 | 28,7216262 | 41,7563546 | 37,5791295 | 25,44425 | 1,326969219 | 27,2614358 | 3,66667807 | 18,1519147 | 5,39348255 | 9,93701978 | 28,02719 | 8,01405172 | 10,29026 | 25,48784 | 14,58473265 | 49,63421 | 49,54257444 | 71,29249 | 98,66513 | 21,25747 | 25,1261494 |
| Upregulated | ENSMUSG00000033952 | 20,09203169 | 36,1679738 | 46,2950888 | 11,5628091 | 1,2722125 | 26,53938437 | 35,2126879 | 31,7778766 | 20,287434 | 0 | 2,20822662 | 1,751699 | 1,00175646 | 0,935478 | 14,33691 | 5,469274744 | 52,94316 | 67,37790124 | 107,414 | 158,6642 | 42,51494 | 53,8417486 |
| Upregulated | ENSMUSG00000027323 | 44,4139648 | 96,802518 | 71,7120004 | 63,59545 | 17,810975 | 30,52029203 | 32,9409016 | 34,2223286 | 50,1847052 | 24,8100197 | 14,353473 | 13,13775 | 25,0439116 | 32,74174 | 11,94742 | 7,292366325 | 119,9493 | 123,8564361 | 187,2616 | 265,9958 | 91,22999 | 46,6628488 |
| Upregulated | ENSMUSG00000017146 | 22,2069824 | 40,4230295 | 35,4021268 | 33,7248598 | 2,544425 | 19,90453828 | 10,2230384 | 22,0000684 | 24,5584728 | 0 | 3,31233993 | 0 | 3,00526939 | 11,22574 | 0,796495 | 0 | 62,04276 | 59,45108933 | 76,99589 | 133,9979 | 28,3433 | 43,0733989 |
| Upregulated | ENSMUSG00000040658 | 30,66678522 | 41,4867934 | 61,7267851 | 40,4698318 | 19,0831875 | 31,84726125 | 20,4460768 | 53,777945 | 32,0327906 | 29,1248057 | 16,5616996 | 16,64114 | 38,0667457 | 43,03201 | 35,04578 | 41,93110637 | 86,85987 | 76,29556464 | 128,3265 | 193,997 | 19,48602 | 33,2024117 |
| Upregulated | ENSMUSG00000041406 | 59,21861973 | 32,976682 | 25,4169115 | 14,4535114 | 15,26655 | 21,2315075 | 18,1742905 | 41,5556847 | 26,6939921 | 10,7869651 | 5,52056654 | 13,13775 | 18,0316164 | 6,548349 | 24,69134 | 48,3119269 | 81,89645 | 84,22237655 | 60,83626 | 124,6647 | 40,74349 | 35,8944991 |
| Upregulated | ENSMUSG00000049097 | 0 | 7,44634754 | 9,0774684 | 4,81783712 | 3,8166375 | 0 | 3,40767947 | 4,88890409 | 16,0163953 | 1,07869651 | 8,83290647 | 5,255098 | 10,0175646 | 1,870957 | 9,557939 | 9,115457906 | 23,16263 | 23,78043573 | 18,06076 | 40,66603 | 8,85728 | 22,4340619 |
| Upregulated | ENSMUSG00000046610 | 1,057475352 | 2,12752787 | 5,44648104 | 7,70853939 | 2,544425 | 17,25059984 | 3,40767947 | 0 | 3,20327906 | 3,23608953 | 8,83290647 | 7,006798 | 11,0193211 | 16,83861 | 6,37196 | 0,911545791 | 56,25211 | 46,57001997 | 75,09475 | 95,9985 | 15,9431 | 56,5338361 |
| Upregulated | ENSMUSG00000041620 | 0 | 0 | 0 | 0 | 0 | 0 | 0 | 0 | 0 | 0 | 0 | 2,627549 | 0 | 0 | 0,796495 | 0 | 7,445132 | 4,954257444 | 19,9619 | 25,33294 | 1,771456 | 17,0498871 |
| Upregulated | ENSMUSG00000048644 | 34,89668663 | 10,6376393 | 24,5091647 | 10,5992417 | 13,9943375 | 1,326969219 | 18,1742905 | 9,77780817 | 8,54207749 | 6,47217906 | 9,93701978 | 11,38605 | 20,0351293 | 36,48366 | 23,09835 | 15,49627844 | 71,96961 | 29,72554466 | 169,2008 | 161,3308 | 44,2864 | 81,6599854 |
| Upregulated | ENSMUSG00000022489 | 25,37940846 | 11,7014033 | 18,1549368 | 9,63567424 | 20,3554 | 21,2315075 | 30,6691153 | 23,2222944 | 14,9486356 | 8,62957207 | 35,3316259 | 21,89624 | 16,0281034 | 37,41914 | 11,94742 | 13,67318686 | 43,01632 | 18,82617829 | 82,69928 | 114,6649 | 27,45757 | 79,8652605 |
| Upregulated | ENSMUSG00000028128 | 1483,637919 | 1944,56047 | 2450,00872 | 2533,21876 | 270,981262 | 262,7399053 | 199,917196 | 1807,67229 | 842,462392 | 728,120144 | 983,764958 | 1375,084 | 2168,80275 | 998,1554 | 366,3877 | 494,9693643 | 4408,345 | 2025,300443 | 5802,258 | 9706,515 | 6498,586 | 3822,76415 |
| Upregulated | ENSMUSG00000044522 | 6,344852114 | 10,6376393 | 11,8007089 | 12,5263765 | 0 | 0 | 5,67946579 | 12,2222602 | 0 | 11,8656616 | 7,72879316 | 14,0136 | 25,0439116 | 3,741914 | 0 | 3,646183163 | 75,27855 | 18,82617829 | 111,2163 | 151,331 | 130,202 | 36,7918616 |
| Upregulated | ENSMUSG00000030017 | 0 | 0 | 0 | 0,96356742 | 0 | 0 | 1,13589316 | 0 | 3,20327906 | 0 | 0 | 0,87585 | 45,0790409 | 20,58052 | 13,54041 | 0 | 168,7563 | 154,5728323 | 543,724 | 738,6551 | 347,2054 | 66,4048233 |
| Upregulated | ENSMUSG00000024598 | 200,9203169 | 101,057574 | 275,047293 | 348,811407 | 63,6106249 | 212,315075 | 89,7355595 | 287,223115 | 108,911488 | 239,470625 | 133,59771 | 252,2447 | 403,707855 | 329,2884 | 75,66702 | 124,8817733 | 555,0759 | 627,2089924 | 1373,569 | 2095,967 | 1291,391 | 561,748911 |
| Upregulated | ENSMUSG00000106820 | 3,172426057 | 2,12752787 | 0 | 8,67210682 | 3,8166375 | 3,980907656 | 2,27178632 | 6,11113011 | 8,54207749 | 1,07869651 | 3,31233993 | 0,87585 | 0 | 12,16122 | 7,168455 | 3,646183163 | 21,50816 | 13,87192084 | 30,41813 | 50,66588 | 35,42912 | 7,17889982 |
| Upregulated | ENSMUSG00000040907 | 0 | 1,06376393 | 0,90774684 | 6,74497197 | 0 | 0 | 0 | 2,44445204 | 3,20327906 | 2,15739302 | 0 | 3,503399 | 21,0368858 | 2,806435 | 3,982475 | 0,911545791 | 43,01632 | 0,990851489 | 93,15552 | 212,6633 | 143,4879 | 9,87098725 |
| Upregulated | ENSMUSG00000026069 | 2588,699662 | 2375,38486 | 2237,59596 | 455,767392 | 346,0418 | 242,835367 | 229,450418 | 513,334929 | 139,876519 | 335,474614 | 665,780325 | 10827,25 | 2380,17336 | 3107,659 | 8922,336 | 3456,581638 | 21801 | 7220,334799 | 12543,68 | 37284,08 | 14089,28 | 12039,015 |
| Upregulated | ENSMUSG00000032363 | 422,9901409 | 227,645482 | 437,533977 | 203,312726 | 209,915062 | 175,1599369 | 240,80935 | 178,444999 | 168,70603 | 302,035023 | 241,800815 | 495,7309 | 110,193211 | 144,0637 | 388,6895 | 340,0065799 | 1228,447 | 258,6122386 | 1244,292 | 1800,639 | 793,6123 | 803,139417 |
| Upregulated | ENSMUSG00000059810 | 271,7711655 | 169,138465 | 294,109976 | 279,434553 | 235,359312 | 221,6038595 | 180,607012 | 190,667259 | 163,367232 | 247,021501 | 325,713426 | 315,3059 | 346,607737 | 275,0306 | 165,6709 | 146,7588723 | 627,8728 | 336,8895062 | 683,4573 | 1266,647 | 502,2078 | 412,78674 |
| Upregulated | ENSMUSG00000032387 | 1,057475352 | 6,3825836 | 14,5239494 | 13,4899439 | 6,36106249 | 10,61575375 | 14,7666111 | 9,77780817 | 16,0163953 | 17,2591441 | 9,93701978 | 10,5102 | 10,0175646 | 7,483827 | 10,35443 | 10,0270037 | 39,70737 | 13,87192084 | 42,77549 | 69,99891 | 22,1432 | 34,0997741 |
| Upregulated | ENSMUSG00000037010 | 2,114950705 | 11,7014033 | 15,4316963 | 33,7248598 | 30,5331 | 35,8281689 | 30,6691153 | 12,2222602 | 10,6775969 | 136,994457 | 16,5616996 | 52,55098 | 79,1387607 | 56,1287 | 39,02825 | 30,08101109 | 81,06921 | 63,41449528 | 186,311 | 360,661 | 146,1451 | 36,7918616 |
| Upregulated | ENSMUSG00000035455 | 3,172426057 | 4,25505574 | 29,9556457 | 29,8705901 | 1,2722125 | 1,326969219 | 10,2230384 | 8,55558215 | 10,6775969 | 40,9904673 | 0 | 2,627549 | 13,022834 | 0 | 7,168455 | 0 | 27,29882 | 23,78043573 | 39,92379 | 77,99878 | 38,97203 | 9,87098725 |
| Upregulated | ENSMUSG00000030774 | 156,5063521 | 164,88341 | 148,870482 | 161,879327 | 91,5992999 | 127,389045 | 57,9305511 | 57,444623 | 82,2174958 | 94,9252928 | 119,244237 | 160,2805 | 122,214289 | 251,6437 | 125,8462 | 178,662975 | 249,8255 | 229,8775454 | 318,4398 | 630,6568 | 248,0038 | 140,885909 |
| Upregulated | ENSMUSG00000019808 | 10,57475352 | 10,6376393 | 11,8007089 | 17,3442136 | 15,26655 | 13,26969219 | 6,81535895 | 17,1111643 | 32,0327906 | 18,3378407 | 9,93701978 | 7,006798 | 17,0298599 | 34,6127 | 21,50536 | 24,61173635 | 35,57118 | 33,68895062 | 36,12153 | 103,9984 | 29,22902 | 23,3314244 |
| Upregulated | ENSMUSG00000070469 | 167,0811057 | 192,541272 | 228,752204 | 209,094131 | 109,410275 | 114,1193528 | 103,366277 | 155,222705 | 162,299472 | 217,896695 | 205,365075 | 175,1699 | 113,19848 | 138,4508 | 149,7411 | 251,5866382 | 361,5025 | 342,8346151 | 236,6911 | 625,3236 | 233,8322 | 139,091184 |
| Upregulated | ENSMUSG00000028073 | 195,6329402 | 157,437062 | 316,803647 | 210,057698 | 99,2325749 | 98,19572218 | 205,596662 | 140,555993 | 107,843728 | 244,864108 | 235,176135 | 245,2379 | 371,651648 | 168,3861 | 98,76537 | 233,3557224 | 598,9195 | 283,3835258 | 401,1391 | 1290,647 | 411,8635 | 274,592918 |
| Upregulated | ENSMUSG00000026475 | 267,5412641 | 192,541272 | 472,936104 | 313,159413 | 185,743025 | 226,9117364 | 223,770952 | 206,556198 | 225,297294 | 304,192416 | 333,442219 | 226,8451 | 386,677995 | 158,0958 | 227,7976 | 253,4097298 | 745,3404 | 455,7916849 | 392,584 | 1590,642 | 476,5217 | 321,255767 |
| Upregulated | ENSMUSG00000034773 | 3,172426057 | 13,8289311 | 48,1105825 | 3,8542697 | 2,544425 | 14,5966614 | 32,9409016 | 7,33335613 | 6,40655811 | 0 | 3,31233993 | 26,27549 | 3,00526939 | 7,483827 | 9,557939 | 20,05400739 | 97,61395 | 75,30471315 | 71,29249 | 207,9968 | 67,31533 | 35,8944991 |
| Upregulated | ENSMUSG00000102543 | 20,09203169 | 23,4028065 | 83,5127093 | 20,2349159 | 25,44425 | 33,17423047 | 34,0767947 | 28,1111985 | 8,54207749 | 34,5182883 | 23,1863795 | 53,42683 | 22,0386422 | 35,54818 | 50,97568 | 44,66574374 | 132,3579 | 78,27726762 | 79,84759 | 313,3284 | 65,54387 | 49,3549363 |
| Upregulated | ENSMUSG00000021569 | 51,81629226 | 47,869377 | 86,2359498 | 59,7411803 | 22,899825 | 33,17423047 | 38,6203674 | 56,222397 | 46,9814262 | 14,0230546 | 18,7699262 | 32,40644 | 43,075528 | 74,83827 | 73,27754 | 72,92366325 | 193,5734 | 142,6826144 | 140,6838 | 514,6586 | 157,6596 | 101,40196 |
| Upregulated | ENSMUSG00000079685 | 99,40268312 | 42,5505574 | 150,685975 | 104,065282 | 41,9830125 | 49,09786109 | 99,9585979 | 133,222636 | 44,8459068 | 52,8561289 | 58,5180054 | 106,8537 | 40,0702586 | 54,25775 | 135,4041 | 64,71975114 | 329,2403 | 241,7677633 | 298,4779 | 1053,984 | 267,4899 | 219,853807 |
| Upregulated | ENSMUSG00000040936 | 29,60930986 | 35,1042098 | 44,4795952 | 14,4535114 | 10,1777 | 39,80907656 | 26,1255426 | 15,8889383 | 13,8808759 | 11,8656616 | 14,353473 | 13,13775 | 38,0667457 | 25,25792 | 12,74392 | 11,85009528 | 44,67079 | 49,54257444 | 38,02266 | 115,3315 | 61,11523 | 38,5865865 |
| Upregulated | ENSMUSG00000035351 | 132,184419 | 99,9938098 | 161,578938 | 115,628091 | 63,6106249 | 124,7351066 | 62,4741237 | 57,444623 | 52,3202246 | 88,4531138 | 75,079705 | 42,91663 | 116,20375 | 127,2251 | 133,0147 | 115,7663154 | 138,1486 | 117,9113272 | 160,6457 | 330,6615 | 163,8597 | 96,9151476 |
| Upregulated | ENSMUSG00000072621 | 102,5751092 | 18,0839869 | 55,3725572 | 5,78140454 | 5,08884999 | 10,61575375 | 1,13589316 | 6,11113011 | 3,20327906 | 5,39348255 | 17,6658129 | 48,17173 | 84,147543 | 19,64505 | 77,26001 | 34,63874004 | 110,0225 | 61,43279231 | 128,3265 | 238,6629 | 108,0588 | 41,278674 |
| Upregulated | ENSMUSG00000033276 | 93,057831 | 23,4028065 | 56,2803041 | 30,8341576 | 66,1550499 | 26,53938437 | 30,6691153 | 19,5556163 | 6,40655811 | 36,6756813 | 15,4575863 | 31,53059 | 31,0544504 | 17,77409 | 43,80722 | 15,49627844 | 50,46145 | 53,5059804 | 70,34192 | 157,9975 | 96,54435 | 40,3813115 |
| Upregulated | ENSMUSG00000023345 | 57,10366903 | 48,933141 | 69,8965067 | 28,9070227 | 27,988675 | 3,980907656 | 11,3589316 | 35,4445546 | 30,9650309 | 31,2821988 | 30,9151726 | 18,39284 | 37,0649892 | 11,22574 | 49,38269 | 34,63874004 | 57,90658 | 55,48768337 | 83,64985 | 155,3309 | 85,91562 | 53,8417486 |
| Upregulated | ENSMUSG00000052572 | 54,98871832 | 56,3794885 | 68,081013 | 46,2512364 | 25,44425 | 6,634846093 | 4,54357263 | 20,7778424 | 18,1519147 | 57,170915 | 15,4575863 | 21,89624 | 69,1211961 | 5,61287 | 34,24928 | 8,203912116 | 44,67079 | 17,8353268 | 102,6612 | 259,3293 | 131,0877 | 46,6628488 |
| Upregulated | ENSMUSG00000026605 | 47,58639085 | 44,6780852 | 81,6972156 | 5,78140454 | 2,544425 | 7,961815312 | 18,1742905 | 4,88890409 | 17,084155 | 26,9674127 | 2,20822662 | 10,5102 | 1,00175646 | 0 | 7,964949 | 0,911545791 | 13,23579 | 90,16748548 | 44,67663 | 175,3306 | 70,85824 | 87,9415228 |
| Upregulated | ENSMUSG00000037628 | 25,37940846 | 31,912918 | 101,667646 | 26,9798879 | 11,4499125 | 41,13604578 | 32,9409016 | 26,8889725 | 17,084155 | 16,1804476 | 4,41645324 | 2,627549 | 6,01053879 | 7,483827 | 5,575465 | 10,0270037 | 53,77039 | 133,764951 | 56,08342 | 292,6621 | 136,4021 | 96,0177851 |
| Upregulated | ENSMUSG00000026683 | 41,24153874 | 35,1042098 | 170,656406 | 41,4333992 | 3,8166375 | 26,53938437 | 51,1151921 | 44,0001368 | 19,2196743 | 18,3378407 | 2,20822662 | 1,751699 | 0 | 10,29026 | 3,18598 | 0 | 110,8497 | 174,389862 | 87,45212 | 379,3274 | 151,4595 | 113,067672 |
| Upregulated | ENSMUSG00000001517 | 93,057831 | 52,1244328 | 166,117672 | 41,4333992 | 29,2608875 | 58,38664562 | 24,9896495 | 63,5557531 | 33,1005503 | 55,013522 | 16,5616996 | 27,15134 | 13,022834 | 25,25792 | 14,33691 | 13,67318686 | 79,41474 | 227,8958424 | 132,1287 | 487,9924 | 172,717 | 160,627883 |
| Upregulated | ENSMUSG00000005233 | 27,49435916 | 85,1011147 | 141,608507 | 59,7411803 | 2,544425 | 11,94272297 | 62,4741237 | 45,2223628 | 48,0491859 | 5,39348255 | 2,20822662 | 7,882647 | 15,026347 | 6,548349 | 10,35443 | 19,1424616 | 83,55092 | 210,0605156 | 141,6344 | 435,3265 | 116,0304 | 146,270084 |
| Upregulated | ENSMUSG00000098090 | 11,63222888 | 8,51011147 | 37,2176204 | 14,4535114 | 6,36106249 | 9,28878453 | 6,81535895 | 13,4444862 | 5,33879843 | 5,39348255 | 1,10411331 | 15,76529 | 8,01405172 | 7,483827 | 7,168455 | 0,911545791 | 17,37197 | 30,71639615 | 25,6653 | 69,33225 | 22,1432 | 26,9208743 |
| Upregulated | ENSMUSG00000079553 | 29,60930986 | 2,12752787 | 28,140152 | 15,4170788 | 22,899825 | 2,653938437 | 6,81535895 | 9,77780817 | 12,8131162 | 4,31478604 | 7,72879316 | 0 | 0 | 0 | 0 | 9,115457906 | 18,19921 | 34,67980211 | 24,71473 | 136,6645 | 21,25747 | 52,9443862 |
| Upregulated | ENSMUSG00000017737 | 31,72426057 | 6,3825836 | 28,140152 | 15,4170788 | 6,36106249 | 7,961815312 | 15,9025042 | 6,11113011 | 1,06775969 | 1,07869651 | 8,83290647 | 2,627549 | 1,00175646 | 0 | 1,59299 | 0 | 0 | 139,7100599 | 29,46756 | 411,9936 | 33,65766 | 82,5573479 |
| Upregulated | ENSMUSG00000025432 | 44,4139648 | 5,31881967 | 37,2176204 | 26,0163204 | 3,8166375 | 22,55847672 | 6,81535895 | 30,5556505 | 17,084155 | 8,62957207 | 2,20822662 | 20,14454 | 29,0509375 | 26,19339 | 3,982475 | 26,43482793 | 29,78053 | 136,7375055 | 57,03399 | 357,3278 | 110,716 | 69,9942732 |
| Upregulated | ENSMUSG00000109086 | 6,344852114 | 14,8926951 | 47,2028357 | 3,8542697 | 1,2722125 | 0 | 6,81535895 | 25,6667465 | 4,27103874 | 0 | 9,93701978 | 0 | 0 | 0 | 0 | 0 | 3,308947 | 34,67980211 | 35,17096 | 152,6643 | 67,31533 | 59,2259235 |
| Upregulated | ENSMUSG00000023505 | 53,93124297 | 65,9533639 | 186,088102 | 19,2713485 | 10,1777 | 7,961815312 | 54,5228716 | 51,3334929 | 38,4393487 | 18,3378407 | 19,8740396 | 4,379248 | 0 | 3,741914 | 4,77897 | 0,911545791 | 100,0957 | 179,3441195 | 229,0865 | 792,6543 | 377,3201 | 192,932933 |
| Upregulated | ENSMUSG00000029177 | 67,67842255 | 69,1446557 | 205,150786 | 20,2349159 | 49,6162874 | 30,52029203 | 89,7355595 | 80,6669174 | 42,7103874 | 45,3052534 | 23,1863795 | 2,627549 | 6,01053879 | 15,90313 | 18,31938 | 15,49627844 | 65,35171 | 228,8866939 | 106,4634 | 859,9866 | 426,9209 | 198,317108 |
| Upregulated | ENSMUSG00000017716 | 63,44852114 | 89,3561705 | 213,320507 | 76,1218265 | 5,08884999 | 30,52029203 | 46,5716195 | 94,1114037 | 42,7103874 | 30,2035023 | 7,72879316 | 11,38605 | 11,0193211 | 6,548349 | 7,168455 | 7,292366325 | 182,8193 | 303,2005556 | 202,4707 | 1045,984 | 415,4064 | 164,217333 |
| Upregulated | ENSMUSG00000032218 | 114,2073381 | 137,225547 | 326,788862 | 64,5590174 | 29,2608875 | 80,94512234 | 95,4150253 | 128,333732 | 100,36941 | 55,013522 | 32,019286 | 41,16494 | 14,0245905 | 18,70957 | 19,11588 | 13,67318686 | 158,0022 | 470,6544572 | 345,0556 | 2007,302 | 668,7247 | 346,381916 |
| Upregulated | ENSMUSG00000084133 | 14,80465493 | 9,57387541 | 33,5866331 | 7,70853939 | 1,2722125 | 1,326969219 | 13,6307179 | 13,4444862 | 7,4743178 | 4,31478604 | 1,10411331 | 0 | 0 | 0 | 0 | 0 | 13,23579 | 34,67980211 | 33,26983 | 171,9973 | 55,80086 | 29,6129618 |
| Upregulated | ENSMUSG00000028175 | 9,517278171 | 18,0839869 | 50,833823 | 2,89070227 | 1,2722125 | 0 | 4,54357263 | 9,77780817 | 8,54207749 | 0 | 2,20822662 | 0 | 0 | 0 | 0 | 0 | 22,33539 | 19,81702978 | 19,9619 | 119,9981 | 33,65766 | 38,5865865 |
| Upregulated | ENSMUSG00000030867 | 48,64386621 | 53,1881967 | 199,704305 | 20,2349159 | 19,0831875 | 14,5966614 | 15,9025042 | 78,2224654 | 32,0327906 | 25,8887162 | 2,20822662 | 14,0136 | 11,0193211 | 15,90313 | 19,91237 | 10,93854949 | 79,41474 | 120,8838816 | 96,95778 | 681,3227 | 171,8312 | 89,7362477 |
| Upregulated | ENSMUSG00000044201 | 8,459802819 | 6,3825836 | 65,3577725 | 11,5628091 | 0 | 10,61575375 | 27,2614358 | 14,6667123 | 8,54207749 | 0 | 5,52056654 | 0 | 0 | 0 | 2,389485 | 0 | 21,50816 | 35,6706536 | 35,17096 | 211,9967 | 49,60077 | 38,5865865 |
| Upregulated | ENSMUSG00000006398 | 113,1498627 | 117,014033 | 444,795952 | 82,8667985 | 25,44425 | 67,67543015 | 152,209683 | 118,555924 | 107,843728 | 61,485701 | 40,8521924 | 28,02719 | 27,0474245 | 3,741914 | 52,56867 | 22,78864477 | 145,5937 | 233,8409514 | 200,5695 | 1119,316 | 359,6056 | 189,343483 |
| Upregulated | ENSMUSG00000026955 | 15,86213028 | 13,8289311 | 28,140152 | 1,92713485 | 0 | 3,980907656 | 1,13589316 | 4,88890409 | 1,06775969 | 1,07869651 | 0 | 0 | 0 | 0 | 0 | 0,911545791 | 4,963421 | 16,84447531 | 30,41813 | 133,3313 | 35,42912 | 13,4604372 |
| Upregulated | ENSMUSG00000114533 | 7,402327466 | 14,8926951 | 20,8781773 | 2,89070227 | 0 | 0 | 5,67946579 | 2,44445204 | 4,27103874 | 5,39348255 | 1,10411331 | 0 | 0 | 0 | 2,389485 | 0 | 11,58132 | 18,82617829 | 29,46756 | 105,3317 | 31,00048 | 19,7419745 |
| Upregulated | ENSMUSG00000027331 | 169,1960564 | 84,0373508 | 127,992304 | 30,8341576 | 19,0831875 | 31,84726125 | 61,3382305 | 55,000171 | 36,3038293 | 6,47217906 | 11,0411331 | 4,379248 | 23,0403987 | 40,22557 | 21,50536 | 25,52328214 | 106,7136 | 166,4630501 | 158,7446 | 649,9899 | 197,5173 | 166,012058 |
| Upregulated | ENSMUSG00000036777 | 224,1847747 | 206,370203 | 363,098736 | 72,2675568 | 11,4499125 | 41,13604578 | 176,06344 | 161,333835 | 95,030612 | 74,4300591 | 39,7480791 | 30,65474 | 11,0193211 | 0,935478 | 15,1334 | 8,203912116 | 209,2909 | 419,1301798 | 333,6488 | 1753,973 | 459,6928 | 314,97423 |
| Upregulated | ENSMUSG00000048574 | 34,89668663 | 38,2955016 | 123,45357 | 14,4535114 | 5,08884999 | 2,653938437 | 26,1255426 | 37,8890067 | 17,084155 | 2,15739302 | 1,10411331 | 0,87585 | 0 | 0 | 4,77897 | 0 | 56,25211 | 102,0577033 | 83,64985 | 427,3267 | 118,6876 | 92,4283352 |
| Upregulated | ENSMUSG00000040084 | 60,27609508 | 45,7418492 | 185,180355 | 58,7776129 | 15,26655 | 17,25059984 | 54,5228716 | 69,6668832 | 8,54207749 | 8,62957207 | 27,6028327 | 29,77889 | 3,00526939 | 0,935478 | 2,389485 | 0 | 90,99605 | 160,5179412 | 98,85892 | 676,6561 | 167,4026 | 123,836022 |
| Upregulated | ENSMUSG00000027306 | 70,85084861 | 76,5910032 | 120,73033 | 40,4698318 | 5,08884999 | 19,90453828 | 54,5228716 | 67,2224312 | 36,3038293 | 5,39348255 | 16,5616996 | 7,882647 | 2,00351293 | 20,58052 | 7,168455 | 30,08101109 | 54,59763 | 108,9936638 | 71,29249 | 402,6604 | 111,6017 | 87,0441603 |
| Upregulated | ENSMUSG00000020330 | 68,7358979 | 84,0373508 | 177,010634 | 49,1419386 | 0 | 15,92363062 | 57,9305511 | 37,8890067 | 26,6939921 | 4,31478604 | 8,83290647 | 2,627549 | 5,00878232 | 0 | 11,15093 | 1,823091581 | 105,8863 | 210,0605156 | 105,5129 | 619,3237 | 201,0603 | 113,965035 |
| Upregulated | ENSMUSG00000001403 | 143,8166479 | 85,1011147 | 274,139546 | 67,4497197 | 11,4499125 | 30,52029203 | 77,2407348 | 85,5558215 | 39,5071084 | 36,6756813 | 5,52056654 | 16,64114 | 3,00526939 | 0,935478 | 4,77897 | 6,380820534 | 76,10579 | 310,136516 | 164,448 | 864,6532 | 317,0906 | 183,061945 |
| Upregulated | ENSMUSG00000041431 | 109,9774366 | 63,825836 | 253,261368 | 32,7612924 | 6,36106249 | 9,28878453 | 39,7562605 | 55,000171 | 22,4229534 | 22,6526267 | 13,2493597 | 11,38605 | 3,00526939 | 8,419306 | 1,59299 | 0,911545791 | 109,1953 | 329,9535458 | 149,2389 | 896,6527 | 196,6316 | 154,346346 |
| Upregulated | ENSMUSG00000020808 | 47,58639085 | 36,1679738 | 104,390887 | 40,4698318 | 2,544425 | 27,86635359 | 23,8537563 | 19,5556163 | 17,084155 | 7,55087556 | 3,31233993 | 7,006798 | 0 | 0 | 0 | 0 | 54,59763 | 91,15833697 | 75,09475 | 310,6618 | 94,7729 | 69,0969108 |
| Upregulated | ENSMUSG00000024795 | 29,60930986 | 32,976682 | 85,328203 | 16,3806462 | 6,36106249 | 9,28878453 | 29,5332221 | 17,1111643 | 32,0327906 | 8,62957207 | 0 | 0,87585 | 5,00878232 | 16,83861 | 19,91237 | 8,203912116 | 19,85368 | 66,38704975 | 70,34192 | 181,9972 | 40,74349 | 35,8944991 |
| Upregulated | ENSMUSG00000047534 | 7,402327466 | 28,7216262 | 64,4500256 | 18,3077811 | 5,08884999 | 15,92363062 | 20,4460768 | 26,8889725 | 11,7453565 | 5,39348255 | 5,52056654 | 7,006798 | 9,01580818 | 14,03218 | 19,91237 | 0 | 14,89026 | 60,44194082 | 36,12153 | 143,9978 | 23,02893 | 32,3050492 |
| Upregulated | ENSMUSG00000012443 | 89,88540495 | 115,950269 | 156,132456 | 65,5225848 | 19,0831875 | 10,61575375 | 136,307179 | 116,111472 | 32,0327906 | 34,5182883 | 14,353473 | 0 | 6,01053879 | 2,806435 | 8,761444 | 6,380820534 | 119,1221 | 149,6185748 | 133,0793 | 553,3247 | 112,4875 | 133,707009 |
| Upregulated | ENSMUSG00000032254 | 57,10366903 | 49,9969049 | 92,5901777 | 67,4497197 | 3,8166375 | 15,92363062 | 97,6868116 | 88,0002736 | 51,2524649 | 19,4165372 | 8,83290647 | 23,64794 | 6,01053879 | 20,58052 | 36,63877 | 20,96555318 | 41,36184 | 152,5911293 | 117,8702 | 453,3263 | 114,2589 | 121,143934 |
| Upregulated | ENSMUSG00000022021 | 29,60930986 | 32,976682 | 87,1436966 | 35,6519947 | 6,36106249 | 15,92363062 | 77,2407348 | 23,2222944 | 11,7453565 | 18,3378407 | 7,72879316 | 4,379248 | 2,00351293 | 4,677392 | 10,35443 | 8,203912116 | 90,16882 | 144,6643174 | 77,94645 | 417,9935 | 126,6591 | 69,0969108 |
| Upregulated | ENSMUSG00000033031 | 28,55183451 | 76,5910032 | 115,283849 | 47,2148038 | 24,1720375 | 29,19332281 | 52,2510853 | 68,4446572 | 33,1005503 | 11,8656616 | 40,8521924 | 42,04079 | 25,0439116 | 39,29009 | 39,82475 | 35,55028583 | 85,20539 | 138,7192084 | 93,15552 | 394,6605 | 107,1731 | 106,786135 |
| Upregulated | ENSMUSG00000027379 | 30,66678522 | 45,7418492 | 143,424001 | 57,8140454 | 11,4499125 | 7,961815312 | 19,3101837 | 59,8890751 | 35,2360696 | 2,15739302 | 13,2493597 | 10,5102 | 0 | 0 | 0 | 0 | 110,8497 | 175,3807135 | 96,95778 | 637,99 | 124,0019 | 118,451847 |
| Upregulated | ENSMUSG00000022322 | 17,97708099 | 45,7418492 | 140,70076 | 54,9233432 | 5,08884999 | 2,653938437 | 57,9305511 | 45,2223628 | 38,4393487 | 10,7869651 | 4,41645324 | 0 | 0 | 10,29026 | 7,964949 | 0 | 88,51434 | 164,4813471 | 125,4748 | 595,9907 | 155,0024 | 91,5309727 |
| Upregulated | ENSMUSG00000027496 | 24,3219331 | 64,8896 | 117,099342 | 41,4333992 | 27,988675 | 34,50119968 | 43,16394 | 40,3334587 | 57,659023 | 24,8100197 | 8,83290647 | 41,16494 | 8,01405172 | 46,77392 | 33,45279 | 20,05400739 | 92,65053 | 123,8564361 | 87,45212 | 439,9931 | 111,6017 | 80,762623 |
| Upregulated | ENSMUSG00000028068 | 32,78173592 | 39,3592656 | 79,8817219 | 57,8140454 | 2,544425 | 2,653938437 | 39,7562605 | 61,1113011 | 20,287434 | 57,170915 | 19,8740396 | 13,13775 | 1,00175646 | 3,741914 | 0,796495 | 0 | 116,6404 | 175,3807135 | 184,4099 | 445,993 | 168,2883 | 123,836022 |

| | | | | | | | | | | | | | | | | | | | | | | | |
|---|---|---|---|---|---|---|---|---|---|---|---|---|---|---|---|---|---|---|---|---|---|---|---|
| Upregulated | ENSMUSG00000074476 | 37,01163733 | 77,6547672 | 150,685975 | 71,3039894 | 10,1777 | 9,28878453 | 29,5332221 | 52,5557189 | 43,7781471 | 43,1478604 | 9,93701978 | 4,379248 | 9,01580818 | 14,03218 | 9,557939 | 4,557728953 | 205,982 | 313,1090705 | 257,6035 | 612,6571 | 220,5463 | 131,912284 |
| Upregulated | ENSMUSG00000036768 | 9,517278171 | 15,956459 | 17,24719 | 15,4170788 | 5,08884999 | 5,307876874 | 12,4948247 | 40,3334587 | 10,6775969 | 16,1804476 | 6,62467985 | 3,503399 | 7,01229525 | 2,806435 | 7,964949 | 5,469274744 | 43,84355 | 84,22237655 | 58,93512 | 151,331 | 50,4865 | 26,9208743 |
| Upregulated | ENSMUSG00000046057 | 19,03455634 | 21,2752787 | 42,6641015 | 20,2349159 | 0 | 1,326969219 | 9,08714527 | 24,4445204 | 13,8808759 | 8,62957207 | 0 | 3,503399 | 3,00526939 | 1,870957 | 0,796495 | 0 | 30,60776 | 66,38704975 | 42,77549 | 158,6642 | 78,82979 | 19,7419745 |
| Upregulated | ENSMUSG00000040204 | 44,4139648 | 90,4199344 | 157,94795 | 36,6155621 | 3,8166375 | 1,326969219 | 34,0767947 | 96,5558557 | 46,9814262 | 52,8561289 | 12,1452464 | 6,130948 | 8,01405172 | 3,741914 | 7,964949 | 2,734637372 | 145,5937 | 160,5179412 | 142,585 | 451,9929 | 170,9455 | 63,7127359 |
| Upregulated | ENSMUSG00000006715 | 25,37940846 | 56,3794885 | 53,5570636 | 35,6519947 | 6,36106249 | 5,307876874 | 22,7178632 | 59,8890751 | 16,0163953 | 26,9674127 | 4,41645324 | 7,006798 | 3,00526939 | 21,516 | 37,43526 | 12,76164107 | 99,26842 | 108,9936638 | 98,85892 | 267,9958 | 123,1162 | 34,9971366 |
| Upregulated | ENSMUSG00000060509 | 0 | 12,7651672 | 9,98521524 | 4,81783712 | 0 | 0 | 2,27178632 | 6,11113011 | 2,13551937 | 4,31478604 | 4,41645324 | 4,379248 | 3,00526939 | 1,870957 | 3,982475 | 6,380820534 | 21,50816 | 17,8353268 | 20,91246 | 60,66572 | 16,82883 | 17,0498871 |
| Upregulated | ENSMUSG00000037313 | 219,9548733 | 181,903633 | 388,515647 | 176,332839 | 72,5161124 | 157,909337 | 237,40167 | 205,333972 | 139,876519 | 118,656616 | 59,6221187 | 92,84007 | 53,0930926 | 96,35427 | 98,76537 | 76,56984641 | 414,4457 | 532,0872495 | 470,5304 | 1690,64 | 569,5231 | 377,789603 |
| Upregulated | ENSMUSG00000019942 | 127,9545176 | 94,6749901 | 251,445875 | 112,737389 | 2,544425 | 17,25059984 | 89,7355595 | 122,222602 | 33,1005503 | 24,8100197 | 24,2904928 | 7,882647 | 5,00878232 | 9,354784 | 3,982475 | 11,85009528 | 286,2239 | 396,3405955 | 384,0289 | 1157,315 | 429,5781 | 250,364131 |
| Upregulated | ENSMUSG00000035683 | 33,83921127 | 41,4867934 | 80,7894688 | 48,1783712 | 1,2722125 | 0 | 12,4948247 | 28,1111985 | 13,8808759 | 18,3378407 | 3,31233993 | 0 | 3,00526939 | 0 | 11,15093 | 0 | 117,4676 | 75,30471315 | 102,6612 | 353,9945 | 129,3163 | 61,9180109 |
| Upregulated | ENSMUSG00000074026 | 5,287376762 | 10,6376393 | 9,98521524 | 3,8542697 | 0 | 0 | 6,81535895 | 12,2222602 | 2,13551937 | 0 | 2,20822662 | 1,751699 | 0 | 0 | 0,796495 | 0 | 29,78053 | 28,73469318 | 52,28116 | 128,6647 | 38,97203 | 27,8182368 |
| Upregulated | ENSMUSG00000025574 | 51,81629226 | 68,0808918 | 78,0662282 | 45,2876689 | 49,6162874 | 66,34846093 | 93,143239 | 74,5557873 | 52,3202246 | 23,7313232 | 26,4987194 | 63,93703 | 28,049181 | 24,32244 | 33,45279 | 30,08101109 | 217,5633 | 198,1702978 | 231,9382 | 713,3222 | 268,3756 | 127,425472 |
| Upregulated | ENSMUSG00000075266 | 10,57475352 | 34,0404459 | 41,7563546 | 36,6155621 | 16,5387625 | 25,21241515 | 34,0767947 | 52,5557189 | 39,5071084 | 9,70826858 | 25,3946061 | 35,03399 | 29,0509375 | 74,83827 | 50,17918 | 29,1694653 | 81,06921 | 121,8747331 | 93,15552 | 281,9956 | 125,7734 | 56,5338361 |
| Upregulated | ENSMUSG00000068101 | 22,2069824 | 40,4230295 | 46,2950888 | 26,9798879 | 11,4499125 | 1,326969219 | 11,3589316 | 42,7779108 | 18,1519147 | 25,8887162 | 17,6658129 | 37,66154 | 84,147543 | 30,87079 | 23,89485 | 8,203912116 | 104,2318 | 144,6643174 | 176,8054 | 559,9913 | 240,0323 | 73,5837231 |
| Upregulated | ENSMUSG00000045273 | 11,63222888 | 12,7651672 | 30,8633926 | 2,89070227 | 3,8166375 | 0 | 1,13589316 | 6,11113011 | 14,9486356 | 2,15739302 | 17,6658129 | 12,2619 | 17,0298599 | 5,61287 | 1,59299 | 10,93854949 | 45,49803 | 54,49683188 | 47,52832 | 147,9977 | 51,37222 | 20,639337 |
| Upregulated | ENSMUSG00000069910 | 7,402327466 | 23,4028065 | 9,0774684 | 13,4899439 | 0 | 5,307876874 | 13,6307179 | 9,77780817 | 12,8131162 | 0 | 15,4575863 | 14,0136 | 5,00878232 | 21,516 | 10,35443 | 9,115457906 | 59,56105 | 72,33215868 | 50,38002 | 113,9982 | 46,94358 | 40,3813115 |
| Upregulated | ENSMUSG00000032400 | 40,18406339 | 28,7216262 | 10,8929621 | 16,3806462 | 12,722125 | 9,28878453 | 24,9896495 | 24,4445204 | 27,7617518 | 43,1478604 | 12,1452464 | 22,77209 | 10,0175646 | 31,80627 | 21,50536 | 6,380820534 | 68,66066 | 80,25897059 | 62,73739 | 150,6643 | 52,25795 | 50,2522987 |
| Upregulated | ENSMUSG00000026039 | 17,97708099 | 8,51011147 | 10,8929621 | 5,78140454 | 1,2722125 | 3,980907656 | 4,54357263 | 9,77780817 | 1,06775969 | 6,47217906 | 0 | 1,751699 | 1,00175646 | 0,935478 | 6,37196 | 0,911545791 | 18,19921 | 25,76213871 | 17,1102 | 47,33259 | 23,02893 | 25,1261494 |
| Upregulated | ENSMUSG00000029304 | 728,6005177 | 740,379698 | 1393,3914 | 639,80877 | 338,408525 | 122,0811681 | 313,506512 | 250,556334 | 259,465604 | 273,988913 | 227,447342 | 938,9109 | 461,80973 | 122,5477 | 465,1531 | 404,726331 | 1693,354 | 5994,651507 | 1977,178 | 8452,535 | 3403,853 | 2930,78585 |
| Upregulated | ENSMUSG00000063873 | 5,287376762 | 3,1912918 | 1,81549368 | 0,96356742 | 7,63327499 | 18,57756906 | 1,13589316 | 0 | 5,33879843 | 0 | 0 | 0 | 0 | 0,935478 | 0 | 0 | 4,136184 | 5,945108933 | 13,30793 | 29,33288 | 15,9431 | 10,7683497 |
| Upregulated | ENSMUSG00000093738 | 12,68970423 | 4,25505574 | 0 | 0,96356742 | 0 | 31,84726125 | 2,27178632 | 7,33335613 | 0 | 0 | 1,10411331 | 0 | 4,00702586 | 2,806435 | 7,964949 | 7,292366325 | 3,308947 | 0,990851489 | 1,901133 | 58,66575 | 23,02893 | 1,79472495 |
| Upregulated | ENSMUSG00000022070 | 93,057831 | 23,4028065 | 57,1880509 | 44,3241015 | 33,077525 | 45,11695343 | 71,561269 | 55,000171 | 36,3038293 | 18,3378407 | 24,2904928 | 84,08157 | 29,0509375 | 34,6127 | 43,80722 | 28,25791951 | 43,84355 | 83,23152506 | 83,64985 | 231,9964 | 100,973 | 90,6336102 |
| Upregulated | ENSMUSG00000045092 | 44,4139648 | 2,12752787 | 18,1549368 | 22,1620508 | 25,44425 | 35,8281689 | 21,58197 | 31,7778766 | 28,8295115 | 10,7869651 | 24,2904928 | 41,16494 | 23,0403987 | 14,96765 | 37,43526 | 20,96555318 | 22,33539 | 47,56087146 | 34,22039 | 118,6648 | 62,88669 | 43,0733989 |
| Upregulated | ENSMUSG00000032092 | 7,402327466 | 3,1912918 | 5,44648104 | 18,3077811 | 6,36106249 | 2,653938437 | 0 | 3,66667807 | 3,20327906 | 4,31478604 | 1,10411331 | 21,02039 | 14,0245905 | 0 | 7,168455 | 7,292366325 | 17,37197 | 0,990851489 | 10,45623 | 66,66563 | 32,77194 | 11,6657122 |
| Upregulated | ENSMUSG00000006221 | 33,83921127 | 5,31881967 | 6,35422788 | 35,6519947 | 1,2722125 | 0 | 0 | 18,3333903 | 12,8131162 | 11,8656616 | 6,62467985 | 118,2397 | 62,1089008 | 17,77409 | 39,82475 | 0 | 39,70737 | 21,79873275 | 17,1102 | 350,6612 | 62,88669 | 26,0235118 |
| Upregulated | ENSMUSG00000048355 | 5,287376762 | 1,06376393 | 6,35422788 | 12,5263765 | 1,2722125 | 0 | 2,27178632 | 0 | 4,27103874 | 0 | 4,41645324 | 14,88944 | 7,01229525 | 0 | 8,761444 | 2,734637372 | 13,23579 | 9,908514888 | 11,4068 | 63,33234 | 10,62874 | 7,17889982 |
| Upregulated | ENSMUSG00000047747 | 14,80465493 | 5,31881967 | 9,0774684 | 18,3077811 | 0 | 1,326969219 | 3,40767947 | 1,22222602 | 3,20327906 | 0 | 13,2493597 | 3,503399 | 19,0333728 | 24,32244 | 5,575465 | 45,57728953 | 17,37197 | 22,78958424 | 13,30793 | 186,6638 | 53,14368 | 18,844612 |
| Upregulated | ENSMUSG00000075707 | 0 | 4,25505574 | 9,98521524 | 2,89070227 | 3,8166375 | 1,326969219 | 3,40767947 | 19,5556163 | 1,06775969 | 3,23608953 | 0 | 12,2619 | 23,0403987 | 8,419306 | 3,18598 | 3,646183163 | 4,963421 | 8,917663399 | 7,604532 | 130,6646 | 44,2864 | 2,69208743 |
| Upregulated | ENSMUSG00000021806 | 812,1410706 | 285,088734 | 1640,29854 | 324,722222 | 276,070112 | 264,0668745 | 335,088482 | 705,224415 | 463,407704 | 612,699671 | 425,083624 | 225,9692 | 374,656918 | 104,7736 | 118,6777 | 129,4395023 | 541,0129 | 433,0021006 | 1042,771 | 12135,14 | 2832,558 | 1226,69451 |
| Upregulated | ENSMUSG00000027996 | 259,0814613 | 106,376393 | 504,707243 | 155,134355 | 21,6276125 | 25,21241515 | 5,67946579 | 134,444862 | 102,50493 | 510,223449 | 134,701824 | 161,1563 | 213,374127 | 41,16105 | 14,33691 | 31,90410267 | 83,55092 | 356,706536 | 542,7735 | 7961,209 | 1906,087 | 163,319971 |
| Upregulated | ENSMUSG00000082676 | 2,114950705 | 1,06376393 | 6,35422788 | 1,92713485 | 3,8166375 | 1,326969219 | 3,40767947 | 4,88890409 | 1,06775969 | 8,62957207 | 4,41645324 | 9,634347 | 7,01229525 | 6,548349 | 3,18598 | 1,823091581 | 12,40855 | 2,972554466 | 2,851699 | 153,9976 | 34,54339 | 6,28153734 |
| Upregulated | ENSMUSG00000028358 | 2,114950705 | 1,06376393 | 0,90774684 | 4,81783712 | 5,08884999 | 0 | 2,27178632 | 1,22222602 | 8,54207749 | 2,15739302 | 12,1452464 | 0 | 2,00351293 | 24,32244 | 2,389485 | 0,911545791 | 4,963421 | 7,92681191 | 10,45623 | 231,9964 | 43,40067 | 3,58944991 |
| Upregulated | ENSMUSG00000096225 | 30,66678522 | 15,956459 | 30,8633926 | 7,70853939 | 1,2722125 | 127,389045 | 19,3101837 | 61,1113011 | 24,5584728 | 19,4165372 | 12,1452464 | 47,29588 | 21,0544504 | 29,93531 | 118,6777 | 79,30448379 | 39,70737 | 38,64320806 | 101,7106 | 491,9923 | 120,459 | 23,3314244 |
| Upregulated | ENSMUSG00000024063 | 20,09203169 | 30,8491541 | 59,0035446 | 39,5062644 | 30,5331 | 17,25059984 | 52,2510853 | 63,5557531 | 45,9136665 | 33,4395918 | 39,7480791 | 37,66154 | 23,0403987 | 29,93531 | 59,73712 | 87,5083959 | 70,31513 | 50,53342593 | 79,84759 | 265,9958 | 93,00144 | 43,0733989 |
| Upregulated | ENSMUSG00000036136 | 5,287376762 | 0 | 14,5239494 | 0,96356742 | 0 | 5,307876874 | 0 | 0 | 0 | 0 | 0 | 2,627549 | 5,00878232 | 0 | 0,796495 | 0 | 9,926842 | 1,981702978 | 7,604532 | 486,6591 | 59,34378 | 155,243709 |
| Upregulated | ENSMUSG00000048988 | 23,26445775 | 19,1477508 | 12,7084558 | 16,3806462 | 21,6276125 | 23,88544594 | 27,2614358 | 52,5557189 | 40,5748681 | 34,5182883 | 34,2275126 | 21,02039 | 41,072015 | 46,77392 | 58,94063 | 36,46183163 | 19,85368 | 15,85362382 | 38,97323 | 240,6629 | 40,74349 | 73,5837231 |
| Upregulated | ENSMUSG00000004371 | 58,16114438 | 35,1042098 | 87,1436966 | 26,0163204 | 48,344075 | 50,42483031 | 54,5228716 | 41,5556847 | 45,9136665 | 42,0691639 | 33,1233993 | 381,8705 | 653,145215 | 241,3534 | 385,5036 | 331,8026678 | 1596,567 | 593,5200418 | 813,6849 | 5649,212 | 1207,247 | 313,179505 |
| Upregulated | ENSMUSG00000006219 | 61,33357043 | 31,912918 | 37,7711394 | 43,3605341 | 15,26655 | 14,5966614 | 2,27178632 | 51,3334929 | 10,6775969 | 7,55087556 | 51,8933255 | 174,2941 | 137,240636 | 216,0955 | 141,7761 | 133,9972312 | 413,6184 | 160,5179412 | 252,8507 | 1319,313 | 169,1741 | 75,3784481 |
| Upregulated | ENSMUSG00000029371 | 186,115662 | 94,6749901 | 600,020661 | 391,208374 | 188,28745 | 55,73270718 | 56,7946579 | 122,222602 | 123,860124 | 45,3052534 | 363,253279 | 7172,333 | 10847,019 | 10171,46 | 6717,638 | 2313,503217 | 12301,84 | 10289,99271 | 13559,83 | 81537,39 | 11983,01 | 1721,14123 |
| Upregulated | ENSMUSG00000052353 | 9,517278171 | 1,06376393 | 0,90774684 | 10,5992417 | 3,8166375 | 2,653938437 | 5,67946579 | 1,22222602 | 0 | 4,31478604 | 4,41645324 | 113,8605 | 14,0245905 | 22,45148 | 24,69134 | 42,84265216 | 163,7929 | 78,27726762 | 230,9877 | 1036,65 | 30,11475 | 172,293596 |
| Upregulated | ENSMUSG00000026980 | 7,402327466 | 11,7014033 | 6,35422788 | 0,96356742 | 5,08884999 | 0 | 5,67946579 | 2,44445204 | 11,7453565 | 35,5969848 | 32,019286 | 16,64114 | 7,01229525 | 3,741914 | 54,95815 | 20,05400739 | 93,47776 | 63,41449528 | 67,49022 | 692,6559 | 10,62874 | 63,7127359 |
| Upregulated | ENSMUSG00000041605 | 0 | 0 | 0 | 2,89070227 | 0 | 0 | 0 | 0 | 0 | 0 | 1,10411331 | 1,751699 | 4,00702586 | 5,61287 | 8,761444 | 1,823091581 | 22,33539 | 3,963405955 | 17,1102 | 122,6648 | 2,657184 | 8,97362477 |
| Upregulated | ENSMUSG00000000805 | 17,97708099 | 6,3825836 | 10,8929621 | 14,4535114 | 6,36106249 | 0 | 0 | 0 | 0 | 0 | 5,52056654 | 173,4182 | 223,391692 | 173,0635 | 52,56867 | 18,23091581 | 754,44 | 37,65235657 | 919,1978 | 7871,877 | 387,0631 | 567,133086 |
| Upregulated | ENSMUSG00000027843 | 0 | 0 | 0 | 0 | 0 | 0 | 0 | 0 | 0 | 0 | 0 | 4,379248 | 1,00175646 | 7,483827 | 0,796495 | 0 | 18,19921 | 9,908514888 | 22,8136 | 140,6645 | 8,85728 | 10,7683497 |
| Upregulated | ENSMUSG00000020007 | 0 | 1,06376393 | 0 | 0 | 0 | 0 | 0 | 0 | 0 | 0 | 0 | 2,627549 | 3,00526939 | 0 | 11,94742 | 9,115457906 | 21,50816 | 6,935960422 | 14,2585 | 81,33206 | 0 | 7,17889982 |
| Upregulated | ENSMUSG00000074577 | 65,56347184 | 12,7651672 | 109,837368 | 38,542697 | 13,9943375 | 27,86635359 | 19,3101837 | 23,2222944 | 14,9486356 | 99,2400788 | 39,7480791 | 105,102 | 98,1721335 | 98,22523 | 72,48104 | 40,10801479 | 378,0472 | 155,5636837 | 337,4511 | 1455,311 | 76,17261 | 60,123286 |
| Upregulated | ENSMUSG00000019359 | 4,229901409 | 0 | 0 | 8,67210682 | 0 | 2,653938437 | 0 | 1,22222602 | 2,13551937 | 0 | 0 | 0 | 4,00702586 | 0 | 0 | 0 | 28,95329 | 7,92681191 | 19,9619 | 110,6649 | 4,42864 | 2,69208743 |
| Upregulated | ENSMUSG00000062991 | 13,74717958 | 11,7014033 | 61,7267851 | 20,2349159 | 19,0831875 | 7,961815312 | 7,95125211 | 14,6667123 | 10,6775969 | 30,2035023 | 33,1233993 | 54,30268 | 11,0193211 | 30,87079 | 44,60372 | 51,95811007 | 206,8092 | 89,17663399 | 78,89702 | 802,6541 | 70,85824 | 87,9415228 |
| Upregulated | ENSMUSG00000057604 | 15,86213028 | 11,7014033 | 33,5986283 | 52,9962083 | 3,8166375 | 18,57756906 | 36,3485811 | 20,7778424 | 11,7453565 | 10,7869651 | 12,1452464 | 311,8025 | 384,674482 | 389,159 | 278,8394 | 344,5643089 | 2496,601 | 539,0232099 | 1391,629 | 11463,15 | 1495,995 | 726,863607 |
| Upregulated | ENSMUSG00000032332 | 1109,291645 | 816,970701 | 1292,6315 | 1182,29723 | 187,015237 | 165,8711523 | 205,596662 | 361,778902 | 215,687457 | 831,675009 | 441,645324 | 834,6848 | 1113,95319 | 1307,799 | 864,9935 | 1020,931286 | 10562,16 | 4979,028731 | 9524,676 | 44712,64 | 5262,11 | 2161,74621 |
| Upregulated | ENSMUSG00000026070 | 0 | 1,06376393 | 0 | 4,81783712 | 0 | 0 | 0 | 3,66667807 | 3,20327906 | 0 | 0 | 16,64114 | 6,01053879 | 0 | 2,389485 | 0 | 9,926842 | 2,972554466 | 19,9619 | 152,6643 | 29,22902 | 17,9472495 |
| Upregulated | ENSMUSG00000029377 | 52,87376762 | 152,118243 | 112,560608 | 72,2675568 | 188,28745 | 35,8281689 | 27,2614358 | 97,7780817 | 89,6918136 | 152,096208 | 534,390841 | 1632,584 | 339,595441 | 435,9329 | 754,2807 | 773,9023763 | 2115,245 | 767,9099038 | 2039,916 | 13050,46 | 1573,939 | 1171,9554 |
| Upregulated | ENSMUSG00000048040 | 7,402327466 | 8,51011147 | 11,8007089 | 20,2349159 | 2,544425 | 11,94272297 | 4,54357263 | 9,77780817 | 4,27103874 | 7,55087556 | 20,9781529 | 17,51699 | 21,0368858 | 13,0967 | 19,91237 | 24,61173635 | 39,70737 | 35,6706536 | 48,47889 | 301,9953 | 57,57232 | 43,9707614 |
| Upregulated | ENSMUSG00000024173 | 2,114950705 | 0 | 9,98521524 | 0 | 0 | 0 | 0 | 0 | 0 | 15,1017511 | 0 | 0 | 1,00175646 | 0 | 0 | 0 | 8,272368 | 33,68895062 | 35,17096 | 245,9962 | 38,0863 | 10,7683497 |
| Upregulated | ENSMUSG00000038305 | 193,5179895 | 172,329757 | 278,67828 | 263,053907 | 151,393287 | 233,5465825 | 198,781303 | 149,111575 | 142,012038 | 353,812455 | 218,614435 | 201,4454 | 156,274008 | 136,5798 | 207,8852 | 148,5819639 | 316,8317 | 244,7403177 | 514,2565 | 1918,637 | 327,7194 | 182,164583 |
| Upregulated | ENSMUSG00000029859 | 14,80465493 | 6,3825836 | 8,16972156 | 10,5992417 | 8,90548749 | 29,19332281 | 20,4460768 | 23,2222944 | 5,33879843 | 0 | 12,1452464 | 10,5102 | 25,0439116 | 9,354784 | 19,91237 | 11,85009528 | 26,47158 | 20,80788127 | 31,36869 | 273,3291 | 51,37222 | 9,87098725 |
| Upregulated | ENSMUSG00000062300 | 137,4717958 | 109,567685 | 220,582482 | 188,859215 | 179,381962 | 327,761397 | 290,788649 | 211,445102 | 243,449208 | 265,359341 | 226,343228 | 225,9692 | 127,223071 | 236,676 | 189,5658 | 287,1369241 | 400,3826 | 278,4292684 | 370,7209 | 2289,964 | 315,3192 | 192,932933 |
| Upregulated | ENSMUSG00000025069 | 32,78173592 | 40,4230295 | 13,6162026 | 22,1620508 | 20,3554 | 41,13604578 | 26,1255426 | 47,6668148 | 30,9650309 | 35,5969848 | 22,0822662 | 20,14454 | 22,0386422 | 9,354784 | 27,08083 | 12,76164107 | 51,28868 | 25,76213871 | 62,73739 | 380,6607 | 29,22902 | 47,5602113 |
| Upregulated | ENSMUSG00000062609 | 0 | 2,12752787 | 3,63098736 | 1,92713485 | 1,2722125 | 0 | 0 | 7,33335613 | 0 | 0 | 0 | 42,04079 | 76,1334913 | 101,9671 | 8,761444 | 0 | 34,74395 | 0 | 120,7219 | 1124,649 | 102,7444 | 47,5602113 |
| Upregulated | ENSMUSG00000038963 | 0 | 1,06376393 | 10,8929621 | 18,3077811 | 7,63327499 | 1,326969219 | 35,2126879 | 19,5556163 | 25,6262325 | 22,6526267 | 0 | 31,53059 | 14,0245905 | 0 | 1,59299 | 14,58473265 | 28,12605 | 21,79873275 | 53,23172 | 913,9857 | 48,71504 | 111,272947 |
| Upregulated | ENSMUSG00000062184 | 15,86213028 | 18,0839869 | 39,0331141 | 10,5992417 | 31,8053125 | 19,90453828 | 13,6307179 | 45,2223628 | 46,9814262 | 46,3839499 | 66,2467985 | 13,13775 | 8,01405172 | 14,03218 | 17,52289 | 39,196469 | 68,66066 | 91,15833697 | 108,3646 | 987,9846 | 93,00144 | 41,278674 |
| Upregulated | ENSMUSG00000013584 | 2,114950705 | 12,7651672 | 6,35422788 | 1,92713485 | 13,9943375 | 6,634846093 | 0 | 1,22222602 | 4,27103874 | 7,55087556 | 5,52056654 | 50,79928 | 55,0966056 | 27,12887 | 23,89485 | 0 | 50,46145 | 13,87192084 | 103,6117 | 2267,298 | 269,2613 | 17,0498871 |
| Upregulated | ENSMUSG00000021728 | 182,943236 | 154,24577 | 139,793013 | 194,64062 | 167,93205 | 159,2363062 | 147,666111 | 201,667294 | 201,806581 | 131,600974 | 143,53473 | 116,488 | 297,52167 | 202,0633 | 139,3866 | 162,2551507 | 138,1486 | 235,8226543 | 190,1133 | 7940,543 | 736,9257 | 320,358404 |
| Upregulated | ENSMUSG00000027997 | 318,300081 | 241,474413 | 218,766988 | 216,80267 | 162,8432 | 261,4129361 | 171,519867 | 185,778355 | 172,977069 | 183,378407 | 184,386923 | 160,2805 | 130,22834 | 220,7729 | 223,0186 | 145,8473265 | 265,543 | 229,8775454 | 452,4697 | 1135,982 | 321,5193 | 204,598645 |
| Upregulated | ENSMUSG00000034987 | 10,57475352 | 1,06376393 | 10,8929621 | 0,96356742 | 2,544425 | 0 | 15,9025042 | 1,22222602 | 8,54207749 | 0 | 0 | 7,882647 | 13,022834 | 0 | 0 | 2,734637372 | 0,827237 | 0 | 33,26983 | 134,6646 | 19,48602 | 2,69208743 |
| Upregulated | ENSMUSG00000068335 | 151,2189754 | 95,7387541 | 174,287393 | 132,972305 | 183,1986 | 115,446322 | 141,986645 | 96,5558557 | 140,944279 | 148,860118 | 200,948622 | 129,6258 | 225,395205 | 196,4505 | 153,7235 | 132,1741396 | 214,2543 | 218,978179 | 297,5273 | 867,9865 | 259,5183 | 165,114696 |
| Upregulated | ENSMUSG00000022309 | 127,9545176 | 127,651672 | 197,888811 | 198,494889 | 96,6881499 | 72,98330702 | 47,7075126 | 253,000787 | 163,367232 | 98,1613823 | 154,575863 | 95,46762 | 221,388179 | 112,2574 | 93,18991 | 121,2355902 | 242,3804 | 155,5636837 | 354,5613 | 991,3179 | 414,5207 | 333,818842 |
| Upregulated | ENSMUSG00000066829 | 37,01163733 | 40,4230295 | 65,3577725 | 55,8869106 | 71,2438999 | 34,50119968 | 52,2510853 | 56,222397 | 42,7103874 | 26,9674127 | 13,2493597 | 37,66154 | 26,0456681 | 35,54818 | 23,89485 | 20,05400739 | 79,41474 | 47,56087146 | 137,8321 | 354,6611 | 136,4021 | 97,81251 |
| Upregulated | ENSMUSG00000049866 | 19,03455634 | 11,7014033 | 29,9556457 | 9,63567424 | 6,36106249 | 7,961815312 | 11,3589316 | 23,2222944 | 23,4907131 | 21,5739302 | 25,3946061 | 7,882647 | 0 | 6,548349 | 4,77897 | 3,646183163 | 29,78053 | 17,8353268 | 26,61586 | 110,6649 | 37,20058 | 20,639337 |
| Upregulated | ENSMUSG00000022090 | 363,7715212 | 198,923856 | 411,209318 | 330,503626 | 251,898075 | 295,9141358 | 256,711854 | 261,556369 | 280,820797 | 389,40944 | 200,948622 | 76,19892 | 172,302112 | 40,22557 | 100,3584 | 72,92366325 | 269,6792 | 283,3835258 | 389,7323 | 1408,645 | 593,4378 | 256,645669 |
| Upregulated | ENSMUSG00000022652 | 9,517278171 | 15,956459 | 24,5091647 | 7,70853939 | 13,9943375 | 23,88544594 | 2,27178632 | 22,0000684 | 1,06775969 | 9,70826858 | 12,1452464 | 2,627549 | 15,026347 | 0 | 9,557939 | 7,292366325 | 29,78053 | 4,954257444 | 15,20906 | 161,3308 | 62,00096 | 12,5630747 |
| Upregulated | ENSMUSG00000025272 | 3,172426057 | 8,51011147 | 13,6162026 | 5,78140454 | 11,4499125 | 29,19332281 | 3,40767947 | 17,1111643 | 20,287434 | 11,8656616 | 6,62467985 | 4,379248 | 2,00351293 | 0 | 10,35443 | 0 | 26,47158 | 6,935960422 | 19,01133 | 117,9982 | 44,2864 | 21,5366995 |
| Upregulated | ENSMUSG00000054555 | 432,5074191 | 144,671895 | 634,515041 | 285,215958 | 36,8941625 | 79,61815312 | 199,917196 | 139,333766 | 51,2524649 | 577,102632 | 397,480791 | 315,3059 | 53,0930926 | 471,4811 | 207,8852 | 145,8473265 | 590,6471 | 469,6636057 | 423,9527 | 2245,298 | 785,6407 | 358,944991 |
| Upregulated | ENSMUSG00000117629 | 4,229901409 | 3,1912918 | 15,4316963 | 0,96356742 | 0 | 0 | 6,81535895 | 1,22222602 | 0 | 2,15739302 | 7,72879316 | 3,503399 | 1,00175646 | 5,61287 | 2,389485 | 0,911545791 | 5,790658 | 7,92681191 | 6,653965 | 50,66588 | 15,9431 | 5,38417486 |
| Upregulated | ENSMUSG00000035365 | 34,89668663 | 11,7014033 | 40,8486078 | 13,4899439 | 0 | 2,653938437 | 14,7666111 | 4,88890409 | 4,27103874 | 0 | 0 | 0 | 6,01053879 | 0 | 0,796495 | 0 | 4,963421 | 18,82617829 | 13,30793 | 121,9981 | 23,02893 | 18,844612 |
| Upregulated | ENSMUSG00000040856 | 8,459802819 | 7,44634754 | 68,9887598 | 2,89070227 | 1,2722125 | 0 | 0 | 11,0000342 | 5,33879843 | 5,39348255 | 2,20822662 | 7,882647 | 4,00702586 | 0 | 0,796495 | 7,292366325 | 0 | 0 | 4,752832 | 175,3306 | 21,25747 | 11,6657122 |
| Upregulated | ENSMUSG00000020389 | 21,14950705 | 25,5303344 | 67,1732662 | 50,105506 | 26,7164625 | 30,52029203 | 96,5509185 | 35,4445546 | 39,5071084 | 26,9674127 | 40,8521924 | 29,77889 | 59,1036314 | 39,29009 | 19,91237 | 9,115457906 | 46,32526 | 58,46023784 | 102,6612 | 176,6639 | 77,05834 | 108,58086 |
| Upregulated | ENSMUSG00000029093 | 93,057831 | 22,3390426 | 132,531039 | 155,134355 | 71,2438999 | 39,80907656 | 60,2023374 | 85,5558215 | 64,0655811 | 193,086675 | 101,578424 | 7,006798 | 175,307381 | 101,0317 | 29,47031 | 43,75419795 | 195,2279 | 128,8106935 | 135,931 | 727,322 | 370,2343 | 365,226528 |
| Upregulated | ENSMUSG00000079293 | 0 | 0 | 1,81549368 | 0 | 0 | 0 | 0 | 4,88890409 | 2,13551937 | 0 | 2,20822662 | 0 | 13,022834 | 0 | 0 | 0 | 0 | 2,972554466 | 0 | 36,66609 | 15,9431 | 19,7419745 |
| Upregulated | ENSMUSG00000034275 | 41,24153874 | 9,57387541 | 15,4316963 | 7,70853939 | 41,9830125 | 35,8281689 | 12,4948247 | 14,6667123 | 13,8808759 | 25,8887162 | 4,41645324 | 0,87585 | 39,0685021 | 30,87079 | 38,23176 | 21,87709898 | 45,49803 | 10,89936638 | 120,7219 | 190,6637 | 98,31581 | 127,425472 |
| Upregulated | ENSMUSG00000070780 | 12,68970423 | 3,1912918 | 4,5387342 | 8,67210682 | 8,90548749 | 6,634846093 | 0 | 1,22222602 | 4,27103874 | 1,07869651 | 3,31233993 | 7,006798 | 26,0456681 | 4,677392 | 1,59299 | 2,734637372 | 5,790658 | 9,908514888 | 19,9619 | 61,99903 | 41,62922 | 52,9443862 |
| Upregulated | ENSMUSG00000035835 | 189,2880881 | 25,5303344 | 45,387342 | 55,8869106 | 59,7939874 | 63,69452249 | 94,2791321 | 51,3334929 | 23,4907131 | 31,2821988 | 102,682538 | 94,59177 | 133,23361 | 46,77392 | 56,55114 | 40,10801479 | 120,7766 | 126,8289906 | 169,2008 | 357,3278 | 246,2324 | 234,211607 |
| Upregulated | ENSMUSG00000032093 | 88,82792959 | 14,8926951 | 30,8633926 | 54,9233432 | 54,7051374 | 116,7732912 | 78,3766279 | 62,3335271 | 44,8459068 | 7,55087556 | 176,658129 | 200,5696 | 207,363588 | 54,25775 | 73,27754 | 61,07356797 | 158,8295 | 112,9570697 | 158,7446 | 717,9888 | 354,2912 | 325,742579 |
| Upregulated | ENSMUSG00000027239 | 75,08075001 | 26,5940983 | 62,634532 | 78,0489613 | 27,988675 | 41,13604578 | 21,58197 | 73,3335613 | 14,9486356 | 43,1478604 | 52,9974388 | 61,30948 | 130,22834 | 72,03184 | 25,48784 | 20,05400739 | 59,56105 | 48,55172295 | 57,98456 | 219,9966 | 174,4884 | 52,9443862 |
| Upregulated | ENSMUSG00000021508 | 1855,869243 | 1298,85576 | 3239,74847 | 1824,03313 | 389,297025 | 84,92602999 | 251,032388 | 843,335955 | 600,080943 | 594,361777 | 757,42173 | 3241,52 | 24775,4409 | 7399,634 | 980,4853 | 1267,960195 | 10005,43 | 2819,963337 | 11400,14 | 32599,49 | 18727,83 | 6150,52242 |
| Upregulated | ENSMUSG00000022180 | 66,6209472 | 25,5303344 | 51,7415699 | 29,8705901 | 12,722125 | 15,92363062 | 15,9025042 | 44,0001368 | 13,8808759 | 12,9443581 | 5,52056654 | 27,15134 | 236,414526 | 61,74157 | 12,74392 | 6,380820534 | 59,56105 | 79,2681191 | 107,414 | 366,6609 | 225,8606 | 78,967898 |
| Upregulated | ENSMUSG00000084989 | 25,37940846 | 6,3825836 | 35,4021268 | 7,70853939 | 2,544425 | 2,653938437 | 2,27178632 | 12,2222602 | 5,33879843 | 0 | 0 | 27,15134 | 54,0948491 | 18,70957 | 11,15093 | 0,911545791 | 74,45132 | 18,82617829 | 38,02266 | 151,331 | 79,71552 | 45,7654863 |
| Upregulated | ENSMUSG00000000359 | 10,57475352 | 6,3825836 | 20,8781773 | 3,8542697 | 12,722125 | 3,980907656 | 11,3589316 | 12,2222602 | 2,13551937 | 1,07869651 | 4,41645324 | 9,634347 | 36,0632327 | 3,741914 | 15,9299 | 4,557728953 | 32,26224 | 10,89936638 | 19,9619 | 77,99878 | 51,37222 | 33,2024117 |

| | | | | | | | | | | | | | | | | | | | | | | | |
|---|---|---|---|---|---|---|---|---|---|---|---|---|---|---|---|---|---|---|---|---|---|---|---|
| Upregulated | ENSMUSG00000040473 | 7,402327466 | 9,57387541 | 8,16972156 | 1,92713485 | 8,90548749 | 2,653938437 | 4,54357263 | 6,11113011 | 2,13551937 | 4,31478604 | 0 | 28,90304 | 30,0526939 | 36,48366 | 3,18598 | 7,292366325 | 53,77039 | 6,935960422 | 21,86303 | 83,99869 | 42,51494 | 26,9208743 |
| Upregulated | ENSMUSG00000020374 | 6,344852114 | 3,1912918 | 5,44648104 | 0,96356742 | 1,2722125 | 14,5966614 | 0 | 8,55558215 | 4,27103874 | 5,39348255 | 5,52056654 | 18,39284 | 21,0368858 | 5,61287 | 15,1334 | 2,734637372 | 20,68092 | 5,945108933 | 4,752832 | 61,99903 | 40,74349 | 12,5630747 |
| Upregulated | ENSMUSG00000095419 | 5,287376762 | 1,06376393 | 2,72324052 | 5,78140454 | 3,8166375 | 13,26969219 | 11,3589316 | 1,22222602 | 11,7453565 | 4,31478604 | 9,93701978 | 9,634347 | 20,0351293 | 11,22574 | 10,35443 | 13,67318686 | 26,47158 | 6,935960422 | 11,4068 | 50,66588 | 28,3433 | 13,4604372 |
| Upregulated | ENSMUSG00000027533 | 68,7358979 | 59,5707803 | 114,376102 | 76,1218265 | 113,226912 | 138,0047987 | 128,355927 | 130,778184 | 155,892914 | 96,0039893 | 94,9537446 | 109,4812 | 225,395205 | 154,3539 | 153,7235 | 107,5624033 | 272,1609 | 176,371565 | 246,1967 | 729,9886 | 335,6909 | 208,188095 |
| Upregulated | ENSMUSG00000044254 | 69,79337325 | 8,51011147 | 63,5422788 | 13,4899439 | 24,1720375 | 41,13604578 | 101,094491 | 293,334245 | 85,4207749 | 122,971402 | 225,239115 | 319,6851 | 143,251174 | 742,7698 | 238,152 | 360,0605873 | 778,4299 | 552,8951308 | 536,1195 | 1547,976 | 1458,794 | 936,846426 |
| Upregulated | ENSMUSG00000028179 | 2,114950705 | 7,44634754 | 29,9556457 | 6,74497197 | 30,5331 | 27,86635359 | 13,6307179 | 15,8889383 | 9,60983717 | 33,4395918 | 36,4357392 | 56,05438 | 48,0843103 | 37,41914 | 22,30186 | 35,55028583 | 62,04276 | 52,51512891 | 38,97323 | 120,6648 | 97,43008 | 59,2259235 |
| Upregulated | ENSMUSG00000045672 | 78,25317607 | 19,1477508 | 86,2359498 | 24,0891856 | 2,544425 | 13,26969219 | 12,4948247 | 31,7778766 | 44,8459068 | 35,5969848 | 45,2686457 | 40,28909 | 141,247662 | 89,80593 | 43,80722 | 163,1666965 | 196,0551 | 124,8472876 | 148,2884 | 463,3261 | 277,2329 | 80,762623 |
| Upregulated | ENSMUSG00000079138 | 236,8744789 | 86,1648786 | 175,19514 | 155,134355 | 47,0718625 | 112,7923836 | 89,7355595 | 74,5557873 | 26,6939921 | 147,781422 | 83,9126115 | 478,2139 | 451,792166 | 303,095 | 223,0186 | 230,621085 | 597,265 | 317,0724764 | 461,0247 | 1838,638 | 999,1012 | 422,657727 |
| Upregulated | ENSMUSG00000015653 | 368,0014226 | 247,856997 | 386,700154 | 240,891856 | 211,187275 | 63,69452249 | 143,122538 | 238,334074 | 105,708209 | 115,420526 | 92,7455179 | 684,9145 | 712,248846 | 852,2208 | 465,9495 | 473,0922653 | 967,8671 | 553,8859822 | 946,7642 | 2866,622 | 1484,48 | 410,992015 |
| Upregulated | ENSMUSG00000016458 | 0 | 4,25505574 | 1,81549368 | 0,96356742 | 0 | 0 | 0 | 0 | 0 | 0 | 0 | 7,882647 | 17,0298599 | 14,96765 | 4,77897 | 0 | 23,98987 | 21,79873275 | 13,30793 | 108,665 | 77,05834 | 3,58944991 |
| Upregulated | ENSMUSG00000030067 | 179,7708099 | 191,477508 | 129,807798 | 150,316518 | 103,049212 | 108,8114759 | 124,948247 | 156,444931 | 125,995643 | 126,207492 | 173,345789 | 244,3621 | 419,735959 | 400,3848 | 153,7235 | 261,6136419 | 408,655 | 215,0147731 | 259,5047 | 1050,65 | 791,8408 | 242,287869 |
| Upregulated | ENSMUSG00000020486 | 7,402327466 | 14,8926951 | 13,6162026 | 43,3605341 | 26,7164625 | 26,53938437 | 21,58197 | 24,4445204 | 29,8972712 | 24,8100197 | 28,706946 | 38,53739 | 38,0667457 | 39,29009 | 27,87732 | 27,34637372 | 160,4839 | 77,28641613 | 84,60042 | 300,662 | 201,0603 | 61,9180109 |
| Upregulated | ENSMUSG00000079652 | 23,26445775 | 6,3825836 | 13,6162026 | 11,5628091 | 5,08884999 | 29,19332281 | 14,7666111 | 35,4445546 | 14,9486356 | 10,7869651 | 15,4575863 | 37,66154 | 26,0456681 | 14,03218 | 27,87732 | 17,31937002 | 67,83342 | 43,59746551 | 40,87436 | 129,3313 | 83,25843 | 34,9971366 |
| Upregulated | ENSMUSG00000003865 | 277,0585423 | 199,98762 | 249,630381 | 157,06149 | 95,4159374 | 110,1384451 | 123,812354 | 125,88928 | 89,6918136 | 257,808466 | 141,326504 | 284,6511 | 152,266983 | 215,16 | 185,5833 | 159,5205134 | 373,0838 | 276,4475654 | 335,55 | 853,9867 | 691,7536 | 272,798193 |
| Upregulated | ENSMUSG00000032334 | 4356,798452 | 2496,65395 | 3726,30078 | 4990,31569 | 1669,1428 | 2246,558887 | 2236,57363 | 2853,89776 | 2365,0877 | 2745,28262 | 2711,70229 | 4043,798 | 4901,59438 | 3906,558 | 4134,605 | 4864,008339 | 6109,144 | 5440,765525 | 6310,811 | 12760,47 | 10968,86 | 3928,65293 |
| Upregulated | ENSMUSG00000061353 | 21286,97884 | 5884,74208 | 17823,6092 | 7465,7204 | 2698,36271 | 2387,217624 | 2713,64876 | 5874,01826 | 5182,90551 | 3562,93457 | 8188,1043 | 13930,39 | 27272,8197 | 18409,28 | 9305,45 | 10848,30645 | 29021,12 | 18555,67583 | 30994,17 | 88065,96 | 58437,68 | 20812,5279 |
| Upregulated | ENSMUSG00000021260 | 243,219331 | 74,4634754 | 218,766988 | 115,628091 | 29,2608875 | 21,2315075 | 35,2126879 | 18,3333903 | 13,8808759 | 74,4300591 | 59,6221187 | 175,1699 | 95,1668641 | 76,70923 | 134,6076 | 126,7048649 | 262,2341 | 179,3441195 | 407,793 | 1013,984 | 707,6967 | 159,730521 |
| Upregulated | ENSMUSG00000029126 | 670,4393734 | 298,917665 | 705,319295 | 403,734751 | 69,9716874 | 294,5871665 | 161,296828 | 319,000992 | 214,619697 | 283,697182 | 249,529608 | 615,7223 | 852,494751 | 207,6762 | 681,7997 | 811,2757537 | 1020,81 | 802,5897059 | 1115,014 | 3387,28 | 2233,806 | 634,435272 |
| Upregulated | ENSMUSG00000034040 | 63,44852114 | 58,5070164 | 69,8965067 | 114,664523 | 29,2608875 | 49,09786109 | 106,773957 | 47,6668148 | 42,7103874 | 104,633561 | 51,8933255 | 135,7567 | 94,1651077 | 85,12853 | 110,7128 | 92,06612485 | 100,9229 | 76,29556464 | 66,53965 | 267,3292 | 196,6316 | 115,75976 |
| Upregulated | ENSMUSG00000021090 | 0 | 1,06376393 | 0 | 0 | 0 | 0 | 0 | 0 | 0 | 0 | 0 | 0 | 0 | 14,96765 | 0 | 0 | 0 | 0 | 10,45623 | 41,33269 | 15,9431 | 14,3577996 |
| Upregulated | ENSMUSG00000038112 | 248,5067078 | 139,353075 | 329,512103 | 153,20722 | 62,3384124 | 54,40573796 | 23,8537563 | 135,667088 | 130,266682 | 75,5087556 | 126,973031 | 504,4894 | 777,363016 | 598,7062 | 504,1813 | 733,7943615 | 196,0551 | 181,3258225 | 474,3327 | 2155,3 | 1038,073 | 591,361873 |
| Upregulated | ENSMUSG00000041642 | 1,057475352 | 6,3825836 | 2,72324052 | 0,96356742 | 7,63327499 | 6,634846093 | 0 | 6,11113011 | 3,20327906 | 2,15739302 | 1,10411331 | 3,503399 | 7,01229525 | 14,96765 | 5,575465 | 17,31937002 | 1,654474 | 5,945108933 | 11,4068 | 47,33259 | 31,00048 | 12,5630747 |
| Upregulated | ENSMUSG00000111840 | 5,287376762 | 15,956459 | 10,8929621 | 1,92713485 | 0 | 0 | 0 | 12,2222602 | 0 | 0 | 0 | 10,5102 | 16,0281034 | 2,806435 | 0,796495 | 10,93854949 | 9,099605 | 14,86277233 | 6,653965 | 16,66641 | 7,085824 | 10,7683497 |
| Upregulated | ENSMUSG00000095366 | 445,1971233 | 112,758977 | 387,607901 | 104,065282 | 111,9547 | 176,4869061 | 95,4150253 | 744,335647 | 146,283077 | 60,4070045 | 231,863795 | 796,1474 | 454,797435 | 270,3533 | 101,1549 | 374,64532 | 653,5171 | 442,9106155 | 122,6231 | 1016,651 | 132,8592 | 395,736853 |
| Upregulated | ENSMUSG00000087263 | 114,2073381 | 25,5303344 | 99,8521524 | 21,1984833 | 38,166375 | 38,48210734 | 29,5332221 | 200,445068 | 43,7781471 | 25,8887162 | 66,2467985 | 315,3059 | 227,398717 | 97,28975 | 23,09835 | 100,270037 | 119,1221 | 230,8683969 | 41,82493 | 235,3297 | 36,31485 | 150,756896 |
| Upregulated | ENSMUSG00000063018 | 2,114950705 | 2,12752787 | 10,8929621 | 2,89070227 | 2,544425 | 11,94272297 | 11,3589316 | 4,88890409 | 7,4743178 | 11,8656616 | 13,2493597 | 6,130948 | 30,0526939 | 16,83861 | 21,50536 | 21,87709898 | 21,50816 | 21,79873275 | 19,01133 | 37,99941 | 28,3433 | 24,2287869 |
| Upregulated | ENSMUSG00000032776 | 9,517278171 | 0 | 29,9556457 | 2,89070227 | 0 | 0 | 1,13589316 | 0 | 0 | 0 | 0 | 15,76529 | 182,319677 | 80,45114 | 66,10908 | 27,34637372 | 49,63421 | 36,66150509 | 100,76 | 158,6642 | 105,4016 | 35,8944991 |
| Upregulated | ENSMUSG00000011254 | 43,35648945 | 52,1244328 | 94,4056714 | 46,2512364 | 68,6994749 | 47,77089187 | 111,31753 | 51,3334929 | 48,0491859 | 61,485701 | 76,1838183 | 65,68873 | 251,440873 | 197,3859 | 152,927 | 120,3240444 | 119,9493 | 121,8747331 | 222,4326 | 279,9956 | 93,00144 | 137,296459 |
| Upregulated | ENSMUSG00000025321 | 0 | 11,7014033 | 7,26197472 | 17,3442136 | 3,8166375 | 2,653938437 | 10,2230384 | 9,77780817 | 1,06775969 | 7,55087556 | 0 | 17,51699 | 59,1036314 | 1,870957 | 28,67382 | 13,67318686 | 26,47158 | 35,6706536 | 22,8136 | 58,66575 | 18,60029 | 0,89736248 |
| Upregulated | ENSMUSG00000022865 | 8,459802819 | 1,06376393 | 0 | 44,3241015 | 6,36106249 | 19,90453828 | 9,08714527 | 25,6667465 | 18,1519147 | 8,62957207 | 8,83290647 | 9,634347 | 97,1703771 | 20,58052 | 16,72639 | 11,85009528 | 13,23579 | 4,954257444 | 37,07209 | 123,9981 | 39,85776 | 17,0498871 |
| Upregulated | ENSMUSG00000023013 | 27,49435916 | 30,8491541 | 30,8633926 | 5,78140454 | 0 | 0 | 0 | 2,44445204 | 2,13551937 | 4,31478604 | 0 | 3,503399 | 156,274008 | 0 | 0 | 0 | 17,37197 | 21,79873275 | 69,39135 | 203,9968 | 39,85776 | 19,7419745 |
| Upregulated | ENSMUSG00000043461 | 0 | 0 | 0 | 0 | 0 | 0 | 0 | 0 | 0 | 0 | 0 | 11,38605 | 154,270496 | 1,870957 | 6,37196 | 0 | 7,445132 | 11,89021787 | 88,40268 | 122,6648 | 1,771456 | 0,89736248 |
| Upregulated | ENSMUSG00000022040 | 106,8050106 | 92,5474623 | 120,73033 | 88,648203 | 35,62195 | 79,61815312 | 65,8818032 | 134,444862 | 88,6240539 | 45,3052534 | 33,1233993 | 13,13775 | 30,0526939 | 0 | 7,964949 | 18,23091581 | 74,45132 | 45,57916849 | 47,52832 | 113,9982 | 114,2589 | 248,569406 |
| Upregulated | ENSMUSG00000009633 | 29,60930986 | 40,4230295 | 20,8781773 | 42,3969667 | 49,6162874 | 92,8878453 | 30,6691153 | 53,777945 | 34,1683099 | 56,0922185 | 52,9974388 | 2,627549 | 7,01229525 | 3,741914 | 3,18598 | 5,469274744 | 43,01632 | 18,82617829 | 30,41813 | 19,33303 | 54,91514 | 255,748306 |
| Upregulated | ENSMUSG00000040564 | 16,91960564 | 9,57387541 | 7,26197472 | 7,70853939 | 1,2722125 | 11,94272297 | 0 | 18,3333903 | 20,287434 | 4,31478604 | 3,31233993 | 0 | 33,0579633 | 0,935478 | 1,59299 | 2,734637372 | 24,81711 | 3,963405955 | 16,15963 | 19,99969 | 87,68707 | 227,930069 |
| Upregulated | ENSMUSG00000012705 | 3,172426057 | 1,06376393 | 0 | 5,78140454 | 0 | 0 | 0 | 0 | 0 | 0 | 0 | 2,627549 | 0 | 0 | 0 | 0 | 26,47158 | 0 | 0 | 2,666625 | 105,4016 | 209,98282 |
| Upregulated | ENSMUSG00000027559 | 88,82792959 | 5,31881967 | 333,14309 | 416,261127 | 8,90548749 | 19,90453828 | 19,3101837 | 42,7779108 | 9,60983717 | 1,07869651 | 4,41645324 | 0,87585 | 1,00175646 | 3,741914 | 0 | 2,734637372 | 1901,818 | 163,4904957 | 261,4058 | 31,33284 | 3284,279 | 9060,66893 |
| Upregulated | ENSMUSG00000022878 | 2,114950705 | 0 | 12,7084558 | 45,2876689 | 0 | 6,634846093 | 9,08714527 | 12,2222602 | 0 | 1,07869651 | 3,31233993 | 0,87585 | 0 | 0 | 0 | 0 | 158,0022 | 27,74384169 | 14,2585 | 0 | 428,6924 | 1328,99383 |
| Upregulated | ENSMUSG00000061780 | 5,287376762 | 9,57387541 | 4,5387342 | 195,604187 | 0 | 54,40573796 | 18,1742905 | 13,4444862 | 2,13551937 | 5,39348255 | 2,20822662 | 4,379248 | 3,00526939 | 0 | 0,796495 | 5,469274744 | 668,4074 | 29,72554466 | 13,30793 | 27,33291 | 2038,946 | 5534,93176 |
| Upregulated | ENSMUSG00000025479 | 33,83921127 | 15,956459 | 35,4021268 | 72,2675568 | 0 | 21,2315075 | 13,6307179 | 72,1113353 | 5,33879843 | 32,3608953 | 23,1863795 | 0 | 10,0175646 | 0 | 0 | 0 | 48,80697 | 14,86277233 | 12,35736 | 7,999875 | 330,3765 | 1679,86256 |
| Upregulated | ENSMUSG00000030278 | 0 | 0 | 0,90774684 | 75,1582591 | 0 | 2,653938437 | 1,13589316 | 2,44445204 | 1,06775969 | 1,07869651 | 1,10411331 | 0,87585 | 2,00351293 | 0 | 0 | 0 | 441,7445 | 25,76213871 | 28,51699 | 15,33309 | 349,8626 | 1491,41644 |
| Upregulated | ENSMUSG00000036395 | 0 | 2,12752787 | 0,90774684 | 60,7047477 | 2,544425 | 5,307876874 | 20,4460768 | 24,4445204 | 1,06775969 | 16,1804476 | 2,20822662 | 22,77209 | 3,00526939 | 8,419306 | 0,796495 | 0,911545791 | 270,5064 | 23,78043573 | 16,15963 | 33,99947 | 259,5183 | 1310,14922 |
| Upregulated | ENSMUSG00000075296 | 4,229901409 | 3,1912918 | 0 | 11,5628091 | 0 | 1,326969219 | 0 | 0 | 1,06775969 | 0 | 0 | 0 | 27,0474245 | 0 | 0 | 0 | 8,272368 | 0 | 2,851699 | 3,999938 | 16,82883 | 99,607235 |
| Upregulated | ENSMUSG00000047786 | 11,63222888 | 10,6376393 | 13,6162026 | 14,4535114 | 5,08884999 | 3,980907656 | 9,08714527 | 9,77780817 | 11,7453565 | 16,1804476 | 22,0822662 | 12,2619 | 21,0368858 | 4,677392 | 7,964949 | 2,734637372 | 11,58132 | 14,86277233 | 7,604532 | 23,99963 | 33,65766 | 90,6336102 |
| Upregulated | ENSMUSG00000022235 | 11,63222888 | 0 | 9,0774684 | 3,8542697 | 1,2722125 | 3,980907656 | 5,67946579 | 9,77780817 | 4,27103874 | 2,15739302 | 7,72879316 | 21,02039 | 17,0298599 | 11,22574 | 7,168455 | 8,203912116 | 5,790658 | 12,88106935 | 8,555098 | 27,99956 | 38,0863 | 68,1995483 |
| Upregulated | ENSMUSG00000019326 | 24,3219331 | 11,7014033 | 98,9444056 | 9,63567424 | 5,08884999 | 2,653938437 | 0 | 26,8889725 | 9,60983717 | 0 | 9,93701978 | 2,627549 | 36,0632327 | 17,77409 | 0 | 0,911545791 | 102,5774 | 25,76213871 | 82,69928 | 214,6633 | 138,1736 | 414,581465 |
| Upregulated | ENSMUSG00000030895 | 13,74717958 | 3,1912918 | 1,81549368 | 5,78140454 | 5,08884999 | 13,26969219 | 5,67946579 | 3,66667807 | 1,06775969 | 1,07869651 | 3,31233993 | 27,15134 | 30,0526939 | 36,48366 | 6,37196 | 12,76164107 | 34,74395 | 0 | 27,56643 | 109,3316 | 124,8877 | 331,126754 |
| Upregulated | ENSMUSG00000062515 | 250,6216585 | 36,1679738 | 589,127699 | 131,04517 | 6,36106249 | 15,92363062 | 15,9025042 | 28,1111985 | 33,1005503 | 6,47217906 | 34,2275126 | 139,2601 | 234,411013 | 21,516 | 27,08083 | 21,87709898 | 879,3528 | 115,9296242 | 303,2307 | 1037,984 | 1635,94 | 4027,3628 |
| Upregulated | ENSMUSG00000038775 | 30,66678522 | 5,31881967 | 22,693671 | 24,0891856 | 6,36106249 | 0 | 6,81535895 | 11,0000342 | 6,40655811 | 20,4952337 | 30,9151726 | 0,87585 | 32,0562069 | 0 | 6,37196 | 4,557728953 | 52,11592 | 38,64320806 | 4,752832 | 71,99888 | 100,973 | 97,81251 |
| Upregulated | ENSMUSG00000037440 | 7,402327466 | 13,8289311 | 4,5387342 | 5,78140454 | 0 | 0 | 9,08714527 | 1,22222602 | 1,06775969 | 0 | 0 | 4,379248 | 1,00175646 | 0 | 6,37196 | 0,911545791 | 9,099605 | 0 | 1,901133 | 20,66634 | 31,00048 | 33,2024117 |
| Upregulated | ENSMUSG00000003526 | 49,70134156 | 41,4867934 | 55,3725572 | 72,2675568 | 26,7164625 | 34,50119968 | 53,3869784 | 61,1113011 | 43,7781471 | 39,9117708 | 25,3946061 | 77,95062 | 63,1106573 | 52,38679 | 80,44599 | 17,31937002 | 69,48789 | 91,15833697 | 63,68796 | 105,9983 | 164,7454 | 173,190958 |
| Upregulated | ENSMUSG00000051920 | 12,68970423 | 38,2955016 | 11,8007089 | 40,4698318 | 0 | 1,326969219 | 0 | 36,6667807 | 9,60983717 | 16,1804476 | 22,0822662 | 7,882647 | 10,0175646 | 0 | 1,59299 | 0,911545791 | 9,099605 | 7,92681191 | 9,505665 | 25,33294 | 43,40067 | 45,7654863 |
| Upregulated | ENSMUSG00000038235 | 12,68970423 | 10,6376393 | 16,3394431 | 70,340422 | 15,26655 | 34,50119968 | 7,95125211 | 36,6667807 | 19,2196743 | 22,6526267 | 6,62467985 | 16,64114 | 44,0772844 | 18,70957 | 2,389485 | 7,292366325 | 9,926842 | 15,85362382 | 12,35736 | 45,33263 | 98,31581 | 82,5573479 |
| Upregulated | ENSMUSG00000078688 | 50,75881691 | 25,5303344 | 204,243039 | 123,33663 | 1,2722125 | 34,50119968 | 52,2510853 | 106,333664 | 10,6775969 | 11,8656616 | 15,4575863 | 7,006798 | 4,00702586 | 0,935478 | 27,08083 | 2,734637372 | 2428,767 | 281,4018228 | 261,4058 | 15,33309 | 1008,844 | 2430,95495 |
| Upregulated | ENSMUSG00000078675 | 13,74717958 | 3,1912918 | 78,0662282 | 30,8341576 | 2,544425 | 6,634846093 | 11,3589316 | 34,2223286 | 2,13551937 | 1,07869651 | 3,31233993 | 0,87585 | 4,00702586 | 0 | 12,74392 | 0 | 708,942 | 64,40534677 | 45,62719 | 0,666656 | 293,176 | 630,845822 |
| Upregulated | ENSMUSG00000096688 | 25,37940846 | 6,3825836 | 147,054988 | 71,3039894 | 2,544425 | 7,961815312 | 28,397329 | 58,666849 | 4,27103874 | 10,7869651 | 12,1452464 | 0,87585 | 0 | 2,806435 | 29,47031 | 1,823091581 | 1235,065 | 125,8381391 | 113,1174 | 4,666594 | 472,093 | 1088,50069 |
| Upregulated | ENSMUSG00000078687 | 5,287376762 | 5,31881967 | 38,1253673 | 19,2713485 | 0 | 1,326969219 | 17,0383974 | 19,5556163 | 0 | 1,07869651 | 1,10411331 | 0 | 1,00175646 | 0 | 3,982475 | 0,911545791 | 415,2729 | 40,62491104 | 49,42946 | 1,333313 | 166,5169 | 632,640547 |
| Upregulated | ENSMUSG00000078673 | 23,26445775 | 9,57387541 | 141,608507 | 80,9396636 | 5,08884999 | 26,53938437 | 40,8921537 | 80,6669174 | 5,33879843 | 15,1017511 | 13,2493597 | 3,503399 | 1,00175646 | 3,741914 | 20,70887 | 0 | 1326,888 | 199,1611493 | 118,8208 | 1,999969 | 512,8365 | 1996,63151 |
| Upregulated | ENSMUSG00000078674 | 16,91960564 | 9,57387541 | 82,6049624 | 65,5225848 | 1,2722125 | 22,55847672 | 19,3101837 | 74,5557873 | 2,13551937 | 1,07869651 | 1,10411331 | 2,627549 | 3,00526939 | 0 | 18,31938 | 2,734637372 | 995,1659 | 127,8198421 | 113,1174 | 1,999969 | 406,5492 | 1432,19051 |
| Upregulated | ENSMUSG00000078689 | 0 | 3,1912918 | 6,35422788 | 4,81783712 | 0 | 0 | 0 | 3,66667807 | 1,06775969 | 0 | 0 | 0 | 0 | 0 | 0 | 0 | 60,38829 | 4,954257444 | 2,851699 | 1,333313 | 29,22902 | 83,4547104 |
| Upregulated | ENSMUSG00000089873 | 13,74717958 | 4,25505574 | 63,5422788 | 38,542697 | 0 | 10,61575375 | 6,81535895 | 52,5557189 | 6,40655811 | 2,15739302 | 2,20822662 | 0,87585 | 1,00175646 | 0 | 11,15093 | 1,823091581 | 739,5497 | 112,9570697 | 61,78682 | 1,333313 | 376,4344 | 1083,11651 |
| Upregulated | ENSMUSG00000078680 | 28,55183451 | 8,51011147 | 147,962735 | 135,863007 | 0 | 22,55847672 | 35,2126879 | 102,666986 | 8,54207749 | 8,62957207 | 14,353473 | 0 | 7,01229525 | 0 | 42,21423 | 4,557728953 | 2073,883 | 250,6854267 | 229,0865 | 11,33316 | 1059,331 | 3050,13506 |
| Upregulated | ENSMUSG00000078686 | 48,64386621 | 18,0839869 | 210,597267 | 79,9760962 | 10,1777 | 10,61575375 | 20,4460768 | 68,4446572 | 11,7453565 | 3,23608953 | 5,52056654 | 0 | 1,00175646 | 1,870957 | 23,09835 | 0,911545791 | 1335,988 | 190,2434859 | 146,3872 | 4,666594 | 571,2946 | 1518,33731 |
| Upregulated | ENSMUSG00000078683 | 20,09203169 | 6,3825836 | 92,5901777 | 31,797725 | 0 | 10,61575375 | 35,2126879 | 25,6667465 | 1,06775969 | 3,23608953 | 1,10411331 | 0 | 1,00175646 | 0,935478 | 15,9299 | 0,911545791 | 732,1046 | 64,40534677 | 49,42946 | 0,666656 | 250,661 | 947,614776 |
| Upregulated | ENSMUSG00000080738 | 3,172426057 | 0 | 9,0774684 | 8,67210682 | 0 | 0 | 2,27178632 | 3,66667807 | 2,13551937 | 3,23608953 | 0 | 0,87585 | 0 | 0 | 0,796495 | 0,911545791 | 93,47776 | 2,972554466 | 4,752832 | 0,666656 | 30,11475 | 111,272947 |
| Upregulated | ENSMUSG00000032554 | 620,7380318 | 601,026623 | 1160,10046 | 967,421694 | 633,561824 | 325,1074586 | 418,008682 | 492,557087 | 497,576014 | 827,360223 | 1027,92949 | 451,9384 | 1778,11772 | 480,8359 | 516,1287 | 539,6351081 | 2695,138 | 1735,971808 | 1069,387 | 1007,318 | 5950,321 | 7035,32182 |
| Upregulated | ENSMUSG00000025911 | 46,5289155 | 44,6780852 | 43,5718483 | 36,6155621 | 30,5331 | 22,55847672 | 5,67946579 | 26,8889725 | 25,6262325 | 38,8330743 | 27,6028327 | 54,30268 | 83,1457866 | 28,99983 | 23,09835 | 23,70019056 | 81,06921 | 13,87192084 | 41,82493 | 57,99909 | 139,945 | 254,850944 |
| Upregulated | ENSMUSG00000020963 | 0 | 0 | 0 | 0 | 0 | 0 | 0 | 0 | 1,06775969 | 0 | 0 | 2,627549 | 0 | 0 | 0 | 0 | 11,58132 | 13,87192084 | 10,45623 | 11,33316 | 23,91466 | 47,5602113 |
| Upregulated | ENSMUSG00000021215 | 77,19570072 | 21,2752787 | 125,269064 | 53,9597757 | 21,6276125 | 6,634846093 | 56,7946579 | 85,5558215 | 66,2011005 | 94,9252928 | 29,8110593 | 178,6733 | 51,0895797 | 25,25792 | 65,31259 | 48,3119269 | 326,7586 | 97,1034459 | 134,9804 | 183,3305 | 377,3201 | 583,28561 |
| Upregulated | ENSMUSG00000114607 | 8,459802819 | 4,25505574 | 15,4316963 | 12,5263765 | 2,544425 | 1,326969219 | 2,27178632 | 1,22222602 | 6,40655811 | 4,31478604 | 0 | 10,5102 | 5,00878232 | 0,935478 | 1,59299 | 4,557728953 | 23,98987 | 4,954257444 | 8,555098 | 19,99969 | 32,77194 | 58,328561 |
| Upregulated | ENSMUSG00000030546 | 1,057475352 | 0 | 23,6014178 | 3,8542697 | 0 | 0 | 0 | 0 | 0 | 0 | 0 | 0 | 0 | 0 | 0 | 0,911545791 | 58,73382 | 2,972554466 | 5,703399 | 1,333313 | 59,34378 | 147,167446 |
| Upregulated | ENSMUSG00000039956 | 1,057475352 | 0 | 7,26197472 | 5,78140454 | 0 | 1,326969219 | 0 | 1,22222602 | 0 | 0 | 1,10411331 | 5,255098 | 8,01405172 | 0,935478 | 2,389485 | 0 | 122,4311 | 17,8353268 | 30,41813 | 33,33281 | 162,974 | 318,563679 |
| Upregulated | ENSMUSG00000051748 | 0 | 0 | 0,90774684 | 0 | 0 | 1,326969219 | 0 | 0 | 0 | 0 | 0 | 0,87585 | 0 | 3,741914 | 4,77897 | 0 | 84,37816 | 2,972554466 | 19,9619 | 19,99969 | 113,3732 | 255,748306 |
| Upregulated | ENSMUSG00000073834 | 21,14950705 | 9,57387541 | 93,4979245 | 49,1419386 | 0 | 3,980907656 | 26,1255426 | 41,5556847 | 5,33879843 | 8,62957207 | 2,20822662 | 0 | 1,00175646 | 0 | 17,52289 | 0,911545791 | 842,9543 | 117,9113272 | 115,9691 | 0 | 464,1215 | 1648,45487 |
| Upregulated | ENSMUSG00000110439 | 32,78173592 | 21,2752787 | 137,069773 | 168,624299 | 1,2722125 | 18,57756906 | 27,2614358 | 105,111438 | 13,8808759 | 10,7869651 | 17,6658129 | 5,255098 | 7,01229525 | 1,870957 | 19,91237 | 0 | 1983,714 | 234,8318028 | 176,8054 | 3,999938 | 1003,53 | 3640,59957 |
| Upregulated | ENSMUSG00000073830 | 26,43688381 | 1,06376393 | 125,269064 | 76,1218265 | 5,08884999 | 22,55847672 | 28,397329 | 70,8891093 | 6,40655811 | 4,31478604 | 3,31233993 | 1,751699 | 0 | 0 | 33,45279 | 0,911545791 | 1308,689 | 184,2983769 | 134,0299 | 4,666594 | 644,81 | 2289,17168 |
| Upregulated | ENSMUSG00000073842 | 9,517278171 | 6,3825836 | 121,638077 | 59,7411803 | 1,2722125 | 9,28878453 | 11,3589316 | 53,777945 | 6,40655811 | 5,39348255 | 3,31233993 | 0,87585 | 4,00702586 | 0 | 9,557939 | 0,911545791 | 965,3854 | 115,9296242 | 96,95778 | 1,999969 | 465,8929 | 2571,84086 |
| Upregulated | ENSMUSG00000096674 | 19,03455634 | 5,31881967 | 82,6049624 | 18,3077811 | 1,2722125 | 11,94272297 | 10,2230384 | 40,3334587 | 1,06775969 | 4,31478604 | 1,10411331 | 0 | 5,00878232 | 0 | 7,168455 | 0 | 663,4439 | 139,7100599 | 51,33059 | 5,33325 | 279,8901 | 1860,23242 |
| Upregulated | ENSMUSG00000094793 | 19,03455634 | 4,25505574 | 54,4648104 | 26,0163204 | 0 | 7,961815312 | 17,0383974 | 25,6667465 | 0 | 1,07869651 | 1,10411331 | 0,87585 | 0 | 0 | 8,761444 | 0,911545791 | 429,3359 | 45,57916849 | 47,52832 | 1,999969 | 166,5169 | 946,717414 |
| Upregulated | ENSMUSG00000023828 | 0 | 0 | 0,90774684 | 7,70853939 | 3,8166375 | 0 | 0 | 4,88890409 | 1,06775969 | 8,62957207 | 0 | 0 | 0 | 0 | 0 | 0 | 45,49803 | 2,972554466 | 5,703399 | 5,999906 | 24,80038 | 131,014922 |
| Upregulated | ENSMUSG00000049422 | 61,33357043 | 61,6983082 | 55,3725572 | 39,5062644 | 36,8941625 | 34,50119968 | 26,1255426 | 18,3333903 | 85,4207749 | 30,2035023 | 40,8521924 | 98,09517 | 204,358319 | 102,9026 | 160,0955 | 184,1322497 | 206,8092 | 77,28641613 | 93,15552 | 98,66513 | 262,1755 | 620,077472 |
| Upregulated | ENSMUSG00000018796 | 176,5983838 | 182,967397 | 236,014178 | 246,673261 | 239,17595 | 208,3341673 | 266,934892 | 200,445068 | 169,77379 | 244,864108 | 228,551455 | 343,3331 | 277,486541 | 289,9983 | 418,9563 | 333,6257594 | 540,1857 | 267,529902 | 325,0937 | 439,9931 | 505,7507 | 950,306864 |
| Upregulated | ENSMUSG00000033854 | 7,402327466 | 1,06376393 | 6,35422788 | 0 | 0 | 0 | 0 | 0 | 0 | 7,55087556 | 2,20822662 | 26,27549 | 16,0281034 | 16,83861 | 90,00393 | 20,05400739 | 186,9555 | 58,46023784 | 190,1133 | 57,33244 | 49,60077 | 258,440394 |
| Upregulated | ENSMUSG00000050578 | 29,60930986 | 11,7014033 | 40,8486078 | 3,8542697 | 29,2608875 | 6,634846093 | 26,1255426 | 31,7778766 | 55,5235037 | 30,2035023 | 60,726232 | 1320,781 | 327,574364 | 746,5118 | 861,011 | 597,0624929 | 4831,89 | 1397,100599 | 5670,129 | 2769,29 | 396,8062 | 10434,5309 |
| Upregulated | ENSMUSG00000025348 | 84,59802819 | 43,6143213 | 83,5127093 | 48,1783712 | 8,90548749 | 70,32936859 | 37,4844742 | 28,1111985 | 24,5584728 | 93,8465963 | 59,6221187 | 152,3978 | 114,200237 | 65,48349 | 103,5443 | 72,92366325 | 361,5025 | 146,6460203 | 306,0824 | 283,9956 | 220,5463 | 666,740321 |
| Upregulated | ENSMUSG00000045502 | 142,7591726 | 19,1477508 | 81,6972156 | 33,7248598 | 7,63327499 | 33,17423047 | 14,7666111 | 3,66667807 | 14,9486356 | 24,8100197 | 28,706946 | 102,4744 | 87,1528124 | 211,4181 | 96,37589 | 105,7393117 | 237,417 | 95,12174293 | 207,2235 | 116,6648 | 107,1731 | 589,567148 |
| Upregulated | ENSMUSG00000110344 | 0 | 10,6376393 | 9,98521524 | 21,1984833 | 7,63327499 | 18,57756906 | 10,2230384 | 7,33335613 | 7,4743178 | 18,3378407 | 27,6028327 | 20,14454 | 12,0210776 | 25,25792 | 32,65629 | 29,1694653 | 40,53461 | 28,73469318 | 42,77549 | 42,666 | 22,1432 | 115,75976 |

| | | | | | | | | | | | | | | | | | | | | | | | |
|---|---|---|---|---|---|---|---|---|---|---|---|---|---|---|---|---|---|---|---|---|---|---|---|
| Upregulated | ENSMUSG00000019577 | 236,8744789 | 111,695213 | 128,900051 | 33,7248598 | 139,943375 | 127,389045 | 139,714858 | 68,4446572 | 76,8786974 | 104,633561 | 107,098991 | 105,102 | 112,196724 | 46,77392 | 56,55114 | 52,86965586 | 309,3866 | 53,5059804 | 134,9804 | 151,331 | 98,31581 | 575,209348 |
| Upregulated | ENSMUSG00000024892 | 652,4622924 | 345,723279 | 551,910079 | 428,787504 | 228,99825 | 201,6993212 | 228,314525 | 310,44541 | 337,412061 | 319,294167 | 316,88052 | 494,8551 | 505,887015 | 400,3848 | 341,6963 | 430,2496132 | 886,7979 | 476,5995661 | 607,412 | 999,9844 | 597,8664 | 1543,46346 |
| Upregulated | ENSMUSG00000022636 | 54,98871832 | 12,7651672 | 25,4169115 | 32,7612924 | 16,5387625 | 19,90453828 | 13,6307179 | 29,3334245 | 27,7617518 | 0 | 34,2275126 | 9,634347 | 41,072015 | 48,64488 | 24,69134 | 22,78864477 | 76,10579 | 39,63405955 | 26,61586 | 55,33247 | 49,60077 | 127,425472 |
| Upregulated | ENSMUSG00000000440 | 173,4259578 | 47,869377 | 205,150786 | 42,3969667 | 16,5387625 | 17,25059984 | 10,2230384 | 47,6668148 | 21,3551937 | 46,3839499 | 32,019286 | 35,03399 | 131,230097 | 50,51583 | 6,37196 | 16,40782423 | 104,2318 | 83,23152506 | 105,5129 | 106,665 | 76,17261 | 243,185231 |
| Upregulated | ENSMUSG00000002944 | 471,6340071 | 111,695213 | 996,70603 | 162,842895 | 3,8166375 | 10,61575375 | 24,9896495 | 119,77815 | 67,2688602 | 11,8656616 | 140,22239 | 119,1156 | 239,419795 | 0,935478 | 54,95815 | 3,646183163 | 521,9864 | 120,8838816 | 506,6519 | 105,3317 | 565,0945 | 1491,41644 |
| Upregulated | ENSMUSG00000109866 | 29,60930986 | 37,2317377 | 19,0626836 | 52,0326409 | 6,36106249 | 0 | 5,67946579 | 13,4444862 | 1,06775969 | 17,2591441 | 11,0411331 | 11,38605 | 31,0544504 | 33,67722 | 26,28433 | 15,49627844 | 52,11592 | 39,63405955 | 27,56643 | 13,99978 | 16,82883 | 67,3021858 |
| Upregulated | ENSMUSG00000094443 | 27,49435916 | 9,57387541 | 8,16972156 | 40,4698318 | 8,90548749 | 6,634846093 | 1,13589316 | 9,77780817 | 7,4743178 | 0 | 4,41645324 | 8,758497 | 24,0421551 | 13,0967 | 10,35443 | 7,292366325 | 19,85368 | 18,82617829 | 34,22039 | 1,999969 | 21,25747 | 44,8681239 |
| Upregulated | ENSMUSG00000115667 | 14,80465493 | 5,31881967 | 9,98521524 | 0,96356742 | 3,8166375 | 6,634846093 | 2,27178632 | 2,44445204 | 13,8808759 | 1,07869651 | 11,0411331 | 4,379248 | 18,0316164 | 0,935478 | 8,761444 | 7,292366325 | 35,57118 | 21,79873275 | 7,604532 | 10,6665 | 6,200096 | 16,1525246 |
| Upregulated | ENSMUSG00000046280 | 15,86213028 | 10,6376393 | 16,3394431 | 10,5992417 | 1,2722125 | 2,653938437 | 2,27178632 | 6,11113011 | 0 | 1,07869651 | 1,10411331 | 19,26869 | 2,00351293 | 2,806435 | 3,18598 | 0 | 32,26224 | 6,935960422 | 9,505665 | 4,666594 | 0 | 10,7683497 |
| Upregulated | ENSMUSG00000085363 | 29,60930986 | 29,7853902 | 39,0331141 | 8,67210682 | 7,63327499 | 7,961815312 | 29,5332221 | 17,1111643 | 9,60983717 | 94,9252928 | 37,5398525 | 31,53059 | 23,0403987 | 20,58052 | 20,70887 | 31,90410267 | 168,7563 | 16,84447531 | 77,94645 | 23,33297 | 17,71456 | 43,0733989 |
| Upregulated | ENSMUSG00000004609 | 10,57475352 | 3,1912918 | 5,44648104 | 0 | 0 | 10,61575375 | 0 | 1,22222602 | 0 | 0 | 0 | 10,5102 | 4,00702586 | 18,70957 | 18,31938 | 1,823091581 | 78,5875 | 5,945108933 | 24,71473 | 11,99981 | 3,542912 | 9,87098725 |
| Upregulated | ENSMUSG00000047048 | 12,68970423 | 12,7651672 | 11,8007089 | 1,92713485 | 0 | 0 | 0 | 0 | 1,06775969 | 0 | 0 | 25,39964 | 23,0403987 | 0 | 11,15093 | 3,646183163 | 82,72368 | 4,954257444 | 25,6653 | 35,33278 | 10,62874 | 12,5630747 |
| Upregulated | ENSMUSG00000096971 | 19,03455634 | 10,6376393 | 18,1549368 | 7,70853939 | 2,544425 | 7,961815312 | 5,67946579 | 6,11113011 | 2,13551937 | 8,62957207 | 1,10411331 | 3,503399 | 23,0403987 | 9,354784 | 10,35443 | 11,85009528 | 59,56105 | 14,86277233 | 26,61586 | 22,66631 | 21,25747 | 9,87098725 |
| Upregulated | ENSMUSG00000020140 | 881,9344438 | 1188,22431 | 1246,33641 | 1272,87257 | 259,53135 | 252,1241515 | 490,705844 | 673,446538 | 332,073262 | 869,429386 | 351,108032 | 849,5742 | 323,567338 | 1059,897 | 543,2096 | 972,6193586 | 4598,61 | 4238,862669 | 2658,734 | 645,3233 | 297,6046 | 1307,45713 |
| Upregulated | ENSMUSG00000038871 | 581,6114438 | 634,003305 | 813,341169 | 1428,00692 | 442,72995 | 391,4559195 | 767,863775 | 508,446025 | 326,734464 | 814,415864 | 491,330422 | 731,3345 | 249,43736 | 818,5436 | 1671,843 | 2399,188521 | 5563,168 | 2374,080167 | 2027,558 | 1410,645 | 567,7517 | 1675,37575 |
| Upregulated | ENSMUSG00000031673 | 2639,458479 | 2401,97896 | 3449,43799 | 4188,62759 | 1097,91939 | 862,5299921 | 1762,90618 | 1940,89492 | 1639,01112 | 2729,10217 | 2016,1109 | 1948,766 | 2053,60075 | 3310,658 | 2158,501 | 3215,022004 | 4484,451 | 4545,035779 | 3651,126 | 3493,279 | 1697,941 | 3500,61102 |
| Upregulated | ENSMUSG00000063594 | 35,95416198 | 26,5940983 | 62,634532 | 59,7411803 | 21,6276125 | 15,92363062 | 26,1255426 | 41,5556847 | 25,6262325 | 22,6526267 | 28,706946 | 34,15814 | 10,0175646 | 33,67722 | 34,24928 | 32,81564846 | 81,06921 | 52,51512891 | 57,98456 | 47,33259 | 30,11475 | 57,4311986 |
| Upregulated | ENSMUSG00000001021 | 11,63222888 | 19,1477508 | 8,16972156 | 5,78140454 | 6,36106249 | 5,307876874 | 2,27178632 | 14,6667123 | 9,60983717 | 6,47217906 | 7,72879316 | 15,76529 | 10,0175646 | 6,548349 | 8,761444 | 7,292366325 | 28,12605 | 20,80788127 | 25,6653 | 3,999938 | 6,200096 | 17,0498871 |
| Upregulated | ENSMUSG00000011148 | 62,39104579 | 75,5272393 | 72,6197472 | 30,8341576 | 64,8828374 | 35,8281689 | 32,9409016 | 34,2223286 | 57,659023 | 57,170915 | 37,5398525 | 29,77889 | 6,01053879 | 30,87079 | 44,60372 | 67,45438851 | 196,8824 | 107,0119608 | 221,482 | 20,66634 | 8,85728 | 84,3520729 |
| Upregulated | ENSMUSG00000027107 | 565,7493135 | 719,104419 | 753,429877 | 438,423178 | 298,969937 | 66,34846093 | 366,89349 | 195,556163 | 139,876519 | 352,733759 | 167,825223 | 417,7803 | 74,1299784 | 231,9986 | 722,4209 | 504,996368 | 2184,733 | 792,681191 | 1838,396 | 75,99881 | 64,65815 | 852,494354 |
| Upregulated | ENSMUSG00000039601 | 219,9548733 | 509,542924 | 269,600811 | 109,846686 | 68,6994749 | 66,34846093 | 61,3382305 | 162,556061 | 41,6426277 | 126,207492 | 192,115716 | 195,3145 | 546,95903 | 217,031 | 66,90558 | 75,65830062 | 732,9318 | 213,0330701 | 656,8414 | 518,6586 | 169,1741 | 401,121027 |
| Upregulated | ENSMUSG00000019303 | 20,09203169 | 79,782295 | 97,1289119 | 59,7411803 | 58,5217749 | 43,78998421 | 40,8921537 | 20,7778424 | 33,1005503 | 25,8887162 | 19,8740396 | 11,38605 | 150,26347 | 57,99966 | 44,60372 | 41,93110637 | 191,9189 | 74,31386166 | 169,2008 | 98,66513 | 31,88621 | 145,372721 |
| Upregulated | ENSMUSG00000031488 | 48,64386621 | 37,2317377 | 36,3098736 | 27,9434553 | 39,4385875 | 14,5966614 | 28,397329 | 30,5556505 | 29,8972712 | 53,9348255 | 79,4961582 | 137,5084 | 42,0737715 | 45,83844 | 66,90558 | 51,04656428 | 193,5734 | 77,28641613 | 188,2122 | 60,66572 | 13,28592 | 148,064809 |
| Upregulated | ENSMUSG00000054905 | 9,517278171 | 6,3825836 | 11,8007089 | 2,89070227 | 0 | 5,307876874 | 2,27178632 | 12,2222602 | 5,33879843 | 1,07869651 | 0 | 28,90304 | 2,00351293 | 27,12887 | 31,0633 | 25,52328214 | 81,89645 | 12,88106935 | 102,6612 | 15,99975 | 17,71456 | 36,7918616 |
| Upregulated | ENSMUSG00000042284 | 295,0356233 | 462,737311 | 346,759293 | 188,859215 | 71,2438999 | 236,2005209 | 149,937897 | 342,223286 | 256,262325 | 418,534246 | 210,885642 | 352,9674 | 422,741228 | 371,3849 | 754,2807 | 338,1834883 | 937,2593 | 457,7733878 | 1173,95 | 633,9901 | 296,7189 | 709,81372 |
| Upregulated | ENSMUSG00000047344 | 4,229901409 | 6,3825836 | 1,81549368 | 5,78140454 | 1,2722125 | 0 | 0 | 0 | 2,13551937 | 11,8656616 | 13,2493597 | 6,130948 | 13,022834 | 14,03218 | 14,33691 | 7,292366325 | 31,435 | 4,954257444 | 41,82493 | 9,333188 | 4,42864 | 17,9472495 |
| Upregulated | ENSMUSG00000031927 | 21,14950705 | 0 | 6,35422788 | 0,96356742 | 0 | 0 | 0 | 0 | 0 | 0 | 0 | 0 | 0 | 9,354784 | 3,982475 | 3,646183163 | 33,08947 | 18,82617829 | 39,92379 | 7,999875 | 1,771456 | 44,8681239 |
| Upregulated | ENSMUSG00000050737 | 3081,483177 | 2052,00063 | 3037,32093 | 2662,33679 | 2576,23031 | 1958,606567 | 2700,01804 | 3532,2332 | 3212,88889 | 2486,39545 | 2658,70485 | 8855,716 | 4019,04694 | 4272,33 | 3205,892 | 3730,045375 | 11908,07 | 4495,493205 | 18784,14 | 9570,517 | 6826,306 | 13656,9595 |
| Upregulated | ENSMUSG00000070880 | 0 | 0 | 1,81549368 | 0 | 3,8166375 | 3,980907656 | 10,2230384 | 0 | 0 | 0 | 5,52056654 | 22,77209 | 0 | 0 | 36,63877 | 32,81564846 | 70,31513 | 35,6706536 | 158,7446 | 43,99931 | 3,542912 | 92,4283352 |
| Upregulated | ENSMUSG00000031530 | 97,28773241 | 78,7185311 | 97,1289119 | 71,3039894 | 69,9716874 | 41,13604578 | 76,1048416 | 42,7779108 | 76,8786974 | 53,9348255 | 146,84707 | 229,4726 | 29,0509375 | 86,06401 | 185,5833 | 224,2402645 | 509,5779 | 173,3990105 | 454,3708 | 392,6605 | 259,5183 | 426,247177 |
| Upregulated | ENSMUSG00000045349 | 2,114950705 | 6,3825836 | 1,81549368 | 0,96356742 | 0 | 2,653938437 | 1,13589316 | 8,55558215 | 4,27103874 | 0 | 1,10411331 | 0,87585 | 2,00351293 | 0 | 5,575465 | 4,557728953 | 31,435 | 12,88106935 | 31,36869 | 10,6665 | 11,51446 | 17,0498871 |
| Upregulated | ENSMUSG00000055409 | 12,68970423 | 34,0404459 | 24,5091647 | 45,2876689 | 49,6162874 | 15,92363062 | 22,7178632 | 187,000581 | 27,7617518 | 28,0461092 | 32,019286 | 26,27549 | 32,0562069 | 21,516 | 29,47031 | 4,557728953 | 136,4941 | 159,5270897 | 37,07209 | 25,99959 | 19,48602 | 54,7391111 |
| Upregulated | ENSMUSG00000032036 | 12,68970423 | 32,976682 | 25,4169115 | 20,2349159 | 0 | 0 | 15,9025042 | 44,0001368 | 29,8972712 | 3,23608953 | 17,6658129 | 11,38605 | 19,0333728 | 10,29026 | 10,35443 | 11,85009528 | 49,63421 | 51,52427742 | 18,06076 | 11,99981 | 4,42864 | 20,639337 |
| Upregulated | ENSMUSG00000115243 | 0 | 4,25505574 | 0 | 15,4170788 | 3,8166375 | 1,326969219 | 1,13589316 | 19,5556163 | 23,4907131 | 0 | 2,20822662 | 1,751699 | 2,00351293 | 0 | 0,796495 | 1,823091581 | 39,70737 | 1,981702978 | 29,46756 | 30,66619 | 3,542912 | 2,69208743 |
| Upregulated | ENSMUSG00000068762 | 3,172426057 | 13,8289311 | 15,4316963 | 9,63567424 | 11,4499125 | 6,634846093 | 10,2230384 | 14,6667123 | 11,7453565 | 2,15739302 | 5,52056654 | 3,503399 | 4,00702586 | 0 | 0 | 0 | 23,98987 | 6,935960422 | 14,2585 | 7,333219 | 20,37174 | 11,6657122 |
| Upregulated | ENSMUSG00000029088 | 11,63222888 | 8,51011147 | 5,44648104 | 12,5263765 | 3,8166375 | 6,634846093 | 5,67946579 | 23,2222944 | 11,7453565 | 17,2591441 | 18,7699262 | 7,006798 | 22,0386422 | 5,61287 | 12,74392 | 0 | 57,90658 | 17,8353268 | 33,26983 | 17,33306 | 29,22902 | 15,2551621 |
| Upregulated | ENSMUSG00000045691 | 23,26445775 | 38,2955016 | 47,2028357 | 45,2876689 | 5,08884999 | 9,28878453 | 29,5332221 | 35,4445546 | 26,6939921 | 52,8561289 | 19,8740396 | 21,89624 | 23,0403987 | 11,22574 | 31,8598 | 34,63874004 | 95,95947 | 36,66150509 | 46,57776 | 45,33263 | 59,34378 | 46,6628488 |
| Upregulated | ENSMUSG00000055235 | 10,57475352 | 7,44634754 | 10,8929621 | 48,1783712 | 2,544425 | 7,961815312 | 3,40767947 | 14,6667123 | 3,20327906 | 5,39348255 | 16,5616996 | 38,53739 | 18,0316164 | 0 | 21,50536 | 12,76164107 | 52,94316 | 18,82617829 | 46,57776 | 29,33288 | 27,45757 | 8,97362477 |
| Upregulated | ENSMUSG00000028687 | 24,3219331 | 12,7651672 | 30,8633926 | 46,2512364 | 11,4499125 | 7,961815312 | 9,08714527 | 22,0000684 | 5,33879843 | 30,2035023 | 11,0411331 | 20,14454 | 15,026347 | 0,935478 | 24,69134 | 15,49627844 | 33,08947 | 20,80788127 | 38,02266 | 27,99956 | 15,05738 | 13,4604372 |
| Upregulated | ENSMUSG00000040016 | 614,3931797 | 403,166531 | 572,788256 | 756,400428 | 153,937712 | 348,9929045 | 271,478465 | 336,112156 | 257,330084 | 340,868097 | 280,44478 | 178,6733 | 139,244149 | 157,1604 | 100,3584 | 190,5130702 | 972,0033 | 1087,954935 | 647,3358 | 119,3315 | 325,0622 | 993,380263 |
| Upregulated | ENSMUSG00000020044 | 5389,951871 | 5451,79016 | 5150,55557 | 6538,76854 | 3214,88098 | 3168,802494 | 2998,75794 | 4752,01477 | 4820,93498 | 4222,01814 | 2413,59169 | 1616,819 | 2012,52874 | 469,6102 | 1401,831 | 1764,752651 | 7626,296 | 8354,859754 | 3851,695 | 2867,955 | 4955,648 | 4663,5928 |
| Upregulated | ENSMUSG00000020303 | 488,5536128 | 509,542924 | 170,656406 | 528,034948 | 147,57665 | 143,3126756 | 149,937897 | 80,6669174 | 83,2852555 | 242,706715 | 227,447342 | 232,1002 | 206,361832 | 274,0952 | 271,6048 | 331,8026678 | 638,6268 | 626,2181409 | 462,9259 | 177,3306 | 288,7473 | 378,686965 |
| Upregulated | ENSMUSG00000046031 | 6,344852114 | 6,3825836 | 13,6162026 | 7,70853939 | 1,2722125 | 1,326969219 | 1,13589316 | 31,7778766 | 20,287434 | 29,1248057 | 36,4357392 | 4,379248 | 10,0175646 | 5,61287 | 3,982475 | 3,646183163 | 16,54474 | 13,87192084 | 26,61586 | 16,66641 | 18,60029 | 19,7419745 |
| Upregulated | ENSMUSG00000022456 | 3,172426057 | 6,3825836 | 11,8007089 | 5,78140454 | 0 | 5,307876874 | 1,13589316 | 12,2222602 | 9,60983717 | 15,1017511 | 29,8110593 | 3,503399 | 5,00878232 | 0 | 14,33691 | 10,0270037 | 11,58132 | 14,86277233 | 39,92379 | 27,33291 | 19,48602 | 3,58944991 |
| Upregulated | ENSMUSG00000074569 | 5,287376762 | 1,06376393 | 1,81549368 | 1,92713485 | 11,4499125 | 17,25059984 | 3,40767947 | 14,6667123 | 1,06775969 | 10,7869651 | 22,0822662 | 4,379248 | 14,0245905 | 11,22574 | 6,37196 | 15,49627844 | 9,099605 | 17,8353268 | 12,35736 | 7,333219 | 4,42864 | 9,87098725 |
| Upregulated | ENSMUSG00000038132 | 50,75881691 | 29,7853902 | 64,4500256 | 42,3969667 | 94,1437249 | 195,0644751 | 61,3382305 | 114,889246 | 100,36941 | 102,476168 | 324,609313 | 57,80608 | 38,0667457 | 116,9348 | 196,7343 | 118,5009528 | 57,07934 | 126,8289906 | 73,19362 | 15,99975 | 76,17261 | 45,7654863 |
| Upregulated | ENSMUSG00000060183 | 0 | 3,1912918 | 12,7084558 | 4,81783712 | 24,1720375 | 23,88544594 | 0 | 7,33335613 | 12,8131162 | 22,6526267 | 51,8933255 | 12,2619 | 0 | 14,03218 | 3,18598 | 0 | 47,97974 | 10,89936638 | 17,1102 | 0 | 0 | 2,69208743 |
| Upregulated | ENSMUSG00000031549 | 0 | 5,31881967 | 12,7084558 | 2,89070227 | 6,36106249 | 10,61575375 | 14,7666111 | 1,22222602 | 5,33879843 | 5,39348255 | 83,9126115 | 45,54418 | 5,00878232 | 24,32244 | 8,761444 | 31,90410267 | 28,12605 | 24,77128722 | 20,91246 | 19,33303 | 9,743008 | 20,639337 |
| Upregulated | ENSMUSG00000102813 | 7,402327466 | 0 | 6,35422788 | 0 | 5,08884999 | 0 | 2,27178632 | 2,44445204 | 6,40655811 | 6,47217906 | 51,8933255 | 38,53739 | 2,00351293 | 0 | 6,37196 | 10,93854949 | 36,39842 | 28,73469318 | 7,604532 | 6,666563 | 0,885728 | 17,9472495 |
| Upregulated | ENSMUSG00000103085 | 1,057475352 | 1,06376393 | 0,90774684 | 1,92713485 | 6,36106249 | 25,21241515 | 6,81535895 | 0 | 3,20327906 | 5,39348255 | 6,62467985 | 5,255098 | 30,0526939 | 2,806435 | 8,761444 | 10,0270037 | 17,37197 | 10,89936638 | 13,30793 | 5,33325 | 1,771456 | 7,17889982 |
| Upregulated | ENSMUSG00000103220 | 81,42560213 | 17,0202229 | 30,8633926 | 68,4132871 | 96,6881499 | 131,3699526 | 79,5125211 | 79,4446914 | 106,775969 | 55,013522 | 147,951183 | 65,68873 | 162,284547 | 52,38679 | 74,07403 | 77,4813922 | 101,7501 | 114,9387727 | 98,85892 | 5,33325 | 17,71456 | 48,4575738 |
| Upregulated | ENSMUSG00000050141 | 5,287376762 | 2,12752787 | 5,44648104 | 4,81783712 | 10,1777 | 15,92363062 | 4,54357263 | 25,6667465 | 12,8131162 | 2,15739302 | 7,72879316 | 12,2619 | 27,0474245 | 8,419306 | 24,69134 | 10,0270037 | 9,926842 | 13,87192084 | 8,555098 | 9,333188 | 20,37174 | 6,28153734 |
| Upregulated | ENSMUSG00000104524 | 3,172426057 | 6,3825836 | 0 | 4,81783712 | 6,36106249 | 25,21241515 | 1,13589316 | 26,8889725 | 7,4743178 | 2,15739302 | 4,41645324 | 7,882647 | 31,0544504 | 25,25792 | 7,168455 | 13,67318686 | 0,827237 | 6,935960422 | 10,45623 | 11,33316 | 3,542912 | 8,0762623 |
| Upregulated | ENSMUSG00000078894 | 0 | 14,8926951 | 6,35422788 | 22,1620508 | 6,36106249 | 70,32936859 | 39,7562605 | 79,4446914 | 61,9300618 | 63,643094 | 45,2686457 | 24,52379 | 107,187942 | 0,935478 | 51,77217 | 73,83520904 | 35,57118 | 46,57001997 | 71,29249 | 91,33191 | 45,17213 | 66,4048233 |
| Upregulated | ENSMUSG00000078877 | 5,287376762 | 23,4028065 | 6,35422788 | 26,0163204 | 12,722125 | 61,04058406 | 54,5228716 | 89,2224996 | 61,9300618 | 43,1478604 | 68,4550251 | 43,79248 | 105,184429 | 8,419306 | 74,07403 | 84,77375853 | 62,04276 | 62,4236438 | 94,10608 | 91,99856 | 77,94407 | 79,8652605 |
| Upregulated | ENSMUSG00000084853 | 17,97708099 | 5,31881967 | 17,24719 | 17,3442136 | 80,1493874 | 78,2911839 | 26,1255426 | 94,1114037 | 140,944279 | 4,31478604 | 2,20822662 | 45,54418 | 207,363588 | 14,96765 | 68,49857 | 33,72719425 | 57,90658 | 57,46938635 | 85,55098 | 48,66591 | 46,94358 | 84,3520729 |
| Upregulated | ENSMUSG00000047230 | 4,229901409 | 0 | 1,81549368 | 0,96356742 | 0 | 0 | 0 | 13,4444862 | 0 | 8,62957207 | 0 | 0 | 77,1352478 | 0 | 0 | 0 | 10,75408 | 0,990851489 | 13,30793 | 7,333219 | 14,17165 | 20,639337 |
| Upregulated | ENSMUSG00000036225 | 7,402327466 | 6,3825836 | 19,9704305 | 13,4899439 | 17,810975 | 30,52029203 | 6,81535895 | 13,4444862 | 26,6939921 | 9,70826858 | 25,3946061 | 10,5102 | 85,1492995 | 84,19306 | 40,62124 | 27,34637372 | 13,23579 | 7,92681191 | 15,20906 | 7,333219 | 0,885728 | 7,17889982 |
| Upregulated | ENSMUSG00000087201 | 8,459802819 | 6,3825836 | 0 | 5,78140454 | 3,8166375 | 3,980907656 | 7,95125211 | 1,22222602 | 14,9486356 | 1,07869651 | 5,52056654 | 9,634347 | 46,0807974 | 34,6127 | 8,761444 | 14,58473265 | 0 | 11,89021787 | 0,950566 | 0,666656 | 7,085824 | 12,5630747 |
| Upregulated | ENSMUSG00000031757 | 103,6325845 | 112,758977 | 47,2028357 | 49,1419386 | 173,0209 | 45,11695343 | 118,132888 | 47,6668148 | 66,2011005 | 168,276655 | 200,948622 | 183,0526 | 307,539235 | 224,5148 | 242,931 | 170,4590628 | 34,74395 | 129,801545 | 57,03399 | 11,33316 | 77,94407 | 144,475359 |
| Upregulated | ENSMUSG00000108693 | 10,57475352 | 4,25505574 | 3,63098736 | 1,92713485 | 8,90548749 | 26,53938437 | 14,7666111 | 12,2222602 | 13,8808759 | 17,2591441 | 34,2275126 | 22,77209 | 45,0790409 | 20,58052 | 25,48784 | 9,115457906 | 6,617895 | 13,87192084 | 3,802266 | 4,666594 | 6,200096 | 12,5630747 |
| Upregulated | ENSMUSG00000040972 | 25,37940846 | 81,9098229 | 83,5127093 | 108,883119 | 33,077525 | 75,63724546 | 90,8714527 | 56,222397 | 36,3038293 | 61,485701 | 157,888203 | 43,79248 | 128,224827 | 58,93514 | 120,2707 | 60,16202218 | 8,272368 | 13,87192084 | 37,07209 | 0 | 0 | 11,6657122 |
| Upregulated | ENSMUSG00000074227 | 2,114950705 | 9,57387541 | 2,72324052 | 32,7612924 | 2,544425 | 11,94272297 | 10,2230384 | 2,44445204 | 19,2196743 | 7,55087556 | 25,3946061 | 9,634347 | 41,072015 | 17,77409 | 0 | 4,557728953 | 2,481711 | 6,935960422 | 0 | 0 | 2,657184 | 1,79472495 |
| Upregulated | ENSMUSG00000113750 | 6,344852114 | 12,7651672 | 9,0774684 | 5,78140454 | 22,899825 | 11,94272297 | 23,8537563 | 22,0000684 | 21,3551937 | 4,31478604 | 15,4575863 | 28,02719 | 21,0368858 | 26,19339 | 27,08083 | 21,87709898 | 15,7175 | 19,81702978 | 19,01133 | 11,33316 | 23,02893 | 17,9472495 |
| Upregulated | ENSMUSG00000038859 | 62,39104579 | 41,4867934 | 52,6493167 | 55,8869106 | 208,64285 | 66,34846093 | 289,652755 | 188,222807 | 142,012038 | 131,600974 | 164,512883 | 299,5406 | 339,595441 | 154,3539 | 68,49857 | 53,78120165 | 119,1221 | 153,5819808 | 163,4974 | 123,3314 | 265,7184 | 176,780408 |
| Upregulated | ENSMUSG00000031170 | 12,68970423 | 26,5940983 | 2,72324052 | 34,6884273 | 105,593637 | 35,8281689 | 76,1048416 | 154,000479 | 34,1683099 | 90,6105068 | 56,3097787 | 67,44043 | 73,1282219 | 72,96731 | 27,08083 | 51,95811007 | 9,926842 | 77,28641613 | 23,76416 | 11,33316 | 97,43008 | 161,525246 |
| Upregulated | ENSMUSG00000104002 | 2,114950705 | 1,06376393 | 0,90774684 | 1,92713485 | 7,63327499 | 11,94272297 | 18,1742905 | 3,66667807 | 14,9486356 | 11,8656616 | 2,20822662 | 18,39284 | 9,01580818 | 5,61287 | 15,1334 | 10,93854949 | 4,963421 | 13,87192084 | 14,2585 | 8,666531 | 1,771456 | 6,28153734 |
| Upregulated | ENSMUSG00000095432 | 17,97708099 | 27,6578623 | 2,72324052 | 33,7248598 | 55,9773499 | 33,17423047 | 46,5716195 | 52,5557189 | 113,182527 | 49,6200394 | 82,8084982 | 105,102 | 93,1633512 | 13,0967 | 101,1549 | 106,6508575 | 56,25211 | 94,13089144 | 79,84759 | 9,333188 | 38,97203 | 36,7918616 |
| Upregulated | ENSMUSG00000032020 | 45,47144015 | 120,205325 | 120,73033 | 76,1218265 | 264,6202 | 289,2792897 | 243,081136 | 294,556471 | 296,837193 | 242,706715 | 291,485914 | 369,6086 | 239,419795 | 155,2894 | 280,3662 | 399,2570563 | 166,2746 | 200,1520007 | 139,7333 | 162,6641 | 159,431 | 222,545894 |
| Upregulated | ENSMUSG00000117113 | 1,057475352 | 4,25505574 | 1,81549368 | 9,63567424 | 19,0831875 | 29,19332281 | 22,7178632 | 18,3333903 | 18,1519147 | 2,15739302 | 16,5616996 | 31,53059 | 20,0351293 | 5,61287 | 16,72639 | 13,67318686 | 8,272368 | 19,81702978 | 6,653965 | 16,66641 | 7,971552 | 20,639337 |
| Upregulated | ENSMUSG00000102856 | 8,459802819 | 0 | 2,72324052 | 7,70853939 | 24,1720375 | 29,19332281 | 28,397329 | 25,6667465 | 10,6775969 | 4,31478604 | 20,9781529 | 18,39284 | 31,0544504 | 23,38696 | 15,1334 | 12,76164107 | 28,95329 | 51,52427742 | 4,752832 | 1,333313 | 3,542912 | 20,639337 |
| Upregulated | ENSMUSG00000037686 | 912,601229 | 299,981429 | 514,692458 | 192,713485 | 912,176362 | 1145,174436 | 761,048416 | 451,001402 | 660,943245 | 476,783857 | 833,605548 | 1255,093 | 584,024019 | 1024,349 | 523,2972 | 522,315738 | 535,2222 | 1210,820519 | 418,2493 | 57,99909 | 76,17261 | 264,721931 |
| Upregulated | ENSMUSG00000021983 | 0 | 6,3825836 | 5,44648104 | 7,70853939 | 6,36106249 | 26,53938437 | 4,54357263 | 6,11113011 | 5,33879843 | 3,23608953 | 15,4575863 | 13,13775 | 16,0281034 | 12,16122 | 0,796495 | 0 | 0 | 1,981702978 | 0 | 13,99978 | 12,40019 | 7,17889982 |
| Upregulated | ENSMUSG00000100967 | 0 | 0 | 3,63098736 | 3,8542697 | 0 | 15,92363062 | 0 | 13,4444862 | 12,8131162 | 2,15739302 | 8,83290647 | 28,02719 | 2,00351293 | 14,03218 | 7,964949 | 8,203912116 | 9,099605 | 1,981702978 | 2,851699 | 20,66634 | 1,771456 | 3,58944991 |
| Upregulated | ENSMUSG00000108218 | 30,66678522 | 23,4028065 | 15,4316963 | 18,3077811 | 59,7939874 | 91,56087608 | 15,9025042 | 74,5557873 | 59,7945424 | 45,3052534 | 70,6632518 | 101,5986 | 18,0316164 | 21,516 | 18,31938 | 10,93854949 | 19,85368 | 11,89021787 | 5,703399 | 39,99938 | 8,85728 | 16,1525246 |
| Upregulated | ENSMUSG00000046711 | 415,5878135 | 347,850806 | 462,950888 | 190,78635 | 1136,08576 | 751,0645777 | 816,707181 | 1043,78102 | 1110,47007 | 413,140763 | 1762,16484 | 1582,66 | 820,438544 | 1019,671 | 837,9127 | 959,8577175 | 988,548 | 1031,4764 | 374,5232 | 1419,311 | 814,8698 | 447,783876 |
| Upregulated | ENSMUSG00000018543 | 8,459802819 | 4,25505574 | 1,81549368 | 4,81783712 | 6,36106249 | 9,28878453 | 18,1742905 | 6,11113011 | 6,40655811 | 8,62957207 | 17,6658129 | 27,15134 | 9,01580818 | 16,83861 | 21,50536 | 5,469274744 | 12,40855 | 3,963405955 | 0,950566 | 25,33294 | 4,42864 | 4,48681239 |
| Upregulated | ENSMUSG00000085385 | 180,8282852 | 95,7387541 | 132,531039 | 78,0489613 | 162,8432 | 195,0644751 | 238,537563 | 136,889314 | 147,350837 | 189,850586 | 368,773845 | 578,9366 | 270,474245 | 359,2237 | 235,7625 | 283,4907409 | 229,9718 | 409,2216649 | 240,4933 | 219,9966 | 195,7459 | 302,411155 |
| Upregulated | ENSMUSG00000038388 | 121,6096655 | 129,7792 | 102,575393 | 143,571546 | 225,181612 | 165,8711523 | 222,635059 | 117,333698 | 152,689635 | 140,230546 | 470,35227 | 419,532 | 322,565582 | 474,2875 | 285,1452 | 250,6750924 | 253,9617 | 340,8529122 | 227,1854 | 94,66519 | 90,34426 | 316,768955 |
| Upregulated | ENSMUSG00000112823 | 5,287376762 | 9,57387541 | 5,44648104 | 1,92713485 | 16,5387625 | 3,980907656 | 11,3589316 | 6,11113011 | 6,40655811 | 5,39348255 | 33,1233993 | 31,53059 | 21,0368858 | 23,38696 | 8,761444 | 11,85009528 | 17,37197 | 15,85362382 | 20,91246 | 17,33306 | 11,51446 | 21,5366995 |
| Upregulated | ENSMUSG00000034333 | 41,24153874 | 48,933141 | 35,4021268 | 32,7612924 | 39,4385875 | 34,50119968 | 59,0664442 | 59,8890751 | 57,659023 | 53,9348255 | 129,181257 | 172,5424 | 108,189698 | 133,7734 | 96,37589 | 80,21602958 | 91,82329 | 89,17663399 | 164,448 | 78,66544 | 47,82931 | 89,7362477 |
| Upregulated | ENSMUSG00000116237 | 23,26445775 | 11,7014033 | 15,4316963 | 25,052753 | 21,6276125 | 2,653938437 | 23,8537563 | 9,77780817 | 26,6939921 | 16,1804476 | 75,079705 | 97,21932 | 54,0948491 | 53,32227 | 70,88805 | 63,80820534 | 57,07934 | 29,72554466 | 57,03399 | 14,66644 | 37,20058 | 51,1496612 |
| Upregulated | ENSMUSG00000018334 | 227,3572007 | 99,9938098 | 182,457115 | 85,7575007 | 159,026562 | 200,372352 | 157,889149 | 183,333903 | 244,516968 | 125,128795 | 696,695498 | 747,0998 | 249,43736 | 677,2864 | 235,7625 | 327,2449388 | 216,7361 | 248,7037237 | 228,136 | 134,6646 | 277,2329 | 159,730521 |
| Upregulated | ENSMUSG00000028680 | 58,16114438 | 76,5910032 | 52,6493167 | 84,7939333 | 72,5161124 | 149,9475217 | 138,578965 | 85,5558215 | 114,250286 | 96,0039893 | 257,258401 | 508,8687 | 118,207263 | 267,5468 | 86,02145 | 120,3240444 | 68,66066 | 108,9936638 | 40,87436 | 101,3318 | 71,74397 | 61,0206485 |
| Upregulated | ENSMUSG00000028889 | 78,25317607 | 56,3794885 | 45,387342 | 57,8140454 | 36,8941625 | 99,5226914 | 53,3869784 | 77,0002394 | 91,827333 | 66,8791836 | 239,592588 | 479,9656 | 141,247662 | 303,095 | 116,2883 | 98,44694539 | 65,35171 | 104,0394063 | 63,68796 | 80,66541 | 64,65815 | 54,7391111 |
| Upregulated | ENSMUSG00000028124 | 84,59802819 | 75,5272393 | 82,6049624 | 109,846686 | 77,6049624 | 120,7541989 | 74,9689484 | 99,0003078 | 119,589085 | 131,600974 | 379,814978 | 663,8941 | 239,419795 | 359,2237 | 253,2854 | 185,9553413 | 113,3314 | 120,8838816 | 89,35325 | 55,33247 | 57,57232 | 81,6599854 |
| Upregulated | ENSMUSG00000106239 | 33,83921127 | 41,4867934 | 54,4648104 | 32,7612924 | 24,1720375 | 27,86635359 | 11,3589316 | 13,4444862 | 30,9650309 | 48,5413429 | 177,762243 | 405,5184 | 60,1053879 | 173,0635 | 58,94063 | 61,07356797 | 36,39842 | 56,47853486 | 42,77549 | 19,33303 | 27,45757 | 48,4575738 |

| | | | | | | | | | | | | | | | | | | | | | | | |
|---|---|---|---|---|---|---|---|---|---|---|---|---|---|---|---|---|---|---|---|---|---|---|---|
| Upregulated | ENSMUSG00000107092 | 339,4495881 | 240,410649 | 250,538128 | 241,855423 | 118,315762 | 228,2387056 | 163,568615 | 168,667191 | 171,909309 | 368,914206 | 893,227667 | 2258,816 | 517,908092 | 1045,865 | 414,9739 | 425,6918842 | 234,9353 | 347,7888726 | 282,3182 | 201,3302 | 271,0328 | 228,827432 |
| Upregulated | ENSMUSG00000011179 | 541,4273804 | 496,777757 | 489,275547 | 413,370425 | 217,548337 | 406,0525809 | 213,547914 | 249,334108 | 301,108231 | 505,908663 | 1725,7291 | 3892,276 | 827,45084 | 1792,377 | 737,5543 | 739,2636362 | 489,7242 | 547,9408733 | 441,0629 | 445,3264 | 424,2637 | 381,379053 |
| Upregulated | ENSMUSG00000033545 | 151,2189754 | 115,950269 | 96,221165 | 72,2675568 | 90,3270874 | 108,8114759 | 97,6868116 | 70,8891093 | 102,50493 | 121,892706 | 538,807295 | 1135,101 | 305,535722 | 435,9329 | 201,5132 | 251,5866382 | 205,1547 | 148,6277233 | 271,862 | 128,6647 | 77,05834 | 157,038434 |
| Upregulated | ENSMUSG00000071151 | 28,55183451 | 17,0202229 | 23,6014178 | 26,9798879 | 19,0831875 | 19,90453828 | 20,4460768 | 20,7778424 | 20,287434 | 34,5182883 | 129,181257 | 174,2941 | 55,0966056 | 73,90279 | 34,24928 | 40,10801479 | 37,22566 | 10,89936638 | 34,22039 | 38,66606 | 41,62922 | 15,2551621 |
| Upregulated | ENSMUSG00000027583 | 50,75881691 | 32,976682 | 26,3246584 | 15,4170788 | 54,7051374 | 59,71361484 | 43,16394 | 41,5556847 | 44,8459068 | 80,9022382 | 185,491036 | 195,3145 | 84,147543 | 89,80593 | 52,56867 | 38,28492321 | 30,60776 | 58,46023784 | 77,94645 | 31,33284 | 72,6297 | 88,8388853 |
| Upregulated | ENSMUSG00000028982 | 31,72426057 | 55,3157246 | 12,7084558 | 16,3806462 | 38,166375 | 35,8281689 | 55,6587648 | 39,1112327 | 68,3366199 | 45,3052534 | 221,926775 | 211,9556 | 80,1405172 | 72,03184 | 23,09835 | 28,25791951 | 27,29882 | 64,40534677 | 9,505665 | 29,99953 | 36,31485 | 17,0498871 |
| Upregulated | ENSMUSG00000049551 | 6,344852114 | 3,1912918 | 0,90774684 | 0 | 0 | 6,634846093 | 1,13589316 | 4,88890409 | 1,06775969 | 4,31478604 | 67,3509118 | 49,92343 | 8,01405172 | 19,64505 | 4,77897 | 8,203912116 | 0 | 2,972554466 | 3,802266 | 0 | 0 | 5,38417486 |
| Upregulated | ENSMUSG00000060733 | 201,9777923 | 223,390426 | 198,796558 | 102,138147 | 153,937712 | 339,70412 | 236,265777 | 188,222807 | 272,27872 | 319,294167 | 993,701978 | 1292,754 | 246,43209 | 588,4159 | 230,187 | 215,1248066 | 125,74 | 199,1611493 | 136,8816 | 127,3313 | 160,3168 | 167,806783 |
| Upregulated | ENSMUSG00000040128 | 1184,372395 | 1090,35803 | 1074,77226 | 962,603857 | 763,327499 | 1076,172036 | 723,563942 | 908,113934 | 1256,75315 | 1253,44534 | 3897,51998 | 4455,447 | 880,543932 | 1641,765 | 1029,071 | 691,8632551 | 516,1958 | 733,2301017 | 596,0052 | 533,9917 | 698,8394 | 604,82231 |
| Upregulated | ENSMUSG00000034765 | 140,6442219 | 45,7418492 | 149,778229 | 77,0853939 | 54,7051374 | 149,9475217 | 70,4253758 | 39,1112327 | 73,6754183 | 124,050099 | 577,451261 | 606,9638 | 57,1001185 | 181,4828 | 89,20743 | 73,83520904 | 49,63421 | 72,33215868 | 29,46756 | 33,99947 | 46,05786 | 57,4311986 |
| Upregulated | ENSMUSG00000028444 | 114,2073381 | 174,457285 | 56,2803041 | 95,393175 | 47,0718625 | 39,80907656 | 63,6100169 | 52,5557189 | 32,0327906 | 66,8791836 | 209,781529 | 3033,943 | 357,627058 | 2453,76 | 463,5601 | 806,7180247 | 153,8661 | 183,3075254 | 138,7827 | 37,33275 | 3,542912 | 41,278674 |
| Upregulated | ENSMUSG00000033323 | 185,0581867 | 140,416839 | 139,793013 | 138,753709 | 193,3763 | 175,1599369 | 204,460768 | 165,000513 | 286,159596 | 165,040566 | 384,231431 | 864,4636 | 294,516401 | 859,7046 | 293,1101 | 313,571752 | 190,2645 | 204,1154067 | 113,1174 | 153,3309 | 144,3737 | 146,270084 |
| Upregulated | ENSMUSG00000036422 | 1,057475352 | 0 | 0 | 4,81783712 | 3,8166375 | 11,94272297 | 4,54357263 | 0 | 2,13551937 | 2,15739302 | 16,5616996 | 37,66154 | 0 | 31,80627 | 5,575465 | 9,115457906 | 7,445132 | 6,935960422 | 0,950566 | 11,33316 | 0 | 1,79472495 |
| Upregulated | ENSMUSG00000037999 | 0 | 2,12752787 | 1,81549368 | 1,92713485 | 1,2722125 | 6,634846093 | 6,81535895 | 1,22222602 | 17,084155 | 8,62957207 | 15,4575863 | 31,53059 | 10,0175646 | 31,80627 | 3,18598 | 8,203912116 | 7,445132 | 1,981702978 | 0,950566 | 5,999906 | 7,085824 | 7,17889982 |
| Upregulated | ENSMUSG00000037224 | 0 | 1,06376393 | 0,90774684 | 4,81783712 | 1,2722125 | 0 | 0 | 4,88890409 | 1,06775969 | 22,6526267 | 15,4575863 | 35,90984 | 6,01053879 | 18,70957 | 3,982475 | 0,911545791 | 4,136184 | 1,981702978 | 3,802266 | 17,99972 | 18,60029 | 5,38417486 |
| Upregulated | ENSMUSG00000005057 | 26,43688381 | 37,2317377 | 42,6641015 | 6,74497197 | 16,5387625 | 13,26969219 | 5,67946579 | 20,7778424 | 11,7453565 | 57,170915 | 80,6002715 | 167,2873 | 45,0790409 | 94,48332 | 47,7897 | 58,3389306 | 54,59763 | 31,70724764 | 36,12153 | 52,66584 | 50,4865 | 61,9180109 |
| Upregulated | ENSMUSG00000026163 | 5,287376762 | 49,9969049 | 18,1549368 | 24,0891856 | 5,08884999 | 42,463015 | 23,8537563 | 75,7780133 | 18,1519147 | 28,0461092 | 80,6002715 | 225,0934 | 118,207263 | 97,28975 | 74,07403 | 36,46183163 | 4,136184 | 17,8353268 | 30,41813 | 65,99897 | 25,68611 | 20,639337 |
| Upregulated | ENSMUSG00000096351 | 0 | 0 | 7,26197472 | 0 | 0 | 0 | 0 | 7,33335613 | 4,27103874 | 0 | 0 | 79,70232 | 11,0193211 | 22,45148 | 1,59299 | 0,911545791 | 12,40855 | 7,92681191 | 1,901133 | 0,666656 | 9,743008 | 4,48681239 |
| Upregulated | ENSMUSG00000048218 | 35,95416198 | 67,0171278 | 54,4648104 | 3,8542697 | 26,7164625 | 19,90453828 | 21,58197 | 6,11113011 | 18,1519147 | 64,7217906 | 114,827784 | 256,624 | 60,1053879 | 59,87062 | 56,55114 | 41,01956058 | 13,23579 | 12,88106935 | 19,01133 | 7,999875 | 7,085824 | 15,2551621 |
| Upregulated | ENSMUSG00000043439 | 30,66678522 | 14,8926951 | 22,693671 | 1,92713485 | 6,36106249 | 9,28878453 | 2,27178632 | 14,6667123 | 13,8808759 | 25,8887162 | 52,9974388 | 178,6733 | 17,0298599 | 12,16122 | 15,1334 | 8,203912116 | 3,308947 | 3,963405955 | 3,802266 | 7,999875 | 6,200096 | 5,38417486 |
| Upregulated | ENSMUSG00000050896 | 5,287376762 | 9,57387541 | 5,44648104 | 4,81783712 | 0 | 2,653938437 | 2,27178632 | 1,22222602 | 1,06775969 | 6,47217906 | 80,6002715 | 299,5406 | 45,0790409 | 24,32244 | 5,575465 | 8,203912116 | 7,445132 | 1,981702978 | 11,4068 | 3,999938 | 5,314368 | 0,89736248 |
| Upregulated | ENSMUSG00000080950 | 20,09203169 | 21,2752787 | 10,8929621 | 9,63567424 | 6,36106249 | 7,961815312 | 4,54357263 | 4,88890409 | 6,40655811 | 17,2591441 | 70,6632518 | 185,6801 | 27,0474245 | 46,77392 | 17,52289 | 28,25791951 | 9,926842 | 14,86277233 | 16,15963 | 9,999844 | 2,657184 | 14,3577996 |
| Upregulated | ENSMUSG00000091952 | 96,23025706 | 78,7185311 | 49,0183294 | 41,4333992 | 16,5387625 | 54,40573796 | 29,5332221 | 18,3333903 | 40,5748681 | 47,4626464 | 244,009041 | 577,1849 | 90,1580818 | 165,5797 | 66,90558 | 70,18902588 | 61,21553 | 62,4236438 | 58,93512 | 34,66613 | 35,42912 | 36,7918616 |
| Upregulated | ENSMUSG00000080845 | 27,49435916 | 45,7418492 | 32,6788862 | 8,67210682 | 8,90548749 | 15,92363062 | 13,6307179 | 18,3333903 | 8,54207749 | 60,4070045 | 143,53473 | 396,7599 | 70,1229525 | 81,38662 | 23,09835 | 29,1694653 | 17,37197 | 23,78043573 | 19,9619 | 7,333219 | 19,48602 | 9,87098725 |
| Upregulated | ENSMUSG00000073700 | 1477,293067 | 1527,56501 | 1478,7196 | 539,597757 | 810,399362 | 954,0908682 | 822,386647 | 690,557702 | 994,084267 | 1565,18863 | 4592,00725 | 12156,79 | 2133,74127 | 3387,367 | 1327,757 | 1426,569162 | 865,2897 | 1130,561549 | 851,7076 | 586,6575 | 674,039 | 828,265567 |
| Upregulated | ENSMUSG00000014773 | 146,989074 | 193,605036 | 135,254279 | 95,393175 | 398,202512 | 183,1217522 | 407,785644 | 188,222807 | 135,60548 | 207,10973 | 452,686457 | 493,9792 | 138,242392 | 229,1922 | 760,6527 | 419,3110637 | 701,4968 | 297,2554466 | 412,5459 | 161,3308 | 75,28688 | 326,639942 |
| Upregulated | ENSMUSG00000070822 | 11,63222888 | 6,3825836 | 4,5387342 | 5,78140454 | 10,1777 | 18,57756906 | 17,0383974 | 9,77780817 | 6,40655811 | 10,7869651 | 24,2904928 | 31,53059 | 5,00878232 | 14,96765 | 31,8598 | 22,78864477 | 20,68092 | 19,81702978 | 19,9619 | 3,999938 | 6,200096 | 20,639337 |
| Upregulated | ENSMUSG00000025938 | 27,49435916 | 89,3561705 | 66,2655193 | 24,0891856 | 63,6106249 | 87,57996843 | 119,268782 | 92,8891777 | 38,4393487 | 97,0826858 | 220,822662 | 141,8877 | 120,210776 | 79,51566 | 226,2046 | 242,4711803 | 205,1547 | 156,5545352 | 200,5695 | 137,9978 | 93,88717 | 218,956444 |
| Upregulated | ENSMUSG00000063632 | 154,3914014 | 60,6345442 | 129,807798 | 53,9597757 | 260,803562 | 731,1600395 | 130,627713 | 97,7780817 | 109,979248 | 132,679671 | 455,998797 | 451,0626 | 138,242392 | 495,8035 | 621,2661 | 476,7384485 | 707,2875 | 334,9078032 | 390,6828 | 153,3309 | 73,51543 | 233,314244 |
| Upregulated | ENSMUSG00000075302 | 23,26445775 | 52,1244328 | 36,3098736 | 26,0163204 | 150,121075 | 106,1575375 | 87,4637732 | 70,8891093 | 99,3016508 | 40,9904673 | 93,8496312 | 102,4744 | 33,0579633 | 97,28975 | 173,6359 | 83,86221274 | 95,95947 | 58,46023784 | 141,6344 | 28,66622 | 30,11475 | 104,094047 |
| Upregulated | ENSMUSG00000071047 | 8,459802819 | 8,51011147 | 1,81549368 | 5,78140454 | 29,2608875 | 7,961815312 | 9,08714527 | 15,8889383 | 1,06775969 | 14,0230546 | 25,3946061 | 14,88944 | 16,0281034 | 32,74174 | 30,26681 | 17,31937002 | 30,60776 | 9,908514888 | 19,01133 | 5,999906 | 5,314368 | 17,9472495 |
| Upregulated | ENSMUSG00000117234 | 7,402327466 | 13,8289311 | 17,24719 | 15,4170788 | 248,081437 | 58,38664562 | 122,676461 | 26,8889725 | 16,0163953 | 33,4395918 | 114,827784 | 266,2583 | 23,0403987 | 242,2889 | 545,599 | 205,0978029 | 257,2707 | 144,6643174 | 19,9619 | 50,66588 | 155,8881 | 8,97362477 |
| Upregulated | ENSMUSG00000063727 | 420,8751902 | 422,314282 | 459,319901 | 634,027365 | 111,9547 | 67,67543015 | 168,112187 | 100,222534 | 160,163953 | 288,011968 | 216,406209 | 2921,835 | 1104,93738 | 2212,406 | 2920,747 | 3609,721331 | 3096,348 | 1761,733947 | 1173,95 | 1635,974 | 1566,853 | 1132,47145 |
| Upregulated | ENSMUSG00002076348 | 12,68970423 | 11,7014033 | 9,98521524 | 2,89070227 | 5,08884999 | 11,94272297 | 10,2230384 | 0 | 5,33879843 | 2,15739302 | 22,0822662 | 118,2397 | 31,0544504 | 93,54784 | 125,8462 | 76,56984641 | 127,3945 | 46,57001997 | 24,71473 | 33,33281 | 43,40067 | 24,2287869 |
| Upregulated | ENSMUSG00000048279 | 72,96579931 | 68,0808918 | 62,634532 | 37,5791295 | 71,2438999 | 57,0596764 | 64,74591 | 94,1114037 | 50,1847052 | 105,712258 | 143,53473 | 232,976 | 74,1299784 | 241,3534 | 266,8258 | 233,3557224 | 268,0247 | 172,4081591 | 193,9156 | 138,6645 | 69,97251 | 169,601508 |
| Upregulated | ENSMUSG00000023032 | 0 | 0 | 0 | 2,89070227 | 0 | 0 | 1,13589316 | 6,11113011 | 1,06775969 | 0 | 0 | 56,05438 | 3,00526939 | 37,41914 | 93,18991 | 63,80820534 | 39,70737 | 15,85362382 | 47,52832 | 49,99922 | 23,02893 | 35,8944991 |
| Upregulated | ENSMUSG00000051341 | 88,82792959 | 94,6749901 | 98,9444056 | 47,2148038 | 77,6049624 | 34,50119968 | 71,561269 | 57,444623 | 52,3202246 | 86,2957207 | 108,203104 | 471,2071 | 185,324946 | 511,7067 | 504,9778 | 251,5866382 | 396,2464 | 177,3624165 | 273,7632 | 71,33222 | 116,9161 | 222,545894 |
| Upregulated | ENSMUSG00000034796 | 29,60930986 | 25,5303344 | 13,6162026 | 23,1256182 | 35,62195 | 26,53938437 | 23,8537563 | 19,5556163 | 16,0163953 | 7,55087556 | 64,0385719 | 165,5356 | 42,0737715 | 156,2249 | 176,8219 | 72,01211746 | 134,0124 | 33,68895062 | 53,23172 | 19,99969 | 22,1432 | 34,0997741 |
| Upregulated | ENSMUSG00000117914 | 0 | 2,12752787 | 4,5387342 | 4,81783712 | 5,08884999 | 1,326969219 | 0 | 6,11113011 | 0 | 0 | 3,31233993 | 21,89624 | 3,00526939 | 25,25792 | 35,84227 | 5,469274744 | 17,37197 | 6,935960422 | 2,851699 | 0 | 0 | 4,48681239 |
| Upregulated | ENSMUSG00000119009 | 0 | 0 | 0 | 0 | 0 | 0 | 0 | 0 | 0 | 0 | 0 | 23,64794 | 3,00526939 | 21,516 | 19,11588 | 1,823091581 | 8,272368 | 5,945108933 | 11,4068 | 3,999938 | 3,542912 | 6,28153734 |
| Upregulated | ENSMUSG00000119451 | 0 | 0 | 0 | 0 | 0 | 1,326969219 | 0 | 0 | 0 | 0 | 2,20822662 | 24,52379 | 1,00175646 | 23,38696 | 26,28433 | 4,557728953 | 11,58132 | 13,87192084 | 11,4068 | 1,999969 | 0,885728 | 8,97362477 |
| Upregulated | ENSMUSG00000119118 | 0 | 0 | 0,90774684 | 0 | 0 | 2,653938437 | 0 | 1,22222602 | 0 | 0 | 0 | 28,90304 | 7,01229525 | 23,38696 | 43,01073 | 15,49627844 | 18,19921 | 20,80788127 | 29,46756 | 5,33325 | 19,48602 | 9,87098725 |
| Upregulated | ENSMUSG00002075987 | 23,26445775 | 28,7216262 | 29,9556457 | 9,63567424 | 33,077525 | 51,75179953 | 30,6691153 | 22,0000684 | 29,8972712 | 31,2821988 | 73,9755917 | 750,6032 | 117,205506 | 314,3207 | 607,7256 | 342,7412173 | 351,5757 | 161,5087927 | 128,3265 | 158,6642 | 160,3168 | 180,369858 |
| Upregulated | ENSMUSG00002076809 | 53,93124297 | 46,8056131 | 40,8486078 | 19,2713485 | 27,988675 | 84,92602999 | 67,0176963 | 9,77780817 | 44,8459068 | 29,1248057 | 100,474311 | 1086,929 | 167,29333 | 471,4811 | 1012,345 | 431,161159 | 483,1063 | 262,5756445 | 182,5088 | 170,664 | 226,7464 | 253,953581 |
| Upregulated | ENSMUSG00002076161 | 152,2764507 | 134,034256 | 136,162026 | 75,1582591 | 123,404612 | 160,5632755 | 134,035393 | 68,4446572 | 171,909309 | 152,096208 | 314,672293 | 4112,114 | 747,310323 | 1849,441 | 3656,708 | 1676,332709 | 1809,994 | 1168,213905 | 630,2256 | 781,9878 | 922,0429 | 941,333239 |
| Upregulated | ENSMUSG00000118798 | 3,172426057 | 1,06376393 | 0 | 0 | 0 | 0 | 0 | 0 | 0 | 0 | 0 | 35,90984 | 4,00702586 | 21,516 | 33,45279 | 18,23091581 | 24,81711 | 5,945108933 | 19,9619 | 4,666594 | 5,314368 | 8,0762623 |
| Upregulated | ENSMUSG00000118870 | 0 | 0 | 0,90774684 | 0 | 0 | 0 | 0 | 0 | 0 | 1,07869651 | 3,31233993 | 58,68193 | 6,01053879 | 36,48366 | 57,34764 | 11,85009528 | 22,33539 | 18,82617829 | 24,71473 | 2,666625 | 9,743008 | 18,844612 |
| Upregulated | ENSMUSG00000119748 | 0 | 0 | 0 | 0 | 0 | 0 | 0 | 1,22222602 | 0 | 0 | 0 | 22,77209 | 3,00526939 | 14,03218 | 21,50536 | 6,380820534 | 12,40855 | 9,908514888 | 7,604532 | 0 | 6,200096 | 8,97362477 |
| Upregulated | ENSMUSG00000119012 | 0 | 0 | 0 | 1,92713485 | 0 | 0 | 1,13589316 | 1,22222602 | 0 | 0 | 0 | 24,52379 | 6,01053879 | 14,03218 | 35,04578 | 8,203912116 | 16,54474 | 8,917663399 | 10,45623 | 0,666656 | 4,42864 | 7,17889982 |
| Upregulated | ENSMUSG00000119730 | 0 | 0 | 0 | 0 | 0 | 0 | 0 | 0 | 0 | 0 | 0 | 36,78569 | 8,01405172 | 19,64505 | 50,97568 | 12,76164107 | 28,95329 | 16,84447531 | 14,2585 | 1,333313 | 7,971552 | 18,844612 |
| Upregulated | ENSMUSG00000070167 | 6,344852114 | 5,31881967 | 10,8929621 | 10,5992417 | 10,1777 | 14,5966614 | 12,4948247 | 6,11113011 | 19,2196743 | 8,62957207 | 11,0411331 | 27,15134 | 20,0351293 | 27,12887 | 40,62124 | 11,85009528 | 10,75408 | 16,84447531 | 20,91246 | 6,666563 | 2,657184 | 10,7683497 |
| Upregulated | ENSMUSG00000118106 | 0 | 8,51011147 | 4,5387342 | 0 | 1,2722125 | 2,653938437 | 4,54357263 | 3,66667807 | 1,06775969 | 7,55087556 | 13,2493597 | 13,13775 | 18,0316164 | 18,70957 | 28,67382 | 7,292366325 | 8,272368 | 3,963405955 | 10,45623 | 5,999906 | 4,42864 | 9,87098725 |
| Upregulated | ENSMUSG00000119615 | 0 | 0 | 0 | 0 | 0 | 0 | 0 | 1,22222602 | 0 | 1,07869651 | 0 | 35,90984 | 11,0193211 | 4,677392 | 22,30186 | 5,469274744 | 11,58132 | 6,935960422 | 15,20906 | 3,333281 | 4,42864 | 4,48681239 |
| Upregulated | ENSMUSG00000053192 | 31,72426057 | 21,2752787 | 44,4795952 | 31,797725 | 59,7939874 | 30,52029203 | 39,7562605 | 29,3334245 | 35,2360696 | 51,7774324 | 51,8933255 | 162,908 | 92,1615947 | 101,0317 | 137,7936 | 74,74675483 | 104,2318 | 77,28641613 | 133,0793 | 75,33216 | 43,40067 | 72,6863607 |
| Upregulated | ENSMUSG00000031910 | 3,172426057 | 3,1912918 | 0 | 0 | 0 | 6,634846093 | 0 | 1,22222602 | 3,20327906 | 0 | 9,93701978 | 33,28229 | 8,01405172 | 6,548349 | 37,43526 | 14,58473265 | 9,099605 | 9,908514888 | 28,51699 | 15,33309 | 0 | 13,4604372 |
| Upregulated | ENSMUSG00000030069 | 78,25317607 | 61,6983082 | 131,623292 | 47,2148038 | 33,077525 | 33,17423047 | 22,7178632 | 35,4445546 | 19,2196743 | 179,063621 | 98,2660845 | 493,9792 | 130,22834 | 130,0315 | 335,3244 | 240,6480887 | 138,9758 | 120,8838816 | 123,5736 | 111,3316 | 90,34426 | 154,346346 |
| Upregulated | ENSMUSG00000057835 | 17,97708099 | 46,8056131 | 42,6641015 | 13,4899439 | 50,8884999 | 50,42483031 | 11,3589316 | 64,7779792 | 39,5071084 | 107,869651 | 43,060419 | 158,5288 | 81,1422736 | 90,7414 | 117,0848 | 78,39293799 | 80,24197 | 76,29556464 | 69,39135 | 87,99863 | 46,05786 | 101,40196 |
| Upregulated | ENSMUSG00000116679 | 24,3219331 | 17,0202229 | 23,6014178 | 21,1984833 | 49,6162874 | 46,44392265 | 34,0767947 | 29,3334245 | 26,6939921 | 38,8330743 | 47,4768723 | 112,1088 | 56,098362 | 43,03201 | 53,36516 | 57,42738481 | 46,32526 | 22,78958424 | 25,6653 | 27,33291 | 18,60029 | 41,278674 |
| Upregulated | ENSMUSG00000102752 | 12,68970423 | 29,7853902 | 10,8929621 | 9,63567424 | 13,9943375 | 46,44392265 | 10,2230384 | 19,5556163 | 28,8295115 | 11,8656616 | 72,8714784 | 200,5696 | 64,1124137 | 90,7414 | 82,03898 | 98,44694539 | 47,97974 | 32,69809913 | 61,78682 | 15,33309 | 30,11475 | 34,0997741 |
| Upregulated | ENSMUSG00000022237 | 283,4033944 | 114,886505 | 163,394431 | 77,0853939 | 148,848862 | 244,1623362 | 214,683807 | 238,334074 | 156,960674 | 92,7678998 | 428,395964 | 711,1899 | 297,52167 | 333,9658 | 344,0858 | 718,298083 | 166,2746 | 285,3652288 | 109,3151 | 71,99888 | 102,7444 | 182,164583 |
| Upregulated | ENSMUSG00000022479 | 924,2334579 | 738,25217 | 419,37904 | 459,621661 | 618,295274 | 1046,978713 | 885,996663 | 565,890648 | 336,344301 | 527,482593 | 780,608109 | 2624,046 | 1018,78632 | 1516,41 | 2101,154 | 2230,55255 | 1615,594 | 1232,619252 | 942,962 | 529,9917 | 213,4605 | 989,790813 |
| Upregulated | ENSMUSG00000030930 | 168,138581 | 351,042098 | 129,807798 | 96,3567424 | 248,081437 | 47,77089187 | 137,443072 | 145,444897 | 133,469961 | 243,785411 | 550,952541 | 1132,474 | 422,741228 | 653,8994 | 1016,328 | 995,4080034 | 650,2082 | 524,1604376 | 463,8764 | 329,9948 | 247,1181 | 546,493749 |
| Upregulated | ENSMUSG00000026656 | 176,5983838 | 437,206977 | 746,167902 | 469,257335 | 129,765675 | 82,27209155 | 77,2407348 | 58,666849 | 275,481999 | 266,438038 | 809,315055 | 2837,753 | 577,011724 | 3127,304 | 2438,071 | 2965,258457 | 864,4625 | 886,8120825 | 553,2297 | 35,99944 | 51,37222 | 473,807388 |
| Upregulated | ENSMUSG00000037482 | 1,057475352 | 8,51011147 | 5,44648104 | 8,67210682 | 0 | 0 | 0 | 3,66667807 | 2,13551937 | 1,07869651 | 3,31233993 | 125,2465 | 8,01405172 | 52,38679 | 59,73712 | 104,8277659 | 32,26224 | 10,89936638 | 8,555098 | 6,666563 | 5,314368 | 13,4604372 |
| Upregulated | ENSMUSG00000108599 | 21,14950705 | 19,1477508 | 21,7859242 | 6,74497197 | 21,6276125 | 27,86635359 | 34,0767947 | 14,6667123 | 3,20327906 | 43,1478604 | 30,9151726 | 76,19892 | 31,0544504 | 41,16105 | 55,75465 | 41,01956058 | 43,84355 | 47,56087146 | 40,87436 | 8,666531 | 5,314368 | 26,9208743 |
| Upregulated | ENSMUSG00000027313 | 112,0923873 | 440,398269 | 326,788862 | 202,349159 | 571,223412 | 518,8449645 | 430,503507 | 926,447324 | 389,732285 | 940,623356 | 679,029685 | 1804,25 | 1414,48013 | 1435,024 | 1094,384 | 1298,952752 | 1263,191 | 757,0105375 | 783,2668 | 93,99853 | 264,8327 | 690,071745 |
| Upregulated | ENSMUSG00000019997 | 1750,121708 | 5099,6843 | 707,134788 | 7386,70787 | 1067,38629 | 1518,052786 | 1530,04808 | 1227,11493 | 1155,31598 | 2256,6331 | 2098,9194 | 6057,376 | 2379,1716 | 7960,921 | 6439,662 | 9761,743872 | 8319,521 | 8126,963911 | 4921,083 | 2997,287 | 5675,745 | 6854,0546 |
| Upregulated | ENSMUSG00000048376 | 4210,866853 | 7506,98208 | 3414,03587 | 9259,88294 | 4308,98373 | 1682,596969 | 5512,4895 | 6686,79857 | 2702,49976 | 6837,85717 | 5539,33647 | 16043,81 | 6534,45742 | 12446,54 | 8246,112 | 10768,09042 | 13390,48 | 13083,20306 | 8518,026 | 4625,261 | 3575,684 | 8274,5794 |
| Upregulated | ENSMUSG00000030525 | 222,069824 | 479,757534 | 252,353622 | 526,107814 | 393,113662 | 165,8711523 | 252,168281 | 144,222671 | 240,245929 | 417,455549 | 505,683895 | 1202,542 | 251,440873 | 775,5116 | 1299,083 | 1032,781381 | 807,3832 | 1253,427133 | 439,1617 | 5,33325 | 51,37222 | 519,572874 |
| Upregulated | ENSMUSG00000017652 | 93,057831 | 188,286216 | 307,726179 | 255,345367 | 187,015237 | 348,9929045 | 210,140234 | 157,667157 | 274,414239 | 272,910217 | 336,754559 | 776,8787 | 292,512888 | 615,5448 | 986,8572 | 1183,186436 | 877,6983 | 366,6150509 | 407,793 | 109,3316 | 205,4889 | 336,510929 |
| Upregulated | ENSMUSG00000027470 | 28,55183451 | 5,31881967 | 10,8929621 | 21,1984833 | 2,544425 | 11,94272297 | 17,0383974 | 6,11113011 | 3,20327906 | 6,47217906 | 16,5616996 | 64,81288 | 28,049181 | 11,22574 | 113,8988 | 172,2821544 | 72,79684 | 45,57916849 | 15,20906 | 1,999969 | 5,314368 | 48,4575738 |
| Upregulated | ENSMUSG00000022887 | 10753,46686 | 13342,791 | 11698,1335 | 8242,35575 | 6855,95316 | 4335,208437 | 7184,52423 | 5271,46083 | 7017,31665 | 7759,06399 | 8076,58885 | 16035,93 | 7240,69573 | 8040,437 | 19835,91 | 20813,32504 | 20852,99 | 15789,21847 | 18165,33 | 2145,3 | 2262,149 | 8664,93208 |
| Upregulated | ENSMUSG00000053297 | 45,47144015 | 15,956459 | 29,0478989 | 28,9070227 | 22,899825 | 6,634846093 | 23,8537563 | 6,11113011 | 32,0327906 | 10,7869651 | 6,62467985 | 35,03399 | 19,0333728 | 33,67722 | 74,87053 | 53,78120165 | 56,25211 | 64,40534677 | 38,97323 | 5,33325 | 23,02893 | 35,8944991 |
| Upregulated | ENSMUSG00000119562 | 0 | 0 | 0 | 0 | 0 | 1,326969219 | 0 | 0 | 0 | 0 | 1,10411331 | 26,27549 | 8,01405172 | 5,61287 | 31,0633 | 3,646183163 | 32,26224 | 7,92681191 | 11,4068 | 1,999969 | 2,657184 | 0,89736248 |
| Upregulated | ENSMUSG00000118860 | 1,057475352 | 3,1912918 | 0,90774684 | 0 | 0 | 0 | 2,27178632 | 2,44445204 | 3,20327906 | 0 | 2,20822662 | 59,55778 | 18,0316164 | 24,32244 | 64,51609 | 10,0270037 | 57,07934 | 24,77128722 | 28,51699 | 2,666625 | 3,542912 | 24,2287869 |
| Upregulated | ENSMUSG00000049420 | 19,03455634 | 20,2115147 | 18,1549368 | 12,5263765 | 2,544425 | 2,653938437 | 1,13589316 | 12,2222602 | 19,2196743 | 5,39348255 | 9,93701978 | 62,18533 | 10,0175646 | 7,483827 | 73,27754 | 24,61173635 | 110,8497 | 32,69809913 | 12,35736 | 37,99941 | 7,085824 | 34,9971366 |
| Upregulated | ENSMUSG00000072647 | 4,229901409 | 6,3825836 | 1,81549368 | 0,96356742 | 5,08884999 | 10,61575375 | 5,67946579 | 11,0000342 | 5,33879843 | 2,15739302 | 5,52056654 | 21,02039 | 13,022834 | 2,806435 | 15,9299 | 19,1424616 | 19,02645 | 15,85362382 | 9,505665 | 6,666563 | 1,771456 | 14,3577996 |
| Upregulated | ENSMUSG00000020108 | 216,7824472 | 374,444905 | 364,91423 | 264,017474 | 741,699887 | 571,9237332 | 572,490152 | 388,667875 | 186,857945 | 400,196405 | 230,759682 | 934,5316 | 457,802704 | 397,5783 | 1356,431 | 1298,952752 | 1499,78 | 874,9218646 | 987,6386 | 675,3228 | 169,1741 | 585,977698 |
| Upregulated | ENSMUSG00000085609 | 2,114950705 | 5,31881967 | 9,0774684 | 11,5628091 | 11,4499125 | 13,26969219 | 18,1742905 | 11,0000342 | 5,33879843 | 3,23608953 | 6,62467985 | 35,90984 | 9,01580818 | 11,22574 | 30,26681 | 27,34637372 | 30,60776 | 12,88106935 | 15,20906 | 2,666625 | 7,971552 | 7,17889982 |
| Upregulated | ENSMUSG00000058099 | 0 | 7,44634754 | 3,63098736 | 8,67210682 | 5,08884999 | 1,326969219 | 2,27178632 | 18,3333903 | 3,20327906 | 16,1804476 | 1,10411331 | 77,95062 | 14,0245905 | 4,677392 | 52,56867 | 39,196469 | 67,00618 | 24,77128722 | 4,752832 | 0 | 2,657184 | 3,58944991 |
| Upregulated | ENSMUSG00000111116 | 0 | 17,0202229 | 4,5387342 | 3,8542697 | 5,08884999 | 0 | 1,13589316 | 6,11113011 | 3,20327906 | 0 | 12,1452464 | 42,91663 | 1,00175646 | 12,16122 | 27,08083 | 25,52328214 | 43,01632 | 13,87192084 | 12,35736 | 3,999938 | 0 | 0,89736248 |
| Upregulated | ENSMUSG00000119014 | 0 | 0 | 0 | 0 | 1,2722125 | 0 | 0 | 1,22222602 | 1,06775969 | 0 | 0 | 68,31628 | 10,0175646 | 28,99983 | 42,21423 | 26,43482793 | 38,88013 | 21,79873275 | 30,41813 | 2,666625 | 8,85728 | 13,4604372 |
| Upregulated | ENSMUSG00000118907 | 2,114950705 | 1,06376393 | 0 | 0 | 0 | 1,326969219 | 0 | 0 | 1,06775969 | 0 | 1,10411331 | 70,94383 | 8,01405172 | 29,93531 | 41,41774 | 14,58473265 | 33,08947 | 18,82617829 | 25,6653 | 5,33325 | 12,40019 | 6,28153734 |
| Upregulated | ENSMUSG00000119808 | 0 | 1,06376393 | 2,72324052 | 0 | 1,2722125 | 0 | 2,27178632 | 0 | 0 | 2,15739302 | 0 | 100,7227 | 12,0210776 | 39,29009 | 60,53362 | 21,87709898 | 42,18908 | 20,80788127 | 35,17096 | 3,333281 | 10,62874 | 24,2287869 |
| Upregulated | ENSMUSG00000119204 | 0 | 3,1912918 | 0 | 0 | 0 | 0 | 0 | 0 | 0 | 0 | 1,10411331 | 57,80608 | 15,026347 | 19,64505 | 35,84227 | 14,58473265 | 21,50816 | 17,8353268 | 21,86303 | 2,666625 | 8,85728 | 16,1525246 |
| Upregulated | ENSMUSG00000119546 | 2,114950705 | 0 | 0 | 0,96356742 | 0 | 1,326969219 | 2,27178632 | 0 | 2,13551937 | 0 | 0 | 106,8537 | 18,0316164 | 58,93514 | 90,80042 | 32,81564846 | 70,31513 | 41,61576253 | 59,88569 | 11,99981 | 22,1432 | 25,1261494 |
| Upregulated | ENSMUSG00001118664 | 0 | 0 | 0 | 0 | 0 | 1,326969219 | 1,13589316 | 3,66667807 | 4,27103874 | 1,07869651 | 0 | 319,6851 | 40,0702586 | 183,3538 | 195,1413 | 72,01211746 | 183,6466 | 105,0302578 | 149,2389 | 16,66641 | 41,62922 | 84,3520729 |

| | | | | | | | | | | | | | | | | | | | | | | | |
|---|---|---|---|---|---|---|---|---|---|---|---|---|---|---|---|---|---|---|---|---|---|---|---|
| Upregulated | ENSMUSG00000119523 | 1,057475352 | 3,1912918 | 1,81549368 | 0 | 1,2722125 | 0 | 2,27178632 | 4,88890409 | 2,13551937 | 0 | 1,10411331 | 302,1681 | 37,0649892 | 178,6764 | 208,6817 | 92,06612485 | 188,61 | 119,8930301 | 143,5355 | 13,33313 | 31,00048 | 87,0441603 |
| Upregulated | ENSMUSG00000119210 | 0 | 0 | 0,90774684 | 0 | 0 | 1,326969219 | 0 | 0 | 1,06775969 | 0 | 0 | 85,83327 | 7,01229525 | 43,03201 | 60,53362 | 21,87709898 | 59,56105 | 32,69809913 | 41,82493 | 5,33325 | 11,51446 | 28,7155993 |
| Upregulated | ENSMUSG00000119919 | 0 | 2,12752787 | 0,90774684 | 0 | 0 | 2,653938437 | 0 | 0 | 0 | 1,07869651 | 1,10411331 | 93,71592 | 19,0333728 | 53,32227 | 66,90558 | 17,31937002 | 50,46145 | 36,66150509 | 35,17096 | 7,999875 | 15,05738 | 24,2287869 |
| Upregulated | ENSMUSG00000119587 | 2,114950705 | 1,06376393 | 0 | 0 | 0 | 2,653938437 | 0 | 2,44445204 | 4,27103874 | 1,07869651 | 0 | 62,18533 | 12,0210776 | 23,38696 | 29,47031 | 9,115457906 | 26,47158 | 30,71639615 | 22,8136 | 5,999906 | 7,971552 | 7,17889982 |
| Upregulated | ENSMUSG00000119844 | 0 | 0 | 0 | 0 | 0 | 0 | 0 | 0 | 0 | 0 | 1,10411331 | 36,78569 | 3,00526939 | 18,70957 | 16,72639 | 12,76164107 | 21,50816 | 14,86277233 | 23,76416 | 1,333313 | 15,9431 | 8,97362477 |
| Upregulated | ENSMUSG00000119762 | 0 | 0 | 0 | 0 | 0 | 2,653938437 | 0 | 0 | 0 | 0 | 0 | 100,7227 | 9,01580818 | 34,6127 | 49,38269 | 28,25791951 | 40,53461 | 24,77128722 | 67,49022 | 5,33325 | 9,743008 | 9,87098725 |
| Upregulated | ENSMUSG00000070225 | 0 | 2,12752787 | 0 | 0 | 0 | 0 | 1,13589316 | 0 | 2,13551937 | 3,23608953 | 0 | 47,29588 | 11,0193211 | 30,87079 | 43,01073 | 17,31937002 | 55,42487 | 19,81702978 | 44,67663 | 2,666625 | 15,05738 | 15,2551621 |
| Upregulated | ENSMUSG00000118952 | 0 | 1,06376393 | 0 | 1,92713485 | 0 | 0 | 0 | 0 | 0 | 0 | 2,20822662 | 19,26869 | 1,00175646 | 9,354784 | 16,72639 | 1,823091581 | 16,54474 | 8,917663399 | 12,35736 | 1,999969 | 2,657184 | 6,28153734 |
| Upregulated | ENSMUSG00000118727 | 0 | 0 | 0 | 0 | 0 | 1,326969219 | 2,27178632 | 0 | 0 | 0 | 0 | 59,55778 | 3,00526939 | 35,54818 | 39,82475 | 11,85009528 | 53,77039 | 27,74384169 | 34,22039 | 3,333281 | 7,971552 | 17,0498871 |
| Upregulated | ENSMUSG00000119038 | 3,172426057 | 0 | 0,90774684 | 0 | 0 | 0 | 1,13589316 | 0 | 0 | 0 | 0 | 116,488 | 21,0368858 | 50,51583 | 61,33011 | 43,75419795 | 91,82329 | 26,7529902 | 67,49022 | 5,33325 | 23,02893 | 25,1261494 |
| Upregulated | ENSMUSG00000118725 | 0 | 0 | 2,72324052 | 0 | 1,2722125 | 0 | 2,27178632 | 0 | 0 | 0 | 1,10411331 | 75,32307 | 15,026347 | 43,03201 | 40,62124 | 23,70019056 | 60,38829 | 22,78958424 | 37,07209 | 0,666656 | 21,25747 | 16,1525246 |
| Upregulated | ENSMUSG00000119324 | 2,114950705 | 0 | 2,72324052 | 0 | 3,8166375 | 1,326969219 | 0 | 0 | 0 | 1,07869651 | 2,20822662 | 99,84686 | 13,022834 | 66,41897 | 48,58619 | 51,04656428 | 86,03263 | 34,67980211 | 53,23172 | 15,99975 | 20,37174 | 34,0997741 |
| Upregulated | ENSMUSG00000119299 | 0 | 0 | 0 | 0 | 0 | 0 | 2,27178632 | 4,88890409 | 1,06775969 | 0 | 1,10411331 | 89,33667 | 9,01580818 | 43,03201 | 35,04578 | 32,81564846 | 94,305 | 26,7529902 | 46,57776 | 8,666531 | 22,1432 | 13,4604372 |
| Upregulated | ENSMUSG00000025137 | 670,4393734 | 442,525797 | 553,725572 | 676,424332 | 506,340574 | 453,8234728 | 550,908182 | 613,557463 | 345,954138 | 504,829966 | 620,51168 | 1850,67 | 915,605409 | 1133,8 | 1077,658 | 784,8409257 | 1941,525 | 1473,396164 | 1100,756 | 443,9931 | 836,1272 | 847,110179 |
| Upregulated | ENSMUSG00000051111 | 30,66678522 | 39,3592656 | 31,7711394 | 21,1984833 | 41,9830125 | 18,57756906 | 45,4357263 | 48,8890409 | 52,3202246 | 52,8561289 | 11,0411331 | 124,3707 | 8,01405172 | 57,06418 | 85,22496 | 31,90410267 | 99,26842 | 122,8655846 | 103,6117 | 60,66572 | 60,22951 | 61,9180109 |
| Upregulated | ENSMUSG00000020120 | 80,36812678 | 59,5707803 | 83,5127093 | 54,9233432 | 139,943375 | 63,69452249 | 114,725209 | 182,111677 | 166,570511 | 223,290177 | 228,551455 | 566,6748 | 143,251174 | 187,0957 | 201,5132 | 278,0214661 | 263,0613 | 269,511605 | 331,7477 | 259,3293 | 119,5733 | 259,337756 |
| Upregulated | ENSMUSG00000072082 | 68,7358979 | 90,4199344 | 129,807798 | 40,4698318 | 66,1550499 | 22,55847672 | 46,5716195 | 58,666849 | 45,9136665 | 76,5874522 | 56,3097787 | 525,5098 | 93,1633512 | 92,61236 | 166,4674 | 237,0019056 | 292,8418 | 210,0605156 | 250,9496 | 249,3294 | 162,0882 | 206,39337 |
| Upregulated | ENSMUSG00000117752 | 6,344852114 | 24,4665705 | 11,8007089 | 3,8542697 | 68,6994749 | 37,15513812 | 24,9896495 | 36,6667807 | 20,287434 | 72,2726661 | 142,430617 | 79,70232 | 101,177403 | 10,29026 | 48,58619 | 46,48883532 | 233,2808 | 117,9113272 | 148,2884 | 7,333219 | 31,88621 | 79,8652605 |
| Upregulated | ENSMUSG00000020696 | 436,7373205 | 571,241233 | 521,954433 | 327,612924 | 660,278287 | 459,1313496 | 722,428049 | 1071,89222 | 1048,54001 | 619,171796 | 812,627395 | 1831,402 | 1568,75062 | 1105,735 | 1030,664 | 1731,025456 | 2278,21 | 1528,883847 | 1652,085 | 1014,651 | 876,8707 | 1530,00302 |
| Upregulated | ENSMUSG00000024087 | 10447,85648 | 9746,20516 | 7478,92621 | 5895,1055 | 8165,05982 | 7112,555012 | 7354,9082 | 16436,4955 | 12482,1107 | 12664,9757 | 13787,0629 | 17826,17 | 12581,0594 | 10489,52 | 12625,24 | 22095,86997 | 27829,07 | 18068,1769 | 19960 | 13099,8 | 9880,296 | 14488,8146 |
| Upregulated | ENSMUSG00000035370 | 26,43688381 | 18,0839869 | 29,0478989 | 11,5628091 | 27,988675 | 9,28878453 | 44,2998332 | 37,8890067 | 26,6939921 | 35,5969848 | 55,2056654 | 122,619 | 44,0772844 | 22,45148 | 41,41774 | 70,18902588 | 134,0124 | 53,5059804 | 95,05665 | 60,66572 | 9,743008 | 25,1261494 |
| Upregulated | ENSMUSG00000027737 | 78,25317607 | 175,521049 | 118,914836 | 190,78635 | 151,393287 | 122,0811681 | 156,753256 | 315,334314 | 149,486356 | 152,096208 | 94,9537446 | 618,3499 | 431,757036 | 307,7724 | 398,2475 | 331,8026678 | 1038,182 | 324,9992883 | 459,1236 | 199,9969 | 130,202 | 339,203016 |
| Upregulated | ENSMUSG00000078956 | 56,04619367 | 27,6578623 | 38,1253673 | 12,5263765 | 59,7939874 | 31,84726125 | 28,397329 | 40,3334587 | 51,2524649 | 9,70826858 | 35,3316259 | 159,4046 | 71,124709 | 30,87079 | 86,81795 | 76,56984641 | 228,3174 | 98,09429739 | 102,6612 | 71,33222 | 86,80135 | 73,5837231 |
| Upregulated | ENSMUSG00000119414 | 0 | 0 | 0 | 0 | 0 | 0 | 2,27178632 | 0 | 0 | 0 | 0 | 26,27549 | 2,00351293 | 14,96765 | 11,94742 | 4,557728953 | 25,64434 | 3,963405955 | 8,555098 | 7,999875 | 0,885728 | 7,17889982 |
| Upregulated | ENSMUSG00000024140 | 753,9799262 | 732,93335 | 1294,44699 | 639,80877 | 700,989087 | 1151,809282 | 1576,6197 | 1389,67099 | 1650,75647 | 1346,21324 | 2594,66628 | 7801,193 | 3670,43569 | 6118,964 | 4723,215 | 4959,720647 | 8920,095 | 3002,280011 | 5277,545 | 4733,259 | 1743,998 | 2560,17515 |
| Upregulated | ENSMUSG00000082243 | 5,287376762 | 3,1912918 | 0,90774684 | 1,92713485 | 0 | 2,653938437 | 6,81535895 | 2,44445204 | 2,13551937 | 6,47217906 | 2,20822662 | 20,14454 | 13,022834 | 15,90313 | 13,54041 | 14,58473265 | 23,98987 | 7,92681191 | 9,505665 | 4,666594 | 5,314368 | 7,17889982 |
| Upregulated | ENSMUSG00000020017 | 22,2069824 | 11,7014033 | 11,8007089 | 28,9070227 | 43,255225 | 46,44392265 | 35,2126879 | 6,11113011 | 17,084155 | 3,23608953 | 0 | 285,527 | 61,1071443 | 212,3536 | 164,078 | 217,859444 | 474,8339 | 124,8472876 | 92,20495 | 41,33269 | 94,7729 | 120,246572 |
| Upregulated | ENSMUSG00000018774 | 222,069824 | 106,376393 | 160,671191 | 41,4333992 | 89,0548749 | 132,6969219 | 80,6484142 | 80,6669174 | 69,4043796 | 36,6756813 | 93,8496312 | 513,2479 | 188,330215 | 287,1919 | 319,3945 | 376,4684115 | 624,5638 | 215,0147731 | 244,2956 | 120,6648 | 51,37222 | 88,8388853 |
| Upregulated | ENSMUSG00000032502 | 75,08075001 | 196,796328 | 21,7859242 | 336,285031 | 137,39895 | 10,61575375 | 61,3382305 | 22,0000684 | 12,8131162 | 78,7448452 | 112,619557 | 371,3603 | 158,277521 | 337,7077 | 552,7675 | 353,6797668 | 710,5964 | 249,6945752 | 376,4243 | 25,33294 | 42,51494 | 437,015527 |
| Upregulated | ENSMUSG00000016524 | 83,54055283 | 95,7387541 | 195,165571 | 23,1256182 | 29,2608875 | 84,92602999 | 104,502171 | 81,8891435 | 178,315868 | 66,8791836 | 160,09643 | 1594,922 | 1608,82088 | 2387,341 | 1044,205 | 1635,313148 | 3932,684 | 1019,586182 | 2017,102 | 299,3287 | 87,68707 | 1253,61538 |
| Upregulated | ENSMUSG00000017002 | 913,6587044 | 961,642596 | 1680,2394 | 1412,58984 | 136,126737 | 87,57996843 | 257,847747 | 138,11154 | 120,656844 | 291,248057 | 209,781529 | 4584,197 | 3001,26237 | 10973,16 | 7389,88 | 4627,917979 | 15689,37 | 2723,850743 | 11507,56 | 3135,284 | 1386,164 | 6519,3384 |
| Upregulated | ENSMUSG00000022231 | 859,7274614 | 1148,86505 | 861,451751 | 1643,84603 | 605,573149 | 301,2220126 | 798,53289 | 836,002599 | 742,092982 | 867,271993 | 853,479588 | 1931,249 | 1455,55214 | 2724,113 | 2010,353 | 2732,81428 | 3634,879 | 2107,541117 | 2605,503 | 1473,977 | 519,9223 | 1555,12917 |
| Upregulated | ENSMUSG00000071036 | 8,459802819 | 21,2752787 | 26,3246584 | 2,89070227 | 2,544425 | 10,61575375 | 5,67946579 | 12,2222602 | 17,084155 | 19,4165372 | 71,7673651 | 72,69552 | 87,1528124 | 109,451 | 94,7829 | 44,66574374 | 180,3376 | 40,62491104 | 55,13286 | 36,66609 | 19,48602 | 60,123286 |
| Upregulated | ENSMUSG00000035270 | 2,114950705 | 5,31881967 | 9,98521524 | 11,5628091 | 21,6276125 | 9,28878453 | 3,40767947 | 1,22222602 | 29,8972712 | 12,9443581 | 45,2686457 | 49,04758 | 43,075528 | 72,03184 | 26,28433 | 20,96555318 | 77,76026 | 38,64320806 | 9,505665 | 21,333 | 8,85728 | 23,3314244 |
| Upregulated | ENSMUSG00000082449 | 1,057475352 | 12,7651672 | 0,90774684 | 0,96356742 | 5,08884999 | 10,61575375 | 6,81535895 | 3,66667807 | 7,4743178 | 8,62957207 | 9,93701978 | 24,52379 | 12,0210776 | 28,06435 | 9,557939 | 18,23091581 | 30,60776 | 17,8353268 | 15,20906 | 6,666563 | 15,05738 | 13,4604372 |
| Upregulated | ENSMUSG00000083945 | 34,89668663 | 76,5910032 | 43,5718483 | 44,3241015 | 129,765675 | 124,7351066 | 68,1535895 | 39,1112327 | 49,1169455 | 36,6756813 | 98,2660845 | 214,5832 | 171,300355 | 274,0952 | 153,7235 | 243,3827261 | 366,4659 | 158,5362382 | 253,8013 | 107,9983 | 143,4879 | 123,836022 |
| Upregulated | ENSMUSG00000005800 | 71,90832396 | 52,1244328 | 110,745114 | 94,4296076 | 72,5161124 | 27,86635359 | 35,2126879 | 30,5556505 | 16,0163953 | 112,184437 | 101,578424 | 255,7481 | 158,277521 | 344,256 | 108,3233 | 174,105246 | 455,8075 | 444,8923185 | 273,7632 | 87,33197 | 62,00096 | 157,935796 |
| Upregulated | ENSMUSG00000031169 | 473,7489578 | 399,975239 | 324,973369 | 395,062644 | 343,497375 | 331,7423047 | 407,785644 | 293,334245 | 297,904952 | 345,182883 | 544,327861 | 882,8565 | 598,048609 | 936,4139 | 557,5465 | 817,6565742 | 1145,723 | 930,409548 | 993,342 | 525,3251 | 452,607 | 569,825173 |
| Upregulated | ENSMUSG00000041245 | 2,114950705 | 4,25505574 | 4,5387342 | 0,96356742 | 11,4499125 | 2,653938437 | 0 | 3,66667807 | 10,6775969 | 50,6987359 | 2,20822662 | 23,64794 | 14,0245905 | 18,70957 | 11,94742 | 19,1424616 | 29,78053 | 30,71639615 | 29,46756 | 21,333 | 15,05738 | 18,844612 |
| Upregulated | ENSMUSG00000022860 | 0 | 10,6376393 | 20,8781773 | 46,2512364 | 3,8166375 | 18,57756906 | 4,54357263 | 2,44445204 | 13,8808759 | 139,15185 | 24,2904928 | 28,02719 | 22,0386422 | 0 | 23,09835 | 40,10801479 | 86,03263 | 74,31386166 | 40,87436 | 1,333313 | 7,971552 | 7,17889982 |
| Upregulated | ENSMUSG00000042256 | 12,68970423 | 61,6983082 | 7,26197472 | 9,63567424 | 3,8166375 | 11,94272297 | 9,08714527 | 9,77780817 | 20,287434 | 9,70826858 | 15,4575863 | 21,02039 | 21,0368858 | 46,77392 | 18,31938 | 62,89665955 | 62,87 | 55,48768337 | 57,98456 | 54,66581 | 4,42864 | 48,4575738 |
| Upregulated | ENSMUSG00000059974 | 1,057475352 | 11,7014033 | 0,90774684 | 0 | 1,2722125 | 0 | 0 | 1,22222602 | 0 | 19,4165372 | 2,20822662 | 0,87585 | 6,01053879 | 5,61287 | 5,575465 | 18,23091581 | 11,58132 | 15,85362382 | 10,45623 | 6,666563 | 1,771456 | 15,2551621 |
| Upregulated | ENSMUSG00000025857 | 19,03455634 | 10,6376393 | 13,6162026 | 27,9434553 | 6,36106249 | 42,463015 | 20,4460768 | 9,77780817 | 19,2196743 | 7,55087556 | 13,2493597 | 49,04758 | 69,1211961 | 81,38662 | 44,60372 | 33,72719425 | 33,08947 | 64,40534677 | 24,71473 | 29,99953 | 26,57184 | 17,0498871 |
| Upregulated | ENSMUSG00000114338 | 0 | 6,3825836 | 7,26197472 | 9,63567424 | 2,544425 | 22,55847672 | 13,6307179 | 7,33335613 | 12,8131162 | 8,62957207 | 7,72879316 | 14,0136 | 19,0333728 | 31,80627 | 15,9299 | 15,49627844 | 4,136184 | 12,88106935 | 8,555098 | 16,66641 | 7,085824 | 4,48681239 |
| Upregulated | ENSMUSG00000064010 | 8,459802819 | 2,12752787 | 0 | 3,8542697 | 0 | 1,326969219 | 1,13589316 | 0 | 1,06775969 | 0 | 3,31233993 | 17,51699 | 34,0597198 | 106,6445 | 9,557939 | 30,99255688 | 11,58132 | 29,72554466 | 0 | 0 | 11,51446 | 7,17889982 |
| Upregulated | ENSMUSG00000074115 | 109,9774366 | 121,269088 | 160,671191 | 502,018628 | 45,79965 | 43,78998421 | 20,4460768 | 69,6668832 | 112,114767 | 47,4626464 | 532,182615 | 5239,333 | 8923,64659 | 22601,16 | 5050,574 | 4441,051092 | 1693,354 | 254,6488326 | 826,0423 | 269,3291 | 1553,567 | 641,614171 |
| Upregulated | ENSMUSG00000057465 | 35,95416198 | 38,2955016 | 98,0366587 | 74,1946916 | 2,544425 | 37,15513812 | 3,40767947 | 26,8889725 | 30,9650309 | 3,23608953 | 242,904928 | 1440,773 | 3874,794 | 11354,84 | 1777,777 | 1778,425838 | 320,1407 | 85,21322804 | 165,3986 | 47,33259 | 410,0921 | 232,416882 |
| Upregulated | ENSMUSG00000040896 | 0 | 8,51011147 | 4,5387342 | 0,96356742 | 0 | 5,307876874 | 1,13589316 | 1,22222602 | 0 | 5,39348255 | 2,20822662 | 21,89624 | 6,01053879 | 65,48349 | 14,33691 | 7,292366325 | 4,136184 | 0,990851489 | 2,851699 | 4,666594 | 0 | 0,89736248 |
| Upregulated | ENSMUSG00000069792 | 11,63222888 | 61,6983082 | 59,9112914 | 81,903231 | 6,36106249 | 5,307876874 | 2,27178632 | 7,33335613 | 9,60983717 | 7,55087556 | 58,5180054 | 643,7495 | 320,562069 | 4578,231 | 915,1727 | 766,6100099 | 1139,932 | 317,0724764 | 446,7663 | 361,9944 | 207,2604 | 157,038434 |
| Upregulated | ENSMUSG00000042845 | 894,6241481 | 614,855554 | 1190,05611 | 2676,7903 | 388,024812 | 384,8210734 | 521,37496 | 366,667807 | 402,545401 | 334,395918 | 309,151726 | 3209,989 | 2812,93215 | 14469,98 | 5553,163 | 3064,616948 | 2761,317 | 1503,121709 | 2514,248 | 52,66584 | 447,2926 | 777,115905 |
| Upregulated | ENSMUSG00000040154 | 37,01163733 | 19,1477508 | 36,3098736 | 38,542697 | 0 | 0 | 11,3589316 | 3,66667807 | 2,13551937 | 25,8887162 | 9,93701978 | 70,94383 | 68,1194396 | 228,2567 | 157,706 | 125,7933191 | 46,32526 | 69,35960422 | 43,72606 | 1,333313 | 7,085824 | 8,0762623 |
| Upregulated | ENSMUSG00000054863 | 3,172426057 | 3,1912918 | 0 | 12,5263765 | 0 | 0 | 4,54357263 | 0 | 7,4743178 | 0 | 1,10411331 | 33,28229 | 14,0245905 | 85,12853 | 40,62124 | 41,01956058 | 8,272368 | 25,76213871 | 12,35736 | 11,33316 | 4,42864 | 4,48681239 |
| Upregulated | ENSMUSG00000024679 | 0 | 1,06376393 | 6,35422788 | 20,2349159 | 8,90548749 | 2,653938437 | 4,54357263 | 0 | 1,06775969 | 4,31478604 | 2,20822662 | 79,70232 | 88,1545689 | 234,8051 | 103,5443 | 74,74675483 | 25,64434 | 42,60661402 | 70,34192 | 2,666625 | 20,37174 | 26,9208743 |
| Upregulated | ENSMUSG00000054793 | 0 | 1,06376393 | 0,90774684 | 8,67210682 | 2,544425 | 14,5966614 | 2,27178632 | 0 | 2,13551937 | 8,62957207 | 8,83290647 | 12,2619 | 12,0210776 | 28,99983 | 19,91237 | 18,23091581 | 8,272368 | 3,963405955 | 0 | 3,333281 | 0,885728 | 2,69208743 |
| Upregulated | ENSMUSG00000087006 | 13,74717958 | 10,6376393 | 39,940861 | 17,3442136 | 27,988675 | 34,50119968 | 52,2510853 | 36,6667807 | 35,2360696 | 29,1248057 | 96,0578579 | 80,57817 | 33,0579633 | 183,3538 | 83,63197 | 99,35849118 | 63,69724 | 24,77128722 | 33,26983 | 21,99966 | 23,02893 | 26,0235118 |
| Upregulated | ENSMUSG00000016528 | 454,7144015 | 345,723279 | 364,91423 | 412,406857 | 412,19685 | 569,2697948 | 516,831387 | 426,556882 | 737,821943 | 433,635997 | 713,257197 | 770,7477 | 913,601896 | 2117,923 | 911,9867 | 926,1305233 | 861,9808 | 764,9373494 | 632,1267 | 422,6601 | 553,58 | 656,869333 |
| Upregulated | ENSMUSG00000008730 | 903,0839509 | 774,420144 | 587,312205 | 766,99967 | 1100,46381 | 1195,599266 | 939,383642 | 1130,55907 | 1336,83513 | 939,544659 | 1489,44885 | 1909,352 | 1699,98072 | 3459,399 | 1788,928 | 2143,044154 | 1365,768 | 1638,868362 | 1442,96 | 1386,645 | 1049,588 | 1547,05291 |
| Upregulated | ENSMUSG00000024789 | 440,9672219 | 429,760629 | 366,729723 | 282,325255 | 320,59755 | 505,5752723 | 455,493156 | 469,334792 | 654,536687 | 379,701171 | 694,487271 | 1079,923 | 645,131163 | 1815,764 | 874,5515 | 966,2385381 | 840,4726 | 789,7086366 | 399,2379 | 841,3202 | 426,0352 | 476,499476 |
| Upregulated | ENSMUSG00000028755 | 11,63222888 | 49,9969049 | 19,9704305 | 154,170788 | 34,3497375 | 15,92363062 | 32,9409016 | 12,2222602 | 16,0163953 | 24,8100197 | 16,5616996 | 171,6665 | 92,1615947 | 359,2237 | 119,4742 | 123,0586817 | 172,0653 | 74,31386166 | 111,2163 | 89,33194 | 62,00096 | 76,2758106 |
| Upregulated | ENSMUSG00000043155 | 1,057475352 | 4,25505574 | 0 | 5,78140454 | 1,2722125 | 5,307876874 | 4,54357263 | 2,44445204 | 0 | 11,8656616 | 1,10411331 | 17,51699 | 8,01405172 | 25,25792 | 9,557939 | 14,58473265 | 14,89026 | 6,935960422 | 10,45623 | 7,333219 | 6,200096 | 6,28153734 |
| Upregulated | ENSMUSG00000092203 | 189,2880881 | 129,7792 | 206,96628 | 171,515001 | 236,631525 | 264,0668745 | 310,098832 | 220,000684 | 264,804402 | 167,197959 | 237,384361 | 475,5864 | 411,721907 | 819,4791 | 439,6652 | 371,9106826 | 433,4721 | 327,9718428 | 382,1277 | 466,6594 | 356,9484 | 318,563679 |
| Upregulated | ENSMUSG00000048234 | 92,00035565 | 45,7418492 | 66,2655193 | 51,0690735 | 62,3384124 | 78,2911839 | 72,6971621 | 59,8890751 | 37,371589 | 61,485701 | 144,638843 | 278,5202 | 149,261713 | 346,127 | 169,6534 | 112,1201322 | 153,8661 | 116,9204757 | 117,8702 | 178,6639 | 96,54435 | 120,246572 |
| Upregulated | ENSMUSG00000032350 | 151,2189754 | 145,735659 | 138,885267 | 171,515001 | 319,325337 | 408,7065193 | 264,663106 | 301,889827 | 268,007681 | 235,155839 | 450,47823 | 951,1728 | 599,050366 | 1136,606 | 688,1716 | 491,3231812 | 335,0309 | 496,4165959 | 288,9722 | 219,9966 | 286,9759 | 372,405428 |
| Upregulated | ENSMUSG00000038068 | 0 | 1,06376393 | 1,81549368 | 5,78140454 | 13,9943375 | 23,88544594 | 1,13589316 | 6,11113011 | 1,06775969 | 1,07869651 | 24,2904928 | 6,130948 | 8,01405172 | 44,90296 | 0 | 15,49627844 | 0 | 0,990851489 | 1,901133 | 8,666531 | 6,200096 | 17,0498871 |
| Upregulated | ENSMUSG00000071226 | 3,172426057 | 13,8289311 | 7,26197472 | 13,4899439 | 49,6162874 | 22,55847672 | 17,0383974 | 34,2223286 | 25,6262325 | 20,4952337 | 24,2904928 | 13,13775 | 19,0333728 | 81,38662 | 26,28433 | 62,89665955 | 18,19921 | 25,76213871 | 24,71473 | 20,66634 | 7,971552 | 15,2551621 |
| Upregulated | ENSMUSG00000021596 | 11,63222888 | 49,9969049 | 15,4316963 | 58,7776129 | 83,9660249 | 17,25059984 | 53,3869784 | 123,444828 | 76,8786974 | 56,0922185 | 72,8714784 | 76,19892 | 24,0421551 | 296,5466 | 130,6252 | 71,10057167 | 58,73382 | 78,27726762 | 106,4634 | 7,333219 | 14,17165 | 83,4547104 |
| Upregulated | ENSMUSG00000043811 | 57,10366903 | 9,57387541 | 34,4943799 | 54,9233432 | 45,79965 | 42,463015 | 47,7075126 | 58,666849 | 49,1169455 | 50,6987359 | 83,9126115 | 40,28909 | 33,0579633 | 110,3864 | 39,02825 | 80,21602958 | 28,95329 | 47,56087146 | 26,61586 | 11,99981 | 12,40019 | 23,3314244 |
| Upregulated | ENSMUSG00000066902 | 27,49435916 | 73,3997114 | 38,1253673 | 32,7612924 | 10,1777 | 23,88544594 | 18,1742905 | 30,5556505 | 55,5235037 | 15,1017511 | 24,2904928 | 31,53059 | 43,075528 | 57,99966 | 51,77217 | 25,52328214 | 61,21553 | 36,66150509 | 49,42946 | 37,33275 | 40,74349 | 12,5630747 |
| Upregulated | ENSMUSG00000021071 | 365,8864719 | 313,810361 | 177,010634 | 244,746126 | 76,3327499 | 46,44392265 | 54,5228716 | 95,3336297 | 73,6754183 | 22,6526267 | 107,098991 | 330,1953 | 443,778114 | 188,9666 | 311,4295 | 265,2598251 | 244,0349 | 114,9387727 | 243,345 | 101,3318 | 15,9431 | 354,458179 |
| Upregulated | ENSMUSG00000026527 | 204,092743 | 232,964302 | 107,114127 | 14,4535114 | 7,63327499 | 2,653938437 | 3,40767947 | 3,66667807 | 3,20327906 | 9,70826858 | 50,7892122 | 151,522 | 207,363588 | 111,3219 | 104,3408 | 110,2970407 | 48,80697 | 16,84447531 | 46,57776 | 0,666656 | 0,885728 | 23,3314244 |
| Upregulated | ENSMUSG00000043263 | 94,11530636 | 54,2519606 | 53,5570636 | 0 | 2,544425 | 0 | 0 | 0 | 0 | 0 | 1,10411331 | 14,88944 | 33,0579633 | 46,77392 | 34,24928 | 29,1694653 | 43,84355 | 6,935960422 | 38,97323 | 3,333281 | 5,314368 | 11,6657122 |
| Upregulated | ENSMUSG00000057829 | 8,459802819 | 6,3825836 | 32,6788862 | 46,2512364 | 80,1493874 | 128,7160142 | 69,2894827 | 45,2223628 | 62,9978215 | 45,3052534 | 73,9755917 | 70,06798 | 67,1176831 | 68,28992 | 54,95815 | 104,8277659 | 76,93303 | 49,54257444 | 67,49022 | 46,66594 | 39,85776 | 21,5366995 |
| Upregulated | ENSMUSG00000093548 | 1,057475352 | 3,1912918 | 11,8007089 | 5,78140454 | 16,5387625 | 42,463015 | 34,0767947 | 14,6667123 | 24,5584728 | 11,8656616 | 25,3946061 | 19,26869 | 33,0579633 | 43,03201 | 15,9299 | 25,52328214 | 32,26224 | 14,86277233 | 14,2585 | 8,666531 | 14,17165 | 12,5630747 |
| Upregulated | ENSMUSG00000083553 | 1,057475352 | 5,31881967 | 5,44648104 | 4,81783712 | 11,4499125 | 22,55847672 | 12,4948247 | 9,77780817 | 17,084155 | 2,15739302 | 6,62467985 | 8,758497 | 13,022834 | 14,96765 | 13,54041 | 21,87709898 | 22,33539 | 17,8353268 | 3,802266 | 17,99972 | 15,05738 | 7,17889982 |
| Upregulated | ENSMUSG00000066620 | 7,402327466 | 11,7014033 | 17,24719 | 36,6155621 | 62,3384124 | 99,5226914 | 44,2998332 | 28,1111985 | 28,8295115 | 26,9674127 | 29,8110593 | 43,79248 | 64,1124137 | 52,38679 | 31,0633 | 71,10057167 | 76,10579 | 31,70724764 | 57,98456 | 69,33225 | 77,94407 | 13,4604372 |
| Upregulated | ENSMUSG00000081675 | 3,172426057 | 2,12752787 | 5,44648104 | 0,96356742 | 2,544425 | 13,26969219 | 17,0383974 | 3,66667807 | 7,4743178 | 10,7869651 | 6,62467985 | 7,882647 | 11,0193211 | 14,03218 | 19,91237 | 10,93854949 | 13,23579 | 1,981702978 | 6,653965 | 13,99978 | 10,62874 | 7,17889982 |
| Upregulated | ENSMUSG00000094238 | 5,287376762 | 18,0839869 | 6,35422788 | 7,70853939 | 12,722125 | 45,11695343 | 12,4948247 | 15,8889383 | 25,6262325 | 10,7869651 | 25,3946061 | 27,15134 | 20,0351293 | 57,06418 | 25,48784 | 30,99255688 | 28,95329 | 7,92681191 | 39,92379 | 33,33281 | 38,97203 | 14,3577996 |
| Upregulated | ENSMUSG00000080811 | 21,14950705 | 45,7418492 | 37,2176204 | 37,5791295 | 72,5161124 | 124,7351066 | 47,7075126 | 53,777945 | 54,455744 | 24,8100197 | 54,1015521 | 77,07477 | 81,1422736 | 169,3216 | 74,07403 | 103,9162201 | 86,85987 | 71,34130719 | 83,64985 | 102,6651 | 114,2589 | 43,0733989 |
| Upregulated | ENSMUSG00000080054 | 24,3219331 | 9,57387541 | 19,9704305 | 15,4170788 | 29,2608875 | 61,04058406 | 20,4460768 | 35,4445546 | 38,4393487 | 9,70826858 | 28,706946 | 23,64794 | 48,0843103 | 87,93497 | 39,82475 | 55,60429323 | 83,55092 | 18,82617829 | 60,83626 | 78,66544 | 62,88669 | 25,1261494 |
| Upregulated | ENSMUSG00000021262 | 13,74717958 | 7,44634754 | 15,4316963 | 10,5992417 | 24,1720375 | 49,09786109 | 32,9409016 | 2,44445204 | 19,2196743 | 12,9443581 | 28,706946 | 27,15134 | 17,0298599 | 66,41897 | 29,47031 | 30,08101109 | 33,91671 | 23,78043573 | 32,31926 | 65,99897 | 32,77194 | 34,0997741 |
| Upregulated | ENSMUSG00000026620 | 82,48307748 | 107,440157 | 52,6493167 | 99,2474447 | 132,3101 | 216,2959826 | 106,773957 | 112,444794 | 154,825154 | 102,476168 | 187,699262 | 254,8723 | 117,205506 | 366,7075 | 254,8784 | 221,5056271 | 185,3011 | 184,2983769 | 138,7827 | 443,3264 | 149,688 | 132,809647 |
| Upregulated | ENSMUSG00000081076 | 13,74717958 | 11,7014033 | 9,0774684 | 11,5628091 | 21,6276125 | 33,17423047 | 27,2614358 | 11,0000342 | 18,1519147 | 12,9443581 | 16,5616996 | 19,26869 | 30,0526939 | 41,16105 | 56,55114 | 48,3119269 | 52,11592 | 41,61576253 | 43,72606 | 47,99925 | 72,6297 | 21,5366995 |
| Upregulated | ENSMUSG00000114688 | 21,14950705 | 11,7014033 | 26,3246584 | 67,4497197 | 47,0718625 | 67,67543015 | 48,8434058 | 22,0000684 | 70,4721393 | 39,9117708 | 54,1015521 | 70,06798 | 90,1580818 | 120,6767 | 122,6602 | 91,15457906 | 96,78671 | 83,23152506 | 82,69928 | 119,3315 | 171,8312 | 122,041297 |
| Upregulated | ENSMUSG00000081695 | 19,03455634 | 27,6578623 | 29,0478989 | 83,8303659 | 97,9603624 | 141,9857064 | 95,4150253 | 40,3334587 | 121,724604 | 83,0596312 | 110,411331 | 124,3707 | 231,405743 | 243,2244 | 243,7275 | 195,0707992 | 223,3539 | 123,8564361 | 199,619 | 235,3297 | 395,0347 | 218,059082 |
| Upregulated | ENSMUSG00000032518 | 318,300081 | 381,891252 | 373,083951 | 1183,2608 | 1183,15762 | 1277,871358 | 950,742573 | 512,112703 | 1282,37938 | 902,868978 | 1178,0889 | 1312,899 | 2069,62886 | 2774,629 | 2517,721 | 1969,850454 | 2188,041 | 1547,710026 | 1728,13 | 2463,295 | 3738,658 | 2142,9016 |
| Upregulated | ENSMUSG00000045055 | 291,8631972 | 311,682833 | 307,726179 | 739,056214 | 656,461649 | 727,1791318 | 594,072122 | 363,001128 | 745,296261 | 476,783857 | 659,155645 | 775,127 | 1399,45378 | 1517,346 | 1330,147 | 1181,363345 | 1177,158 | 985,8972314 | 1096,003 | 1363,979 | 2301,121 | 1228,48923 |
| Upregulated | ENSMUSG00000082128 | 9,517278171 | 19,1477508 | 7,26197472 | 56,850478 | 35,62195 | 78,2911839 | 46,5716195 | 14,6667123 | 48,0491859 | 31,2821988 | 35,3316259 | 45,54418 | 80,1405172 | 102,9026 | 121,0672 | 88,41994169 | 133,1851 | 51,52427742 | 67,49022 | 119,3315 | 195,7459 | 83,4547104 |

| | | | | | | | | | | | | | | | | | | | | | | | |
|---|---|---|---|---|---|---|---|---|---|---|---|---|---|---|---|---|---|---|---|---|---|---|---|
| Upregulated | ENSMUSG00000082826 | 2,114950705 | 1,06376393 | 3,63098736 | 16,3806462 | 3,8166375 | 10,61575375 | 11,3589316 | 3,66667807 | 3,20327906 | 1,07869651 | 2,20822662 | 7,006798 | 4,00702586 | 25,25792 | 14,33691 | 10,0270037 | 20,68092 | 10,89936638 | 15,20906 | 23,33297 | 25,68611 | 8,97362477 |
| Upregulated | ENSMUSG00000083378 | 4,229901409 | 9,57387541 | 14,5239494 | 20,2349159 | 15,26655 | 27,86635359 | 19,3101837 | 8,55558215 | 12,8131162 | 4,31478604 | 5,52056654 | 11,38605 | 20,0351293 | 41,16105 | 23,89485 | 18,23091581 | 18,19921 | 22,78958424 | 19,9619 | 25,33294 | 47,82931 | 18,844612 |
| Upregulated | ENSMUSG00000041220 | 120,5521902 | 45,7418492 | 126,176811 | 204,276294 | 63,6106249 | 61,04058406 | 86,32788 | 151,556027 | 164,434992 | 174,748834 | 199,844509 | 197,942 | 523,918631 | 517,3195 | 141,7761 | 108,4739491 | 308,5593 | 286,3560803 | 239,5428 | 253,996 | 519,0366 | 203,701282 |
| Upregulated | ENSMUSG00000022747 | 5,287376762 | 18,0839869 | 10,8929621 | 27,9434553 | 6,36106249 | 13,26969219 | 5,67946579 | 26,8889725 | 26,6939921 | 9,70826858 | 32,019286 | 28,90304 | 49,0860668 | 39,29009 | 14,33691 | 5,469274744 | 38,05289 | 43,59746551 | 18,06076 | 49,33256 | 48,71504 | 46,6628488 |
| Upregulated | ENSMUSG00000037095 | 72,96579931 | 36,1679738 | 101,667646 | 198,494889 | 13,9943375 | 2,653938437 | 53,3869784 | 15,8889383 | 26,6939921 | 5,39348255 | 68,4550251 | 1233,196 | 3457,06156 | 2642,726 | 741,5368 | 1162,220883 | 3000,388 | 705,48626 | 1487,637 | 1013,318 | 2111,576 | 4228,37199 |
| Upregulated | ENSMUSG00000046805 | 16,91960564 | 10,6376393 | 17,24719 | 44,3241015 | 10,1777 | 9,28878453 | 22,7178632 | 9,77780817 | 25,6262325 | 24,8100197 | 7,72879316 | 44,66833 | 175,307381 | 124,4186 | 30,26681 | 100,270037 | 114,1587 | 37,65235657 | 33,26983 | 15,99975 | 105,4016 | 66,4048233 |
| Upregulated | ENSMUSG00000117604 | 46,5289155 | 23,4028065 | 28,140152 | 24,0891856 | 44,5274375 | 26,53938437 | 21,58197 | 37,8890067 | 41,6426277 | 22,6526267 | 73,9755917 | 181,3009 | 487,855398 | 164,6442 | 125,0497 | 195,982345 | 190,2645 | 169,4356046 | 188,2122 | 84,66535 | 384,406 | 242,287869 |
| Upregulated | ENSMUSG00000037820 | 618,6230811 | 456,354728 | 568,249522 | 641,735904 | 226,453825 | 153,9284294 | 272,614358 | 552,446162 | 384,393487 | 539,348255 | 806,002715 | 1351,436 | 2786,88648 | 988,8007 | 539,2271 | 937,0690728 | 891,7613 | 485,5172295 | 1247,143 | 627,3235 | 2338,322 | 1663,71003 |
| Upregulated | ENSMUSG00000118283 | 93,057831 | 126,587908 | 107,114127 | 105,992417 | 73,7883249 | 80,94512234 | 102,230384 | 102,666986 | 74,743178 | 90,6105068 | 105,994878 | 202,3213 | 311,54626 | 271,2887 | 140,9796 | 171,3706086 | 151,3843 | 152,5911293 | 243,345 | 185,3304 | 422,4923 | 206,39337 |
| Upregulated | ENSMUSG00000020077 | 33,83921127 | 29,7853902 | 40,8486078 | 79,0125288 | 67,4272624 | 83,59906077 | 45,4357263 | 116,111472 | 83,2852555 | 89,5318103 | 128,077144 | 183,0526 | 1227,15167 | 639,8672 | 150,5375 | 142,2011433 | 224,1812 | 133,764951 | 222,4326 | 434,6599 | 1399,45 | 587,772423 |
| Upregulated | ENSMUSG00000115602 | 135,3568451 | 130,842964 | 177,918381 | 163,806462 | 156,482137 | 107,4845067 | 156,753256 | 162,556061 | 222,094015 | 103,554865 | 131,389484 | 303,9198 | 578,01348 | 299,3531 | 293,1101 | 299,8985651 | 368,1204 | 267,529902 | 284,2194 | 366,6609 | 723,6398 | 370,610703 |
| Upregulated | ENSMUSG00000035376 | 269,6562148 | 332,958111 | 448,426939 | 403,734751 | 291,336662 | 277,3365667 | 281,701503 | 380,112293 | 478,356339 | 229,762356 | 244,009041 | 520,2547 | 1242,17802 | 608,9964 | 577,4588 | 556,0429323 | 770,9847 | 524,1604376 | 631,1762 | 729,9886 | 1472,08 | 841,726004 |
| Upregulated | ENSMUSG00000106720 | 4,229901409 | 0 | 0,90774684 | 0 | 0 | 2,653938437 | 9,08714527 | 7,33335613 | 2,13551937 | 7,55087556 | 0 | 0,87585 | 19,0333728 | 15,90313 | 6,37196 | 21,87709898 | 0 | 22,78958424 | 13,30793 | 1,999969 | 7,085824 | 17,0498871 |
| Upregulated | ENSMUSG00000101609 | 258,023986 | 342,531987 | 183,364862 | 264,017474 | 422,37455 | 485,670734 | 338,496161 | 542,668354 | 288,295115 | 179,063621 | 361,045052 | 451,0626 | 1701,98423 | 1076,736 | 543,2096 | 398,3455105 | 332,5492 | 762,9556464 | 394,4851 | 285,3289 | 201,946 | 459,449588 |
| Upregulated | ENSMUSG00000113650 | 1,057475352 | 3,1912918 | 0 | 1,92713485 | 3,8166375 | 2,653938437 | 0 | 0 | 2,13551937 | 0 | 0 | 2,627549 | 35,0614763 | 17,77409 | 5,575465 | 4,557728953 | 8,272368 | 26,7529902 | 3,802266 | 3,333281 | 0 | 5,38417486 |
| Upregulated | ENSMUSG00000101414 | 63,44852114 | 55,3157246 | 36,3098736 | 39,5062644 | 125,949037 | 138,0047987 | 55,6587648 | 72,1113353 | 87,5562942 | 43,1478604 | 102,682538 | 87,58497 | 282,495323 | 197,3859 | 54,16166 | 116,6778612 | 154,6933 | 223,9324365 | 153,9918 | 31,33284 | 46,05786 | 148,962171 |
| Upregulated | ENSMUSG00000096433 | 43,35648945 | 40,4230295 | 29,0478989 | 30,8341576 | 19,0831875 | 19,90453828 | 10,2230384 | 36,6667807 | 4,27103874 | 21,5739302 | 17,6658129 | 50,79928 | 46,0807974 | 58,93514 | 30,26681 | 35,55028583 | 62,04276 | 43,59746551 | 50,38002 | 5,33325 | 42,51494 | 68,1995483 |
| Upregulated | ENSMUSG00000097125 | 37,01163733 | 27,6578623 | 29,0478989 | 23,1256182 | 17,810975 | 11,94272297 | 18,1742905 | 18,3333903 | 10,6775969 | 23,7313232 | 20,9781529 | 83,20572 | 80,1405172 | 57,06418 | 9,557939 | 20,05400739 | 71,96961 | 71,34130719 | 96,00722 | 19,33303 | 18,60029 | 32,3050492 |
| Upregulated | ENSMUSG00000028494 | 4217,211705 | 2578,56378 | 3190,73014 | 1224,6942 | 1203,51302 | 1330,950126 | 1430,08949 | 1450,78229 | 1377,40999 | 1107,82131 | 2167,37443 | 3673,314 | 7029,32511 | 5121,744 | 2633,212 | 2922,415805 | 4590,337 | 3146,944328 | 6057,96 | 2737,291 | 2338,322 | 4734,48443 |
| Upregulated | ENSMUSG00000044288 | 100,4601585 | 71,2721836 | 68,081013 | 21,1984833 | 12,722125 | 18,57756906 | 6,81535895 | 40,3334587 | 17,084155 | 10,7869651 | 24,2904928 | 137,5084 | 192,337241 | 52,38679 | 34,24928 | 77,4813922 | 90,99605 | 120,8838816 | 159,6952 | 73,33219 | 93,00144 | 140,885909 |
| Upregulated | ENSMUSG00000050345 | 9,517278171 | 0 | 1,81549368 | 3,8542697 | 0 | 1,326969219 | 1,13589316 | 0 | 1,06775969 | 0 | 0 | 2,627549 | 13,022834 | 21,516 | 0 | 0 | 31,435 | 6,935960422 | 19,01133 | 3,999938 | 6,200096 | 1,79472495 |
| Upregulated | ENSMUSG00000074183 | 6,344852114 | 5,31881967 | 1,81549368 | 0,96356742 | 0 | 0 | 0 | 0 | 1,06775969 | 0 | 0 | 23,64794 | 10,0175646 | 38,35461 | 22,30186 | 8,203912116 | 38,05289 | 20,80788127 | 51,33059 | 1,999969 | 17,71456 | 13,4604372 |
| Upregulated | ENSMUSG00000111709 | 2,114950705 | 6,3825836 | 0 | 5,78140454 | 0 | 3,980907656 | 0 | 1,22222602 | 0 | 1,07869651 | 0 | 13,13775 | 14,0245905 | 23,38696 | 15,1334 | 7,292366325 | 25,64434 | 32,69809913 | 31,36869 | 4,666594 | 10,62874 | 17,0498871 |
| Upregulated | ENSMUSG00000031654 | 2,114950705 | 0 | 0,90774684 | 9,63567424 | 0 | 0 | 1,13589316 | 0 | 0 | 0 | 1,10411331 | 14,88944 | 2,00351293 | 50,51583 | 26,28433 | 1,823091581 | 38,05289 | 48,55172295 | 62,73739 | 17,99972 | 4,42864 | 7,17889982 |
| Upregulated | ENSMUSG00000091844 | 7,402327466 | 1,06376393 | 8,16972156 | 5,78140454 | 17,810975 | 7,961815312 | 10,2230384 | 8,55558215 | 10,6775969 | 14,0230546 | 6,62467985 | 16,64114 | 18,0316164 | 61,74157 | 6,37196 | 9,115457906 | 24,81711 | 24,77128722 | 20,91246 | 34,66613 | 4,42864 | 19,7419745 |
| Upregulated | ENSMUSG00000111865 | 10,57475352 | 6,3825836 | 5,44648104 | 8,67210682 | 8,90548749 | 21,2315075 | 3,40767947 | 8,55558215 | 9,60983717 | 3,23608953 | 4,41645324 | 9,634347 | 28,049181 | 50,51583 | 15,9299 | 21,87709898 | 39,70737 | 22,78958424 | 11,4068 | 3,333281 | 8,85728 | 0 |
| Upregulated | ENSMUSG00000025491 | 232,6445775 | 146,799423 | 293,202229 | 347,84784 | 188,28745 | 321,1265509 | 298,739901 | 234,667396 | 222,094015 | 378,622475 | 432,812417 | 718,1967 | 462,811487 | 1986,021 | 681,0032 | 532,3427417 | 1731,407 | 670,8064579 | 700,5675 | 983,9846 | 533,2083 | 454,962776 |
| Upregulated | ENSMUSG00000071042 | 0 | 5,31881967 | 3,63098736 | 37,5791295 | 24,1720375 | 22,55847672 | 7,95125211 | 3,66667807 | 9,60983717 | 9,70826858 | 0 | 8,758497 | 1,00175646 | 361,0947 | 136,2006 | 63,80820534 | 199,3641 | 58,46023784 | 50,38002 | 22,66631 | 38,0863 | 33,2024117 |
| Upregulated | ENSMUSG00000045008 | 10,57475352 | 8,51011147 | 14,5239494 | 5,78140454 | 1,2722125 | 1,326969219 | 5,67946579 | 7,33335613 | 27,7617518 | 22,6526267 | 9,93701978 | 8,758497 | 8,01405172 | 30,87079 | 30,26681 | 21,87709898 | 21,50816 | 14,86277233 | 29,46756 | 20,66634 | 29,22902 | 17,9472495 |
| Upregulated | ENSMUSG00000087366 | 10,57475352 | 8,51011147 | 9,0774684 | 10,5992417 | 1,2722125 | 5,307876874 | 1,13589316 | 9,77780817 | 11,7453565 | 16,1804476 | 14,353473 | 13,13775 | 7,01229525 | 21,516 | 9,557939 | 8,203912116 | 26,47158 | 9,908514888 | 14,2585 | 21,99966 | 19,48602 | 21,5366995 |
| Upregulated | ENSMUSG00000102525 | 27,49435916 | 37,2317377 | 130,715545 | 115,628091 | 86,5104499 | 54,40573796 | 68,1535895 | 262,778595 | 238,11041 | 111,10574 | 152,367637 | 274,141 | 229,40223 | 239,4825 | 234,966 | 206,0093487 | 277,9516 | 264,5573475 | 223,3831 | 89,33194 | 242,6895 | 201,906557 |
| Upregulated | ENSMUSG00000032193 | 1579,868176 | 1480,7594 | 3186,19141 | 4200,1904 | 1778,55307 | 1967,895351 | 1745,86778 | 3809,67851 | 2653,38282 | 2677,32474 | 2659,80896 | 6453,261 | 4987,74544 | 4365,878 | 4603,741 | 5924,136093 | 7981,181 | 5579,484733 | 5654,92 | 2074,634 | 5495,057 | 3897,24524 |
| Upregulated | ENSMUSG00000069713 | 1,057475352 | 1,06376393 | 5,44648104 | 7,70853939 | 1,2722125 | 0 | 1,13589316 | 4,88890409 | 5,33879843 | 4,31478604 | 7,72879316 | 4,379248 | 7,01229525 | 7,483827 | 12,74392 | 10,0270037 | 16,54474 | 9,908514888 | 15,20906 | 6,666563 | 7,971552 | 4,48681239 |
| Upregulated | ENSMUSG00000032018 | 303,4954261 | 457,418492 | 650,854484 | 695,69568 | 381,66375 | 453,8234728 | 447,541904 | 910,558386 | 650,265649 | 596,51917 | 628,240473 | 966,0622 | 699,226012 | 1062,703 | 935,0851 | 775,7254678 | 1307,861 | 1550,68258 | 941,0608 | 604,6572 | 1036,302 | 722,376794 |
| Upregulated | ENSMUSG00000026675 | 297,150574 | 257,430872 | 269,600811 | 483,710847 | 293,881087 | 399,4177348 | 350,990986 | 529,223867 | 469,814262 | 435,79339 | 313,56818 | 696,3005 | 686,203178 | 673,5444 | 632,417 | 508,6425512 | 1207,766 | 942,2997659 | 858,3615 | 700,6557 | 679,3534 | 611,103847 |
| Upregulated | ENSMUSG00000021273 | 189,2880881 | 204,242675 | 255,984609 | 575,249752 | 152,6655 | 141,9857064 | 116,996995 | 232,222944 | 215,687457 | 222,211481 | 79,4961582 | 703,3073 | 618,083739 | 1440,637 | 1041,019 | 711,9172625 | 1547,76 | 1361,429946 | 920,1484 | 625,3236 | 1030,987 | 480,088925 |
| Upregulated | ENSMUSG00000035129 | 21,14950705 | 36,1679738 | 27,2324052 | 68,4132871 | 20,3554 | 17,25059984 | 9,08714527 | 33,0001026 | 25,6262325 | 20,4952337 | 16,5616996 | 86,70912 | 83,1457866 | 169,3216 | 114,6953 | 72,92366325 | 221,6995 | 161,5087927 | 123,5736 | 86,66531 | 113,3732 | 51,1496612 |
| Upregulated | ENSMUSG00000024640 | 425,1050916 | 643,57718 | 559,172053 | 433,605341 | 818,032637 | 530,7876874 | 868,958266 | 1070,67 | 826,445997 | 400,196405 | 559,785448 | 1452,159 | 1818,18798 | 1235,767 | 1522,102 | 1380,991873 | 1493,163 | 1452,588283 | 1242,39 | 1524,643 | 1481,823 | 1729,21749 |
| Upregulated | ENSMUSG00000078768 | 7,402327466 | 19,1477508 | 31,7711394 | 15,4170788 | 27,988675 | 7,961815312 | 26,1255426 | 44,0001368 | 26,6939921 | 29,1248057 | 14,353473 | 56,05438 | 70,1229525 | 63,61253 | 38,23176 | 43,75419795 | 74,45132 | 54,49683188 | 39,92379 | 41,33269 | 32,77194 | 59,2259235 |
| Upregulated | ENSMUSG00000003541 | 9492,956238 | 11805,6521 | 11788,9082 | 10716,7969 | 21751,0171 | 14527,65901 | 14172,5389 | 21522,178 | 31601,4157 | 12941,122 | 16333,1482 | 32803,2 | 27463,1535 | 29391,8 | 34244,5 | 31534,01508 | 35479,36 | 28737,66573 | 33118,69 | 30874,18 | 34918,06 | 21439,7843 |
| Upregulated | ENSMUSG00000032487 | 4065,99273 | 9693,01697 | 6947,89431 | 5689,86564 | 5432,34737 | 5621,04161 | 4734,40268 | 24088,8527 | 22775,3141 | 6886,39851 | 6486,66569 | 39813,5 | 21150,0842 | 19878,92 | 28116,27 | 45464,25785 | 40273,2 | 29022,04011 | 26036,02 | 30000,2 | 39475,13 | 17905,9709 |
| Upregulated | ENSMUSG00000113689 | 0 | 12,7651672 | 0 | 1,92713485 | 5,08884999 | 5,307876874 | 2,27178632 | 11,0000342 | 11,7453565 | 0 | 6,62467985 | 21,89624 | 15,026347 | 24,32244 | 11,15093 | 28,25791951 | 52,11592 | 27,74384169 | 14,2585 | 20,66634 | 50,4865 | 16,1525246 |
| Upregulated | ENSMUSG00000001467 | 429,334993 | 670,171278 | 890,49965 | 1388,50066 | 548,323587 | 427,2840884 | 516,831387 | 1485,00462 | 1021,84602 | 838,147188 | 740,86003 | 2362,167 | 2247,94151 | 3141,336 | 2290,719 | 2266,102836 | 4818,655 | 3877,201876 | 2756,643 | 2095,967 | 3663,371 | 2327,75827 |
| Upregulated | ENSMUSG00000022351 | 288,6907712 | 343,595751 | 616,360104 | 1129,30102 | 208,64285 | 225,5847672 | 245,352922 | 777,33575 | 554,167277 | 495,121698 | 485,809856 | 2533,833 | 1629,85777 | 2468,727 | 1528,474 | 1687,271258 | 3724,22 | 2895,26805 | 1879,27 | 1419,978 | 3224,05 | 1486,03226 |
| Upregulated | ENSMUSG00000021670 | 637,6576374 | 676,553862 | 1054,80183 | 915,389053 | 676,817049 | 554,6731334 | 689,487147 | 1164,7814 | 842,462392 | 900,711585 | 937,392199 | 1722,796 | 1425,49945 | 1668,893 | 1303,066 | 1486,731185 | 2659,566 | 2002,510859 | 1796,571 | 1708,64 | 3005,275 | 1603,58675 |
| Upregulated | ENSMUSG00000031604 | 243,219331 | 284,02497 | 467,489623 | 840,230794 | 268,436837 | 148,6205525 | 197,64541 | 733,335613 | 524,270006 | 288,011968 | 465,935816 | 1361,946 | 1423,49594 | 2394,825 | 1153,325 | 1049,189205 | 2599,178 | 2176,900721 | 1586,495 | 1007,318 | 2796,243 | 1521,92676 |
| Upregulated | ENSMUSG00000058258 | 131,1269437 | 205,306439 | 295,017723 | 522,253544 | 123,404612 | 112,7923836 | 123,812354 | 693,002154 | 421,765076 | 265,359341 | 234,072021 | 915,2629 | 1002,75822 | 1484,604 | 883,3129 | 535,9889249 | 1833,984 | 1358,457391 | 1019,958 | 723,322 | 1970,745 | 921,591264 |
| Upregulated | ENSMUSG00000083287 | 49,70134156 | 74,4634754 | 103,48314 | 218,729805 | 41,9830125 | 30,52029203 | 32,9409016 | 250,556334 | 185,790185 | 130,522278 | 70,6632518 | 341,5814 | 321,563825 | 483,6423 | 338,5104 | 186,8668871 | 626,2183 | 469,6636057 | 298,4779 | 238,6629 | 729,8399 | 355,355541 |
| Upregulated | ENSMUSG00000093930 | 920,0035565 | 1049,935 | 1794,6155 | 2363,63089 | 992,325749 | 862,5299921 | 1034,79867 | 3036,00944 | 1946,52591 | 1471,34204 | 1602,06841 | 3120,652 | 3608,32679 | 5294,808 | 2875,347 | 2593,347774 | 5634,31 | 4751,132889 | 3186,299 | 3107,951 | 6813,02 | 3519,45564 |
| Upregulated | ENSMUSG00000042719 | 664,0945213 | 672,298806 | 561,895294 | 516,472139 | 819,304849 | 594,4822099 | 714,476797 | 827,447017 | 684,433959 | 636,43094 | 938,496312 | 2129,191 | 1689,96316 | 1509,862 | 1412,186 | 1827,64931 | 1070,444 | 1121,643885 | 1031,365 | 709,3223 | 764,3833 | 1096,57695 |
| Upregulated | ENSMUSG00000011008 | 1909,800486 | 2022,21524 | 1876,31272 | 1357,6665 | 2775,96767 | 2912,697435 | 2087,77162 | 2625,34149 | 3133,87468 | 1651,48436 | 5202,58191 | 12542,17 | 11004,2948 | 10563,42 | 10204,69 | 11499,15015 | 4996,511 | 3895,037203 | 3843,14 | 1385,312 | 1478,28 | 4126,97003 |
| Upregulated | ENSMUSG00000079012 | 2051,502183 | 1988,17479 | 699,872814 | 3390,79377 | 1602,98775 | 1905,527798 | 1685,66545 | 1965,33944 | 1545,04827 | 1405,54155 | 3047,35273 | 6989,281 | 6221,9094 | 7538,085 | 3569,89 | 6372,616622 | 1977,096 | 2557,387693 | 2357,405 | 995,3178 | 2994,646 | 3376,775 |
| Upregulated | ENSMUSG00000038457 | 13,74717958 | 5,31881967 | 10,8929621 | 17,3442136 | 2,544425 | 19,90453828 | 17,0383974 | 15,8889383 | 21,3551937 | 31,2821988 | 29,8110593 | 210,2039 | 189,331972 | 335,8367 | 172,0429 | 218,7709898 | 79,41474 | 53,5059804 | 48,47889 | 2,666625 | 29,22902 | 24,2287869 |
| Upregulated | ENSMUSG00000052560 | 953,8427678 | 803,14177 | 975,827853 | 691,84141 | 681,905899 | 932,8593607 | 619,061771 | 1040,11434 | 1139,29958 | 620,250493 | 892,123554 | 2829,87 | 2986,23602 | 3162,852 | 2568,696 | 2699,998632 | 1446,01 | 1615,087927 | 1594,1 | 989,3179 | 1308,22 | 1264,38373 |
| Upregulated | ENSMUSG00000039519 | 1840,007113 | 1439,2726 | 1805,50846 | 1721,89499 | 966,881499 | 1194,272297 | 527,054425 | 799,335818 | 1407,30727 | 503,75127 | 2158,54152 | 5469,681 | 9111,9768 | 10423,1 | 6512,143 | 9620,454274 | 4475,351 | 3303,498864 | 4051,314 | 2042,635 | 1239,133 | 2039,70491 |
| Upregulated | ENSMUSG00000073530 | 143,8166479 | 521,244328 | 255,984609 | 635,9545 | 220,092762 | 61,04058406 | 134,035393 | 188,222807 | 155,892914 | 266,438038 | 205,365075 | 1258,596 | 1744,058 | 1287,218 | 576,6623 | 1343,618495 | 1058,036 | 767,9099038 | 360,2647 | 136,6645 | 434,8925 | 684,68757 |
| Upregulated | ENSMUSG00000032204 | 0 | 13,8289311 | 3,63098736 | 0,96356742 | 2,544425 | 2,653938437 | 0 | 4,88890409 | 0 | 0 | 0 | 47,29588 | 52,0913362 | 55,19322 | 19,11588 | 20,96555318 | 36,39842 | 19,81702978 | 22,8136 | 3,333281 | 0,885728 | 6,28153734 |
| Upregulated | ENSMUSG00000001056 | 541,4273804 | 648,896 | 678,086889 | 1003,07369 | 703,533512 | 684,7161168 | 631,556596 | 886,113866 | 759,177136 | 604,070045 | 694,487271 | 1748,196 | 2642,63355 | 2167,503 | 1776,184 | 2056,447304 | 1799,24 | 1623,014739 | 1515,203 | 1403,311 | 1533,195 | 1275,15208 |
| Upregulated | ENSMUSG00000001707 | 230,5296268 | 294,66261 | 204,243039 | 261,126772 | 293,881087 | 175,1599369 | 240,80935 | 360,556676 | 308,582549 | 238,391929 | 276,028327 | 749,7273 | 1028,80389 | 924,2526 | 552,7675 | 719,2096288 | 588,1654 | 585,5932299 | 393,5345 | 497,9922 | 762,6118 | 559,954186 |
| Upregulated | ENSMUSG00000027076 | 176,5983838 | 270,196039 | 189,71909 | 234,146884 | 284,9756 | 248,1432439 | 157,889149 | 300,667601 | 270,1432 | 199,558854 | 415,146604 | 641,9978 | 863,514072 | 781,1245 | 514,5357 | 560,6006612 | 519,5047 | 468,6727542 | 407,793 | 333,3281 | 632,4098 | 463,039038 |
| Upregulated | ENSMUSG00000045455 | 123,7246162 | 105,312629 | 108,021874 | 95,393175 | 124,676825 | 134,0238911 | 127,220034 | 180,889451 | 194,332263 | 117,57792 | 224,135002 | 482,5932 | 552,969568 | 512,6422 | 366,3877 | 349,1220378 | 292,0146 | 314,099922 | 254,7518 | 217,9966 | 333,0337 | 279,07973 |
| Upregulated | ENSMUSG00000048007 | 188,2306127 | 181,903633 | 186,088102 | 145,498681 | 237,903737 | 230,892644 | 157,889149 | 317,778766 | 314,989107 | 161,804476 | 323,505199 | 668,2733 | 873,531637 | 764,2858 | 523,2972 | 595,2394013 | 438,4355 | 453,8099819 | 320,3409 | 315,3284 | 404,7777 | 442,399701 |
| Upregulated | ENSMUSG00000117241 | 4,229901409 | 5,31881967 | 2,72324052 | 8,67210682 | 2,544425 | 7,961815312 | 6,81535895 | 4,88890409 | 2,13551937 | 4,31478604 | 3,31233993 | 14,0136 | 23,0403987 | 14,03218 | 14,33691 | 10,93854949 | 11,58132 | 3,963405955 | 12,35736 | 13,99978 | 5,314368 | 12,5630747 |
| Upregulated | ENSMUSG00000117720 | 0 | 0 | 1,81549368 | 7,70853939 | 2,544425 | 13,26969219 | 3,40767947 | 9,77780817 | 5,33879843 | 5,39348255 | 7,72879316 | 14,0136 | 21,0368858 | 15,90313 | 11,15093 | 12,76164107 | 9,099605 | 1,981702978 | 7,604532 | 9,999844 | 6,200096 | 8,0762623 |
| Upregulated | ENSMUSG00000024378 | 40,18406339 | 78,7185311 | 126,176811 | 252,454665 | 34,3497375 | 96,86875296 | 47,7075126 | 221,22291 | 137,740999 | 65,8004871 | 88,3290647 | 345,0848 | 455,799191 | 819,4791 | 332,1384 | 371,9106826 | 565,83 | 474,6178631 | 266,1586 | 307,3285 | 608,4951 | 344,587191 |
| Upregulated | ENSMUSG00000069125 | 33,83921127 | 78,7185311 | 72,6197472 | 68,4132871 | 33,077525 | 63,69452249 | 30,6691153 | 53,777945 | 58,7267827 | 38,8330743 | 47,4768723 | 155,9012 | 160,281034 | 378,8687 | 189,5658 | 130,3510481 | 162,9657 | 134,7558025 | 230,9877 | 211,9967 | 269,2613 | 114,862397 |
| Upregulated | ENSMUSG00000113059 | 2,114950705 | 7,44634754 | 9,98521524 | 10,5992417 | 8,90548749 | 7,961815312 | 5,67946579 | 8,55558215 | 19,2196743 | 5,39348255 | 5,52056654 | 14,88944 | 26,0456681 | 35,54818 | 18,31938 | 20,05400739 | 23,16263 | 15,85362382 | 19,9619 | 22,66631 | 34,54339 | 11,6657122 |
| Upregulated | ENSMUSG00000040026 | 2910,17217 | 2299,85763 | 4974,45268 | 4147,19419 | 346,0418 | 390,1289503 | 298,739901 | 3895,23433 | 1130,75751 | 460,603409 | 13772,7094 | 51899,35 | 58938,3416 | 227763,7 | 32428,5 | 27774,80024 | 39231,71 | 5426,893604 | 44066,36 | 49229,9 | 91979,31 | 17894,3052 |
| Upregulated | ENSMUSG00000026822 | 112,0923873 | 58,5070164 | 59,9112914 | 55,8869106 | 2,544425 | 3,980907656 | 9,08714527 | 36,6667807 | 19,2196743 | 1,07869651 | 129,181257 | 1423,256 | 4700,24133 | 7270,538 | 821,1863 | 601,6202218 | 3059,949 | 202,1337037 | 1115,965 | 785,9877 | 3374,624 | 798,652605 |
| Upregulated | ENSMUSG00000081565 | 9,517278171 | 20,2115147 | 8,16972156 | 12,5263765 | 11,4499125 | 33,17423047 | 18,1742905 | 7,33335613 | 5,33879843 | 9,70826858 | 18,7699262 | 25,39964 | 39,0685021 | 50,51583 | 41,41774 | 22,78864477 | 26,47158 | 19,81702978 | 21,86303 | 21,99966 | 34,54339 | 12,5630747 |
| Upregulated | ENSMUSG00000095499 | 3,172426057 | 4,25505574 | 6,35422788 | 2,89070227 | 3,8166375 | 14,5966614 | 2,27178632 | 4,88890409 | 6,40655811 | 1,07869651 | 7,72879316 | 10,5102 | 18,0316164 | 27,12887 | 12,74392 | 9,115457906 | 12,40855 | 4,954257444 | 9,505665 | 13,33313 | 9,743008 | 7,17889982 |
| Upregulated | ENSMUSG00000081491 | 11,63222888 | 24,4665705 | 13,6162026 | 21,1984833 | 16,5387625 | 42,463015 | 17,0383974 | 41,5556847 | 35,2360696 | 23,7313232 | 46,372759 | 30,65474 | 63,1106573 | 74,83827 | 61,33011 | 26,43482793 | 51,28868 | 22,78958424 | 29,46756 | 31,9995 | 47,82931 | 15,2551621 |
| Upregulated | ENSMUSG00000071369 | 38,06911268 | 15,956459 | 51,7415699 | 18,3077811 | 24,1720375 | 53,07876874 | 21,58197 | 46,4445888 | 37,371589 | 21,5739302 | 104,890764 | 82,32987 | 113,19848 | 152,483 | 65,31259 | 75,65830062 | 58,73382 | 90,16748548 | 69,39135 | 46,66594 | 103,6302 | 75,3784481 |
| Upregulated | ENSMUSG00000097552 | 2,114950705 | 4,25505574 | 12,7084558 | 8,67210682 | 3,8166375 | 19,90453828 | 9,08714527 | 2,44445204 | 9,60983717 | 11,8656616 | 16,5616996 | 28,90304 | 23,0403987 | 26,19339 | 18,31938 | 26,43482793 | 11,58132 | 13,87192084 | 11,4068 | 10,6665 | 25,68611 | 13,4604372 |
| Upregulated | ENSMUSG00000030109 | 34,89668663 | 26,5940983 | 13,6162026 | 18,3077811 | 8,90548749 | 11,94272297 | 15,9025042 | 30,5556505 | 16,0163953 | 11,8656616 | 65,1426852 | 275,0168 | 311,54626 | 249,7727 | 122,6602 | 148,5819639 | 316,0045 | 127,8198421 | 193,9156 | 43,99931 | 301,1475 | 148,064809 |
| Upregulated | ENSMUSG00000025591 | 144,8741233 | 102,121338 | 111,652861 | 164,77003 | 123,404612 | 70,32936859 | 87,4637732 | 143,000445 | 91,827333 | 139,15185 | 177,762243 | 488,7241 | 509,89404 | 366,7075 | 206,2922 | 346,3874004 | 273,8154 | 248,7037237 | 231,9382 | 347,3279 | 457,9214 | 287,155993 |
| Upregulated | ENSMUSG00000034203 | 168,138581 | 208,497731 | 142,516254 | 120,445928 | 189,559662 | 128,7160142 | 162,432722 | 183,333903 | 187,925705 | 121,892706 | 229,655568 | 475,5864 | 670,175075 | 554,7387 | 250,8959 | 453,9498037 | 340,8216 | 462,7276453 | 408,7436 | 389,3273 | 521,6938 | 363,431803 |
| Upregulated | ENSMUSG00000049950 | 9,517278171 | 41,4867934 | 17,24719 | 19,2713485 | 26,7164625 | 31,84726125 | 6,81535895 | 35,4445546 | 61,9300618 | 40,9904673 | 121,452464 | 122,619 | 126,221315 | 97,28975 | 36,63877 | 82,03912116 | 114,9859 | 105,0302578 | 45,62719 | 63,999 | 111,6017 | 65,5074609 |
| Upregulated | ENSMUSG00000073599 | 58,16114438 | 73,3997114 | 154,316963 | 176,332839 | 73,7883249 | 118,1002605 | 121,540568 | 64,7779792 | 166,570511 | 129,443581 | 453,79057 | 561,4197 | 588,031045 | 1115,09 | 957,3869 | 865,9685011 | 278,7788 | 302,2097041 | 324,1432 | 799,3209 | 926,4715 | 354,458179 |
| Upregulated | ENSMUSG00000116725 | 1,057475352 | 5,31881967 | 6,35422788 | 9,63567424 | 0 | 1,326969219 | 0 | 0 | 7,4743178 | 1,07869651 | 0 | 14,88944 | 21,0368858 | 65,48349 | 41,41774 | 59,25047639 | 19,02645 | 21,79873275 | 14,2585 | 27,33291 | 41,62922 | 6,28153734 |
| Upregulated | ENSMUSG00000067351 | 13,74717958 | 2,12752787 | 17,24719 | 17,3442136 | 17,810975 | 13,26969219 | 14,7666111 | 17,1111643 | 13,8808759 | 11,8656616 | 19,8740396 | 20,14454 | 36,0632327 | 66,41897 | 41,41774 | 32,81564846 | 33,08947 | 32,69809913 | 36,12153 | 34,66613 | 35,42912 | 29,6129618 |
| Upregulated | ENSMUSG00000082878 | 8,459802819 | 2,12752787 | 9,0774684 | 2,89070227 | 5,08884999 | 6,634846093 | 12,4948247 | 9,77780817 | 13,8808759 | 4,31478604 | 5,52056654 | 13,13775 | 24,0421551 | 30,87079 | 15,9299 | 33,72719425 | 20,68092 | 11,89021787 | 13,30793 | 22,66631 | 13,28592 | 8,0762623 |
| Upregulated | ENSMUSG00000046598 | 10,57475352 | 5,31881967 | 10,8929621 | 12,5263765 | 25,44425 | 6,634846093 | 12,4948247 | 1,22222602 | 14,9486356 | 6,47217906 | 30,9151726 | 59,55778 | 43,075528 | 77,64471 | 103,5443 | 144,0242349 | 50,46145 | 45,57916849 | 72,24305 | 49,99922 | 36,31485 | 34,0997741 |
| Upregulated | ENSMUSG00000027409 | 0 | 2,12752787 | 0 | 0,96356742 | 0 | 0 | 0 | 1,22222602 | 1,06775969 | 0 | 8,83290647 | 14,0136 | 23,0403987 | 40,22557 | 35,84227 | 50,13501849 | 20,68092 | 4,954257444 | 12,35736 | 12,66647 | 15,05738 | 4,48681239 |
| Upregulated | ENSMUSG00000049303 | 4,229901409 | 5,31881967 | 20,8781773 | 7,70853939 | 1,2722125 | 6,634846093 | 0 | 3,66667807 | 2,13551937 | 3,23608953 | 15,4575863 | 112,1088 | 103,180916 | 136,5798 | 156,113 | 175,0167918 | 63,69724 | 36,66150509 | 68,44079 | 42,666 | 30,11475 | 40,3813115 |
| Upregulated | ENSMUSG00000091083 | 34,89668663 | 21,2752787 | 24,5091647 | 31,797725 | 10,1777 | 10,61575375 | 11,3589316 | 17,1111643 | 35,2360696 | 15,1017511 | 38,6439658 | 44,66833 | 46,0807974 | 69,2254 | 58,94063 | 61,07356797 | 47,1525 | 38,64320806 | 30,41813 | 34,66613 | 40,74349 | 31,4076867 |
| Upregulated | ENSMUSG00000034981 | 2,114950705 | 15,956459 | 12,7084558 | 80,9396636 | 48,344075 | 218,9499211 | 14,7666111 | 70,8891093 | 79,0142167 | 155,332297 | 136,91005 | 277,6444 | 99,17389 | 433,1265 | 365,5912 | 386,4954152 | 193,5734 | 304,1914071 | 227,1854 | 549,9914 | 551,8086 | 138,193822 |

| | | | | | | | | | | | | | | | | | | | | | | | |
|---|---|---|---|---|---|---|---|---|---|---|---|---|---|---|---|---|---|---|---|---|---|---|---|
| Upregulated | ENSMUSG00000021280 | 23,26445775 | 19,1477508 | 18,1549368 | 2,89070227 | 29,2608875 | 7,961815312 | 10,2230384 | 6,11113011 | 12,8131162 | 18,3378407 | 43,060419 | 52,55098 | 34,0597198 | 53,32227 | 53,36516 | 61,07356797 | 45,49803 | 36,66150509 | 81,74872 | 82,66538 | 82,37271 | 46,6628488 |
| Upregulated | ENSMUSG00000022074 | 118,4372395 | 52,1244328 | 82,6049624 | 67,4497197 | 41,9830125 | 53,07876874 | 27,2614358 | 34,2223286 | 40,5748681 | 49,6200394 | 128,077144 | 134,8809 | 113,19848 | 152,483 | 159,299 | 174,105246 | 107,5408 | 134,7558025 | 139,7333 | 210,6634 | 148,8023 | 61,0206485 |
| Upregulated | ENSMUSG00000087070 | 60,27609508 | 57,4432524 | 30,8633926 | 17,3442136 | 54,7051374 | 41,13604578 | 28,397329 | 37,8890067 | 39,5071084 | 29,1248057 | 58,5180054 | 141,8877 | 186,326702 | 130,0315 | 125,8462 | 241,5596345 | 90,16882 | 133,764951 | 111,2163 | 121,9981 | 162,974 | 142,680634 |
| Upregulated | ENSMUSG00000002489 | 56,04619367 | 31,912918 | 61,7267851 | 42,3969667 | 39,4385875 | 47,77089187 | 31,8050084 | 84,3335955 | 74,743178 | 80,9022382 | 37,5398525 | 182,1767 | 119,209019 | 154,3539 | 116,2883 | 209,6555318 | 100,0957 | 175,3807135 | 122,6231 | 198,6636 | 170,0598 | 137,296459 |
| Upregulated | ENSMUSG00000017756 | 66,6209472 | 114,886505 | 54,4648104 | 88,648203 | 52,1607124 | 55,73270718 | 35,2126879 | 46,4445888 | 42,7103874 | 103,554865 | 50,7892122 | 124,3707 | 123,216045 | 141,2572 | 200,7167 | 249,7635466 | 120,7766 | 149,6185748 | 124,5242 | 250,6628 | 139,945 | 151,654259 |
| Upregulated | ENSMUSG00000028885 | 19,03455634 | 26,5940983 | 47,2028357 | 82,8667985 | 58,5217749 | 33,17423047 | 68,1535895 | 110,000342 | 75,8109377 | 165,040566 | 132,493597 | 119,9914 | 113,19848 | 133,7734 | 152,1305 | 176,8398834 | 120,7766 | 112,9570697 | 127,3759 | 111,3316 | 73,51543 | 104,99141 |
| Upregulated | ENSMUSG00000109489 | 9,517278171 | 8,51011147 | 9,0774684 | 10,5992417 | 25,44425 | 6,634846093 | 27,2614358 | 14,6667123 | 24,5584728 | 24,8100197 | 22,0822662 | 24,52379 | 21,0368858 | 32,74174 | 30,26681 | 52,86965586 | 52,94316 | 16,84447531 | 50,38002 | 26,66625 | 15,9431 | 21,5366995 |
| Upregulated | ENSMUSG00000082766 | 4,229901409 | 1,06376393 | 0 | 0,96356742 | 1,2722125 | 0 | 5,67946579 | 2,44445204 | 2,13551937 | 4,31478604 | 9,93701978 | 20,14454 | 8,01405172 | 7,483827 | 20,70887 | 10,93854949 | 16,54474 | 6,935960422 | 31,36869 | 7,333219 | 5,314368 | 8,0762623 |
| Upregulated | ENSMUSG00000052512 | 63,44852114 | 142,544367 | 51,7415699 | 77,0853939 | 99,2325749 | 42,463015 | 138,578965 | 19,5556163 | 22,4229534 | 47,4626464 | 68,4550251 | 340,7055 | 214,375883 | 236,676 | 421,3458 | 487,676998 | 432,6449 | 227,8958424 | 436,31 | 83,99869 | 30,11475 | 248,569406 |
| Upregulated | ENSMUSG00000030096 | 796,2789403 | 905,263108 | 818,78765 | 748,691888 | 918,537424 | 674,1003631 | 858,735228 | 705,224415 | 725,008827 | 729,19884 | 981,556732 | 3002,413 | 1041,82672 | 1728,764 | 2855,434 | 3501,247382 | 3912,83 | 2457,311692 | 3614,054 | 2598,626 | 1200,161 | 2323,27145 |
| Upregulated | ENSMUSG00000039747 | 416,6452888 | 485,076354 | 403,947344 | 372,900593 | 161,570987 | 138,0047987 | 366,89349 | 331,223252 | 178,315868 | 347,340276 | 388,647885 | 1192,031 | 398,699073 | 731,5441 | 813,2213 | 1216,002085 | 1321,097 | 732,2392502 | 1220,527 | 591,3241 | 437,5496 | 741,221406 |
| Upregulated | ENSMUSG00000029267 | 132,184419 | 197,860092 | 167,025419 | 135,863007 | 260,803562 | 216,2959826 | 197,64541 | 350,778868 | 272,27872 | 193,086675 | 348,899806 | 722,576 | 582,020506 | 590,2869 | 510,5533 | 451,2151664 | 544,3218 | 433,0021006 | 514,2565 | 366,6609 | 217,0034 | 519,572874 |
| Upregulated | ENSMUSG00000028420 | 138,5292712 | 265,940983 | 140,70076 | 170,551434 | 202,281787 | 167,1981215 | 135,171286 | 216,334006 | 356,631735 | 138,073153 | 291,485914 | 585,0676 | 579,015237 | 589,3514 | 498,6058 | 510,4656428 | 478,9701 | 465,7001997 | 523,7621 | 389,3273 | 374,663 | 398,42894 |
| Upregulated | ENSMUSG00000034394 | 518,1629226 | 499,969049 | 558,264307 | 375,791295 | 1320,55657 | 891,7233149 | 1025,71152 | 1712,33866 | 3243,85393 | 735,671019 | 2030,46437 | 5018,619 | 5013,79111 | 5014,164 | 2768,616 | 6159,314907 | 3064,085 | 4197,246907 | 3383,066 | 1811,972 | 2156,748 | 1197,08154 |
| Upregulated | ENSMUSG00000113175 | 54,98871832 | 79,782295 | 43,5718483 | 13,4899439 | 49,6162874 | 5,307876874 | 29,5332221 | 18,3333903 | 14,9486356 | 9,70826858 | 47,4768723 | 183,9284 | 162,284547 | 214,2246 | 129,0322 | 223,3287187 | 91,82329 | 168,4447531 | 95,05665 | 69,33225 | 74,40115 | 106,786135 |
| Upregulated | ENSMUSG00000028634 | 486,4386621 | 420,186754 | 524,677673 | 390,244807 | 498,707299 | 598,4631176 | 510,016028 | 551,223936 | 483,695138 | 399,117708 | 1130,61203 | 1312,899 | 1561,73833 | 1910,247 | 1402,628 | 1803,037574 | 1285,526 | 1088,945786 | 1109,311 | 774,6546 | 621,7811 | 1138,75298 |
| Upregulated | ENSMUSG00000045362 | 1020,463715 | 992,49175 | 1464,19565 | 1068,59627 | 1003,77566 | 1070,864159 | 1422,13823 | 1311,44852 | 1180,94221 | 1333,26889 | 1759,95661 | 3913,296 | 1919,36539 | 3972,041 | 3219,433 | 4240,511018 | 3582,763 | 3168,743061 | 2707,213 | 947,3185 | 797,1552 | 1744,47266 |
| Upregulated | ENSMUSG00000027596 | 35,95416198 | 46,8056131 | 42,6641015 | 58,7776129 | 25,44425 | 33,17423047 | 45,4357263 | 9,77780817 | 11,7453565 | 23,7313232 | 44,1645324 | 193,5628 | 111,194968 | 368,5785 | 265,2328 | 246,1173635 | 205,982 | 132,7740995 | 149,2389 | 72,66553 | 16,82883 | 61,9180109 |
| Upregulated | ENSMUSG00000021765 | 661,9795706 | 751,017337 | 896,853878 | 834,449389 | 910,904149 | 976,6493449 | 663,361604 | 620,890819 | 685,501718 | 621,329189 | 815,939735 | 2936,724 | 1152,01993 | 3969,235 | 3567,501 | 2564,178309 | 2631,44 | 2330,482702 | 1615,963 | 957,985 | 1170,047 | 1442,0615 |
| Upregulated | ENSMUSG00000028583 | 4835,834786 | 4909,27056 | 6080,08833 | 5877,76129 | 6008,65963 | 5000,020016 | 6249,68416 | 5876,46271 | 9919,48748 | 3778,67387 | 7768,54124 | 13357,58 | 13574,8019 | 22574,03 | 20628,42 | 22214,37092 | 16701,91 | 16197,44929 | 13392,53 | 7557,215 | 8482,617 | 10049,5624 |
| Upregulated | ENSMUSG00000117869 | 70,85084861 | 34,0404459 | 99,8521524 | 64,5590174 | 58,5217749 | 74,31027624 | 94,2791321 | 130,778184 | 70,4721393 | 119,735313 | 111,515444 | 245,2379 | 179,314407 | 276,9016 | 254,8784 | 222,4171729 | 153,8661 | 246,7220207 | 161,5963 | 181,9972 | 100,973 | 187,548758 |
| Upregulated | ENSMUSG00000030077 | 3850,267758 | 1893,4998 | 2846,69409 | 2218,13221 | 622,111912 | 578,5585793 | 503,200669 | 636,779757 | 755,973857 | 652,611388 | 1850,49391 | 11076,87 | 13944,45 | 20024,85 | 14025,48 | 13120,79011 | 14717,37 | 7164,847116 | 9790,835 | 13125,8 | 4715,616 | 3822,76415 |
| Upregulated | ENSMUSG00000022218 | 153,3339261 | 60,6345442 | 78,0662282 | 99,2474447 | 25,44425 | 35,8281689 | 85,1919869 | 44,0001368 | 58,7267827 | 38,8330743 | 94,9537446 | 702,4315 | 777,363016 | 797,9631 | 452,4091 | 678,1900682 | 618,7732 | 404,2674074 | 341,2534 | 379,9941 | 74,40115 | 126,528109 |
| Upregulated | ENSMUSG00000039037 | 709,5659614 | 344,659515 | 336,774078 | 192,713485 | 145,032225 | 277,3365667 | 174,927546 | 143,000445 | 344,886378 | 66,8791836 | 419,563057 | 1199,038 | 1057,85483 | 1710,054 | 1294,304 | 819,4796658 | 1139,932 | 747,1020226 | 1327,941 | 324,6616 | 457,0357 | 442,399701 |
| Upregulated | ENSMUSG00000021591 | 674,6692748 | 675,490098 | 1236,3512 | 626,318826 | 718,800062 | 863,8569613 | 745,145912 | 755,335681 | 928,950927 | 669,870532 | 1010,26368 | 3440,338 | 1839,22487 | 4421,071 | 2837,115 | 3093,786413 | 4216,426 | 1793,441195 | 2826,034 | 1814,638 | 1178,018 | 2054,96007 |
| Upregulated | ENSMUSG00000006818 | 5182,686702 | 6579,37993 | 5572,65785 | 4630,90504 | 6719,82642 | 5399,437751 | 5147,86779 | 6730,7987 | 8069,05994 | 5146,46105 | 8198,04132 | 18438,39 | 16945,7124 | 20392,49 | 16361,6 | 15572,84829 | 18840,32 | 10246,39525 | 15634,92 | 12414,47 | 8852,852 | 10682,2029 |
| Upregulated | ENSMUSG00000091613 | 136,4143204 | 168,074702 | 156,132456 | 132,008737 | 291,336662 | 149,9475217 | 226,042738 | 178,444999 | 213,551937 | 94,9252928 | 241,800815 | 364,3535 | 478,83959 | 417,2234 | 457,1881 | 577,0084855 | 419,4091 | 334,9078032 | 499,0474 | 207,3301 | 326,8336 | 396,634215 |
| Upregulated | ENSMUSG00000094242 | 362,7140458 | 473,374951 | 370,360711 | 380,609132 | 721,344487 | 356,9547198 | 567,946579 | 470,557018 | 493,304975 | 314,979381 | 524,453822 | 791,7681 | 1179,06736 | 1114,155 | 1311,827 | 1200,505806 | 1120,079 | 1026,522142 | 1097,904 | 534,6583 | 715,6682 | 991,585538 |
| Upregulated | ENSMUSG00000022094 | 5141,445163 | 3676,36816 | 4221,02281 | 2372,303 | 3798,82652 | 3836,268011 | 3515,58932 | 4327,90234 | 4240,07371 | 2112,08776 | 4344,68587 | 8962,57 | 8601,081 | 12718,76 | 15667,06 | 11023,32325 | 13713,93 | 6412,790836 | 9583,611 | 6665,896 | 6461,386 | 5925,28444 |
| Upregulated | ENSMUSG00000106752 | 23,26445775 | 21,2752787 | 18,1549368 | 11,5628091 | 43,255225 | 27,86635359 | 26,1255426 | 31,7778766 | 25,6262325 | 19,4165372 | 36,4357392 | 51,67513 | 53,0930926 | 65,48349 | 70,09156 | 64,71975114 | 81,06921 | 20,80788127 | 61,78682 | 32,66616 | 25,68611 | 32,3050492 |
| Upregulated | ENSMUSG00000110118 | 6,344852114 | 6,3825836 | 0 | 1,92713485 | 2,544425 | 5,307876874 | 3,40767947 | 6,11113011 | 6,40655811 | 1,07869651 | 2,20822662 | 20,14454 | 17,0298599 | 7,483827 | 9,557939 | 21,87709898 | 14,89026 | 8,917663399 | 17,1102 | 3,333281 | 7,971552 | 6,28153734 |
| Upregulated | ENSMUSG00000039929 | 338,3921127 | 401,039003 | 254,169115 | 260,163204 | 428,735612 | 301,2220126 | 403,242071 | 422,890204 | 274,414239 | 367,83551 | 404,105471 | 1263,851 | 772,354234 | 779,2535 | 704,898 | 1130,31678 | 791,6657 | 681,7058243 | 613,1154 | 356,6611 | 263,9469 | 664,945596 |
| Upregulated | ENSMUSG00000050244 | 461,0592536 | 557,412301 | 379,438179 | 402,771183 | 524,151549 | 249,4702131 | 503,200669 | 509,668251 | 471,949781 | 463,839499 | 258,362514 | 1885,704 | 1362,38879 | 761,4794 | 1041,019 | 1515,90065 | 1121,733 | 945,2723203 | 853,6087 | 700,6557 | 485,379 | 962,869938 |
| Upregulated | ENSMUSG00000032834 | 335,2196867 | 282,961206 | 183,364862 | 260,163204 | 245,537012 | 185,7756906 | 212,412021 | 237,111848 | 223,161774 | 223,290177 | 224,135002 | 1406,615 | 864,515829 | 416,2879 | 703,305 | 968,9731754 | 662,6167 | 465,7001997 | 448,6674 | 439,3265 | 415,4064 | 533,033312 |
| Upregulated | ENSMUSG00000035049 | 724,3706163 | 744,634754 | 828,772865 | 722,675568 | 606,845362 | 338,3771507 | 669,04107 | 498,668217 | 382,257967 | 739,985805 | 630,448699 | 3528,798 | 1721,01761 | 1308,734 | 1715,65 | 2524,98184 | 1772,769 | 1256,399688 | 1429,652 | 972,6515 | 764,3833 | 1422,31953 |
| Upregulated | ENSMUSG00000028331 | 41,24153874 | 30,8491541 | 34,4943799 | 26,0163204 | 53,4329249 | 55,73270718 | 45,4357263 | 51,3334929 | 11,7453565 | 57,170915 | 54,1015521 | 188,3077 | 109,191455 | 130,0315 | 134,6076 | 158,6089676 | 177,8559 | 88,1857825 | 101,7106 | 53,3325 | 83,25843 | 121,143934 |
| Upregulated | ENSMUSG00000032860 | 463,1742043 | 354,23339 | 414,840306 | 183,077811 | 321,869762 | 217,6229519 | 349,855093 | 350,778868 | 407,8842 | 431,478604 | 797,169809 | 1880,449 | 1674,93681 | 1622,12 | 1624,85 | 2101,113047 | 1631,311 | 879,8761221 | 1087,448 | 685,3226 | 1043,388 | 1165,67386 |
| Upregulated | ENSMUSG00000038279 | 676,7842255 | 645,704708 | 570,065015 | 535,743488 | 622,111912 | 622,3485635 | 507,744242 | 778,557976 | 616,097339 | 650,453995 | 801,586262 | 2698,493 | 2071,63237 | 1265,702 | 1271,206 | 1666,305705 | 1312,825 | 1275,225866 | 1074,14 | 925,9855 | 1046,045 | 1406,167 |
| Upregulated | ENSMUSG00000028540 | 109,9774366 | 55,3157246 | 60,8190383 | 80,9396636 | 110,682487 | 95,54178374 | 81,7843074 | 102,666986 | 68,3366199 | 118,656616 | 105,994878 | 525,5098 | 426,748254 | 291,8693 | 267,6223 | 407,4609684 | 280,4333 | 321,0358824 | 210,0752 | 121,3314 | 171,8312 | 260,235118 |
| Upregulated | ENSMUSG00000029246 | 248,5067078 | 186,158688 | 145,239494 | 187,895648 | 242,992587 | 199,0453828 | 189,694157 | 339,778834 | 249,855766 | 236,234536 | 204,260962 | 738,3413 | 596,045096 | 537,9001 | 432,4968 | 514,1118259 | 424,3725 | 513,2610712 | 358,3636 | 424,66 | 303,8047 | 352,663454 |
| Upregulated | ENSMUSG00000021694 | 53,93124297 | 44,6780852 | 43,5718483 | 53,9597757 | 83,9660249 | 57,0596764 | 69,2894827 | 85,5558215 | 73,6754183 | 66,8791836 | 66,2467985 | 205,8247 | 174,305625 | 113,1929 | 168,0604 | 166,8128797 | 114,1587 | 142,6826144 | 70,34192 | 81,33206 | 84,14416 | 96,9151476 |
| Upregulated | ENSMUSG00000022704 | 132,184419 | 147,863187 | 124,361317 | 117,555226 | 146,304437 | 98,19572218 | 212,412021 | 182,111677 | 117,453565 | 141,309243 | 145,742957 | 474,7105 | 485,851885 | 289,9983 | 274,7908 | 409,28406 | 230,7991 | 200,1520007 | 209,1246 | 265,3292 | 197,5173 | 297,02698 |
| Upregulated | ENSMUSG00000053801 | 326,7598839 | 344,659515 | 318,619141 | 365,192054 | 457,9965 | 493,6325493 | 358,942238 | 477,890375 | 479,424099 | 266,438038 | 531,078502 | 2090,653 | 1875,2881 | 1207,703 | 959,7764 | 1575,151126 | 962,0764 | 923,4735876 | 688,2101 | 585,3242 | 844,0988 | 845,315454 |
| Upregulated | ENSMUSG00000041506 | 348,9668663 | 359,55221 | 295,92547 | 444,204582 | 337,136312 | 429,9380268 | 318,050084 | 386,223423 | 327,802223 | 264,280645 | 432,812417 | 1481,062 | 1511,65051 | 1116,026 | 865,79 | 1288,925748 | 819,7917 | 709,449666 | 673,0011 | 674,6561 | 722,7541 | 714,300532 |
| Upregulated | ENSMUSG00000001305 | 234,7595282 | 247,856997 | 228,752204 | 285,215958 | 278,614537 | 287,9523204 | 235,129884 | 240,778526 | 342,750859 | 181,221014 | 267,195421 | 812,7885 | 915,605409 | 759,6085 | 591,7957 | 802,1602958 | 434,2993 | 477,5904176 | 454,3708 | 315,3284 | 549,1514 | 447,783876 |
| Upregulated | ENSMUSG00000028741 | 217,8399226 | 267,004747 | 275,955039 | 301,596604 | 388,024812 | 371,5513812 | 373,708849 | 341,00106 | 290,430634 | 298,798933 | 428,395964 | 886,3599 | 803,408685 | 763,3504 | 589,4063 | 771,1677389 | 616,2914 | 462,7276453 | 465,7776 | 392,6605 | 473,8645 | 402,915752 |
| Upregulated | ENSMUSG00000091105 | 26,43688381 | 31,912918 | 39,0331141 | 40,4698318 | 38,166375 | 37,15513812 | 42,0280469 | 64,7779792 | 69,4043796 | 31,2821988 | 38,6439658 | 147,1427 | 132,231853 | 103,8381 | 100,3584 | 143,1126891 | 87,68711 | 73,32301017 | 64,63852 | 98,66513 | 77,05834 | 55,6364736 |
| Upregulated | ENSMUSG00000032841 | 71,90832396 | 305,300249 | 137,97752 | 157,06149 | 408,380212 | 221,6038595 | 303,283473 | 37,8890067 | 527,473285 | 213,581909 | 373,190298 | 565,7989 | 461,80973 | 561,287 | 454,7986 | 563,3352986 | 503,7872 | 411,2033679 | 565,5871 | 237,9963 | 366,6914 | 442,399701 |
| Upregulated | ENSMUSG00000028599 | 1576,69575 | 1693,51218 | 1022,12294 | 647,517309 | 1982,10707 | 1296,448927 | 2121,84842 | 1791,78335 | 1909,15432 | 2340,77142 | 2666,43364 | 5566,901 | 2465,32266 | 3338,722 | 2285,144 | 5063,636867 | 3896,286 | 4187,338392 | 4460,058 | 1631,308 | 3128,391 | 4735,38179 |
| Upregulated | ENSMUSG00000027800 | 523,4502994 | 1083,97545 | 1048,4476 | 1003,07369 | 1236,59055 | 688,6970245 | 1191,55192 | 867,780475 | 928,950927 | 787,448452 | 1132,82025 | 5206,051 | 2431,26294 | 5444,484 | 3037,035 | 4101,956058 | 3374,299 | 3790,006945 | 4156,827 | 2788,623 | 4769,645 | 4950,74879 |
| Upregulated | ENSMUSG00000006724 | 2,114950705 | 8,51011147 | 3,63098736 | 2,89070227 | 5,08884999 | 1,326969219 | 7,95125211 | 14,6667123 | 3,20327906 | 6,47217906 | 12,1452464 | 58,68193 | 14,0245905 | 57,06418 | 38,23176 | 43,75419795 | 42,18908 | 25,76213871 | 48,47889 | 13,99978 | 18,60029 | 18,844612 |
| Upregulated | ENSMUSG00000118747 | 0 | 0 | 0 | 0 | 0 | 0 | 0 | 0 | 0 | 2,15739302 | 0 | 42,04079 | 8,01405172 | 31,80627 | 20,70887 | 12,76164107 | 24,81711 | 18,82617829 | 23,76416 | 1,333313 | 11,51446 | 24,2287869 |
| Upregulated | ENSMUSG00000061175 | 456,8293522 | 521,244328 | 404,855091 | 206,203429 | 914,720787 | 634,2912865 | 924,617031 | 721,113353 | 846,733431 | 410,98337 | 915,309933 | 2988,399 | 1466,57146 | 2187,148 | 1678,215 | 2267,014381 | 1494,817 | 1765,697353 | 1880,221 | 882,6529 | 1283,42 | 1740,88321 |
| Upregulated | ENSMUSG00000043613 | 29044,61803 | 66506,5211 | 42245,6302 | 52980,7912 | 101336,814 | 60135,59105 | 52672,5016 | 43315,6902 | 52236,9393 | 46144,4793 | 82287,3567 | 207098,2 | 131832,152 | 231725,5 | 128760,6 | 187110,2698 | 220319,6 | 188106,2192 | 219266,2 | 85735,33 | 113323,6 | 182744,279 |
| Upregulated | ENSMUSG00000027200 | 890,3942467 | 741,443462 | 831,496105 | 510,690735 | 564,862349 | 505,5752723 | 707,661438 | 613,557463 | 742,092982 | 782,054969 | 808,210942 | 2142,328 | 1727,02814 | 1860,667 | 1585,025 | 1939,769442 | 2042,448 | 2222,479889 | 2277,557 | 1317,979 | 1077,931 | 1620,63663 |
| Upregulated | ENSMUSG00000039629 | 10,57475352 | 8,51011147 | 6,35422788 | 11,5628091 | 6,36106249 | 5,307876874 | 29,5332221 | 3,66667807 | 18,1519147 | 33,4395918 | 15,4575863 | 70,06798 | 80,1405172 | 15,90313 | 23,89485 | 15,49627844 | 9,926842 | 23,78043573 | 72,24305 | 24,66628 | 26,57184 | 55,6364736 |
| Upregulated | ENSMUSG00000111341 | 14,80465493 | 11,7014033 | 11,8007089 | 8,67210682 | 10,1777 | 23,88544594 | 13,6307179 | 12,2222602 | 18,1519147 | 8,62957207 | 25,3946061 | 55,17853 | 43,075528 | 18,70957 | 30,26681 | 23,70019056 | 9,926842 | 14,86277233 | 40,87436 | 1,333313 | 5,314368 | 24,2287869 |
| Upregulated | ENSMUSG00000107971 | 5,287376762 | 8,51011147 | 0 | 0,96356742 | 3,8166375 | 7,961815312 | 20,4460768 | 7,33335613 | 11,7453565 | 1,07869651 | 24,2904928 | 21,89624 | 44,0772844 | 5,61287 | 12,74392 | 19,1424616 | 7,445132 | 8,917663399 | 13,30793 | 5,999906 | 5,314368 | 31,4076867 |
| Upregulated | ENSMUSG00000005413 | 2414,216229 | 4176,3372 | 2964,70118 | 2275,94626 | 4913,28467 | 4067,160655 | 4169,86378 | 3275,56574 | 4479,25188 | 2894,14273 | 5237,91354 | 24560,58 | 37631,9833 | 9233,172 | 13948,22 | 11939,42677 | 10222,99 | 7377,880186 | 6180,583 | 3455,279 | 5932,606 | 8735,82372 |
| Upregulated | ENSMUSG00000028766 | 127,9545176 | 147,863187 | 195,165571 | 73,2311242 | 97,9603624 | 39,80907656 | 73,8330553 | 226,111814 | 306,44703 | 51,7774324 | 147,951183 | 948,5452 | 2826,95674 | 828,8339 | 1511,747 | 1662,659522 | 1300,416 | 530,1055465 | 1011,403 | 808,654 | 511,9508 | 572,517261 |
| Upregulated | ENSMUSG00000028977 | 0 | 8,51011147 | 1,81549368 | 2,89070227 | 2,544425 | 1,326969219 | 3,40767947 | 4,88890409 | 18,1519147 | 2,15739302 | 12,1452464 | 16,64114 | 27,0474245 | 9,354784 | 10,35443 | 20,96555318 | 9,099605 | 11,89021787 | 10,45623 | 8,666531 | 13,28592 | 12,5630747 |
| Upregulated | ENSMUSG00000000182 | 286,5758205 | 35,1042098 | 54,4648104 | 123,33663 | 3,8166375 | 0 | 1,13589316 | 0 | 0 | 1,07869651 | 16,5616996 | 1452,159 | 4187,34202 | 644,5446 | 350,4578 | 352,768221 | 1789,313 | 868,9767557 | 1933,452 | 1879,971 | 1309,992 | 229,724794 |
| Upregulated | ENSMUSG00000021696 | 26,43688381 | 35,1042098 | 16,3394431 | 6,74497197 | 11,4499125 | 45,11695343 | 7,95125211 | 18,3333903 | 17,084155 | 19,4165372 | 5,52056654 | 84,08157 | 203,356562 | 72,96731 | 69,29506 | 74,74675483 | 73,62408 | 39,63405955 | 93,15552 | 95,33185 | 46,05786 | 36,7918616 |
| Upregulated | ENSMUSG00000039058 | 54,98871832 | 32,976682 | 9,0774684 | 45,2876689 | 0 | 2,653938437 | 12,4948247 | 13,4444862 | 14,9486356 | 0 | 7,72879316 | 255,7481 | 359,630571 | 117,8703 | 19,91237 | 49,22347269 | 86,85987 | 33,68895062 | 212,9269 | 120,6648 | 117,8018 | 75,3784481 |
| Upregulated | ENSMUSG00000033107 | 584,7838698 | 246,793233 | 400,316356 | 181,150676 | 75,0605374 | 118,1002605 | 46,5716195 | 86,7780475 | 136,67324 | 125,128795 | 269,403647 | 637,6186 | 824,44557 | 485,5133 | 488,2514 | 534,1658333 | 453,3258 | 412,1942193 | 381,1772 | 595,9907 | 573,066 | 575,209348 |
| Upregulated | ENSMUSG00000110251 | 90,9428803 | 41,4867934 | 72,6197472 | 22,1620508 | 12,722125 | 11,94272297 | 5,67946579 | 23,2222944 | 21,3551937 | 25,8887162 | 35,3316259 | 81,45402 | 112,196724 | 59,87062 | 55,75465 | 66,54284272 | 63,69724 | 54,49683188 | 69,39135 | 83,33203 | 65,54387 | 68,1995483 |
| Upregulated | ENSMUSG00000038296 | 161,7937289 | 32,976682 | 181,549368 | 36,6155621 | 16,5387625 | 19,90453828 | 12,4948247 | 114,889246 | 76,8786974 | 49,6200394 | 64,0385719 | 557,0404 | 609,06793 | 711,8991 | 238,9485 | 432,9842506 | 76,93303 | 106,0211093 | 235,7405 | 228,6631 | 461,4643 | 271,003468 |
| Upregulated | ENSMUSG00000021062 | 32,78173592 | 17,0202229 | 40,8486078 | 9,63567424 | 16,5387625 | 0 | 6,81535895 | 4,88890409 | 9,60983717 | 33,4395918 | 37,5398525 | 78,82647 | 169,296843 | 103,8381 | 12,74392 | 26,43482793 | 14,89026 | 22,78958424 | 26,61586 | 35,99944 | 68,20106 | 38,5865865 |
| Upregulated | ENSMUSG00000025314 | 26,43688381 | 19,1477508 | 35,4021268 | 11,5628091 | 7,63327499 | 6,634846093 | 15,9025042 | 13,4444862 | 14,9486356 | 14,0230546 | 75,079705 | 108,6054 | 121,212532 | 75,77375 | 38,23176 | 32,81564846 | 62,87 | 19,81702978 | 43,72606 | 79,3321 | 139,0593 | 50,2522987 |
| Upregulated | ENSMUSG00000027832 | 2791,73493 | 943,558609 | 3420,39009 | 653,298713 | 265,892412 | 554,6731334 | 212,412021 | 306,778731 | 375,851409 | 804,707596 | 4727,81319 | 10793,97 | 6951,18811 | 2668,92 | 766,2281 | 741,0867278 | 991,857 | 1536,810659 | 1232,885 | 5821,242 | 6162,01 | 1218,61824 |
| Upregulated | ENSMUSG00000030098 | 50,75881691 | 12,7651672 | 19,9704305 | 16,3806462 | 13,9943375 | 21,2315075 | 5,67946579 | 24,4445204 | 22,4229534 | 33,4395918 | 3,31233993 | 75,32307 | 51,0895797 | 8,419306 | 46,19671 | 86,59685011 | 36,39842 | 26,7529902 | 115,0185 | 22,66631 | 52,25795 | 54,7391111 |
| Upregulated | ENSMUSG00000072572 | 8,459802819 | 0 | 6,35422788 | 0,96356742 | 0 | 14,5966614 | 0 | 0 | 0 | 0 | 0 | 14,88944 | 2,00351293 | 0 | 0,796495 | 17,31937002 | 0 | 5,945108933 | 10,45623 | 10,6665 | 7,971552 | 17,0498871 |
| Upregulated | ENSMUSG00000050022 | 2,114950705 | 3,1912918 | 4,5387342 | 0,96356742 | 2,544425 | 6,634846093 | 9,08714527 | 14,6667123 | 10,6775969 | 0 | 9,93701978 | 37,66154 | 21,0368858 | 0 | 16,72639 | 49,22347269 | 7,445132 | 25,76213871 | 5,703399 | 8,666531 | 21,25747 | 19,7419745 |
| Upregulated | ENSMUSG00000043089 | 2,114950705 | 1,06376393 | 0,90774684 | 0 | 2,544425 | 3,980907656 | 14,7666111 | 1,22222602 | 2,13551937 | 0 | 5,52056654 | 23,64794 | 2,00351293 | 0 | 6,37196 | 17,31937002 | 13,23579 | 21,79873275 | 1,901133 | 8,666531 | 5,314368 | 17,0498871 |
| Upregulated | ENSMUSG00000031847 | 10,57475352 | 11,7014033 | 3,63098736 | 1,92713485 | 3,8166375 | 11,94272297 | 27,2614358 | 4,88890409 | 0 | 5,39348255 | 1,10411331 | 37,66154 | 16,0281034 | 14,03218 | 17,52289 | 17,31937002 | 31,435 | 10,89936638 | 18,06076 | 21,333 | 10,62874 | 17,9472495 |
| Upregulated | ENSMUSG00000066042 | 12,68970423 | 15,956459 | 21,7859242 | 6,74497197 | 15,26655 | 21,2315075 | 38,6203674 | 8,55558215 | 12,8131162 | 7,55087556 | 11,0411331 | 69,19213 | 40,0702586 | 39,29009 | 43,01073 | 41,01956058 | 39,70737 | 32,69809913 | 50,38002 | 11,33316 | 37,20058 | 34,9971366 |
| Upregulated | ENSMUSG00000024589 | 274,9435916 | 273,387331 | 266,877571 | 111,773821 | 435,096675 | 222,9308287 | 311,234725 | 243,222978 | 234,907131 | 273,988913 | 342,275126 | 1320,781 | 783,373555 | 542,5775 | 631,6205 | 604,3548592 | 502,1328 | 474,6178631 | 630,2256 | 583,3242 | 445,5212 | 595,848685 |
| Upregulated | ENSMUSG00000106188 | 5,287376762 | 4,25505574 | 8,16972156 | 4,81783712 | 10,1777 | 13,26969219 | 10,2230384 | 9,77780817 | 3,20327906 | 8,62957207 | 3,31233993 | 31,53059 | 22,0386422 | 9,354784 | 15,1334 | 16,40782423 | 19,02645 | 10,89936638 | 14,2585 | 14,66644 | 9,743008 | 17,9472495 |
| Upregulated | ENSMUSG00000020692 | 86,71297889 | 30,8491541 | 66,2655193 | 77,0853939 | 111,9547 | 161,8902447 | 107,90985 | 79,4446914 | 59,7945424 | 60,4070045 | 97,1619712 | 217,2107 | 191,335485 | 169,3216 | 166,4674 | 255,2328214 | 169,5836 | 196,1885948 | 125,4748 | 175,3306 | 89,45853 | 91,5309727 |
| Upregulated | ENSMUSG00000103367 | 26,43688381 | 7,44634754 | 27,2324052 | 21,1984833 | 19,0831875 | 9,28878453 | 9,08714527 | 9,77780817 | 19,2196743 | 2,15739302 | 9,93701978 | 84,08157 | 32,0562069 | 89,80593 | 47,7897 | 91,15457906 | 52,11592 | 43,59746551 | 39,92379 | 61,99903 | 54,91514 | 20,639337 |
| Upregulated | ENSMUSG00000115368 | 35,95416198 | 28,7216262 | 11,8007089 | 7,70853939 | 34,3497375 | 9,28878453 | 11,3589316 | 8,55558215 | 9,60983717 | 7,55087556 | 17,6658129 | 98,97102 | 33,0579633 | 81,38662 | 86,81795 | 91,15457906 | 38,05289 | 74,31386166 | 28,51699 | 87,33197 | 68,20106 | 32,3050492 |
| Upregulated | ENSMUSG00000096157 | 10,57475352 | 25,5303344 | 19,9704305 | 16,3806462 | 17,810975 | 19,90453828 | 20,4460768 | 15,8889383 | 12,8131162 | 25,8887162 | 22,0822662 | 60,43363 | 57,1001185 | 45,83844 | 35,84227 | 56,51583902 | 67,00618 | 38,64320806 | 38,02266 | 44,66597 | 21,25747 | 28,7155993 |
| Upregulated | ENSMUSG00000049553 | 293,9781479 | 211,689023 | 209,68952 | 268,835311 | 281,158962 | 123,4081373 | 201,053089 | 294,556471 | 176,180348 | 247,021501 | 269,403647 | 997,5928 | 637,117111 | 333,0303 | 665,8698 | 688,2170719 | 772,6392 | 467,6819027 | 534,2184 | 720,6554 | 294,0617 | 477,396838 |
| Upregulated | ENSMUSG00000021868 | 97,28773241 | 46,8056131 | 108,929621 | 108,883119 | 97,9603624 | 98,19572218 | 112,453423 | 122,222602 | 54,455744 | 45,3052534 | 132,493597 | 383,6222 | 254,446142 | 221,7084 | 245,3204 | 266,1713709 | 310,2138 | 174,389862 | 245,2462 | 277,329 | 199,2888 | 178,575133 |
| Upregulated | ENSMUSG00000026558 | 567,8642642 | 572,304996 | 670,824915 | 723,639135 | 470,718625 | 334,3962431 | 517,96728 | 345,889964 | 407,8842 | 680,657497 | 1022,40892 | 2193,128 | 1495,6224 | 1360,186 | 1752,289 | 1923,361618 | 1944,834 | 1077,055568 | 1844,099 | 2218,632 | 1103,617 | 1146,82925 |
| Upregulated | ENSMUSG00000022797 | 158,6213028 | 241,474413 | 234,198685 | 227,401912 | 290,06445 | 305,2029203 | 343,039734 | 464,445888 | 463,407704 | 269,674127 | 196,532169 | 981,8275 | 535,939709 | 483,6423 | 567,9009 | 699,1556214 | 760,2307 | 539,0232099 | 683,4573 | 805,3208 | 459,6928 | 719,684707 |
| Upregulated | ENSMUSG00000061436 | 62,39104579 | 80,846059 | 63,5422788 | 47,2148038 | 72,5161124 | 132,6969219 | 112,453423 | 56,222397 | 80,0819764 | 78,7448452 | 145,742957 | 344,2089 | 127,223071 | 242,2889 | 133,8112 | 202,3631655 | 251,48 | 176,371565 | 218,6303 | 227,9964 | 196,6316 | 185,754033 |

| | | | | | | | | | | | | | | | | | | | | | | | |
|---|---|---|---|---|---|---|---|---|---|---|---|---|---|---|---|---|---|---|---|---|---|---|---|
| Upregulated | ENSMUSG00000024754 | 873,474641 | 1027,59596 | 1110,17439 | 706,294922 | 1197,15196 | 882,4345304 | 1055,24474 | 1074,33667 | 1074,16624 | 1147,73309 | 1624,15068 | 3724,989 | 2503,3894 | 2644,597 | 2242,93 | 3259,687747 | 3549,673 | 2829,871852 | 3325,082 | 3567,944 | 2685,527 | 2847,33114 |
| Upregulated | ENSMUSG00000092500 | 30,66678522 | 24,4665705 | 9,98521524 | 2,89070227 | 30,5331 | 34,50119968 | 28,397329 | 18,3333903 | 27,7617518 | 7,55087556 | 56,3097787 | 135,7567 | 86,151056 | 108,5155 | 74,07403 | 100,270037 | 149,7299 | 77,28641613 | 107,414 | 111,3316 | 83,25843 | 76,2758106 |
| Upregulated | ENSMUSG00000037443 | 245,3342817 | 211,689023 | 187,903596 | 192,713485 | 179,381962 | 134,0238911 | 216,955593 | 300,667601 | 232,771611 | 140,230546 | 384,231431 | 689,2937 | 427,75001 | 384,4816 | 336,9174 | 454,8613495 | 712,2509 | 489,4806355 | 538,9712 | 655,3231 | 387,9489 | 395,736853 |
| Upregulated | ENSMUSG00000095134 | 241,1043803 | 102,121338 | 131,623292 | 83,8303659 | 329,503037 | 356,9547198 | 326,001336 | 367,890033 | 179,383627 | 153,174904 | 306,9435 | 1391,725 | 621,089008 | 304,0305 | 152,927 | 562,4237528 | 921,5418 | 699,5411511 | 403,9908 | 971,3182 | 246,2324 | 734,939869 |
| Upregulated | ENSMUSG00000035299 | 251,6791339 | 113,822741 | 200,612052 | 211,984833 | 296,425512 | 391,4559195 | 347,583306 | 519,446059 | 239,17817 | 145,624029 | 405,209584 | 1442,524 | 755,324374 | 304,966 | 191,9553 | 547,8390202 | 605,5374 | 939,3272114 | 417,2987 | 963,3183 | 214,3462 | 840,828641 |
| Upregulated | ENSMUSG00000086695 | 42,29901409 | 27,6578623 | 25,4169115 | 15,4170788 | 49,6162874 | 55,73270718 | 53,3869784 | 83,1113695 | 34,1683099 | 31,2821988 | 77,2879316 | 282,0236 | 132,231853 | 84,19306 | 43,01073 | 75,65830062 | 137,3213 | 169,4356046 | 76,99589 | 206,6634 | 24,80038 | 159,730521 |
| Upregulated | ENSMUSG00000097180 | 24,3219331 | 26,5940983 | 48,1105825 | 49,1419386 | 57,2495624 | 75,63724546 | 128,355927 | 86,7780475 | 45,9136665 | 31,2821988 | 102,682538 | 169,9148 | 111,194968 | 85,12853 | 84,42846 | 106,6508575 | 96,78671 | 137,7283569 | 93,15552 | 131,3313 | 57,57232 | 91,5309727 |
| Upregulated | ENSMUSG00000104801 | 23,26445775 | 12,7651672 | 2,72324052 | 8,67210682 | 12,722125 | 11,94272297 | 5,67946579 | 46,4445888 | 18,1519147 | 22,6526267 | 35,3316259 | 85,83327 | 62,1089008 | 38,35461 | 11,94742 | 22,78864477 | 34,74395 | 86,20407953 | 63,68796 | 59,99906 | 39,85776 | 84,3520729 |
| Upregulated | ENSMUSG00000016181 | 46,5289155 | 34,0404459 | 72,6197472 | 79,9760962 | 129,765675 | 163,2172139 | 84,0560937 | 53,777945 | 66,2011005 | 159,647083 | 135,805937 | 292,5338 | 271,476002 | 178,6764 | 132,2182 | 175,0167918 | 43,84355 | 235,8226543 | 155,8929 | 111,9983 | 108,0588 | 237,801057 |
| Upregulated | ENSMUSG00000105801 | 1,057475352 | 0 | 0,90774684 | 0,96356742 | 6,36106249 | 3,980907656 | 14,7666111 | 6,11113011 | 8,54207749 | 31,2821988 | 11,0411331 | 35,03399 | 15,026347 | 10,29026 | 8,761444 | 2,734637372 | 5,790658 | 22,78958424 | 12,35736 | 16,66641 | 1,771456 | 29,6129618 |
| Upregulated | ENSMUSG00000097804 | 17,97708099 | 8,51011147 | 19,9704305 | 15,4170788 | 81,4215999 | 35,8281689 | 36,3485811 | 67,2224312 | 82,2174958 | 23,7313232 | 78,3920449 | 69,19213 | 32,0562069 | 41,16105 | 82,03898 | 45,57728953 | 55,42487 | 182,3166739 | 39,92379 | 75,33216 | 25,68611 | 100,504597 |
| Upregulated | ENSMUSG00000106121 | 3,172426057 | 8,51011147 | 4,5387342 | 6,74497197 | 10,1777 | 6,634846093 | 13,6307179 | 2,44445204 | 2,13551937 | 1,07869651 | 35,3316259 | 13,13775 | 34,0597198 | 20,58052 | 6,37196 | 25,52328214 | 10,75408 | 40,62491104 | 31,36869 | 35,33278 | 20,37174 | 25,1261494 |
| Upregulated | ENSMUSG00000044037 | 17,97708099 | 8,51011147 | 18,1549368 | 21,1984833 | 52,1607124 | 19,90453828 | 19,3101837 | 4,88890409 | 25,6262325 | 21,5739302 | 29,8110593 | 46,42003 | 15,026347 | 29,93531 | 39,82475 | 69,27748009 | 46,32526 | 30,71639615 | 42,77549 | 80,66541 | 29,22902 | 119,349209 |
| Upregulated | ENSMUSG00000032327 | 0 | 0 | 0 | 1,92713485 | 1,2722125 | 0 | 0 | 2,44445204 | 2,13551937 | 5,39348255 | 0 | 21,89624 | 0 | 0 | 0 | 0 | 0 | 9,908514888 | 2,851699 | 26,66625 | 14,17165 | 17,9472495 |
| Upregulated | ENSMUSG00000029368 | 0 | 0 | 9,98521524 | 8,67210682 | 25,44425 | 11,94272297 | 3,40767947 | 8,55558215 | 8,54207749 | 24,8100197 | 2,20822662 | 24,52379 | 2,00351293 | 7,483827 | 0 | 5,469274744 | 17,37197 | 45,57916849 | 9,505665 | 87,33197 | 15,9431 | 48,4575738 |
| Upregulated | ENSMUSG00000041235 | 15,86213028 | 0 | 9,0774684 | 11,5628091 | 21,6276125 | 13,26969219 | 6,81535895 | 0 | 3,20327906 | 3,23608953 | 8,83290647 | 41,16494 | 22,0386422 | 16,83861 | 7,964949 | 4,557728953 | 26,47158 | 36,66150509 | 45,62719 | 51,99919 | 29,22902 | 44,8681239 |
| Upregulated | ENSMUSG00000111690 | 21,14950705 | 3,1912918 | 4,5387342 | 0,96356742 | 7,63327499 | 9,28878453 | 10,2230384 | 12,2222602 | 5,33879843 | 1,07869651 | 7,72879316 | 41,16494 | 37,0649892 | 2,806435 | 19,11588 | 11,85009528 | 22,33539 | 23,78043573 | 22,8136 | 73,33219 | 14,17165 | 50,2522987 |
| Upregulated | ENSMUSG00000021803 | 0 | 4,25505574 | 5,44648104 | 2,89070227 | 0 | 0 | 0 | 1,22222602 | 0 | 0 | 0 | 0 | 17,0298599 | 22,45148 | 0,796495 | 0 | 6,617895 | 7,92681191 | 19,9619 | 15,33309 | 2,657184 | 5,38417486 |
| Upregulated | ENSMUSG00000114432 | 20,09203169 | 11,7014033 | 4,5387342 | 24,0891856 | 13,9943375 | 2,653938437 | 15,9025042 | 7,33335613 | 2,13551937 | 6,47217906 | 7,72879316 | 20,14454 | 57,1001185 | 92,61236 | 2,389485 | 3,646183163 | 28,12605 | 10,89936638 | 124,5242 | 0,666656 | 13,28592 | 10,7683497 |
| Upregulated | ENSMUSG00000028076 | 320,4150318 | 269,132275 | 319,526888 | 229,329047 | 134,854525 | 213,6420442 | 164,704508 | 129,555958 | 122,792364 | 128,364885 | 194,323942 | 387,1256 | 449,788653 | 455,578 | 306,6506 | 318,1294809 | 532,7405 | 219,9690305 | 742,3924 | 351,3279 | 198,4031 | 314,076867 |
| Upregulated | ENSMUSG00000041750 | 31,72426057 | 15,956459 | 50,833823 | 34,6884273 | 15,26655 | 25,21241515 | 20,4460768 | 8,55558215 | 4,27103874 | 5,39348255 | 32,019286 | 41,16494 | 72,1264654 | 43,96748 | 26,28433 | 30,08101109 | 51,28868 | 31,70724764 | 107,414 | 43,99931 | 8,85728 | 37,6892241 |
| Upregulated | ENSMUSG00000075270 | 168,138581 | 141,480603 | 129,807798 | 107,919551 | 19,0831875 | 22,55847672 | 24,9896495 | 55,000171 | 24,5584728 | 42,0691639 | 47,4768723 | 141,8877 | 479,841347 | 132,8379 | 50,17918 | 125,7933191 | 196,0551 | 87,19493102 | 592,2029 | 421,9934 | 285,2044 | 172,293596 |
| Upregulated | ENSMUSG00000097534 | 12,68970423 | 10,6376393 | 3,63098736 | 4,81783712 | 6,36106249 | 9,28878453 | 7,95125211 | 12,2222602 | 4,27103874 | 5,39348255 | 5,52056654 | 13,13775 | 29,0509375 | 20,58052 | 17,52289 | 11,85009528 | 20,68092 | 18,82617829 | 49,42946 | 21,99966 | 25,68611 | 16,1525246 |
| Upregulated | ENSMUSG00000009654 | 4,229901409 | 5,31881967 | 7,26197472 | 15,4170788 | 0 | 0 | 0 | 8,55558215 | 0 | 8,62957207 | 0 | 3,503399 | 2,00351293 | 8,419306 | 8,761444 | 1,823091581 | 19,02645 | 9,908514888 | 27,56643 | 19,99969 | 11,51446 | 12,5630747 |
| Upregulated | ENSMUSG00000051279 | 6,344852114 | 4,25505574 | 8,16972156 | 14,4535114 | 0 | 7,961815312 | 0 | 0 | 1,06775969 | 6,47217906 | 0 | 3,503399 | 0 | 8,419306 | 0,796495 | 0 | 16,54474 | 1,981702978 | 48,47889 | 23,99963 | 23,91466 | 15,2551621 |
| Upregulated | ENSMUSG00000099470 | 35,95416198 | 4,25505574 | 17,24719 | 19,2713485 | 22,899825 | 14,5966614 | 28,397329 | 11,0000342 | 13,8808759 | 22,6526267 | 17,6658129 | 18,39284 | 6,01053879 | 16,83861 | 30,26681 | 32,81564846 | 62,87 | 32,69809913 | 94,10608 | 69,99891 | 23,02893 | 24,2287869 |
| Upregulated | ENSMUSG00000018849 | 41,24153874 | 18,0839869 | 65,3577725 | 14,4535114 | 21,6276125 | 7,961815312 | 6,81535895 | 11,0000342 | 6,40655811 | 25,8887162 | 8,83290647 | 43,79248 | 26,0456681 | 27,12887 | 57,34764 | 30,08101109 | 97,61395 | 68,36875273 | 177,7559 | 111,3316 | 31,88621 | 82,5573479 |
| Upregulated | ENSMUSG00000056054 | 0 | 0 | 0 | 0 | 0 | 1,326969219 | 0 | 0 | 0 | 1,07869651 | 0 | 6,130948 | 4,00702586 | 17,77409 | 11,94742 | 3,646183163 | 15,7175 | 6,935960422 | 118,8208 | 57,33244 | 0 | 28,7155993 |
| Upregulated | ENSMUSG00000079597 | 0 | 2,12752787 | 0 | 0 | 0 | 0 | 0 | 0 | 0 | 0 | 0 | 2,627549 | 0 | 11,22574 | 14,33691 | 1,823091581 | 99,26842 | 9,908514888 | 226,2348 | 125,998 | 2,657184 | 111,272947 |
| Upregulated | ENSMUSG00000023232 | 6,344852114 | 0 | 24,5091647 | 21,1984833 | 0 | 0 | 2,27178632 | 17,1111643 | 0 | 17,2591441 | 4,41645324 | 6,130948 | 5,00878232 | 0 | 3,982475 | 0 | 54,59763 | 1,981702978 | 123,5736 | 43,99931 | 4,42864 | 49,3549363 |
| Upregulated | ENSMUSG00000005983 | 14,80465493 | 11,7014033 | 9,0774684 | 15,4170788 | 6,36106249 | 13,26969219 | 2,27178632 | 6,11113011 | 9,60983717 | 14,0230546 | 18,7699262 | 5,255098 | 18,0316164 | 30,87079 | 12,74392 | 5,469274744 | 35,57118 | 25,76213871 | 52,28116 | 50,66588 | 15,9431 | 14,3577996 |
| Upregulated | ENSMUSG00000082163 | 25,37940846 | 15,956459 | 9,0774684 | 9,63567424 | 5,08884999 | 3,980907656 | 11,3589316 | 18,3333903 | 8,54207749 | 8,62957207 | 5,52056654 | 21,89624 | 8,01405172 | 9,354784 | 12,74392 | 7,292366325 | 38,88013 | 27,74384169 | 47,52832 | 42,666 | 12,40019 | 29,6129618 |
| Upregulated | ENSMUSG00000022422 | 10,57475352 | 17,0202229 | 34,4943799 | 27,9434553 | 2,544425 | 11,94272297 | 5,67946579 | 6,11113011 | 8,54207749 | 0 | 4,41645324 | 8,758497 | 5,00878232 | 0 | 1,59299 | 1,823091581 | 54,59763 | 27,74384169 | 62,73739 | 57,33244 | 11,51446 | 28,7155993 |
| Upregulated | ENSMUSG00000071516 | 4,229901409 | 2,12752787 | 1,81549368 | 3,8542697 | 0 | 2,653938437 | 1,13589316 | 3,66667807 | 1,06775969 | 1,07869651 | 0 | 0 | 1,00175646 | 0 | 2,389485 | 0,911545791 | 16,54474 | 9,908514888 | 19,01133 | 12,66647 | 2,657184 | 4,48681239 |
| Upregulated | ENSMUSG00000021799 | 2,114950705 | 0 | 18,1549368 | 0 | 0 | 0 | 0 | 4,88890409 | 1,06775969 | 0 | 4,41645324 | 13,13775 | 0 | 17,77409 | 11,94742 | 19,1424616 | 11,58132 | 11,89021787 | 34,22039 | 28,66622 | 18,60029 | 2,69208743 |
| Upregulated | ENSMUSG00000109284 | 14,80465493 | 3,1912918 | 17,24719 | 22,1620508 | 0 | 3,980907656 | 9,08714527 | 4,88890409 | 4,27103874 | 1,07869651 | 3,31233993 | 5,255098 | 11,0193211 | 16,83861 | 6,37196 | 30,99255688 | 13,23579 | 10,89936638 | 38,97323 | 49,33256 | 27,45757 | 11,6657122 |
| Upregulated | ENSMUSG00000017412 | 7,402327466 | 1,06376393 | 3,63098736 | 3,8542697 | 0 | 0 | 0 | 7,33335613 | 2,13551937 | 0 | 1,10411331 | 7,006798 | 3,00526939 | 12,16122 | 0,796495 | 10,0270037 | 12,40855 | 7,92681191 | 15,20906 | 11,33316 | 17,71456 | 3,58944991 |
| Upregulated | ENSMUSG00000041625 | 21,14950705 | 11,7014033 | 25,4169115 | 15,4170788 | 6,36106249 | 23,88544594 | 14,7666111 | 50,1112669 | 13,8808759 | 1,07869651 | 15,4575863 | 19,26869 | 43,075528 | 69,2254 | 38,23176 | 28,25791951 | 70,31513 | 44,588317 | 58,93512 | 83,33203 | 73,51543 | 48,4575738 |
| Upregulated | ENSMUSG00000074780 | 16,91960564 | 13,8289311 | 23,6014178 | 15,4170788 | 5,08884999 | 5,307876874 | 11,3589316 | 17,1111643 | 20,287434 | 7,55087556 | 9,93701978 | 21,89624 | 19,0333728 | 34,6127 | 15,1334 | 9,115457906 | 27,29882 | 28,73469318 | 28,51699 | 34,66613 | 30,11475 | 23,3314244 |
| Upregulated | ENSMUSG00000119079 | 10,57475352 | 11,7014033 | 9,98521524 | 9,63567424 | 0 | 1,326969219 | 0 | 0 | 0 | 6,47217906 | 2,20822662 | 21,02039 | 7,01229525 | 25,25792 | 19,91237 | 1,823091581 | 33,08947 | 0,990851489 | 21,86303 | 17,33306 | 11,51446 | 1,79472495 |
| Upregulated | ENSMUSG00000064899 | 59,21861973 | 115,950269 | 162,486684 | 63,59545 | 22,899825 | 69,00239937 | 64,74591 | 25,6667465 | 13,8808759 | 25,8887162 | 18,7699262 | 225,9692 | 29,0509375 | 188,0312 | 237,3555 | 62,89665955 | 159,6567 | 69,35960422 | 157,794 | 121,3314 | 110,716 | 42,1760364 |
| Upregulated | ENSMUSG00000078485 | 45,47144015 | 41,4867934 | 29,9556457 | 26,9798879 | 20,3554 | 31,84726125 | 12,4948247 | 19,5556163 | 7,4743178 | 50,6987359 | 50,7892122 | 169,9148 | 17,0298599 | 30,87079 | 38,23176 | 35,55028583 | 105,8863 | 34,67980211 | 142,585 | 87,33197 | 6,200096 | 52,9443862 |
| Upregulated | ENSMUSG00000021234 | 8,459802819 | 3,1912918 | 12,7084558 | 20,2349159 | 1,2722125 | 6,634846093 | 3,40767947 | 2,44445204 | 6,40655811 | 15,1017511 | 12,1452464 | 35,03399 | 4,00702586 | 11,22574 | 10,35443 | 9,115457906 | 16,54474 | 8,917663399 | 37,07209 | 18,66638 | 8,85728 | 13,4604372 |
| Upregulated | ENSMUSG00000062661 | 234,7595282 | 170,202229 | 511,969218 | 200,422024 | 40,7108 | 53,07876874 | 168,112187 | 108,778116 | 92,8950927 | 100,318775 | 347,795692 | 614,8465 | 406,713125 | 375,1268 | 177,6184 | 411,1071516 | 586,5109 | 216,0056246 | 366,9187 | 299,3287 | 227,6321 | 103,196685 |
| Upregulated | ENSMUSG00000031805 | 419,8177149 | 159,56459 | 290,478989 | 137,790142 | 100,504787 | 173,8329676 | 138,578965 | 30,5556505 | 45,9136665 | 69,0365766 | 214,197982 | 935,4075 | 181,31792 | 322,74 | 131,4217 | 620,7626834 | 505,4417 | 184,2983769 | 562,7354 | 170,664 | 228,5178 | 135,501734 |
| Upregulated | ENSMUSG00000043017 | 517,1054473 | 212,752787 | 521,954433 | 238,964721 | 147,57665 | 237,5274901 | 260,119533 | 267,667499 | 355,563975 | 472,469071 | 545,431975 | 1403,111 | 552,969568 | 676,3509 | 685,7822 | 1137,609147 | 1332,679 | 850,1505774 | 1074,14 | 697,3225 | 418,0636 | 439,707614 |
| Upregulated | ENSMUSG00000037656 | 575,2665917 | 452,099672 | 389,423394 | 284,25239 | 309,147637 | 110,1384451 | 149,937897 | 215,11178 | 150,554116 | 427,163818 | 440,54121 | 1685,135 | 715,254116 | 687,5766 | 991,6362 | 1352,733953 | 1182,949 | 788,7177851 | 1211,022 | 703,3224 | 585,4662 | 946,717414 |
| Upregulated | ENSMUSG00000099835 | 6,344852114 | 5,31881967 | 6,35422788 | 2,89070227 | 3,8166375 | 9,28878453 | 5,67946579 | 12,2222602 | 1,06775969 | 5,39348255 | 9,93701978 | 14,88944 | 10,0580818 | 24,32244 | 14,33691 | 16,40782423 | 9,926842 | 11,89021787 | 20,91246 | 13,99978 | 19,48602 | 11,6657122 |
| Upregulated | ENSMUSG00000049723 | 0 | 7,44634754 | 3,63098736 | 0 | 0 | 2,653938437 | 0 | 0 | 1,06775969 | 9,70826858 | 6,62467985 | 109,4812 | 22,0386422 | 149,6765 | 12,74392 | 24,61173635 | 26,47158 | 59,45108933 | 177,7559 | 98,66513 | 100,973 | 88,8388853 |
| Upregulated | ENSMUSG00000031709 | 0 | 2,12752787 | 0 | 1,92713485 | 0 | 1,326969219 | 5,67946579 | 0 | 1,06775969 | 2,15739302 | 6,62467985 | 53,42683 | 7,01229525 | 43,96748 | 13,54041 | 12,76164107 | 12,40855 | 10,89936638 | 8,555098 | 24,66628 | 18,60029 | 21,5366995 |
| Upregulated | ENSMUSG00000023473 | 2,114950705 | 2,12752787 | 10,8929621 | 5,78140454 | 6,36106249 | 0 | 6,81535895 | 2,44445204 | 0 | 1,07869651 | 11,0411331 | 38,53739 | 3,00526939 | 50,51583 | 26,28433 | 13,67318686 | 4,136184 | 2,972554466 | 20,91246 | 23,33297 | 24,80038 | 26,9208743 |
| Upregulated | ENSMUSG00000016526 | 13,74717958 | 1,06376393 | 9,98521524 | 0,96356742 | 0 | 0 | 1,13589316 | 1,22222602 | 10,6775969 | 0 | 0 | 53,42683 | 5,00878232 | 38,35461 | 20,70887 | 10,0270037 | 4,963421 | 8,917663399 | 26,61586 | 42,666 | 8,85728 | 25,1261494 |
| Upregulated | ENSMUSG00000058427 | 1078,624859 | 871,222662 | 1971,62614 | 951,041048 | 583,945537 | 524,1528414 | 303,283473 | 1015,66982 | 997,287546 | 1224,32054 | 2361,69837 | 13541,51 | 8637,14424 | 7144,248 | 4441,256 | 3975,251193 | 4411,654 | 7818,809098 | 6598,833 | 17451,06 | 18455,91 | 3059,10869 |
| Upregulated | ENSMUSG00000042515 | 8,459802819 | 19,1477508 | 54,4648104 | 32,7612924 | 22,899825 | 35,8281689 | 13,6307179 | 41,5556847 | 40,5748681 | 30,2035023 | 35,3316259 | 94,59177 | 87,1528124 | 79,51566 | 81,24248 | 91,15457906 | 62,87 | 51,52427742 | 36,12153 | 164,6641 | 133,7449 | 62,8153734 |
| Upregulated | ENSMUSG00000028655 | 17,97708099 | 20,2115147 | 28,140152 | 35,6519947 | 26,7164625 | 31,84726125 | 23,8537563 | 66,0002052 | 21,3551937 | 11,8656616 | 46,372759 | 105,9778 | 90,1580818 | 108,5155 | 45,40021 | 35,55028583 | 42,18908 | 43,59746551 | 35,17096 | 134,6646 | 100,0873 | 43,0733989 |
| Upregulated | ENSMUSG00000021130 | 1210,809278 | 840,373508 | 1149,2075 | 1407,77201 | 662,822712 | 1137,21262 | 929,160603 | 1219,78157 | 1377,40999 | 1126,15916 | 1268,62619 | 5475,812 | 2890,0674 | 3184,368 | 2624,451 | 3532,239939 | 1198,666 | 2186,809236 | 1829,841 | 3657,943 | 5950,321 | 2639,14305 |
| Upregulated | ENSMUSG00000052942 | 338,3921127 | 173,393521 | 275,955039 | 100,211012 | 45,79965 | 67,67543015 | 77,2407348 | 45,2223628 | 54,455744 | 165,040566 | 136,91005 | 1661,487 | 845,482456 | 851,2853 | 555,9535 | 745,6444567 | 700,6696 | 390,3954866 | 584,5984 | 1139,982 | 902,5568 | 470,217938 |
| Upregulated | ENSMUSG00000112825 | 702,1636339 | 877,605246 | 1084,75747 | 1108,10254 | 1043,21425 | 893,0502841 | 1117,71887 | 1423,89332 | 1241,80451 | 749,694074 | 767,35875 | 2463,765 | 2894,07443 | 3110,466 | 2332,934 | 2533,185752 | 2159,088 | 2040,163215 | 1795,62 | 3163,951 | 2625,298 | 1830,61945 |
| Upregulated | ENSMUSG00000057113 | 1115,636497 | 1244,6038 | 1441,50198 | 1454,98681 | 1475,7665 | 1560,515801 | 1691,34491 | 2301,4516 | 2025,54012 | 1064,67345 | 1137,23671 | 2998,033 | 4259,46849 | 4972,068 | 3713,259 | 3015,393475 | 2937,518 | 2760,512248 | 2582,689 | 5581,913 | 2901,645 | 2272,12179 |
| Upregulated | ENSMUSG00000028633 | 728,6005177 | 820,161993 | 698,965067 | 1172,66156 | 686,994749 | 414,0143962 | 634,964275 | 942,336263 | 827,513756 | 724,884054 | 1029,0336 | 2834,25 | 2516,41224 | 3014,111 | 2463,559 | 2322,618675 | 2437,867 | 1906,398264 | 2207,215 | 2733,291 | 2289,607 | 1550,64236 |
| Upregulated | ENSMUSG00000090602 | 133,2418944 | 121,269088 | 150,685975 | 162,842895 | 139,943375 | 140,6587372 | 145,394324 | 156,444931 | 167,638271 | 127,286188 | 143,53473 | 401,1392 | 348,61125 | 539,771 | 309,04 | 429,3380674 | 386,3196 | 231,8592484 | 219,5809 | 426,66 | 401,2348 | 211,777545 |
| Upregulated | ENSMUSG00000022325 | 15,86213028 | 20,2115147 | 24,5091647 | 26,0163204 | 27,988675 | 25,21241515 | 24,9896495 | 23,2222944 | 38,4393487 | 23,7313232 | 5,52056654 | 119,1156 | 132,231853 | 143,1282 | 54,95815 | 113,031678 | 114,9859 | 64,40534677 | 88,40268 | 124,6647 | 70,85824 | 118,451847 |
| Upregulated | ENSMUSG00000022015 | 12,68970423 | 2,12752787 | 9,0774684 | 4,81783712 | 3,8166375 | 0 | 2,27178632 | 4,88890409 | 2,13551937 | 33,4395918 | 18,7699262 | 518,503 | 685,201422 | 838,1886 | 275,5873 | 277,1099204 | 600,5739 | 259,6030901 | 557,032 | 913,9857 | 335,6909 | 136,399097 |
| Upregulated | ENSMUSG00000101625 | 37,01163733 | 45,7418492 | 23,6014178 | 15,4170788 | 38,166375 | 22,55847672 | 62,4741237 | 68,4446572 | 57,659023 | 33,4395918 | 71,7673651 | 140,136 | 180,316164 | 174,9345 | 115,4918 | 164,9897881 | 188,61 | 141,6917629 | 180,6076 | 206,6634 | 93,88717 | 92,4283352 |
| Upregulated | ENSMUSG00000001995 | 28,55183451 | 38,2955016 | 48,1105825 | 45,2876689 | 50,8884999 | 63,69452249 | 19,3101837 | 88,0002736 | 59,7945424 | 43,1478604 | 78,3920449 | 345,9606 | [illegible] | 271,2887 | 202,3097 | 235,1788614 | 244,0349 | 150,6094263 | 110,2657 | 265,3292 | 148,8023 | 99,6077235 |
| Upregulated | ENSMUSG00000114642 | 6,344852114 | 0 | 9,0774684 | 1,92713485 | 11,4499125 | 6,634846093 | 2,27178632 | 9,77780817 | 4,27103874 | 8,62957207 | 5,52056654 | 25,39964 | 19,0333728 | 22,45148 | 13,54041 | 20,96555318 | 23,16263 | 14,86277233 | 6,653965 | 21,333 | 9,743008 | 9,87098725 |
| Upregulated | ENSMUSG00000026107 | 103,6325845 | 141,480603 | 173,379646 | 127,1909 | 181,926387 | 147,2935853 | 136,307179 | 242,000752 | 194,332263 | 184,457103 | 153,47175 | 548,2819 | 368,646379 | 420,9653 | 532,8551 | 368,2644994 | 469,0433 | 208,0788127 | 231,9382 | 765,988 | 323,2907 | 256,645669 |
| Upregulated | ENSMUSG00000066952 | 12,68970423 | 13,8289311 | 12,7084558 | 17,3442136 | 7,63327499 | 9,28878453 | 10,2230384 | 13,4444862 | 13,8808759 | 26,9674127 | 23,1863795 | 45,54418 | 26,0456681 | 42,09653 | 26,28433 | 14,58473265 | 24,81711 | 38,64320806 | 28,51699 | 39,33272 | 28,3433 | 22,4340619 |
| Upregulated | ENSMUSG00000043079 | 104,6900599 | 102,121338 | 163,394431 | 108,883119 | 54,7051374 | 54,40573796 | 70,4253758 | 143,000445 | 154,825154 | 139,15185 | 211,989755 | 261,8791 | 197,346024 | 161,8378 | 198,3272 | 195,0707992 | 241,5532 | 174,389862 | 236,6911 | 299,9953 | 201,946 | 138,193822 |
| Upregulated | ENSMUSG00000035268 | 209,3801198 | 265,940983 | 177,010634 | 161,879327 | 90,3270874 | 106,1575375 | 141,986645 | 118,555924 | 137,740999 | 206,031033 | 252,841948 | 356,4708 | 354,621788 | 326,482 | 258,0644 | 345,4758547 | 360,6753 | 289,3286347 | 347,9073 | 453,9929 | 291,4045 | 178,575133 |

**Supplementary Table 7 related to Fig. 6g**

| Order | Ensembl ID | Gene |
|---|---|---|
| 1 | ENSMUSG00000020897 | *Ube2c* |
| 2 | ENSMUSG00000021835 | *Spag5* |
| 3 | ENSMUSG00000012443 | *Kifc2* |
| 4 | ENSMUSG00000023015 | *Cdc20* |
| 5 | ENSMUSG00000001403 | *Kif11* |
| 6 | ENSMUSG00000028873 | *Ncapg* |
| 7 | ENSMUSG00000071176 | *Cdc6* |
| 8 | ENSMUSG00000038943 | *Mybl2* |
| 9 | ENSMUSG00000031004 | *Aurkb* |
| 10 | ENSMUSG00000036777 | *Trip13* |
| 11 | ENSMUSG00000041431 | *Cenpk* |
| 12 | ENSMUSG00000051378 | *Bmp4* |
| 13 | ENSMUSG00000058290 | *Bora* |
| 14 | ENSMUSG00000026622 | *Dscc1* |
| 15 | ENSMUSG00000026683 | *Cep97* |
| 16 | ENSMUSG00000017861 | *Racgap1* |
| 17 | ENSMUSG00000034311 | *Pkmyt1* |
| 18 | ENSMUSG00000028678 | *Sgo1* |
| 19 | ENSMUSG00000045328 | *Ndc80* |
| 20 | ENSMUSG00000037313 | *Dusp1* |
| 21 | ENSMUSG00000027469 | *Cdca5* |
| 22 | ENSMUSG00000006398 | *Kif20b* |
| 23 | ENSMUSG00000041117 | *Cep55* |
| 24 | ENSMUSG00000043065 | *Sgo2a* |
| 25 | ENSMUSG00000004187 | *Cenpf* |
| 26 | ENSMUSG00000022604 | *Nek2* |
| 27 | ENSMUSG00000030867 | *Nuf2* |
| 28 | ENSMUSG00000044201 | *Nek6* |
| 29 | ENSMUSG00000027306 | *Nusap1* |
| 30 | ENSMUSG00000015880 | *Knstrn* |
| 31 | ENSMUSG00000031906 | *Bub1* |
| 32 | ENSMUSG00000002055 | *Tpx2* |
| 33 | ENSMUSG00000027331 | *Aurka* |
| 34 | ENSMUSG00000026605 | *Edn3* |
| 35 | ENSMUSG00000024056 | *Dsn1* |
| 36 | ENSMUSG00000062510 | *Smc2* |
| 37 | ENSMUSG00000029414 | *Kif2c* |
| 38 | ENSMUSG00000032254 | *Cdca8* |
| 39 | ENSMUSG00000021714 | *Rcc1* |
| 40 | ENSMUSG00000051235 | *Ereg* |
| 41 | ENSMUSG00000030677 | *Kntc1* |
| 42 | ENSMUSG00000038379 | *Chek2* |
| 43 | ENSMUSG00000022422 | *Mad2l1* |
| 44 | ENSMUSG00000034906 | *Kif22* |
| 45 | ENSMUSG00000029521 | *Plk1* |
| 46 | ENSMUSG00000040084 | *Mki67* |
| 47 | ENSMUSG00000027379 | *Smpd3* |
| 48 | ENSMUSG00000024795 | *Chek1* |
| 49 | ENSMUSG00000024791 | *Kif23* |
| 50 | ENSMUSG00000050107 | *Zwilch* |
| 51 | ENSMUSG00000024989 | *Cdc14b* |
| 52 | ENSMUSG00000026039 | *Ctdp1* |
| 53 | ENSMUSG00000079553 | *Kif4* |
| 54 | ENSMUSG00000026749 | *Ncaph* |
| 55 | ENSMUSG00000029377 | *Kif15* |
| 56 | ENSMUSG00000033323 | *Anln* |
| 57 | ENSMUSG00000033102 | *Tacc3* |
| 58 | ENSMUSG00000027524 | *Cep85* |
| 59 | ENSMUSG00000028896 | *Dlgap5* |
| 60 | ENSMUSG00000037443 | *Ncapd2* |
| 61 | ENSMUSG00000021569 | *Ttk* |
| 62 | ENSMUSG00000042029 | *Prc1* |
| 63 | ENSMUSG00000027635 | *Bub1b* |
| 64 | ENSMUSG00000023908 | *Ccdc8* |
| 65 | ENSMUSG00000029910 | *Ccnb1* |
| 66 | ENSMUSG00000028312 | *Ncapg2* |
| 67 | ENSMUSG00000037544 | *Spice1* |
| 68 | ENSMUSG00000036768 | *Cdc25c* |
| 69 | ENSMUSG00000017499 | *Cenpe* |
| 70 | ENSMUSG00000069910 | *Phf13* |
| 71 | ENSMUSG00000032400 | *Cdca2* |
| 72 | ENSMUSG00000027496 | *Haspin* |
| 73 | ENSMUSG00000032113 | *Gen1* |
| 74 | ENSMUSG00000023940 | *Kif18b* |
| 75 | ENSMUSG00000024190 | *Espl1* |
| 76 | ENSMUSG00000047777 | *Nsl1* |
| 77 | ENSMUSG00000063060 | *Sox7* |
| 78 | ENSMUSG00000022070 | *Spdl1* |
| 79 | ENSMUSG00000038252 | *Arhgef10* |
| 80 | ENSMUSG00000048922 | *Kifc1* |

**Supplementary Table 8 related to Supplementary Fig. 6l REACTOME_SIGNALING_BY_WNT heatmap genes**

| Order | Ensembl I | Gene |
|---|---|---|
| 1 | ENSMUSG | *Gnao1* |
| 2 | ENSMUSG | *Psmb9* |
| 3 | ENSMUSG | *H2bc4* |
| 4 | ENSMUSG | *H2bc6* |
| 5 | ENSMUSG | *Camk2a* |
| 6 | ENSMUSG | *H2bc7* |
| 7 | ENSMUSG | *H2ac23* |
| 8 | ENSMUSG | *Rspo4* |
| 9 | ENSMUSG | *Wnt2* |
| 10 | ENSMUSG | *H2bc8* |
| 11 | ENSMUSG | *H2bc21* |
| 12 | ENSMUSG | *Gng13* |
| 13 | ENSMUSG | *H2ac11* |
| 14 | ENSMUSG | *Rspo2* |
| 15 | ENSMUSG | *Wnt10a* |
| 16 | ENSMUSG | *Rspo1* |
| 17 | ENSMUSG | *Wnt11* |
| 18 | ENSMUSG | *Wnt6* |
| 19 | ENSMUSG | *Lgr5* |
| 20 | ENSMUSG | *Sfrp1* |
| 21 | ENSMUSG | *Porcn* |
| 22 | ENSMUSG | *Sfrp2* |
| 23 | ENSMUSG | *Fzd4* |
| 24 | ENSMUSG | *Gng8* |
| 25 | ENSMUSG | *H2ac13* |
| 26 | ENSMUSG | *Kremen1* |
| 27 | ENSMUSG | *Ror2* |
| 28 | ENSMUSG | *Tcf4* |
| 29 | ENSMUSG | *Wnt5a* |
| 30 | ENSMUSG | *Nfatc1* |
| 31 | ENSMUSG | *Prickle1* |
| 32 | ENSMUSG | *Tle1* |
| 33 | ENSMUSG | *Axin2* |
| 34 | ENSMUSG | *Dkk2* |
| 35 | ENSMUSG | *Gng4* |
| 36 | ENSMUSG | *Sox9* |
| 37 | ENSMUSG | *Sox7* |

**Hashing primers**

| Primer | Sequence (5'-3') |
|---|---|
| HTO primer | GTGACTGGAGTTCAGACGTGTGC*T*C |
| 10x Genomics SI-PCR primer | AATGATACGGCGACCACCGAGATCTACACTCTTTCCCTACACGACGC*T*C |
| Illumina TruSeq D701_S | CAAGCAGAAGACGGCATACGAGATCGAGTAATGTGACTGGAGTTCAGACGTGT*G*C |
| Illumina TruSeq D702_S | CAAGCAGAAGACGGCATACGAGATTCTCCGGAGTGACTGGAGTTCAGACGTGT*G*C |
| Illumina TruSeq D703_S | CAAGCAGAAGACGGCATACGAGATAATGAGCGGTGACTGGAGTTCAGACGTGT*G*C |
| Illumina TruSeq D704_S | CAAGCAGAAGACGGCATACGAGATGGAATCTCGTGACTGGAGTTCAGACGTGT*G*C |
| Illumina TruSeq D705_S | CAAGCAGAAGACGGCATACGAGATTTCTGAATGTGACTGGAGTTCAGACGTGT*G*C |
| Illumina TruSeq D706_S | CAAGCAGAAGACGGCATACGAGATACGAATTCGTGACTGGAGTTCAGACGTGT*G*C |

**Xenium gene panel**

| Gene | Ensembl ID | Probesets | Panel |
|---|---|---|---|
| *Ar* | ENSMUSG00000046532 | 8 | Custom |
| *Atf3* | ENSMUSG00000026628 | 8 | Custom |
| *Atf6* | ENSMUSG00000026663 | 8 | Custom |
| *Atp5g1* | ENSMUSG00000006057 | 8 | Custom |
| *Axin2* | ENSMUSG00000000142 | 8 | Custom |
| *Ccbe1* | ENSMUSG00000046318 | 8 | Custom |
| *Ccn1* | ENSMUSG00000028195 | 8 | Custom |
| *Ccn2* | ENSMUSG00000019997 | 8 | Custom |
| *Ccnd2* | ENSMUSG00000000184 | 8 | Custom |
| *Cd34* | ENSMUSG00000016494 | 7 | Custom |
| *Cd9* | ENSMUSG00000030342 | 8 | Custom |
| *Cebpd* | ENSMUSG00000071637 | 7 | Custom |
| *Clmp* | ENSMUSG00000032024 | 8 | Custom |
| *Col5a1* | ENSMUSG00000026837 | 8 | Custom |
| *Col5a2* | ENSMUSG00000026042 | 8 | Custom |
| *Corin* | ENSMUSG00000005220 | 8 | Custom |
| *Cpz* | ENSMUSG00000036596 | 8 | Custom |
| *Creb5* | ENSMUSG00000053007 | 8 | Custom |
| *Cyb5r3* | ENSMUSG00000018042 | 8 | Custom |
| *Dkk2* | ENSMUSG00000028031 | 8 | Custom |
| *Egr1* | ENSMUSG00000038418 | 8 | Custom |
| *Egr3* | ENSMUSG00000033730 | 8 | Custom |
| *Ehd2* | ENSMUSG00000074364 | 8 | Custom |
| *Elf1* | ENSMUSG00000036461 | 8 | Custom |
| *Esd* | ENSMUSG00000021996 | 8 | Custom |
| *Fmod* | ENSMUSG00000041559 | 8 | Custom |
| *Fn1* | ENSMUSG00000026193 | 8 | Custom |
| *Fst* | ENSMUSG00000021765 | 8 | Custom |
| *Fxyd5* | ENSMUSG00000009687 | 8 | Custom |
| *Gatad1* | ENSMUSG00000007415 | 8 | Custom |
| *Gfpt2* | ENSMUSG00000020363 | 7 | Custom |
| *Gpx3* | ENSMUSG00000018339 | 8 | Custom |
| *Gsn* | ENSMUSG00000026879 | 8 | Custom |
| *Hmga2* | ENSMUSG00000056758 | 8 | Custom |
| *Igfbp5* | ENSMUSG00000026185 | 8 | Custom |
| *Il13ra1* | ENSMUSG00000017057 | 8 | Custom |
| *Il17ra* | ENSMUSG00000002897 | 8 | Custom |
| *Inhba* | ENSMUSG00000041324 | 8 | Custom |
| *Itgb1* | ENSMUSG00000025809 | 8 | Custom |
| *Jun* | ENSMUSG00000052684 | 8 | Custom |
| *Jund* | ENSMUSG00000071076 | 8 | Custom |
| *Klf10* | ENSMUSG00000037465 | 8 | Custom |
| *Klf2* | ENSMUSG00000055148 | 8 | Custom |
| *Krt1* | ENSMUSG00000046834 | 8 | Custom |
| *Krt10* | ENSMUSG00000019761 | 8 | Custom |
| *Krt14* | ENSMUSG00000045545 | 8 | Custom |
| *Krt15* | ENSMUSG00000054146 | 8 | Custom |
| *Lamb3* | ENSMUSG00000026639 | 8 | Custom |
| *Lef1* | ENSMUSG00000027985 | 8 | Custom |
| *Lgr5* | ENSMUSG00000020140 | 8 | Custom |
| *Lrig1* | ENSMUSG00000030029 | 5 | Custom |
| *Lrp1* | ENSMUSG00000040249 | 8 | Custom |
| *Ly6a* | ENSMUSG00000075602 | 8 | Custom |

| | | | |
|---|---|---|---|
| *Maf* | ENSMUSG00000055435 | 8 | Custom |
| *Mafb* | ENSMUSG00000074622 | 8 | Custom |
| *Mgst1* | ENSMUSG00000008540 | 8 | Custom |
| *Mme* | ENSMUSG00000027820 | 8 | Custom |
| *Mpz* | ENSMUSG00000056569 | 8 | Custom |
| *Myod1* | ENSMUSG00000009471 | 8 | Custom |
| *Nfib* | ENSMUSG00000008575 | 8 | Custom |
| *Nhp2* | ENSMUSG00000001056 | 8 | Custom |
| *Nme1* | ENSMUSG00000037601 | 8 | Custom |
| *Notum* | ENSMUSG00000042988 | 8 | Custom |
| *Nr1d2* | ENSMUSG00000021775 | 8 | Custom |
| *Nr2f2* | ENSMUSG00000030551 | 8 | Custom |
| *Nrp1* | ENSMUSG00000025810 | 8 | Custom |
| *Pdgfra* | ENSMUSG00000029231 | 8 | Custom |
| *Pdpn* | ENSMUSG00000028583 | 8 | Custom |
| *Pecam1* | ENSMUSG00000020717 | 8 | Custom |
| *Phldb2* | ENSMUSG00000033149 | 8 | Custom |
| *Plagl1* | ENSMUSG00000019817 | 8 | Custom |
| *Ppp1r14a* | ENSMUSG00000037166 | 8 | Custom |
| *Prrx1* | ENSMUSG00000026586 | 8 | Custom |
| *Ptprc* | ENSMUSG00000026395 | 8 | Custom |
| *Rbm3* | ENSMUSG00000031167 | 8 | Custom |
| *Scd1* | ENSMUSG00000037071 | 8 | Custom |
| *Sh3bgrl3* | ENSMUSG00000028843 | 5 | Custom |
| *Slc6a8* | ENSMUSG00000019558 | 8 | Custom |
| *Slit3* | ENSMUSG00000056427 | 8 | Custom |
| *Smoc2* | ENSMUSG00000023886 | 8 | Custom |
| *Sod2* | ENSMUSG00000006818 | 8 | Custom |
| *Sprr1a* | ENSMUSG00000050359 | 8 | Custom |
| *Srm* | ENSMUSG00000006442 | 8 | Custom |
| *Stat5b* | ENSMUSG00000020919 | 8 | Custom |
| *Tef* | ENSMUSG00000022389 | 8 | Custom |
| *Tfap2a* | ENSMUSG00000021359 | 8 | Custom |
| *Tfap2c* | ENSMUSG00000028640 | 8 | Custom |
| *Thbs1* | ENSMUSG00000040152 | 7 | Custom |
| *Timp1* | ENSMUSG00000001131 | 8 | Custom |
| *Tnc* | ENSMUSG00000028364 | 8 | Custom |
| *Uap1* | ENSMUSG00000026670 | 8 | Custom |
| *Ugdh* | ENSMUSG00000029201 | 8 | Custom |
| *Wnt5a* | ENSMUSG00000021994 | 8 | Custom |
| *Xdh* | ENSMUSG00000024066 | 8 | Custom |
| *Zbtb4* | ENSMUSG00000018750 | 8 | Custom |
| *0610005C* | ENSMUSG00000109644 | 6 | Mouse Tissue Atlassing |
| *1110017D* | ENSMUSG00000028441 | 7 | Mouse Tissue Atlassing |
| *2610528A* | ENSMUSG00000096001 | 8 | Mouse Tissue Atlassing |
| *5330417C* | ENSMUSG00000040412 | 8 | Mouse Tissue Atlassing |
| *6330403K* | ENSMUSG00000018451 | 8 | Mouse Tissue Atlassing |
| *A330076H* | ENSMUSG00000109321 | 8 | Mouse Tissue Atlassing |
| *Aadat* | ENSMUSG00000057228 | 8 | Mouse Tissue Atlassing |
| *Abca13* | ENSMUSG00000004668 | 8 | Mouse Tissue Atlassing |
| *Abcb11* | ENSMUSG00000027048 | 8 | Mouse Tissue Atlassing |
| *Acan* | ENSMUSG00000030607 | 8 | Mouse Tissue Atlassing |
| *Ace2* | ENSMUSG00000015405 | 8 | Mouse Tissue Atlassing |
| *Ackr3* | ENSMUSG00000044337 | 8 | Mouse Tissue Atlassing |
| *Actl6b* | ENSMUSG00000029712 | 8 | Mouse Tissue Atlassing |
| *Actn3* | ENSMUSG00000006457 | 8 | Mouse Tissue Atlassing |

| | | | |
|---|---|---|---|
| *Ager* | ENSMUSG00000015452 | 6 | Mouse Tissue Atlassing |
| *Aif1l* | ENSMUSG00000001864 | 8 | Mouse Tissue Atlassing |
| *Aldh1b1* | ENSMUSG00000035561 | 8 | Mouse Tissue Atlassing |
| *Alox5ap* | ENSMUSG00000060063 | 8 | Mouse Tissue Atlassing |
| *Ambp* | ENSMUSG00000028356 | 8 | Mouse Tissue Atlassing |
| *Angpt2* | ENSMUSG00000031465 | 8 | Mouse Tissue Atlassing |
| *Anxa2* | ENSMUSG00000032231 | 8 | Mouse Tissue Atlassing |
| *Anxa8* | ENSMUSG00000021950 | 8 | Mouse Tissue Atlassing |
| *Aplp1* | ENSMUSG00000006651 | 8 | Mouse Tissue Atlassing |
| *Apoc2* | ENSMUSG00000002992 | 6 | Mouse Tissue Atlassing |
| *Apoc4* | ENSMUSG00000074336 | 2 | Mouse Tissue Atlassing |
| *Aqp1* | ENSMUSG00000004655 | 8 | Mouse Tissue Atlassing |
| *Aqp3* | ENSMUSG00000028435 | 8 | Mouse Tissue Atlassing |
| *Aqp4* | ENSMUSG00000024411 | 8 | Mouse Tissue Atlassing |
| *Aqp5* | ENSMUSG00000044217 | 5 | Mouse Tissue Atlassing |
| *Aqp7* | ENSMUSG00000028427 | 8 | Mouse Tissue Atlassing |
| *Areg* | ENSMUSG00000029378 | 8 | Mouse Tissue Atlassing |
| *Arpp21* | ENSMUSG00000032503 | 8 | Mouse Tissue Atlassing |
| *Asb9* | ENSMUSG00000031384 | 8 | Mouse Tissue Atlassing |
| *Ascl1* | ENSMUSG00000020052 | 8 | Mouse Tissue Atlassing |
| *Aspn* | ENSMUSG00000021388 | 8 | Mouse Tissue Atlassing |
| *Asprv1* | ENSMUSG00000033508 | 7 | Mouse Tissue Atlassing |
| *Asz1* | ENSMUSG00000010796 | 8 | Mouse Tissue Atlassing |
| *Atp6v0d2* | ENSMUSG00000028238 | 8 | Mouse Tissue Atlassing |
| *Atp6v1b1* | ENSMUSG00000006269 | 8 | Mouse Tissue Atlassing |
| *AU021092* | ENSMUSG00000051669 | 8 | Mouse Tissue Atlassing |
| *B4galnt2* | ENSMUSG00000013418 | 8 | Mouse Tissue Atlassing |
| *Bgn* | ENSMUSG00000031375 | 5 | Mouse Tissue Atlassing |
| *Bmp2* | ENSMUSG00000027358 | 8 | Mouse Tissue Atlassing |
| *Bst1* | ENSMUSG00000029082 | 8 | Mouse Tissue Atlassing |
| *C3* | ENSMUSG00000024164 | 5 | Mouse Tissue Atlassing |
| *C5ar1* | ENSMUSG00000049130 | 8 | Mouse Tissue Atlassing |
| *Calb1* | ENSMUSG00000028222 | 8 | Mouse Tissue Atlassing |
| *Calcrl* | ENSMUSG00000059588 | 8 | Mouse Tissue Atlassing |
| *Calm4* | ENSMUSG00000033765 | 8 | Mouse Tissue Atlassing |
| *Calm5* | ENSMUSG00000099269 | 7 | Mouse Tissue Atlassing |
| *Capns2* | ENSMUSG00000078144 | 8 | Mouse Tissue Atlassing |
| *Car2* | ENSMUSG00000027562 | 4 | Mouse Tissue Atlassing |
| *Car3* | ENSMUSG00000027559 | 8 | Mouse Tissue Atlassing |
| *Cav1* | ENSMUSG00000007655 | 8 | Mouse Tissue Atlassing |
| *Cavin2* | ENSMUSG00000045954 | 8 | Mouse Tissue Atlassing |
| *Ccdc153* | ENSMUSG00000070306 | 7 | Mouse Tissue Atlassing |
| *Ccdc80* | ENSMUSG00000022665 | 8 | Mouse Tissue Atlassing |
| *Ccl12* | ENSMUSG00000035352 | 4 | Mouse Tissue Atlassing |
| *Ccl9* | ENSMUSG00000019122 | 8 | Mouse Tissue Atlassing |
| *Ccn3* | ENSMUSG00000037362 | 8 | Mouse Tissue Atlassing |
| *Cd300lf* | ENSMUSG00000047798 | 8 | Mouse Tissue Atlassing |
| *Cd36* | ENSMUSG00000002944 | 8 | Mouse Tissue Atlassing |
| *Cd3d* | ENSMUSG00000032094 | 8 | Mouse Tissue Atlassing |
| *Cd5l* | ENSMUSG00000015854 | 8 | Mouse Tissue Atlassing |
| *Cd8a* | ENSMUSG00000053977 | 8 | Mouse Tissue Atlassing |
| *Cdh16* | ENSMUSG00000031881 | 8 | Mouse Tissue Atlassing |
| *Cdhr5* | ENSMUSG00000025497 | 8 | Mouse Tissue Atlassing |
| *Cdx1* | ENSMUSG00000024619 | 7 | Mouse Tissue Atlassing |
| *Celf3* | ENSMUSG00000028137 | 8 | Mouse Tissue Atlassing |
| *Cenpf* | ENSMUSG00000026605 | 8 | Mouse Tissue Atlassing |

| | | | |
|---|---|---|---|
| *Cfh* | ENSMUSG00000026365 | 8 | Mouse Tissue Atlassing |
| *Chgb* | ENSMUSG00000027350 | 8 | Mouse Tissue Atlassing |
| *Chl1* | ENSMUSG00000030077 | 8 | Mouse Tissue Atlassing |
| *Chodl* | ENSMUSG00000022860 | 8 | Mouse Tissue Atlassing |
| *Clcnka* | ENSMUSG00000033770 | 8 | Mouse Tissue Atlassing |
| *Clcnkb* | ENSMUSG00000006216 | 8 | Mouse Tissue Atlassing |
| *Cldn11* | ENSMUSG00000037625 | 8 | Mouse Tissue Atlassing |
| *Cldn2* | ENSMUSG00000047230 | 8 | Mouse Tissue Atlassing |
| *Cldn3* | ENSMUSG00000070473 | 1 | Mouse Tissue Atlassing |
| *Cldn4* | ENSMUSG00000047501 | 7 | Mouse Tissue Atlassing |
| *Cldn7* | ENSMUSG00000018569 | 6 | Mouse Tissue Atlassing |
| *Clec14a* | ENSMUSG00000045930 | 8 | Mouse Tissue Atlassing |
| *Clec4a3* | ENSMUSG00000043832 | 8 | Mouse Tissue Atlassing |
| *Clec4f* | ENSMUSG00000014542 | 8 | Mouse Tissue Atlassing |
| *Cndp2* | ENSMUSG00000024644 | 8 | Mouse Tissue Atlassing |
| *Cnfn* | ENSMUSG00000063651 | 1 | Mouse Tissue Atlassing |
| *Cnn1* | ENSMUSG00000001349 | 8 | Mouse Tissue Atlassing |
| *Coch* | ENSMUSG00000020953 | 8 | Mouse Tissue Atlassing |
| *Col17a1* | ENSMUSG00000025064 | 8 | Mouse Tissue Atlassing |
| *Col2a1* | ENSMUSG00000022483 | 8 | Mouse Tissue Atlassing |
| *Col8a1* | ENSMUSG00000068196 | 8 | Mouse Tissue Atlassing |
| *Colec11* | ENSMUSG00000036655 | 8 | Mouse Tissue Atlassing |
| *Comp* | ENSMUSG00000031849 | 8 | Mouse Tissue Atlassing |
| *Cox8b* | ENSMUSG00000025488 | 2 | Mouse Tissue Atlassing |
| *Cp* | ENSMUSG00000003617 | 8 | Mouse Tissue Atlassing |
| *Cpa3* | ENSMUSG00000001865 | 8 | Mouse Tissue Atlassing |
| *Cplx2* | ENSMUSG00000025867 | 8 | Mouse Tissue Atlassing |
| *Crct1* | ENSMUSG00000027913 | 5 | Mouse Tissue Atlassing |
| *Cryab* | ENSMUSG00000032060 | 7 | Mouse Tissue Atlassing |
| *Csf3r* | ENSMUSG00000028859 | 8 | Mouse Tissue Atlassing |
| *Cstdc4* | ENSMUSG00000079597 | 5 | Mouse Tissue Atlassing |
| *Ctla2a* | ENSMUSG00000044258 | 8 | Mouse Tissue Atlassing |
| *Ctla4* | ENSMUSG00000026011 | 8 | Mouse Tissue Atlassing |
| *Ctse* | ENSMUSG00000004552 | 8 | Mouse Tissue Atlassing |
| *Ctsg* | ENSMUSG00000040314 | 8 | Mouse Tissue Atlassing |
| *Ctss* | ENSMUSG00000038642 | 8 | Mouse Tissue Atlassing |
| *Cubn* | ENSMUSG00000026726 | 8 | Mouse Tissue Atlassing |
| *Cux2* | ENSMUSG00000042589 | 8 | Mouse Tissue Atlassing |
| *Cxcr2* | ENSMUSG00000026180 | 8 | Mouse Tissue Atlassing |
| *Cybb* | ENSMUSG00000015340 | 8 | Mouse Tissue Atlassing |
| *Cyp11a1* | ENSMUSG00000032323 | 8 | Mouse Tissue Atlassing |
| *Cyp4b1* | ENSMUSG00000028713 | 8 | Mouse Tissue Atlassing |
| *Dao* | ENSMUSG00000042096 | 8 | Mouse Tissue Atlassing |
| *Dbpht2* | ENSMUSG00000029878 | 8 | Mouse Tissue Atlassing |
| *Dcdc2a* | ENSMUSG00000035910 | 8 | Mouse Tissue Atlassing |
| *Dcpp2* | ENSMUSG00000096278 | 7 | Mouse Tissue Atlassing |
| *Des* | ENSMUSG00000026208 | 8 | Mouse Tissue Atlassing |
| *Dnase1l3* | ENSMUSG00000025279 | 8 | Mouse Tissue Atlassing |
| *Dpt* | ENSMUSG00000026574 | 8 | Mouse Tissue Atlassing |
| *Dsc3* | ENSMUSG00000059898 | 8 | Mouse Tissue Atlassing |
| *Dynlrb2* | ENSMUSG00000034467 | 6 | Mouse Tissue Atlassing |
| *Ecrg4* | ENSMUSG00000026051 | 6 | Mouse Tissue Atlassing |
| *Emb* | ENSMUSG00000021728 | 8 | Mouse Tissue Atlassing |
| *Eng* | ENSMUSG00000026814 | 8 | Mouse Tissue Atlassing |
| *Epcam* | ENSMUSG00000045394 | 8 | Mouse Tissue Atlassing |
| *Epsti1* | ENSMUSG00000022014 | 8 | Mouse Tissue Atlassing |

| | | | |
|---|---|---|---|
| *Etv1* | ENSMUSG00000004151 | 8 | Mouse Tissue Atlassing |
| *F13a1* | ENSMUSG00000039109 | 8 | Mouse Tissue Atlassing |
| *F3* | ENSMUSG00000028128 | 8 | Mouse Tissue Atlassing |
| *Fam183b* | ENSMUSG00000049154 | 5 | Mouse Tissue Atlassing |
| *Fam25c* | ENSMUSG00000043681 | 2 | Mouse Tissue Atlassing |
| *Fbln5* | ENSMUSG00000021186 | 8 | Mouse Tissue Atlassing |
| *Fcnb* | ENSMUSG00000026835 | 8 | Mouse Tissue Atlassing |
| *Fech* | ENSMUSG00000024588 | 8 | Mouse Tissue Atlassing |
| *Fibin* | ENSMUSG00000074971 | 8 | Mouse Tissue Atlassing |
| *Fmo5* | ENSMUSG00000028088 | 8 | Mouse Tissue Atlassing |
| *Folr2* | ENSMUSG00000032725 | 8 | Mouse Tissue Atlassing |
| *Fxyd1* | ENSMUSG00000036570 | 1 | Mouse Tissue Atlassing |
| *Fxyd3* | ENSMUSG00000057092 | 8 | Mouse Tissue Atlassing |
| *Fxyd4* | ENSMUSG00000004988 | 8 | Mouse Tissue Atlassing |
| *Fxyd6* | ENSMUSG00000066705 | 8 | Mouse Tissue Atlassing |
| *Gabrp* | ENSMUSG00000020159 | 8 | Mouse Tissue Atlassing |
| *Gad1* | ENSMUSG00000070880 | 8 | Mouse Tissue Atlassing |
| *Gamt* | ENSMUSG00000020150 | 3 | Mouse Tissue Atlassing |
| *Gap43* | ENSMUSG00000047261 | 8 | Mouse Tissue Atlassing |
| *Gatm* | ENSMUSG00000027199 | 8 | Mouse Tissue Atlassing |
| *Gc* | ENSMUSG00000035540 | 8 | Mouse Tissue Atlassing |
| *Gfap* | ENSMUSG00000020932 | 8 | Mouse Tissue Atlassing |
| *Gjb4* | ENSMUSG00000046623 | 8 | Mouse Tissue Atlassing |
| *Gm13889* | ENSMUSG00000087006 | 3 | Mouse Tissue Atlassing |
| *Gm94* | ENSMUSG00000071858 | 7 | Mouse Tissue Atlassing |
| *Gng11* | ENSMUSG00000032766 | 5 | Mouse Tissue Atlassing |
| *Gp2* | ENSMUSG00000030954 | 8 | Mouse Tissue Atlassing |
| *Gpa33* | ENSMUSG00000000544 | 8 | Mouse Tissue Atlassing |
| *Gpihbp1* | ENSMUSG00000022579 | 8 | Mouse Tissue Atlassing |
| *Gpx2* | ENSMUSG00000042808 | 8 | Mouse Tissue Atlassing |
| *Gpx6* | ENSMUSG00000004341 | 8 | Mouse Tissue Atlassing |
| *Gria2* | ENSMUSG00000033981 | 8 | Mouse Tissue Atlassing |
| *Guca2b* | ENSMUSG00000032978 | 5 | Mouse Tissue Atlassing |
| *Hap1* | ENSMUSG00000006930 | 8 | Mouse Tissue Atlassing |
| *Hc* | ENSMUSG00000026874 | 8 | Mouse Tissue Atlassing |
| *Hemgn* | ENSMUSG00000028332 | 8 | Mouse Tissue Atlassing |
| *Higd1b* | ENSMUSG00000020928 | 3 | Mouse Tissue Atlassing |
| *Hmbs* | ENSMUSG00000032126 | 8 | Mouse Tissue Atlassing |
| *Hmgcs2* | ENSMUSG00000027875 | 8 | Mouse Tissue Atlassing |
| *Hp* | ENSMUSG00000031722 | 5 | Mouse Tissue Atlassing |
| *Hpgd* | ENSMUSG00000031613 | 4 | Mouse Tissue Atlassing |
| *Hpx* | ENSMUSG00000030895 | 8 | Mouse Tissue Atlassing |
| *Hrc* | ENSMUSG00000038239 | 8 | Mouse Tissue Atlassing |
| *Hrg* | ENSMUSG00000022877 | 8 | Mouse Tissue Atlassing |
| *Hsd11b2* | ENSMUSG00000031891 | 8 | Mouse Tissue Atlassing |
| *Hsd3b4* | ENSMUSG00000095143 | 8 | Mouse Tissue Atlassing |
| *Htra3* | ENSMUSG00000029096 | 8 | Mouse Tissue Atlassing |
| *Ifitm6* | ENSMUSG00000059108 | 8 | Mouse Tissue Atlassing |
| *Igfbp6* | ENSMUSG00000023046 | 5 | Mouse Tissue Atlassing |
| *Il1r2* | ENSMUSG00000026073 | 8 | Mouse Tissue Atlassing |
| *Insm1* | ENSMUSG00000068154 | 8 | Mouse Tissue Atlassing |
| *Isca1* | ENSMUSG00000044792 | 8 | Mouse Tissue Atlassing |
| *Isg20* | ENSMUSG00000039236 | 3 | Mouse Tissue Atlassing |
| *Itga8* | ENSMUSG00000026768 | 8 | Mouse Tissue Atlassing |
| *Ivl* | ENSMUSG00000049128 | 8 | Mouse Tissue Atlassing |
| *Iyd* | ENSMUSG00000019762 | 8 | Mouse Tissue Atlassing |

| | | | |
|---|---|---|---|
| *Kcna1* | ENSMUSG00000047976 | 8 | Mouse Tissue Atlassing |
| *Kcnj1* | ENSMUSG00000041248 | 8 | Mouse Tissue Atlassing |
| *Kcnj16* | ENSMUSG00000051497 | 8 | Mouse Tissue Atlassing |
| *Kdr* | ENSMUSG00000062960 | 8 | Mouse Tissue Atlassing |
| *Kl* | ENSMUSG00000058488 | 8 | Mouse Tissue Atlassing |
| *Klra8* | ENSMUSG00000089727 | 8 | Mouse Tissue Atlassing |
| *Kng2* | ENSMUSG00000060459 | 8 | Mouse Tissue Atlassing |
| *Krt13* | ENSMUSG00000044041 | 8 | Mouse Tissue Atlassing |
| *Krt16* | ENSMUSG00000053797 | 8 | Mouse Tissue Atlassing |
| *Krt19* | ENSMUSG00000020911 | 8 | Mouse Tissue Atlassing |
| *Krt79* | ENSMUSG00000061397 | 8 | Mouse Tissue Atlassing |
| *Krt8* | ENSMUSG00000049382 | 8 | Mouse Tissue Atlassing |
| *Lamp3* | ENSMUSG00000041247 | 8 | Mouse Tissue Atlassing |
| *Laptm5* | ENSMUSG00000028581 | 8 | Mouse Tissue Atlassing |
| *Lce1m* | ENSMUSG00000027912 | 4 | Mouse Tissue Atlassing |
| *Ldhb* | ENSMUSG00000030246 | 8 | Mouse Tissue Atlassing |
| *Lhx2* | ENSMUSG00000000247 | 8 | Mouse Tissue Atlassing |
| *Lipg* | ENSMUSG00000053846 | 8 | Mouse Tissue Atlassing |
| *Lor* | ENSMUSG00000043165 | 7 | Mouse Tissue Atlassing |
| *Lrp2* | ENSMUSG00000027070 | 8 | Mouse Tissue Atlassing |
| *Lst1* | ENSMUSG00000073412 | 4 | Mouse Tissue Atlassing |
| *Ltc4s* | ENSMUSG00000020377 | 2 | Mouse Tissue Atlassing |
| *Lum* | ENSMUSG00000036446 | 8 | Mouse Tissue Atlassing |
| *Ly6g6c* | ENSMUSG00000092586 | 8 | Mouse Tissue Atlassing |
| *Ly6h* | ENSMUSG00000022577 | 8 | Mouse Tissue Atlassing |
| *Lyve1* | ENSMUSG00000030787 | 8 | Mouse Tissue Atlassing |
| *Mamdc2* | ENSMUSG00000033207 | 8 | Mouse Tissue Atlassing |
| *Mapk10* | ENSMUSG00000046709 | 8 | Mouse Tissue Atlassing |
| *Marco* | ENSMUSG00000026390 | 8 | Mouse Tissue Atlassing |
| *Mettl7a2* | ENSMUSG00000056487 | 8 | Mouse Tissue Atlassing |
| *Mfap4* | ENSMUSG00000042436 | 6 | Mouse Tissue Atlassing |
| *Mfap5* | ENSMUSG00000030116 | 8 | Mouse Tissue Atlassing |
| *Mgat4c* | ENSMUSG00000019888 | 8 | Mouse Tissue Atlassing |
| *Mgll* | ENSMUSG00000033174 | 8 | Mouse Tissue Atlassing |
| *Mlc1* | ENSMUSG00000035805 | 8 | Mouse Tissue Atlassing |
| *Mmp8* | ENSMUSG00000005800 | 8 | Mouse Tissue Atlassing |
| *Mmp9* | ENSMUSG00000017737 | 8 | Mouse Tissue Atlassing |
| *Mmrn1* | ENSMUSG00000054641 | 8 | Mouse Tissue Atlassing |
| *Mpeg1* | ENSMUSG00000046805 | 8 | Mouse Tissue Atlassing |
| *Mpo* | ENSMUSG00000009350 | 8 | Mouse Tissue Atlassing |
| *Mrgpra2a* | ENSMUSG00000093973 | 8 | Mouse Tissue Atlassing |
| *Ms4a6b* | ENSMUSG00000024677 | 8 | Mouse Tissue Atlassing |
| *Ms4a6c* | ENSMUSG00000079419 | 8 | Mouse Tissue Atlassing |
| *Ms4a7* | ENSMUSG00000024672 | 8 | Mouse Tissue Atlassing |
| *Muc1* | ENSMUSG00000042784 | 8 | Mouse Tissue Atlassing |
| *Mustn1* | ENSMUSG00000042485 | 7 | Mouse Tissue Atlassing |
| *Myf6* | ENSMUSG00000035923 | 8 | Mouse Tissue Atlassing |
| *Myh11* | ENSMUSG00000018830 | 8 | Mouse Tissue Atlassing |
| *Myl9* | ENSMUSG00000067818 | 8 | Mouse Tissue Atlassing |
| *Mylk* | ENSMUSG00000022836 | 8 | Mouse Tissue Atlassing |
| *Mylk3* | ENSMUSG00000031698 | 8 | Mouse Tissue Atlassing |
| *Myoz1* | ENSMUSG00000068697 | 3 | Mouse Tissue Atlassing |
| *Myoz2* | ENSMUSG00000028116 | 8 | Mouse Tissue Atlassing |
| *Myt1l* | ENSMUSG00000061911 | 8 | Mouse Tissue Atlassing |
| *Nap1l5* | ENSMUSG00000055430 | 8 | Mouse Tissue Atlassing |
| *Nbl1* | ENSMUSG00000041120 | 8 | Mouse Tissue Atlassing |

| | | | |
|---|---|---|---|
| *Ncf4* | ENSMUSG00000071715 | 8 | Mouse Tissue Atlassing |
| *Ndrg1* | ENSMUSG00000005125 | 8 | Mouse Tissue Atlassing |
| *Ndufa4l2* | ENSMUSG00000040280 | 5 | Mouse Tissue Atlassing |
| *Ndufs8* | ENSMUSG00000059734 | 8 | Mouse Tissue Atlassing |
| *Neurod1* | ENSMUSG00000034701 | 8 | Mouse Tissue Atlassing |
| *Nnat* | ENSMUSG00000067786 | 3 | Mouse Tissue Atlassing |
| *Nox4* | ENSMUSG00000030562 | 8 | Mouse Tissue Atlassing |
| *Nts* | ENSMUSG00000019890 | 8 | Mouse Tissue Atlassing |
| *Nupr1* | ENSMUSG00000030717 | 4 | Mouse Tissue Atlassing |
| *Oit1* | ENSMUSG00000021749 | 8 | Mouse Tissue Atlassing |
| *Oit3* | ENSMUSG00000009654 | 8 | Mouse Tissue Atlassing |
| *Omp* | ENSMUSG00000074006 | 8 | Mouse Tissue Atlassing |
| *Otor* | ENSMUSG00000027416 | 8 | Mouse Tissue Atlassing |
| *Pax8* | ENSMUSG00000026976 | 8 | Mouse Tissue Atlassing |
| *Pck1* | ENSMUSG00000027513 | 8 | Mouse Tissue Atlassing |
| *Pcolce2* | ENSMUSG00000015354 | 8 | Mouse Tissue Atlassing |
| *Pf4* | ENSMUSG00000029373 | 5 | Mouse Tissue Atlassing |
| *Pgam2* | ENSMUSG00000020475 | 5 | Mouse Tissue Atlassing |
| *Pglyrp1* | ENSMUSG00000030413 | 5 | Mouse Tissue Atlassing |
| *Pi16* | ENSMUSG00000024011 | 8 | Mouse Tissue Atlassing |
| *Plbd1* | ENSMUSG00000030214 | 8 | Mouse Tissue Atlassing |
| *Pln* | ENSMUSG00000038583 | 8 | Mouse Tissue Atlassing |
| *Plp1* | ENSMUSG00000031425 | 2 | Mouse Tissue Atlassing |
| *Plvap* | ENSMUSG00000034845 | 8 | Mouse Tissue Atlassing |
| *Pmp22* | ENSMUSG00000018217 | 8 | Mouse Tissue Atlassing |
| *Pnmal2* | ENSMUSG00000070802 | 8 | Mouse Tissue Atlassing |
| *Podxl* | ENSMUSG00000025608 | 8 | Mouse Tissue Atlassing |
| *Pou3f3* | ENSMUSG00000045515 | 8 | Mouse Tissue Atlassing |
| *Ppargc1a* | ENSMUSG00000029167 | 8 | Mouse Tissue Atlassing |
| *Ppbp* | ENSMUSG00000029372 | 8 | Mouse Tissue Atlassing |
| *Ppp1r1a* | ENSMUSG00000022490 | 8 | Mouse Tissue Atlassing |
| *Prap1* | ENSMUSG00000025467 | 4 | Mouse Tissue Atlassing |
| *Prg2* | ENSMUSG00000027073 | 8 | Mouse Tissue Atlassing |
| *Prom2* | ENSMUSG00000027376 | 8 | Mouse Tissue Atlassing |
| *Prox1* | ENSMUSG00000010175 | 8 | Mouse Tissue Atlassing |
| *Prss23* | ENSMUSG00000039405 | 8 | Mouse Tissue Atlassing |
| *Prss3* | ENSMUSG00000071519 | 7 | Mouse Tissue Atlassing |
| *Prx* | ENSMUSG00000053198 | 8 | Mouse Tissue Atlassing |
| *Psmb8* | ENSMUSG00000024338 | 8 | Mouse Tissue Atlassing |
| *Ptn* | ENSMUSG00000029838 | 8 | Mouse Tissue Atlassing |
| *Ptprn2* | ENSMUSG00000056553 | 8 | Mouse Tissue Atlassing |
| *Pvalb* | ENSMUSG00000005716 | 4 | Mouse Tissue Atlassing |
| *Pygm* | ENSMUSG00000032648 | 8 | Mouse Tissue Atlassing |
| *Rab3b* | ENSMUSG00000003411 | 8 | Mouse Tissue Atlassing |
| *Rac2* | ENSMUSG00000033220 | 8 | Mouse Tissue Atlassing |
| *Rasd1* | ENSMUSG00000049892 | 8 | Mouse Tissue Atlassing |
| *Rbp1* | ENSMUSG00000046402 | 8 | Mouse Tissue Atlassing |
| *Rbp2* | ENSMUSG00000032454 | 4 | Mouse Tissue Atlassing |
| *Rbp7* | ENSMUSG00000028996 | 4 | Mouse Tissue Atlassing |
| *Reg3g* | ENSMUSG00000030017 | 8 | Mouse Tissue Atlassing |
| *Retn* | ENSMUSG00000012705 | 8 | Mouse Tissue Atlassing |
| *Retnla* | ENSMUSG00000061100 | 8 | Mouse Tissue Atlassing |
| *Rgs5* | ENSMUSG00000026678 | 8 | Mouse Tissue Atlassing |
| *Rhcg* | ENSMUSG00000030549 | 8 | Mouse Tissue Atlassing |
| *Rhd* | ENSMUSG00000028825 | 8 | Mouse Tissue Atlassing |
| *Rho* | ENSMUSG00000030324 | 8 | Mouse Tissue Atlassing |

| | | | |
|---|---|---|---|
| *Riiad1* | ENSMUSG00000028139 | 6 | Mouse Tissue Atlassing |
| *Rsad2* | ENSMUSG00000020641 | 8 | Mouse Tissue Atlassing |
| *Runx2* | ENSMUSG00000039153 | 8 | Mouse Tissue Atlassing |
| *S100a14* | ENSMUSG00000042306 | 7 | Mouse Tissue Atlassing |
| *S100a4* | ENSMUSG00000001020 | 6 | Mouse Tissue Atlassing |
| *Satb1* | ENSMUSG00000023927 | 8 | Mouse Tissue Atlassing |
| *Sbsn* | ENSMUSG00000046056 | 8 | Mouse Tissue Atlassing |
| *Scg2* | ENSMUSG00000050711 | 8 | Mouse Tissue Atlassing |
| *Scg3* | ENSMUSG00000032181 | 8 | Mouse Tissue Atlassing |
| *Scg5* | ENSMUSG00000023236 | 8 | Mouse Tissue Atlassing |
| *Scgb3a2* | ENSMUSG00000038791 | 8 | Mouse Tissue Atlassing |
| *Scin* | ENSMUSG00000002565 | 8 | Mouse Tissue Atlassing |
| *Sct* | ENSMUSG00000038580 | 1 | Mouse Tissue Atlassing |
| *Sdc1* | ENSMUSG00000020592 | 8 | Mouse Tissue Atlassing |
| *Selenbp1* | ENSMUSG00000068874 | 8 | Mouse Tissue Atlassing |
| *Serpina10* | ENSMUSG00000061947 | 8 | Mouse Tissue Atlassing |
| *Serpina3n* | ENSMUSG00000021091 | 8 | Mouse Tissue Atlassing |
| *Serpinb11* | ENSMUSG00000026327 | 8 | Mouse Tissue Atlassing |
| *Serpinb5* | ENSMUSG00000067006 | 8 | Mouse Tissue Atlassing |
| *Serpinf1* | ENSMUSG00000000753 | 8 | Mouse Tissue Atlassing |
| *Sez6l* | ENSMUSG00000058153 | 8 | Mouse Tissue Atlassing |
| *Sfrp1* | ENSMUSG00000031548 | 8 | Mouse Tissue Atlassing |
| *Sfta2* | ENSMUSG00000090509 | 4 | Mouse Tissue Atlassing |
| *Sftpd* | ENSMUSG00000021795 | 8 | Mouse Tissue Atlassing |
| *Slc10a1* | ENSMUSG00000021135 | 8 | Mouse Tissue Atlassing |
| *Slc14a1* | ENSMUSG00000059336 | 8 | Mouse Tissue Atlassing |
| *Slc14a2* | ENSMUSG00000024552 | 8 | Mouse Tissue Atlassing |
| *Slc17a7* | ENSMUSG00000070570 | 8 | Mouse Tissue Atlassing |
| *Slc22a6* | ENSMUSG00000024650 | 8 | Mouse Tissue Atlassing |
| *Slc22a8* | ENSMUSG00000063796 | 8 | Mouse Tissue Atlassing |
| *Slc25a37* | ENSMUSG00000034248 | 8 | Mouse Tissue Atlassing |
| *Slc25a47* | ENSMUSG00000048856 | 8 | Mouse Tissue Atlassing |
| *Slc27a2* | ENSMUSG00000027359 | 8 | Mouse Tissue Atlassing |
| *Slc34a2* | ENSMUSG00000029188 | 8 | Mouse Tissue Atlassing |
| *Slc4a1* | ENSMUSG00000006574 | 8 | Mouse Tissue Atlassing |
| *Slc4a4* | ENSMUSG00000060961 | 8 | Mouse Tissue Atlassing |
| *Slc5a1* | ENSMUSG00000011034 | 8 | Mouse Tissue Atlassing |
| *Snap25* | ENSMUSG00000027273 | 8 | Mouse Tissue Atlassing |
| *Sod3* | ENSMUSG00000072941 | 8 | Mouse Tissue Atlassing |
| *Sostdc1* | ENSMUSG00000036169 | 8 | Mouse Tissue Atlassing |
| *Sox17* | ENSMUSG00000025902 | 8 | Mouse Tissue Atlassing |
| *Sox2* | ENSMUSG00000074637 | 8 | Mouse Tissue Atlassing |
| *Sox9* | ENSMUSG00000000567 | 8 | Mouse Tissue Atlassing |
| *Stfa2l1* | ENSMUSG00000059657 | 4 | Mouse Tissue Atlassing |
| *Stmn1* | ENSMUSG00000028832 | 8 | Mouse Tissue Atlassing |
| *Sypl2* | ENSMUSG00000027887 | 8 | Mouse Tissue Atlassing |
| *Tagln* | ENSMUSG00000032085 | 4 | Mouse Tissue Atlassing |
| *Tat* | ENSMUSG00000001670 | 8 | Mouse Tissue Atlassing |
| *Tbx1* | ENSMUSG00000009097 | 6 | Mouse Tissue Atlassing |
| *Tcf15* | ENSMUSG00000068079 | 2 | Mouse Tissue Atlassing |
| *Tfcp2l1* | ENSMUSG00000026380 | 8 | Mouse Tissue Atlassing |
| *Tff2* | ENSMUSG00000024028 | 5 | Mouse Tissue Atlassing |
| *Thbs4* | ENSMUSG00000021702 | 8 | Mouse Tissue Atlassing |
| *Them5* | ENSMUSG00000028148 | 8 | Mouse Tissue Atlassing |
| *Tie1* | ENSMUSG00000033191 | 8 | Mouse Tissue Atlassing |
| *Timp3* | ENSMUSG00000020044 | 8 | Mouse Tissue Atlassing |

| | | | |
|---|---|---|---|
| *Timp4* | ENSMUSG00000030317 | 8 | Mouse Tissue Atlassing |
| *Tm4sf20* | ENSMUSG00000026149 | 8 | Mouse Tissue Atlassing |
| *Tm4sf4* | ENSMUSG00000027801 | 8 | Mouse Tissue Atlassing |
| *Tmem100* | ENSMUSG00000069763 | 8 | Mouse Tissue Atlassing |
| *Tmem147* | ENSMUSG00000006315 | 4 | Mouse Tissue Atlassing |
| *Tmem212* | ENSMUSG00000043164 | 8 | Mouse Tissue Atlassing |
| *Tmem213* | ENSMUSG00000029829 | 3 | Mouse Tissue Atlassing |
| *Tmem59l* | ENSMUSG00000035964 | 8 | Mouse Tissue Atlassing |
| *Tnfaip6* | ENSMUSG00000053475 | 8 | Mouse Tissue Atlassing |
| *Tnnc1* | ENSMUSG00000091898 | 5 | Mouse Tissue Atlassing |
| *Trdn* | ENSMUSG00000019787 | 8 | Mouse Tissue Atlassing |
| *Trim29* | ENSMUSG00000032013 | 8 | Mouse Tissue Atlassing |
| *Try10* | ENSMUSG00000071521 | 8 | Mouse Tissue Atlassing |
| *Tspan8* | ENSMUSG00000034127 | 8 | Mouse Tissue Atlassing |
| *Ttc36* | ENSMUSG00000039438 | 1 | Mouse Tissue Atlassing |
| *Ttn* | ENSMUSG00000051747 | 8 | Mouse Tissue Atlassing |
| *Tubb3* | ENSMUSG00000062380 | 8 | Mouse Tissue Atlassing |
| *Ube2c* | ENSMUSG00000001403 | 8 | Mouse Tissue Atlassing |
| *Ucma* | ENSMUSG00000026668 | 8 | Mouse Tissue Atlassing |
| *Uox* | ENSMUSG00000028186 | 8 | Mouse Tissue Atlassing |
| *Upk1a* | ENSMUSG00000006313 | 6 | Mouse Tissue Atlassing |
| *Upk1b* | ENSMUSG00000049436 | 8 | Mouse Tissue Atlassing |
| *Upk3a* | ENSMUSG00000022435 | 7 | Mouse Tissue Atlassing |
| *Upk3b* | ENSMUSG00000042985 | 8 | Mouse Tissue Atlassing |
| *Vsnl1* | ENSMUSG00000054459 | 8 | Mouse Tissue Atlassing |
| *Vsx2* | ENSMUSG00000021239 | 8 | Mouse Tissue Atlassing |
| *Vwf* | ENSMUSG00000001930 | 8 | Mouse Tissue Atlassing |
| *Wif1* | ENSMUSG00000020218 | 8 | Mouse Tissue Atlassing |
| *Wnt3* | ENSMUSG00000000125 | 8 | Mouse Tissue Atlassing |
